# BloodVariome: a high-resolution atlas of inherited genetic effects in human immune cells

**DOI:** 10.64898/2026.03.30.715213

**Authors:** Aitzkoa Lopez de Lapuente Portilla, Ludvig Ekdahl, Gudmar Thorleifsson, Zain Ali, Antton Lamarca Arrizabalaga, Caterina Cafaro, Saedis Saevarsdottir, Gudny E. Thorlacius, Gisli H Halldorsson, Lilja Stefansdottir, Pall Melsted, Nerea Ugidos Damboriena, Maroulio Pertesi, Patrick Sulem, Daniel F Gudbjartsson, Kari Stefansson, Unnur Thorsteinsdottir, Ingileif Jonsdottir, Thorunn Olafsdottir, Björn Nilsson

## Abstract

Genome-wide association studies have linked thousands of sequence variants to immune-mediated diseases, yet their cellular mechanisms remain largely unresolved. Here we present BloodVariome, a high-resolution atlas of genetic effects across the human immune cell hierarchy. Combining deep immunophenotyping with automated pattern-recognition, we quantified 1,533 traits across 127 immune cell populations in 11,983 individuals. We identified 259 significant associations, the vast majority of which are not captured by conventional bulk blood trait studies. Most associations were restricted to single immune lineages or cell populations, revealing a fine-grained genetic compartmentalization of the immune system. By linking known disease risk alleles to specific immune cell phenotypes, BloodVariome illuminates cellular mechanisms underlying autoimmunity, immunodeficiency, and hematologic malignancy. Moreover, we implicate novel regulators of human immune cell development and function. By bridging the gap between cohort size and phenotypic depth, BloodVariome establishes a high-resolution framework for interpreting how genetic variation shapes cellular immunity at population-scale.

## INTRODUCTION

The human immune system comprises a hierarchically organized network of specialized cell populations that maintain host defense and tissue homeostasis^1^. This functional versatility reflects extensive heterogeneity across lineages, differentiation stages, and activation programs. Many immune cell traits remain stable within individuals yet vary substantially across the population, shaping differences in immune function and disease susceptibility. Twin and population studies indicate that a substantial fraction of this variation is heritable^2–4^.

Genome-wide association studies (GWAS) have identified thousands of DNA sequence variants linked to human diseases, many mapping to immune-related loci^5^. Yet, the cellular mechanisms through which these variants act remain unclear. Large-scale GWAS using electronic health records have connected genetic variation to broad blood cell traits, including total white blood cell counts, erythrocyte indices, and platelet measures^6–8^. Despite their power – cohorts of more than half a million individuals – these studies lack the resolution to assess specific immune cell subsets. In contrast, high-resolution flow cytometry studies have demonstrated that genetic effects can be resolved at the level of immune cell subpopulations^9–14^, but sample sizes have remained limited (≤3,757 individuals).

To address this gap, we developed BloodVariome, a population-scale framework for mapping inherited genetic effects across the immune cell hierarchy. By combining deep flow cytometry with automated pattern-recognition gating, we quantified 1,533 traits across 127 immune cell populations in 11,983 individuals. Our data reveal an unexpectedly fine-grained genetic compartmentalization of the human immune system, unveil specific immune cell endophenotypes mediating disease risk, and uncover novel regulators of human immune cell development. In summary, BloodVariome establishes a high-resolution framework for interpreting how inherited genetic variation shapes human cellular immunity.

## RESULTS

### A population-scale genetic atlas of human immune cell variation

#### Population-scale immune cell phenotyping

To systematically map variation in the human immune cell hierarchy, we analyzed peripheral blood from 11,983 individuals of Swedish ancestry aged 18-72 years (**Figures 1A** and **S1**; **Table S1A**). Using high-resolution flow cytometry, we quantified up to 127 immune cell populations spanning T-cell, B-cell, natural killer cell, monocyte, and dendritic cell lineages (**Figures 1B** and **S2**; **Tables S1B**-**S1D**). Because many subsets are rare, we applied a high sampling depth (up to 1 million events per sample; **Table S2A**).

**Figure 1:**
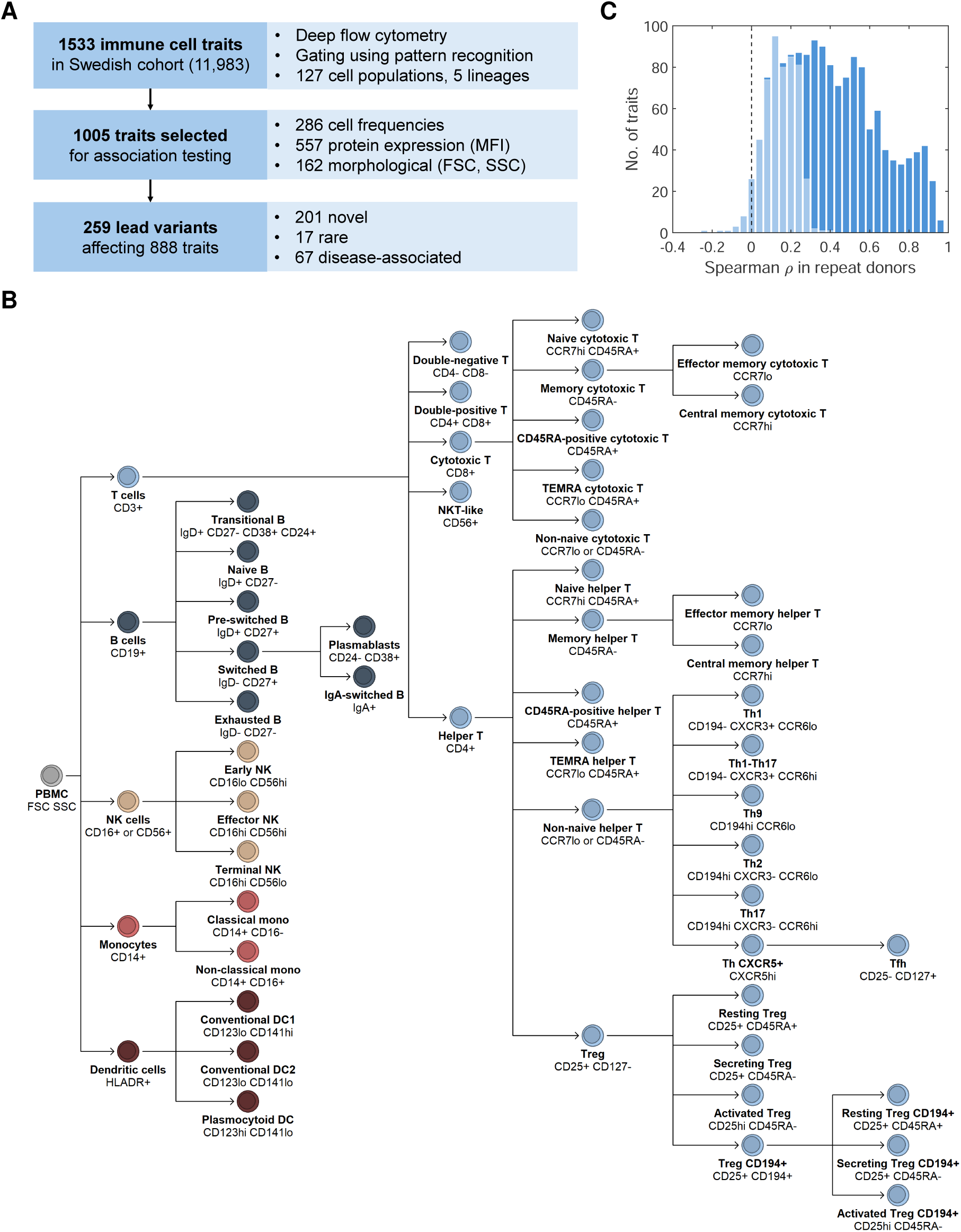
Study design and high-resolution immune phenotyping framework. **(A)** Overview of the BloodVariome study design, including cohort selection, deep immunophenotyping, and genetic findings. Traits were derived from cell frequencies, surface protein expression (median fluorescence intensity, MFI), and morphology (forward scatter, FSC; side scatter, SSC). **(B)** We analyzed 127 immune cell types belonging to five lineages: T cells (light blue), B cells (dark blue), NK cells (beige), monocytes (red), and dendritic cells (brown). This dendrogram illustrates the hierarchical relationships among major populations within the immune cell landscape. Additional subpopulations were defined by HLA-DR (across lineages) and CD39 (within T cell populations). **(C)** Trait reproducibility assessed using duplicate samples from the same donors collected 3 to 9 months apart (n = 263). Protein expression and morphology traits showing significant Spearman correlation were retained together with all frequency traits, yielding 1,005 traits for downstream analysis (dark blue); remaining traits were excluded (light blue). **Abbreviations:** peripheral blood mononuclear cells (PBMC); natural killer cells (NK); dendritic cells (DC); terminal effector memory RA-positive (TEMRA); helper T cells (Th); follicular helper T cells (Tfh); regulatory T cells (Tregs); monocytes (mono); forward scatter (FSC); side scatter (SSC); cluster of differentiation (CD); immunoglobulin A and D (IgA and IgD); C-C chemokine receptor (CCR); human leukocyte antigen DR isotype (HLADR).

To enable operator-independent analysis of large-scale, high-dimensional flow cytometry data, we developed bespoke software (AliGater) for automated pattern-recognition gating^15^. Gating of 127 populations across nearly 12,000 samples requires hundreds of thousands of individual gating decisions, making automation essential for population-scale analysis. All gating results were subsequently reviewed by manual quality control.

For each cell population, we quantified three complementary trait classes capturing cellular abundance (absolute and relative frequencies), surface protein expression (median fluorescence intensity of antibody targets), and cell morphology (forward scatter for size; side scatter for complexity), yielding 1,533 traits (**Figure 1A**; **Table S2A**). Longitudinal resampling of 263 individuals after 3-9 months confirmed reproducibility for 977 traits (**Figure 1C**; **Table S2B**). We retained all frequency traits together with reproducible protein and morphology traits, resulting in 1,005 traits for downstream analysis (**Figure 1A**).

Age and sex exerted widespread, biologically coherent effects across the hierarchy: 546 traits (54.3%) were associated with age and 435 (43.3%) with sex (**Figure 2**; **Table S3**). Age-related changes were consistent with immunosenescence, including increased memory-to-naive T-cell ratios, expansion of secreting *vs.* resting regulatory T cells, elevated Th2 frequencies^16–19^, and reduced plasmablast and pre-switched B cell frequencies^20^. Sex differences were also pronounced: women exhibited higher T cell frequencies, consistent with slower thymic involution^13,21^, whereas men had higher frequencies of innate cells.

**Figure 2:**
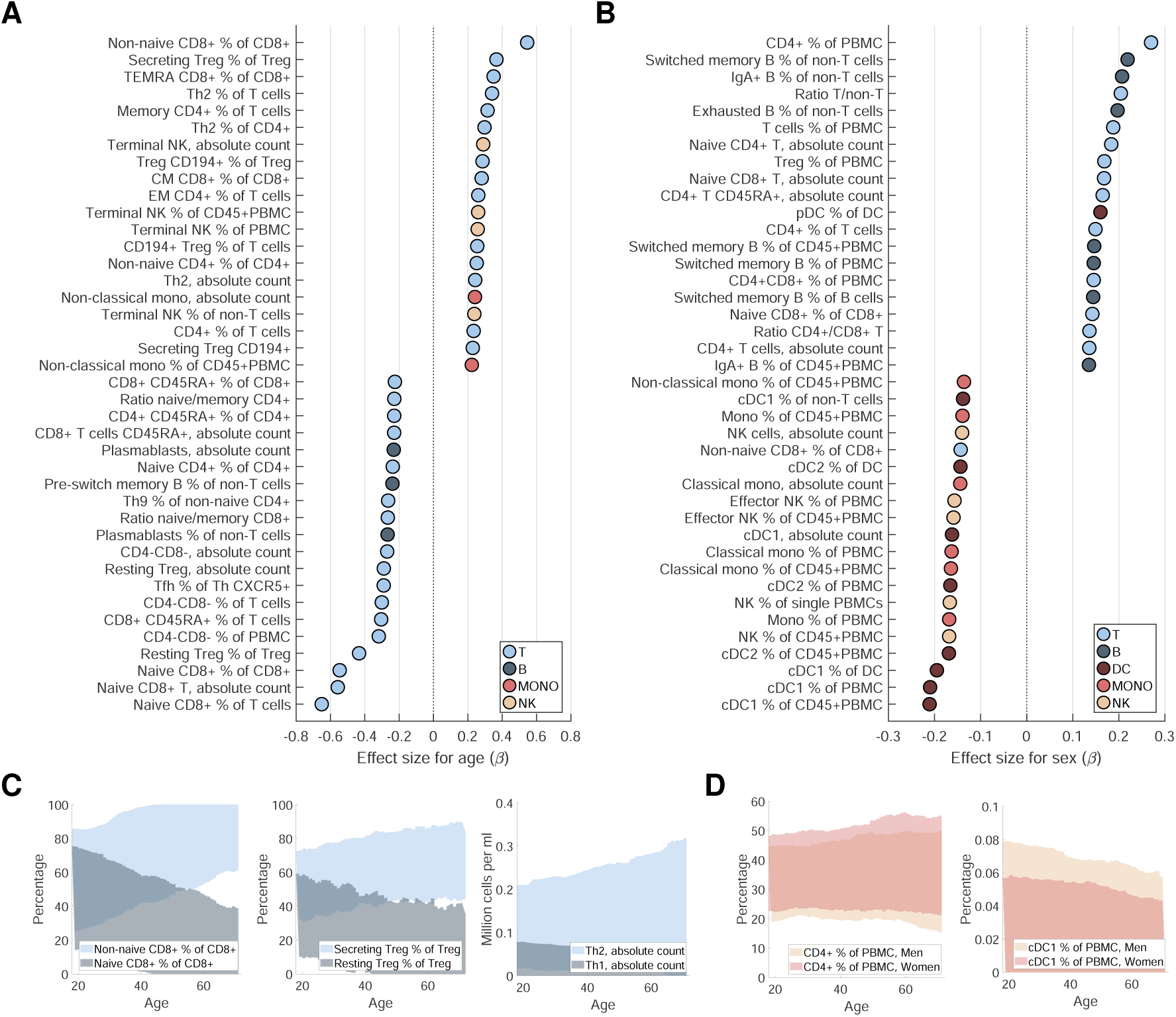
Age and sex are major determinants of immune cell composition. **(A)** Immune cell traits showing the strongest associations with age. Effect sizes (multivariable regression β) are plotted on the *x*-axis; positive values indicate increasing trait levels with age. Colors denote lineage: T cells (light blue), B cells (dark blue), NK cells (beige), monocytes (red), and dendritic cells (brown). **(B)** Immune cell traits showing the strongest association with sex. Positive values indicate higher trait levels in women than in men. **(C)** Age-associated immune remodeling includes increased proportions of non-naïve CD8^+^ and CD4^+^ T cells relative to their naïve counterparts, an increased fraction of secreting versus resting regulatory T cells, and elevated T_h_2 frequencies with stable T_h_1 levels. **(D)** Sex dimorphism in immune composition, with women showing higher frequencies of multiple T cell populations, whereas men exhibit higher levels of innate immune cell types. A complete list of association statistics for the age and sex is provided in **Table S3**. **Abbreviations:** peripheral blood mononuclear cells (PBMC); natural killer cells (NK); dendritic cells (DC); conventional dendritic cells (cDC); terminal effector memory RA-positive (TEMRA); helper T cells (Th); follicular helper T cells (Tfh); regulatory T cells (Tregs); effector memory (EM); central memory (CM); monocytes (mono); cluster of differentiation (CD); immunoglobulin A and D (IgA and IgD); C-C chemokine receptor (CCR); human leukocyte antigen DR isotype (HLA-DR).

These data establish BloodVariome as a population-scale, high-resolution framework for genetic dissection of immune cell traits.

#### Fine-grained genetic architecture of immune cell traits

Genome-wide association testing of 1,005 traits against 32 million sequence variants (**Methods**) revealed a highly structured genetic architecture of immune cell traits. After correction for multiple testing across variant classes^22^ and the effective number of independent traits, we identified 259 genome-wide significant signals affecting 888 traits (**Tables S4A-E; Figure 1A** and **S3**), including 174 that also reached study-wide significance. Stepwise conditional analyses identified independent association signals and loci shared across traits (**Methods**), and we constructed 95% credible sets of candidate causal variants for all non-HLA autosomal signals (**Tables S5A-B**).

Genetic effects were highly lineage-specific: 225 (87%) were confined to a single lineage (B, T, NK, DC, or monocyte) and 90 (35%) affected only one trait (**Figures 3A-C**). Only 71 (27%) signals overlapped bulk white blood cell traits from electronic health records (**Table S6A**)^6-8^. Most associations (199, 77%) were previously unreported, whereas 23% replicated known loci (**Tables S6B-C**)^10–14^.

**Figure 3:**
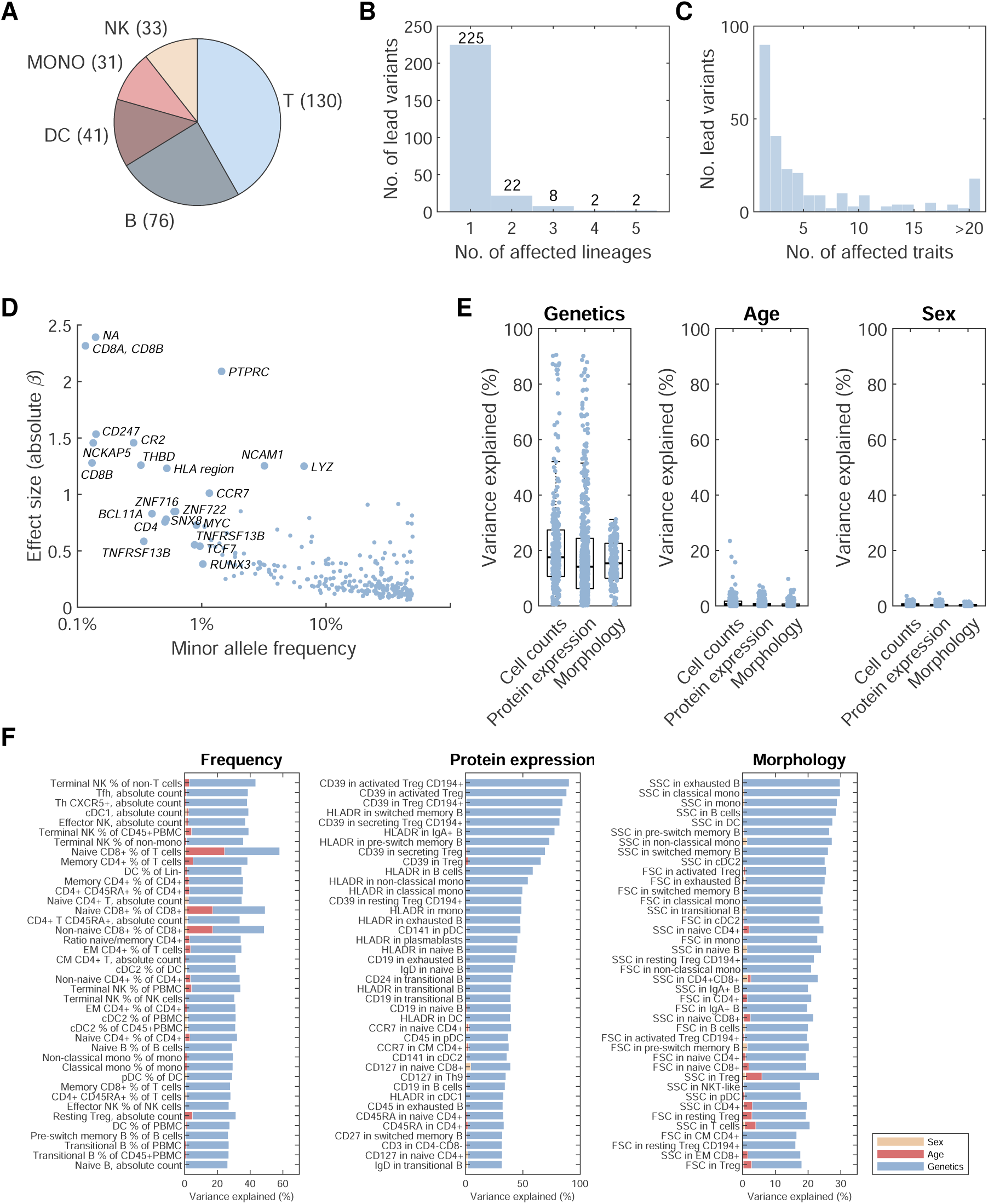
Genetic architecture of immune cell traits revealed by genome-wide association analysis. We tested the 1,005 immune cell traits for association with ∼32 million variants and performed stepwise conditional analyses to identify independent signals and shared effects across traits. **(A)** Number of significant associations observed within each immune cell lineage. **(B, C)** Pleiotropy of lead variants, shown as the number of lineages and individual traits affected by each variant. **(D)** Relationship between minor allele frequencies (MAF) and effect size (absolute β) for lead variant—trait associations (most significant trait per variant). Variants with MAF ≤1% and effect size >1 are highlighted and annotated with candidate genes. **(E)** Variance explained by genetic effects, age, and sex for frequency, protein expression, and morphology traits. **(F)** Traits showing the strongest genetic contribution (*h*^2^) across frequency, protein expression, and morphology categories. Blue, red, and beige indicate variance explained by genetics, age, and sex, respectively. Complete lists of lead variants and their associations are provided in **Table S4B-C**. Variance components for all traits are provided in **Table S3**. Visualizations of all lead variant effects are available in **Data S1**. **Abbreviations:** regulatory T cells (Treg); conventional dendritic cells type 1 and 2 (cDC1, cDC2); natural killer cells (NK); plasmacytoid dendritic cells (pDC); effector memory (EM); central memory (CM); cluster of differentiation (CD); immunoglobulin D (IgD); C-C chemokine receptor (CCR); human leukocyte antigen DR isotype (HLADR).

Seventeen signals involved rare variants (allele frequency, AF, < 1%; **Figure 3D**), and nearly all lead variants were polymorphic across ancestries (**Table S6D**). Effect sizes varied widely, explaining up to 95.4% of trait variance (median 5.1%) and often exceeding contributions from age and sex (**Figures 3E-F** and **S4**; **Table S3**). Linkage disequilibrium score regression^23^ estimated total SNP heritability at 0 to 90.5% (median 16.1%; **Table S3**).

Together, these results reveal a highly compartmentalized genetic architecture of immune cell variation, in which most variants act within specific immune lineages and often on individual cellular phenotypes. This pronounced specificity highlights the power of deep immunophenotyping to resolve genetic effects that remain invisible in bulk blood traits.

#### Genetic prioritization links variants to immune-relevant candidate genes

To assign candidate genes, we integrated credible sets (**Tables S5A-B**) with coding annotations, regulatory QTLs, and epigenomic features (**Tables S7A-D**; **Methods**). This approach nominated 167 genes across 230 signals (**Figure S6**).

Prioritized genes showed lineage-specific expression consistent with their associated traits (**Table S8A**)^24^ and were significantly enriched for immune phenotypes in knockout mice (86 genes; 51% *vs* 27% genome-wide; binomial test *P* = 5.8×10^−11^; **Table S8B**)^25^. In addition, 43 genes underlie Mendelian immune disorders (**Table S8C**)^26^; 24 are recurrently mutated in immune cell malignancies, particularly lymphomas and acute leukemias (binomial test *P* = 2.0×10^−5^ to 4.6×10^−10^; **Table S8D-E**)^27,28^, and 64 show DepMap essentiality in immunological cell lines (**Table S8E**)^29^. These orthogonal enrichments provide convergent genetic support for the immune relevance of the prioritized genes.

Together, these analyses delineate a finely resolved genetic landscape of immune cell regulation and establish BloodVariome as a resource for interrogating human immune variation. To facilitate community use, we developed an interactive web portal (https://bloodvariome.medfak.lu.se) that enables visualization of variant effects across the immune cell hierarchy and gene-centric exploration of significant associations, with comprehensive plots provided for all 259 lead variants (**Data S1**). Additionally, full summary statistics have been deposited in the GWAS Catalog (accession no. pending).

### Mapping disease risk variants to immune cell states

Having established a high-resolution genetic map of immune cell variation, we next asked through which specific immune cell states disease-associated variants exert their effects. By integrating BloodVariome with the GWAS catalog^5^ and ClinVar^30^, we identified 67 credible sets overlapping risk variants for 60 disease outcomes. These included allergic and autoimmune disorders (n = 50 variants), primary immunodeficiencies and chronic infections (n = 14), hematologic malignancies (n = 3), and a broader set of non-immunological conditions (n = 21; **Figure 4**; **Tables S9A-B**).

**Figure 4:**
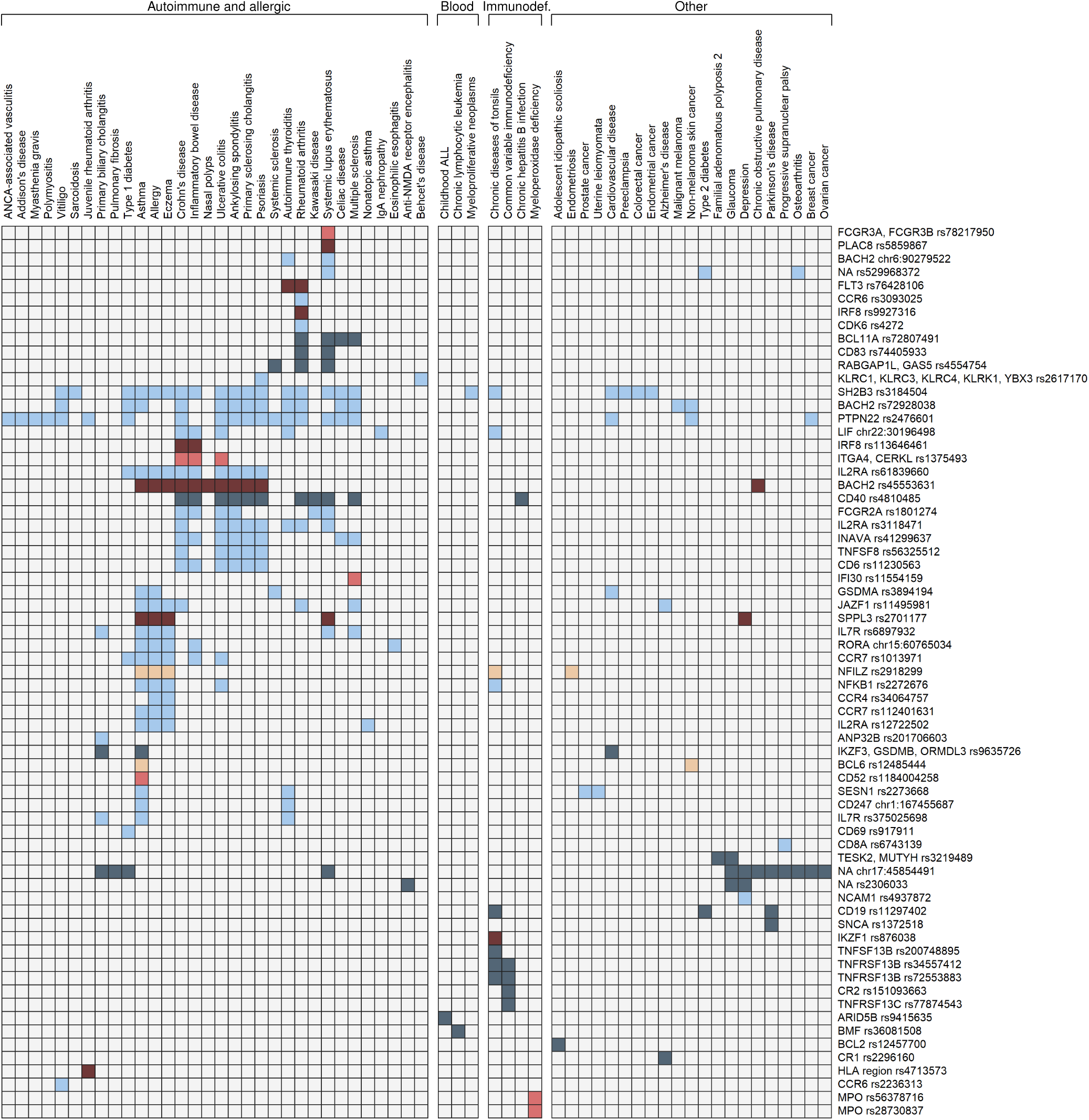
Immune cell endophenotypes link genetic variants to human disease risk. Among the 259 genome-wide significant signals, 67 overlapped previously reported disease-associated variants in the GWAS Catalog or ClinVar (linkage disequilibrium *r*^2^ > 0.8 between catalog and BloodVariome lead variants). The heatmap illustrates the relationship between genetic effects on immune cell traits and disease susceptibility. Colors indicate the lineage of the most strongly associated immune cell trait in BloodVariome using the same scheme as in Figure 1: T cells (light blue), B cells (dark blue), NK cells (beige), monocytes (red), and dendritic cells (brown). Overlapping loci were linked to 60 disease outcomes, including autoimmune and allergic disorders, primary immunodeficiencies and chronic infections, hematologic malignancies, and a broader set of non-immunological conditions. Complete lists of overlapping disease associations are provided in **Table S9A**.

Below we highlight representative examples that illustrate how BloodVariome resolves disease-associated genetic effects at subpopulation resolution.

#### An FLT3 variant predisposes to autoimmunity by expanding cDC2s and monocytes

The low-frequency variant rs76428106-C in *FLT3* (AF = 1.6%; **Figures 5A** and **S6**) showed strong myeloid-restricted effects, marked by expansion of type 2 conventional dendritic cells (cDC2; *P* = 9.2×10^−17^, β = 0.63), accompanied by increased total dendritic cell frequency (*P* = 6.1×10^−18^, β = 0.66), elevated monocyte frequency (*P* = 6.1×10^−12^, β = 0.39), and a modest reduction in T cell frequency (*P* = 6.0×10^−11^, β = −0.34; **Figure S6A**). This allele is the strongest known risk factor for autoimmune thyroiditis and is also associated with rheumatoid arthritis and acute myeloid leukemia (**Figure S6B**; **Table S9A**)^31^.

**Figure 5:**
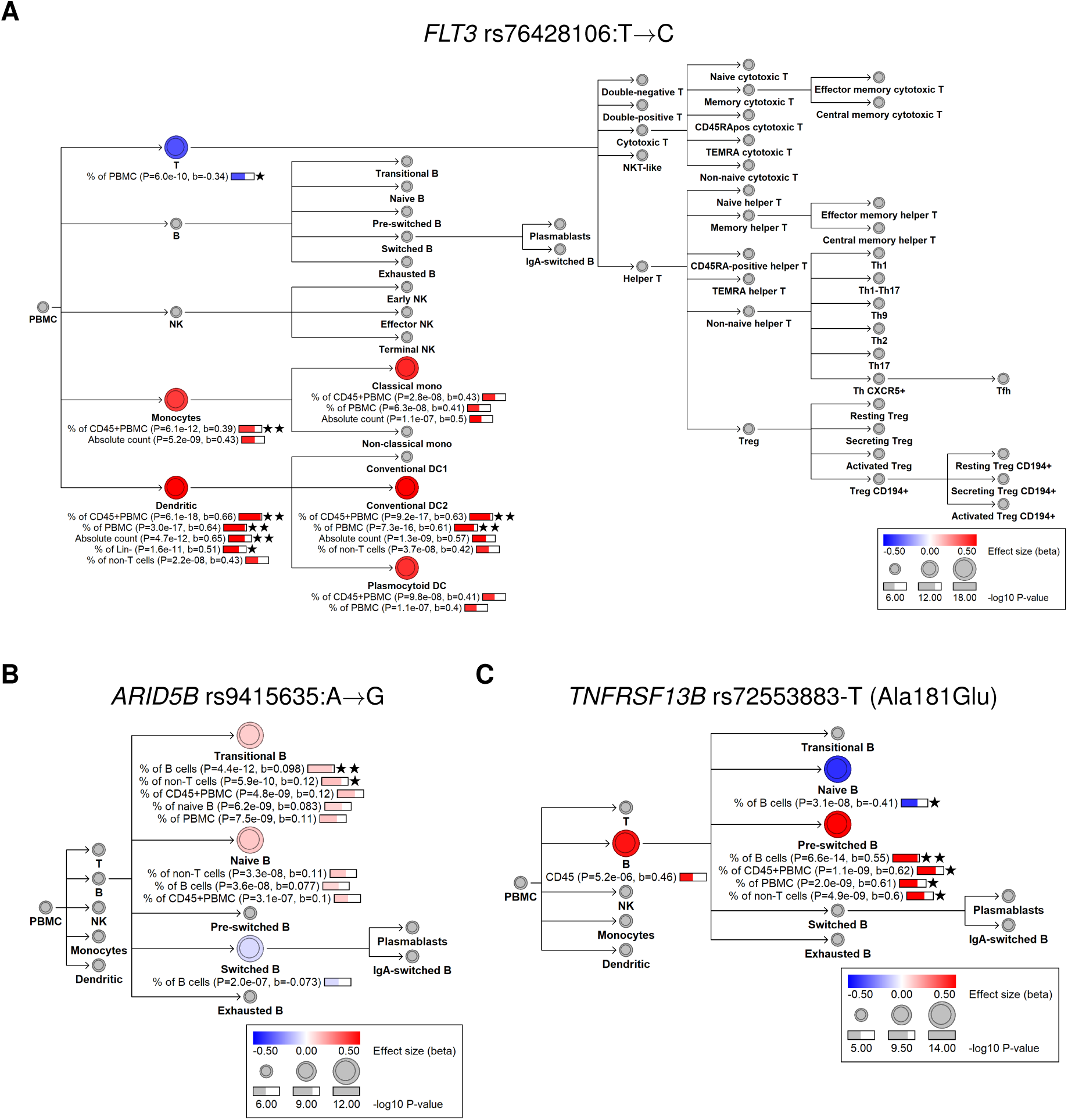
Disease-associated variants map to lineage-specific immune cell phenotypes. Hierarchy plots illustrating cellular phenotypes associated with three disease risk variants. Circular icons denote immune cell populations, and labels beneath each node indicate the associated traits. Bar lengths reflect −log_10_(P) for association, and color denotes effect size (β). Two stars indicate study-wide significance, one star indicates genome-wide significance, and no stars denote sub-significant associations (within two orders of magnitude from the genome-wide threshold). The color of each cell icon represents the effect size of the most strongly associated trait for that population. **(A)** At *FLT3*, the autoimmune thyroiditis risk allele rs76428106-C is associated with increased dendritic cell and monocyte frequencies and reduced T cell frequencies, highlighting a shift toward myeloid and dendritic cell compartments, particularly conventional dendritic cells type 2. **(B)** At *ARID5B*, the childhood B-cell precursor acute lymphoblastic leukemia risk variant is associated with expansion of transitional B cells relative to total B cells, consistent with altered early B-cell development. The illustration is restricted to the B-cell lineage because no significant effects were observed in other lineages. **(C)** At *TNFRSF13B*, a well-characterized CVID risk variant (rs72553883-T; Ala181Glu) is associated with increased pre-switched B cells within the B-cell compartment, consistent with impaired B-cell development beyond the class-switching stage. The illustration is restricted to the B-cell lineage because no significant effects were observed in other lineages. Additional supporting data are provided in **Figures S6**, **S9**, and **S10**.

*FLT3* encodes a receptor tyrosine kinase essential for hematopoietic progenitor, dendritic cell, and monocyte development (**Figure S6C**)^32^. rs76428106-C, located in intron 15 (**Figure 6D**), introduces a cryptic splice site that truncates ∼30% of transcripts^31^. Despite the predicted reduction of normal transcript levels, the variant elevates plasma FLT3 ligand levels and monocyte counts, consistent with a net gain-of-function effect^31^. BloodVariome clarifies the cellular interpretation of this locus by identifying cDC2s as the predominant myeloid endophenotype linking rs76428106-C to autoimmunity, rather than this susceptibility being mediated by increased plasmacytoid dendritic cells which we proposed previously as a tentative mechanism^31^.

**Figure 6:**
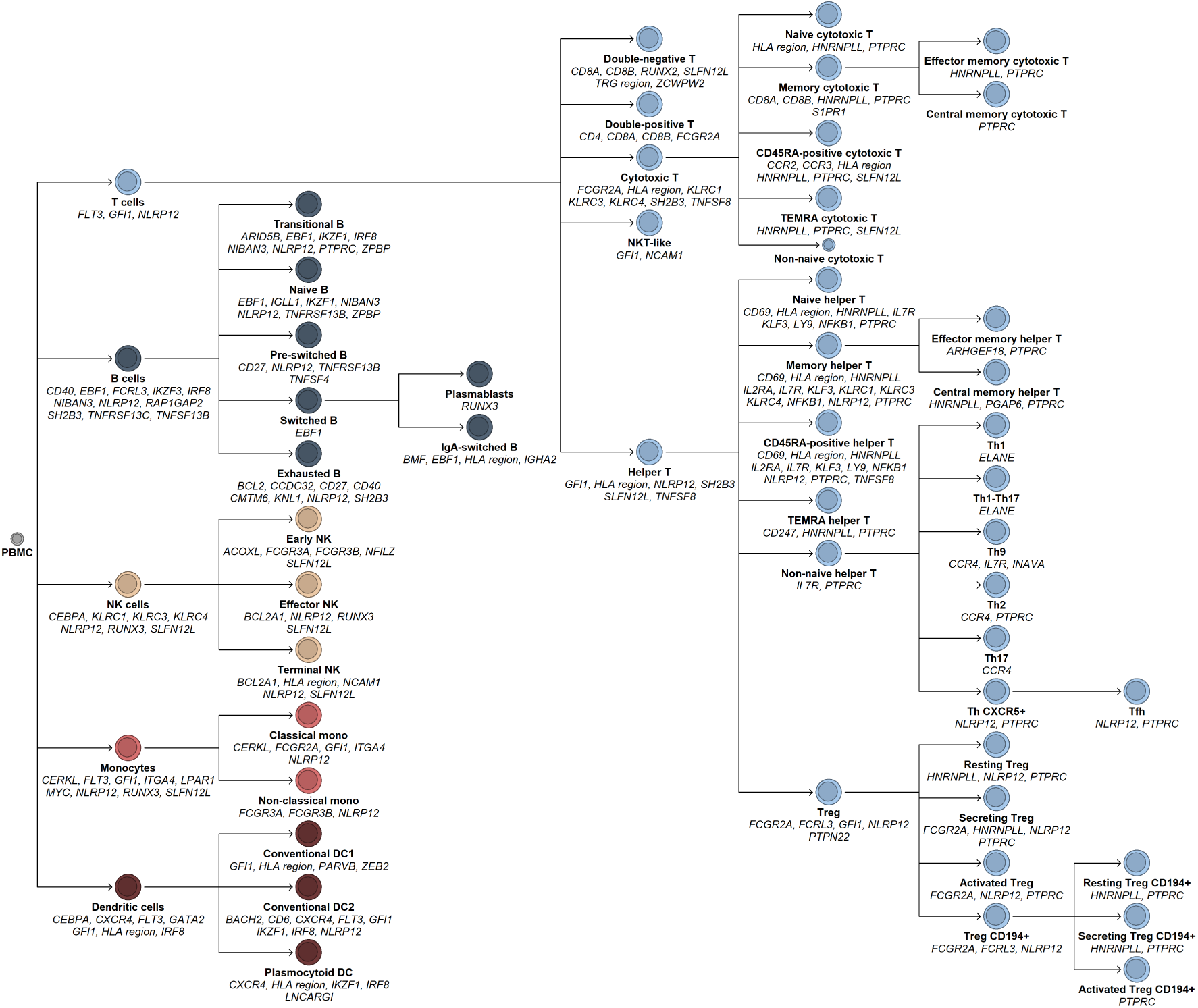
Frequency traits highlight candidate regulators of immune cell development. **(A)** Overview of candidate genes associated with frequency traits across the five lineages analyzed: T cells (light blue), B cells (dark blue), NK cells (beige), monocytes (red), and dendritic cells (brown). Gene names are positioned at the cell population whose frequency is altered by the corresponding variant. Frequency-based associations recapitulate established regulators of immune cell development and nominate genes previously characterized primarily in model systems whose roles in human immune cell development remain incompletely defined. Detailed association data are provided in **Tables S4A** and **S4B**. **(B)** At *NIBAN3,* rs11666267:T>C (p.Ser572Gly) is associated with reduced B-cell frequencies, genetically implicating *NIBAN3* in normal human B-cell development. Additional data are shown in **Figure S14**. **(C)** At *SLFN12L*, two independent variants are associated with frequencies of NK cells, dendritic cells, and T cells. Shown is the effect of the 3’-UTR variant rs58745116:G>A. Additional data are provided in **Figure S15**. **Abbreviations:** regulatory T cells (Treg); conventional dendritic cells (cDC1, cDC2); natural killer cells (NK); plasmacytoid dendritic cells (pDC); effector memory (EM); central memory (CM); cluster of differentiation (CD); immunoglobulin D (IgD); C-C chemokine receptor (CCR); human leukocyte antigen DR isotype (HLA-DR).

#### An IL7R autoimmunity-protective variant alters the naïve-to-memory CD4^+^ T cell balance through alternative splicing

*IL7R* encodes the α-subunit of the IL-7 receptor (IL-7Rα; CD127)^33^, which is critical for T cell development. The missense variant rs6897932-T (Thr244Ile; AF = 26.7%) increases the naïve-to-memory CD4^+^ T cell frequency (*P* = 3.8×10^−15^ to 1.8×10^−8^; β = 0.12; **Figure S7A**) and is associated with protection against multiple sclerosis, primary biliary cholangitis, systemic lupus erythematosus, and allergy (**Figure S7B**). Consistent with our finding, we noted reduced plasma levels of other T cell-related proteins (**Figures S7C-D**) in proteomics data from Iceland and the UK Biobank^34–37^, consistent with attenuated T-cell responses.

IL-7Rα has two main isoforms: one soluble and one membrane bound. Thr244Ile shifts splicing of *IL7R*, reducing the expression of the soluble isoform as detected both on mRNA (*P* < 1.0×10^⁻300^; β = −0.84) and protein (*P* < 1×10^−300^; β = −0.85) while we did not detect any change in membrane-bound IL-7Rα (CD127) surface protein expression (**Figure S7E**; **Table S7A,C**)^38^. These data link a common autoimmunity-protective *IL7R* variant to altered T cell function. Notably, these effects are far milder than the severe immunodeficiency caused by biallelic *IL7R* loss-of-function mutations^39–41^, highlighting the spectrum of phenotypic consequences of graded IL-7Rα perturbation. These data illustrate how BloodVariome resolved fine-grained genetic effects within the T-cell lineage.

*BACH2 variants drive distinct lineage-specific mechanisms of autoimmunity* Immune regulators frequently act across multiple lineages, raising the possibility that variation at a single locus can influence disease risk through distinct cellular routes. BloodVariome enables these effects to be resolved with cell-type resolution.

*BACH2* encodes a broadly expressed transcription factor involved in B cell, T cell, and hematopoietic progenitor biology^42–44^. We identified three independent signals at *BACH2*, each associated with inflammatory disease (**Figure S8A-D**).

Two signals converge on T follicular helper (Tfh) cells. The first, rs72928038-A (AF = 15.6%) increased CCR4 (CD194) surface expression (P = 5.5×10^−19^, β = 0.28) and conferred increased risk of multiple autoimmune diseases (**Figure S8E**). Conversely, rs10637718-C (AF = 43.1%) decreased CCR4 (β = −0.18; *P* = 4.8×10^−15^) and was protective against systemic lupus erythematosus and autoimmune thyroiditis. In both cases, the autoimmunity risk allele increased CCR4 surface expression on Tfh cells while reducing *BACH2* mRNA in T cells or blood (**Figure S8F**), consistent with BACH2’s role as a repressor of *CCR4* transcription^45,46^. CCR4-high IL-4–producing Tfh cells provide potent B cell help^45^, suggesting that elevated CCR4 expression in risk-allele carriers may promote autoantibody generation. Consistently, *Bach2* ablation in murine T cells expands IL-4–producing Tfh cells and induces lupus-like humoral autoimmunity^46^.

In contrast, a third variant, rs45553631-T, reduced cDC2 frequency (*P* = 8.0×10^−11^, β = −0.19) and decreased risk of allergy, eczema, and inflammatory bowel disease (**Figure S8E**). *Bach2^−/-^* mice similarly show reduced cDC2 levels^47^, and rs45553631-T lowers *BACH2* mRNA in blood (**Figure S8F**). Concordantly, carriers exhibit reduced plasma levels of dendritic cell markers in proteomic data from Iceland and the UK Biobank (**Figure S8G**)^34–37^.

This allelic series reveals allele- and lineage-specific pathogenic mechanisms at *BACH2*: some alleles drive autoimmunity by upregulating CCR4 in Tfh cells, whereas others modulate cDC2 frequencies. The example illustrates how BloodVariome can disentangle distinct cellular routes to disease at a single pleiotropic locus.

#### Childhood leukemia risk allele at ARID5B perturbs early B-cell development

The variant rs9415635-G at *ARID5B* selectively increased transitional B-cell frequency (*P* = 4.4×10^−12^, β = 0.098, AF = 31.5%; **Figures 5B** and **S9A-B**). The credible set includes rs7090445-C (*r*^2^ = 0.87 with rs9415635), the functionally fine-mapped causal variant at the strongest GWAS locus for childhood B cell precursor acute lymphoblastic leukemia (BCP-ALL)^48,49^. rs7090445-C upregulates *ARID5B* in B cells and is preferentially retained in BCP-ALL blasts^49^, yet its impact on normal human B-cell development remains unclear.

Transitional B cells are newly egressed immature cells from the bone marrow representing the earliest circulating stage of the B-cell lineage. The *ARID5B* risk allele shifts peripheral B-cell composition toward this immature population, indicating increased early B-cell output or delayed maturation. Consistent with this model, plasma proteomic data from Iceland and the UK Biobank^34–37^ showed elevated levels of proteins expressed in B cells and lymphoid progenitors in risk-allele carriers (**Figures S9D-E**), with the strongest effect on IGLL1 (*P* = 7.4×10^−49^; β = 0.14), a core component of the pre-B cell receptor complex.

Together, these data link the *ARID5B* leukemia risk allele to increased early B-cell output and, to our knowledge, provide the first population-scale resolution of a childhood leukemia susceptibility locus to a specific stage of human B-cell development.

#### Genetic variation at CR2 perturbs the B-cell co-receptor complex

At *CR2*, a rare stop-gain variant, rs151093663-A (p.Trp766Ter; AF = 0.283%) markedly increased CD19 surface expression across B-cell subsets (*P* = 3.1×10^−16^ to 2.5×10^−11^, β = 1.2 to 1.5; **Figure S10A**). *CR2* encodes complement receptor 2 (CD21), a component of the B-cell co-receptor complex^50^. Biallelic *CR2* loss-of-function mutations, including rs151093663-A, cause an immunodeficiency characterized by low IgG1 and IgG4, absent surface CR2, and reduced switched memory B cells^51^. CD19 effects have not previously been reported for this variant. Prior studies in CVID describe an inverse relationship between CR2 and CD19 surface expression^52,53^. Consistent with this model, *Cr2*^−/-^ mice show elevated CD19 expression and impaired B-cell responses that are partially corrected by CD19 deletion^54,55^.

An independent common variant at *CR2* (rs767535979-CA, AF = 11.3%) produced a nearly identical CD19 upregulation pattern (*P* = 1.4×10^−43^ to 2.5×10^−20^, β = 0.26 to 0.39; **Figure S10B**) together with reduced CR2 levels in plasma proteomic data (*P* = 9.5×10^−169^, β = −0.29; **Figure S10D**)^34–37^, consistent with a milder hypomorphic effect. Together, these results support a *CR2* dosage–dependent model in which reduced CR2 drives compensatory CD19 upregulation, revealing convergent genetic control of the CR2–CD19 co-receptor axis.

#### Morphological traits resolve genetic effects on immune cell ultrastructure

Whereas frequency and protein traits resolved genetic effects on developmental and regulatory programs, morphology traits reveal alterations in immune cell ultrastructure.

Two independent variants associated with reduced monocyte SSC – rs28730837-A (*P* = 1.6×10^−12^, β = −0.38) and rs56378716-G (*P* = 1.3×10^−9^, β = −0.46; **Figures S11**) – underlie myeloperoxidase (MPO) deficiency^56–58^. The concordant SSC reduction provides genetic confirmation that SSC reflects variation in granule content.

Similarly, the multiple sclerosis risk allele rs11554159-A (p.Arg76Gln) in *IFI30* was associated with decreased monocyte SSC (*P* = 3.13×10^−24^; β = −0.22; **Figure S12**). *IFI30* encodes the lysosomal thiol reductase GILT, which catalyzes disulfide bond reduction during MHC class II antigen processing^59^. The SSC reduction localizes the effect of this allele to the lysosome–granule compartment in monocytes, consistent with perturbed lysosome-dependent immune regulation, a pathway implicated in multiple sclerosis susceptibility^60^.

Collectively, these examples demonstrate how BloodVariome can map disease-associated variants to discrete immune cell states, revealing lineage- and stage-specific mechanisms of human immune-mediated disease.

### BloodVariome as a discovery platform for immune cell biology

In addition to illuminating disease mechanisms, BloodVariome provides a high-resolution view of normal human immune cell biology *in vivo*. Given the complexity of the immune system and the limitations of model systems in recapitulating its dynamics, the dataset enables the identification of novel genes and regulatory mechanisms, as illustrated in the examples below.

#### Frequency traits expose regulators of immune cell development

Analysis of frequency traits recapitulated established regulators of immune cell development while also nominating genes poorly characterized in humans (**Figure 6**).

At *NIBAN3,* the missense variant rs11666267:G>A (p.Ser572Gly) was associated with reduced frequencies of total, transitional, and naïve B cells (*P* = 1.4×10^−12^ to 2.4×10^−8^, β = −0.13 to −0.074; **Figure S13**) and lower IgM levels in GWAS data for 114,697 individuals from Iceland, Sweden, and the UK (*P* = 1.0×10^−12^, β = −0.04; Ali *et al.*, companion manuscript; **Table S7C**). NIBAN3 was originally identified as a membrane protein on chronic lymphocytic leukemia cells^61^, but its role in normal B cell biology remains unclear.

Functional studies in mice show that *Niban3* deletion enhances B cell proliferation and IgM/IgG3 production via reduced apoptosis, whereas overexpression promotes apoptosis^62,63^. Consistent with these observations, the rs11666267-A allele was associated with reduced naïve B-cell abundance in humans, genetically implicating *NIBAN3* as a regulator of B-cell homeostasis and suggesting a gain-of-function model at this locus.

#### Cis- and trans-genetic control of CD141 in dendritic cells

Protein expression traits enable genetic dissection of immune receptor regulation at cell-subpopulation resolution. In total, we identified 64 *cis*- and 205 *trans*-associations affecting 28 distinct proteins (**Table S10**), most of them previously uncharacterized.

CD141 provides a notable example. This receptor, encoded by *THBD*, promotes anticoagulation and has been linked to thrombophilia, atypical hemolytic uremic syndrome^64^, and – in dendritic cells – to immune regulation^65^. Sixteen variants influenced surface CD141 expression on dendritic cells (**Figure S14A**; **Table S10**). Nine localized within a 500-kb region surrounding *THBD* and overlapped dendritic cell open chromatin, consistent with *cis*-regulation (**Figure S14B**). The remaining seven mapped to distant loci (*IKZF1*, *IRF8*, *NFKBIA*, *LILRB3-LILRA6-LILRB3-LILRB5*, *NLRP12*, and *NBEAL2*), indicating *trans*- regulation. Mechanistically, *IKZF1* and *IRF8* encode transcription factors that bind the *THBD* promoter and associated *cis*-elements (**Figure S14C**); *NFKBIA* and *NLRP12* modulate NF-κB signaling, which also targets *THBD* (**Figure S14C**); and *NBEAL2* mediates membrane trafficking^66^. Co-expression of the *LILR* family receptors with *THBD* suggests previously unrecognized membrane-level interactions. These findings delineate a multilayered *cis*- and *trans*-regulatory network governing CD141 surface expression and illustrate how BloodVariome can resolve genetic control of immune receptor expression.

#### Endosomal trafficking genes converge on BCR–IgD surface regulation

Immunoglobulin D (IgD) is prominently expressed on naïve and transitional B cells and on a subset of unswitched memory B cells, where membrane IgD forms the antigen-binding component of the B-cell receptor (BCR) together with the signaling heterodimer CD79A and CD79B. Surface IgD levels therefore reflect regulatory mechanisms governing IgD-containing BCR complexes in these populations^67^. Engagement of the BCR triggers antigen internalization and endosomal trafficking required for antigen processing and presentation, making tight control of BCR surface abundance essential for B-cell activation^68^. Yet, the genetic mechanisms governing BCR surface homeostasis remain incompletely understood.

Ten variants influenced IgD expression on naïve and transitional B cells (**Figure S15A**; **Table S10**). Notably, five mapped to genes implicated in endocytosis and endosome sorting (*SNX8*, *SNX5-RRBP1, VPS35L*, and *CCDC32*), and several were also associated with circulating Ig levels in our GWAS of 114,697 individuals (Ali *et al.*, companion manuscript), indicating broader effects on B-cell function (**Figure S15B**).

The signal with the largest effect size, rs144787122-G (β = −0.78; *P* = 1.9×10^−10^ for IgD expression; AF = 0.52%), encodes an Ile414Thr substitution in *SNX8* (sorting nexin 8; **Figure 7A**). Sorting nexins regulate endosomal protein trafficking, and SNX8 contains a BAR domain mediating homodimerization and endosomal tubulation^69^. The Ile414Thr substitution lies within this domain, is predicted to be deleterious (AlphaMissense = 0.771; CADD = 24.4), and was associated with marked reductions in surface IgD and serum IgG levels (**Figure 7B** and **S15B**), consistent with impaired SNX8-mediated trafficking (**Figure 7C**). *SNX8* is highly expressed in B cells (**Figure 7D**) but has not previously been implicated in BCR-IgD regulation. A second, independent non-coding variant at *SNX8* also affected IgD levels on naïve and transitional B cells (rs35642219, *P* = 1.4×10^−9^, β = 0.23; **Figure S15A**).

**Figure 7:**
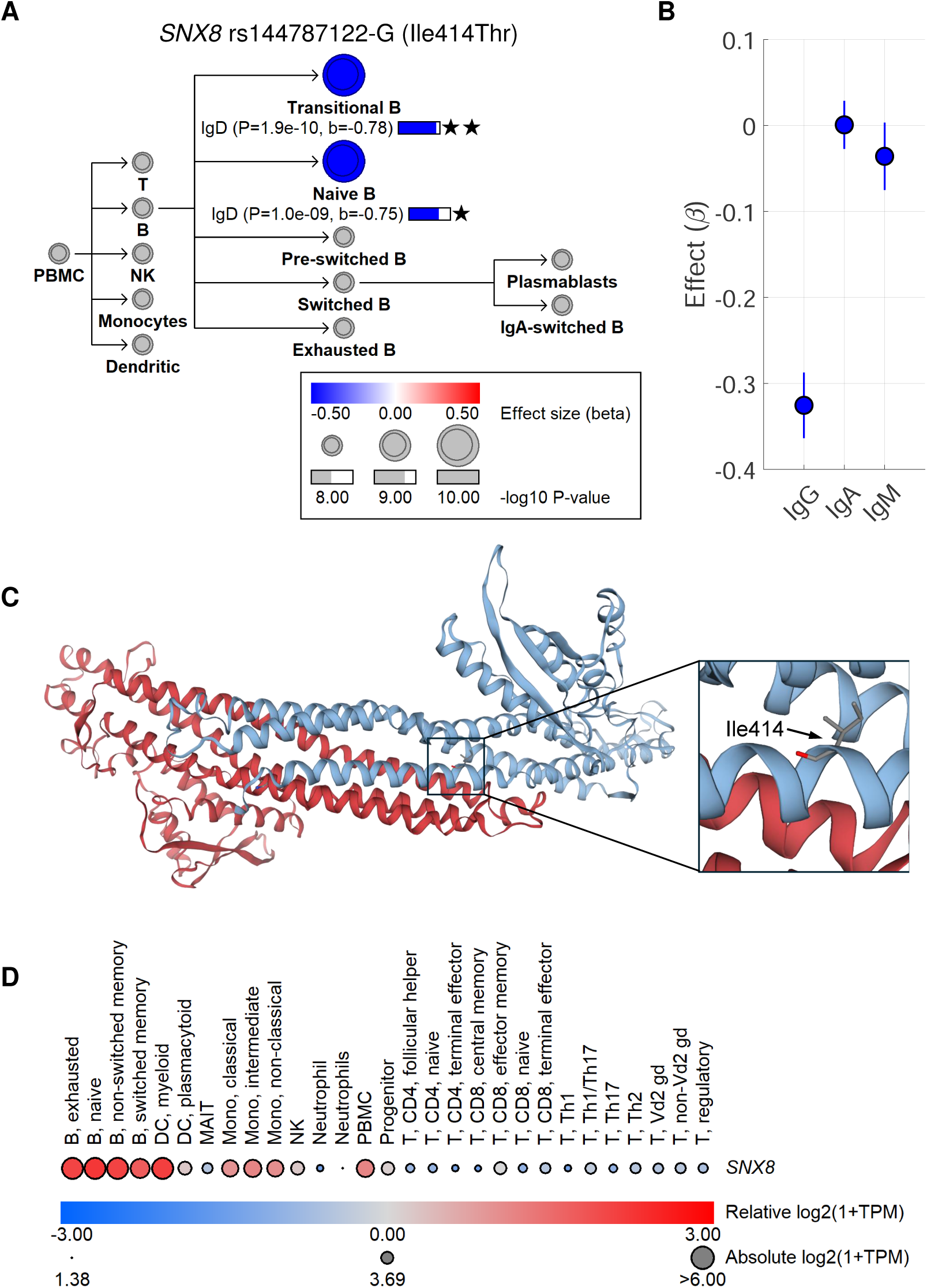
**A rare *SNX8* missense variant is associated with reduced IgD expression in early B cells.** We identified ten variants associated with IgD expression on naïve and transitional B cells (**Figure S17**). The strongest effect was observed for the predicted deleterious missense variant rs144787122-G in *SNX8*. **(B)** Hierarchy plot showing cellular phenotypes associated with rs144787122-G. The illustration is restricted to the B-cell lineage because no significant effects were detected in other lineages. **(C)** Allelic boxplots showing the effect of rs144787122 effect on immunoglobulin levels, including a marked reduction in circulating IgG levels. Data are derived from a large-scale GWAS on IgA, IgG, IgM and six composite Ig traits in 114,697 individuals from Iceland, Sweden, and the UK Biobank (Ali *et al.*, companion manuscript). **(D)** Three-dimensional model of SNX8 highlighting the residue altered by the Ile414Thr substitution. The variant maps to BAR domain of SNX8, which mediates homodimerization and membrane interactions. **(E)** Expression of *SNX8* across hematopoietic cell types in mRNA-sequencing data^96^, showing preferential expression in the B-cell lineage.

Additional signals converged on related trafficking genes. rs2745851-A maps to an intergenic region between *SNX5* and *RRBP1* and upregulates *RRBP1*, an endoplasmic reticulum trafficking factor (**Figure S15C**)^70,71^. rs150300279-T encodes a missense variant in *VPS35L* (p.Arg905Cys), a component of the Retriever complex that cooperates with sorting nexins^68^; rare *VPS35L* loss-of-function variants cause Ritscher-Schinzel syndrome with hypogammaglobulinemia^72^. rs17747633-G downregulates *CCDC32*, a mediator of clathrin-mediated endocytosis^73–75^ (**Figure S15C**) and is accompanied by reduced plasma IgM and surface IgA levels on switched B cells (**Figure S15A-B**).

Collectively, these findings identify endosomal trafficking as a central genetic control axis governing BCR surface homeostasis in human B cells. Notably, we did not observe a comparable convergence of trafficking genes for other proteins measured in our flow cytometry panel. The strong concomitant effects of the rare *SNX8* variant on both surface IgD and circulating IgG levels further suggest that *SNX8* may represent an unrecognized gene contributing to humoral immunodeficiency in humans.

## DISCUSSION

In this study, we introduce BloodVariome, a population-scale atlas of inherited genetic effects across the human immune cell hierarchy. By combining deep flow cytometry with automated pattern-recognition gating, we quantified 1,533 traits spanning 127 immune cell populations in nearly 12,000 individuals. The resource complements existing efforts to annotate genetic variation through effects on gene expression (*e.g*., GTEx and the eQTL Catalog)^76,77^, plasma protein abundance (*e.g.*, UK Biobank Olink)^34–37^, and epigenomic features (*e.g.*, ENCODE)^78–81^ by adding an immune cell phenotypic layer.

Previous studies of hematological and immunological variation have largely relied on bulk blood cell measurements derived from electronic health records^6-8^, whereas high-resolution flow cytometry studies have been limited to comparatively modest cohorts^9-14^.

BloodVariome bridges this gap by combining population-scale statistical power with detailed cellular phenotyping, thereby revealing genetic effects that are largely obscured in bulk blood trait analyses. This combination of scale and phenotypic depth was only possible because we developed high-throughput flow cytometry workflows and bespoke software for automated gating. The latter enabled analysis of the vast amounts of complex flow cytometry data at population scale, something that would have been impossible with manual gating. The framework we have developed thus enables deep immunophenotyping in arbitrary large sample sizes, exposing a previously inaccessible landscape of human phenotypic variation to association studies.

A central finding is that genetic regulation of immune cell phenotypes is highly compartmentalized. The vast majority of variants act within single lineages and often on individual cellular phenotypes, indicating that inherited variation perturbs discrete immune cell states rather than broadly altering hematopoiesis. This compartmentalization has important implications for interpreting disease mechanisms. Many variants identified in BloodVariome overlap previously reported risk alleles for human diseases. By linking these alleles to specific immune cell phenotypes, BloodVariome illuminates candidate cellular mechanisms underlying predisposition to a broad range of diseases, including autoimmunity, immunodeficiency, and hematologic malignancy.

Several limitations should be noted. Despite broad coverage of lineages and subpopulations, the full diversity of immune cell states cannot be captured with a finite marker panel. Our capacity to measure some traits – particularly protein expression and morphology traits and rare cell frequencies – increased during the study, resulting in lower sample sizes for later-added phenotypes. Although the current cohort provides substantial statistical power, further expansion will be required to capture additional associations, including rare-variant effects. Finally, the present cohort is limited to adults of Swedish ancestry. Future expansions of BloodVariome will therefore prioritize increased cohort size, broader ancestry representation, and wider age coverage. As the dataset grows, it will become an increasingly powerful resource for immunogenetics, enabling detailed dissection of the genetic architecture underlying immune cell regulation, disease susceptibility, and variation in immune function across human populations.

## ACKNOWLEDGEMENTS

This work was supported by grants from the European Research Council (CoG-770992 BloodVariome and EU-MSCA-COFUND 754299 CanFaster), the Knut and Alice Wallenberg Foundation (2014.0071 and 2017.0436), the Swedish Research Council (2017-02023, 2018-00424, and 2024-02765), the Swedish Cancer Society (20.0694 and 23-2851), the Swedish Children’s Cancer Fund (PR2018-0118 and PR2023-0067), ARMEC Lindeberg’s Foundation, Inga-Britt and Arne Lundberg’s Foundation (2017-0055), and Region Skåne (ALF 2022-0363). We thank the personnel at Clinical Chemistry and Clinical Immunology and Transfusion Medicine for their assistance with sample collection. We are indebted to the blood donors and primary care patients who participated in the study.

## AUTHOR CONTRIBUTIONS

A.L.d.L.P., L.E., G.T., U.T., and B.N. designed the study. A.L.d.L.P., C.C., N.U.D., and Z.A. carried out flow cytometry phenotyping. L.E. and A.L.A. developed gating algorithms. A.L.d.L.P., L.E., C.C., and A.L.A. analyzed flow cytometry data. A.L.d.L.P., L.E., G.T., A.L.A., Z.A., M.P., G.H.H., L.S., M.O., and B.N. carried out bioinformatic and statistical analyses. U.T., K.S., I.J., and T.O. contributed data and analysis resources. A.L.d.L.P., L.E., and B.N. drafted the manuscript. All authors contributed to the final manuscript.

## DATA AVAILABILITY

Summary statistics for the association studies of the nine Ig traits have been deposited in the GWAS Catalog database (https://www.ebi.ac.uk/gwas/; accession number pending). The BloodVariome immune cell data are available at (https://bloodvariome.medfak.lu.se/). Gene expression datasets are available from the NCBI Gene Expression Omnibus repository (https://www.ncbi.nlm.nih.gov/geo/; accession numbers GSE107011, GSE139369)101,102 and the ProteinAtlas (https://www.proteinatlas.org/). Disease annotations were retrieved from the GWAS Catalog (https://www.ebi.ac.uk/gwas/), ClinVar (https://www.ncbi.nlm.nih.gov/clinvar/), OMIM (https://www.omim.org/), IntOGen (https://www.intogen.org/), and the Mitelman Database (https://mitelmandatabase.isb-cgc.org/). The pooled CRISPR-Cas9 screening data for cancer cell lines is available through the DepMap portal (https://depmap.org).

## METHODS

### Study population

Peripheral blood samples (2-6 ml in EDTA tubes) were collected from 7,773 healthy blood donors and 9,158 primary care patients from Scania in southern Sweden. Blood samples from blood donors were collected with informed consent in the Clinical Immunology and Transfusion Medicine department at Skåne University Hospital in Lund and Malmö. Samples from primary care patients were surplus material from the Clinical Chemistry department at Skåne University Hospital in Lund, where sample-processing robots automatically selected samples from primary care patients aged 18 to 71 years. All samples were collected between 07:00 and 19:00 hours. They were annotated with age and irreversibly anonymized prior to transfer to our laboratory. The collection was performed in three stages: Phase I (November 2015 to April 2016), Phase II (January 2017 to November 2017), and Phase III (August 2018 to April 2019). Duplicate samples from repeat donors were identified using our genetic data. After genetic quality control and ancestry filtering, 11,983 unique individuals remained.

The study was approved by the Lund University Ethical Review Board (no. 2018/2) and conducted in accordance with the Declaration of Helsinki.

### Flow cytometry analysis

Red blood cells were lysed using 0.84% ammonium chloride for 10 min at room temperature, then centrifuged at 1,200 rpm for 5 min. The leukocyte pellet was washed twice with PBS containing 4 mM EDTA and incubated with antibody cocktails (**Table S1B**) for 15 min at room temperature. After staining, cells were washed and resuspended in 250 µL of PBS supplemented with 4 mM EDTA and 0.1% BSA for flow cytometric acquisition.

Flow cytometric analyses were performed using three instruments across study phases: a BD FACS Canto II™ (488, 633, and 405 nm lasers; Phase I), a BD LSR Fortessa™ (488, 650, 405, and 561 nm lasers; Phase II), and a BioRad ZE5™ (488, 650, 405, and 561 nm lasers; Phase III). To ensure robust analysis of rare immune cell subsets, high sampling depths were applied (up to 500,000 events per test in Phases I and II; up to 1,000,000 in Phase III; **Table S1D**). All samples were processed within 36 hours of venipuncture under standardized temperature-controlled conditions to preserve marker stability.

### Flow cytometry data analysis

The antibody panels are described in **Table S1B.** In Phases I and II, we used three panels (lineage, T cell, and B cell) to define 40 immune cell subsets (38 core and 2 intermediate populations). In Phase III, we used two panels (one covering B cells, monocytes, NK cells, and dendritic cell populations, and one covering T cell populations) to define 135 immune cell subsets (127 core and 8 intermediate populations; **Table S1C-D**). To enable scalable, operator-independent analysis of high-dimensional flow cytometry data across nearly 12,000 samples and up to 127 immune cell populations per sample (entailing hundreds of thousands of individual gating decisions across the cohort), we developed bespoke pattern-recognition software for automated gating within the AliGater framework^15^ (https://github.com/LudvigEk/AliGater; and https://github.com/LudvigEk/ImmuneCell). All automated gating outputs were subsequently reviewed to verify gate placement and population fidelity. The gating schemes are shown in **Figure S3** and **Table S1C**.

In all phases, we calculated the relative frequency of core populations relative to relevant reference populations, usually the parent and grandparent populations in the gating hierarchy. In Phase III, we also calculated (1) the absolute counts of 53 core populations, defined as the number of cells per unit blood volume (cells/µL); (2) protein expression traits defined as the median fluorescence intensity, MFI, of antibody targets expressed on core populations; (3) morphology traits defined as the forward and side scatter in core populations (**Table S2A**). Study phase was included as a covariate in all downstream genetic analyses, and reproducibility of all traits was verified prior to association testing (**Table S2B**).

*Summary of gating in Phase I and II:* PBMCs were defined and doublets excluded based on forward and side scatter (FSC/SSC).

In the lineage panel, CD45^+^ single PBMCs were divided into T cells (CD3^+^) and non-T cells (CD3^−^). T cells were classified as helper (CD4^+^) or cytotoxic (CD8^+^). Among non-T, B cells were defined as CD19^+^, monocytes as CD14^+^ within the CD19^−^ fraction, and NK cells as CD16^+^ and/or CD56^+^ within the CD19^−^14^−^ fraction.

In the B cell panel, B cells were gated as CD45^+^CD19^+^ and subdivided into naïve, pre-switched memory, switched memory, and exhausted based on IgD and CD27 expression. Transitional B cells were defined as CD38^+^CD24^+^ within the naïve compartment. Within switched memory B cells, IgA^+^ cells and plasmablasts (CD38^+^CD24^+^) were identified.

In the T cell panel, helper (CD3^+^CD4^+^) and cytotoxic (CD3^+^CD4^−^) cells were gated and classified as naïve (CD45RA^+^CD45RO^−^) and memory (CD45RA^−^CD45RO^+^). Regulatory T cells were defined as CD127^−^CD25^+^ and subdivided into resting, secreting, and activated based on CD45RA and CD25 expression. Additionally, CD194^+^ Tregs (CD194^+^CD45RO^+^) were defined. For each T cell population, we also defined the CD39^+^ subset.

*Summary of gating in Phase III:* Peripheral blood mononuclear cells (PBMCs) were defined and doublets excluded based on forward and side scatter (FSC/SSC).

In the B-NK-MONO-DC panel, cells were divided into CD45^+^CD3^+^ and CD45^+^CD3^−^populations. NKT-like cells were defined as CD3^+^CD56^+^. Among CD45^+^CD3^−^ cells, B cells (CD19^+^) were defined and further categorized as naïve, pre-switched memory, switched memory, or exhausted based on IgD and CD27 expression. Transitional B cells were identified as CD38^+^CD24^+^ within the naïve compartment. Within switched memory B cells, IgA^+^ cells and plasmablasts (CD24^−^CD38^+^) were also identified. Among CD19^−^ cells, monocytes were defined as CD14^+^ and subdivided into classical (CD16^−^) or non-classical (CD16^+^). Within the CD14^−^ fraction, NK cells were identified as CD16^+^ and/or CD56^+^ and classified as early (CD16^low^CD56^high^), effector (CD16^high^CD56^high^), or terminal (CD16^high^CD56^low^). Lineage-negative (CD45^+^CD3^−^CD19^−^CD14^−^CD16^−^CD56^−^) HLA-DR^+^ cells were designated as dendritic cells (DC) and classified as plasmocytoid (pDC; CD123^high^CD141^low^), conventional type 1 (cDC1; CD123^low^CD141^high^), or conventional type 2 (cDC2; CD123^low^CD141^low^). For each B cell panel subset, we defined the HLA-DR^+^ subset.

In the T cell panel, CD3^+^ cells were identified and divided into helper (CD4^+^) and cytotoxic (CD8^+^) subsets. Both were further classified as naïve, central memory, effector memory, and TEMRA based on CD45RA and CCR7 expression. Within the CD4^+^ subset, regulatory T cells (Tregs) were identified as CD127⁻CD25⁺ and subdivided into resting (CD45RA^+^CD25^+^), secreting (CD45RA^−^CD25^+^), and activated (CD45RA^−^CD25^high^) subsets. For completeness, we also defined CD194^+^ Tregs and Treg subpopulations. Follicular helper T cells were defined as CXCR5^high^CD127^+^CD25^−^; Th1, Th2, Th17, and Th9 subsets were identified by expression of CD194, CXCR3, and CCR6. For each T cell subpopulation, we also defined the HLA-DR^+^ and CD39^+^ subsets.

### Analysis of the impact of age and sex

The effects of age and sex on immune cell traits were evaluated using multivariable linear regression. Age, sex, collection phase, and donor type were included as covariates. A separate model was fitted for each trait using the MATLAB mvregress function. To avoid inflation due to non-normality, trait values were inverse-normal transformed.

### Genotyping and association analysis

Samples were genotyped using Illumina OmniExpress-24 and Global Screening microarrays. Samples and variants with < 98% yield were excluded. The microarray genotypes were phased in conjunction with 91,727 Swedish samples using Eagle2^82^. Imputation was done using a phased reference panel of whole-genome sequence data from 50,839 European-ancestry individuals, including 3,697 Swedish individuals, sequenced using HiSeqX and NovaSeq PCR-free Illumina technology to a mean depth of at least 30X^83^. Single-nucleotide polymorphisms (SNPs) and small insertions/deletions (INDELs) were called using Graphtyper^84^ and imputed into the phased microarray genotypes.

To ensure a genetically homogeneous Swedish ancestry cohort, we applied a multi-step ancestry filtering strategy. The population structure was analysed using ADMIXTURE v1.23 in supervised mode with 1,000 Genomes populations CEU, CHB, and YRI as training samples and Swedish individuals as test samples^85,86^. Samples with < 0.9 CEU ancestry were excluded. To identify and exclude individuals of Finnish/Saami origin, we included individuals with < 0.3 CHB and < 0.05 YRI ancestry. The remaining samples were projected onto 20 principal components calculated with a European reference panel. UMAP^87^ was used to reduce test sample coordinates to two dimensions. Additional European samples not in the original reference set were also projected onto the principal components and UMAP components to identify ancestries, and samples with Swedish ancestry were identified. The inclusion of Finnish/Saami individuals allowed us to confirm that we could identify a distinct UMAP cluster of individuals with the following properties: (a) elevated CHB ancestry according to ADMIXTURE; (b) enriched for individuals who we knew to have been born in Finland; (c) on principal component 1 and 2, were positioned in the region occupied by Finnish individuals. Those individuals were excluded from the analysis. After filtering, 11,983 unique samples classified as Swedish remained.

Trait values were first adjusted for covariates (sex, age, day of year, study phase, sample type (blood donor vs primary care), and 12 principal components) and subsequently inverse-normal transformed prior to association testing. The trait values were tested for association with 23.6 million genetic variants using generalized linear regression, which estimated the effects of genotypes under an additive model^88^. Linkage disequilibrium score regression was used to account for distributional inflation in the test statistics due to cryptic relatedness and population stratification^23^. To account for multiple testing, we applied a class-based Bonferroni correction, grouping variants by genomic annotation and adjusting the significance threshold for each category based on the number of variants (**Table S4A**)^22^.

Additionally, study-wide thresholds were defined by further adjusting for the effective number of independent traits, estimated at 146 from the phenotype correlation matrix using the Kaiser-Guttman criterion (number of principal components with eigenvalue > 1).

Stepwise conditional analysis was used to identify independent signals in each associated region, both for all the traits jointly and for the traits grouped into three sets: (i) cell frequency traits, (ii) surface protein expression traits, and (iii) cell morphology traits. The genome was divided into regions of genome-wide association signals for any trait in the set of included traits, with each region separated by at least 1 Mb. Associations for all traits were then adjusted for the strongest association signal in the region. If, after conditioning on the lead signal, any trait in the set still had a significant association in the region, the process was repeated, adjusting both signals. This was repeated until no trait in the set had a significant association in the region. Finally, all identified signals were adjusted for all the other signals.

To define the 95% credible set of plausible causal variants for each association signal, we applied a Bayesian refinement approach^89^ to the trait with the most significant lead variant.

### Heritability estimation

To calculate the total variance explained (*h*^2^) by the identified variants, we used the formula *h*^2^ = ∑ 2 × β_i_^2^ × MAF_i_ × (1 − MAF_i_), where MAF_i_ and β_i_ denote the minor allele frequency and effect size for the *i*-th lead variant. Total SNP heritability was estimated using linkage disequilibrium score regression^23^ with the HLA region excluded. The *h*^2^ of the 20 independent variants identified in the HLA region were then added to the LDSC estimate.

### Expression quantitative locus analysis

To functionally annotate associated variants, we integrated multiple expression quantitative locus (eQTL) datasets for immune cells: (1) RNA-sequencing data for 28 sorted immune cell populations from 416 individuals in the ImmuNexUT compendium (https://www.immunexut.org/)^76^; (2) RNA-sequencing data of cells isolated from peripheral blood with negative selection using magnetic beads from Icelanders^90^: CD19^+^ B cells (n = 758 individuals), CD4^+^ T cells (n = 837), CD8^+^ T cells (n = 807), CD14^+^ monocytes (n = 884), neutrophils (n = 730), and whole blood (n = 17,848); (3) RNA-sequencing data for 198 immune-relevant subsets of the eQTL Catalogue (https://www.ebi.ac.uk/eqtl/)^91^; and (4) RNA-sequencing and microarray for whole blood in eQTLGen (https://www.eqtlgen.org/)^92^. Colocalization analysis using coloc^93^ was performed for ImmuNexUT and eQTL Catalogue signals (posterior colocalization probability > 0.7). For the Icelandic and eQTLGen datasets, we considered correlated eQTLs (*r*^2^ > 0.8 between BloodVariome and eQTL lead variants).

### Splicing quantitative locus analysis

To assess the effects of genetic variants on alternative splicing (sQTLs), we analyzed RNA sequencing data from whole blood collected from 17,848 Icelanders^90^. Alternative splicing was quantified using an algorithm based on LeafCutter^90^. To reduce the likelihood of false-positive associations, we curated significant lead sQTLs in high linkage disequilibrium (*r*² > 0.8) with lead variants from BloodVariome based on read coverage across the alternatively spliced gene, stratified by variant genotype, and splice-junction graphs depicting the connectivity between junctions used to quantify alternative splicing ratios.

### Plasma protein quantitative trait analysis

To detect effects of genetic variants on plasma protein levels (protein quantitative trait loci, pQTLs), we used two datasets measuring proteins with 4,807 aptamers in plasma samples from 36,136 Icelanders with SomaScan v4 and 2,941 immunoassays using the Olink Explore 3,072 in plasma samples from 54,265 participants in the UK Biobank^34–37^. A BloodVariome lead variant was assigned to a pQTL if the variant, or a highly correlated variant (*r*^2^ > 0.8), was identified as a pQTL in either dataset.

### Plasma immunoglobulin quantitative trait analysis

To annotate variants with effects on immunoglobulin (Ig) levels, we used GWAS data on IgA, IgG, IgM, and six composite Ig traits, for 114,697 individuals from Iceland, Sweden, and the UK Biobank, generated in a parallel project (Ali *et al.*, companion manuscript).

### ATAC-sequencing vs mRNA-sequencing correlation analysis

To identify correlations between chromatin accessibility and gene expression across immune cell types, we used mRNA- and ATAC-seq data for 144 samples from 25 sorted blood cell types (Gene Expression Omnibus accession no. GSE118165; GSE118189)^94^. For ATAC-seq peaks within 200 bp of credible set variants, we identified all genes within a 500-kb window to generate peak-gene pairs. For each pair, we calculated the Pearson correlation coefficient between log-normalized RNA-seq and ATAC-seq transcript counts across cell types. After Bonferroni correction, we mapped genes whose expression correlated significantly with ATAC-seq peak intensity to credible set variants proximal to the ATAC-seq peaks.

### Analysis of candidate gene expression across tissues

To test for enrichment of candidate genes in tissues, we used the ProteinAtlas single-cell type dataset, which encompasses mRNA sequencing data for 81 cell types across a broad range of human tissues, including the immune cell lineages analyzed in our study. Using a one-sided Student’s t-test, we compared normalized expression values for candidate vs other genes in the genome. To annotate candidate genes with expression across blood and immune cells at high cell type resolution, we used publicly available mRNA-sequencing data from Monaco *et al.* and Granja *et al.* (Gene Expression Omnibus accession no. GSE107011 and GSE139369)^95,96^.

### Candidate gene assignment

Within each credible set, we assigned candidate genes using the following criteria, in order of priority: (1) the gene encodes an antibody target whose expression on the surface of an immune cell population is affected by the variant (*i.e.*, the variant is a surface *cis*-pQTL); (2) the gene product is known to interact with, or regulate the expression of, an antibody target whose expression on the surface of an immune cell population is affected by the variant (*i.e.*, the variant is a surface *trans*-pQTL). For example, we assigned *CD247* – encoding the T cell receptor ζ chain – as a candidate gene for variants at 1q24.2 influencing CD3 surface expression, and *HNRNPLL* – encoding a CD45RA splicing factor^97^ – for variants at 2p22.1 influencing CD45RA surface expression; (3) the credible set overlaps a non-synonymous coding variant with fine-mapping posterior probability > 0.1; (4) the credible set overlaps an eQTL or an sQTL in sorted immune cells or whole blood, with priority given to eQTLs rather than sQTLs and to effects in sorted immune cells rather than whole blood. If no gene could be identified based on these data, we considered plasma pQTLs and correlations between RNA-sequencing and ATAC-sequencing signals across blood cell types. If multiple genes were identified, we considered statistical significance and relevance to the associated immune cell trait. If no gene could be identified genetically, we assigned the closest relevant gene. In unclear cases – typically association signals spanning multiple genes, none of which could be assigned genetically – we did not assign a candidate gene, nor did we do so in the HLA and T cell receptor gamma regions due to the complexity of these regions.

### Additional annotation of candidate genes

To identify immunological knockout mouse phenotypes, we used the Mouse Phenome Database (https://phenome.jax.org/; phenotype clades MP:0005387 and MP:0005397)^98^. To find links to Mendelian disorders characterized by abnormal immune cell function, we used OMIM (https://www.omim.org/)^26^. To investigate the overlap with somatically mutated genes across tumor types, we considered genes listed as driver genes in IntOGen (https://www.intogen.org) and fusion gene partners that were recurrently reported (five or more times) in the Mitelman database^28^ (https://mitelmandatabase.isb-cgc.org). To identify effects of gene knockdown on the growth of cell lines derived from immune cell malignancies, we used the Dependency Map version 24Q4 (https://depmap.org)^29^. To annotate the *THBD* locus (**Figure S16B**), we used public chromatin immunoprecipitation sequencing data for three transcription factors in cell lines from the Gene Expression Omnibus (GSE129618, sample GSM3717130 for NFKB; GSE123872, sample GSM3514948 for IRF8) and from ENCODE (experiment ENCFF012ZPU for IKZF1). Genomic coordinates were harmonized to hg38.

### Pleiotropy analysis

To assess genetic overlap between the identified variants and previously reported variants associated with diseases and other quantitative traits, we queried the GWAS catalog using the LDTrait webtool (https://ldlink.nci.nih.gov/?tab=ldtrait), with default settings and a linkage disequilibrium threshold of *r*^2^ > 0.8 in Europeans^99,100^.

### Protein modeling

The three-dimensional model of SNX8 (UniProt identifier Q9Y5X2) was created using the SWISS-MODEL webtool (https://swissmodel.expasy.org)^101,102^.

