## Supplementary material for "BloodVariome: a high-resolution atlas of inherited genetic effects in human immune cells": Figure S1-S11

Age distribution in the study population.

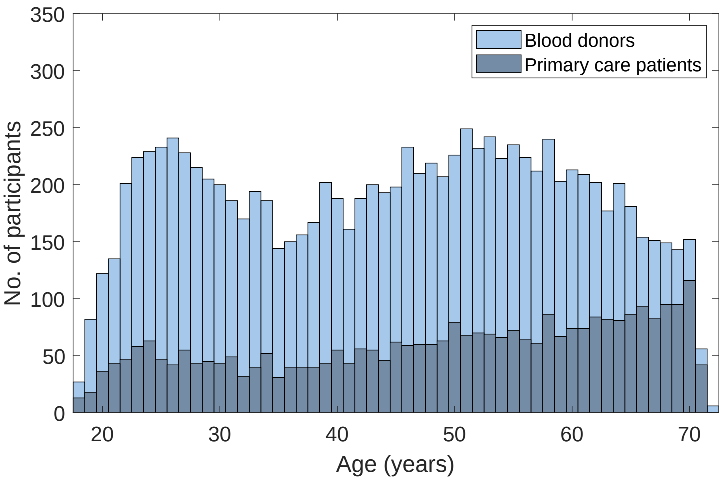

#### Figure S2

Representative gating images generated using AliGater, illustrating sequential gating across immune cell populations. The label above each plot indicates the parent population used for the current gating step. Red lines represent gating boundaries that distinguish the target population from the remaining events. For HLA-DR and CD39, the gating thresholds were fixed for all samples. The absence of a red line in the plot indicates that there were no positive events for the corresponding population.

**(A)** Gating strategy in Phase I and Phase II. **(B)** Gating strategy for the B-NK-monocyte-dendritic cell panel in Phase III. **(C)** Gating strategy for the T cell panel in Phase III

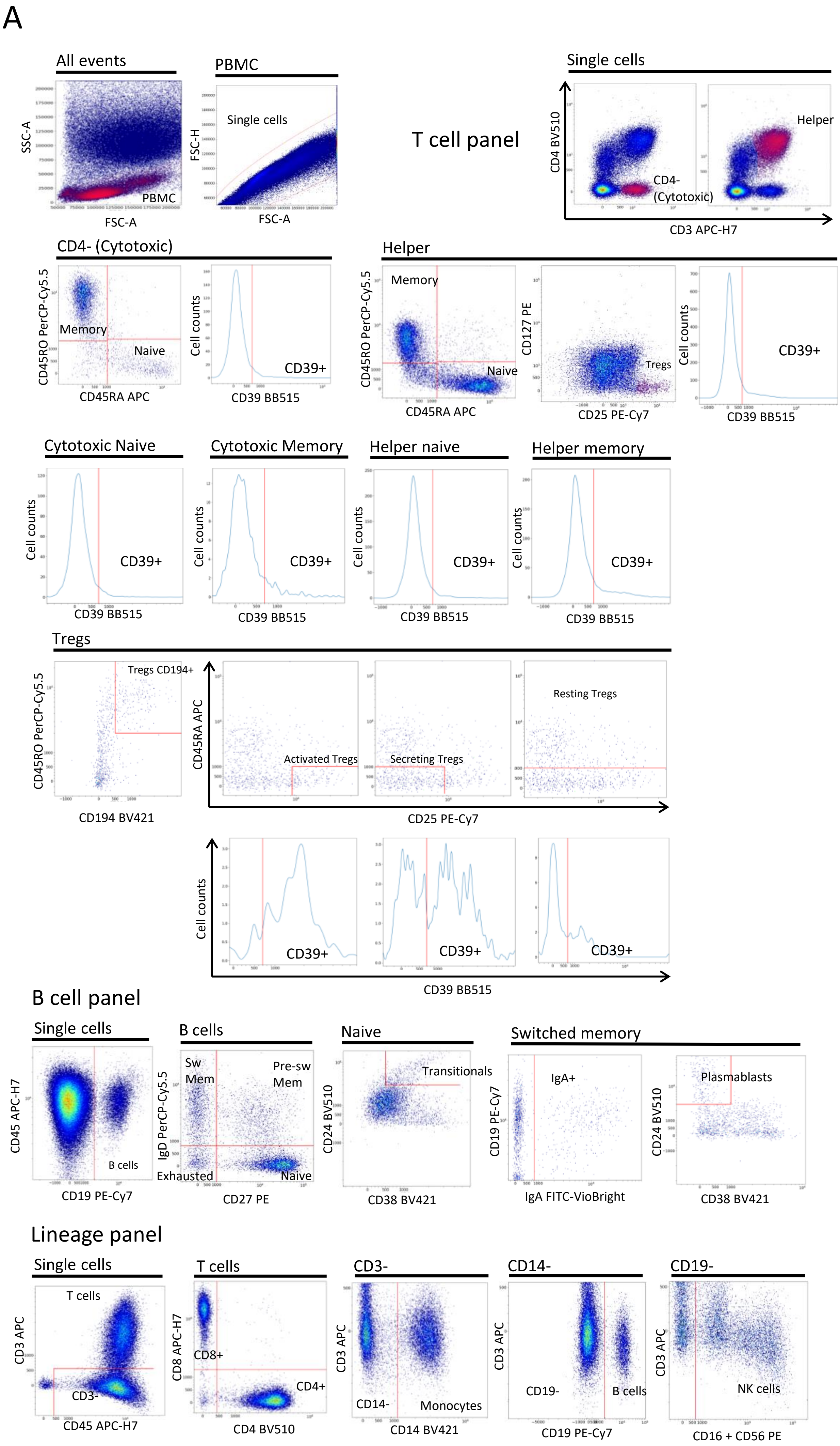
A
All events
SSC-A
FSC-A
PBMC
FSC-H
Single cells
T cell panel
Single cells
CD4 BV510
CD3 APC-H7
CD4-
(Cytotoxic)
Helper
CD4- (Cytotoxic)
CD45RO PerCP-Cy5.5
CD45RA APC
Memory
Naive
Cell counts
CD39 BB515
CD39+
Helper
Memory
Naive
CD127 PE
CD25 PE-Cy7
Tregs
Cytotoxic Naive
Cytotoxic Memory
Helper naive
Helper memory
Tregs
Tregs CD194+
CD194 BV421
CD45RA APC
Activated Tregs
Secreting Tregs
Resting Tregs
CD25 PE-Cy7
B cell panel
Single cells
CD45 APC-H7
CD19 PE-Cy7
B cells
IgD PerCP-Cy5.5
CD27 PE
Sw
Mem
Pre-sw
Mem
Exhausted
Naive
CD24 BV510
CD38 BV421
Transitionals
Switched memory
CD19 PE-Cy7
IgA FITC-VioBright
IgA+
Plasmablasts
Lineage panel
Single cells
T cells
CD3-
CD14-
CD19-
CD3 APC
CD45 APC-H7
CD8 APC-H7
CD4 BV510
CD8+
CD4+
CD14 BV421
Monocytes
CD19 PE-Cy7
B cells
CD16 + CD56 PE
NK cells

B

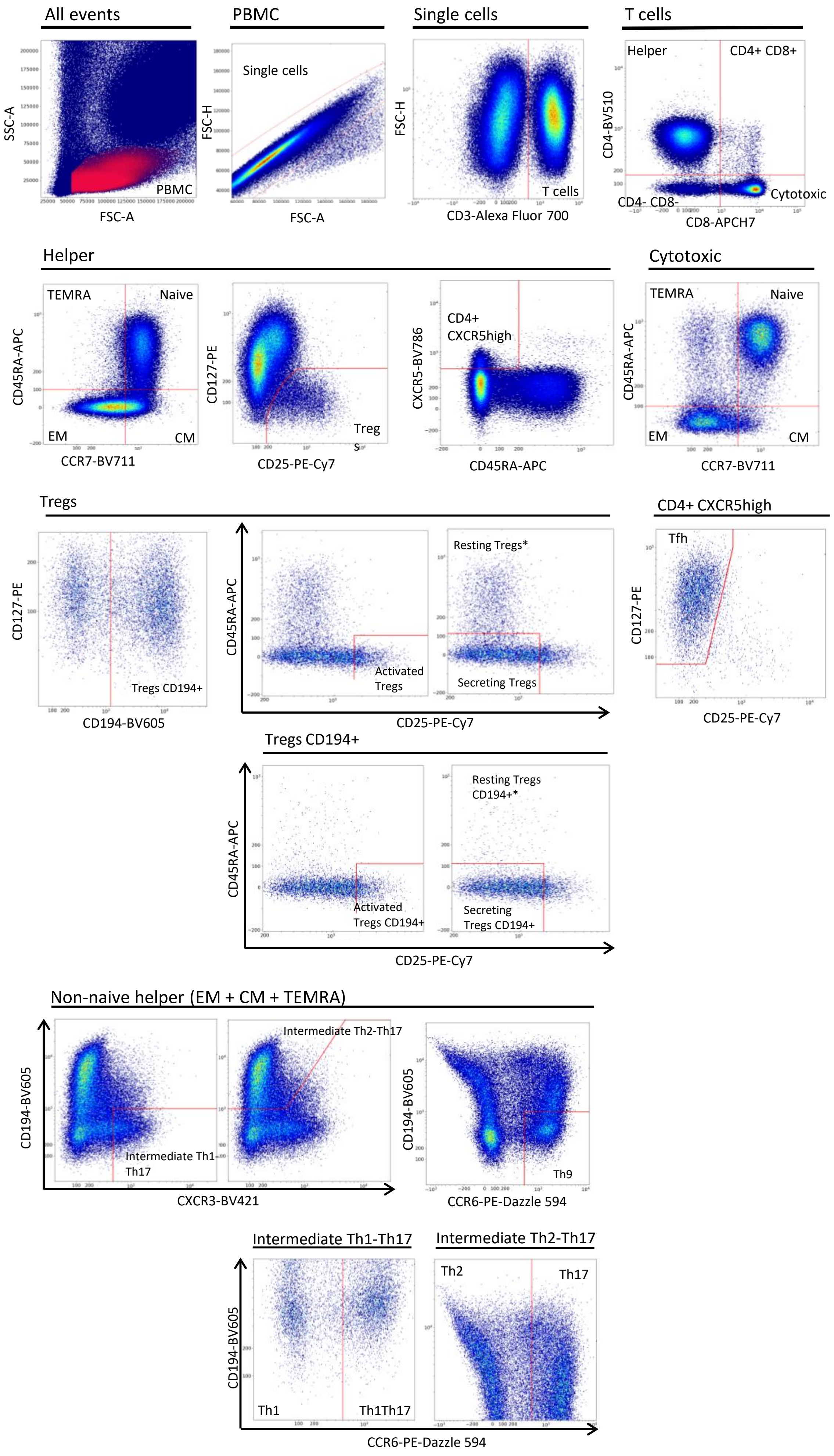
All events
SSC-A
FSC-A
PBMC
Single cells
FSC-H
FSC-A
Single cells
FSC-H
T cells
CD3-Alexa Fluor 700
T cells
Helper
CD4+ CD8+
CD4-BV510
CD4- CD8-
Cytotoxic
CD8-APCH7
Helper
TEMRA
Naive
CD45RA-APC
EM
CM
CCR7-BV711
CD127-PE
Treg
s
CD25-PE-Cy7
CD4+
CXCR5high
CXCR5-BV786
CD45RA-APC
Cytotoxic
TEMRA
Naive
CD45RA-APC
EM
CM
CCR7-BV711
Tregs
CD127-PE
Tregs CD194+
CD194-BV605
CD45RA-APC
Activated
Tregs
Resting Tregs*
Secreting Tregs
CD25-PE-Cy7
CD4+ CXCR5high
Tfh
CD127-PE
CD25-PE-Cy7
Tregs CD194+
CD45RA-APC
Activated
Tregs CD194+
Resting Tregs
CD194+*
Secreting
Tregs CD194+
CD25-PE-Cy7
Non-naive helper (EM + CM + TEMRA)
CD194-BV605
Intermediate Th1-
Th17
Intermediate Th2-Th17
CXCR3-BV421
CD194-BV605
Th9
CCR6-PE-Dazzle 594
Intermediate Th1-Th17
Intermediate Th2-Th17
CD194-BV605
Th1
Th1Th17
Th2
Th17
CCR6-PE-Dazzle 594

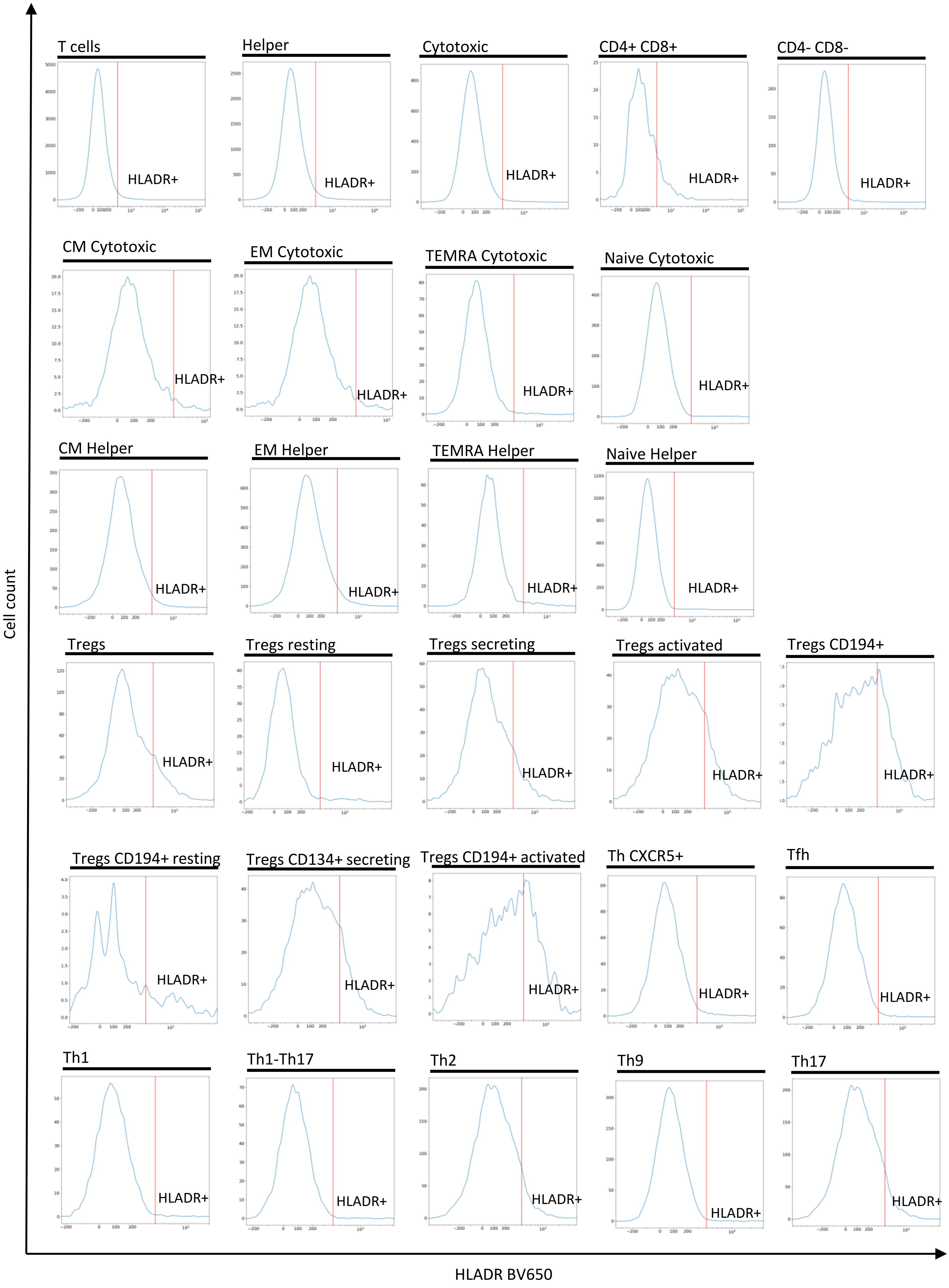
Cell count
T cells
Helper
Cytotoxic
CD4+ CD8+
CD4- CD8-
CM Cytotoxic
EM Cytotoxic
TEMRA Cytotoxic
Naive Cytotoxic
CM Helper
EM Helper
TEMRA Helper
Naive Helper
Tregs
Tregs resting
Tregs secreting
Tregs activated
Tregs CD194+
Tregs CD194+ resting
Tregs CD134+ secreting
Tregs CD194+ activated
Th CXCR5+
Tfh
Th1
Th1-Th17
Th2
Th9
Th17
HLADR+
HLADR BV650

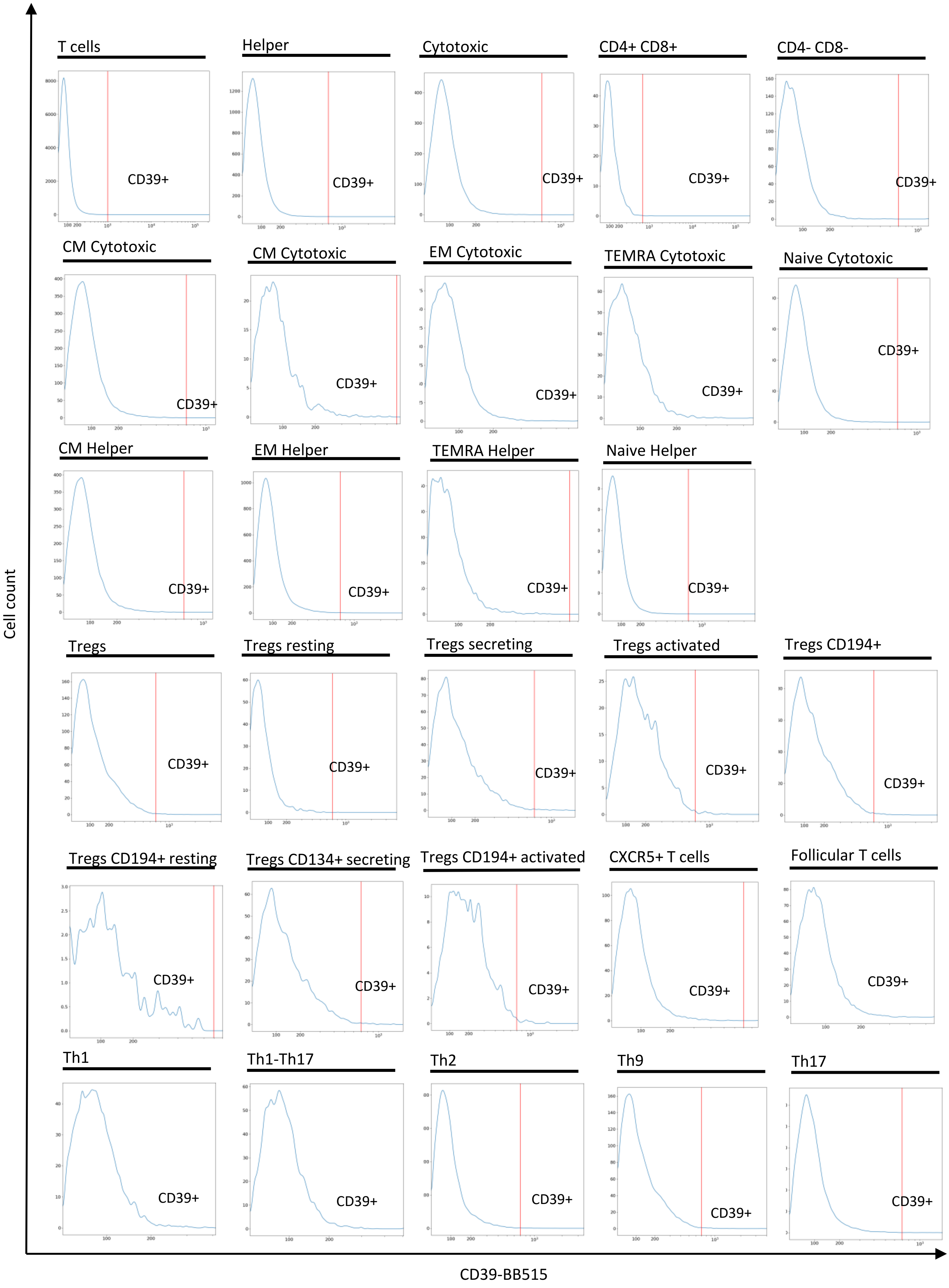

Cell count
CD39-BB515
T cells
Helper
Cytotoxic
CD4+ CD8+
CD4- CD8-
CM Cytotoxic
CM Cytotoxic
EM Cytotoxic
TEMRA Cytotoxic
Naive Cytotoxic
CM Helper
EM Helper
TEMRA Helper
Naive Helper
Tregs
Tregs resting
Tregs secreting
Tregs activated
Tregs CD194+
Tregs CD194+ resting
Tregs CD134+ secreting
Tregs CD194+ activated
CXCR5+ T cells
Follicular T cells
Th1
Th1-Th17
Th2
Th9
Th17
CD39+

C

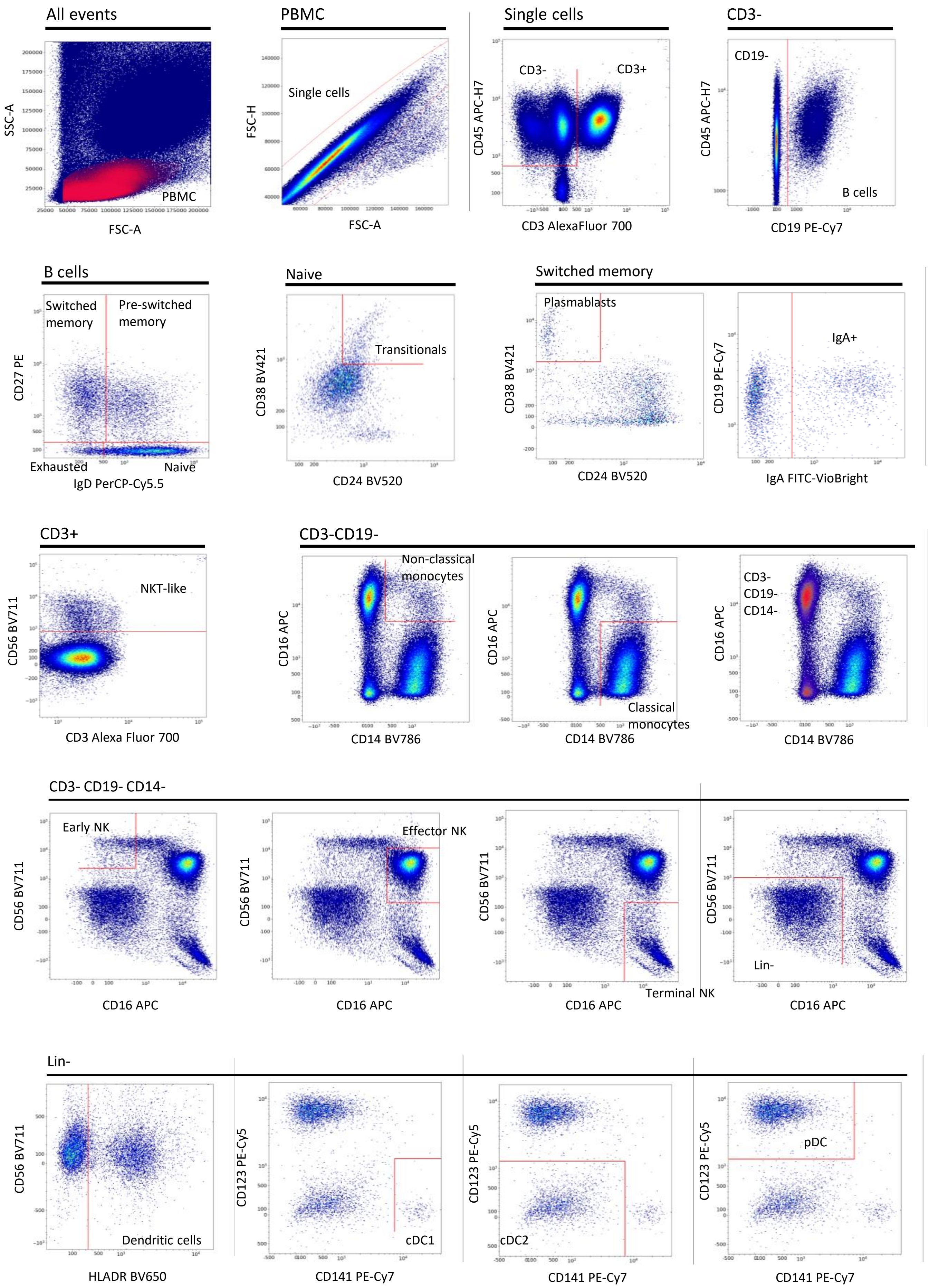
All events
SSC-A
FSC-A
PBMC
PBMC
FSC-H
FSC-A
Single cells
Single cells
CD45 APC-H7
CD3 AlexaFluor 700
CD3-
CD3+
CD3-
CD45 APC-H7
CD19 PE-Cy7
CD19-
B cells
B cells
CD27 PE
IgD PerCP-Cy5.5
Switched memory
Pre-switched memory
Exhausted
Naive
Naive
CD38 BV421
CD24 BV520
Transitionals
Switched memory
CD38 BV421
CD24 BV520
Plasmablasts
CD19 PE-Cy7
IgA FITC-VioBright
IgA+
CD3+
CD56 BV711
CD3 Alexa Fluor 700
NKT-like
CD3-CD19-
CD16 APC
CD14 BV786
Non-classical monocytes
Classical monocytes
CD3-
CD19-
CD14-
CD3- CD19- CD14-
CD56 BV711
CD16 APC
Early NK
Effector NK
Terminal NK
Lin-
Lin-
CD56 BV711
HLADR BV650
Dendritic cells
CD123 PE-Cy5
CD141 PE-Cy7
cDC1
cDC2
pDC

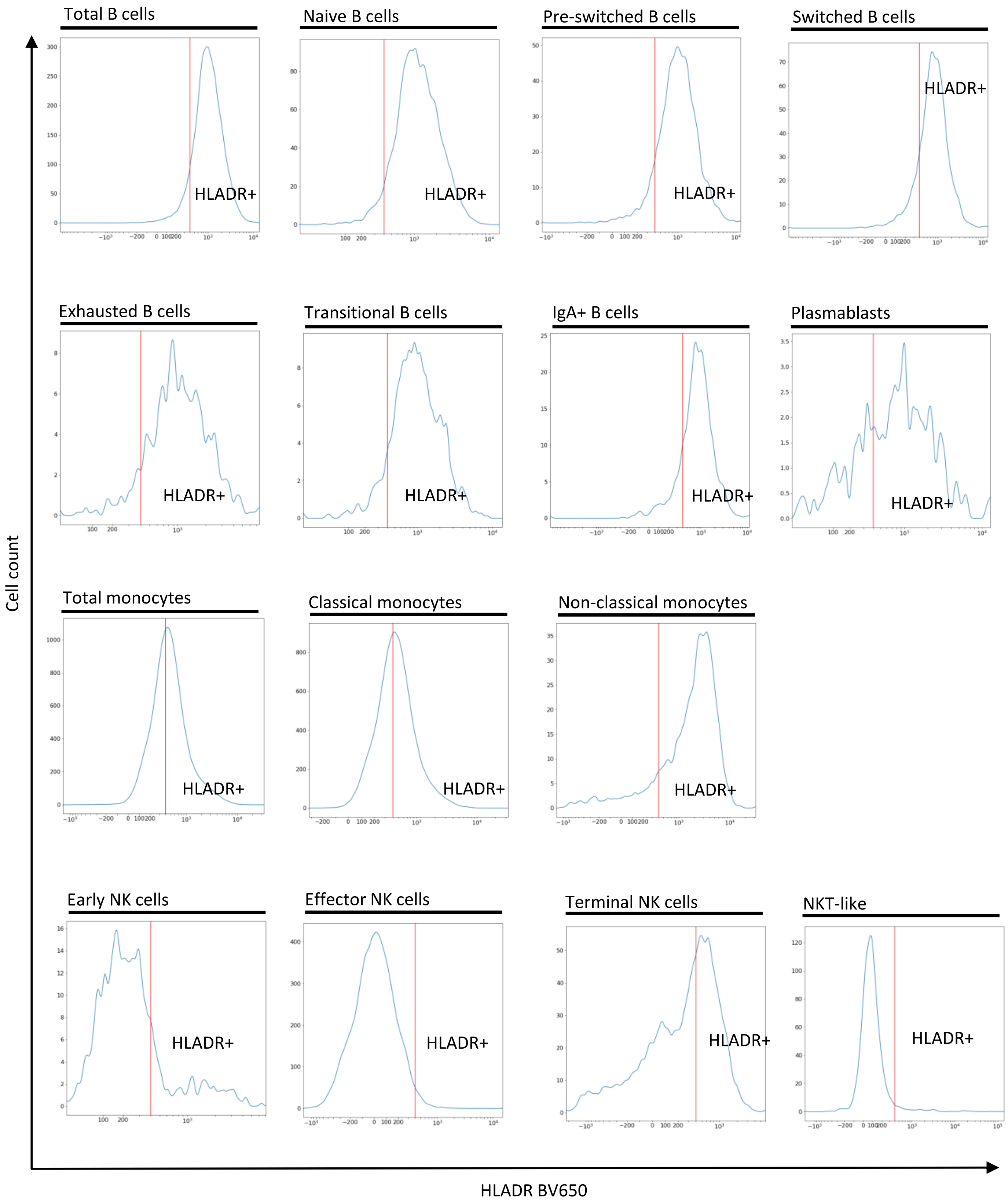

Total B cells
Naive B cells
Pre-switched B cells
Switched B cells
Exhausted B cells
Transitional B cells
IgA+ B cells
Plasmablasts
Total monocytes
Classical monocytes
Non-classical monocytes
Early NK cells
Effector NK cells
Terminal NK cells
NKT-like
HLADR+
Cell count
HLADR BV650

### Figure S3

Heatmaps illustrating correlations between immune cell traits analyzed. **(A)** cell frequency traits; **(B)** protein expression traits; and **(C)** morphology traits. The color scale indicates the magnitude and direction of the Pearson correlation coeffiecients between pairs of traits.

**A**

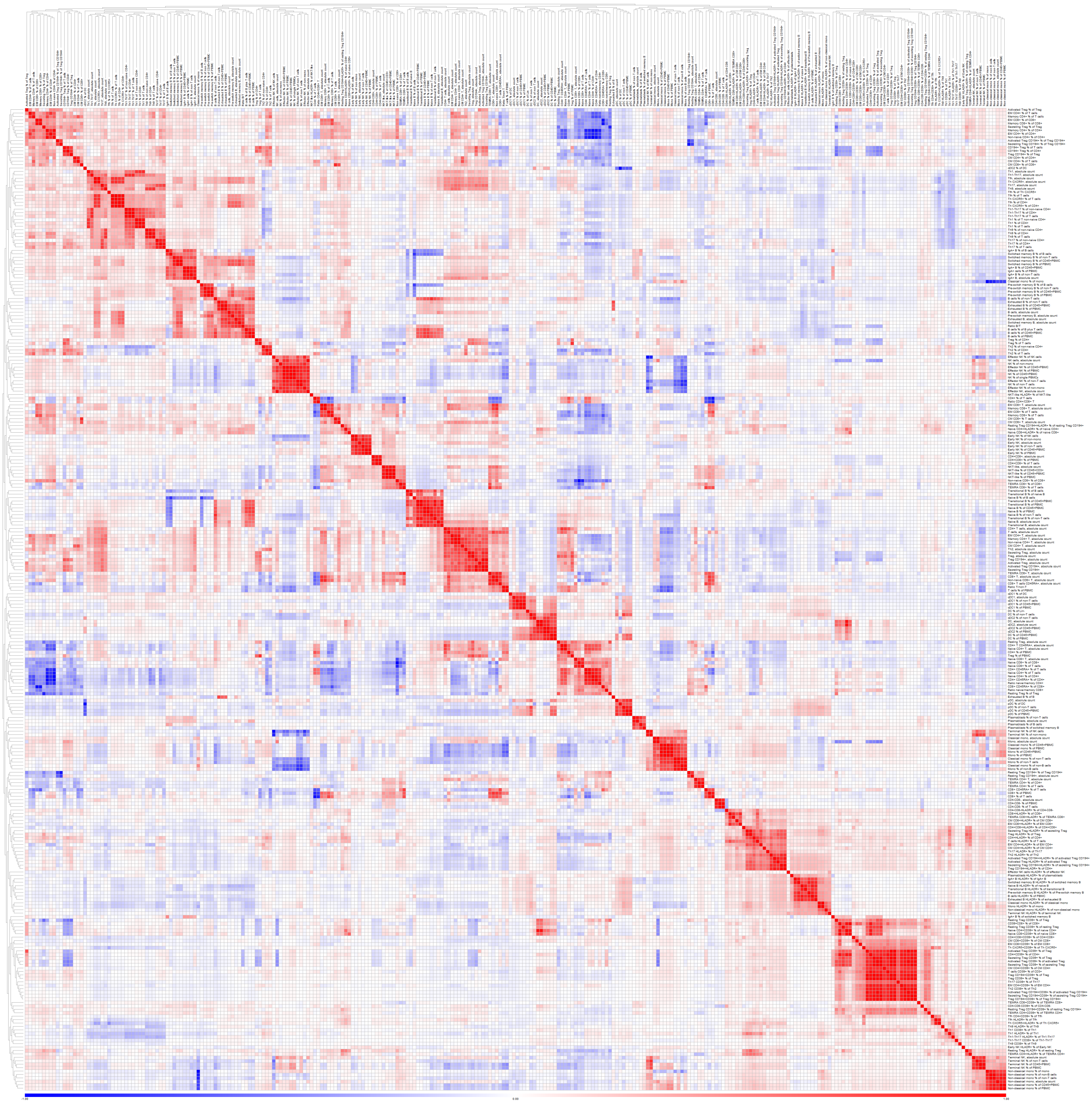

B

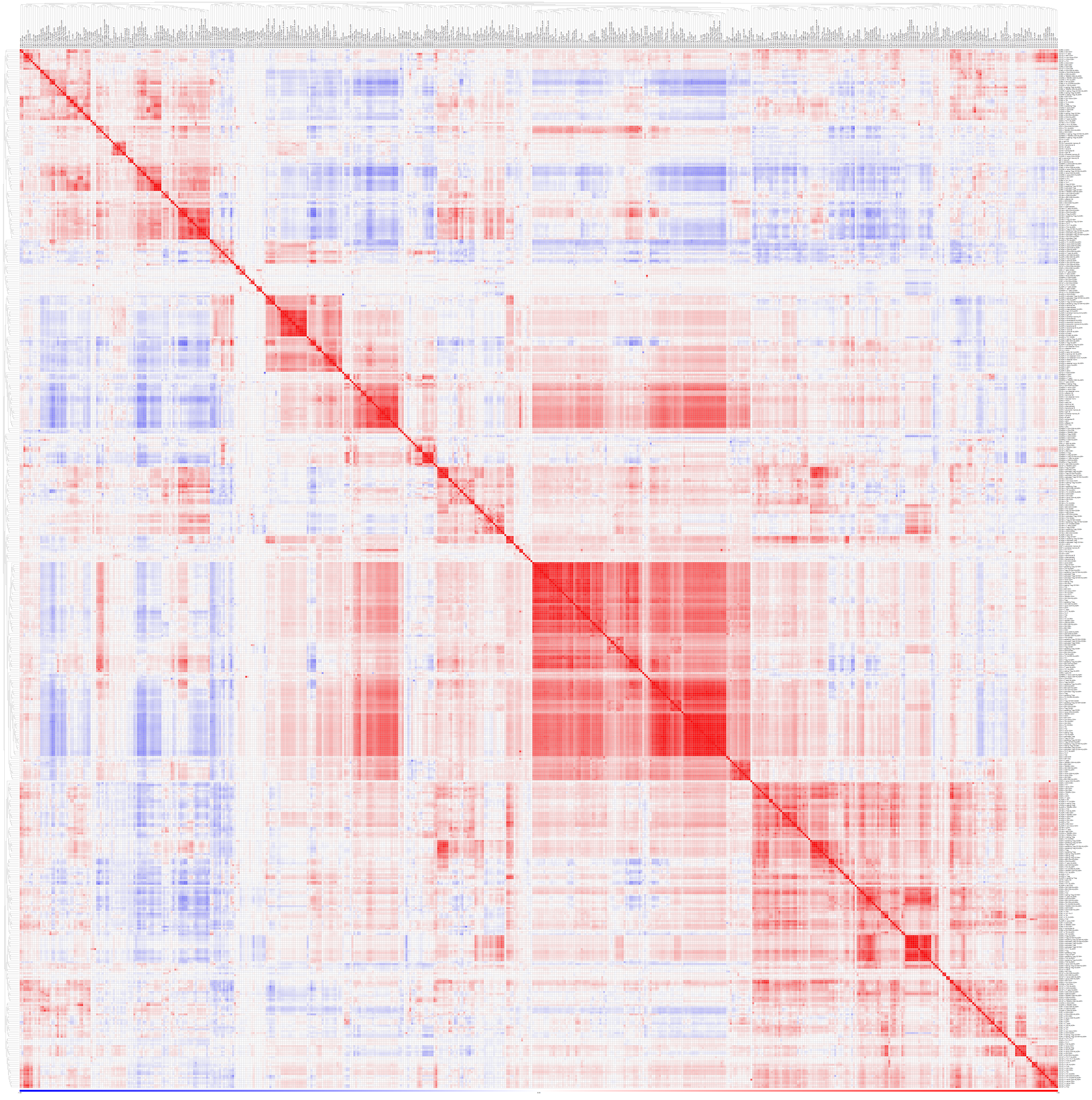

C

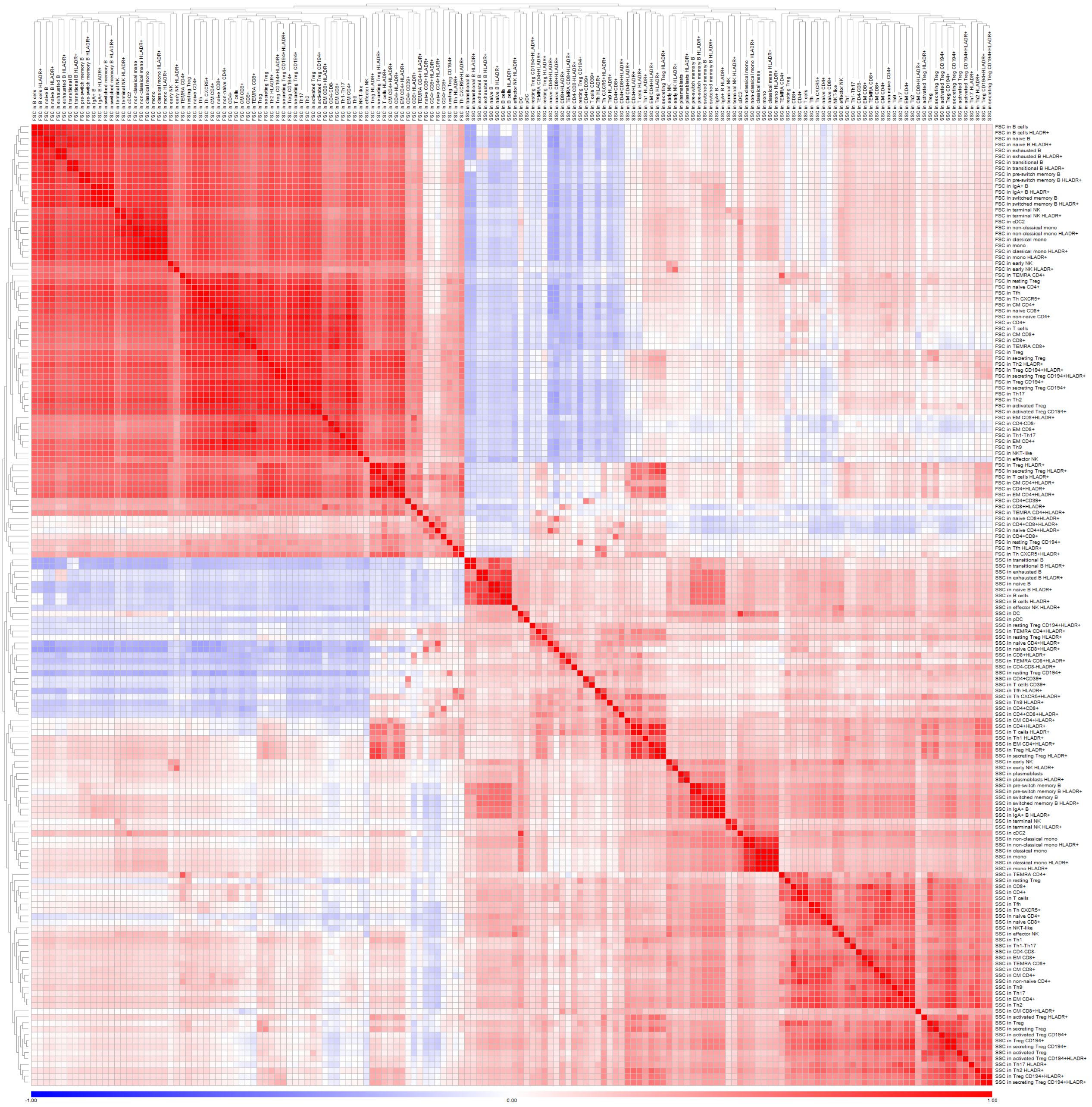

#### Figure S4

Variance explained by genetics, sex, and age for different lineages and trait types. For the sake of visualization, the traits belonging to CD39$^{+}$ and HLADR$^{+}$ subpopulations have been omitted from these plots, but are provided in **Table S6**.

### Frequency traits, T

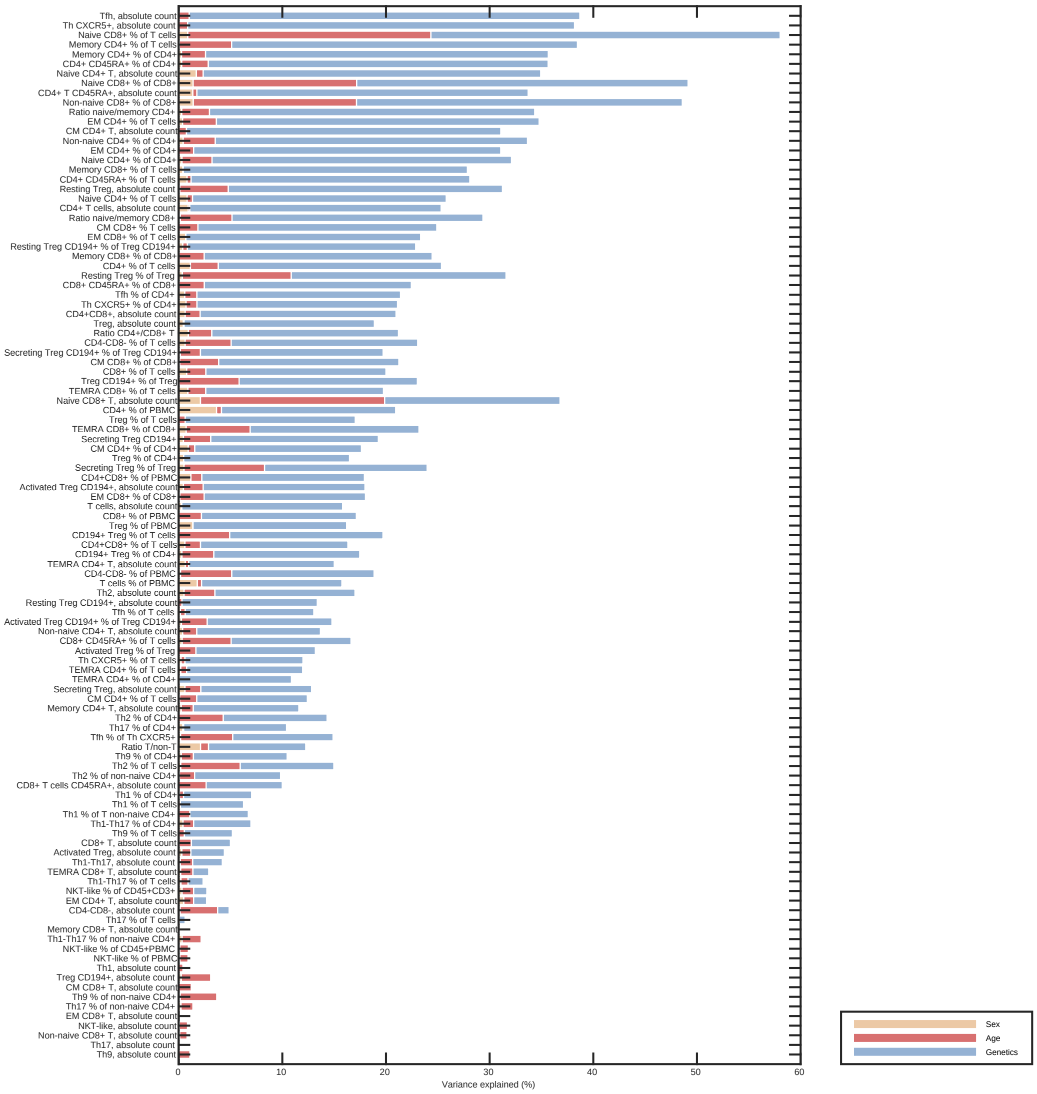

### Frequency traits, B

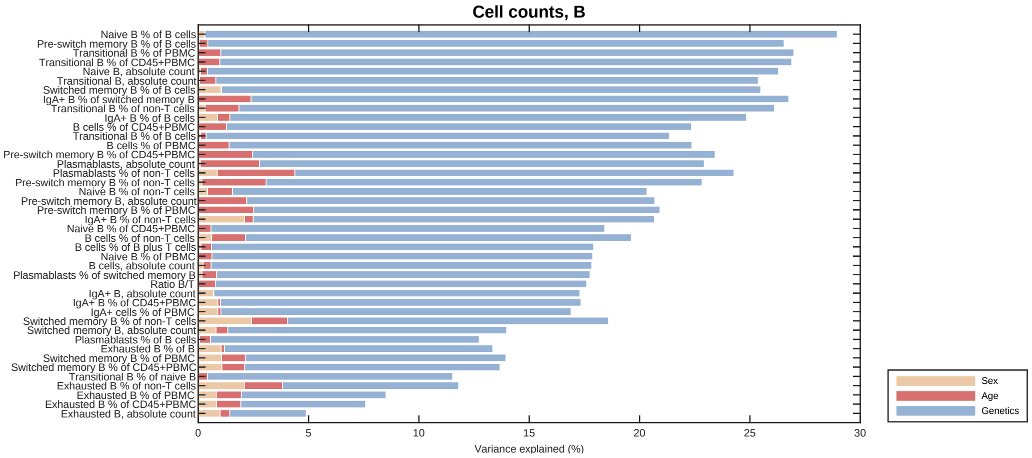


### Frequency traits, MONO-NK-DC

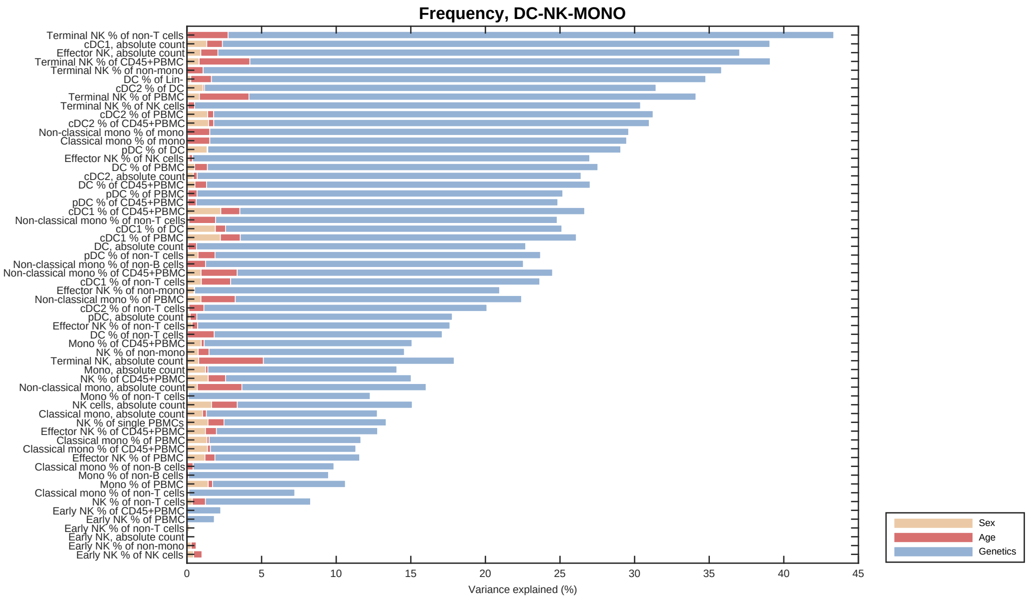

### Protein expression traits, T

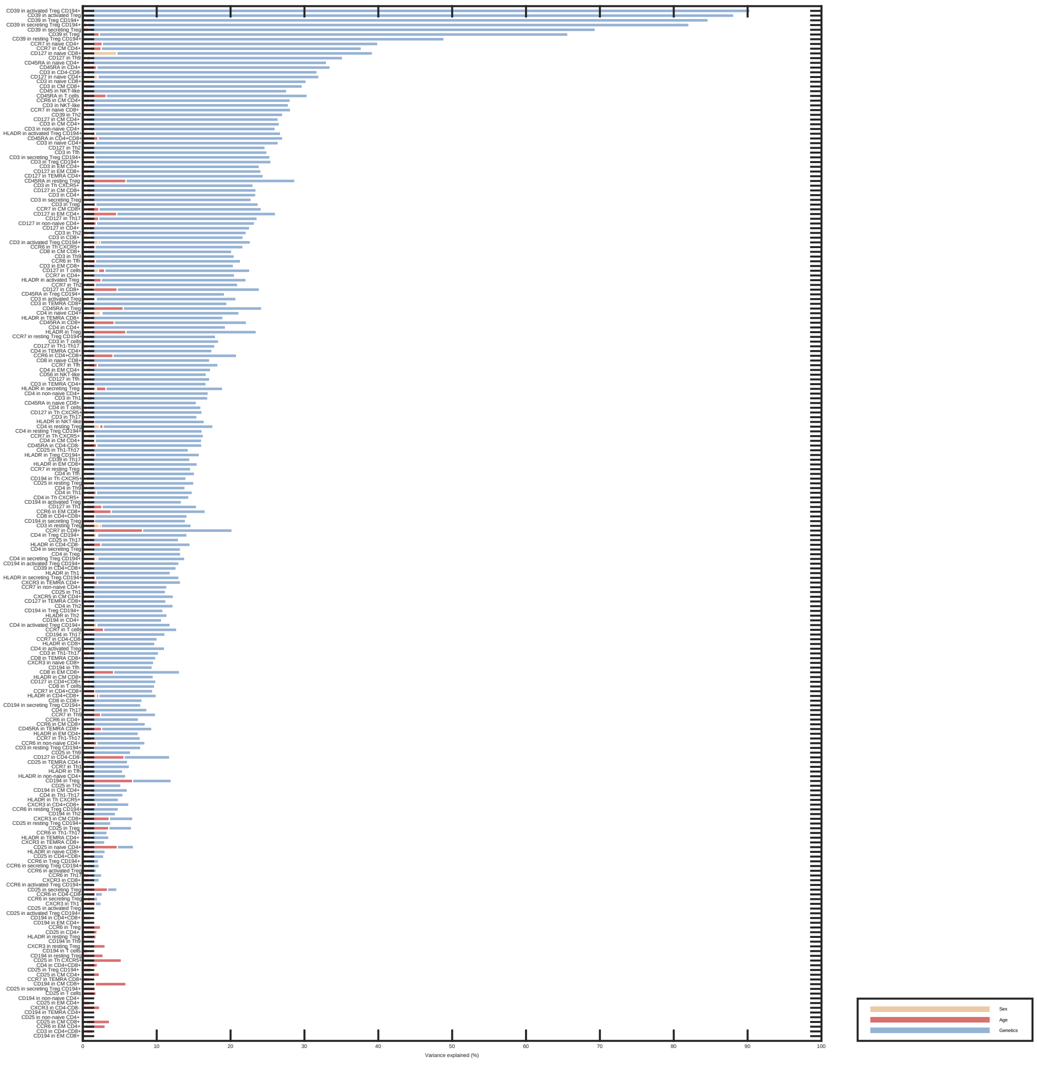

### Protein expression traits, B

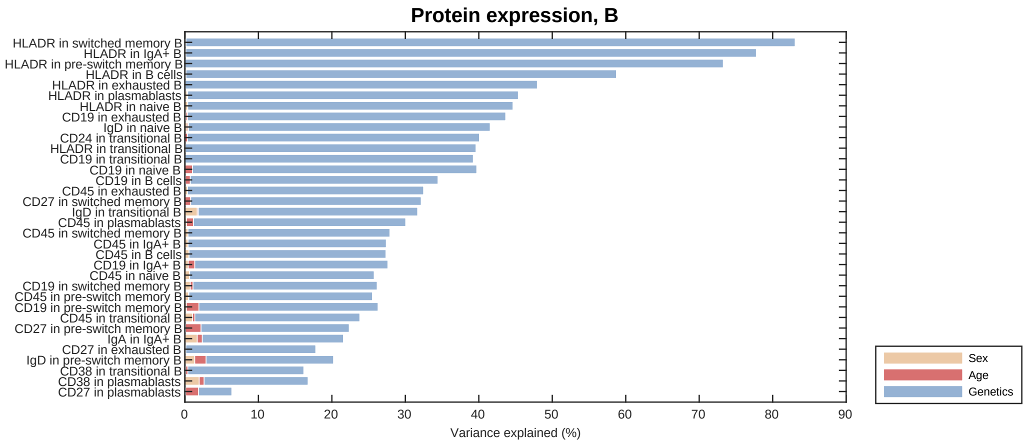


### Protein expression traits, MONO-NK-DC

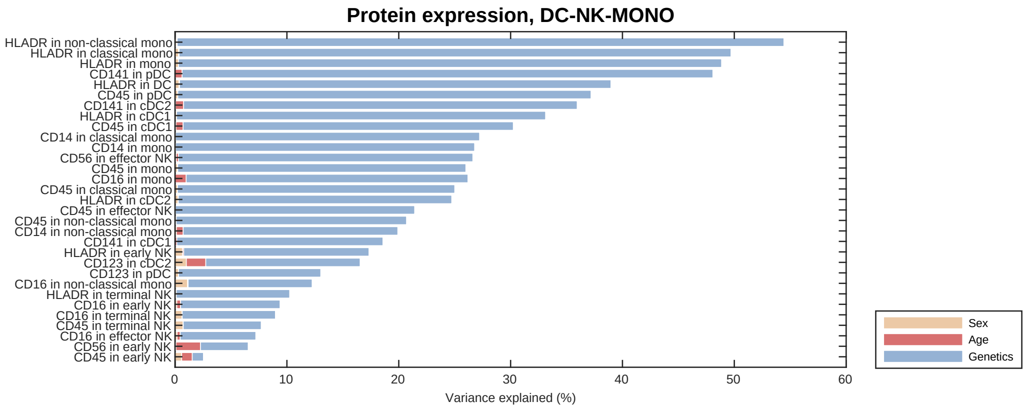

### Morphology, T

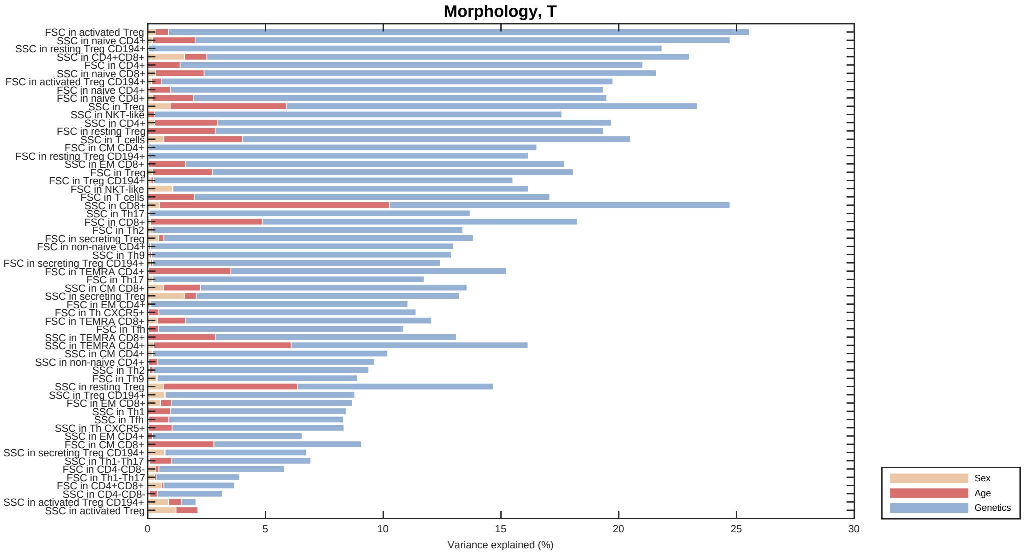


### Morphology, B

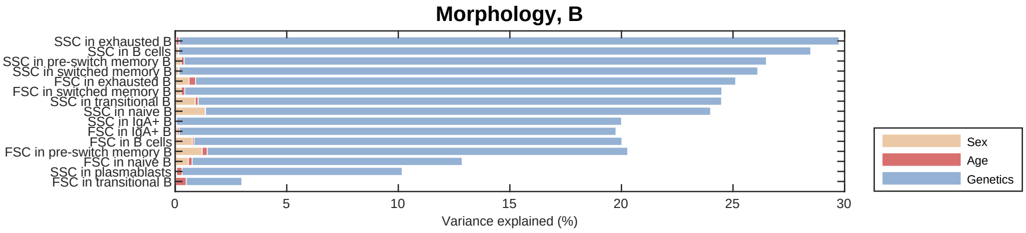


### Morphology, MONO-NK-DC

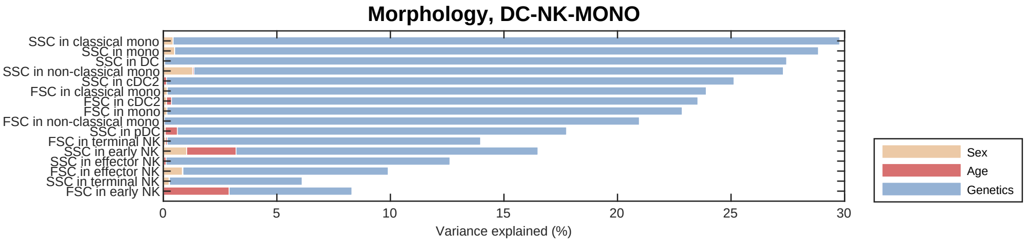

### Figure S5

Overview of data underlying the candidate gene assignment. Abbreviations: Expression quantitative trait locus (eQTL); splice quantitative trait locus (sQTL); protein quantitative trait locus (pQTL); RNAseq-ATACseq-correlation (RNA_ATAC).

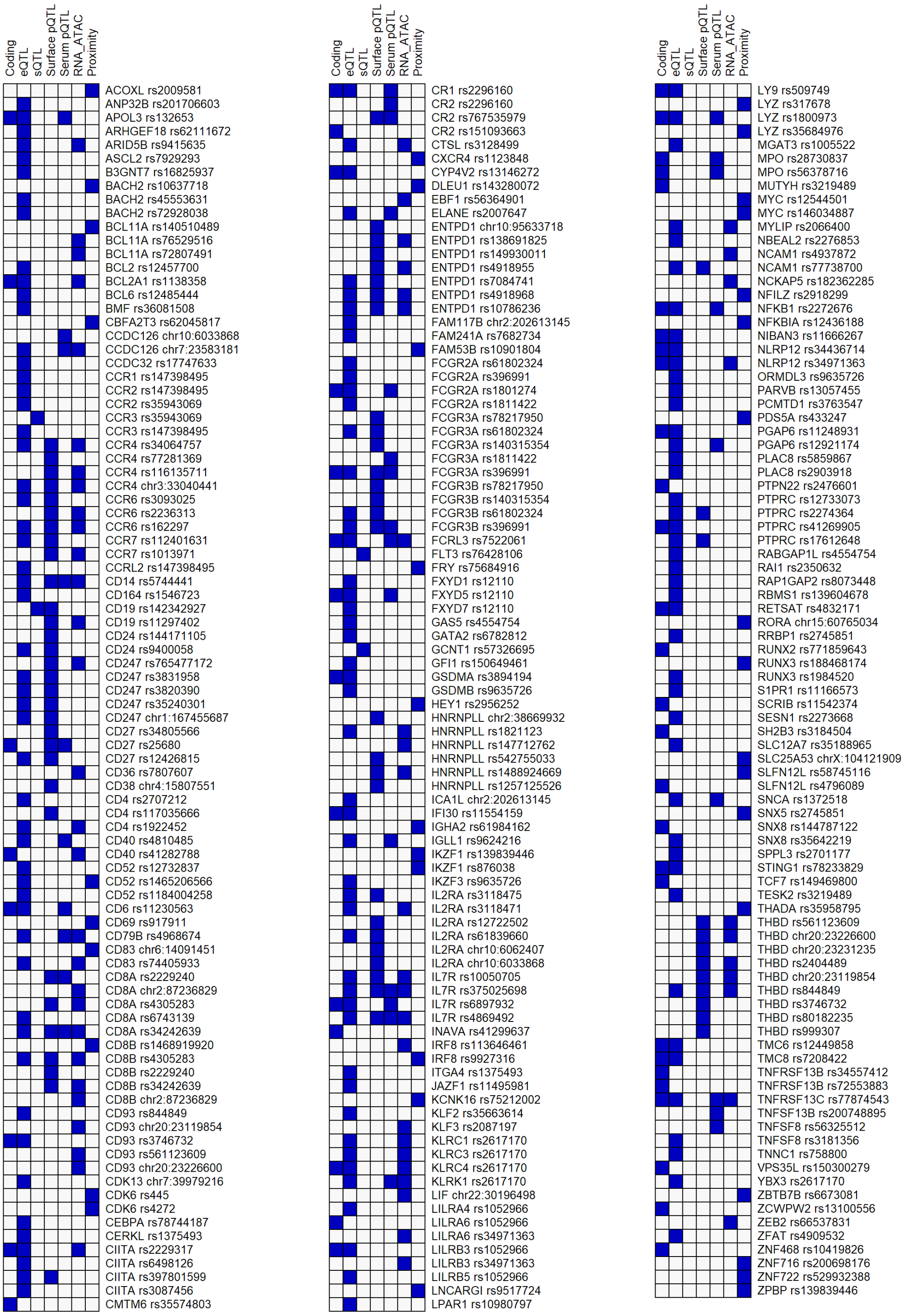

#### Figure S6

Autoimmunity and AML risk variant rs76428106-C perturbs specific myeloid subsets. **(A)** Immune cell traits associated with rs76428106-C, highlighting specific effects on dendritic cells and monocytes. **(B)** Diseases associations reported for rs76428106-C (**Table S9A**; Saevarsdottir *et al.* 2020). **(C)** Expression of *FLT3* across hematopoietic progenitors and immune cells based on RNA-seq data (Monaco *et al.* 2019; Granja *et al.* 2019). **(D)** Locus plot showing the position of rs76428106-C in *FLT3* intron 15. $x$-axis shows genomic position, the $y$-axis -$\log_{10}$(P) for association with the most significant immune cell trait. Introns are shown as lines, exons as boxes.

**A**

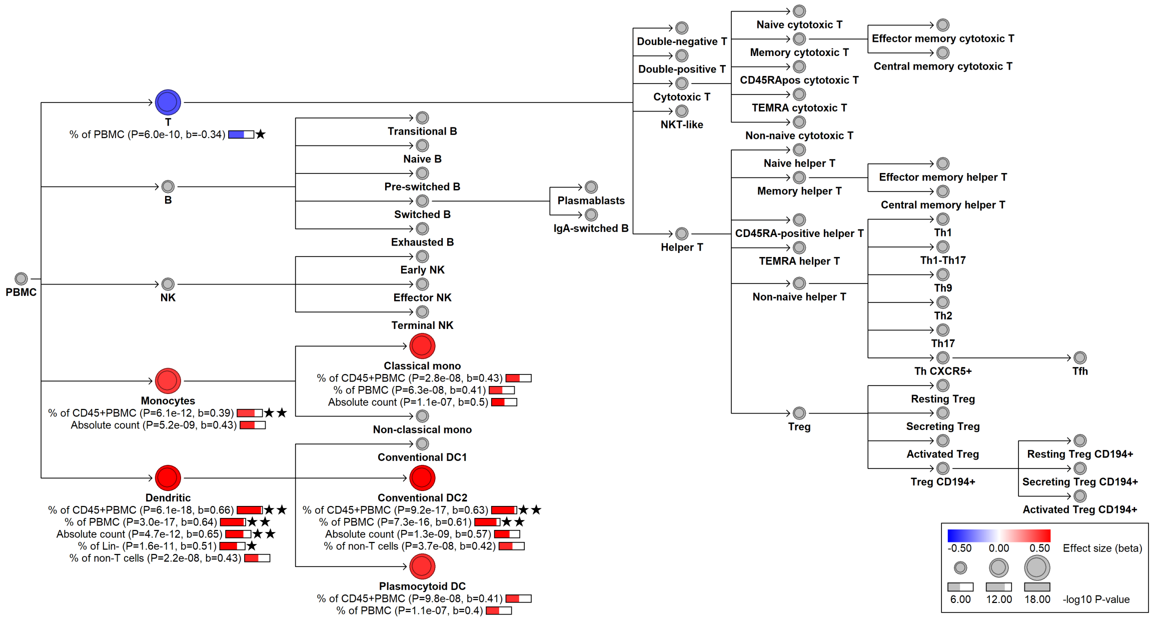


**B**

| Phenotype | rsID | Risk allele | Odds ratio | P-value | PMID | Effect of BloodVariome effect allele |
|---|---|---|---|---|---|---|
| Autoimmune thyroiditis | rs76428106 | C | 1.46 | 2.0e-24 | 32581359 | Increased risk |
| Rheumatoid arthritis | rs76428106 | C | 1.35 | 7.0e-11 | 35470158 | Increased risk |
| Acute myeloid leukemia | rs76428106 | C | 1.90 | 5.4e-3 | 32581359 | Increased risk |

**C**

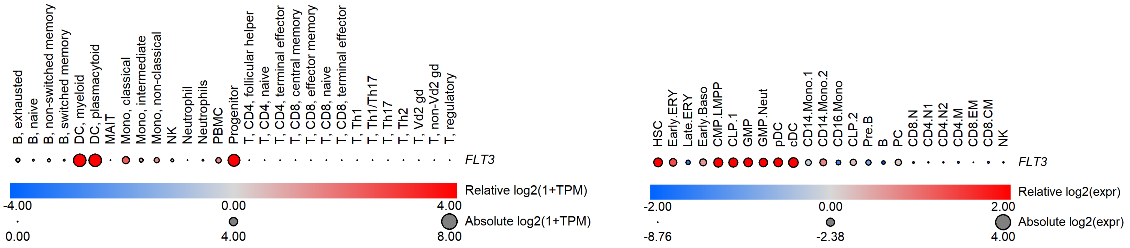


**D**

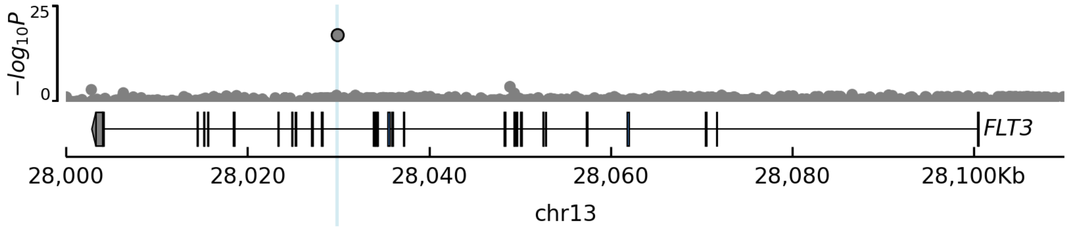

### Figure S7

The *IL7R* variant rs6897932 alters the naïve-to-memory CD4$^+$ T cell balance through alternative splicing. **(A)** rs6897932-T increases the frequency of naive relative to memory CD4$^+$T cells, while leaving membrane-bound IL7R$\alpha$ (CD127) expression unchanged. **(B)** Overlapping diseases associations (**Table S9A**). **(C)** Overlapping plasma pQTLs (**Table S7C**). **(D)** Expression of *IL7R* and genes encoding pQTL proteins in hematopoiesis (Monaco *et al.*, 2019, Granja *et al.*, 2019). TPM: transcripts per kilobase million. **(E)** Locus plot showing the location of credible set variants (blue). The $x$-axis indicates genomic position. The $y$-axis -log$_{10}$ indicates P-value for association with the most significant trait. Introns are shown as continuous lines, exons as boxes. rs6897932 maps to *IL7R* exon 6 and affects differential splicing between the two main isoforms: membrane-bound *IL7R* (with exon 6) and soluble *IL7R* (without exon 6).

**A**

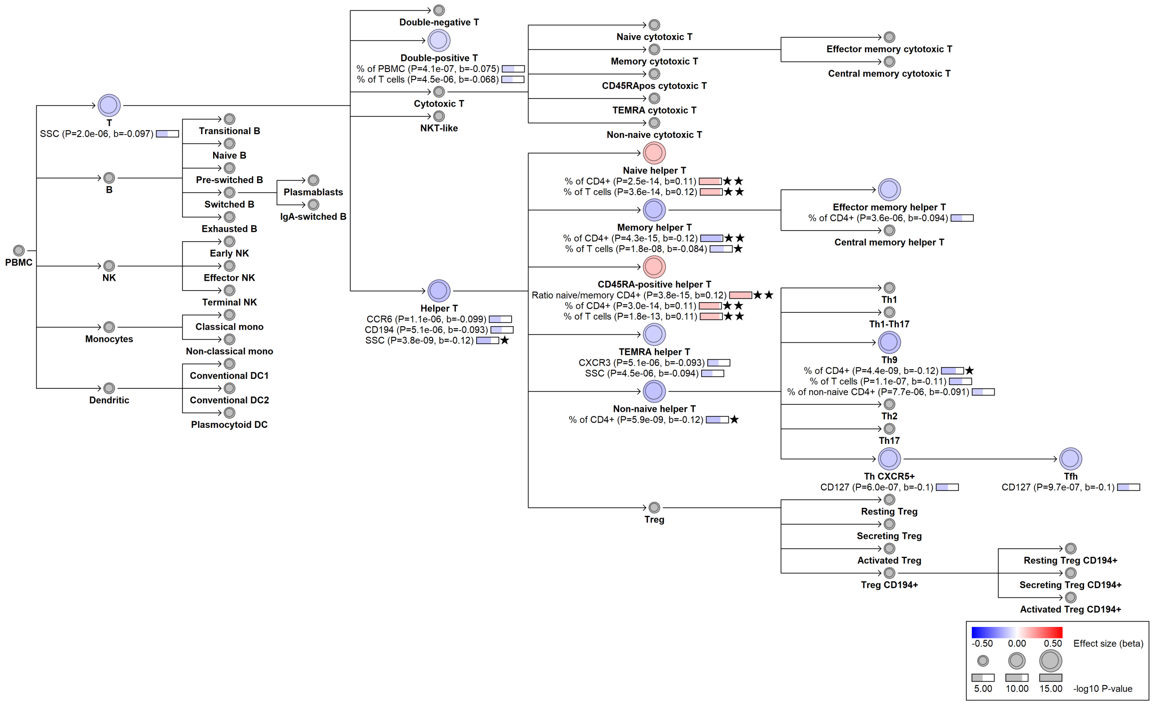


**B**

| Phenotype | rsID | Risk allele | Odds ratio | P-value | PMID | Effect of BloodVariome effect allele |
|---|---|---|---|---|---|---|
| Allergy | rs6881270 | C | 1.10 | 1.00E-48 | 31361310 | Decreased risk |
| Asthma | rs6881270 | C | 1.09 | 3.00E-44 | 31361310 | Decreased risk |
| Eczema | rs6881270 | C | 1.10 | 1.00E-48 | 31361310 | Decreased risk |
| Multiple sclerosis | rs6881706 | C | 1.12 | 4.00E-17 | 24076602 | Decreased risk |
| Primary biliary cholangitis | rs11742270 | A | 0.80 | 4.00E-24 | 34033851 | Decreased risk |
| SLE | rs6897932 | C | 1.07 | 2.00E-14 | 36750564 | Decreased risk |

**C**

| Protein | rsID | Effect allele | Beta | P-value | Source | Effect of BloodVariome effect allele |
|---|---|---|---|---|---|---|
| CCL21 | rs10214273 | G | -0.045 | 1.2e-09 | UKBB | Decreased level |
| CD48 | rs11567694 | G | -0.057 | 4.3e-12 | UKBB | Decreased level |
| CD5 | rs10214237 | C | -0.076 | 3.4e-24 | UKBB | Decreased level |
| CD6 | rs10214237 | C | -0.053 | 3.9e-14 | UKBB | Decreased level |
| GZMA | rs10213865 | C | -0.045 | 5.7e-10 | UKBB | Decreased level |
| IL7R | rs11742270 | A | -0.852 | 0 | UKBB | Decreased level |
| LY9 | rs11406102 | GT | -0.055 | 5.0e-14 | UKBB | Decreased level |
| PAG1 | rs10214237 | C | -0.054 | 4.5e-13 | UKBB | Decreased level |
| SELPLG | rs6871748 | C | -0.059 | 7.4e-15 | UKBB | Decreased level |
| SIT1 | rs33950349 | T | -0.05 | 2.3e-11 | UKBB | Decreased level |

**D**

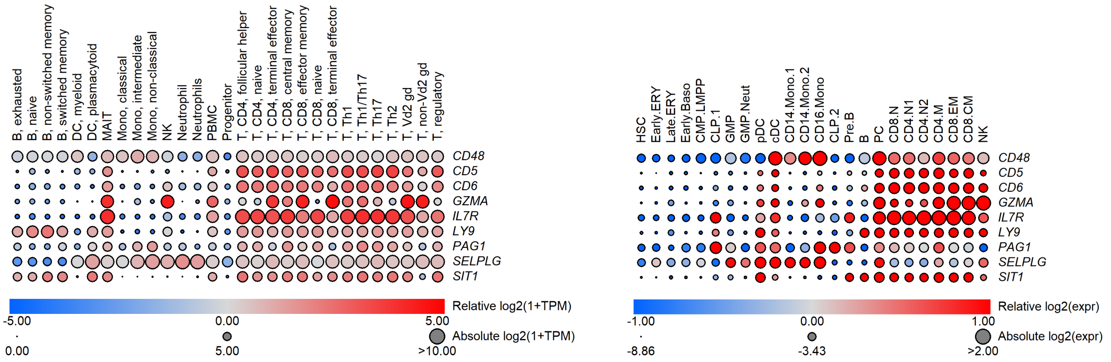
B, exhausted
B, naive
B, non-switched memory
B, switched memory
DC, myeloid
DC, plasmacytoid
MAIT
Mono, classical
Mono, intermediate
Mono, non-classical
NK
Neutrophil
Neutrophils
PBMC
Progenitor
T, CD4, follicular helper
T, CD4, naive
T, CD4, terminal effector
T, CD8, central memory
T, CD8, effector memory
T, CD8, naive
T, CD8, terminal effector
T, Th1
T, Th1/Th17
T, Th17
T, Th2
T, Vd2 gd
T, non-Vd2 gd
T, regulatory
CD48
CD5
CD6
GZMA
IL7R
LY9
PAG1
SELPLG
SIT1
Relative log2(1+TPM)
-5.00
0.00
5.00
Absolute log2(1+TPM)
0.00
5.00
>10.00
HSC
Early.ERY
Late.ERY
Early.Baso
CMP.LMPP
CLP.1
GMP
GMP.Neut
pDC
cDC
CD14.Mono.1
CD14.Mono.2
CD16.Mono
CLP.2
Pre.B
B
PC
CD8.N
CD4.N1
CD4.N2
CD4.M
CD8.EM
CD8.CM
NK
CD48
CD5
CD6
GZMA
IL7R
LY9
PAG1
SELPLG
SIT1
Relative log2(expr)
-1.00
0.00
1.00
Absolute log2(expr)
-8.86
-3.43
>2.00

**E**

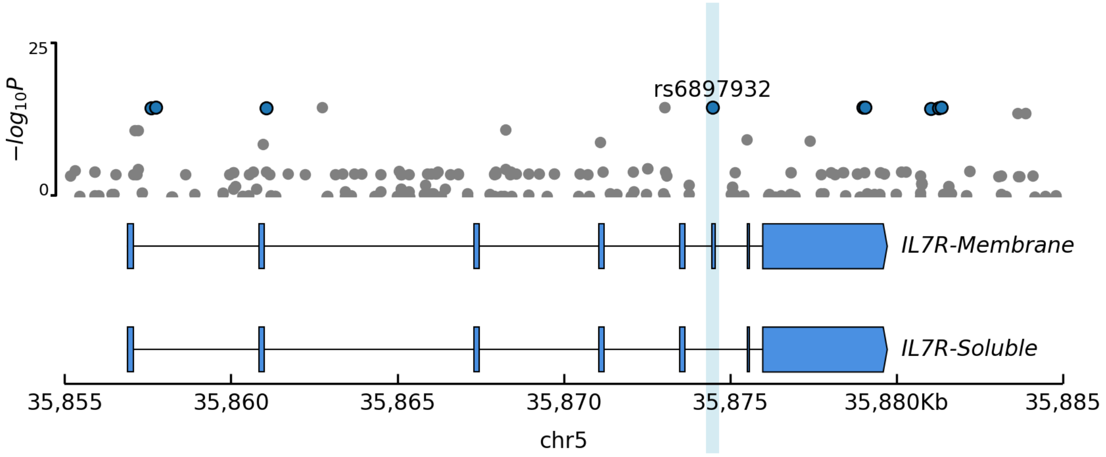
25
0
$-log_{10}P$
rs6897932
IL7R-Membrane
IL7R-Soluble
35,855
35,860
35,865
35,870
35,875
35,880Kb
35,885
chr5

### Figure S8

Independent *BACH2* variants exert lineage-specific effects on immune cell traits. **(A)** Locus plot showing the three independent credible sets at *BACH2* (cs_206 in green, cs_68 in blue, cs_191 in orange). $x$-axis indicates location, $y$-axis -$\log_{10}$ P-value for the most significant trait. Introns are shown as lines, exons as boxes. ATAC-seq tracks for relevant cell types are shown below the locus plots. **(B)** rs72928038-A (green) increases and **(C)** chr6:90279522-C (blue) decreases CCR4 (CD194) surface expression on follicular helper T (Tfh) cells; **(D)** rs45553631-T (orange) reduces cDC2s frequency. **(E)** Overlapping disease associations (**Table S9A**). **(F)** Effects on gene expression. The cs_206 and cs_191 signals have co-localized eQTLs, while cs_68 has a coincidental eQTL. **(G)** Plasma pQTLs for dendritic cell markers (**Table S7C**).

**A**

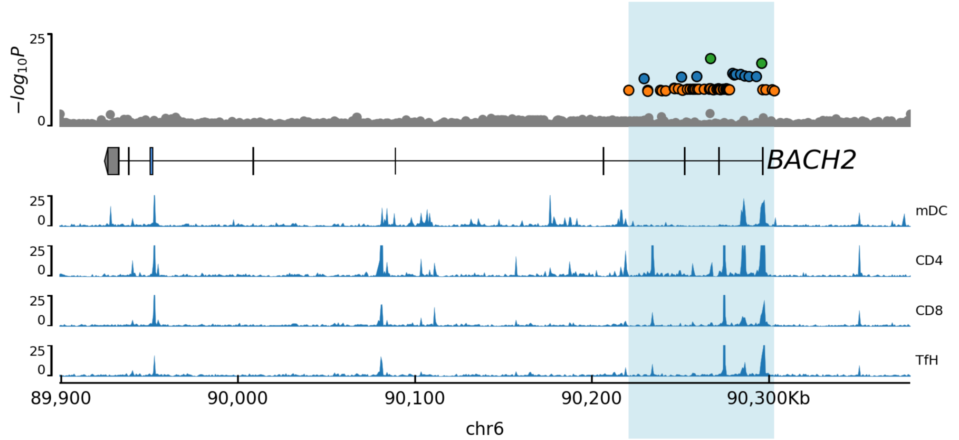


**B**

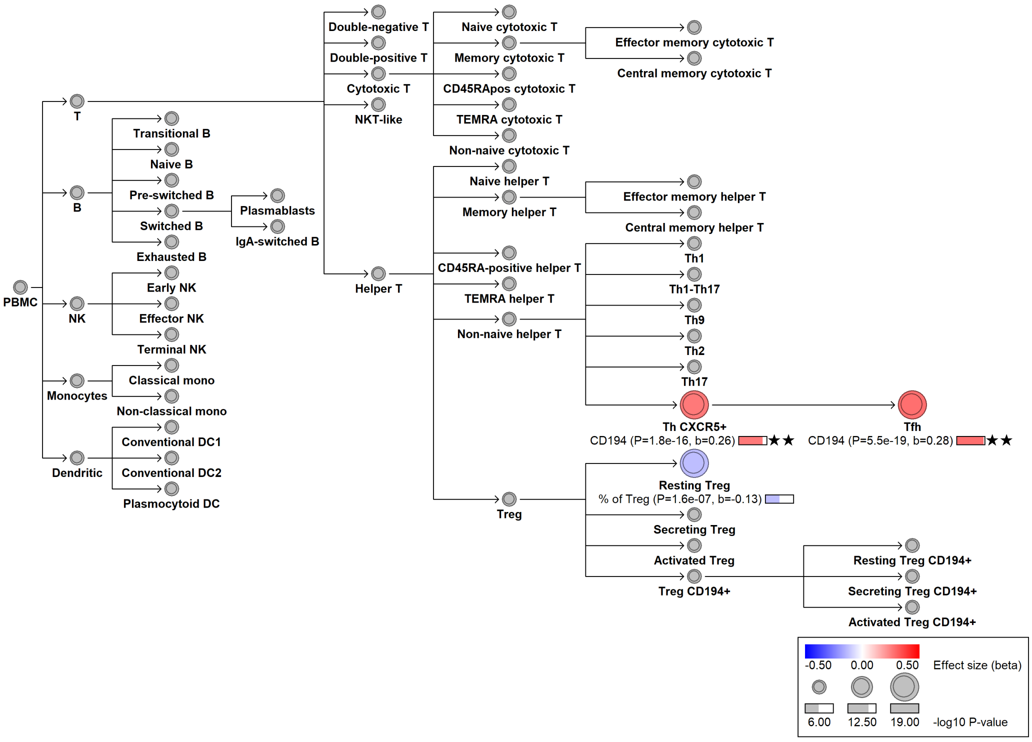

C

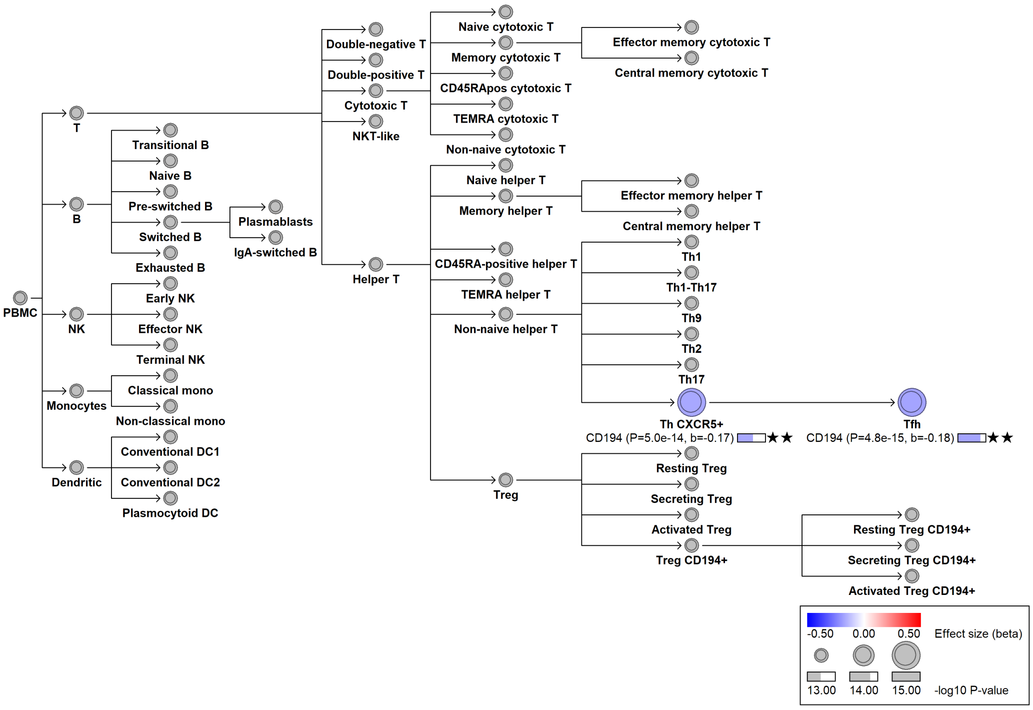
PBMC
T
B
NK
Monocytes
Dendritic
Transitional B
Naive B
Pre-switched B
Switched B
Exhausted B
Plasmablasts
IgA-switched B
Early NK
Effector NK
Terminal NK
Classical mono
Non-classical mono
Conventional DC1
Conventional DC2
Plasmocytoid DC
Double-negative T
Double-positive T
Cytotoxic T
NKT-like
Helper T
Naive cytotoxic T
Memory cytotoxic T
CD45RApos cytotoxic T
TEMRA cytotoxic T
Non-naive cytotoxic T
Effector memory cytotoxic T
Central memory cytotoxic T
Naive helper T
Memory helper T
CD45RA-positive helper T
TEMRA helper T
Non-naive helper T
Effector memory helper T
Central memory helper T
Th1
Th1-Th17
Th9
Th2
Th17
Th CXCR5+
CD194 (P=5.0e-14, b=-0.17) ★★
Tfh
CD194 (P=4.8e-15, b=-0.18) ★★
Treg
Resting Treg
Secreting Treg
Activated Treg
Treg CD194+
Resting Treg CD194+
Secreting Treg CD194+
Activated Treg CD194+
-0.50 0.00 0.50 Effect size (beta)
13.00 14.00 15.00 -log10 P-value

D

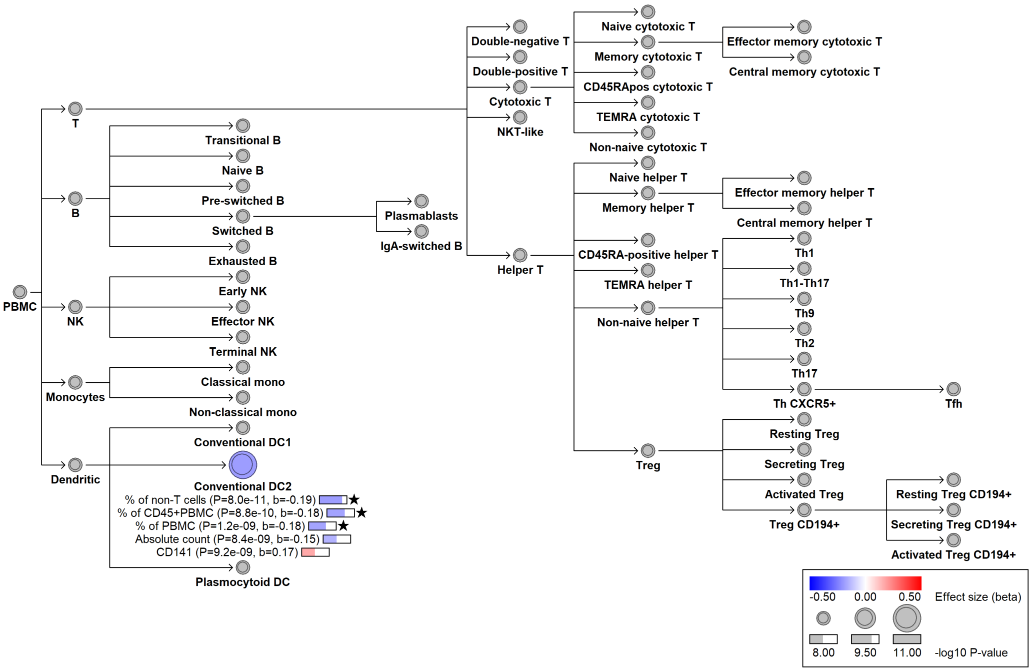
PBMC
T
B
NK
Monocytes
Dendritic
Transitional B
Naive B
Pre-switched B
Switched B
Exhausted B
Plasmablasts
IgA-switched B
Early NK
Effector NK
Terminal NK
Classical mono
Non-classical mono
Conventional DC1
Conventional DC2
% of non-T cells (P=8.0e-11, b=-0.19) ★
% of CD45+PBMC (P=8.8e-10, b=-0.18) ★
% of PBMC (P=1.2e-09, b=-0.18) ★
Absolute count (P=8.4e-09, b=-0.15)
CD141 (P=9.2e-09, b=0.17)
Plasmocytoid DC
Double-negative T
Double-positive T
Cytotoxic T
NKT-like
Helper T
Naive cytotoxic T
Memory cytotoxic T
CD45RApos cytotoxic T
TEMRA cytotoxic T
Non-naive cytotoxic T
Effector memory cytotoxic T
Central memory cytotoxic T
Naive helper T
Memory helper T
CD45RA-positive helper T
TEMRA helper T
Non-naive helper T
Effector memory helper T
Central memory helper T
Th1
Th1-Th17
Th9
Th2
Th17
Th CXCR5+
Tfh
Treg
Resting Treg
Secreting Treg
Activated Treg
Treg CD194+
Resting Treg CD194+
Secreting Treg CD194+
Activated Treg CD194+
-0.50 0.00 0.50 Effect size (beta)
8.00 9.50 11.00 -log10 P-value

**E**

| cs_id | Disease outcome | rsID | RA | OR | Beta | P | PMID | Effect of BloodVariome EA |
|---|---|---|---|---|---|---|---|---|
| cs_206 | Autoimmune thyroiditis | rs6908626 | T | nan | 0.14 | 2.0E-24 | 34594039 | Increased risk |
| cs_206 | Multiple sclerosis | rs72928038 | A | 1.15 | nan | 8.0E-29 | 31604244 | Increased risk |
| cs_206 | Rheumatoid arthritis | rs72928038 | A | 1.11 | nan | 6.0E-09 | 36333501 | Increased risk |
| cs_206 | Type 1 diabetes | rs72928038 | A | 1.20 | nan | 6.0E-14 | 25751624 | Increased risk |
| cs_206 | Vitiligo | rs72928038 | A | 1.27 | nan | 1.0E-14 | 27723757 | Increased risk |
| cs_206 | Malignant melanoma | rs6908626 | G | 1.09 | nan | 4.0E-09 | 32341527 | Decreased risk |
| cs_206 | Non-melanoma skin cancer | rs6908626 | T | 0.91 | nan | 2.0E-24 | 38182794 | Decreased risk |
| cs_68 | Autoimmune thyroiditis | rs654537 | A | 1.14 | nan | 6.0E-42 | 32581359 | Decreased risk |
| cs_68 | SLE | rs597325 | G | 1.12 | nan | 4.0E-12 | 27399966 | Decreased risk |
| cs_191 | Allergy | rs905670 | G | 1.06 | nan | 8.0E-23 | 31361310 | Decreased risk |
| cs_191 | Asthma | rs62408222 | G | nan | -0.06 | 1.0E-38 | 36777996 | Decreased risk |
| cs_191 | COPD | rs56353819 | C | nan | 0.02 | 6.0E-20 | 37069358 | Decreased risk |
| cs_191 | Crohn's disease | rs1847472 | C | 1.09 | nan | 1.0E-10 | 26192919 | Decreased risk |
| cs_191 | Eczema | rs72925996 | C | 0.96 | nan | 2.0E-44 | 37794016 | Decreased risk |
| cs_191 | Inflammatory bowel disease | rs1847472 | C | 1.06 | nan | 2.0E-10 | 23128233 | Decreased risk |
| cs_191 | Nasal polyps | rs62408224 | G | nan | -0.19 | 1.0E-17 | 34594039 | Decreased risk |

Abbreviations: Chronic obstructive pulmonary disease (COPD); Systemic lupus erythematosus (SLE).

**F**

| cs_id | rsID | OA | EA | EAF | Phenotypic effect | eQTL Z | eQTL P | eQTL effect |
|---|---|---|---|---|---|---|---|---|
| cs_206 | rs72928038 | G | A | 15.6 | Increased CCR4 expression on Tfh | -20.9 | 9.0E-97 | *BACH2* downregulation |
| cs_68 | rs10637718 | ! | C | 43.1 | Decreased CCR4 expression on Tfh | 13.7 | 1.3E-42 | *BACH2* upregulation |
| cs_191 | rs45553631 | C | T | 31.2 | Reduced cDC2 frequency | -15.1 | 8.7E-52 | *BACH2* downregulation |

**G**

| cs_id | pQTL pos | pQTL lead variant | r2 | Amin | -log10 P | Effect ($\beta$) | Protein |
|---|---|---|---|---|---|---|---|
| cs_191 | 90303148 | rs56353819 | 0.91 | T | 9.387 | -0.042 | CCL17 |
| cs_191 | 90291613 | rs367983479 | 0.84 | C | 11.36 | -0.06 | CCL22 |
| cs_191 | 90304027 | . | 0.89 | ! | 18.605 | -0.062 | CCL22 |
| cs_191 | 90291613 | rs367983479 | 0.84 | C | 16.26 | -0.07 | CCL22 |
| cs_191 | 90237621 | rs36048311 | 0.90 | G | 48.427 | -0.103 | CD207 |
| cs_191 | 90291613 | rs367983479 | 0.84 | C | 13.91 | -0.06 | FCGR1A |
| cs_191 | 90268664 | rs62408235 | 0.98 | C | 18.937 | -0.062 | FLT3 |

### Figure S9

*ARID5B* childhood leukemia risk variants perturbs early B-cell development. **(A)** rs9415635-G increases the frequency of transitional B cells among total B cells. **(B)** Locus plot showing credible set (blue). The $x$-axis represents the genomic position, the $y$-axis $-\log_{10}$ P-value for the most significant trait. The lead variant rs9415635 and the functionally fine-mapped childhood BCP-ALL risk variant rs7090445 are indicated. **(C)** Associations with BCP-ALL reported in the GWAS Catalog (**Table S9A**). **(D)** Overlapping plasma pQTLs (**Table S7C**). **(E)** Expression of *ARID5B* and genes encoding proteins implicated by pQTLs in blood and immune cells (Monaco *et al.*, 2019; Granja *et al.*, 2019). Counts per million (CPM); Transcripts per million (TPM).

**A**

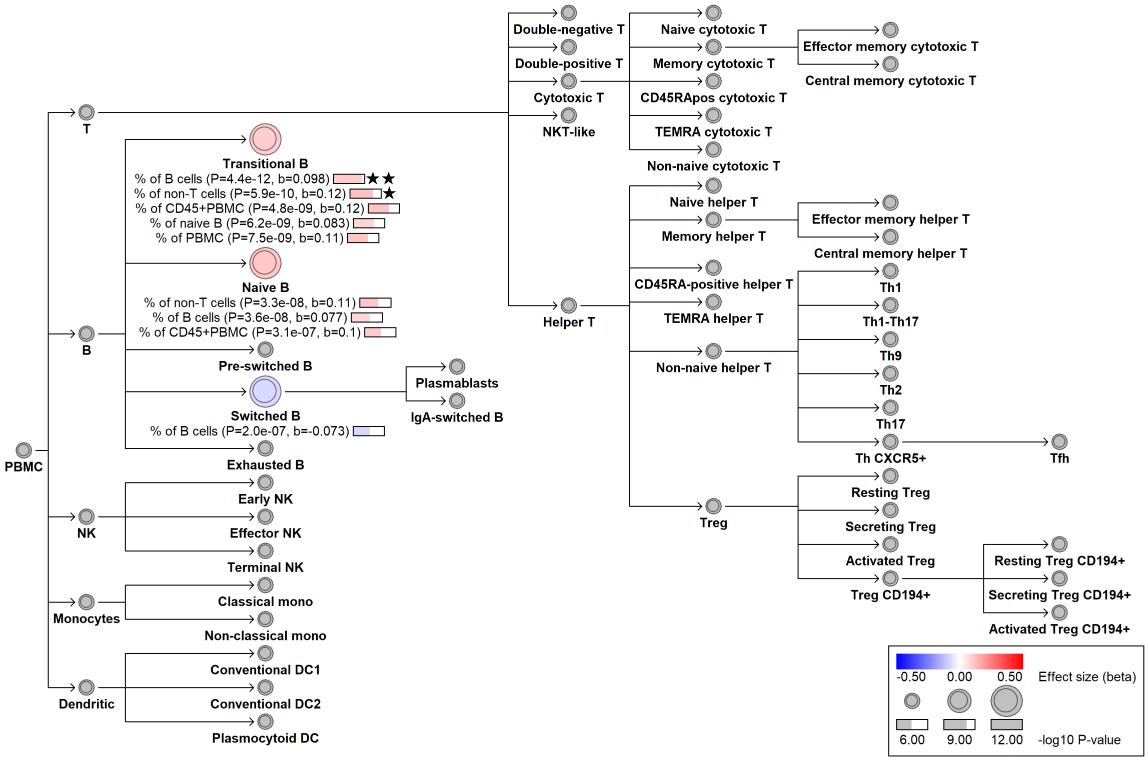


**B**

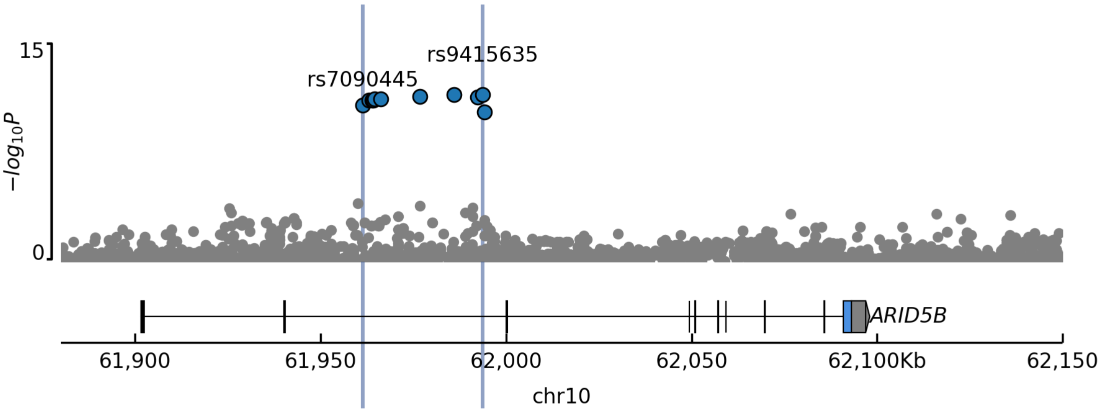

**C**

| GWAS Catalog rsID | Amaj | Amin | MAF | RA | OR | P | PMID | Effect of BloodVariome EA |
|---|---|---|---|---|---|---|---|---|
| rs10821936 | T | C | 30.8 | C | 1.80 | 1.00E-106 | 31767839 | Increased risk |
| rs10821936 | T | C | 30.8 | C | 1.42 | 1.00E-11 | 22076464 | Increased risk |
| rs10821936 | T | C | 30.8 | C | 1.91 | 1.00E-15 | 19684603 | Increased risk |
| rs10821936 | T | C | 30.8 | C | 2.31 | 2.00E-08 | 28817678 | Increased risk |
| rs4245595 | T | C | 30.7 | C | 1.63 | 2.00E-09 | 25310577 | Increased risk |
| rs7089424 | T | G | 31.5 | T | 1.64 | 2.00E-62 | 29348612 | Increased risk |
| rs7089424 | T | G | 31.5 | ? | 1.89 | 2.00E-73 | 29632299 | Increased risk |
| rs7090445 | T | C | 30.7 | ? | 1.96 | 3.00E-14 | 33891562 | Increased risk |
| rs10821936 | T | C | 30.8 | C | 1.46 | 4.00E-15 | 22076464 | Increased risk |
| rs7090445 | T | C | 30.7 | ? | nan | 5.00E-54 | 23996088 | Increased risk |
| rs10821936 | T | C | 30.8 | C | 1.86 | 6.00E-46 | 23512250 | Increased risk |
| rs7089424 | T | G | 31.5 | C | 1.65 | 7.00E-19 | 19684604 | Increased risk |

**D**

| Protein | rsID | Effect allele | Beta | P-value | Source | Effect of BloodVariome effect allele |
|---|---|---|---|---|---|---|
| BLNK | rs7087507 | G | 0.050 | 4.2e-10 | deCODE | Increased level |
| CD22 | rs4245597 | G | 0.071 | 7.7e-24 | UKBB | Increased level |
| CD72 | rs7090445 | C | 0.050 | 1.2e-09 | deCODE | Increased level |
| CD79B | rs9415635 | G | 0.059 | 3.7e-18 | UKBB | Increased level |
| DRAXIN | . | CTTTT | 0.098 | 3.1e-41 | UKBB | Increased level |
| FCER2 | rs4948492 | C | 0.080 | 3.3e-17 | deCODE | Increased level |
| FCRL1 | rs4948492 | C | 0.114 | 3.6e-64 | UKBB | Increased level |
| IGLL1 | rs7090445 | C | 0.140 | 7.4e-49 | deCODE | Increased level |
| PCDH9 | rs7087507 | G | 0.060 | 1.9e-11 | deCODE | Increased level |
| TCL1A | rs4245597 | G | 0.087 | 9.6e-36 | UKBB | Increased level |
| TNFRSF13C | rs7087507 | G | 0.063 | 8.4e-21 | UKBB | Increased level |

**E**

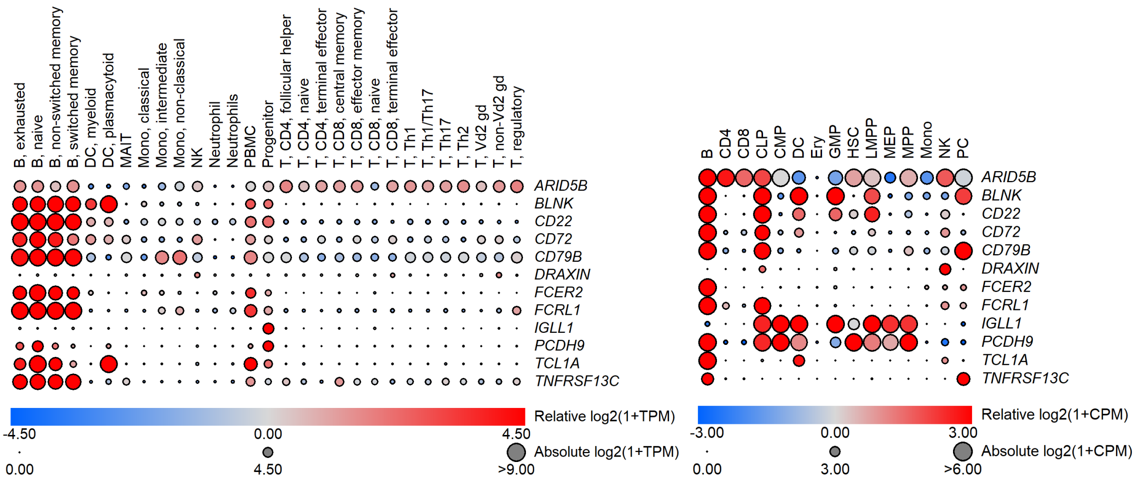

### Figure S10

*CR2* variants modulate the CR2–CD19 co-receptor axis in human B cells. **(A)** Effects of the rare *CR2* variant rs151093663-A; and **(B)** the common variant rs767535979-CA. **(C)** Expression of *CR2* and *CD19* in hematopoietic cell types (Monaco *et al.*, 2019; Granja *et al.*, 2019). **(D)** Plasma pQTL of rs767535979-CA (**Table S7C**).

**A**

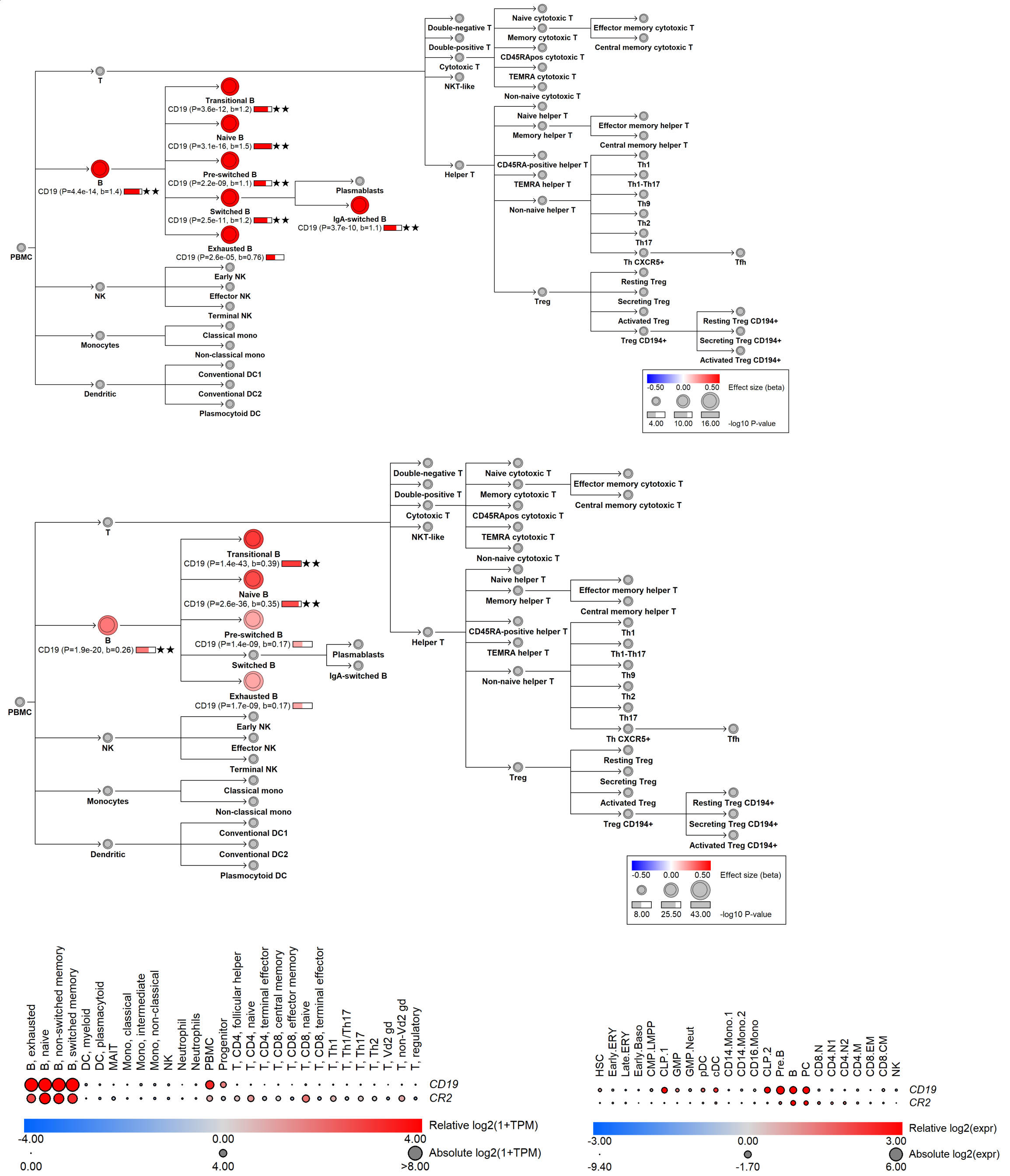


**D**

| Protein | rsID | Effect allele | Beta | P-value | Source | Effect of BloodVariome effect allele |
|---|---|---|---|---|---|---|
| CR2 | rs17044525 | G | -0.29 | 9.5e-169 | UKBB | Decreased risk |

### Figure S11

*MPO* deficiency variants reduce monocyte granule content. **(A,B)** The variants rs28730837-A and rs56378716-G at *MPO* are associated with reduced monocyte side scatter (SSC), consistent with the role of myeloperoxidase as a major antimicrobial effector protein stored in monocytic granules. **(C)** Clinical annotations for MPO variants associated with myeloperoxidase deficiency retrieved from ClinVar.

**A**

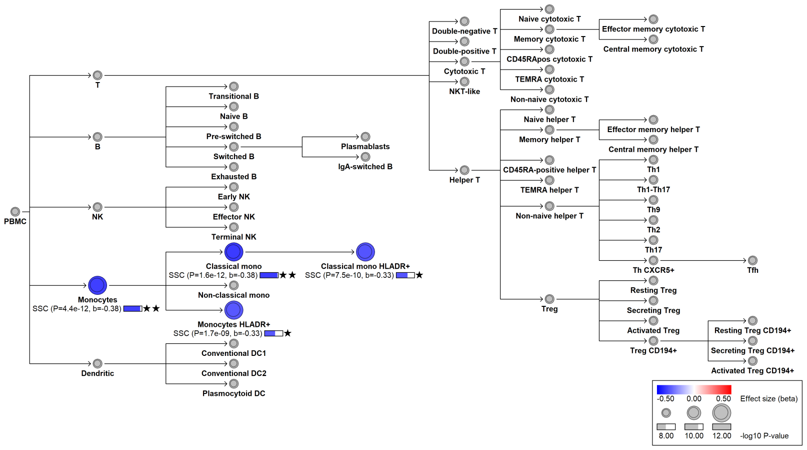


**B**

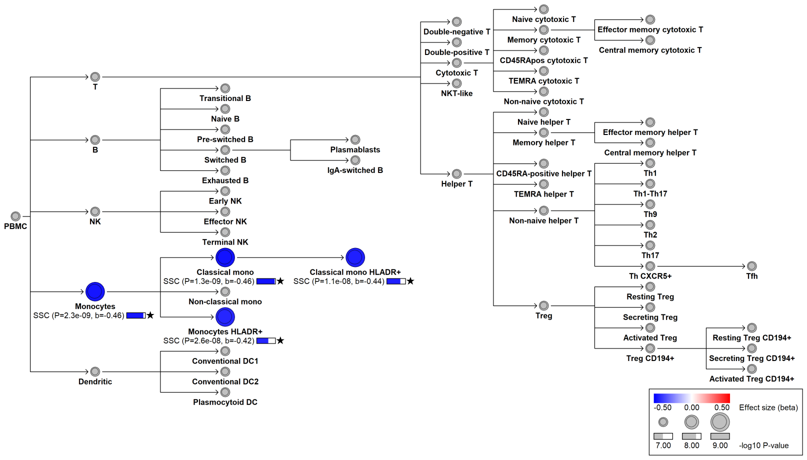


**C**

| csID | Phenotype | rsID | EA | Beta | P-value | Source | Effect of BloodVariome EA |
|---|---|---|---|---|---|---|---|
| cs_20 | MPO deficiency | rs56378716 | G | - | - | ClinVar | Increased risk |
| cs_212 | MPO deficiency | rs28730837 | A | - | - | ClinVar | Increased risk |

### Figure S12

An *IFI30* missense variant alters monocyte lysosomal morphology. **(A)** The *IFI30* variant rs11554159-A (p.Arg76Gln) is associated with reduced monocyte side scatter (SSC), consistent with IFI30's role in lysosomal antigen processing. **(B)** Overlapping disease associations (**Table S9A**). **(C)** Expression of IFI30 across hematopoietic cell types based on bulk and single-cell RNA-sequencing datasets (Monaco et al., 2019; Granja et al., 2019).

**C**

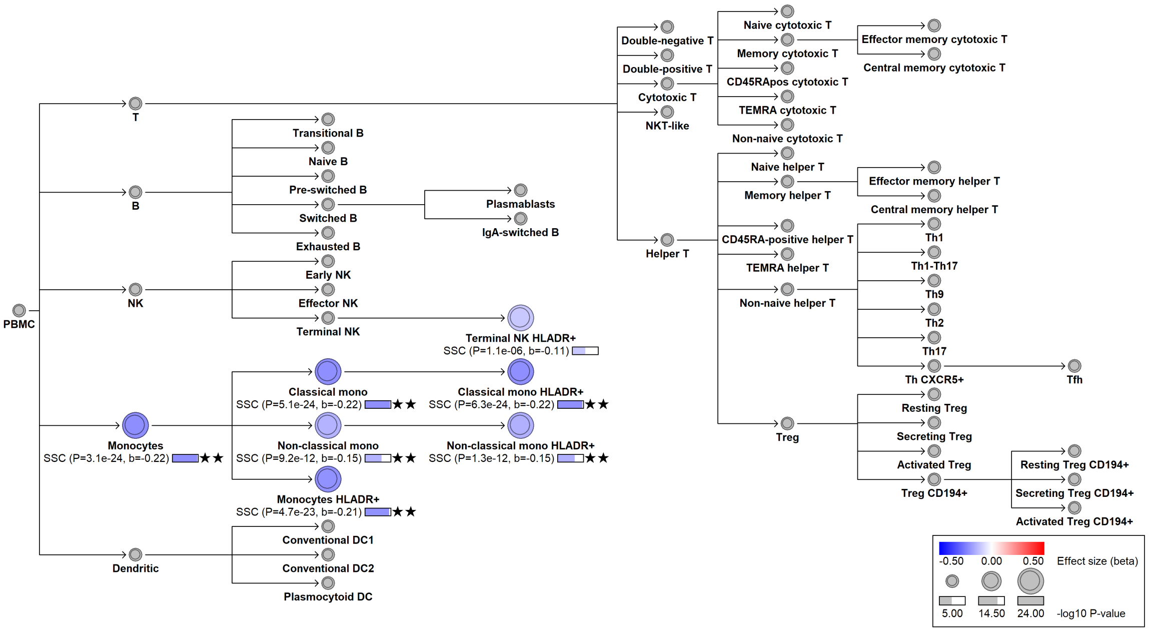


**D**

| csID | Phenotype | rsID | EA | Beta | P-value | Source | Effect of BloodVariome EA |
|---|---|---|---|---|---|---|---|
| cs_226 | Multiple sclerosis | rs874628 | A | 1.11 | 1.00E-08 | 21833088 | Decreased risk |

**E**

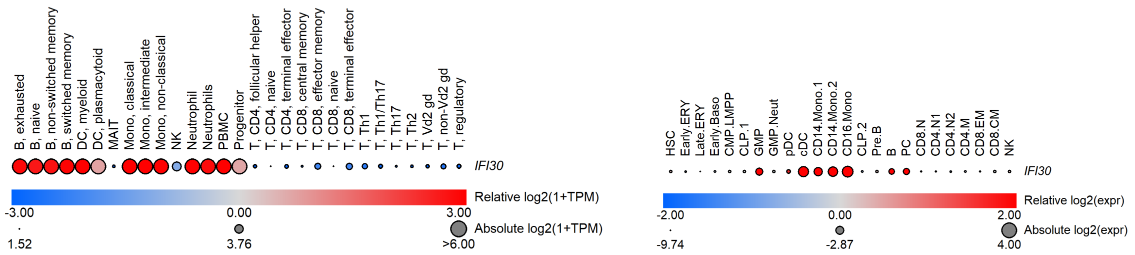

### Figure S13

A missense variant in *NIBAN3* influences B-cell abundance. **(A)** Effects of the missense variant rs11666267 (p.Ser572Gly) on immune cell traits. **(B)** Expression of *NIBAN3* across hematopoietic progenitors and immune cell types based on bulk and single-cell RNA-sequencing datasets (Monaco et al., 2019; Granja et al., 2019). **(C)** Plasma protein quantitative trait loci (pQTLs) associated with variants in the credible set (**Table S7C**).

**A**

chr19:17549491[A→G; EAF 46.672%] (rs11666267, cs129)

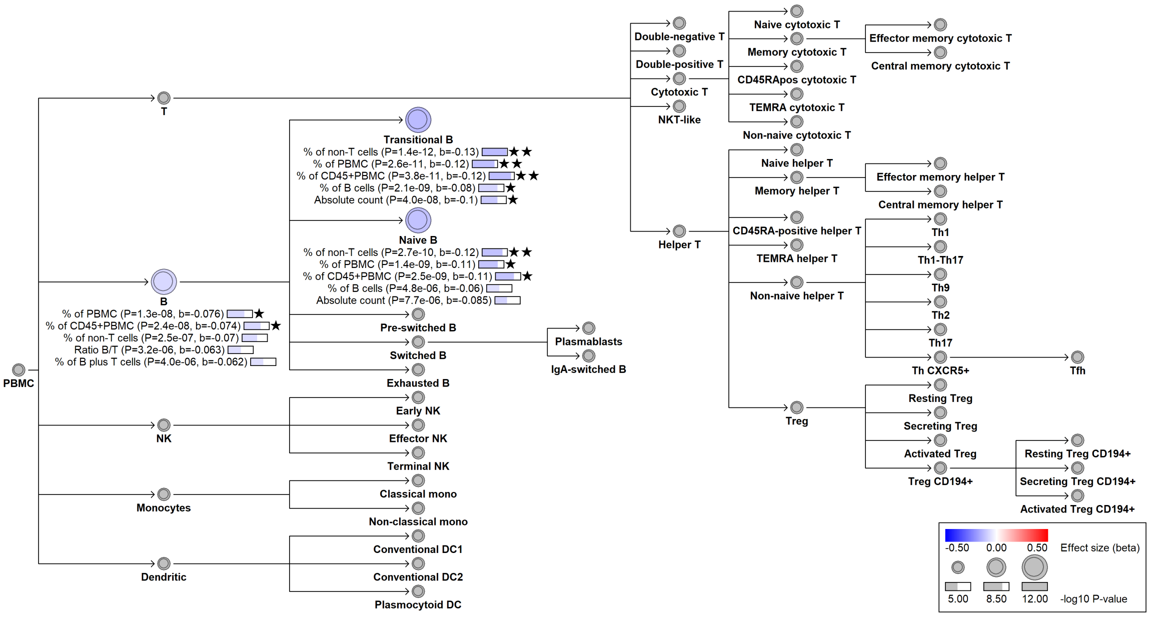


**B**

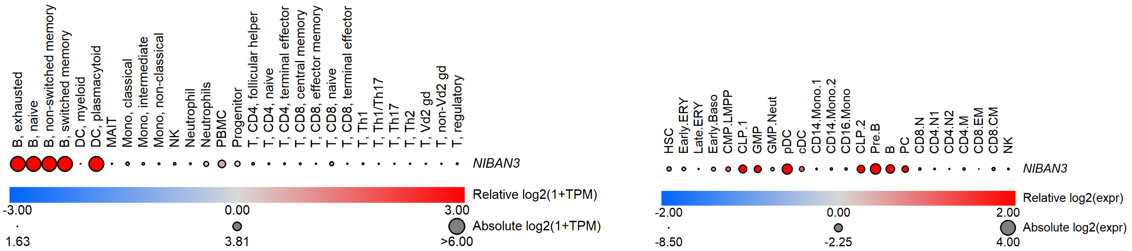


**C**

| Protein | rsID | EA | Beta | P-value | Source | Effect of BloodVariome EA |
|---|---|---|---|---|---|---|
| CD5L | rs11666267 | G | -0.04 | 1.2e-09 | UKBB | Decreased level |
| FCER2 | rs11666267 | G | -0.045 | 3.6e-12 | UKBB | Decreased level |
| IgM | rs11666267 | G | -0.036 | 1.0E-12 | Ali *et al.*, manuscript | Decreased level |
| TCL1A | rs1967330 | C | -0.049 | 2.0e-13 | UKBB | Decreased level |
| TNFRSF13C | rs1967330 | C | -0.055 | 5.9e-17 | UKBB | Decreased level |

#### Figure S14

Regulatory variants influencing THBD (CD141) expression on dendritic cells. **(A)** Identified variants (details in **Table S11**). **(B)** Locus plot of the nine *cis*-acting variants influencing THBD expression (blue, pDC; red, cDC; yellow, both pDC and cDC). The $x$-axis genomic position. The $y$-axis shows the -$\log_{10}$ P for association with the most significant trait. Tracks below the locus plot show transcription factor occupancy from reMap (Hammal *et al.*, 2022) and chromatin accessibility in pDC and mDC (ATAC-seq; Bao *et al.*, 2019). **(C)** Close-up of the *THBD* locus showing with chromatin accessibility in pDC and mDC (Bao *et al.*, 2019) together with transcription factor binding for NF-kB, IRF8, and IKZF1 (ChIP-seq; Wu L *et al.*, 2020, Mohaghegh N *et al.*, 2019, ENCODE, 2012).

**A**

| csID | Lead variant | candidate gene | EA | EAF | P-value | Beta | Action |
|---|---|---|---|---|---|---|---|
| cs_2 | rs75153485 | *THBD* | G | 7.3 | 9.25E-111 | 0.79 | *cis* |
| cs_18 | rs844849 | *THBD/CD93* | A | 10.0 | 3.03E-23 | 0.39 | *cis* |
| cs_31 | rs34290000 | *THBD/CD93* | ! | 27.7 | 4.23E-57 | -0.31 | *cis* |
| cs_61 | rs80182235 | *THBD* | E | 2.6 | 1.18E-09 | 0.38 | *cis* |
| cs_63 | rs999307 | *THBD* | C | 48.7 | 5.91E-84 | 0.40 | *cis* |
| cs_73 | rs2404489 | *THBD* | T | 27.8 | 5.98E-80 | 0.35 | *cis* |
| cs_91 | rs561123609 | *THBD* | C | 0.3 | 1.41E-17 | 1.26 | *cis* |
| cs_153 | chr20:23231235 | *THBD* | ! | 14.4 | 5.31E-52 | -0.38 | *cis* |
| cs_183 | rs3746732 | *THBD* | A | 25.6 | 1.61E-13 | 0.19 | *cis* |
| cs_78 | rs113646461 | *IRF8* | C | 10.0 | 6.39E-15 | -0.23 | *trans* |
| cs_136 | rs9927316 | *IRF8* | G | 23.2 | 3.81E-17 | -0.19 | *trans* |
| cs_30 | rs12436188 | *NFKBIA* | C | 18.2 | 2.64E-09 | -0.14 | *trans* |
| cs_210 | rs876038 | *IKZF1* | T | 29.7 | 4.95E-144 | -0.48 | *trans* |
| cs_9 | rs34436714 | *NLRP12* | A | 22.7 | 1.11E-173 | -0.45 | *trans* |
| cs_154 | rs1052966 | *LILR- family* | T | 13.6 | 2.20E-12 | 0.21 | *trans* |
| cs_98 | rs2276853 | *NBEAL2* | ! | 39.95 | 3.49E-10 | 0.117 | *trans* |

**B**

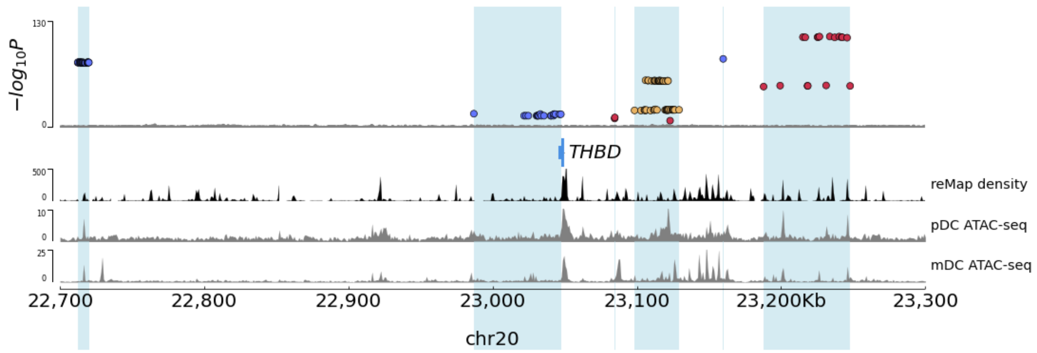


**C**

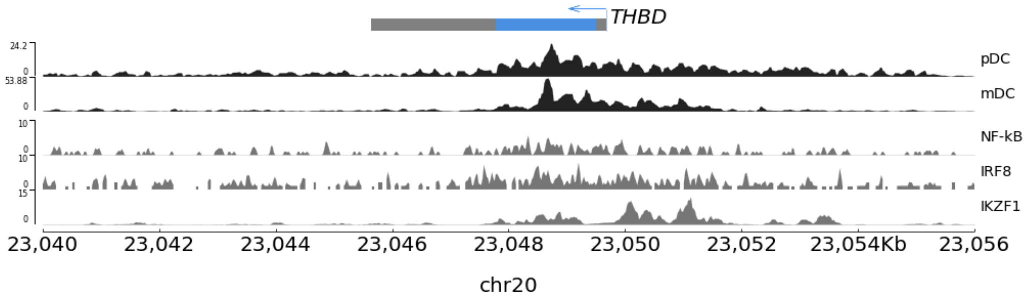

### Figure S15

Genetic variants influencing BCR-IgD expression. **(A)** Variants associated with IgD surface expression on naïve and transitional B cells (**Tables S4B-C**). **(B)** Overlapping associations with circulating Ig levels (**Table S7C**). **(C)** Overlapping *cis*-eQTLs (**Table S7A**).

**A**

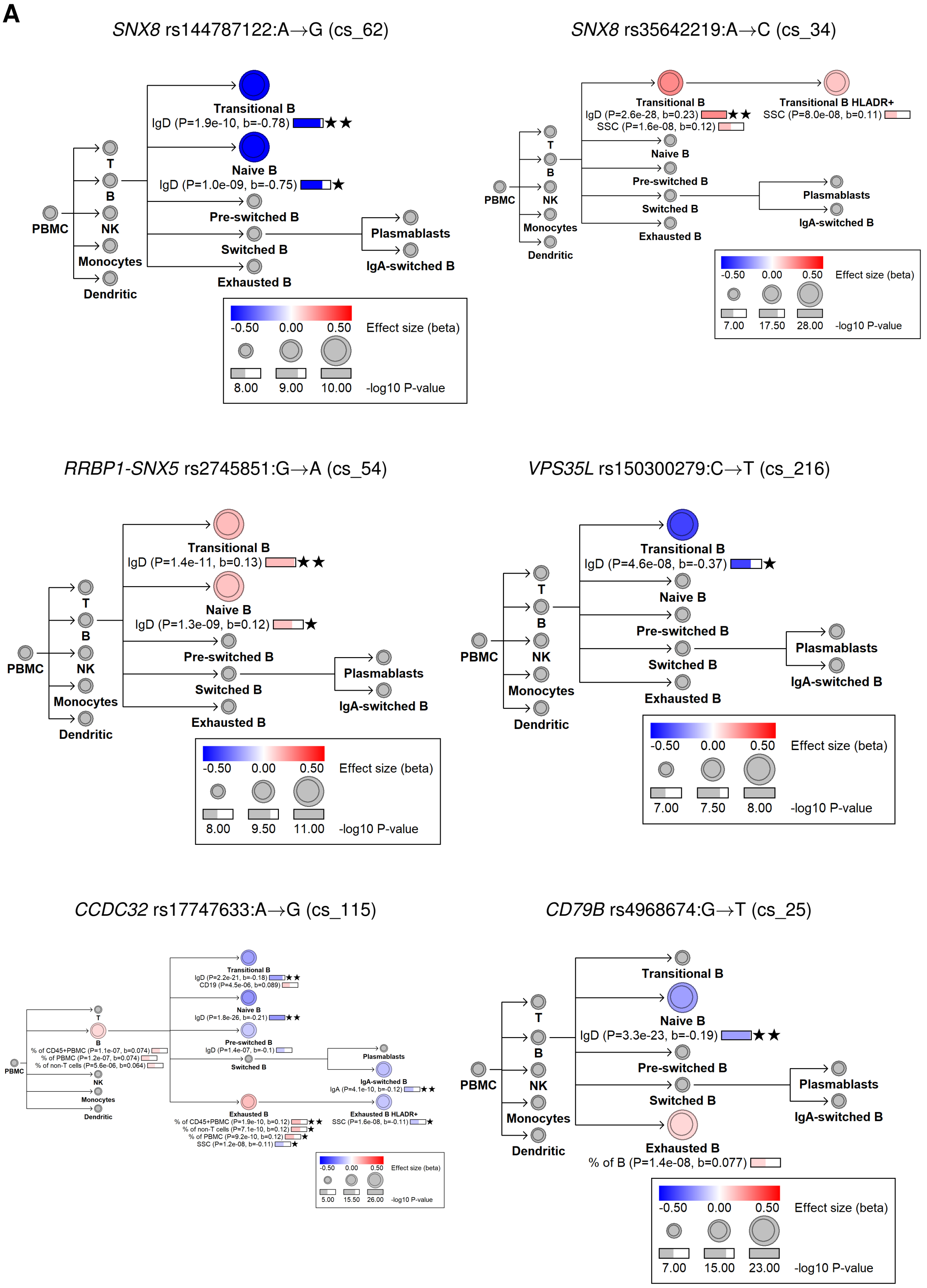

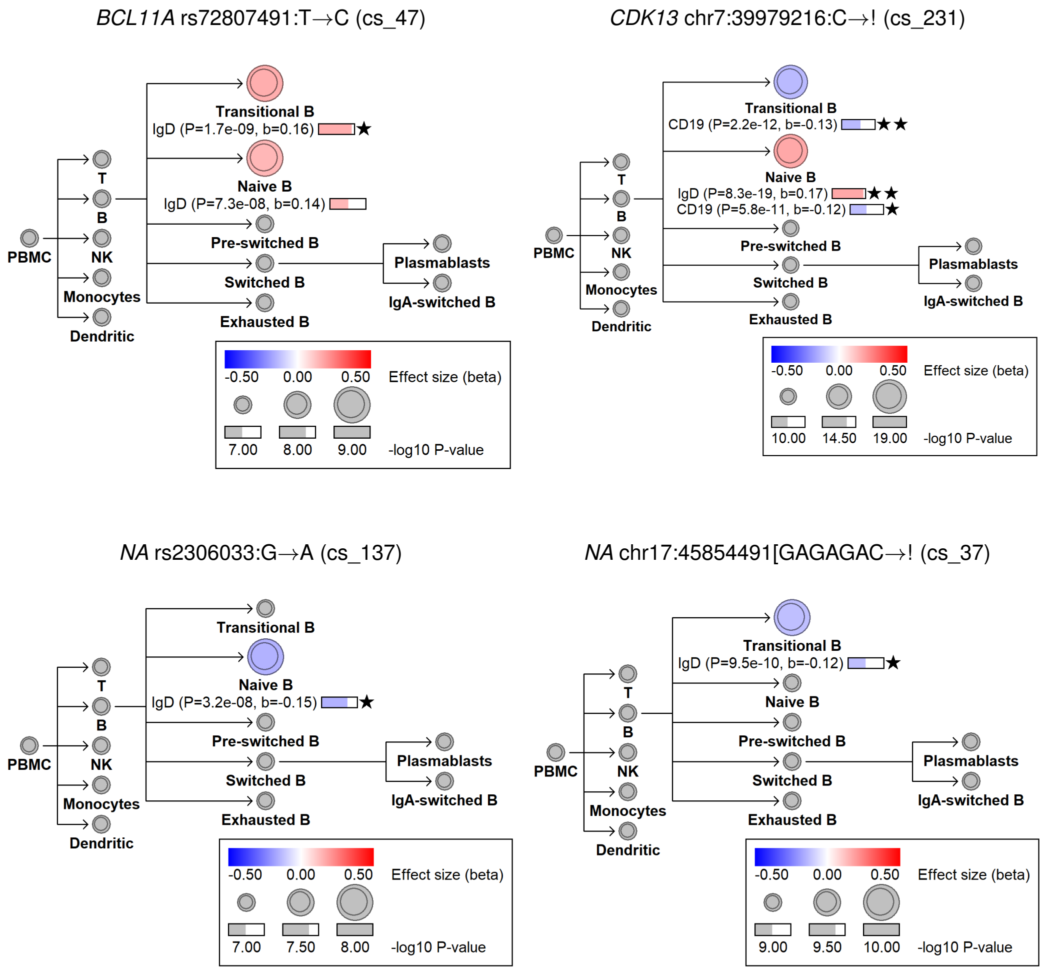


**B**

| cs_ID | Protein | pQTL rsID | EA | P-value | Beta | Effect of BloodVariome EA |
|---|---|---|---|---|---|---|
| cs_115 | Composite Ig trait IgG/IgM | rs10152546 | A | 0.030 | 1.0E-08 | Increase |
| cs_115 | IgM | rs10152546 | A | -0.029 | 8.4E-09 | Decrease |
| cs_25 | Composite Ig trait IgA/IgG | rs12452767 | A | 0.054 | 1.6E-25 | Increase |
| cs_25 | IgA | rs12452767 | A | 0.034 | 2.3E-11 | Increase |
| cs_25 | IgG | rs12452767 | A | -0.032 | 1.3E-10 | Decrease |
| cs_62 | IgG | rs144787122 | G | -0.33 | 1.5E-17 | Decrease |

Abbreviations: Credible set ID (cs_ID); Effect allele (EA).

**C**

| cs_ID | eQTL rsID | Coloc/$r^2$ | Gene | Cell type | EA | Beta | P | Effect of BV EA |
|---|---|---|---|---|---|---|---|---|
| cs_54 | rs2745849 | 1.0 | *RRBP1* | B cells | G | 0.68 | 3.2E-52 | Increae |
| cs_115 | chr15:40583386 | 0.73 | *CCDC32* | Sw. mem. B | A | -0.49 | 2.5E-37 | Decrease |
| cs_115 | chr15:40585318 | 0.79 | *CCDC32* | B cells | A | -0.55 | 1.7E-13 | Decrease |
| cs_115 | chr15:40579459 | 0.80 | *CCDC32* | B cells | CTT | -0.71 | 3.0E-19 | Decrease |

Abbreviations: BloodVariome (BV); Credible set ID (cs_ID); Effect allele (EA).
