## Supplementary material for "BloodVariome: a high-resolution atlas of inherited genetic effects in human immune cells": Data S1

Hierarchy plots illustrating the effects of lead variants for each credible set on different immune cell types.

The circular icons represent cell populations. The labels below describe the traits associated with that cell population. The length of the bars to the right of each trait indicate -$\log_{10}$ $P$-value for the associations. Color indicates the effect size ($\beta$) of the association. One star indicates trait-wide significance; two stars represent study-wide significance. Association signals that do not reach the significance threshold but are within two logarithms of trait-wide significance (suggestive signals) are also indicated, with no stars. The color of the cell icon indicates the effect size of the most significant trait associated with that cell type. In hard brackets we show the other allele with an arrow pointing to the effect allele. For multi-allelic variants, exclamation marks denotes all other alleles.

Also shown are the disease associations and serum pQTL associations of each credible set (details in **Table S9A, S7C**).

### chr1:24911495[C→T; EAF 40.131%] (rs1984520, cs_100)

Candidate genes: *RUNX3*
Diseases associations: NA
Plasma pQTLs: NA

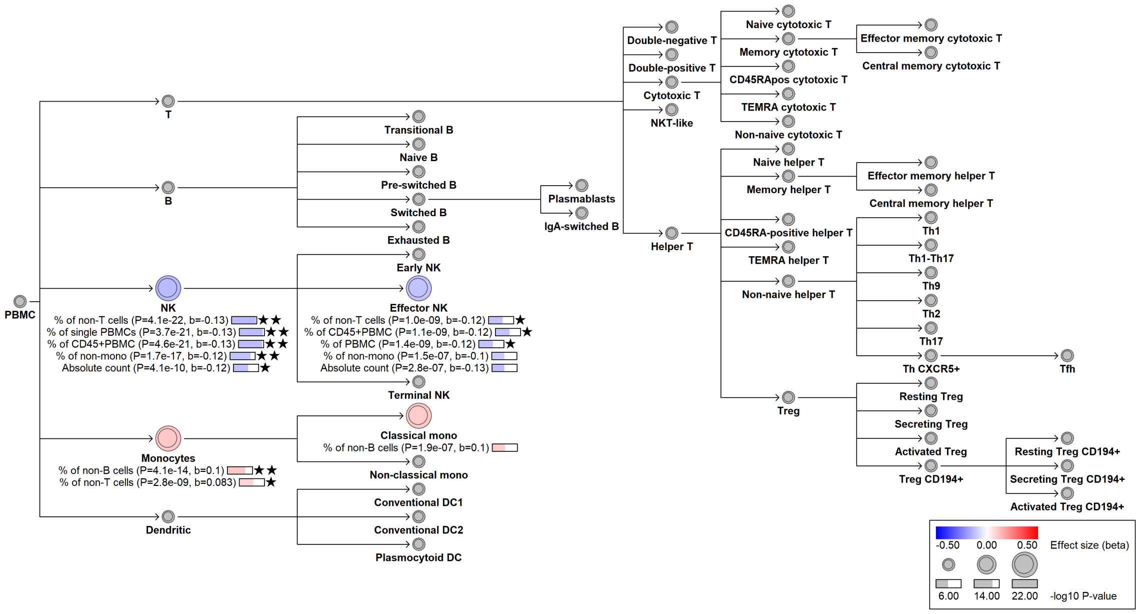

#### chr1:24965206[C→T; EAF 1.021%] (rs188468174, cs_116)

Candidate genes: *RUNX3*
Diseases associations: NA
Plasma pQTLs: Composite immunoglobulin trait (IgA*IgG*IgM), Composite immunoglobulin trait IgA*IgG, Composite immunoglobulin trait IgA/IgG, Composite immunoglobulin trait IgA/IgM, Composite immunoglobulin trait IgM/(IgA*IgG), Immunoglobulin A

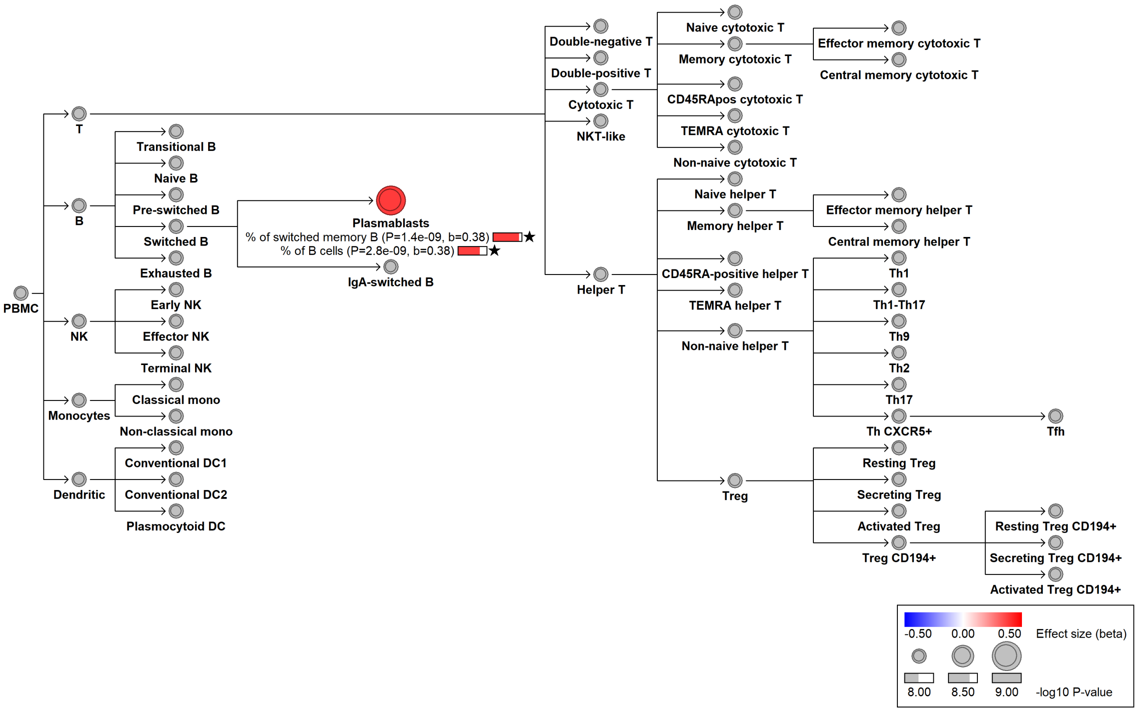

### chr1:26276065[C→G; EAF 34.553%] (rs12732837, cs_29)

Candidate genes: *CD52*
Diseases associations: NA
Plasma pQTLs: NA

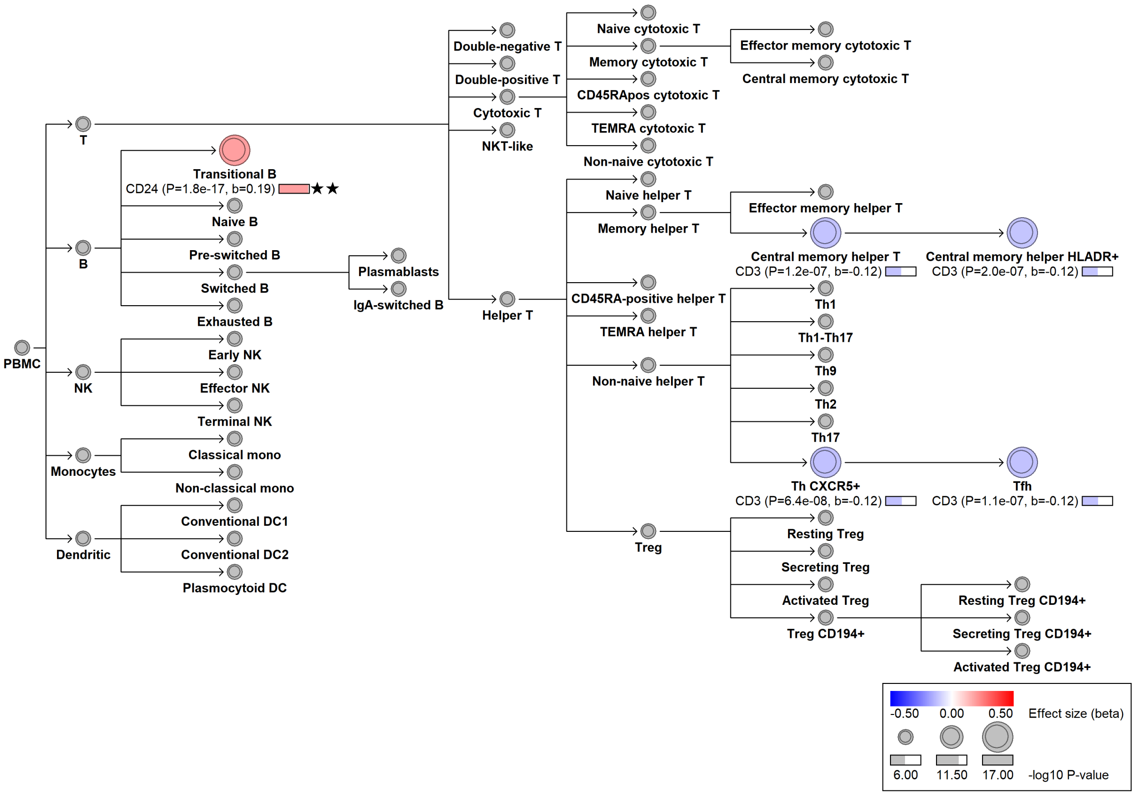

#### chr1:26282344[!→ACCGGGACCGGGACCGGGACTGGGG; EAF 17.518%] (rs1184004258, cs_60)

Candidate genes: *CD52*
Diseases associations: Asthma
Plasma pQTLs: NA

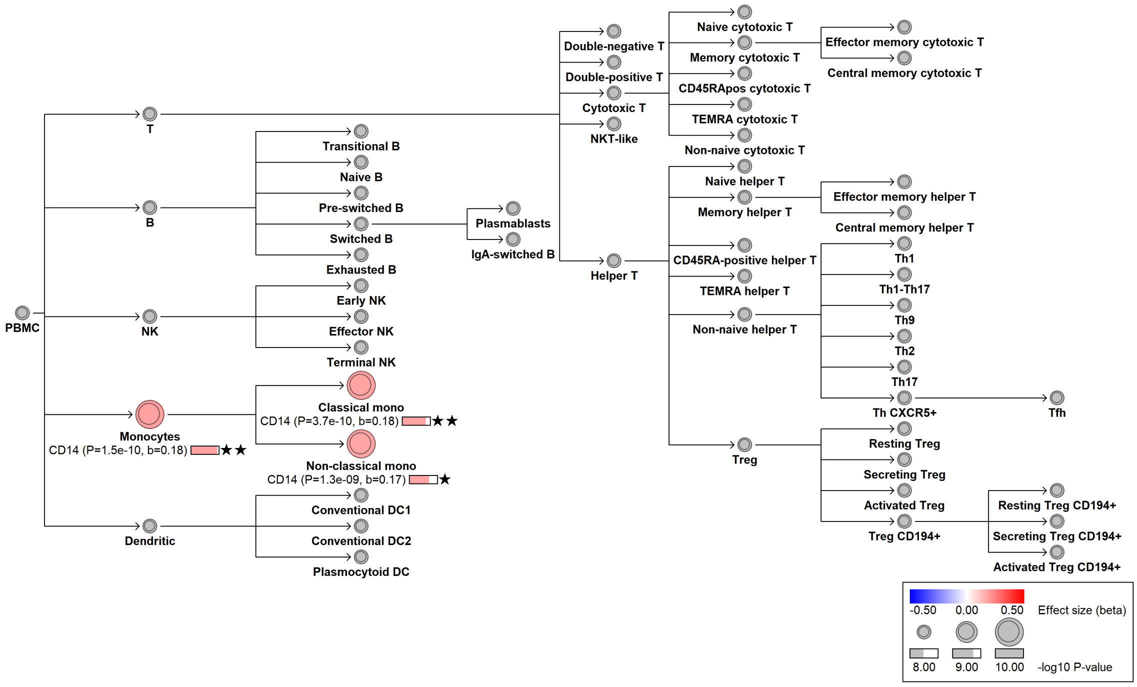

### chr1:26309245[ATTTTTT→!; EAF 34.647%] (rs1465206566, cs_19)

Candidate genes: *CD52*
Diseases associations: NA
Plasma pQTLs: NA

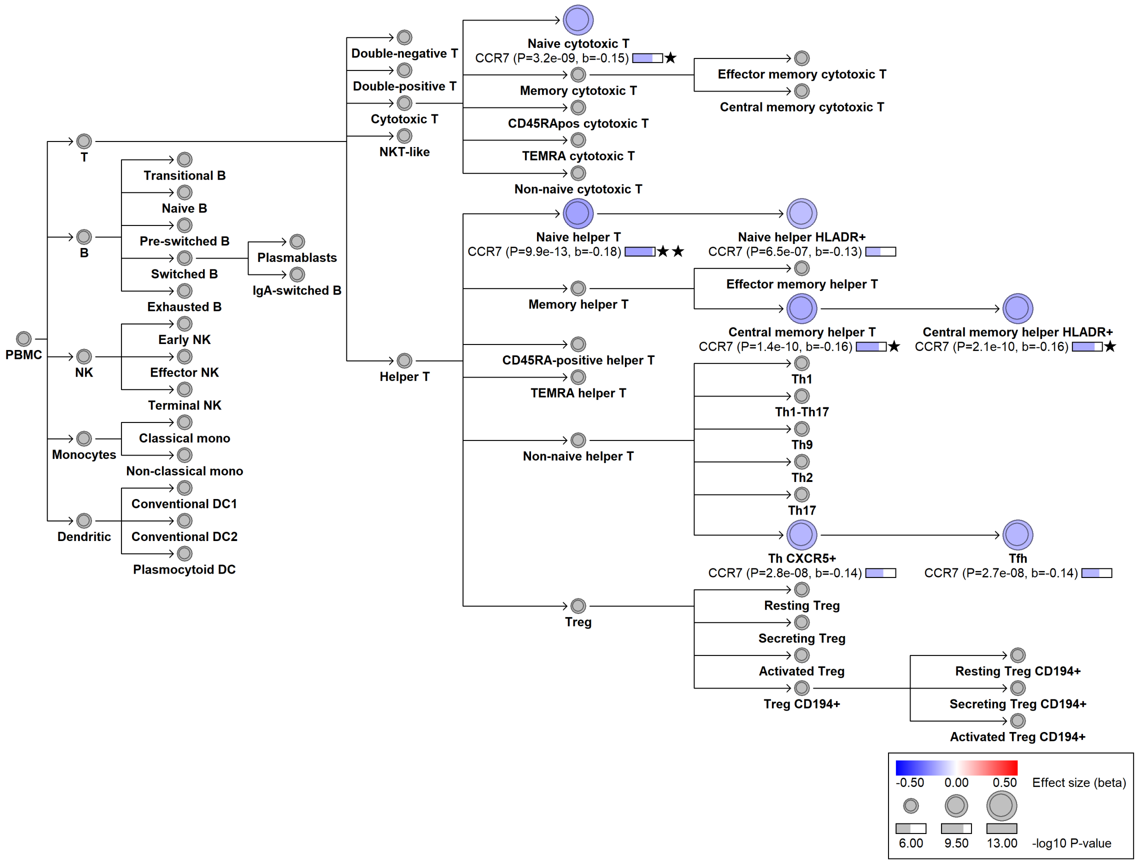

#### chr1:45331833[C→G; EAF 24.413%] (rs3219489, cs_171)

Candidate genes: *TESK2, MUTYH*
Diseases associations: Familial adenomatous polyposis 2, Glaucoma
Plasma pQTLs: CD5L, CDH3, Composite immunoglobulin trait IgA/IgM, Composite immunoglobulin trait IgG/IgM, Composite immunoglobulin trait IgM/(IgA*IgG), EDA2R, HS6ST2, Immunoglobulin A, Immunoglobulin M, TCL1A

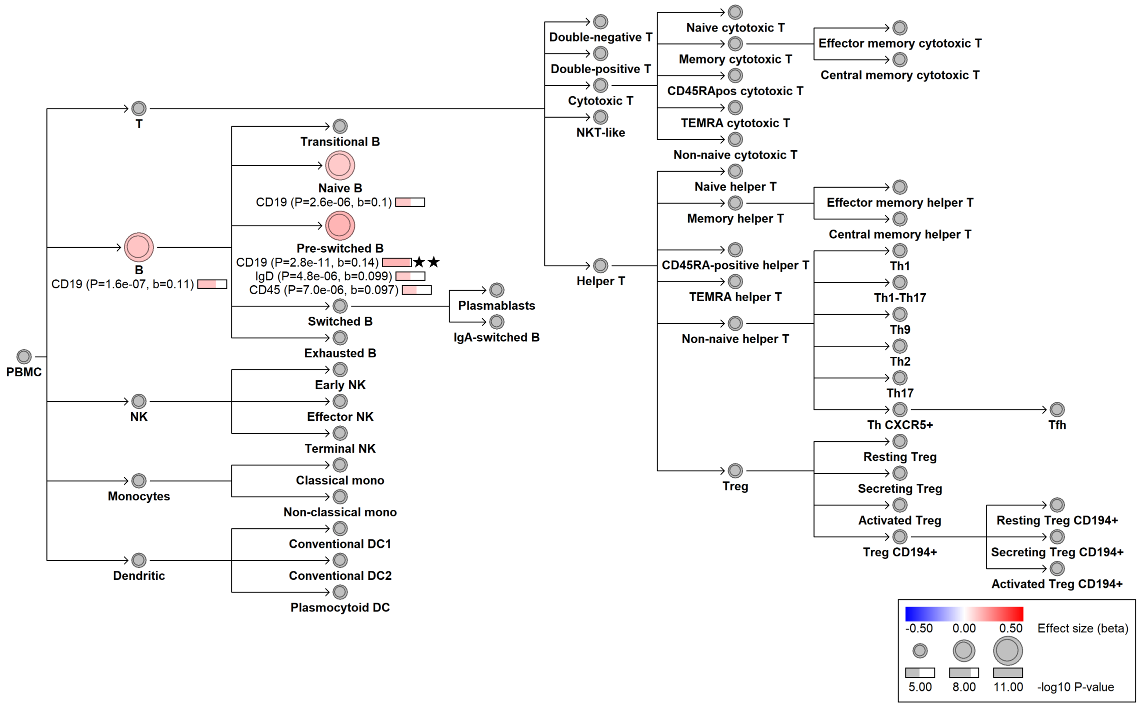

### chr1:92460097[G→C; EAF 2.88%] (rs150649461, cs_38)

Candidate genes: *GFI1*
Diseases associations: NA
Plasma pQTLs: CD177, CD86, IL1R2, IL22RA2, LILRA3, LILRA5, MANSC1, RETN, SEMA4A, TNFSF8

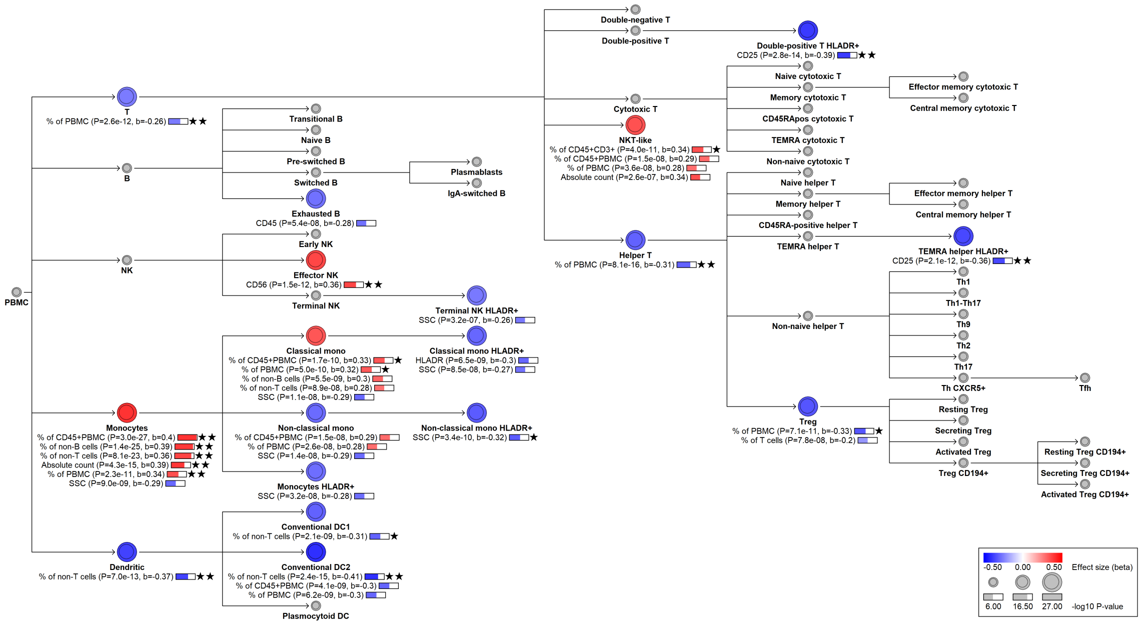

#### chr1:101268305[G→A; EAF 38.375%] (rs11166573, cs_149)

Candidate genes: *S1PR1*
Diseases associations: NA
Plasma pQTLs: GZMK

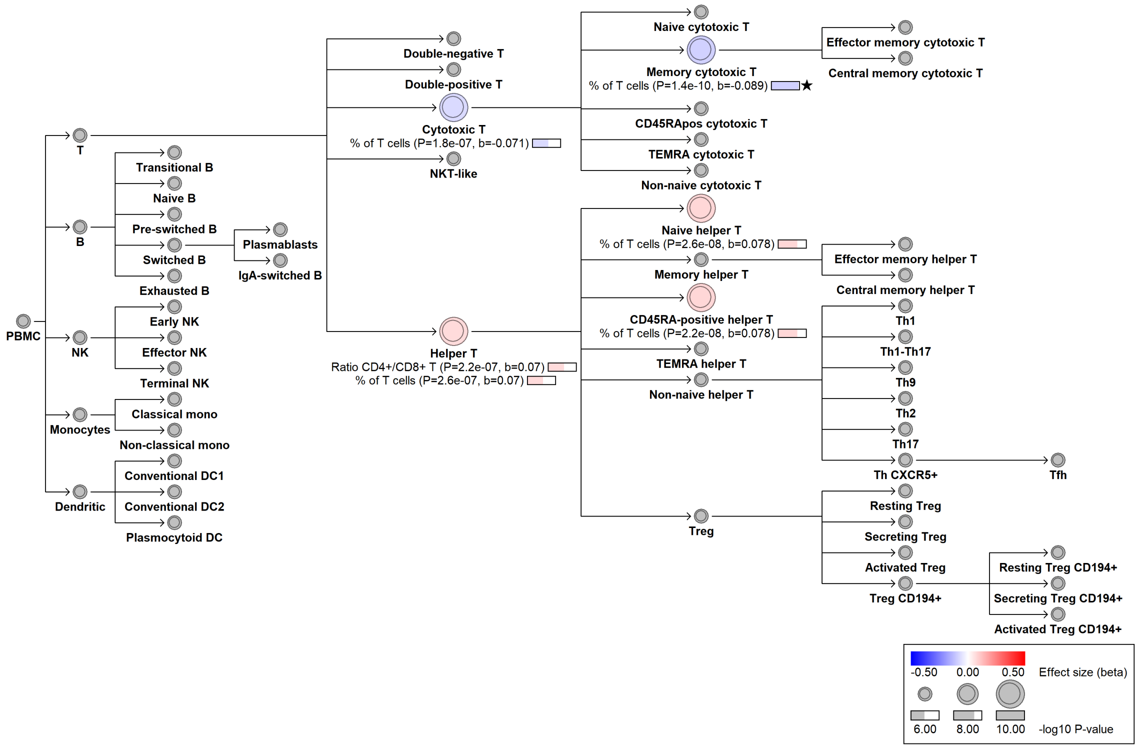

##### chr1:113834946[G→A; EAF 11.711%] (rs2476601, cs_40)

Candidate genes: *PTPN22*
Diseases associations: ANCA-associated vasculitis, Addison's disease, Ankylosing spondylitis, Autoimmune thyroiditis, Breast cancer, Cardiovascular disease, Celiac disease, Crohn's disease, Juvenile rheumatoid arthritis, Multiple sclerosis, Myasthenia gravis, Non-melanoma skin cancer, Polymyositis, Primary sclerosing cholangitis, Psoriasis, Rheumatoid arthritis, Systemic lupus erythematosus, Systemic sclerosis, Type 1 diabetes, Ulcerative colitis, Vitiligo
Plasma pQTLs: CAST, CCL19, CD5, CD5L, CHAD, COL1A1, CXCL10, CXCL9, Composite immunoglobulin trait (IgA*IgG*IgM), Composite immunoglobulin trait IgA/IgM, Composite immunoglobulin trait IgG/IgM, Composite immunoglobulin trait IgM/(IgA*IgG), GIF, GRID2, IL10, IL12A, IL12B, Immunoglobulin M, JCHAIN, KLRF1, PCDHA4, PDCD1

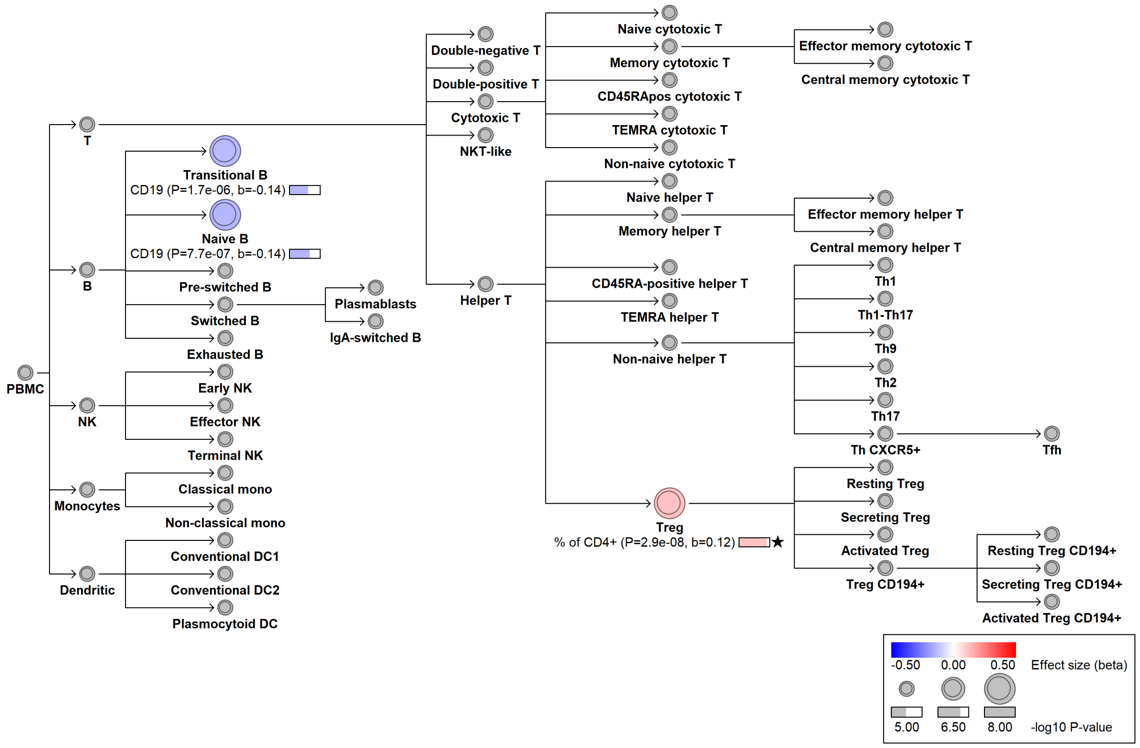

### chr1:155017119[C→T; EAF 44.534%] (rs6673081, cs_102)

Candidate genes: *ZBTB7B*
Diseases associations: NA
Plasma pQTLs: NA

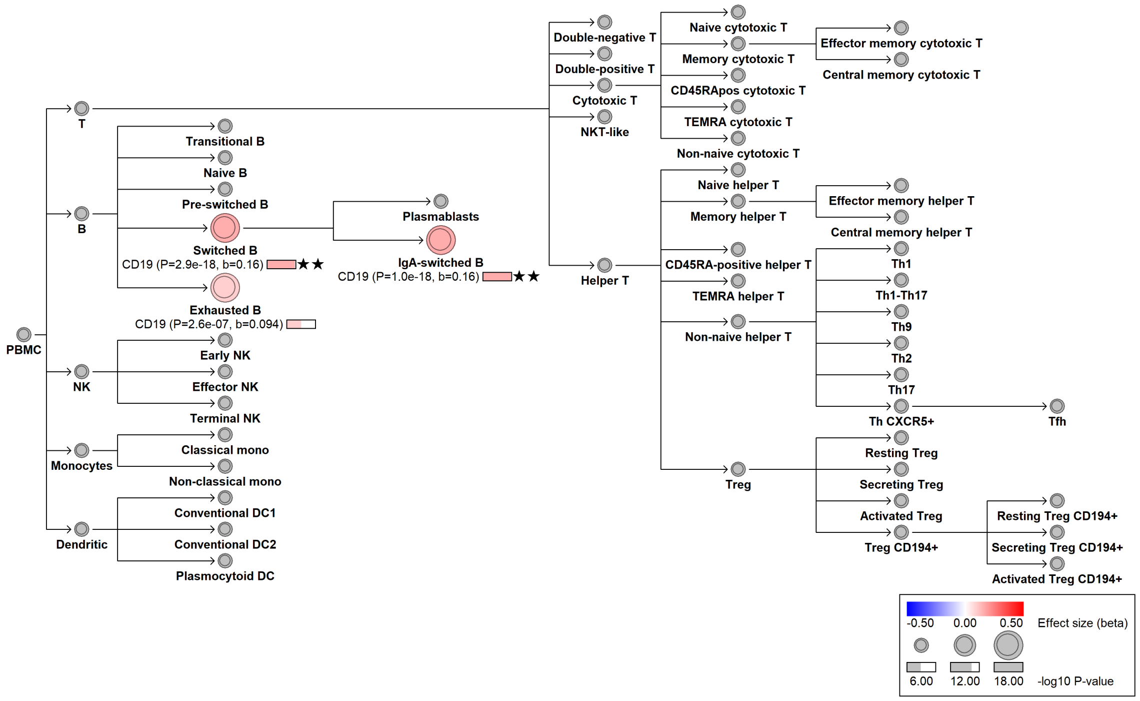

### chr1:157698600[T→C; EAF 45.624%] (rs7522061, cs_23)

Candidate genes: *FCRL3*
Diseases associations: NA
Plasma pQTLs: FCRL3, IGSF11

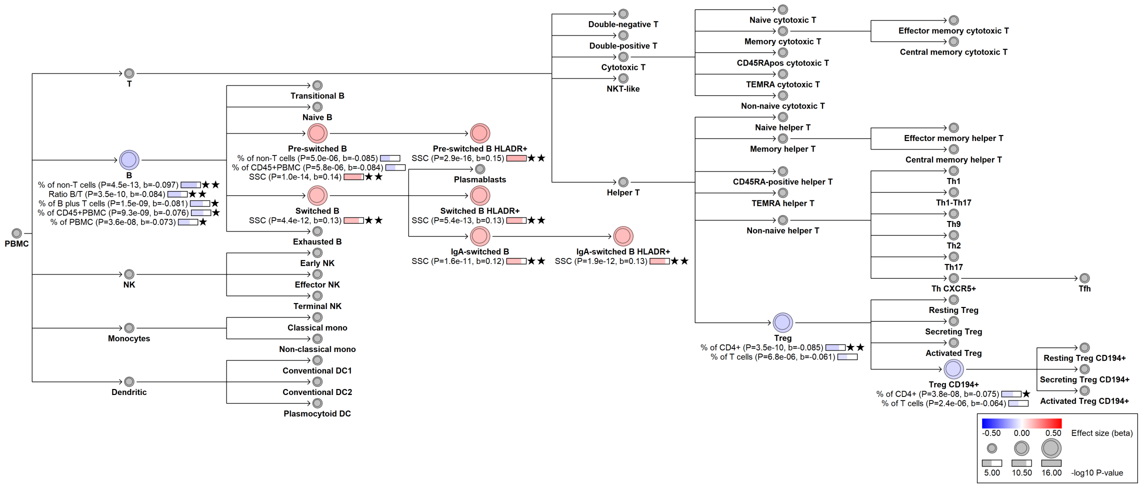

#### chr1:160823770[A→G; EAF 45.502%] (rs509749, cs_21)

Candidate genes: *LY9*
Diseases associations: NA
Plasma pQTLs: NA

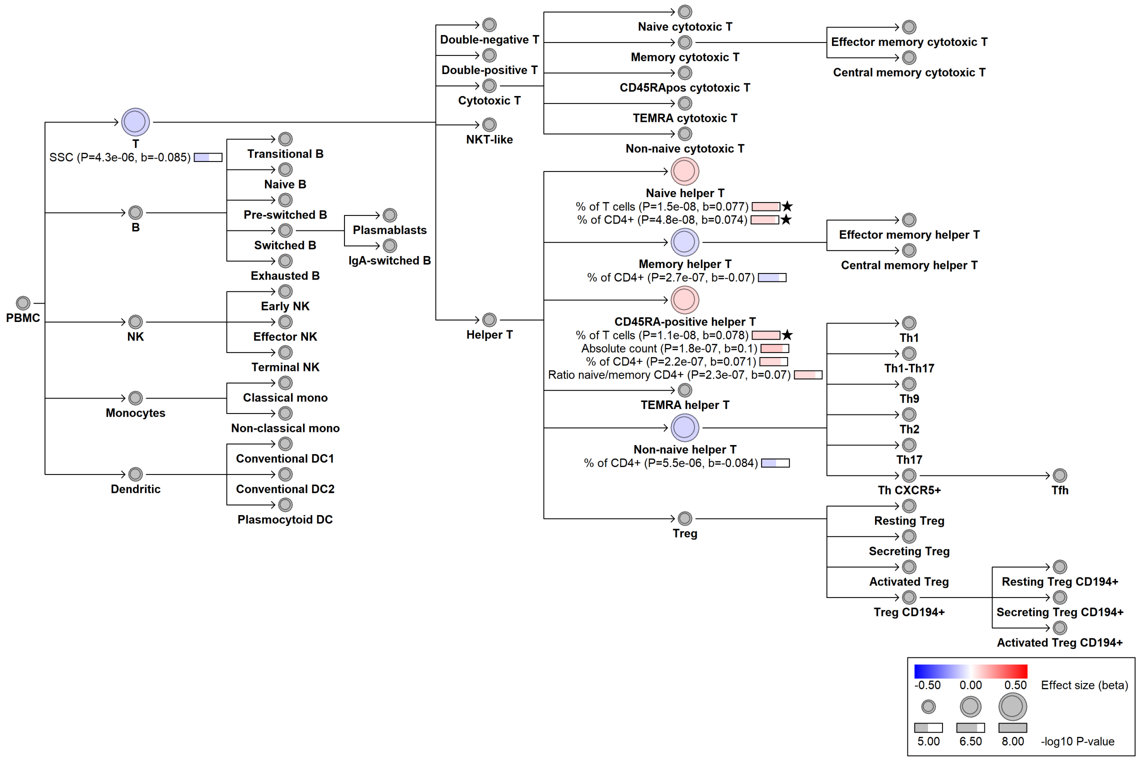

##### chr1:161509955[G$\rightarrow$A; EAF 48.936%] (rs1801274, cs_169)

Candidate genes: *FCGR2A*
Diseases associations: Ankylosing spondylitis, Crohn's disease, Inflammatory bowel disease, Kawasaki disease, Systemic lupus erythematosus, Ulcerative colitis
Plasma pQTLs: CD274, CILP2, FCGR2A, FCGR2B, MCEMP1, TYRP1

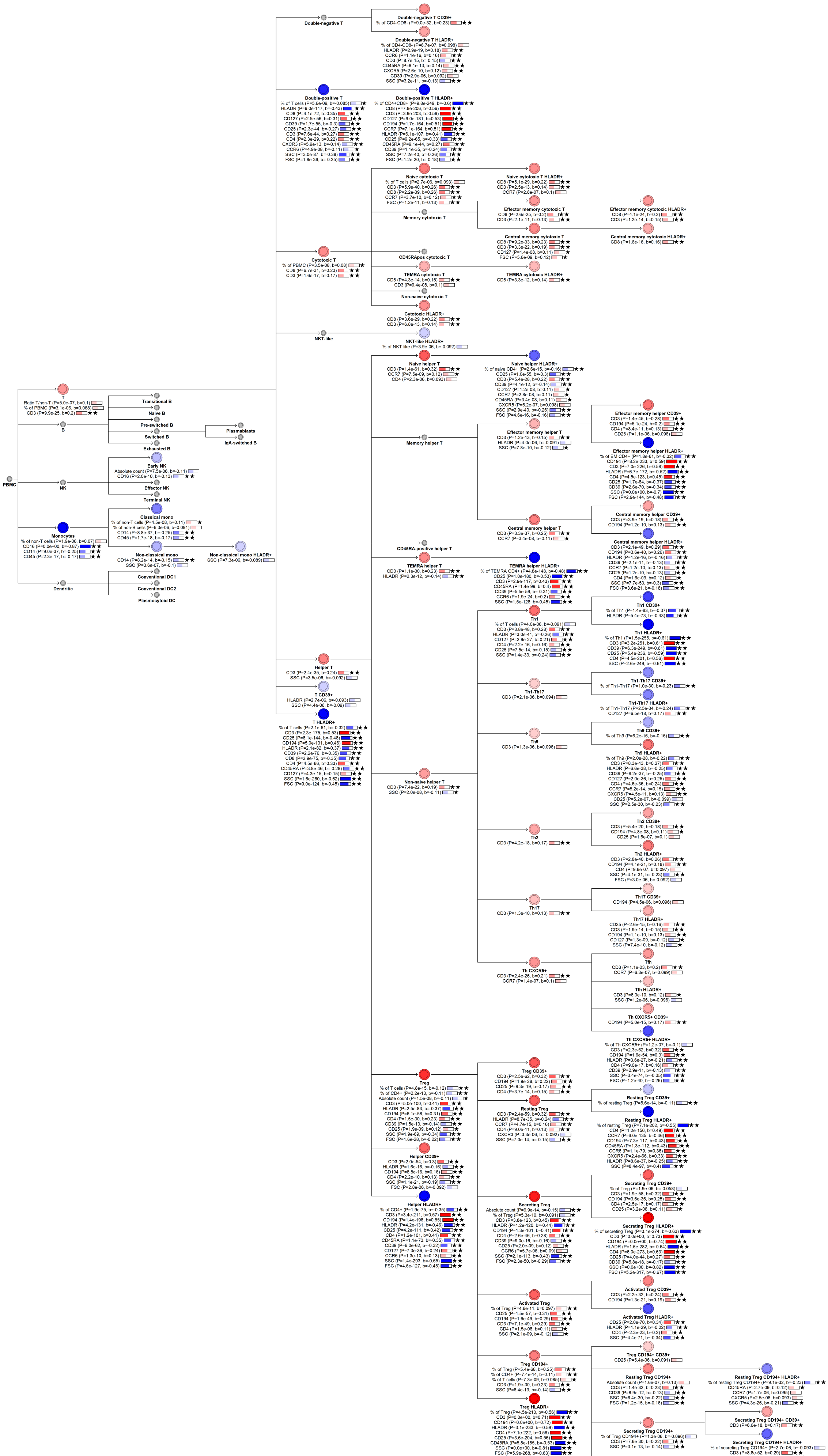

PBMC
T
B
NK
Monocytes
Dendritic
Transitional B
Naive B
Pre-switched B
Switched B
Exhausted B
Plasmablasts
IgA-switched B
Early NK
Effector NK
Terminal NK
Classical mono
Non-classical mono
Non-classical mono HLADR+
Conventional DC1
Conventional DC2
Plasmacytoid DC
Double-negative T
Double-negative T CD39+
Double-negative T HLADR+
Double-positive T
Double-positive T HLADR+
Cytotoxic T
Naive cytotoxic T
Naive cytotoxic T HLADR+
Memory cytotoxic T
Effector memory cytotoxic T
Effector memory cytotoxic HLADR+
Central memory cytotoxic T
Central memory cytotoxic HLADR+
CD45RApos cytotoxic T
TEMRA cytotoxic T
TEMRA cytotoxic HLADR+
Non-naive cytotoxic T
Cytotoxic HLADR+
NKT-like
NKT-like HLADR+
Helper T
T CD39+
T HLADR+
Naive helper T
Naive helper HLADR+
Memory helper T
Effector memory helper T
Effector memory helper CD39+
Effector memory helper HLADR+
Central memory helper T
Central memory helper CD39+
Central memory helper HLADR+
CD45RA-positive helper T
TEMRA helper T
TEMRA helper HLADR+
Non-naive helper T
Th1
Th1 CD39+
Th1 HLADR+
Th1-Th17
Th1-Th17 CD39+
Th1-Th17 HLADR+
Th9
Th9 CD39+
Th9 HLADR+
Th2
Th2 CD39+
Th2 HLADR+
Th17
Th17 CD39+
Th17 HLADR+
Th CXCR5+
Tfh
Tfh HLADR+
Th CXCR5+ CD39+
Th CXCR5+ HLADR+
Treg
Treg CD39+
Resting Treg
Resting Treg CD39+
Resting Treg HLADR+
Secreting Treg
Secreting Treg CD39+
Secreting Treg HLADR+
Activated Treg
Activated Treg CD39+
Activated Treg HLADR+
Treg CD194+
Treg CD194+ CD39+
Resting Treg CD194+
Resting Treg CD194+ HLADR+
Secreting Treg CD194+
Secreting Treg CD194+ CD39+
Secreting Treg CD194+ HLADR+
Treg HLADR+
Helper CD39+
Helper HLADR+

#### chr1:161521688[C→T; EAF 35.677%] (rs1811422, cs_95)

Candidate genes: *FCGR3A*
Diseases associations: NA
Plasma pQTLs: NA

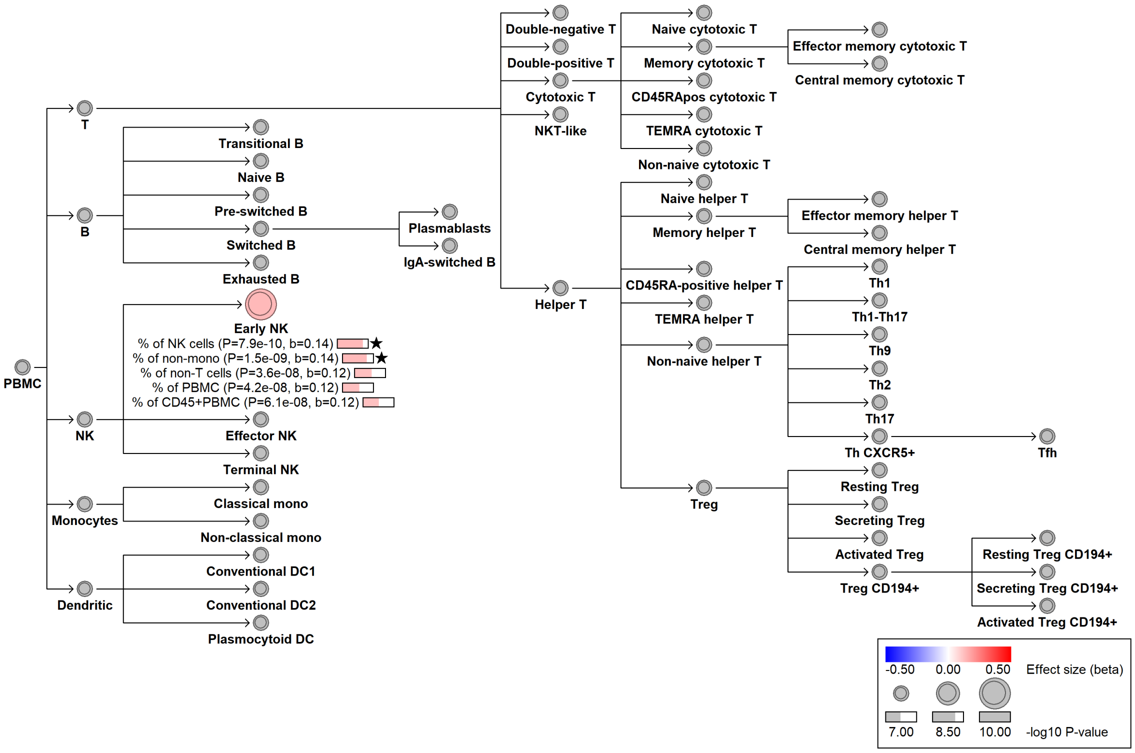

### chr1:161544752[A→C; EAF 32.848%] (rs396991, cs_151)

Candidate genes: *FCGR3A, FCGR3B*
Diseases associations: NA
Plasma pQTLs: ADGRE1, CD163, FCGR3A, FCGR3B, IDO1

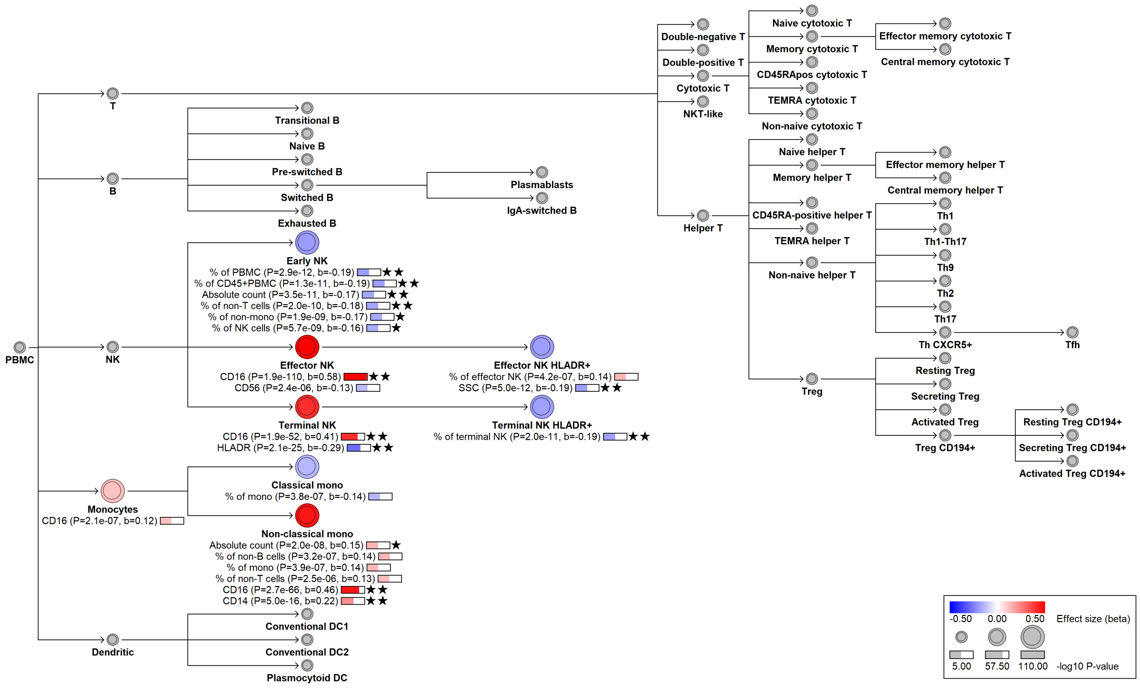

#### chr1:161618290[T→A; EAF 37.054%] (rs61802324, cs_48)

Candidate genes: *FCGR3A, FCGR3B*
Diseases associations: NA
Plasma pQTLs: NA

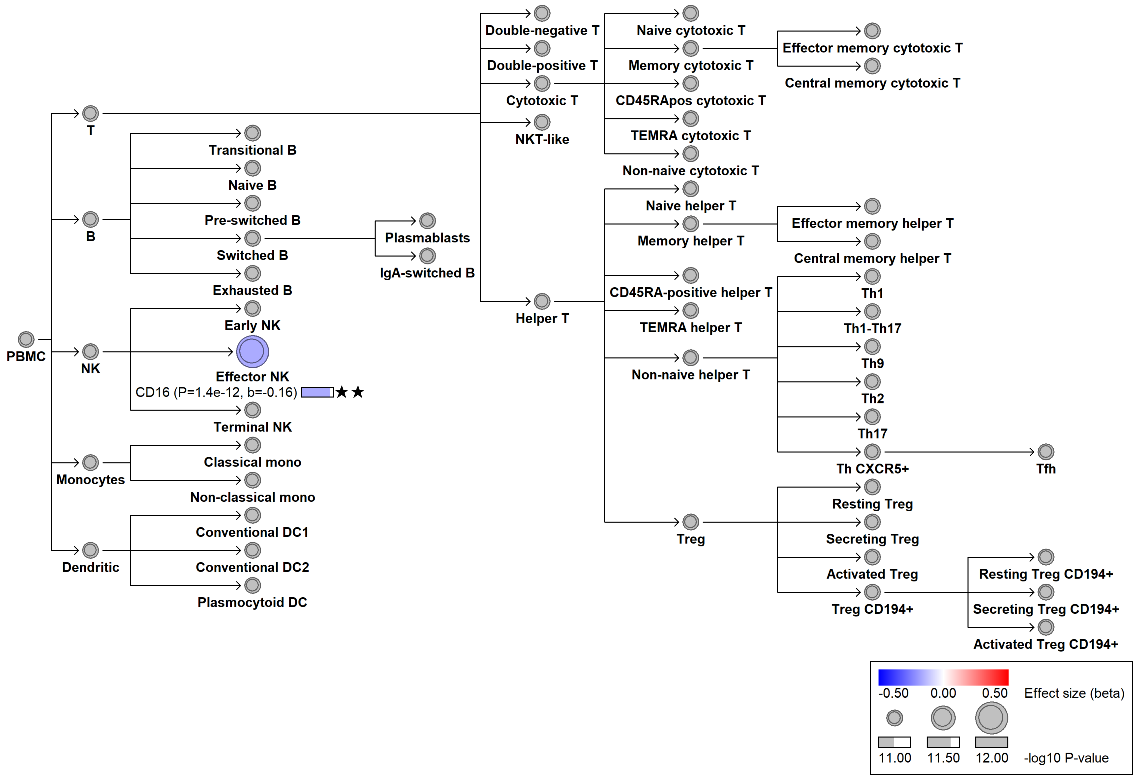

### chr1:161647883[G→A; EAF 7.842%] (rs140315354, cs_7)

Candidate genes: *FCGR3A, FCGR3B*
Diseases associations: NA
Plasma pQTLs: NA

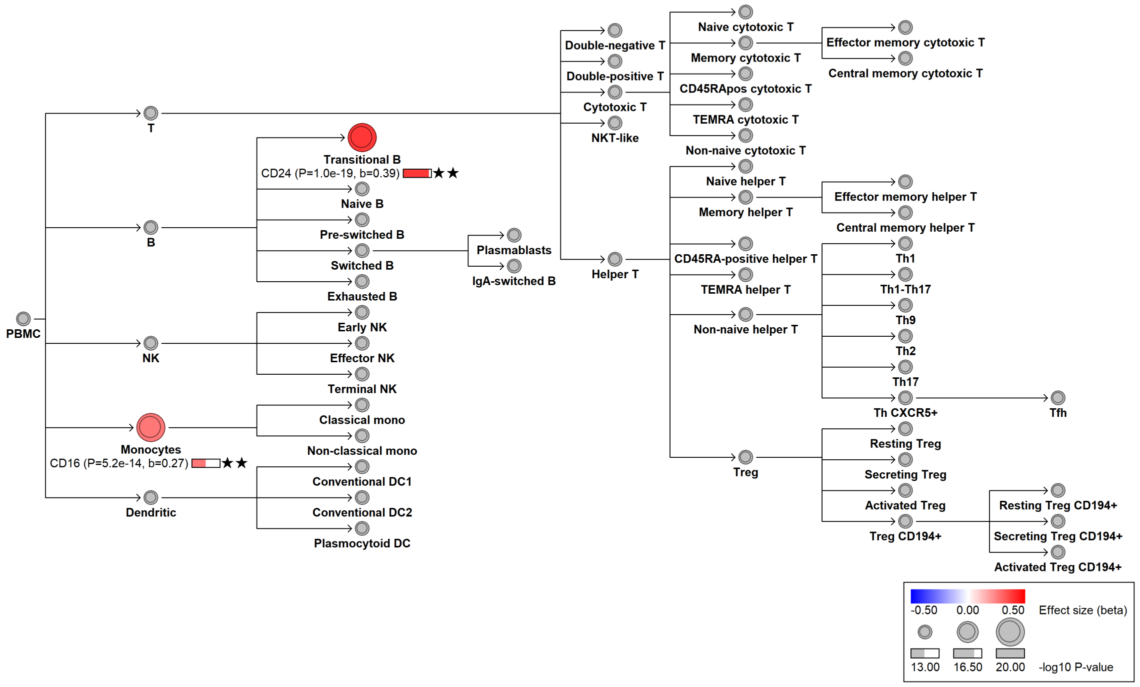

#### chr1:161656087[G→C; EAF 29.798%] (rs78217950, cs_1)

Candidate genes: *FCGR3A, FCGR3B*
Diseases associations: Systemic lupus erythematosus
Plasma pQTLs: NA

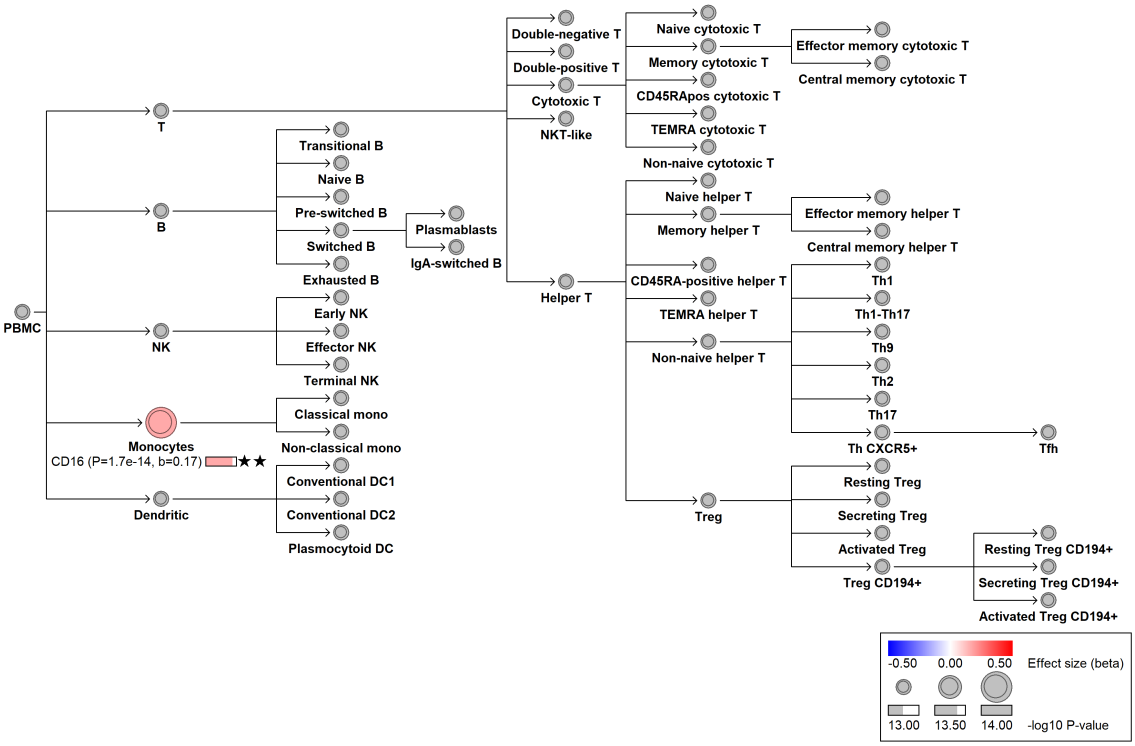

#### chr1:165941996[C→A; EAF 0.141%] (rs765477172, cs_110)

Candidate genes: *CD247*
Diseases associations: NA
Plasma pQTLs: NA

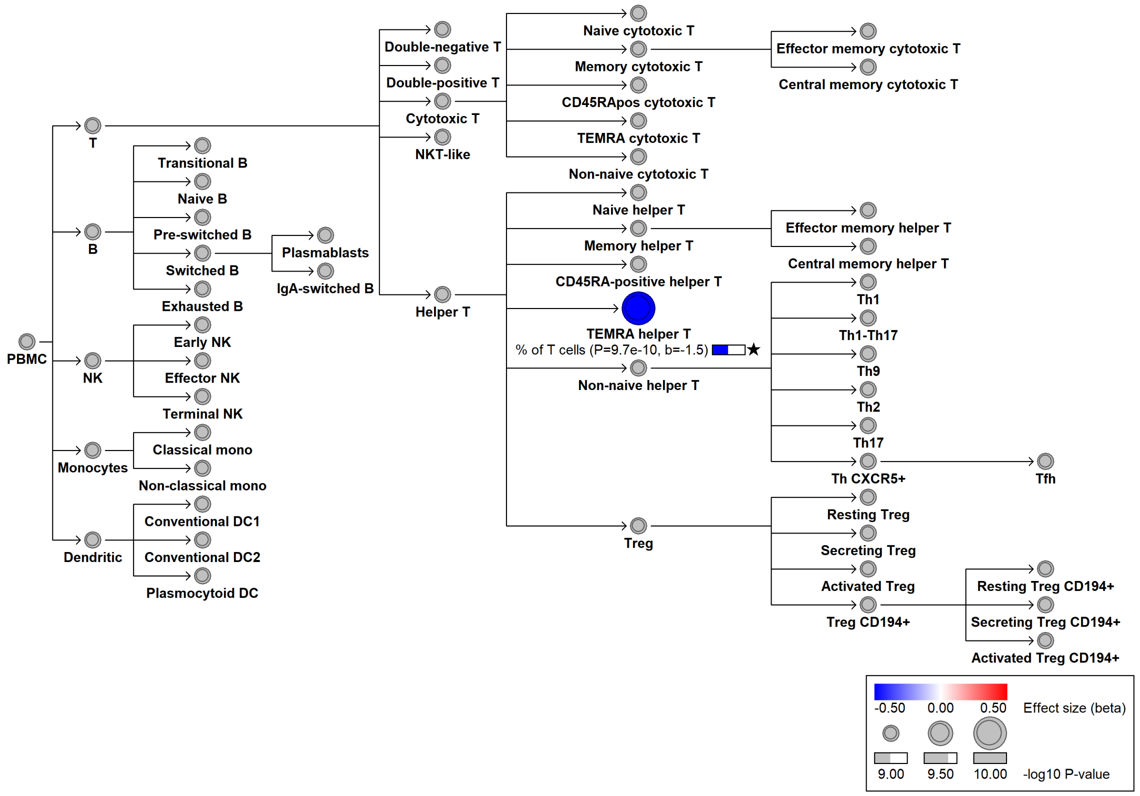

### chr1:167430954[A→AC; EAF 11.19%] (rs3831958, cs_109)

Candidate genes: *CD247*
Diseases associations: NA
Plasma pQTLs: NA

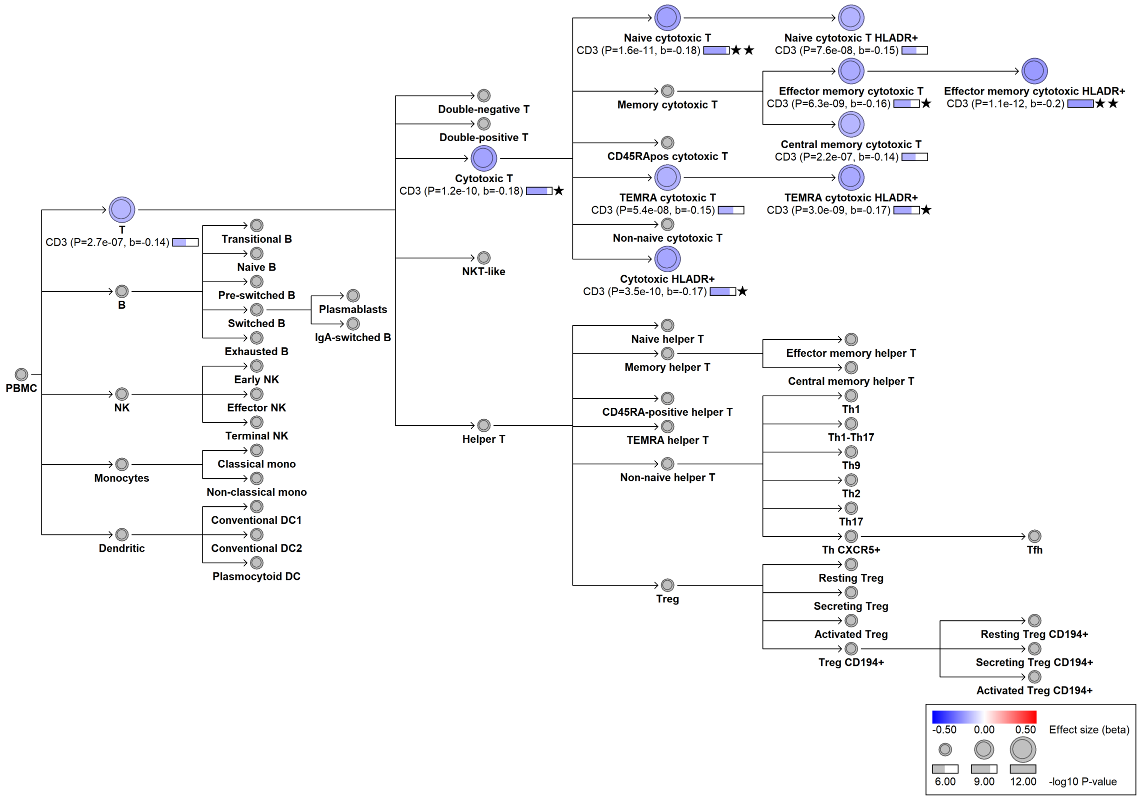

### chr1:167444249[C→G; EAF 2.185%] (rs35240301, cs_67)

Candidate genes: *CD247*
Diseases associations: NA
Plasma pQTLs: NA

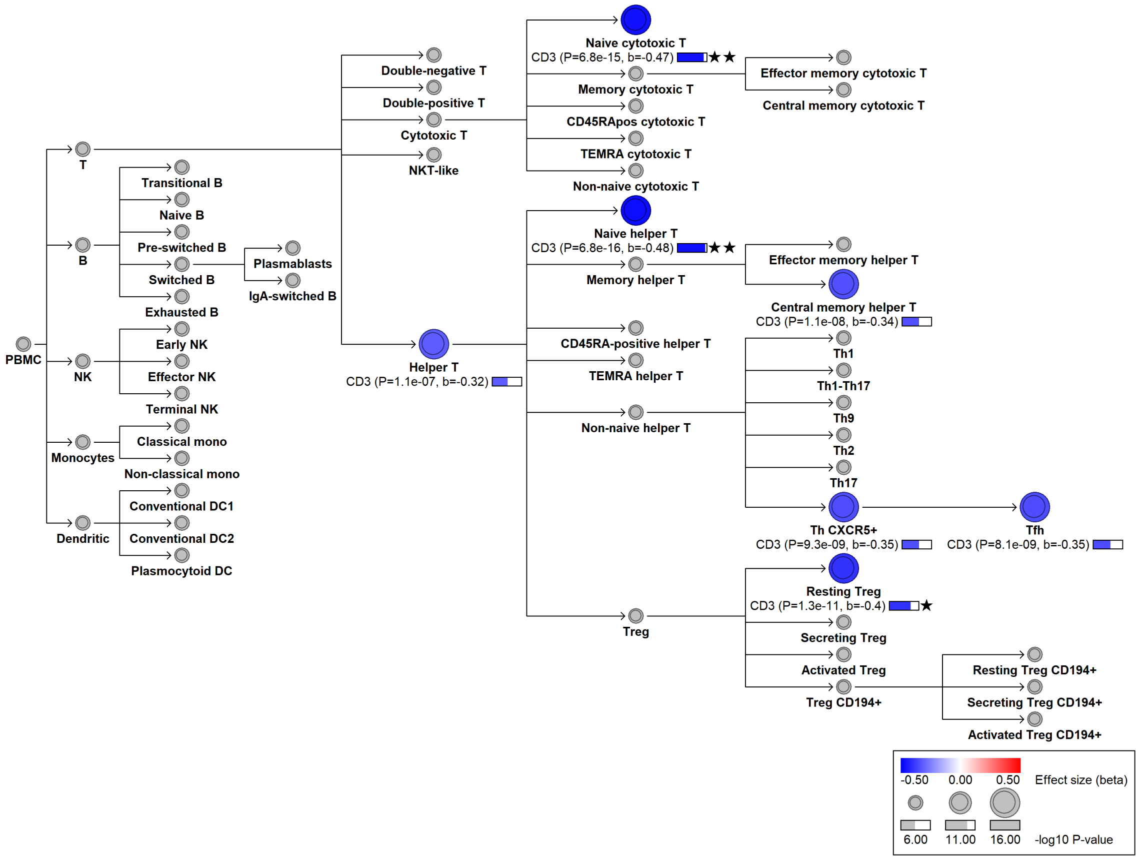

### chr1:167455687[C→!; EAF 35.298%] (chr1:167455687, cs_177)

Candidate genes: *CD247*
Diseases associations: Asthma, Autoimmune thyroiditis
Plasma pQTLs: NA

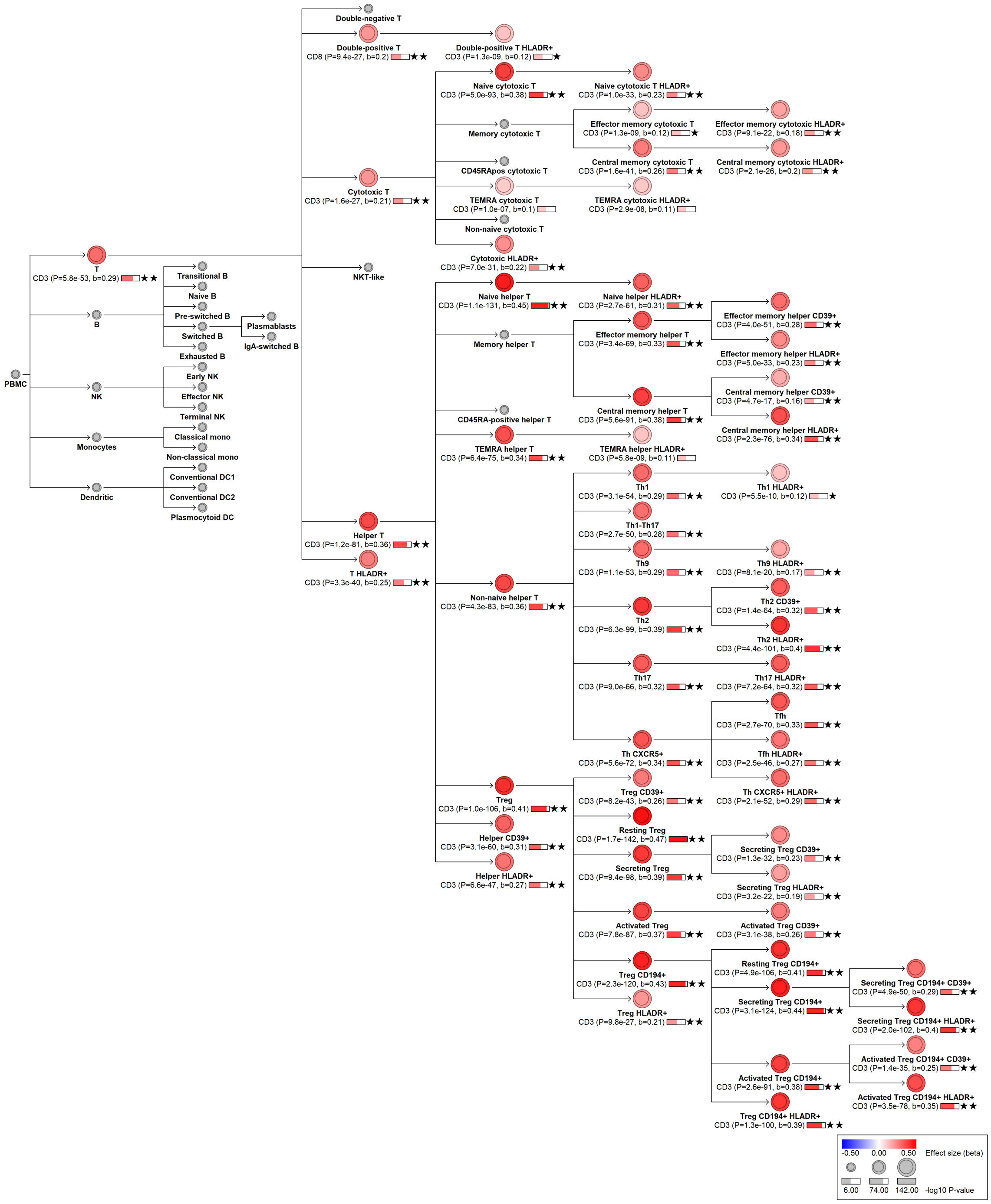

### chr1:167518643[!→T; EAF 7.148%] (rs3820390, cs_127)

Candidate genes: *CD247*
Diseases associations: NA
Plasma pQTLs: NA

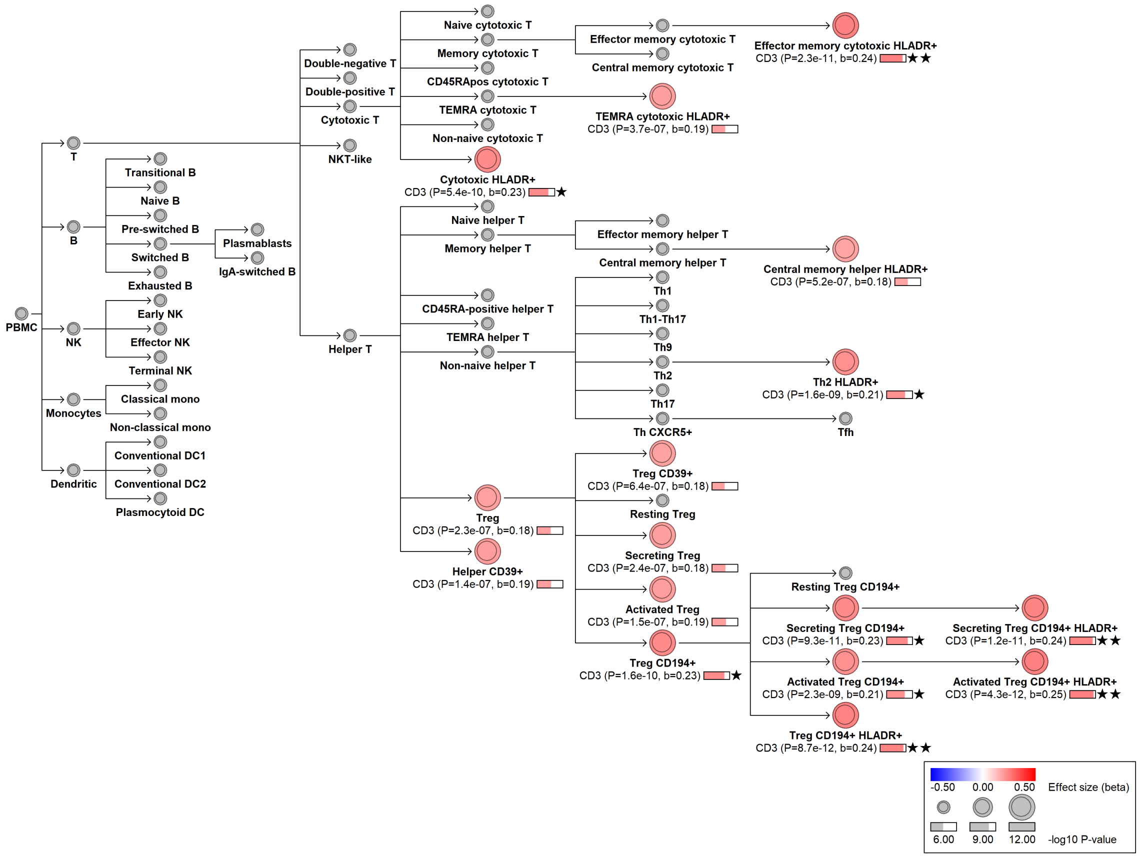

#### chr1:173416334[A→G; EAF 19.792%] (rs4554754, cs_39)

Candidate genes: *RABGAP1L, GAS5*
Diseases associations: Rheumatoid arthritis, Systemic lupus erythematosus, Systemic sclerosis
Plasma pQTLs: FCRL2, JCHAIN

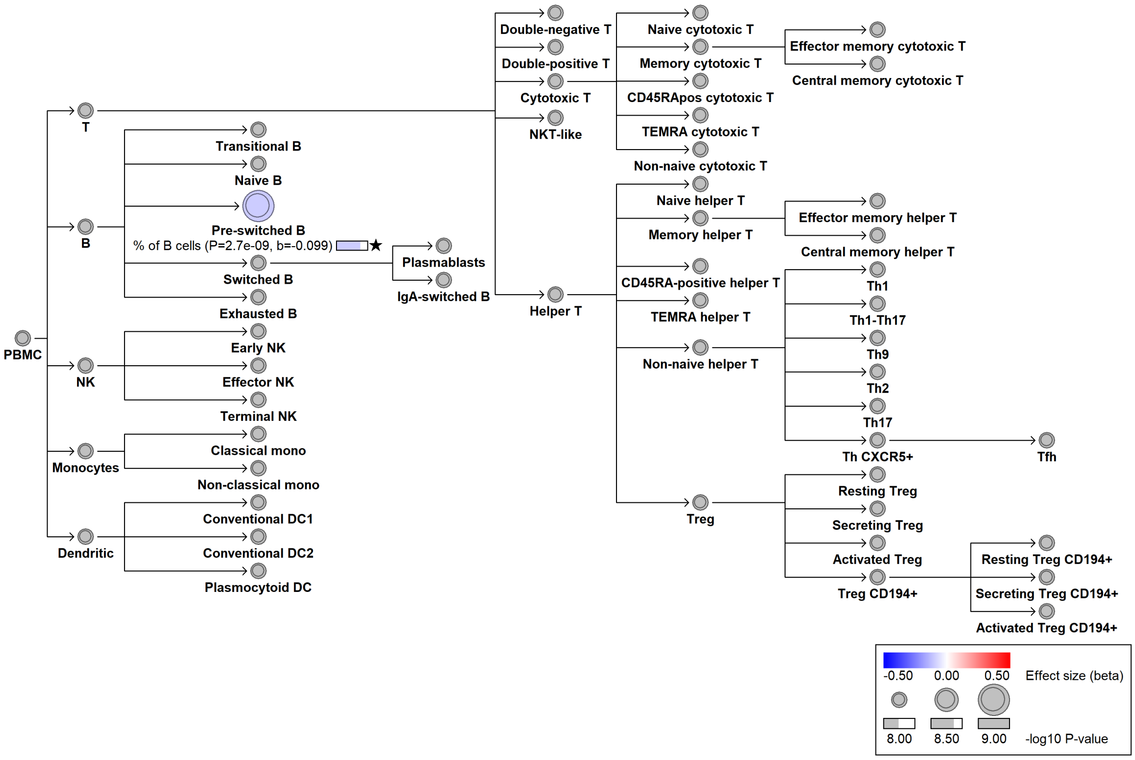

#### chr1:198638824[T→A; EAF 3.119%] (rs2274364, cs_6)

Candidate genes: *PTPRC*
Diseases associations: NA
Plasma pQTLs: NA

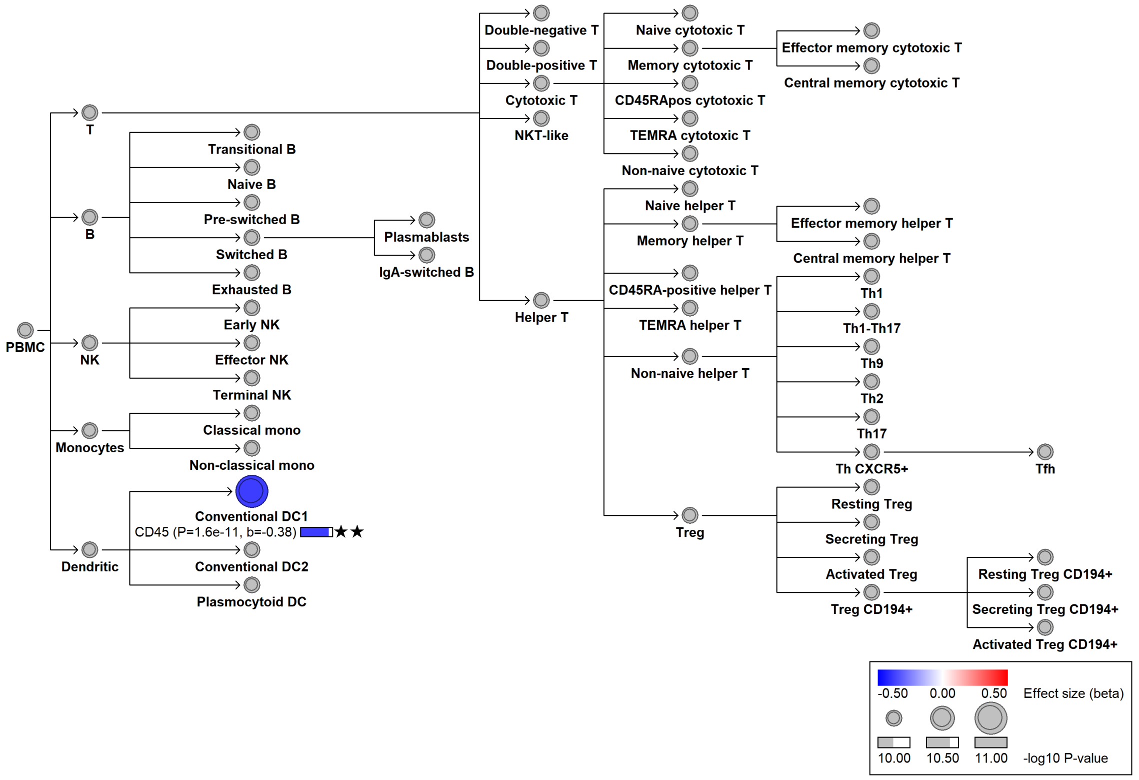

**chr1:198696788[!→G; EAF 1.444%] (rs17612648, cs_104)**

Candidate genes: *PTPRC*
Diseases associations: NA
Plasma pQTLs: NA

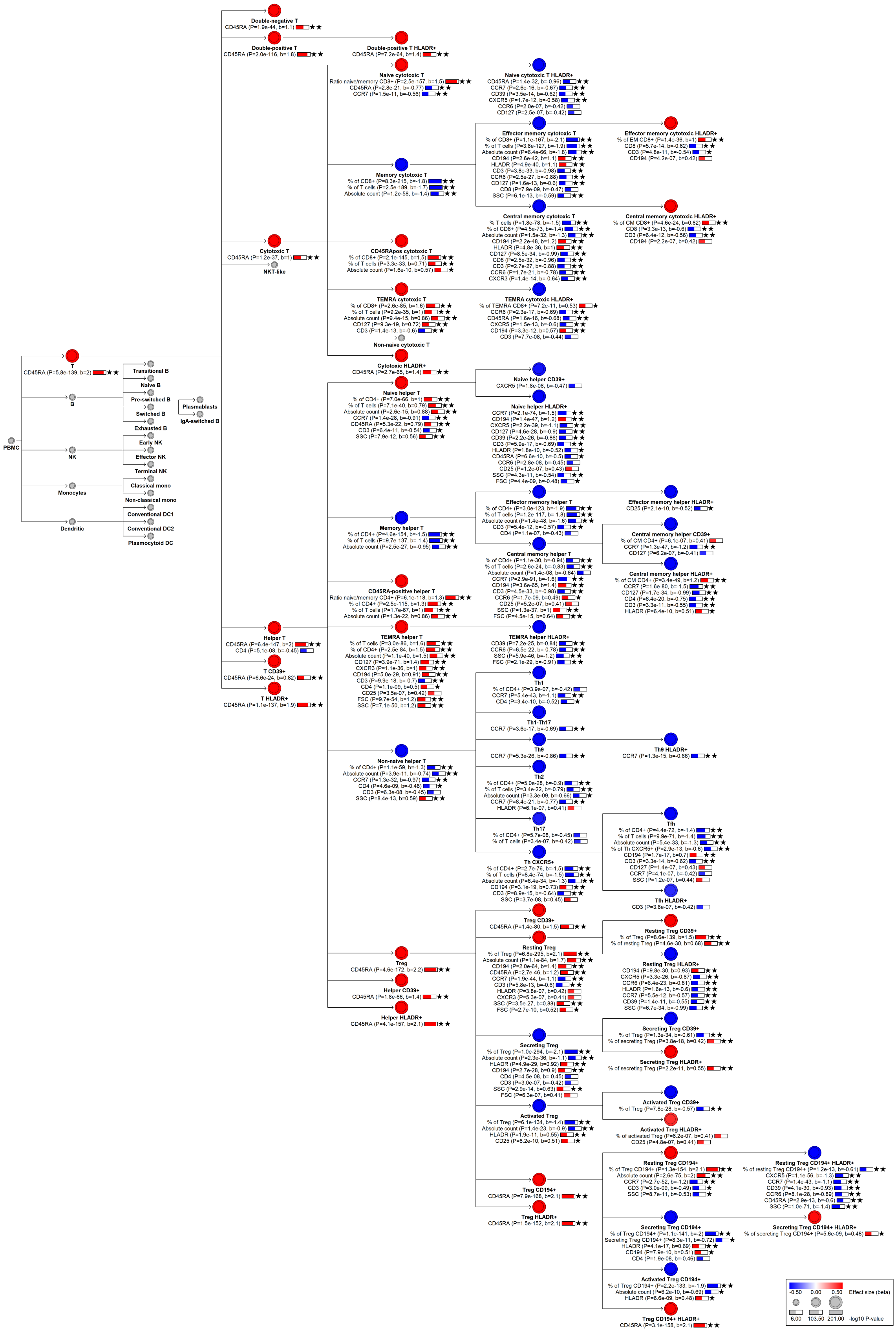

PBMC
T
CD45RA (P=5.8e-139, b=2)
B
NK
Monocytes
Dendritic
Transitional B
Naive B
Pre-switched B
Switched B
Exhausted B
Plasmablasts
IgA-switched B
Early NK
Effector NK
Terminal NK
Classical mono
Non-classical mono
Conventional DC1
Conventional DC2
Plasmacytoid DC
Double-negative T
Double-positive T
Double-positive T HLADR+
Cytotoxic T
NKT-like
Helper T
T CD39+
T HLADR+
Naive cytotoxic T
Naive cytotoxic T HLADR+
Effector memory cytotoxic T
Effector memory cytotoxic HLADR+
Memory cytotoxic T
Central memory cytotoxic T
Central memory cytotoxic HLADR+
CD45RApos cytotoxic T
TEMRA cytotoxic T
TEMRA cytotoxic HLADR+
Non-naive cytotoxic T
Cytotoxic HLADR+
Naive helper T
Naive helper CD39+
Naive helper HLADR+
Memory helper T
Effector memory helper T
Effector memory helper HLADR+
Central memory helper T
Central memory helper CD39+
Central memory helper HLADR+
CD45RA-positive helper T
TEMRA helper T
TEMRA helper HLADR+
Non-naive helper T
Th1
Th1-Th17
Th9
Th9 HLADR+
Th2
Th17
Tfh
Tfh HLADR+
Th CXCR5+
Treg
Helper CD39+
Helper HLADR+
Treg CD39+
Resting Treg
Resting Treg CD39+
Resting Treg HLADR+
Secreting Treg
Secreting Treg CD39+
Secreting Treg HLADR+
Activated Treg
Activated Treg CD39+
Activated Treg HLADR+
Treg CD194+
Resting Treg CD194+
Resting Treg CD194+ HLADR+
Secreting Treg CD194+
Secreting Treg CD194+ HLADR+
Activated Treg CD194+
Treg HLADR+
Treg CD194+ HLADR+
-0.50 0.00 0.50 Effect size (beta)
6.00 103.50 201.00 -log10 P-value

#### chr1:198699632[G→C; EAF 1.389%] (rs41269905, cs_125)

Candidate genes: *PTPRC*
Diseases associations: NA
Plasma pQTLs: NA

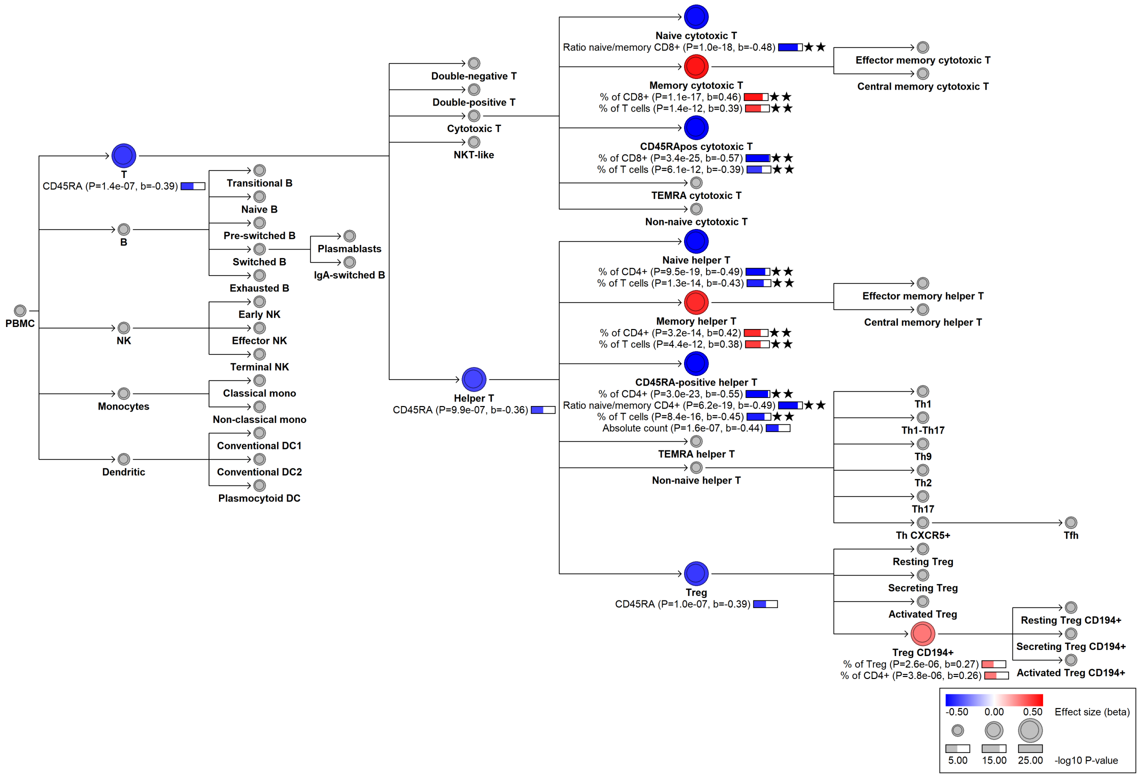

#### chr1:198710886[G→A; EAF 1.192%] (rs12733073, cs_139)

Candidate genes: *PTPRC*
Diseases associations: NA
Plasma pQTLs: FCER2, FLT3LG, IL22RA2, LTA, TCL1A

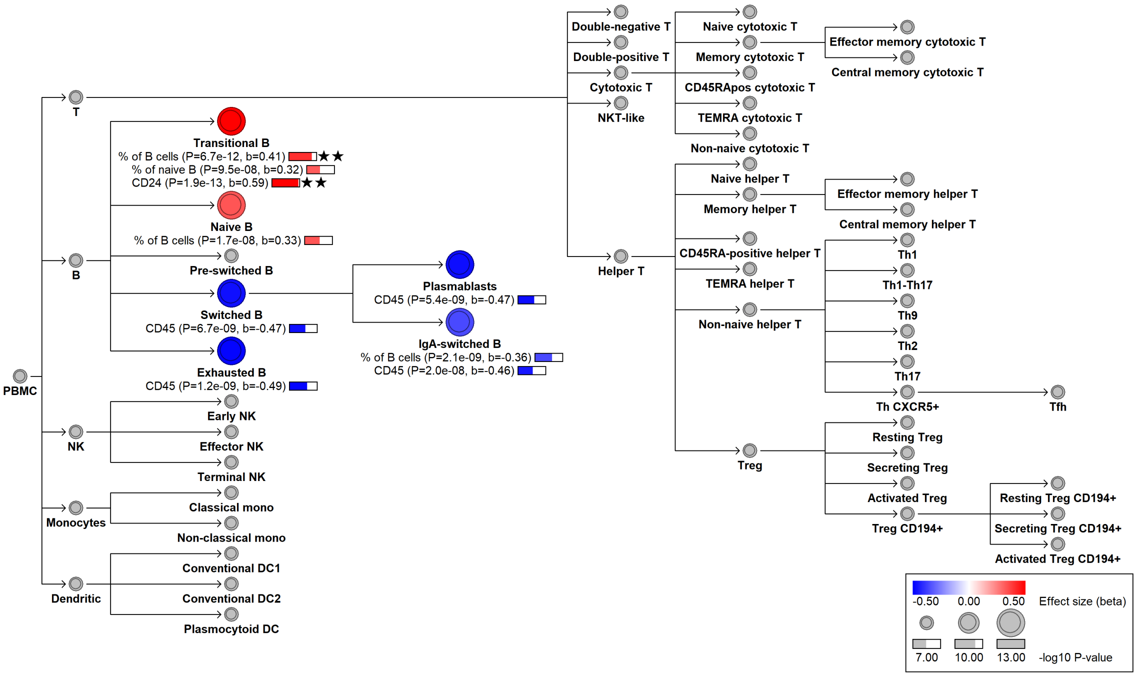

**chr1:200908722[!→G; EAF 27.404%] (rs41299637, cs_141)**

Candidate genes: *INAVA*
Diseases associations: Ankylosing spondylitis, Celiac disease, Crohn's disease, Inflammatory bowel disease, Multiple sclerosis, Primary sclerosing cholangitis, Psoriasis, Ulcerative colitis
Plasma pQTLs: Composite immunoglobulin trait IgA/IgG, Composite immunoglobulin trait IgA/IgM, Composite immunoglobulin trait IgM/(IgA*IgG), Immunoglobulin A

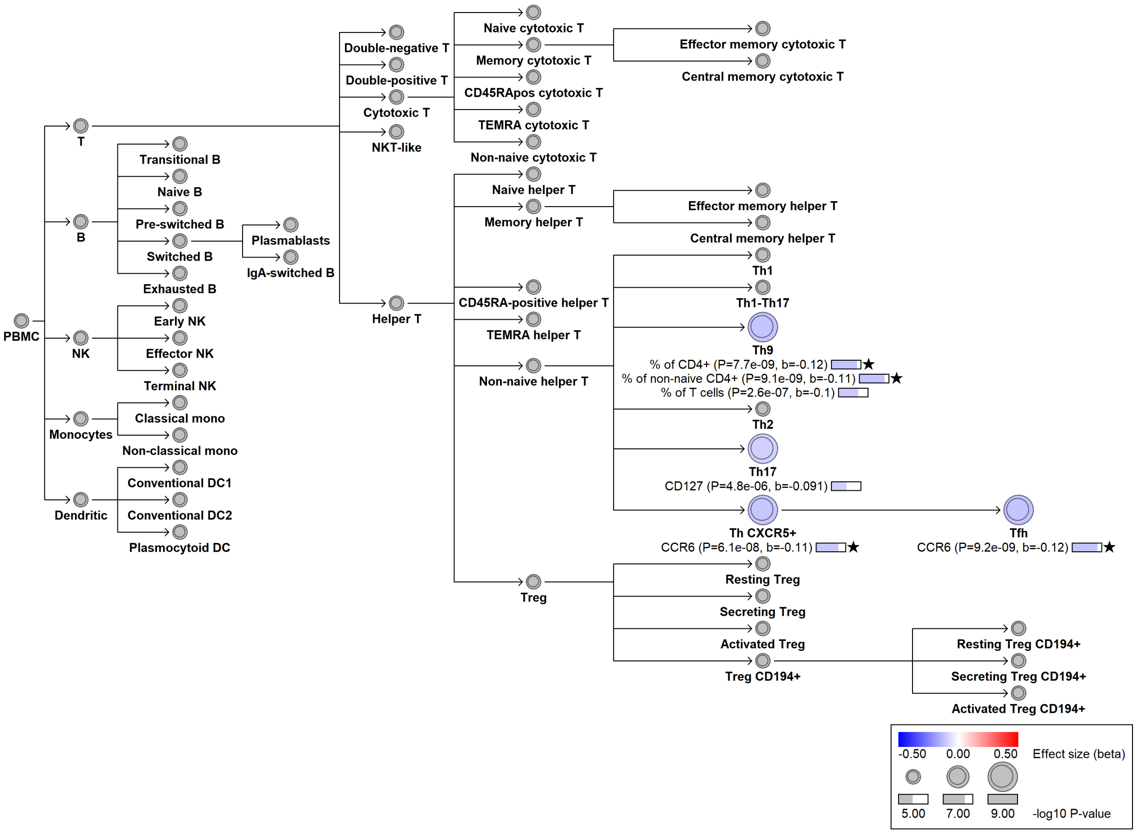

#### chr1:207416965[!→CA; EAF 11.28%] (rs767535979, cs_173)

Candidate genes: *CR2*
Diseases associations: NA
Plasma pQTLs: CR2

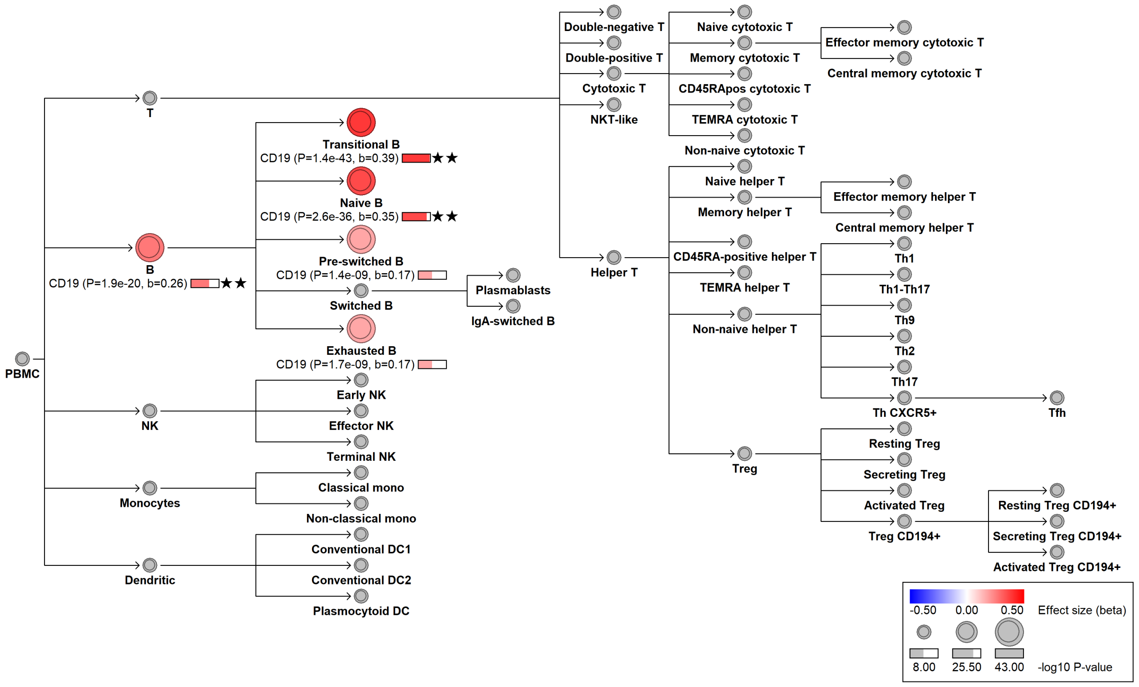

#### chr1:207474298[G→A; EAF 0.283%] (rs151093663, cs_223)

Candidate genes: *CR2*
Diseases associations: Common variable immunodeficiency
Plasma pQTLs: NA

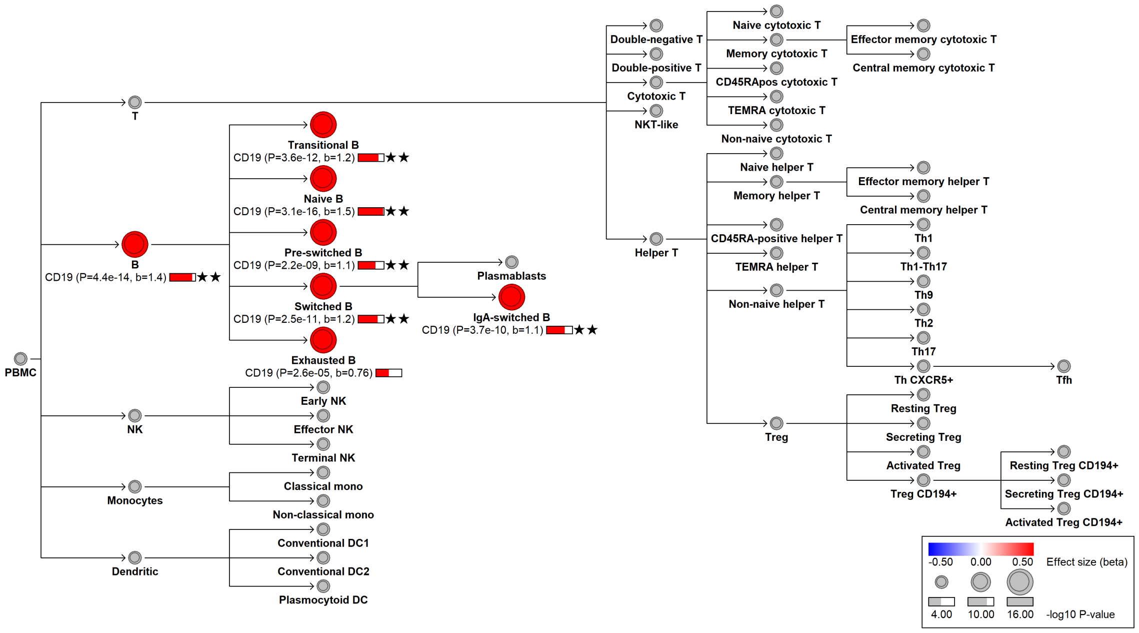

#### chr1:207621975[G→A; EAF 20.456%] (rs2296160, cs_49)

Candidate genes: *CR1*
Diseases associations: Alzheimer's disease
Plasma pQTLs: CR1

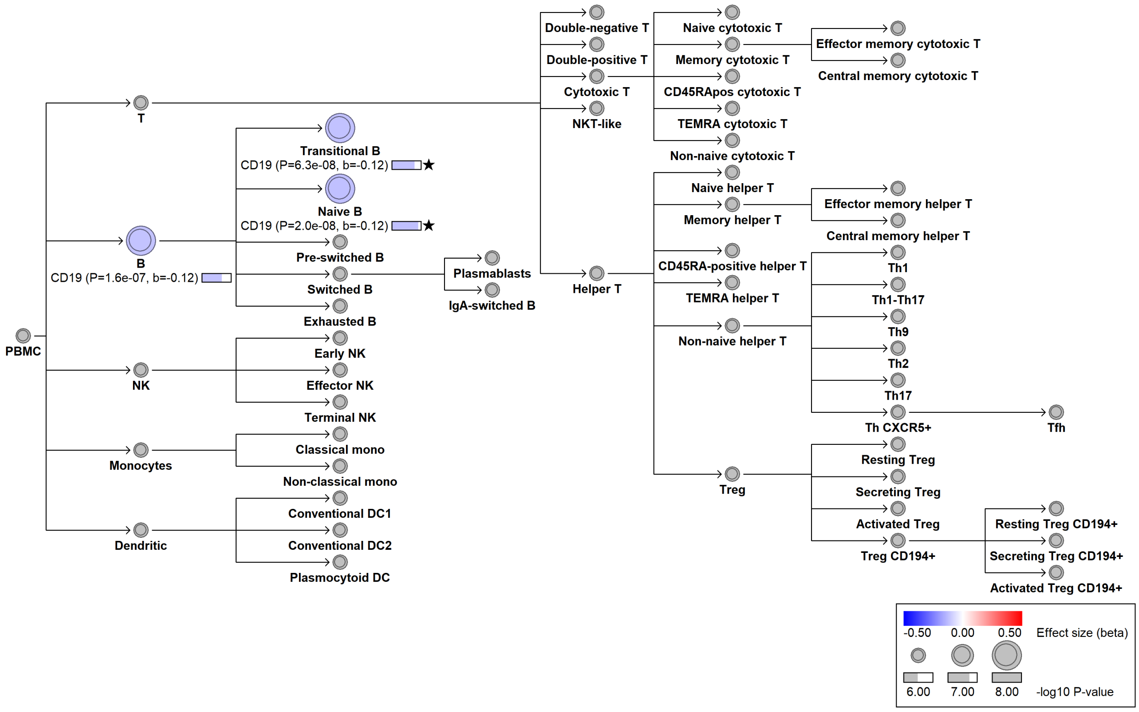

### chr2:38597691[!→AT; EAF 4.091%] (rs1257125526, cs_157)

Candidate genes: *HNRNPLL*
Diseases associations: NA
Plasma pQTLs: NA

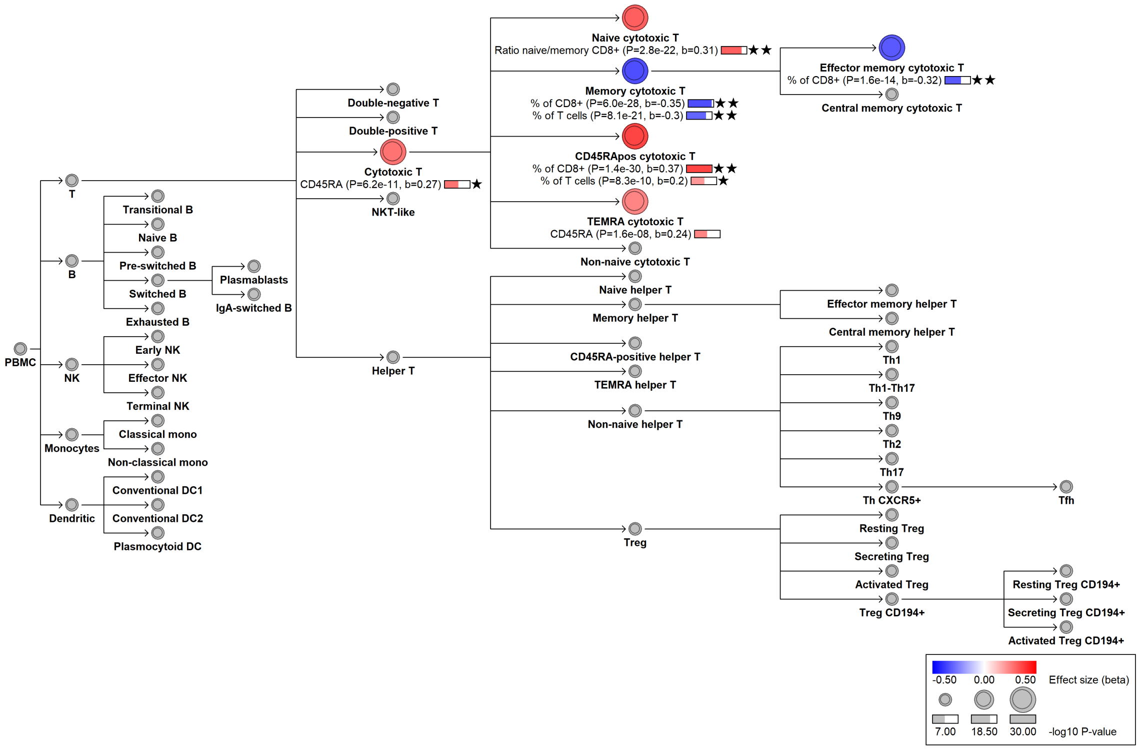

**chr2:38603057[!→GC; EAF 1.852%] (rs542755033, cs_193)**

Candidate genes: *HNRNPLL*
Diseases associations: NA
Plasma pQTLs: NA

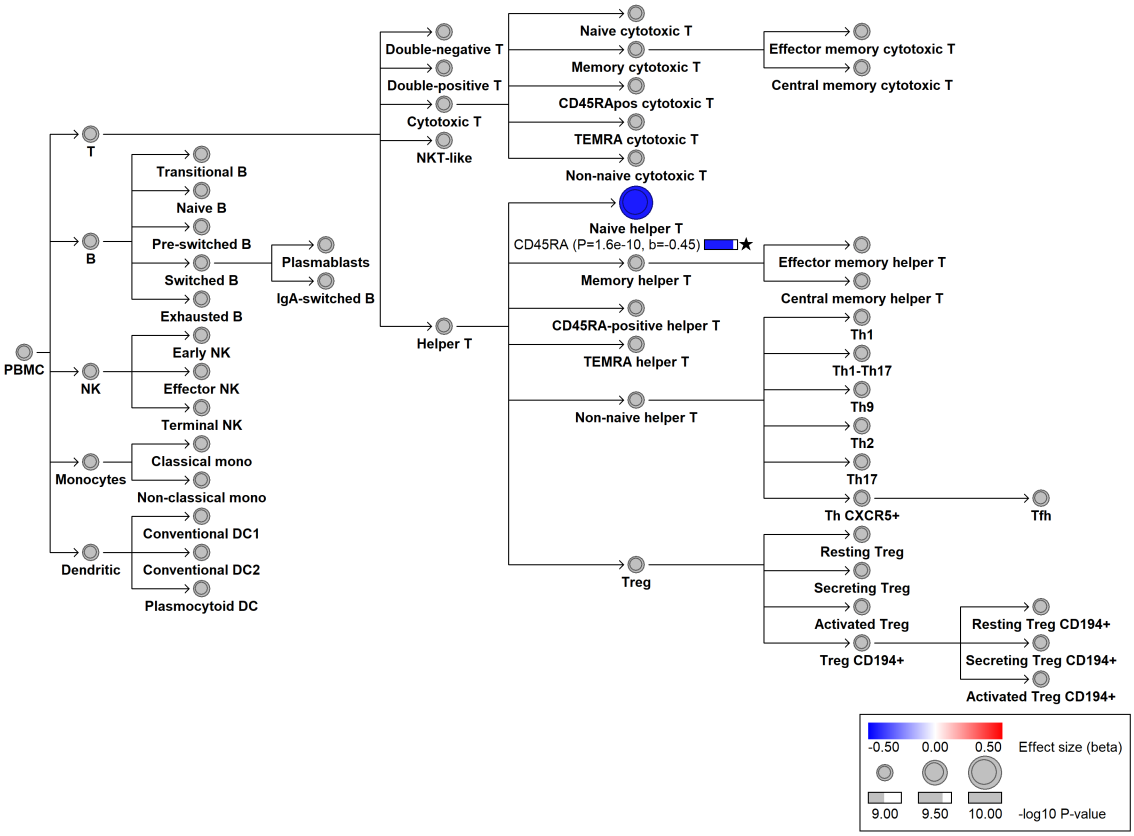

### chr2:38669932[G→!; EAF 10.811%] (chr2:38669932, cs_44)

Candidate genes: *HNRNPLL*
Diseases associations: NA
Plasma pQTLs: NA

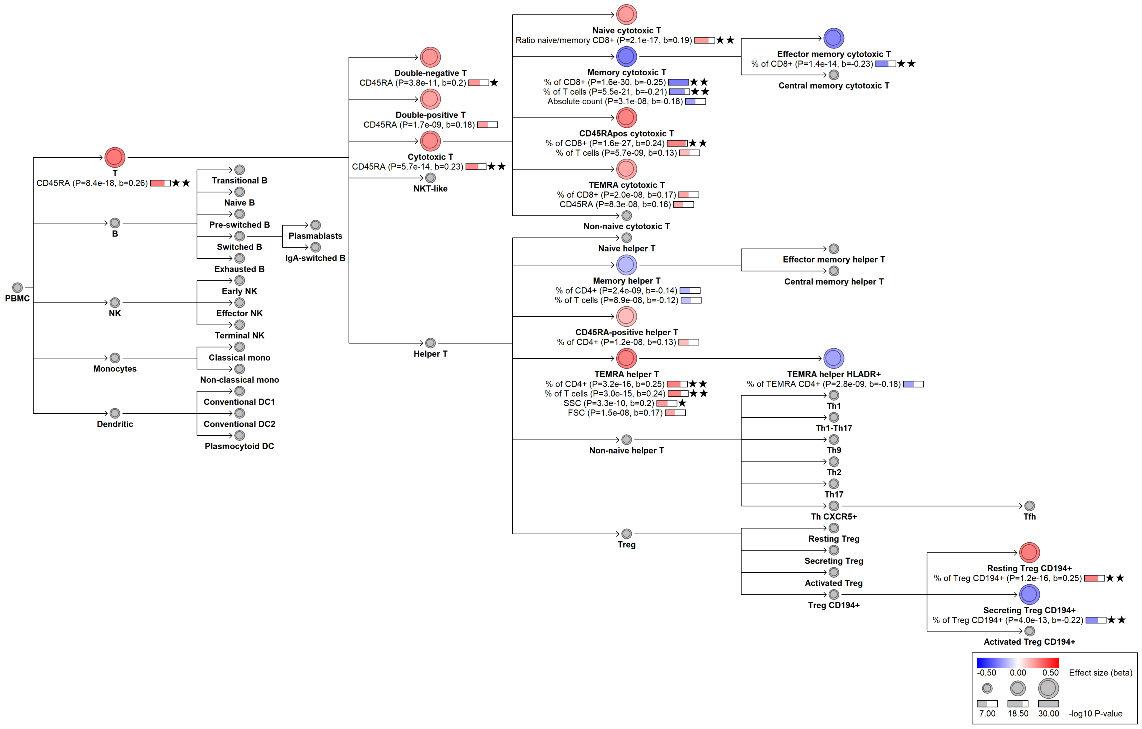

### chr2:38701574[C→G; EAF 49.184%] (rs1821123, cs_190)

Candidate genes: *HNRNPLL*
Diseases associations: NA
Plasma pQTLs: NA

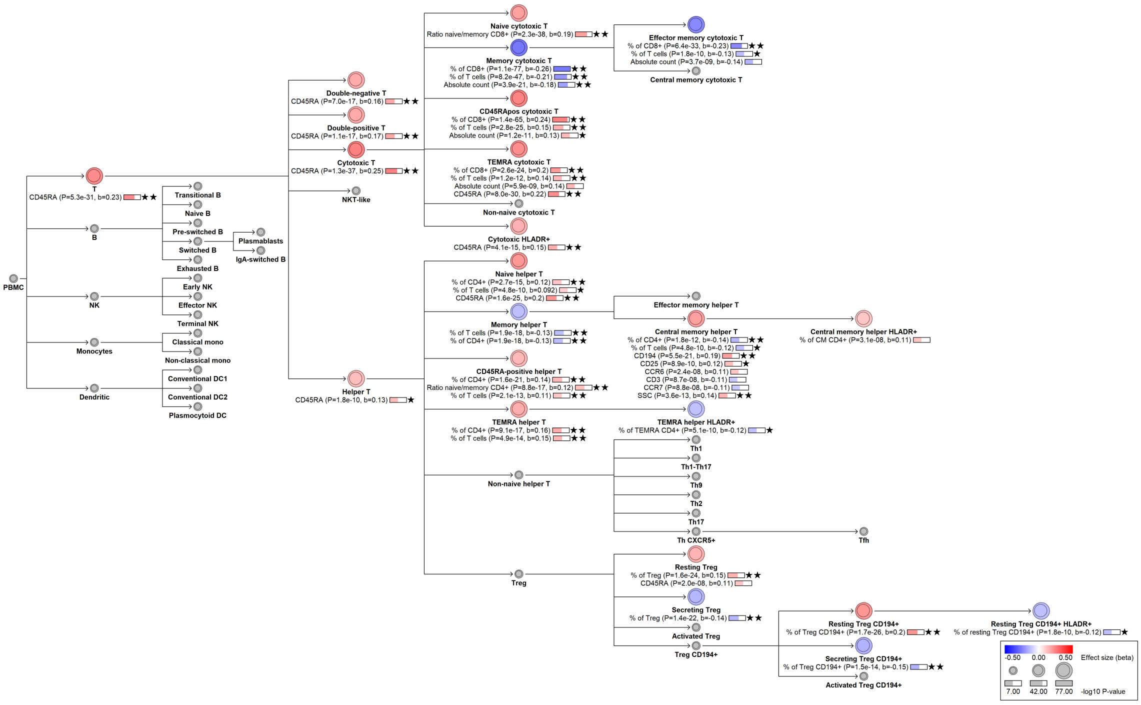

### chr2:38706511[!→CA; EAF 10.45%] (rs1488924669, cs_162)

Candidate genes: *HNRNPLL*
Diseases associations: NA
Plasma pQTLs: NA

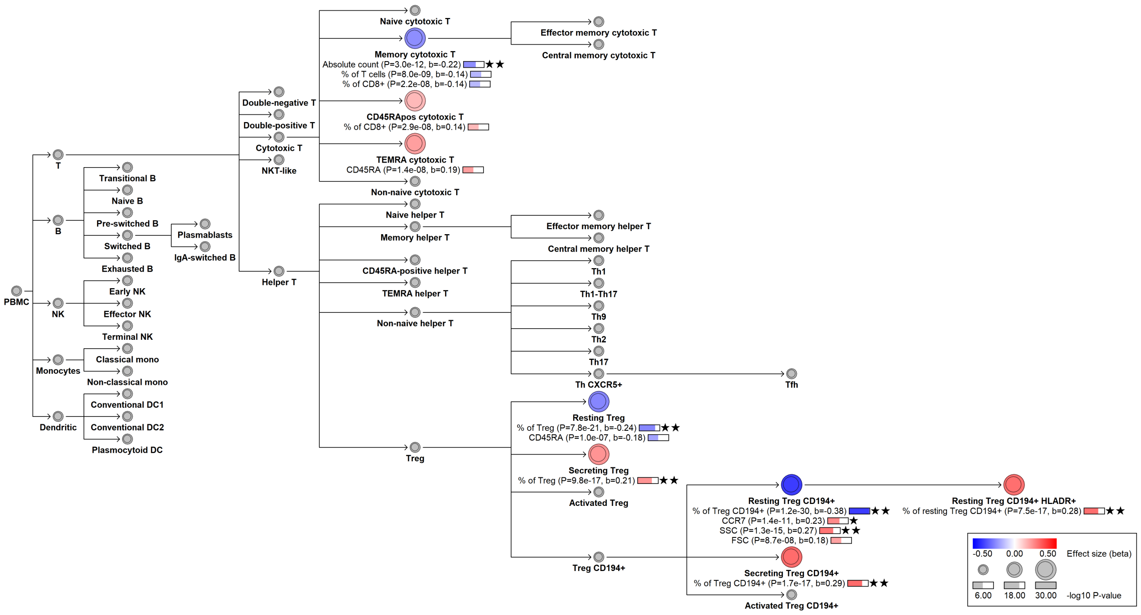

#### chr2:38709640[T→C; EAF 1.864%] (rs147712762, cs_41)

Candidate genes: *HNRNPLL*
Diseases associations: NA
Plasma pQTLs: NA

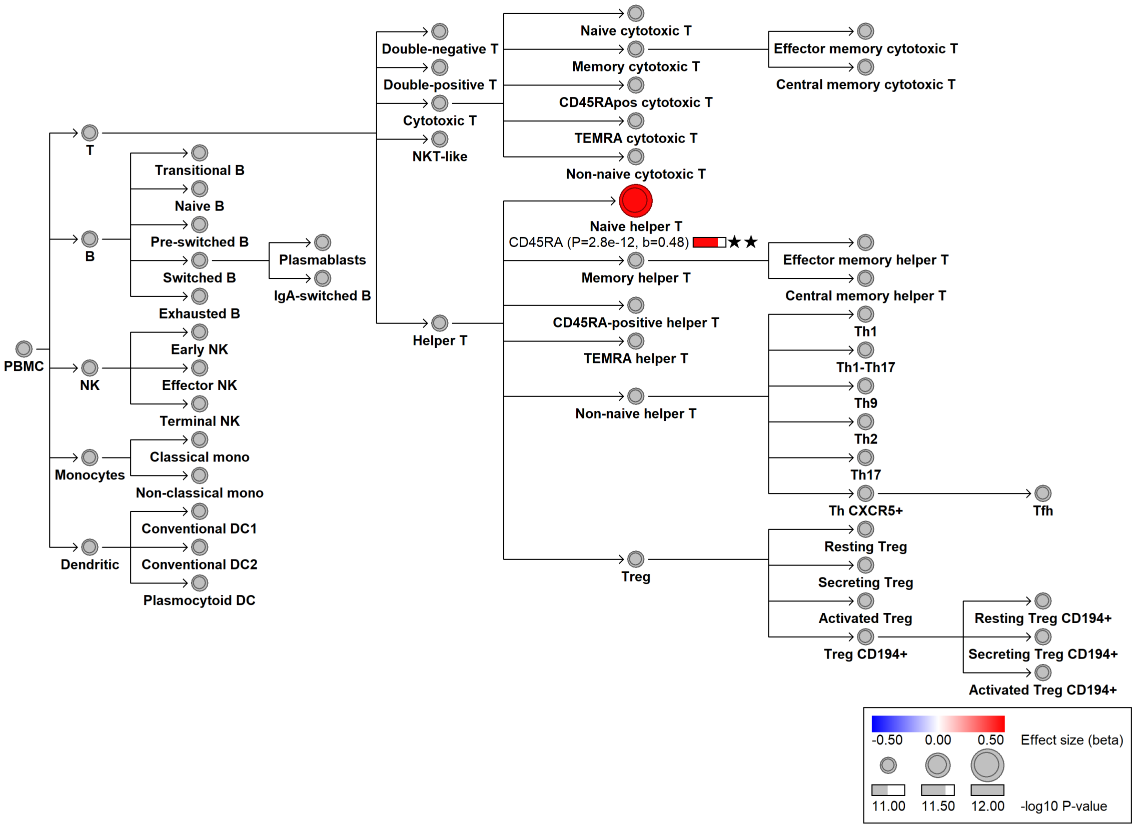

#### chr2:43455518[!→TCACACACACA; EAF 42.285%] (rs35958795, cs_99)

Candidate genes: *THADA*
Diseases associations: NA
Plasma pQTLs: RBP7

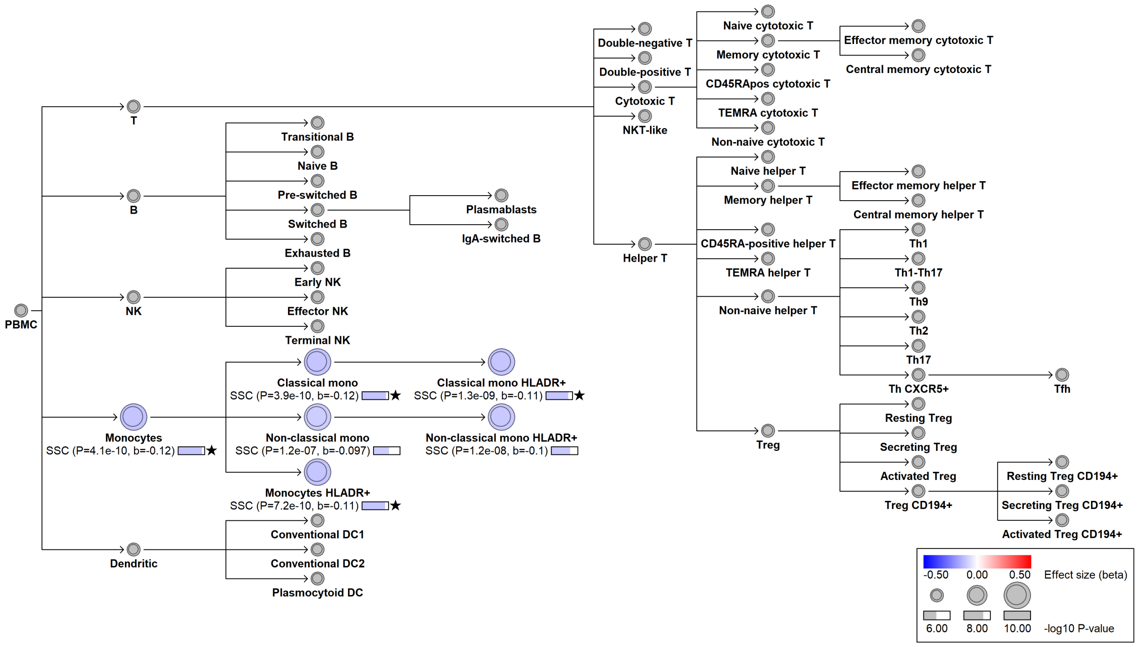

#### chr2:59588718[G→A; EAF 0.398%] (rs140510489, cs_74)

Candidate genes: *BCL11A*
Diseases associations: NA
Plasma pQTLs: NA

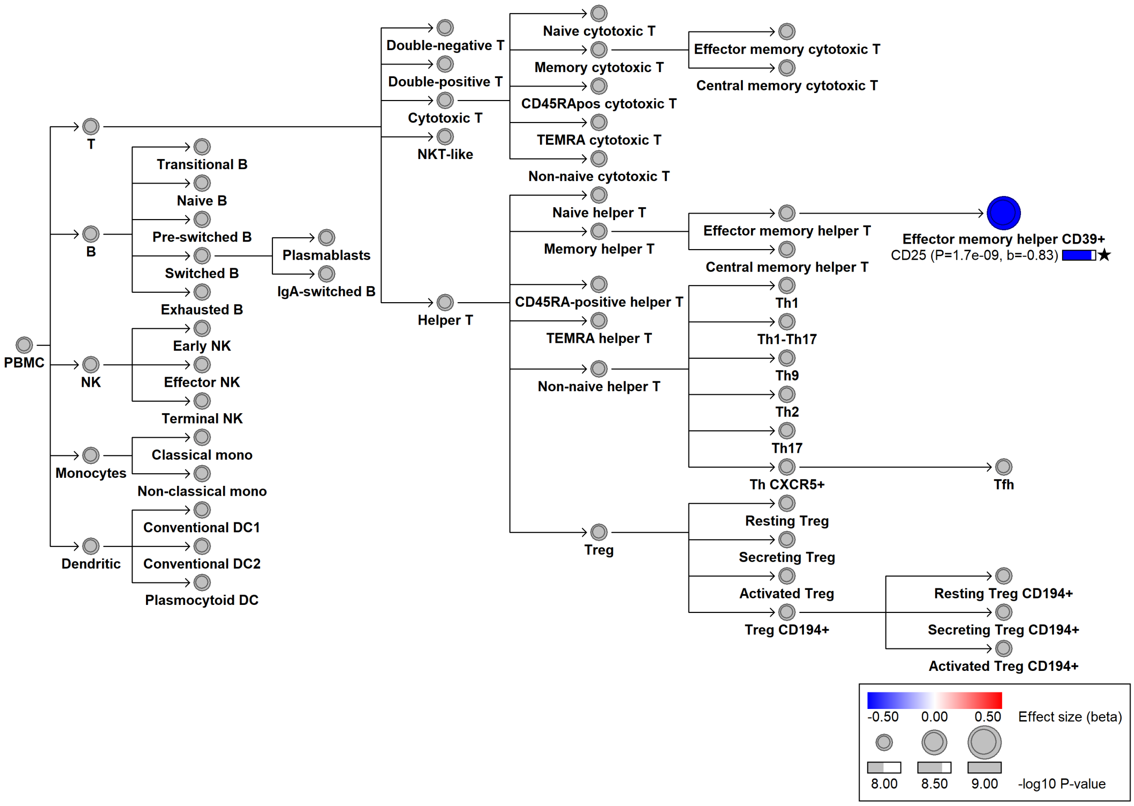

#### chr2:60356235[A→G; EAF 3.196%] (rs76529516, cs_114)

Candidate genes: *BCL11A*
Diseases associations: NA
Plasma pQTLs: CD72, Composite immunoglobulin trait IgA/IgG, Composite immunoglobulin trait IgG/IgM, DRAXIN, FASLG, FCRL1, IGLL1, Immunoglobulin M, NCR1

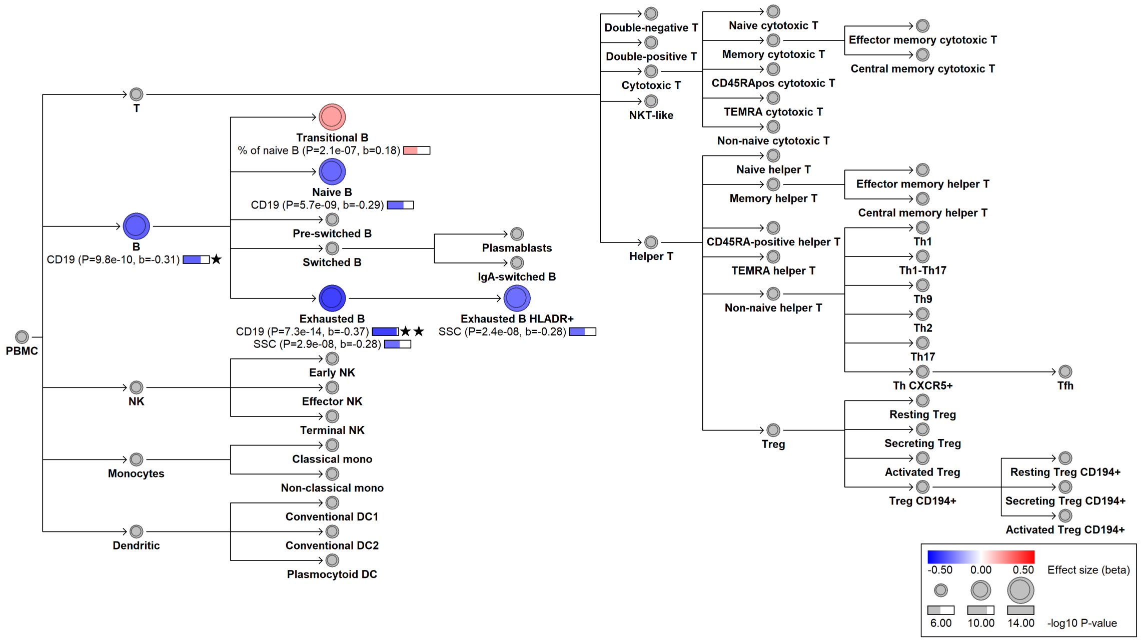

#### chr2:60927375[T→C; EAF 14.109%] (rs72807491, cs_47)

Candidate genes: *BCL11A*
Diseases associations: Celiac disease, Multiple sclerosis, Rheumatoid arthritis, Systemic lupus erythematosus
Plasma pQTLs: FCER2, PCDHA4

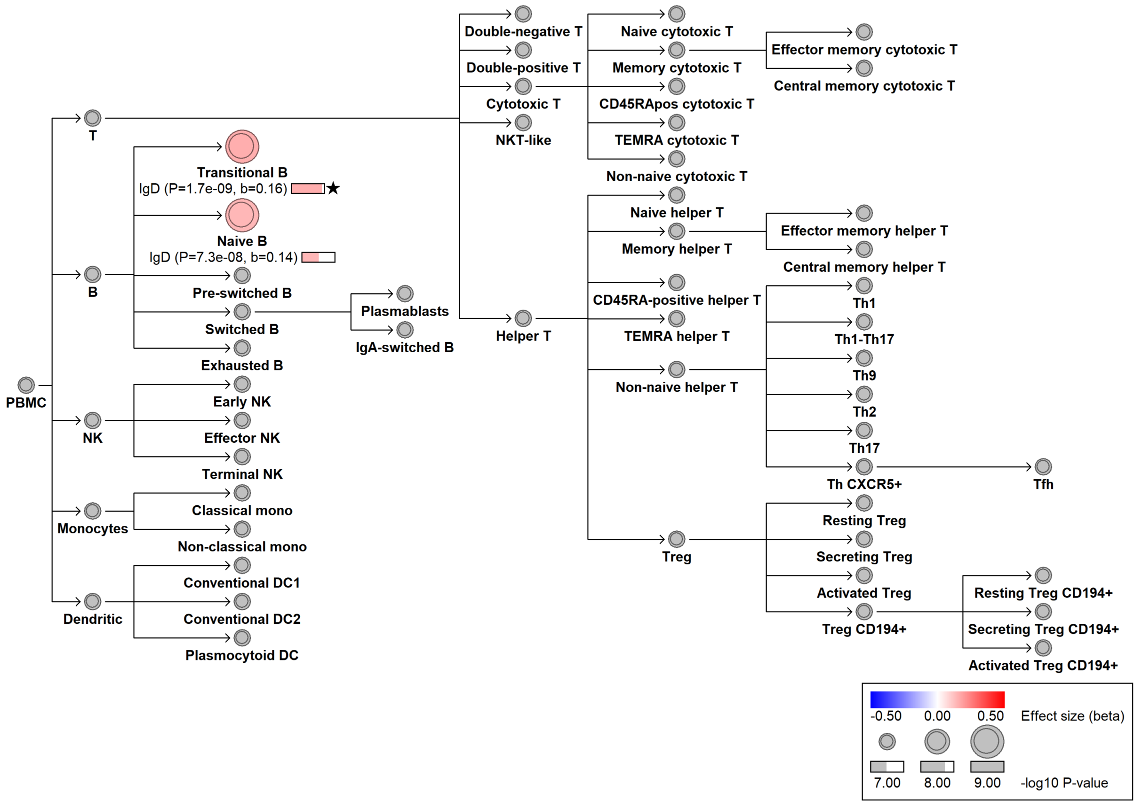

#### chr2:85344587[G→A; EAF 27.328%] (rs4832171, cs_66)

Candidate genes: *RETSAT*
Diseases associations: NA
Plasma pQTLs: NA

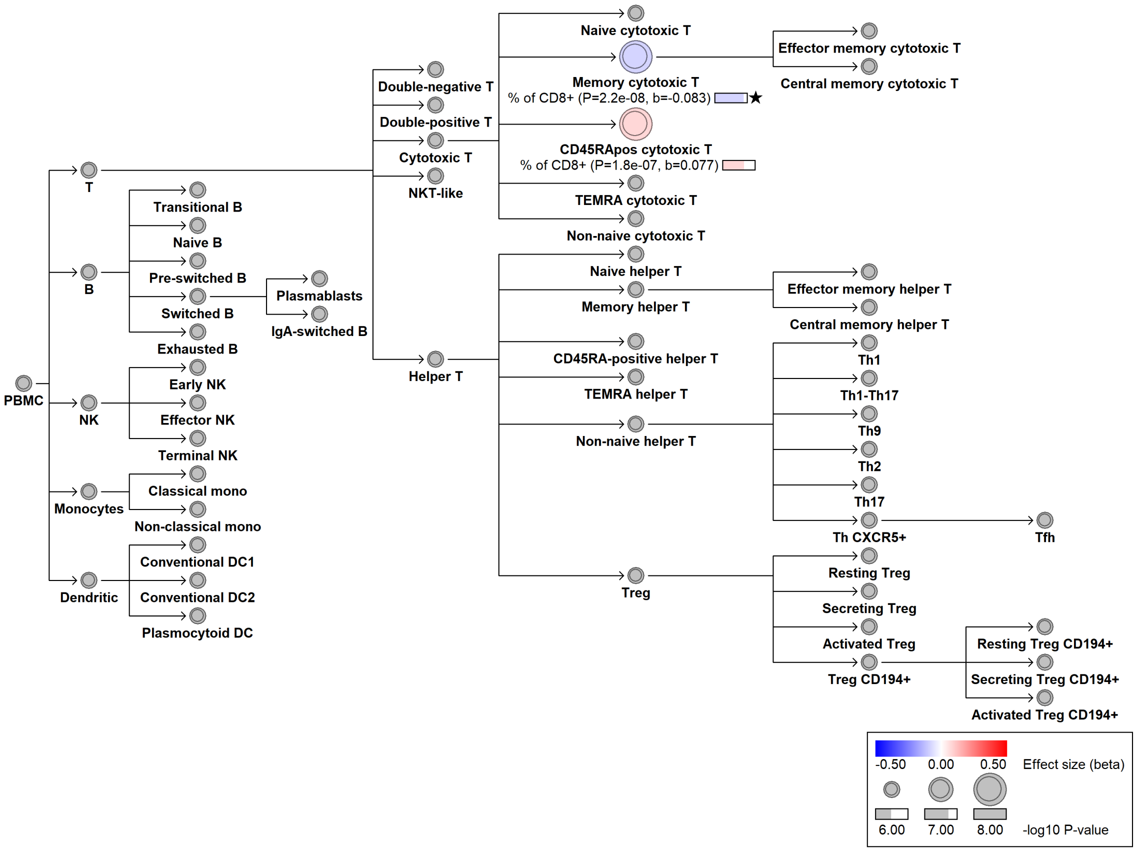

### chr2:86790452[!→T; EAF 9.176%] (rs2229240, cs_165)

Candidate genes: *CD8A*
Diseases associations: NA
Plasma pQTLs: NA

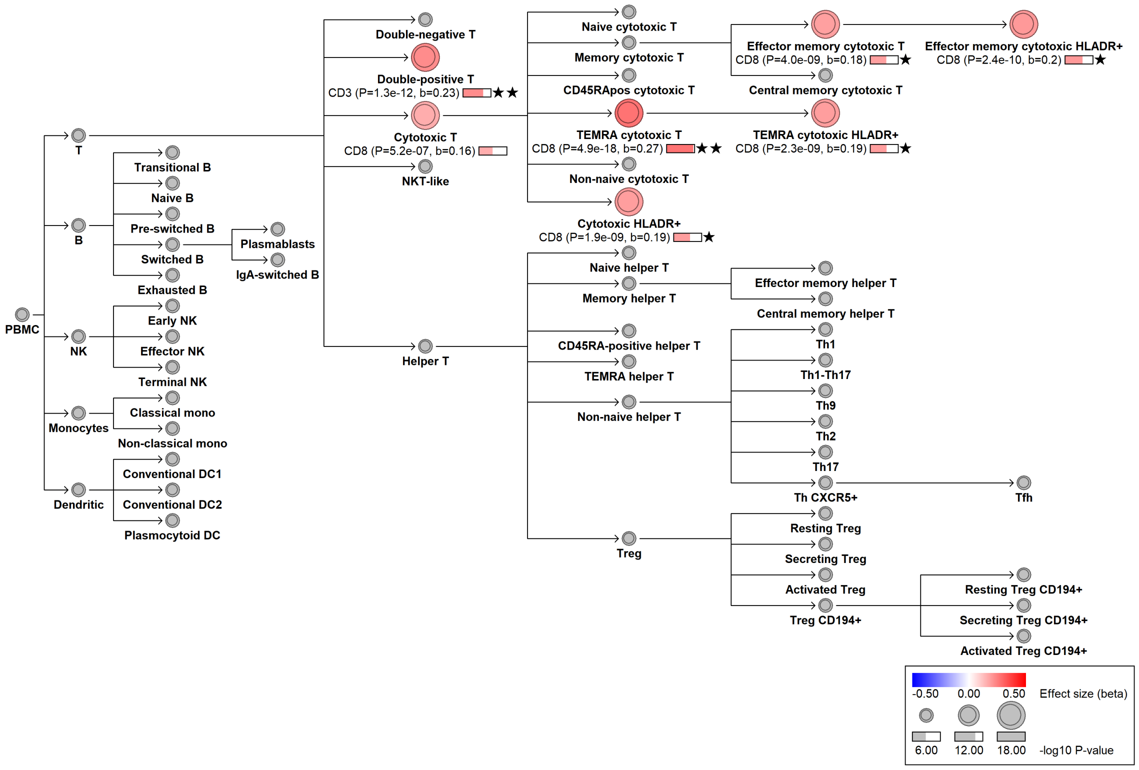

#### chr2:86791329[G→A; EAF 21.383%] (rs6743139, cs_138)

Candidate genes: *CD8A*
Diseases associations: Progressive supranuclear palsy
Plasma pQTLs: NA

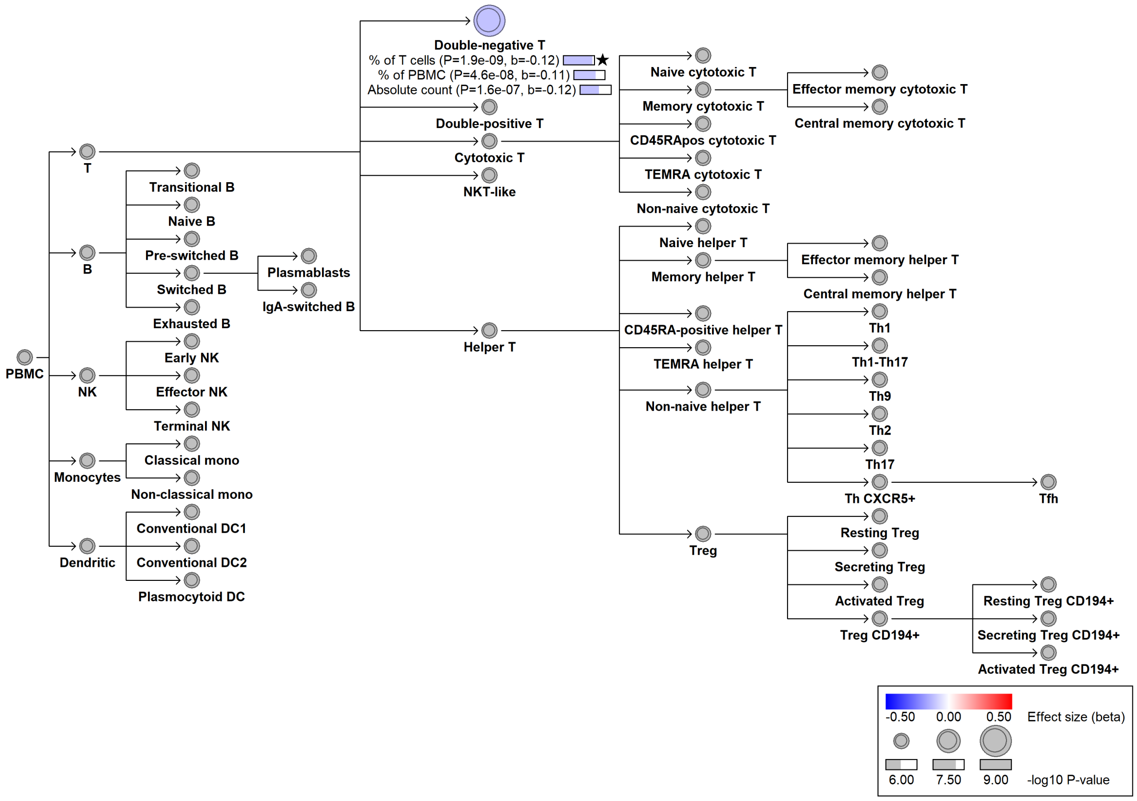

#### chr2:86798528[!→CT; EAF 16.462%] (rs34242639, cs_64)

Candidate genes: *CD8A*
Diseases associations: NA
Plasma pQTLs: NA

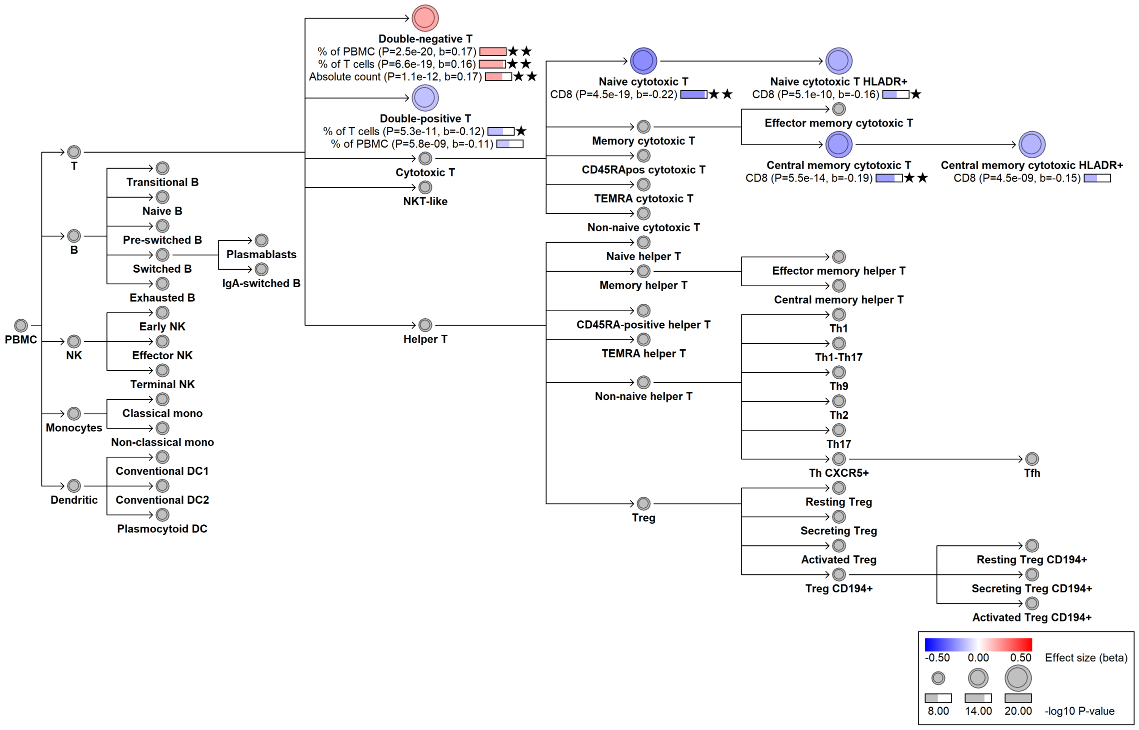

#### chr2:86840231[T→C; EAF 31.155%] (rs4305283, cs_45)

Candidate genes: *CD8B*
Diseases associations: NA
Plasma pQTLs: NA

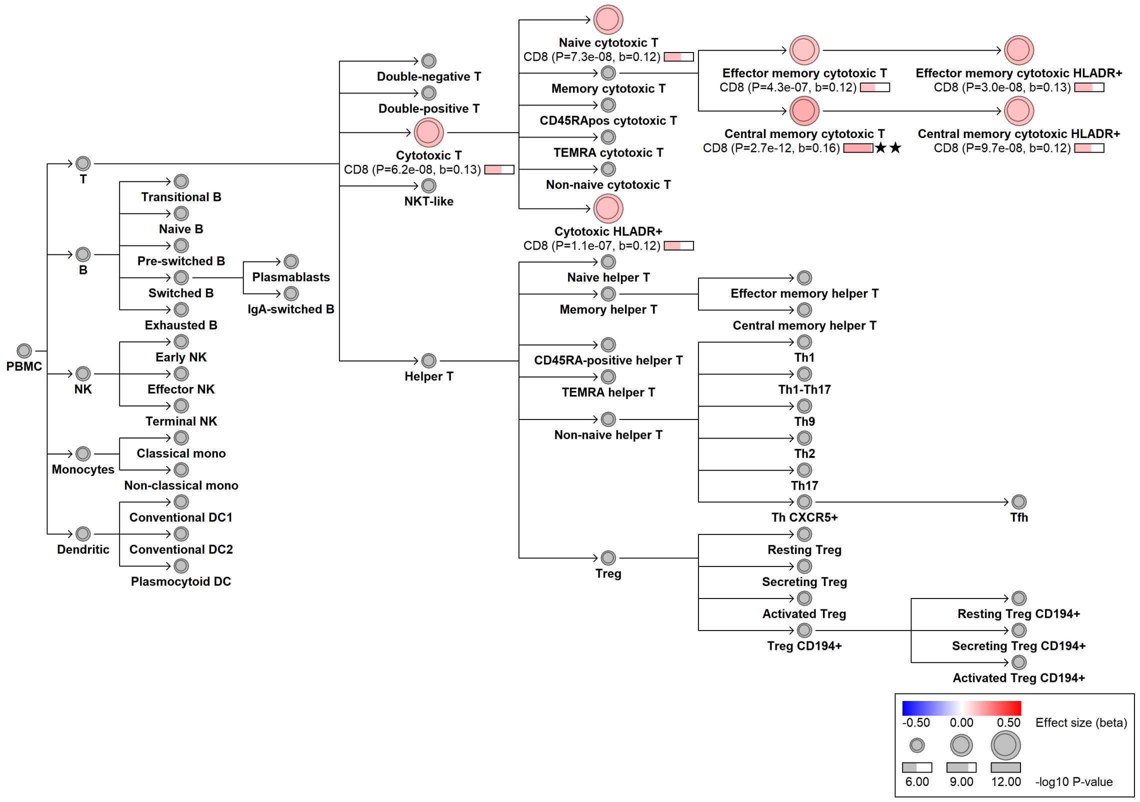

### chr2:86865607[G→A; EAF 0.131%] (rs1468919920, cs_93)

Candidate genes: *CD8B*
Diseases associations: NA
Plasma pQTLs: NA

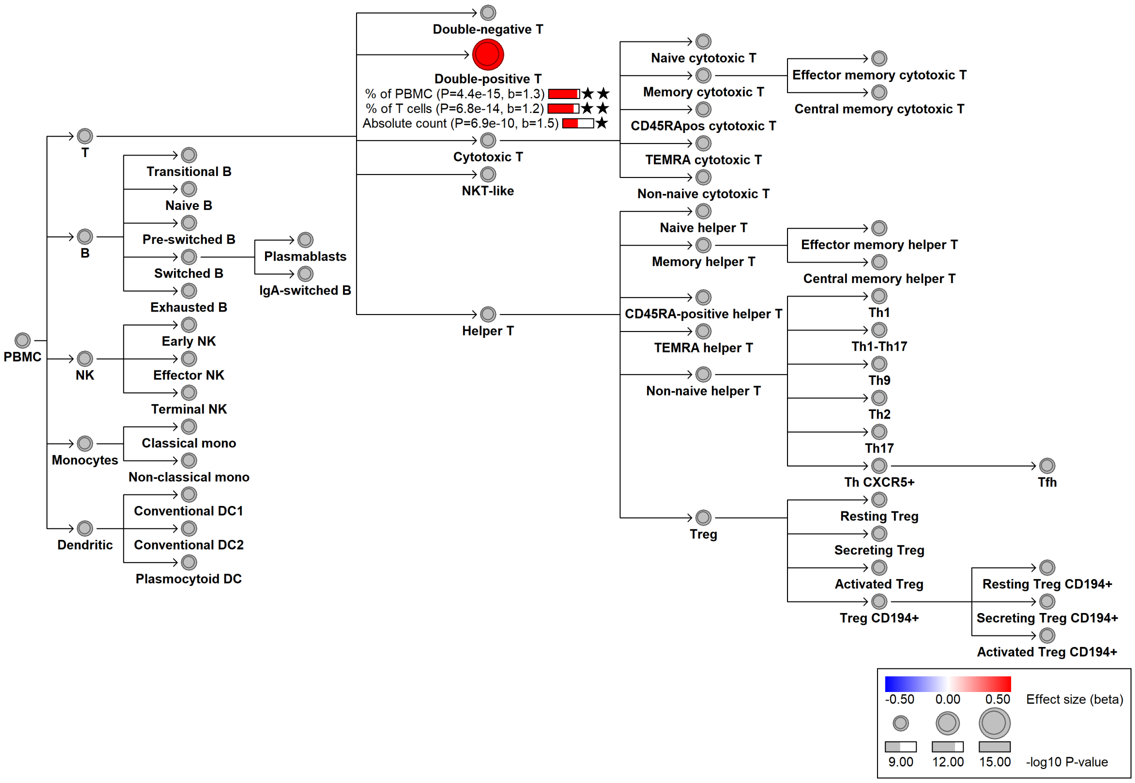

### chr2:87236829[G→!; EAF 0.116%] (chr2:87236829, cs_119)

Candidate genes: *CD8A, CD8B*
Diseases associations: NA
Plasma pQTLs: NA

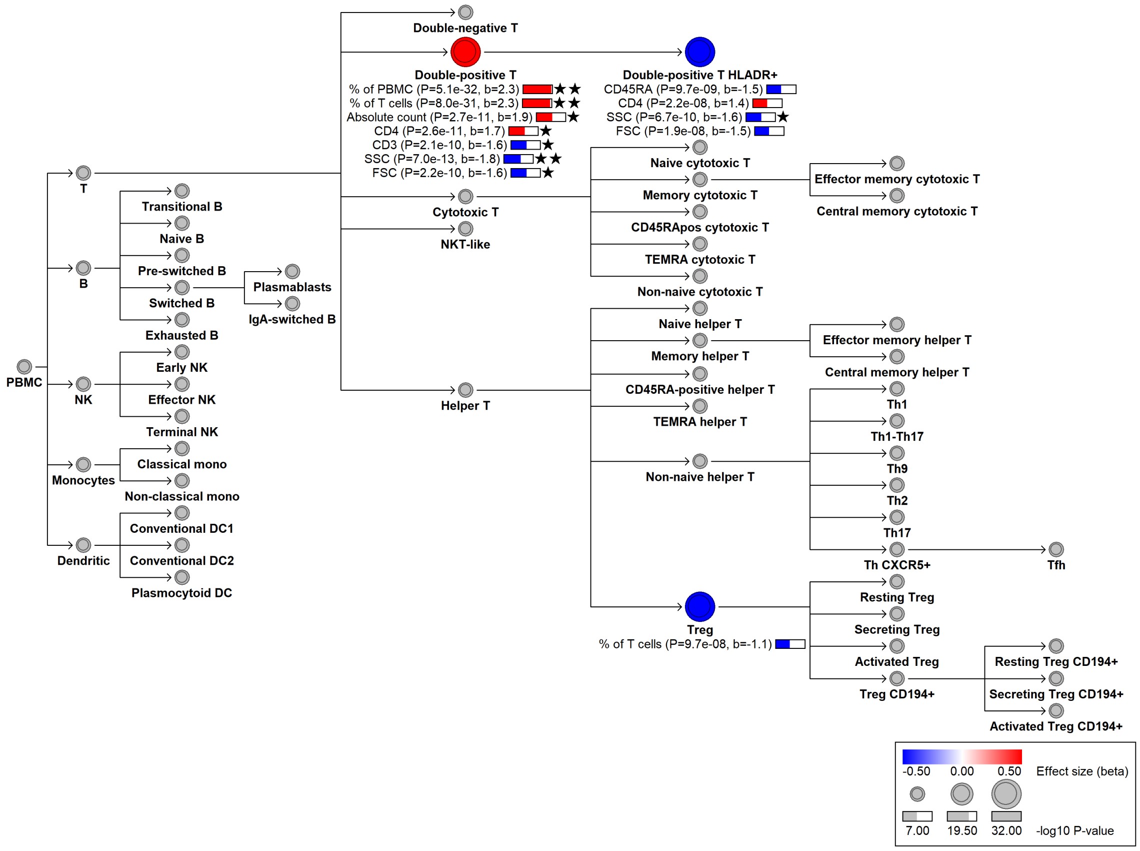

#### chr2:111050100[G→A; EAF 25.07%] (rs2009581, cs_11)

Candidate genes: *ACOXL*
Diseases associations: NA
Plasma pQTLs: ADAMTS13, ADGRE5, CD160, CD22, CD244, CD72, CSF2, CXCL16, ESM1, FCRL1, FCRL3, GCNT1, IL15, IL2RB, KLRB1, KLRD1, KLRF1, KLRK1, LTA, LY9, NCR1, SEMA7A, SLAMF8, TNC, XCL1

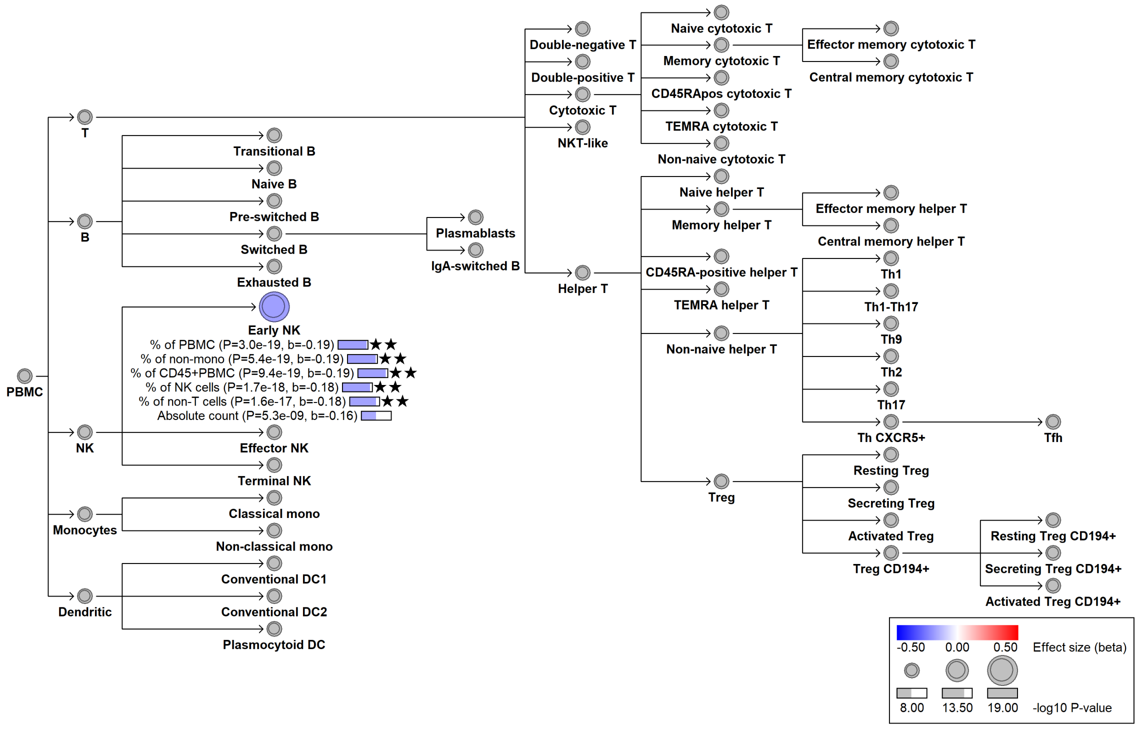

#### chr2:132998726[G→A; EAF 0.134%] (rs182362285, cs_112)

Candidate genes: *NCKAP5*
Diseases associations: NA
Plasma pQTLs: NA

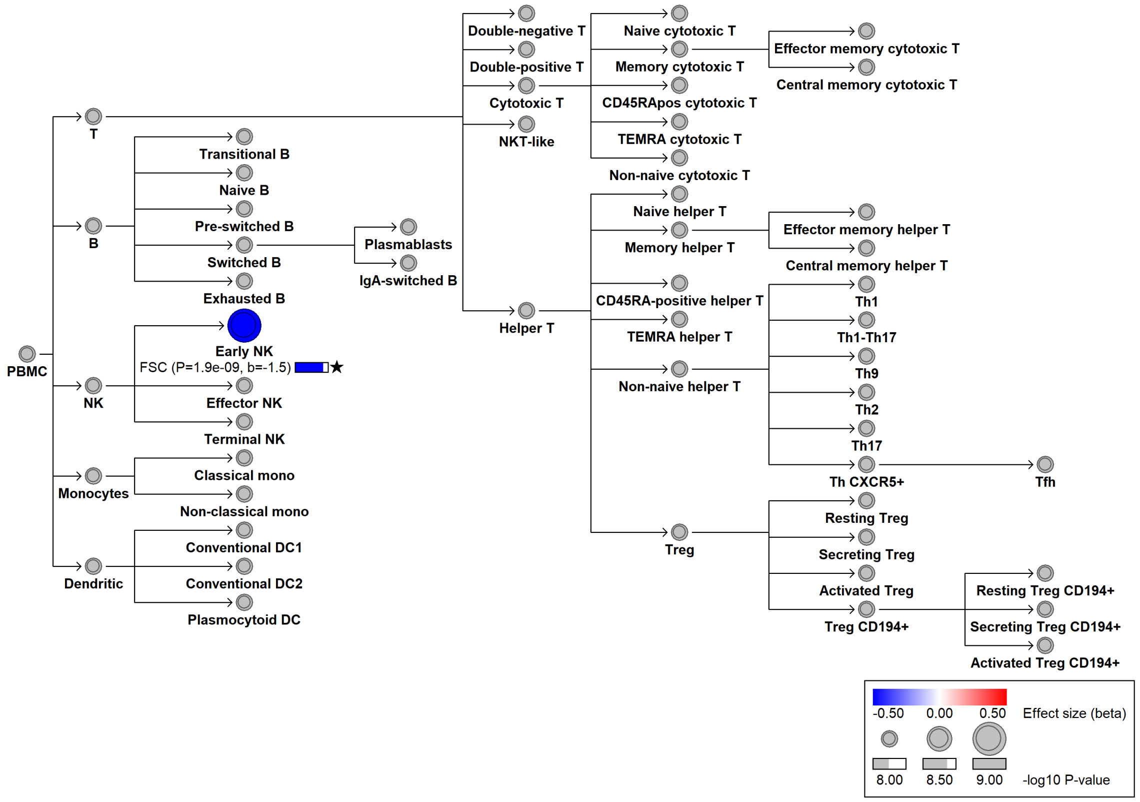

### chr2:136187459[C→T; EAF 27.197%] (rs1123848, cs_172)

Candidate genes: *CXCR4*
Diseases associations: NA
Plasma pQTLs: BLNK, TCL1A

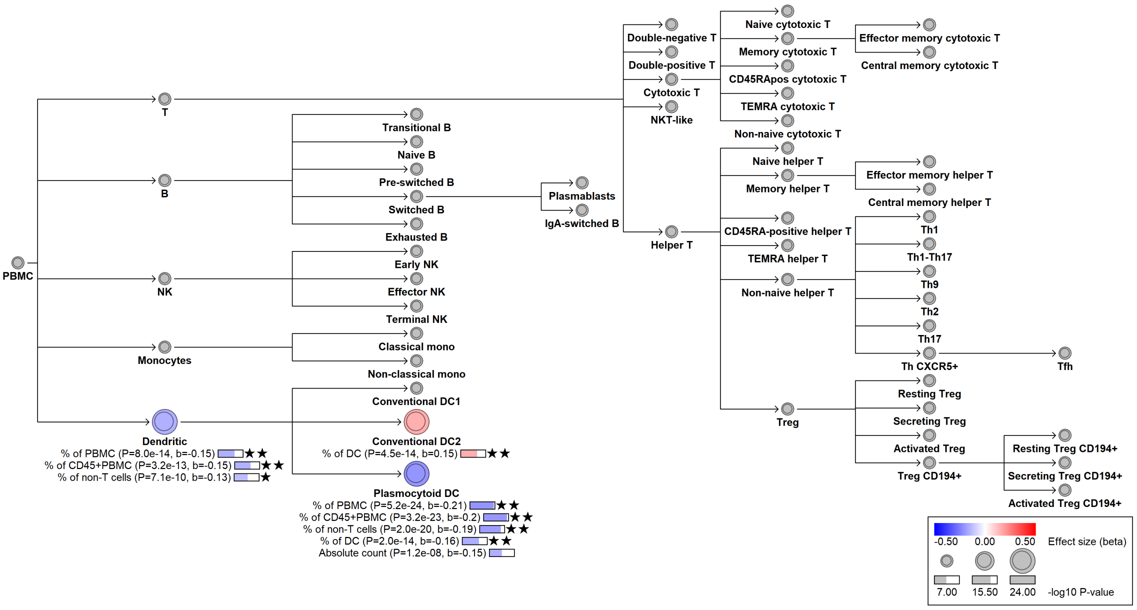

#### chr2:144688891[C→T; EAF 8.262%] (rs66537831, cs_232)

Candidate genes: *ZEB2*
Diseases associations: NA
Plasma pQTLs: CD1C, CD83, CLEC4C, IL12B, SIGLEC6

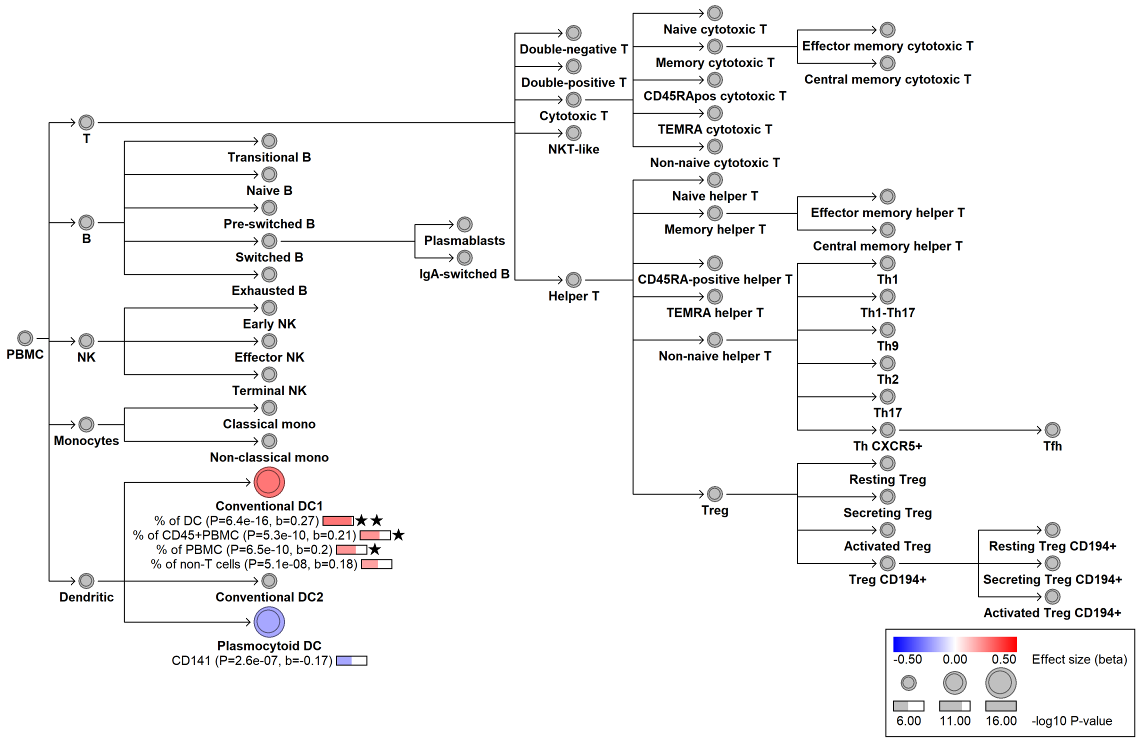

### chr2:160457953[!→TTTG; EAF 20.53%] (rs139604678, cs_233)

Candidate genes: *RBMS1*
Diseases associations: NA
Plasma pQTLs: NA

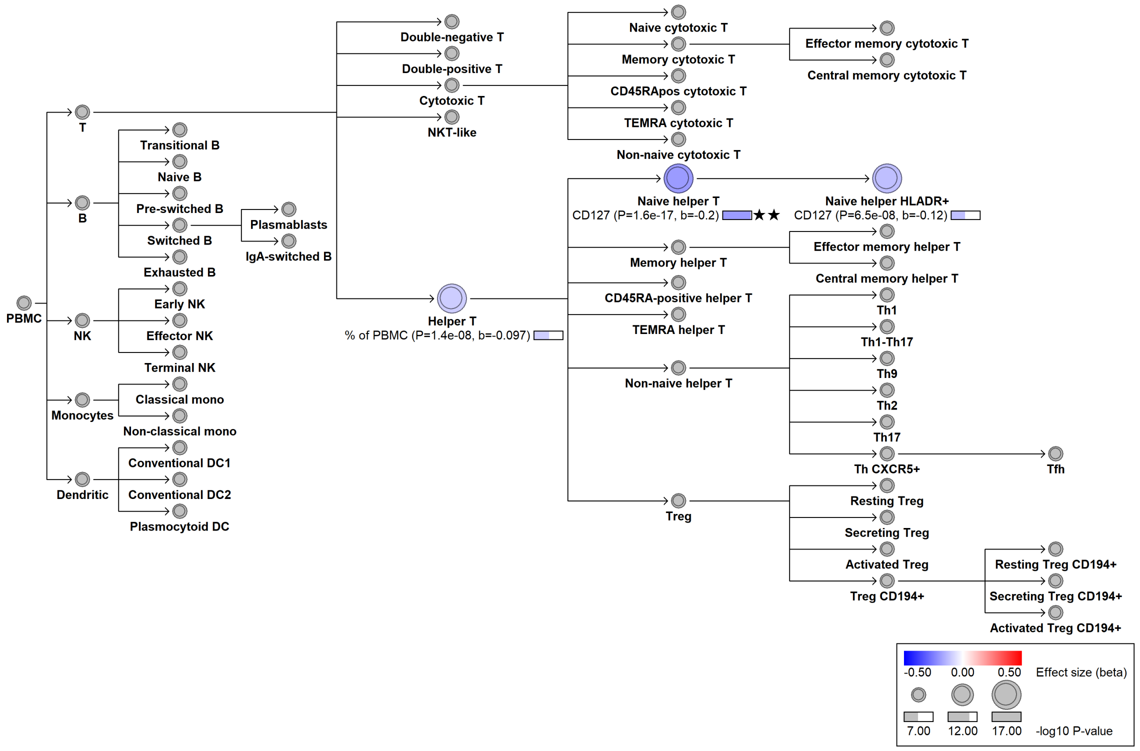

### chr2:181459039[A→G; EAF 43.412%] (rs1375493, cs_92)

Candidate genes: *ITGA4, CERKL*
Diseases associations: Crohn’s disease, Inflammatory bowel disease, Ulcerative colitis
Plasma pQTLs: ITGA5, LILRA5, VCAM1

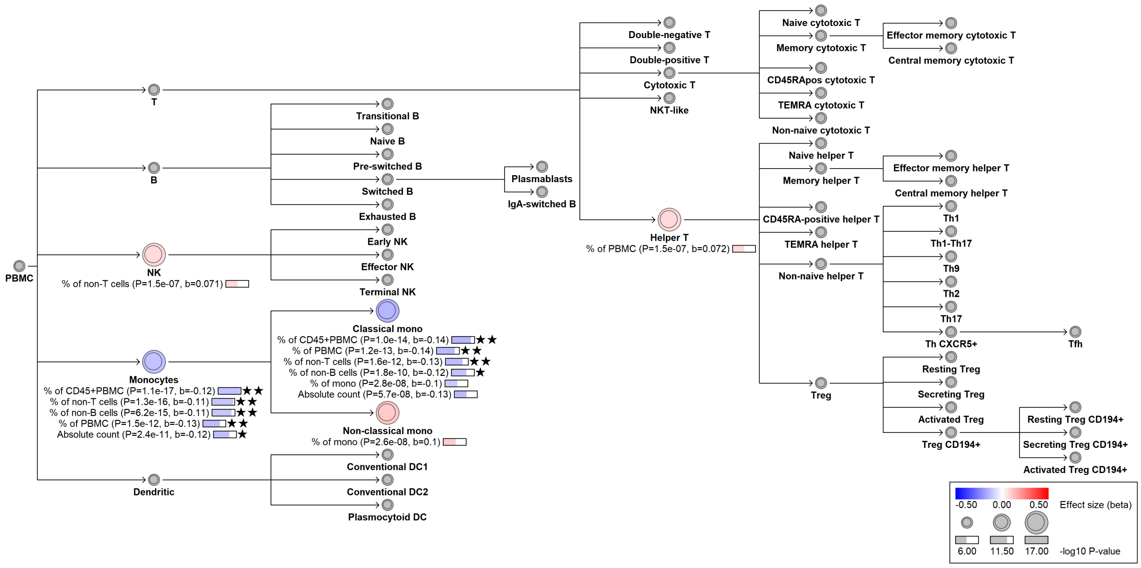

#### chr2:202613145[C→!; EAF 29.593%] (chr2:202613145, cs_126)

Candidate genes: *FAM117B, ICA1L*
Diseases associations: NA
Plasma pQTLs: CCL18, CD163, CD1C, CECR1, HMOX1, MMP19, PCSK2, TIMD4

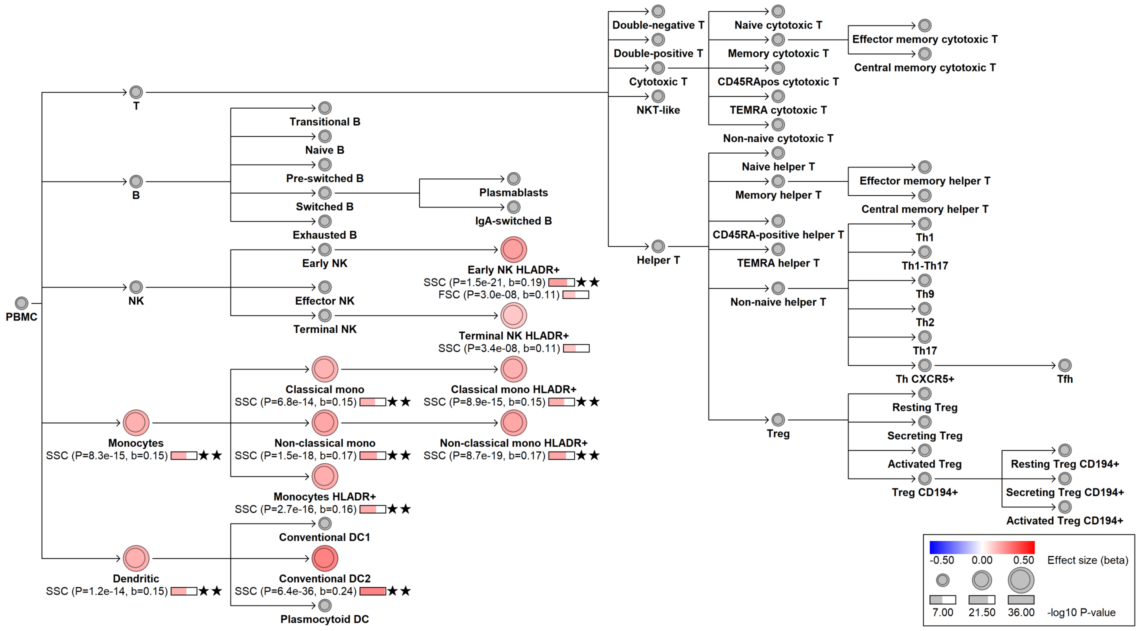

#### chr2:231426182[G→A; EAF 26.163%] (rs16825937, cs_188)

Candidate genes: *B3GNT7*
Diseases associations: NA
Plasma pQTLs: NA

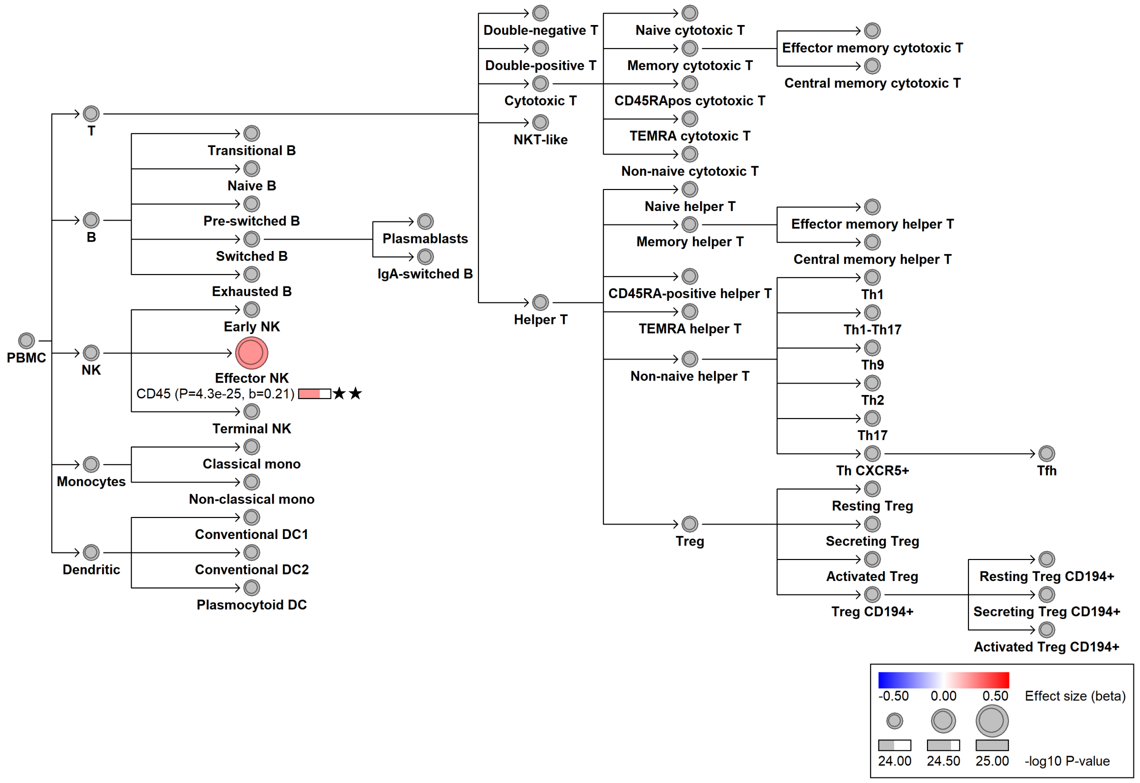

#### chr3:28492167[G→A; EAF 13.922%] (rs13100556, cs_265)

Candidate genes: *ZCWPW2*
Diseases associations: NA
Plasma pQTLs: NA

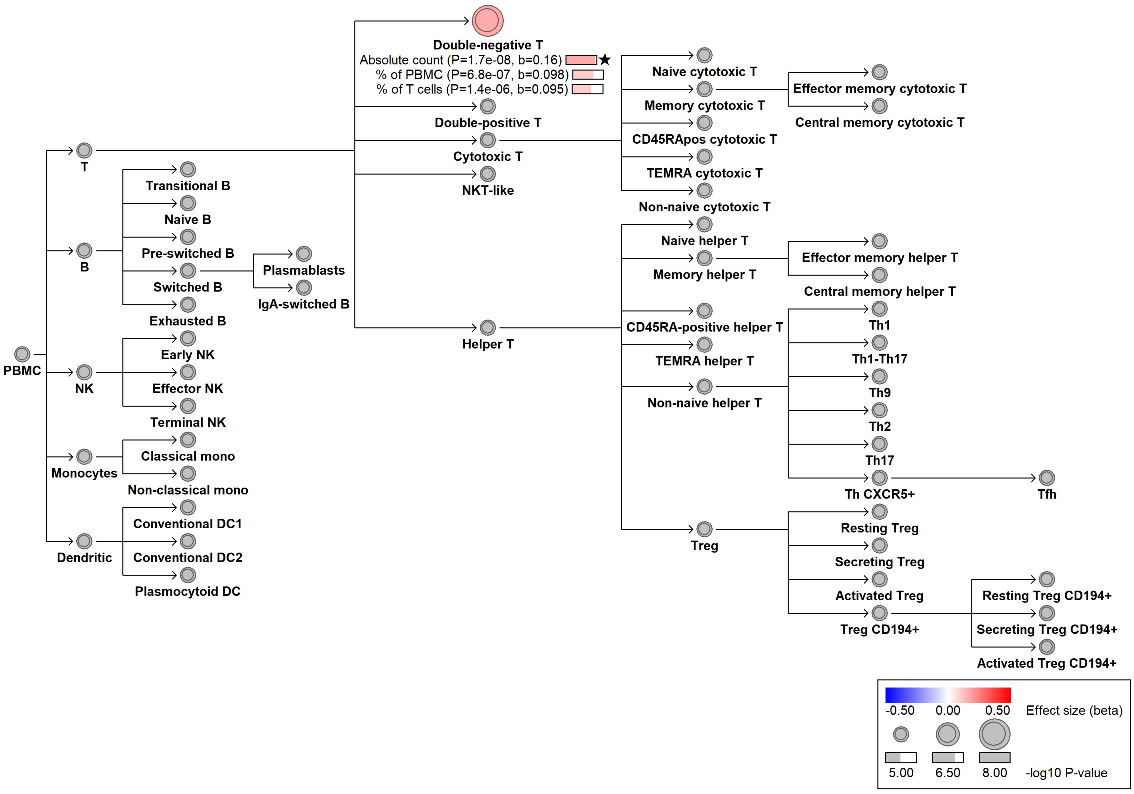

#### chr3:32491754[T→C; EAF 2.057%] (rs35574803, cs_225)

Candidate genes: *CMTM6*
Diseases associations: NA
Plasma pQTLs: NA

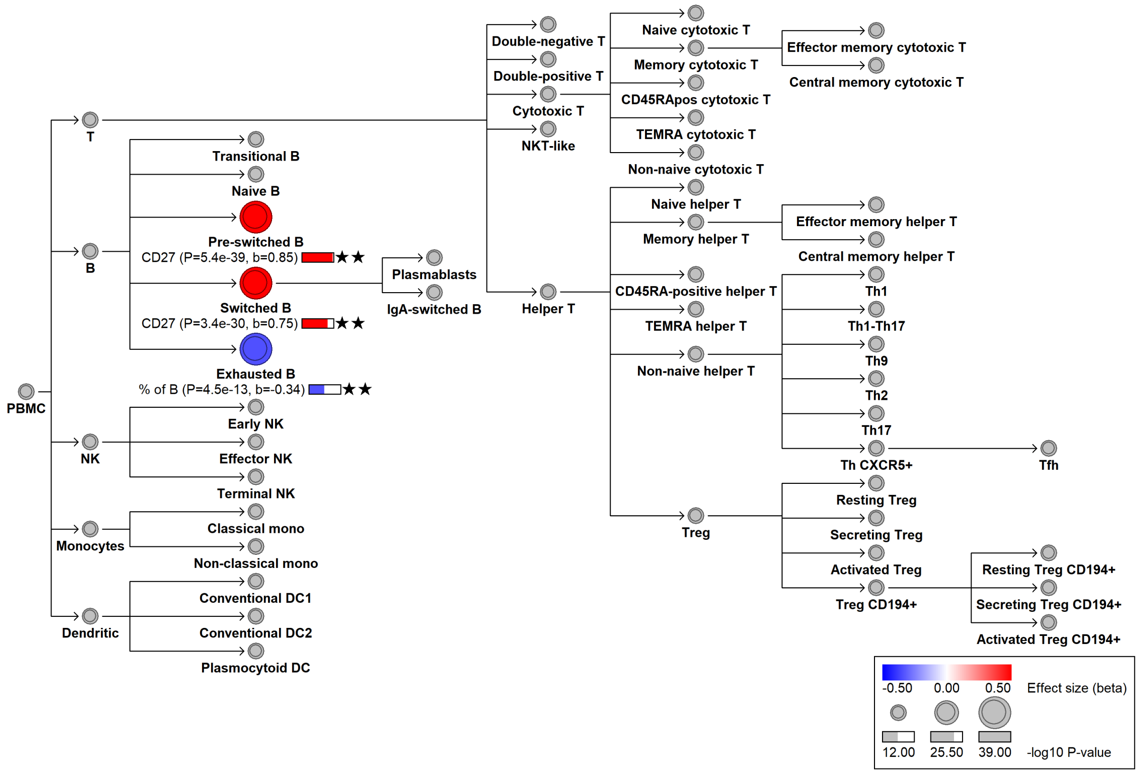

#### chr3:33005791[G→A; EAF 37.866%] (rs34064757, cs_220)

Candidate genes: *CCR4*
Diseases associations: Allergy, Eczema
Plasma pQTLs: NA

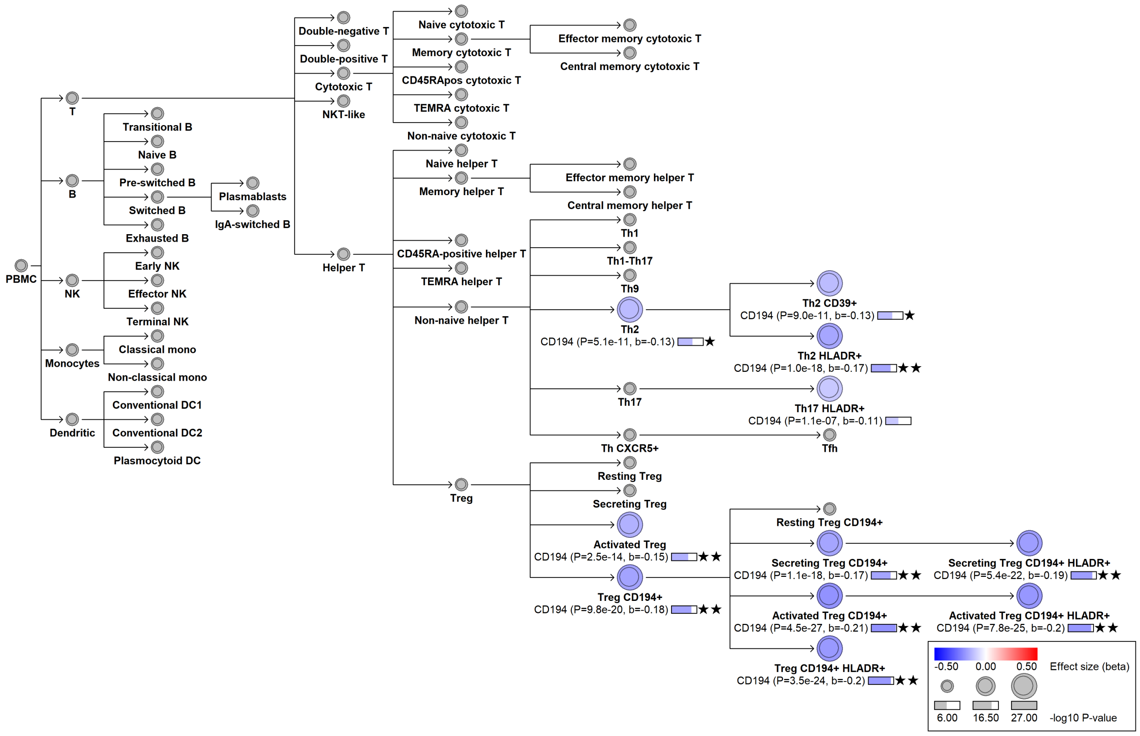

### chr3:33030977[T→C; EAF 4.053%] (rs116135711, cs_146)

Candidate genes: *CCR4*
Diseases associations: NA
Plasma pQTLs: CCL17

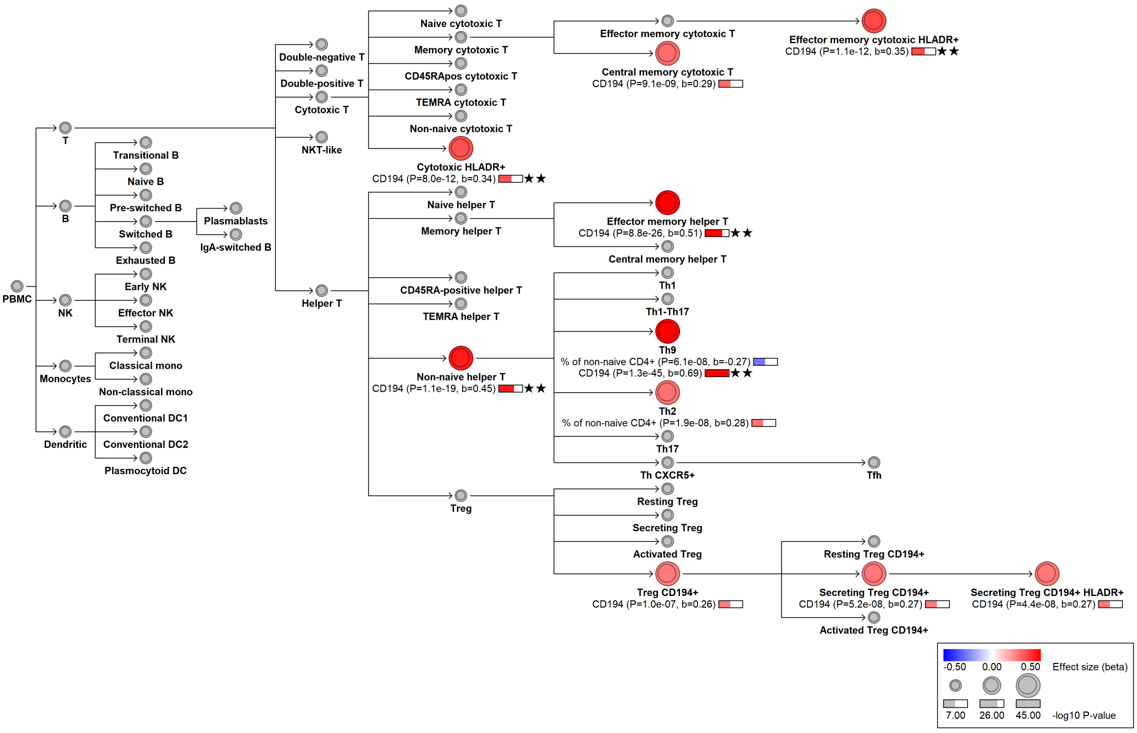

### chr3:33039964[T→C; EAF 10.456%] (rs77281369, cs_259)

Candidate genes: *CCR4*
Diseases associations: NA
Plasma pQTLs: NA

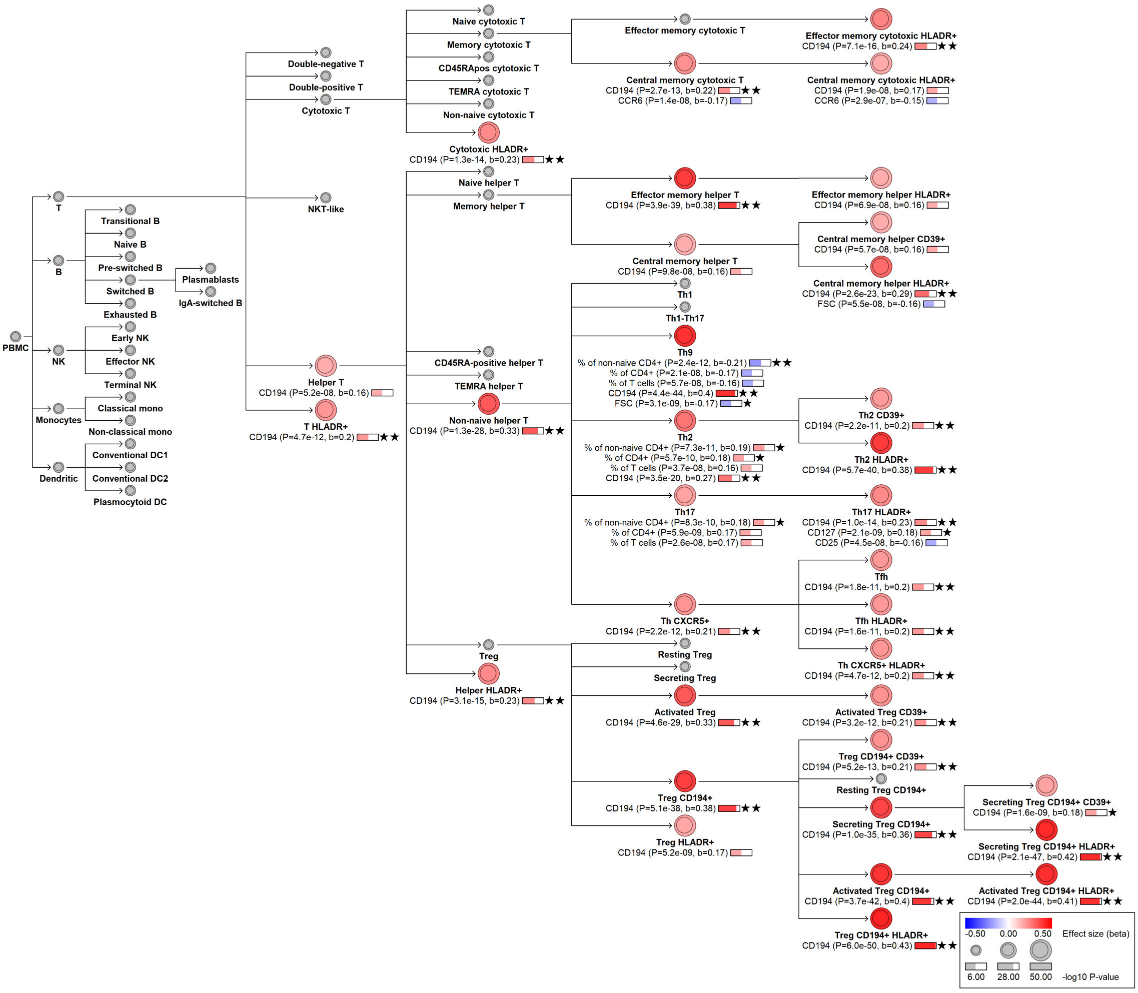

### chr3:33040441[C→!; EAF 18.856%] (chr3:33040441, cs_118)

Candidate genes: *CCR4*
Diseases associations: NA
Plasma pQTLs: NA

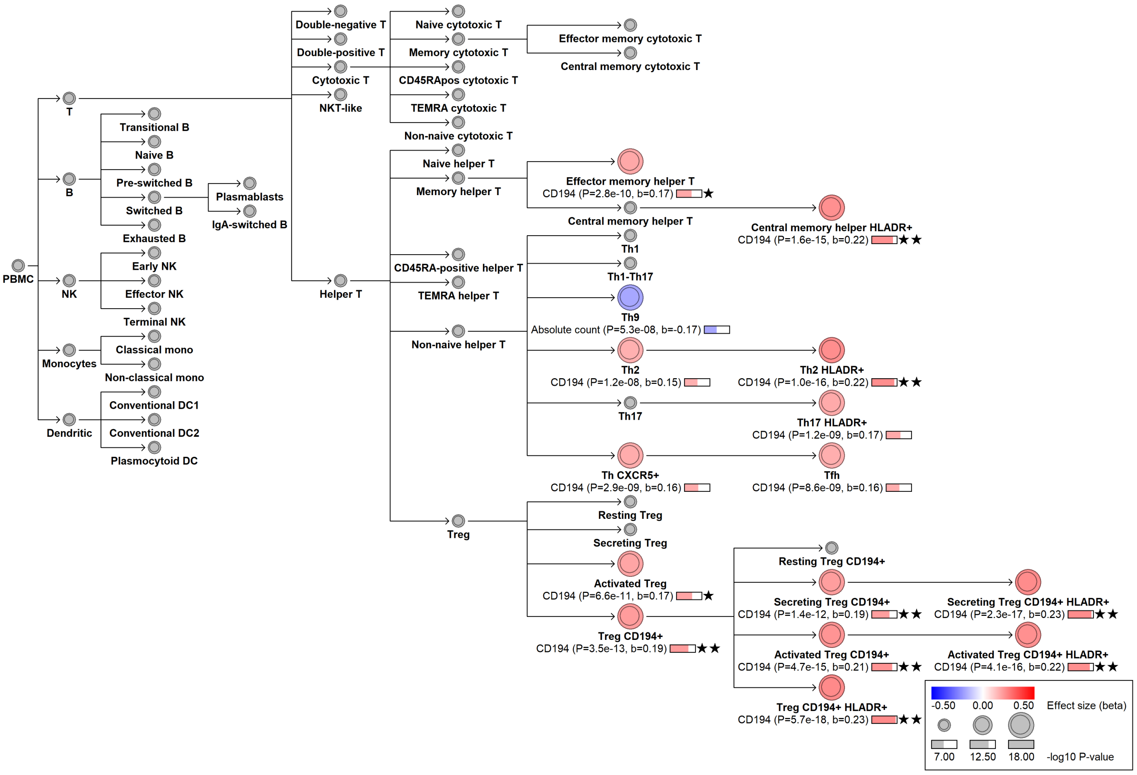

#### chr3:46208452[G→GTTC; EAF 34.809%] (rs147398495, cs_199)

Candidate genes: *CCR1,CCR3, CCRL2*
Diseases associations: NA
Plasma pQTLs: NA

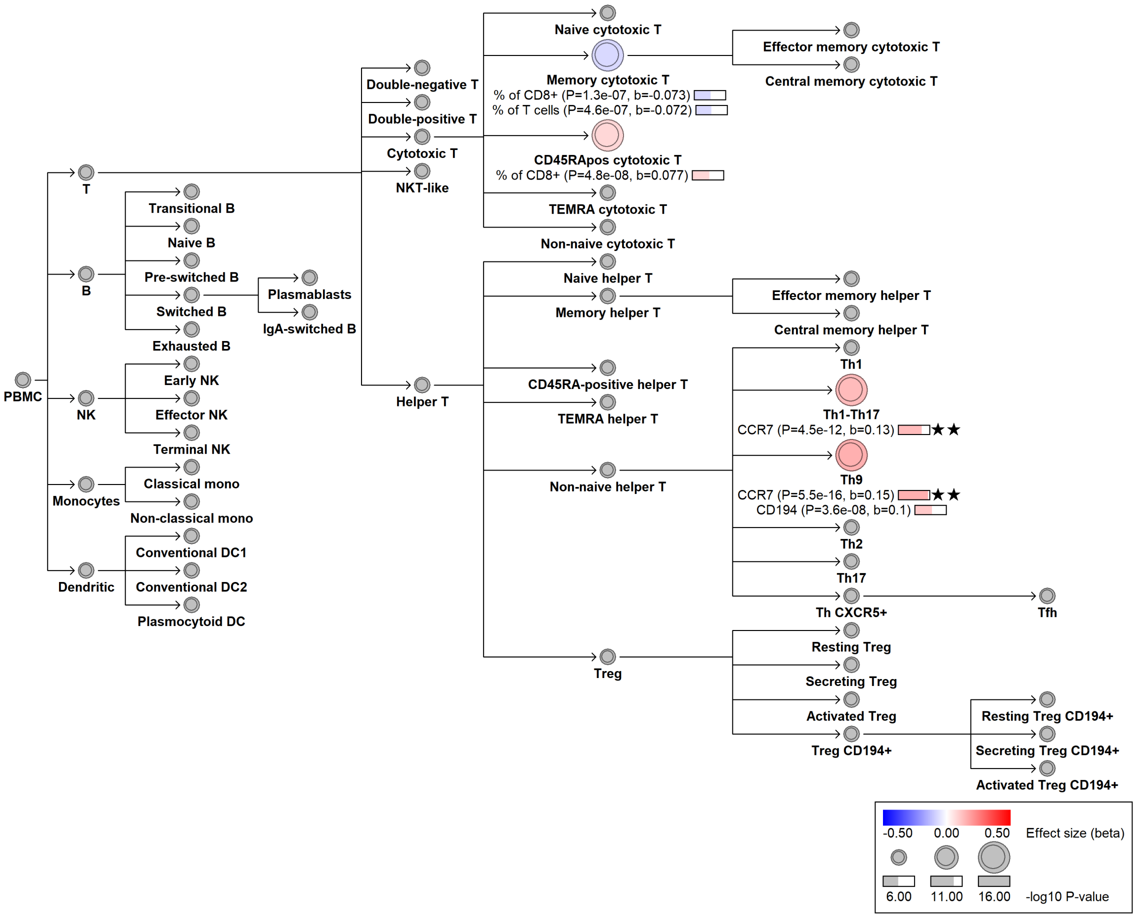

### chr3:46350157[T→G; EAF 8.943%] (rs35943069, cs_260)

Candidate genes: *CCR2*
Diseases associations: NA
Plasma pQTLs: NA

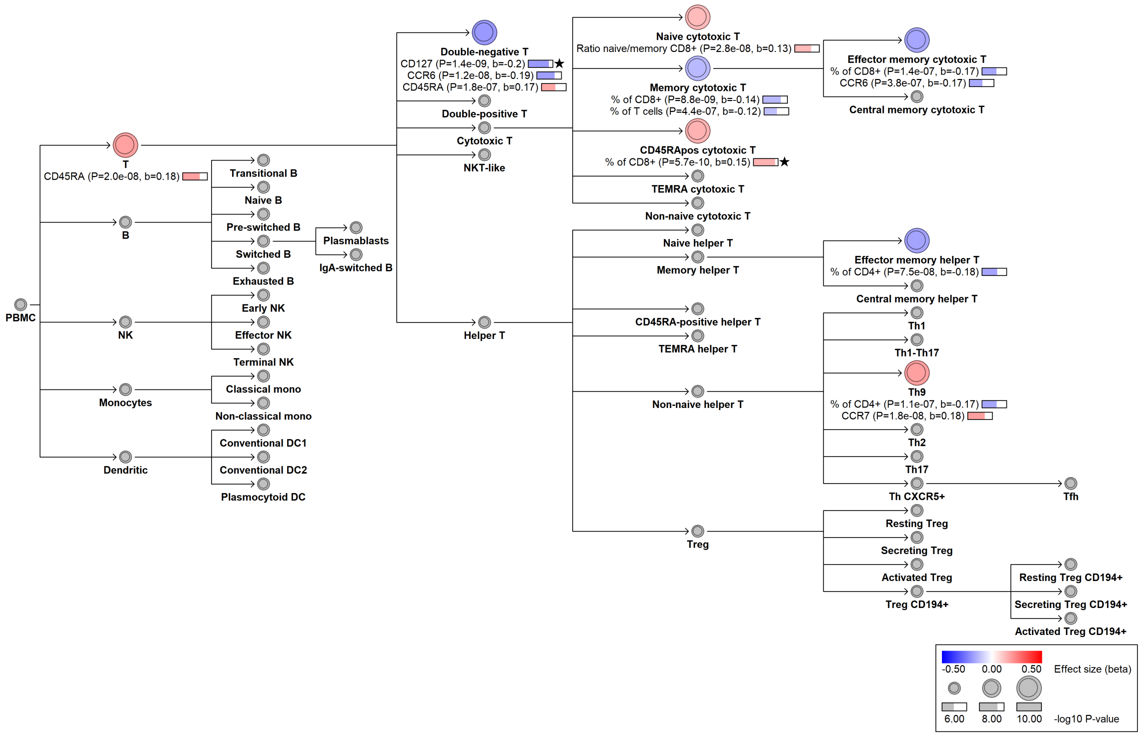

#### chr3:47240813[A→!; EAF 39.95%] (rs2276853, cs_98)

Candidate genes: *NBEAL2*
Diseases associations: NA
Plasma pQTLs: Immunoglobulin G

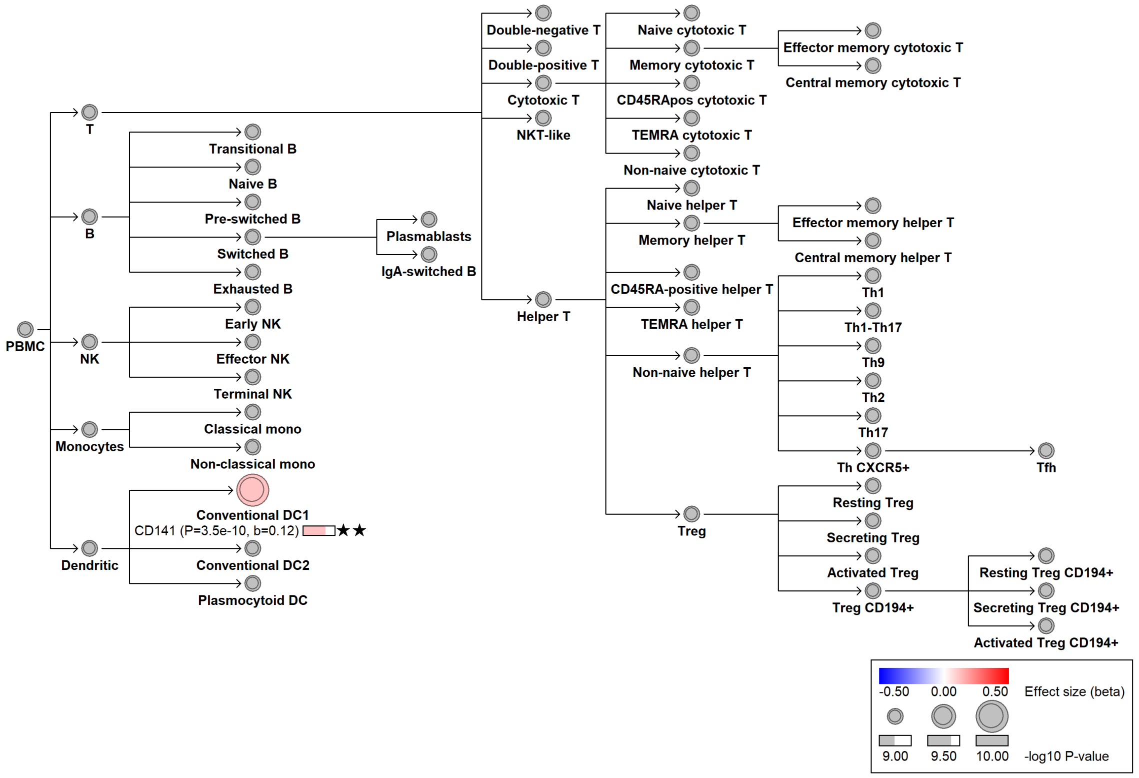

### chr3:52495250[C→T; EAF 4.634%] (rs758800, cs_51)

Candidate genes: *TNNC1*
Diseases associations: NA
Plasma pQTLs: NA

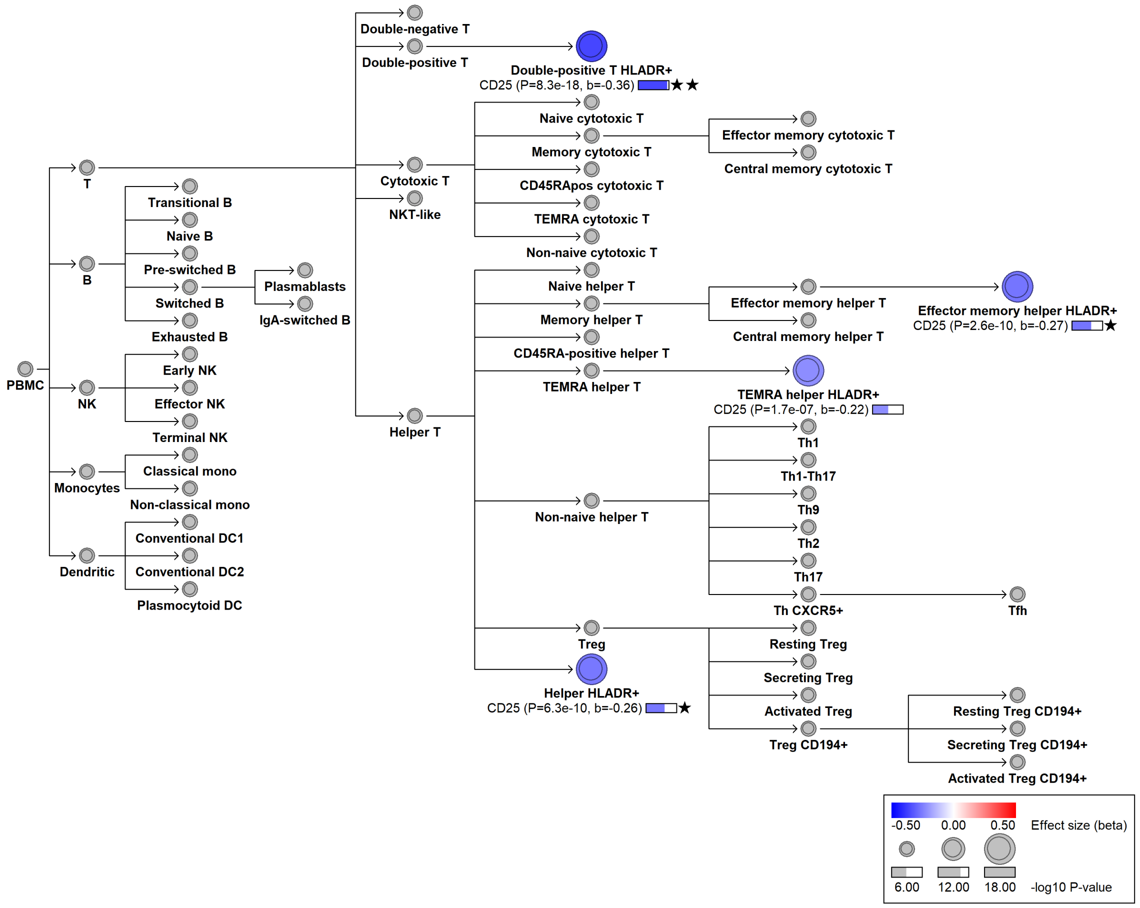

#### chr3:128599154[A→G; EAF 11.499%] (rs6782812, cs_94)

Candidate genes: *GATA2*
Diseases associations: NA
Plasma pQTLs: CAMP, CLC, CRISP3, PAPPA, PRG2, PRG3, RNASE3, VNN2

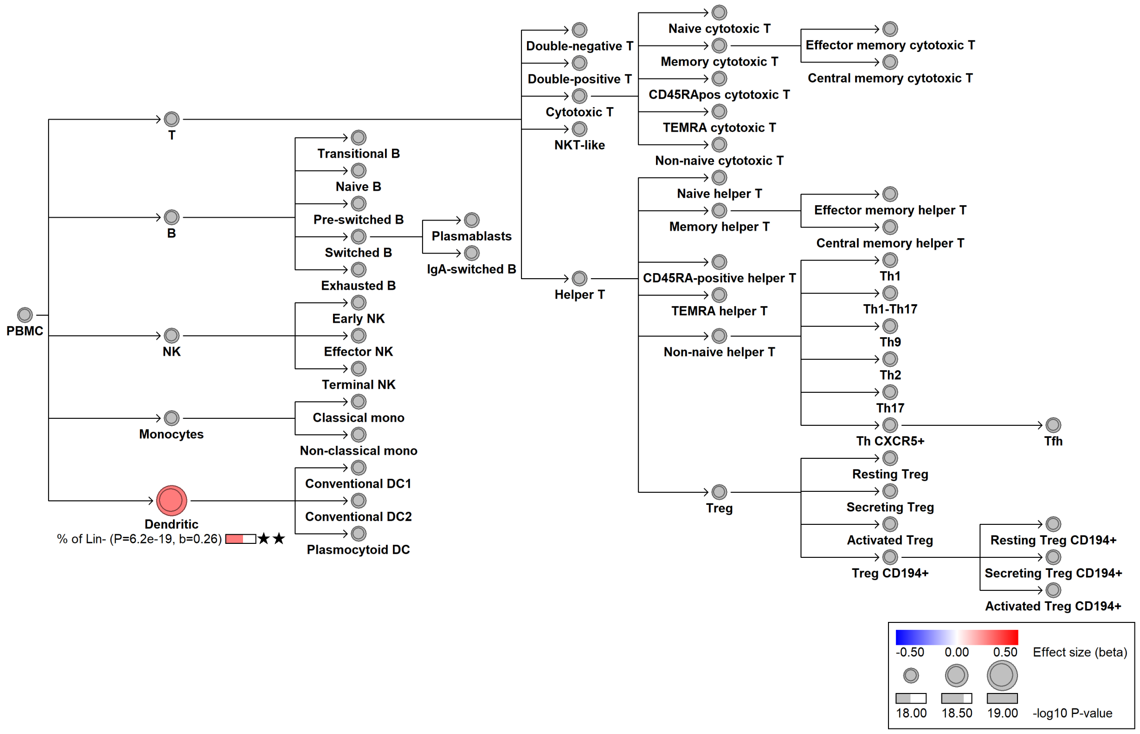

#### chr3:188417995[A→T; EAF 46.856%] (rs12485444, cs_198)

Candidate genes: *BCL6*
Diseases associations: Asthma, Non-melanoma skin cancer
Plasma pQTLs: NA

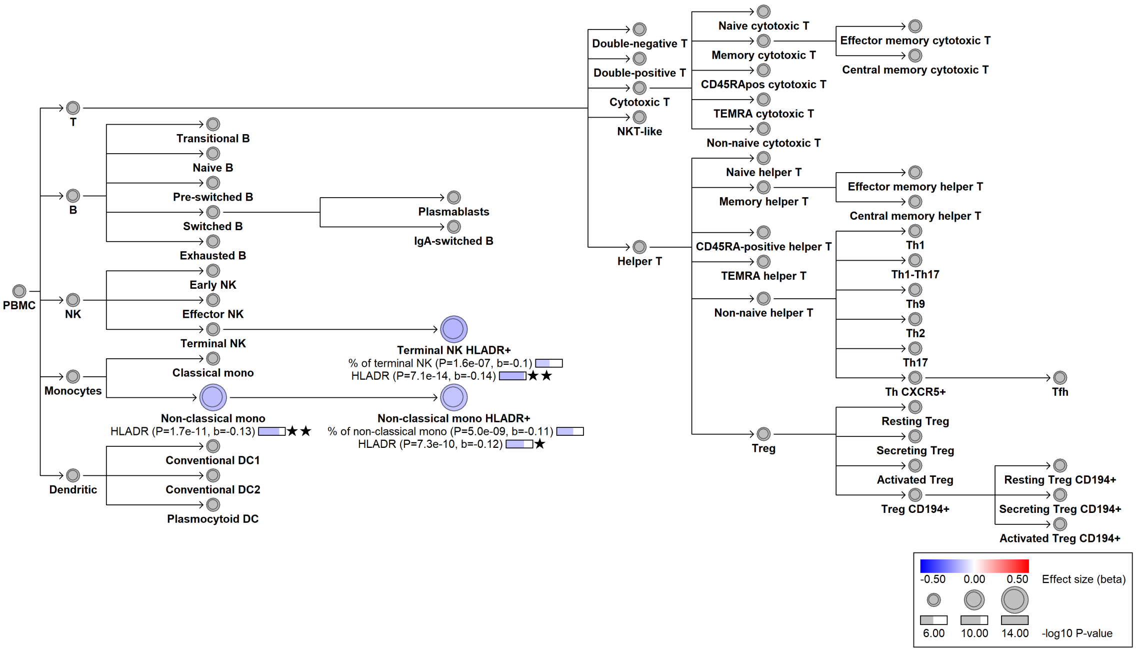

### chr4:15807551[G→!; EAF 31.834%] (chr4:15807551, cs_26)

Candidate genes: *CD38*
Diseases associations: NA
Plasma pQTLs: NA

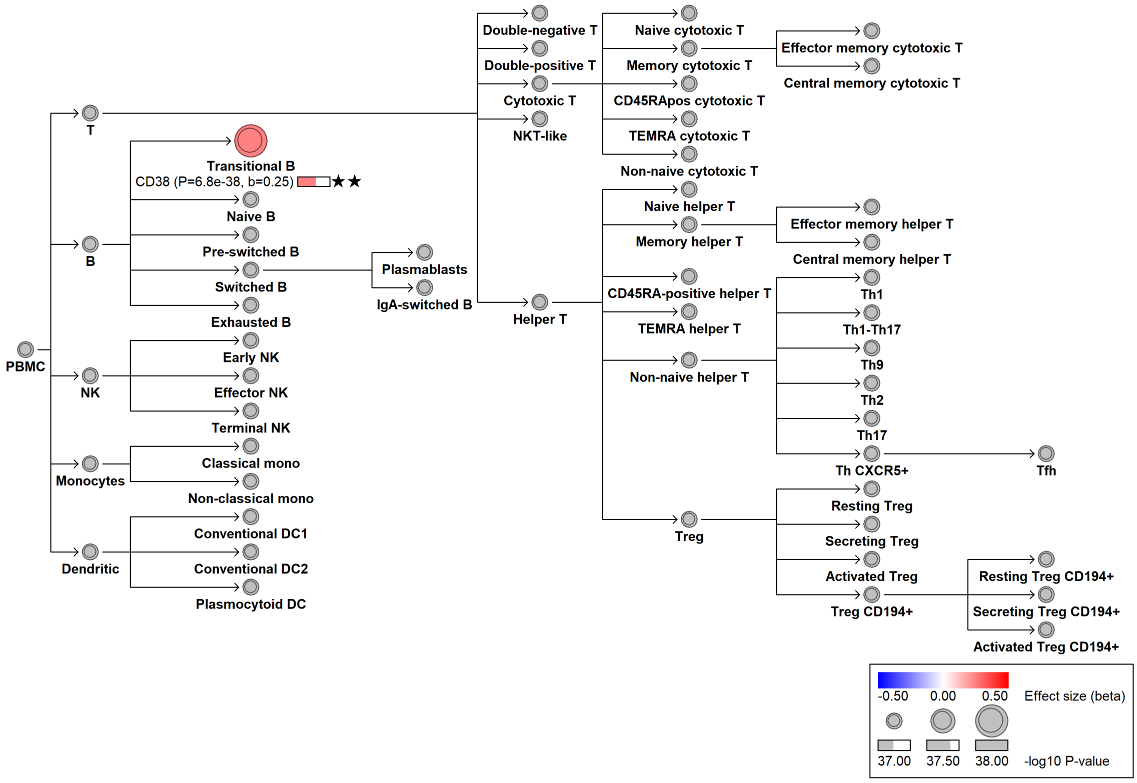

### chr4:38395752[G→T; EAF 46.788%] (rs2087197, cs_230)

Candidate genes: *KLF3*
Diseases associations: NA
Plasma pQTLs: NA

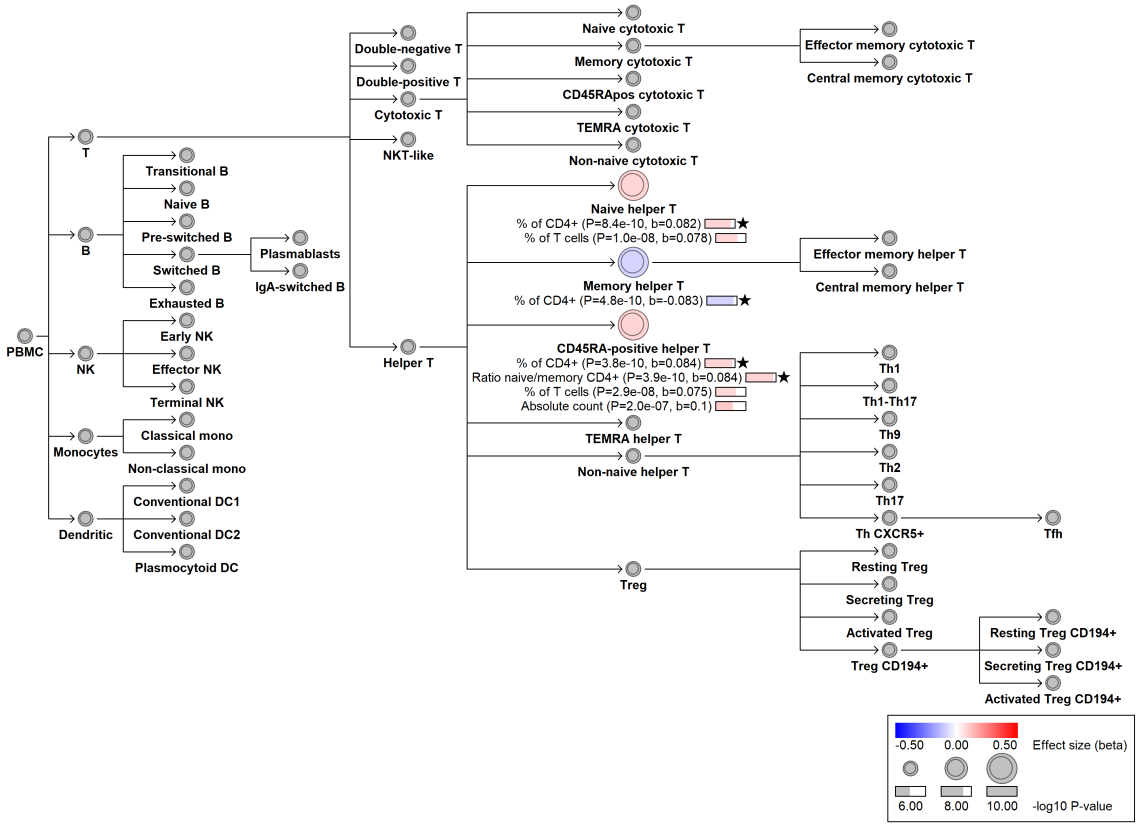

#### chr4:39818403[G→T; EAF 16.056%] (rs433247, cs_263)

Candidate genes: *PDS5A*
Diseases associations: NA
Plasma pQTLs: NA

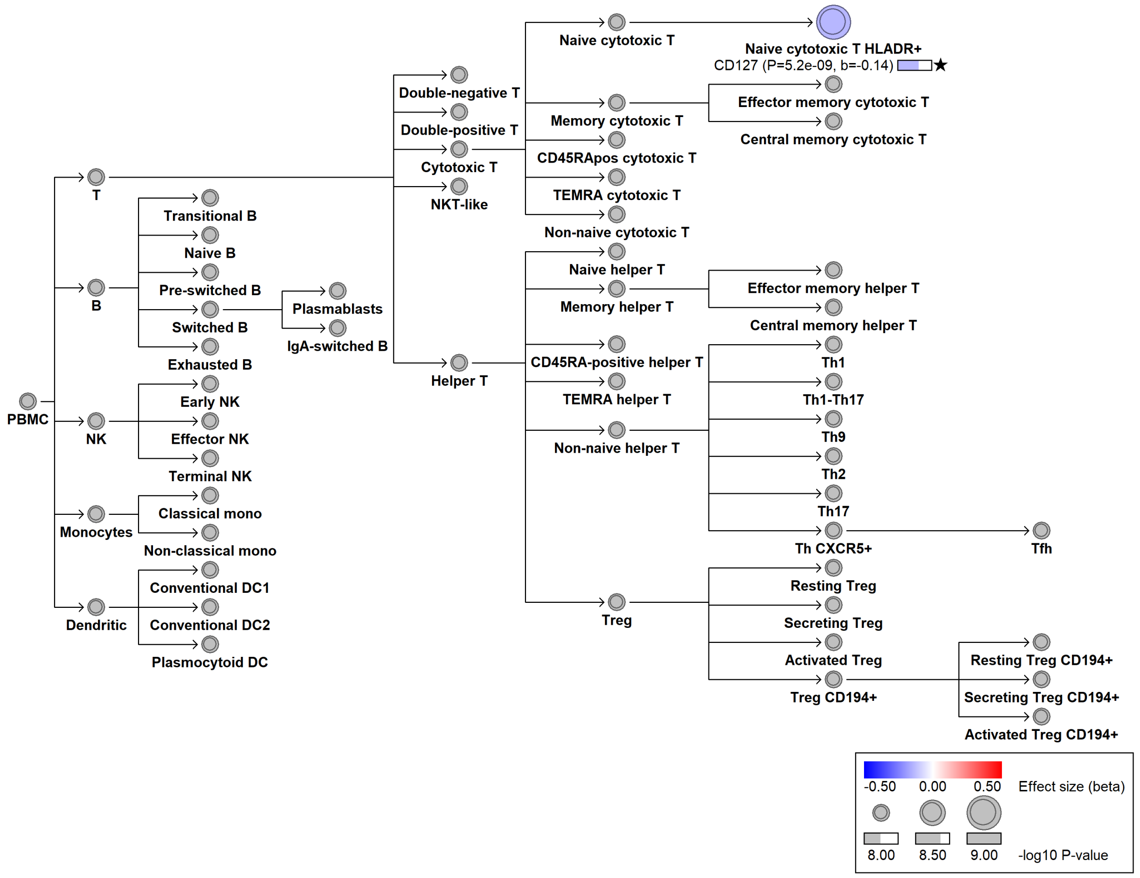

### chr4:83236290[C→G; EAF 34.254%] (rs2903918, cs_215)

Candidate genes: *PLAC8*
Diseases associations: NA
Plasma pQTLs: CLEC4C, ELOA, KCTD5

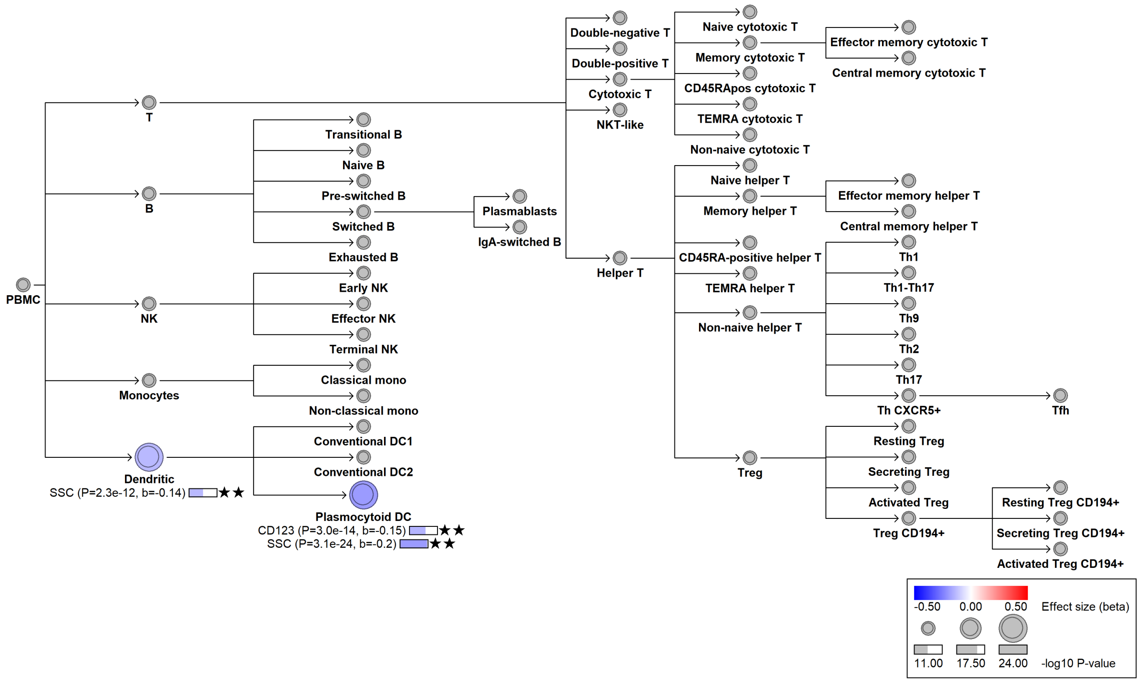

#### chr4:83240767[C→CT; EAF 28.643%] (rs5859867, cs_261)

Candidate genes: *PLAC8*
Diseases associations: Systemic lupus erythematosus
Plasma pQTLs: BLNK, CCDC50

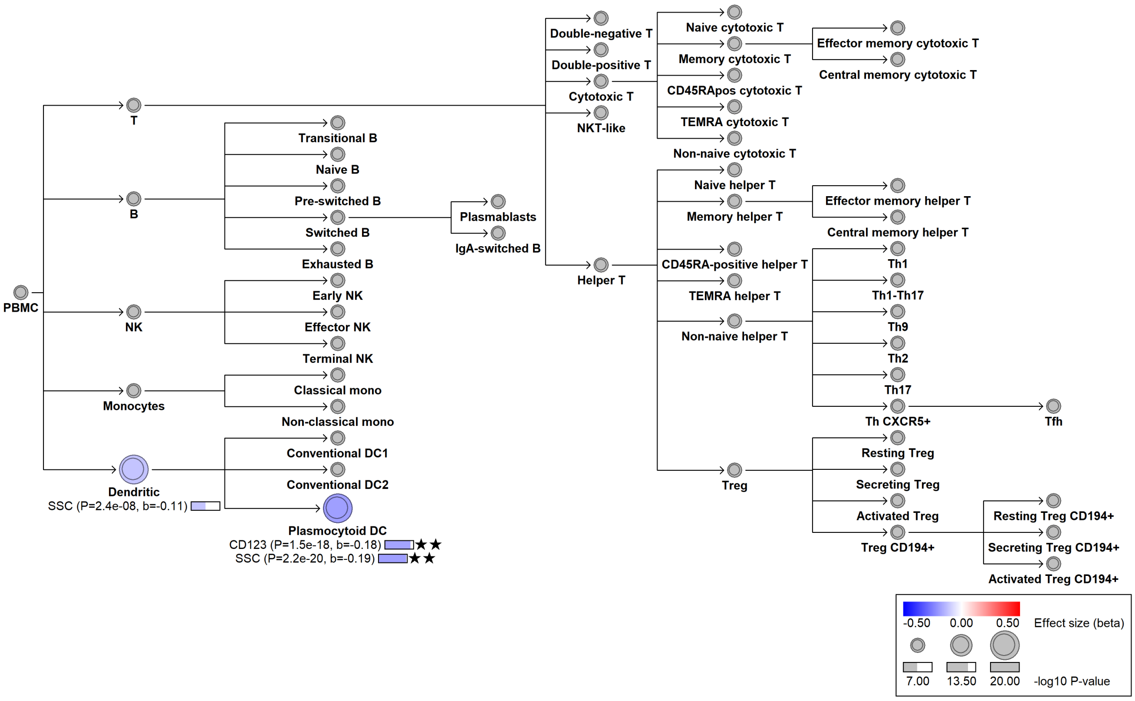

#### chr4:89836143[C→A; EAF 19.696%] (rs1372518, cs_86)

Candidate genes: *SNCA*
Diseases associations: Parkinson's disease
Plasma pQTLs: AKR1C3

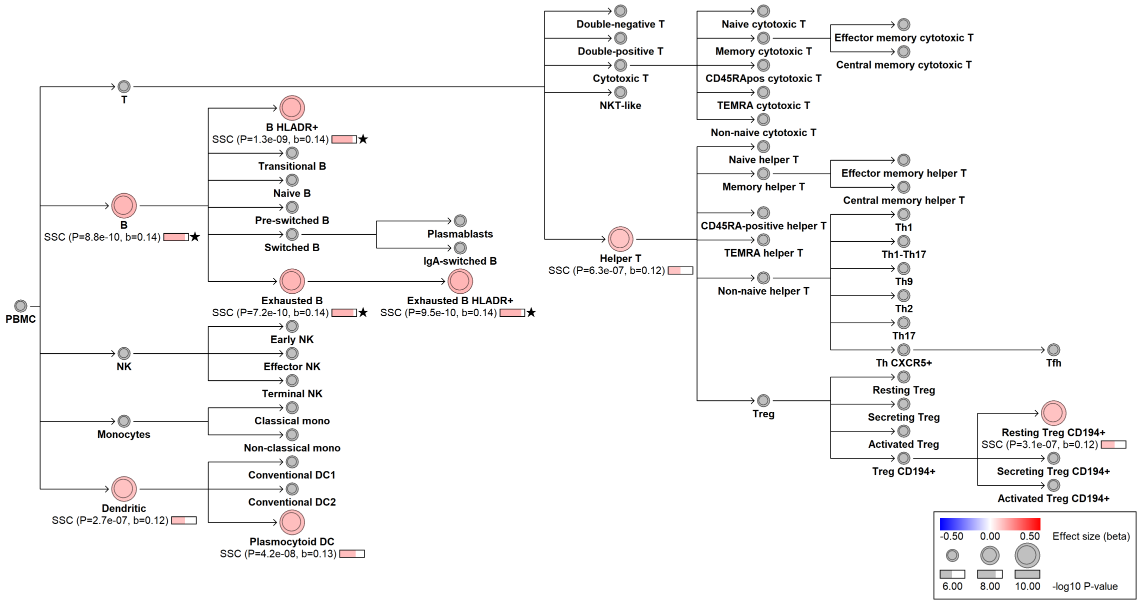

#### chr4:102502169[G→T; EAF 31.991%] (rs2272676, cs_175)

Candidate genes: *NFKB1*
Diseases associations: Allergy, Asthma, Chronic diseases of tonsils, Eczema, Ulcerative colitis
Plasma pQTLs: CA3, IL5RA, NFKB1, RRM1, SKAP1, SPTBN2

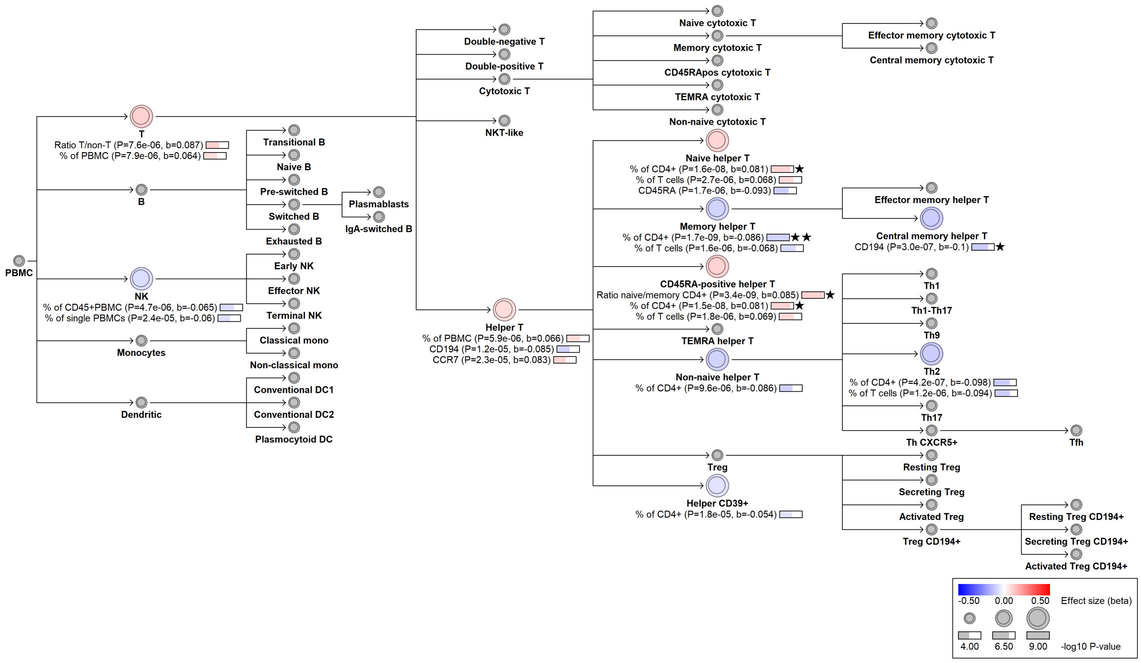

#### chr4:112194663[G→A; EAF 38.117%] (rs7682734, cs_96)

Candidate genes: *FAM241A*
Diseases associations: NA
Plasma pQTLs: COLEC12

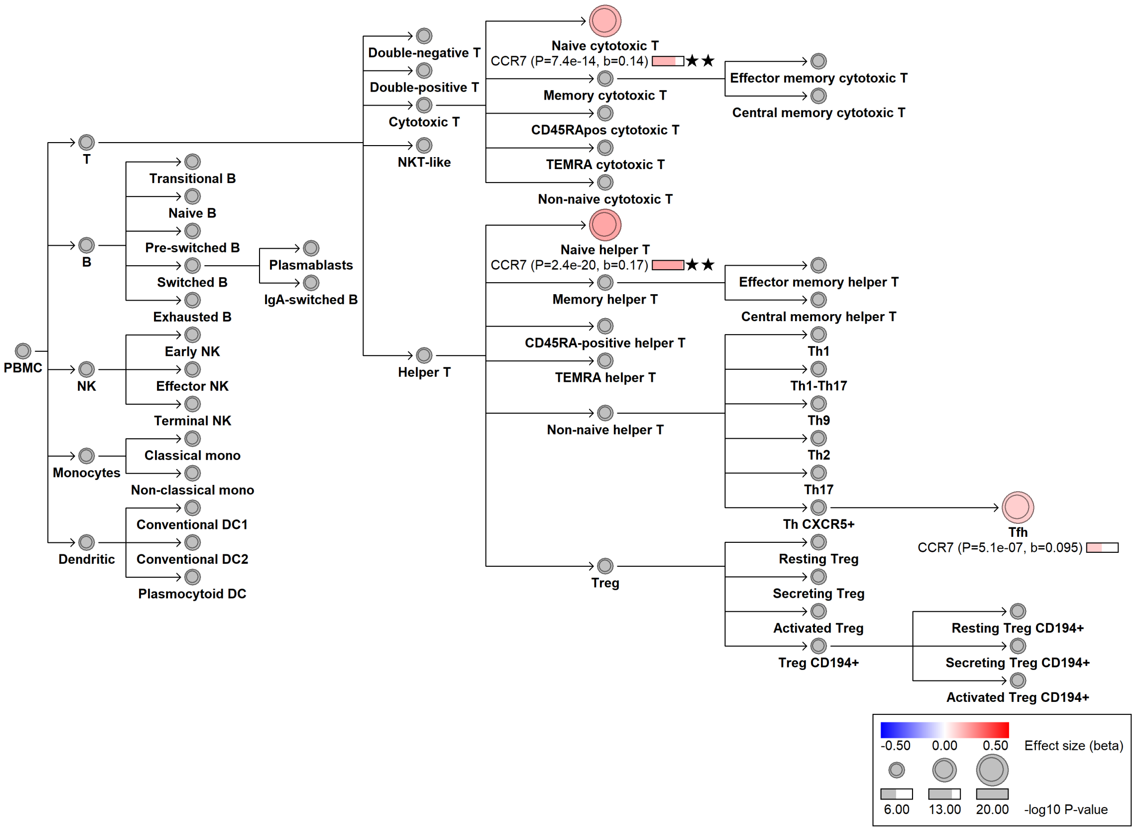

#### chr4:186199057[A→C; EAF 35.304%] (rs13146272, cs_195)

Candidate genes: *CYP4V2*
Diseases associations: NA
Plasma pQTLs: NA

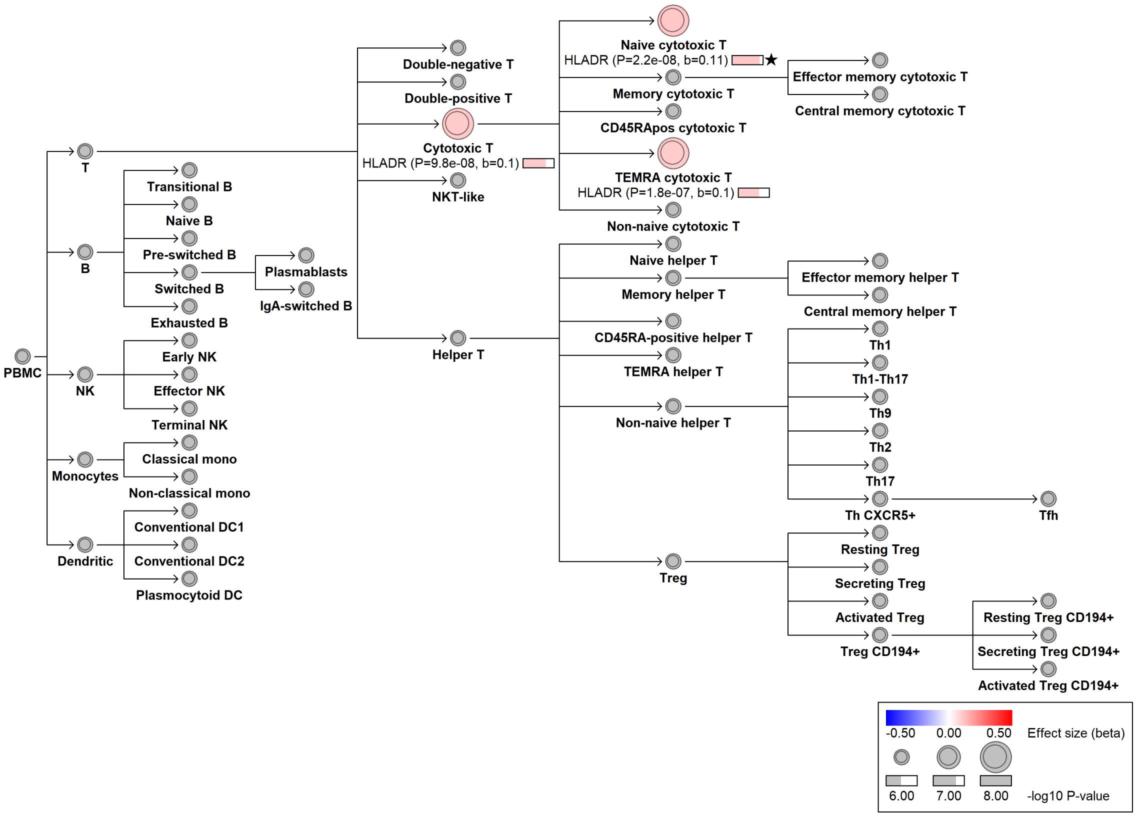

#### chr5:1104823[T→C; EAF 42.828%] (rs35188965, cs_156)

Candidate genes: *SLC12A7*
Diseases associations: NA
Plasma pQTLs: NA

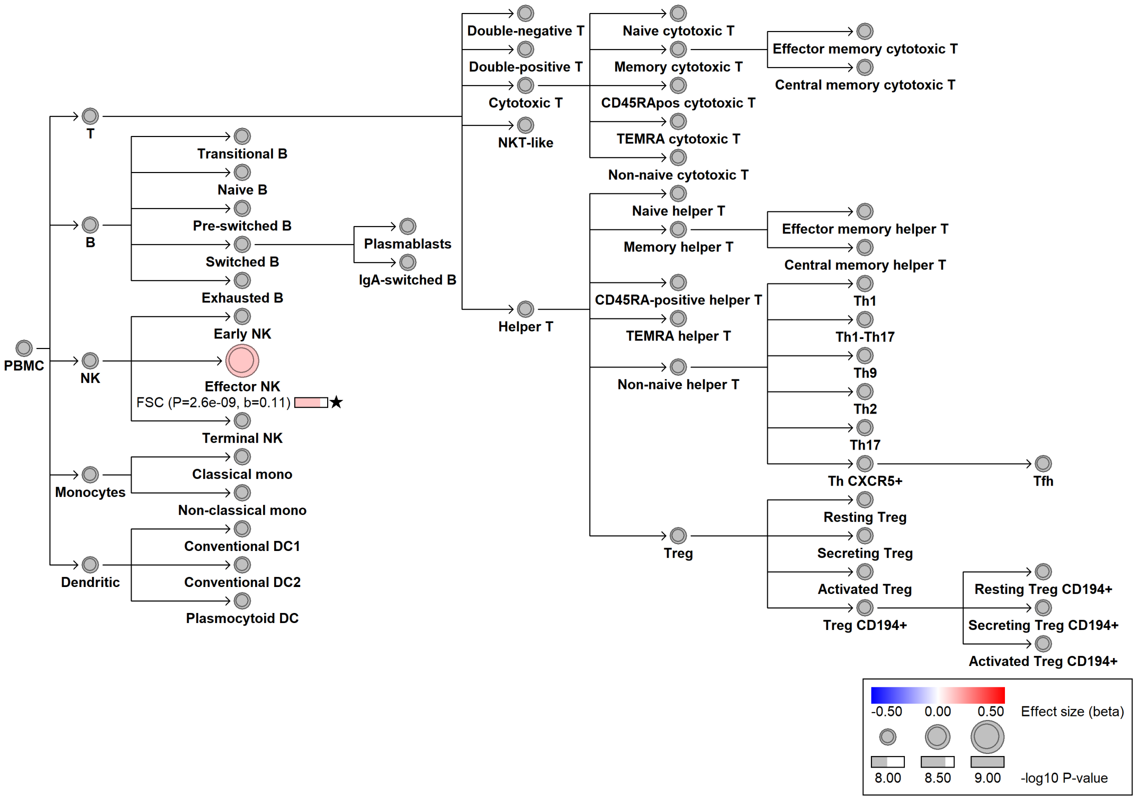

#### chr5:35829849[!→CAA; EAF 34.021%] (rs375025698, cs_87)

Candidate genes: *IL7R*
Diseases associations: Asthma, Autoimmune thyroiditis, Primary biliary cholangitis
Plasma pQTLs: NA

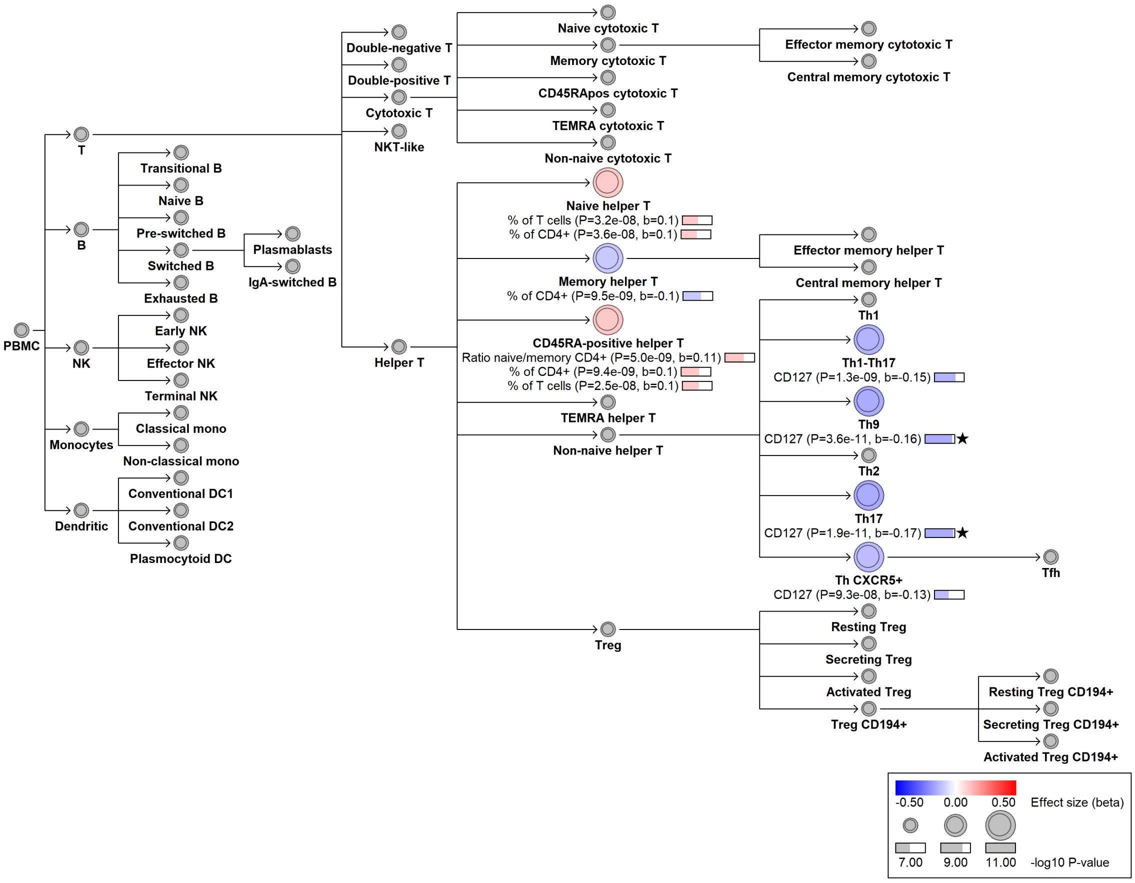

#### chr5:35831056[C→G; EAF 30.824%] (rs10050705, cs_128)

Candidate genes: *IL7R*
Diseases associations: NA
Plasma pQTLs: NA

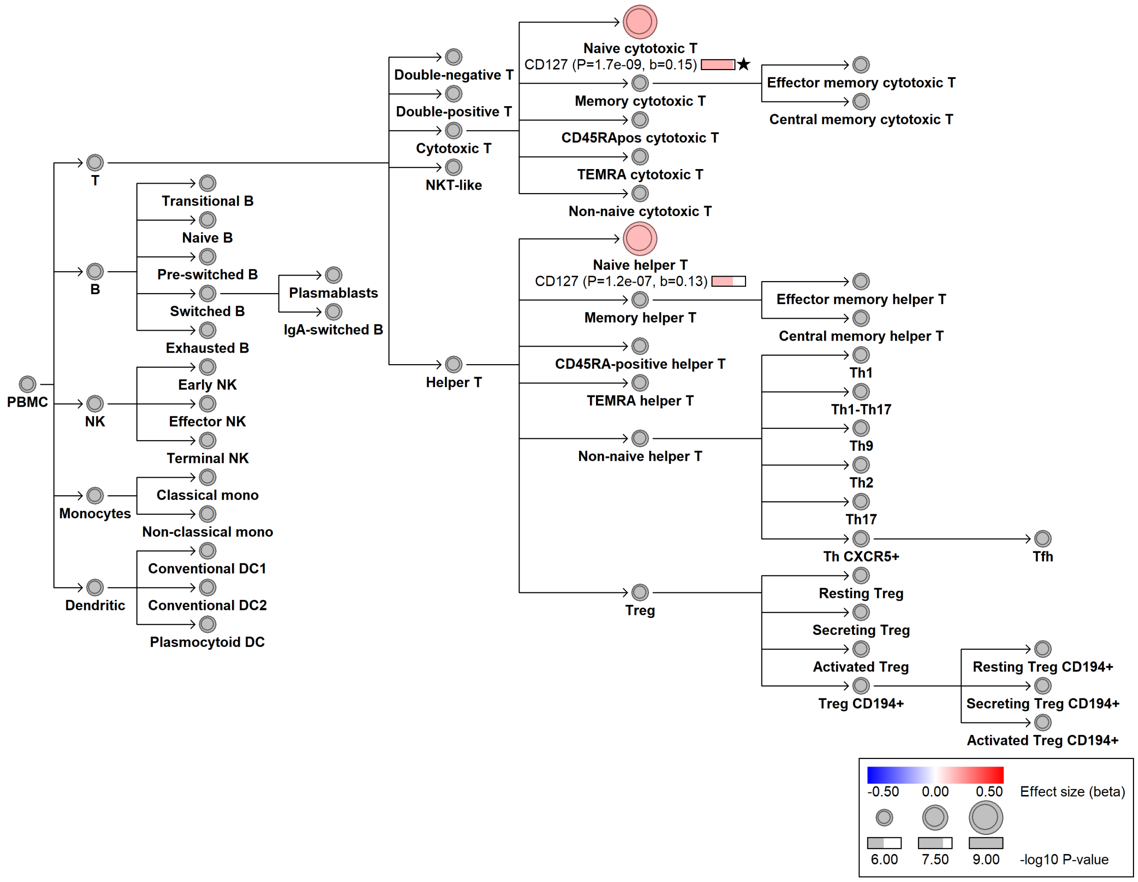

### chr5:35855308[T→C; EAF 13.195%] (rs4869492, cs_46)

Candidate genes: *IL7R*
Diseases associations: NA
Plasma pQTLs: NA

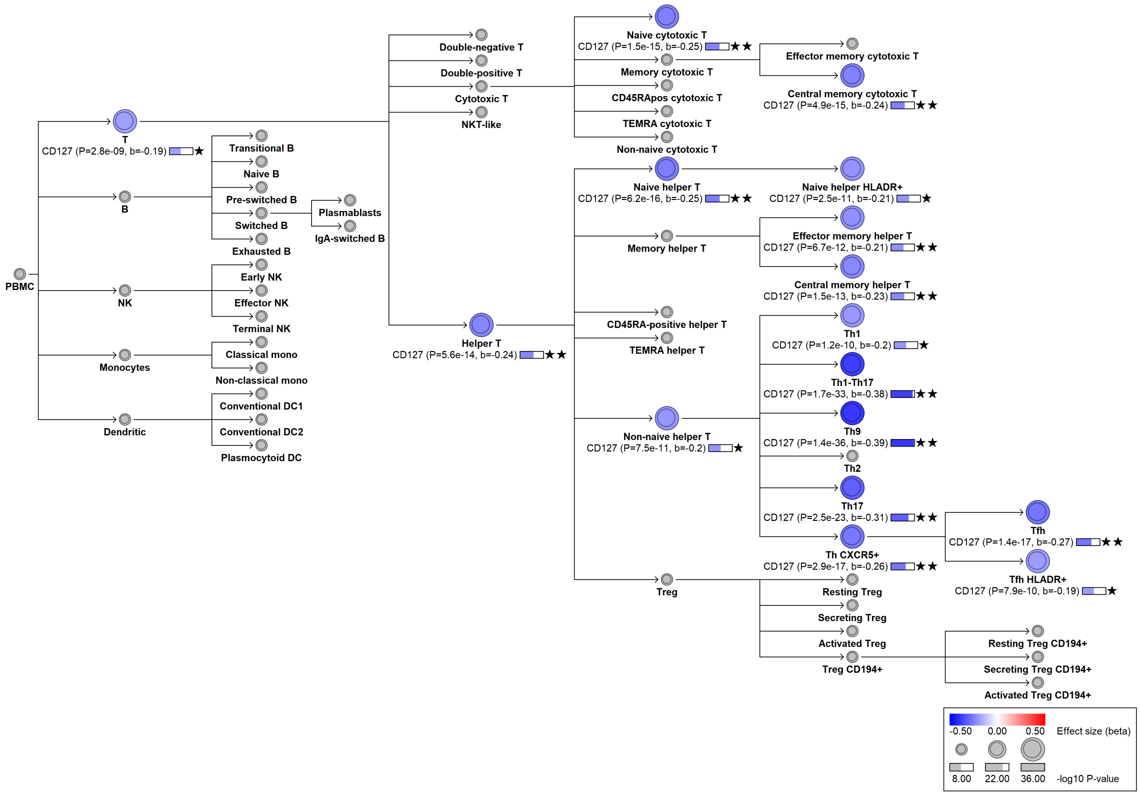

#### chr5:35874473[C→T; EAF 26.763%] (rs6897932, cs_257)

Candidate genes: *IL7R*
Diseases associations: Allergy, Asthma, Eczema, Multiple sclerosis, Primary biliary cholangitis, Systemic lupus erythematosus
Plasma pQTLs: CCL21, CD48, CD5, CD6, GZMA, IL7R, LY9, PAG1, SELPLG, SIT1

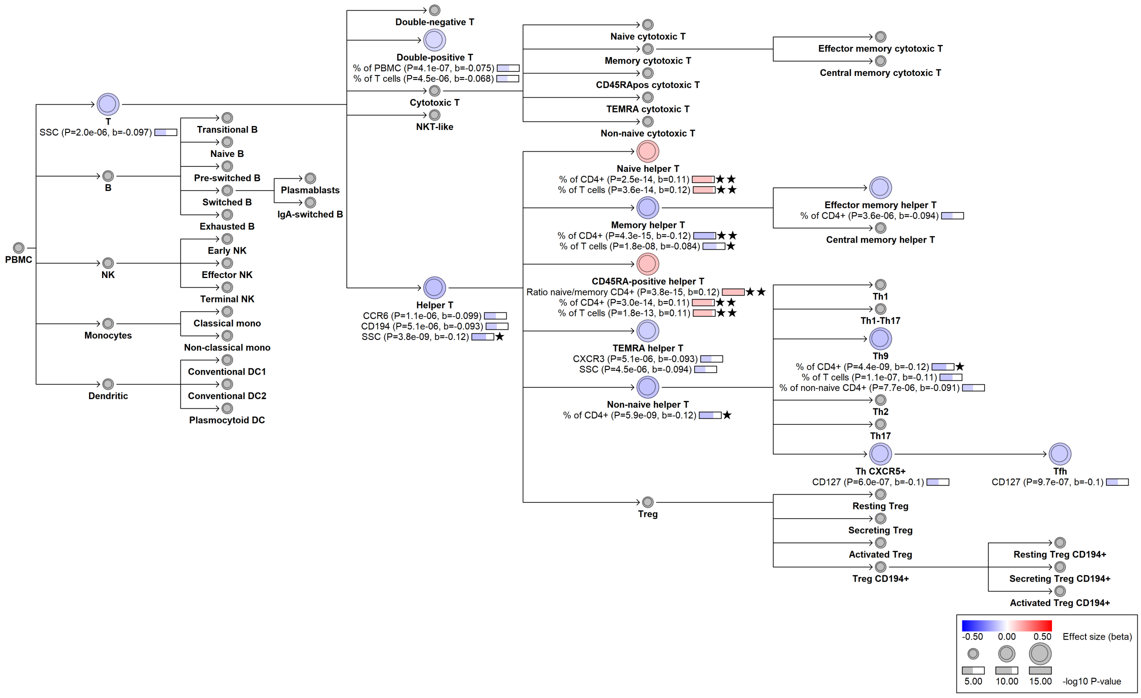

#### chr5:134143625[C→G; EAF 0.967%] (rs149469800, cs_55)

Candidate genes: *TCF7*
Diseases associations: NA
Plasma pQTLs: NA

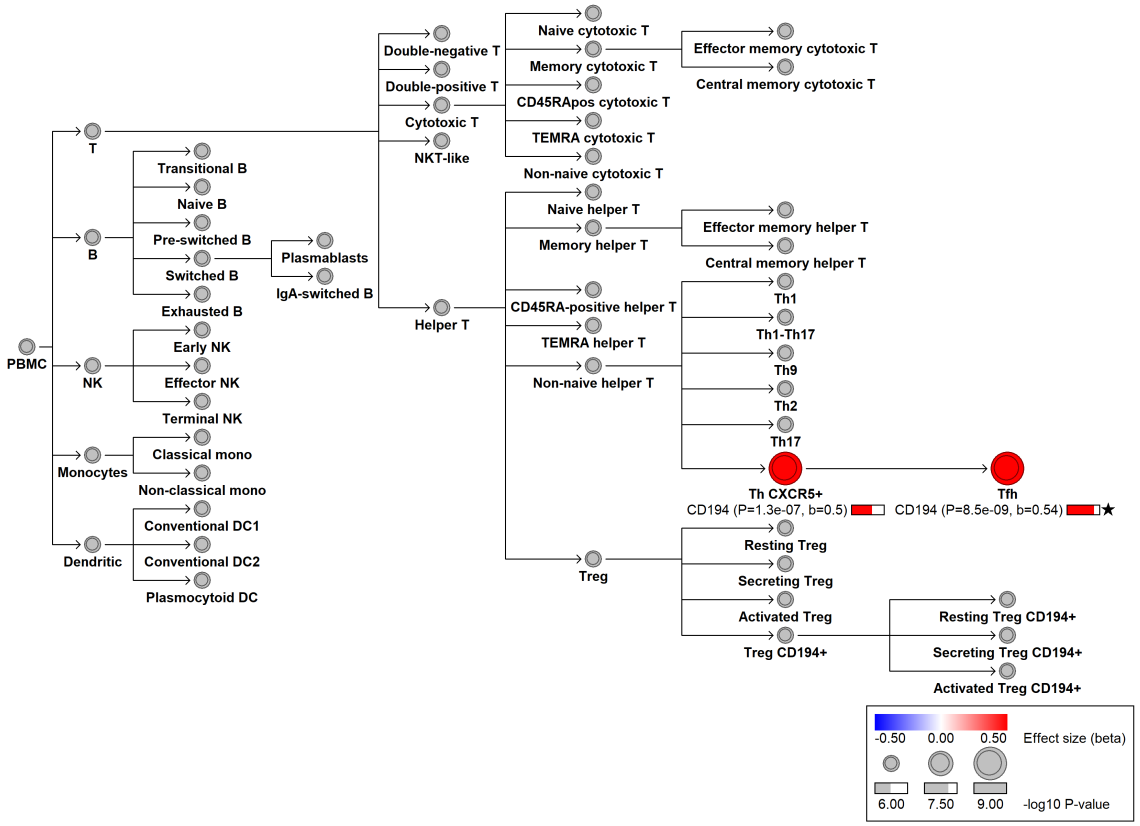

#### chr5:139394718[C→T; EAF 10.538%] (rs73255289, cs_71)

Candidate genes: *STING1*
Diseases associations: NA
Plasma pQTLs: NA

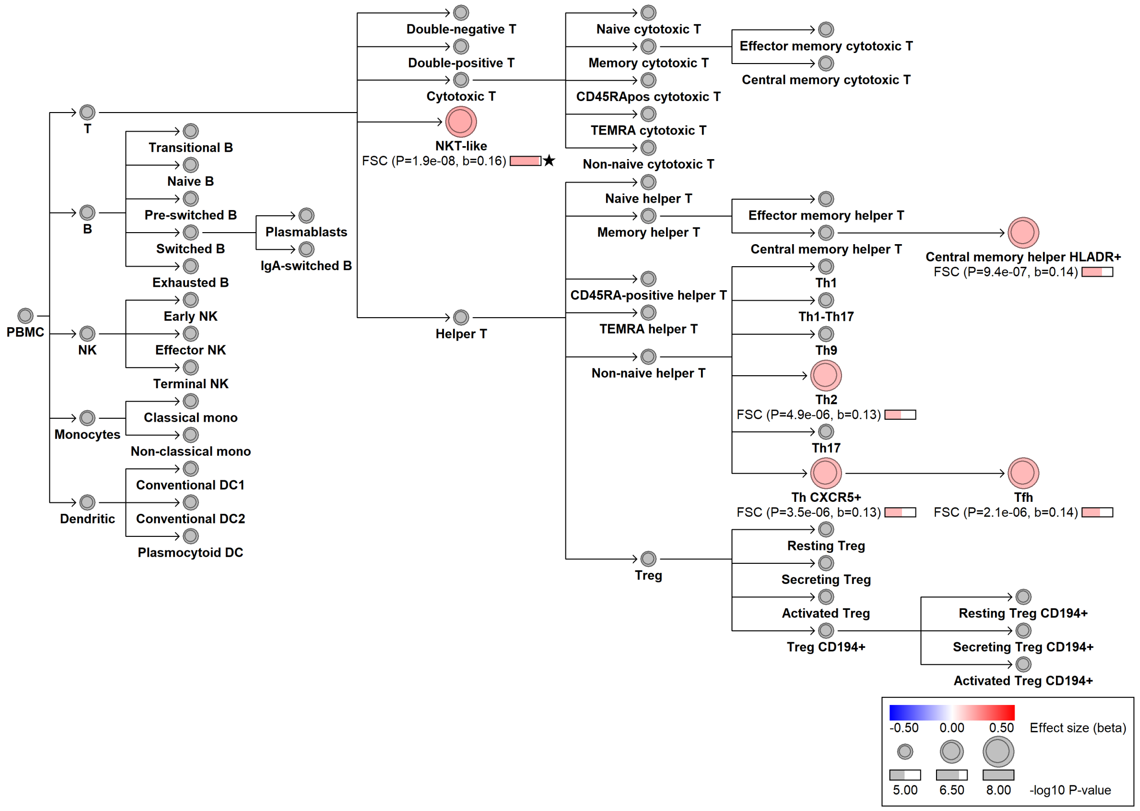

### chr5:139478340[C→G; EAF 12.985%] (rs78233829, cs_131)

Candidate genes: *STING1*
Diseases associations: NA
Plasma pQTLs: NA

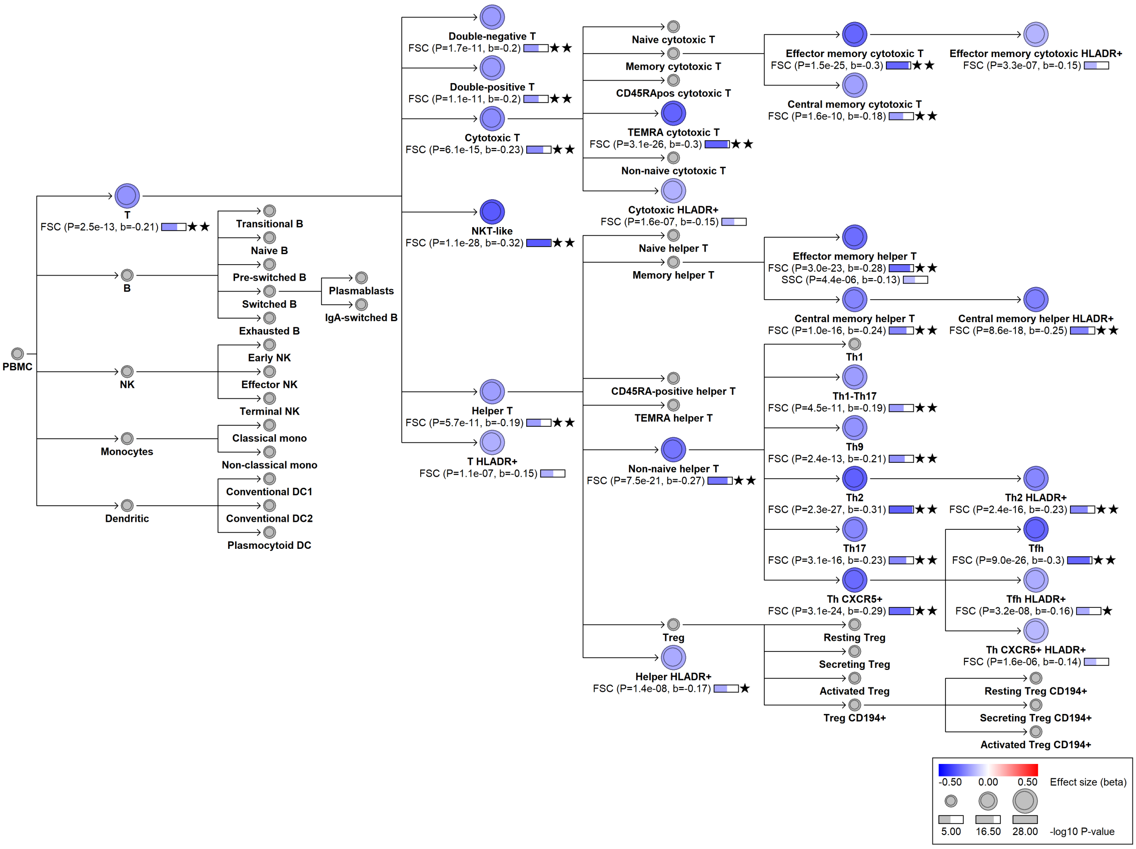

#### chr5:140637262[G→A; EAF 27.269%] (rs5744441, cs_174)

Candidate genes: *CD14*
Diseases associations: NA
Plasma pQTLs: CD14

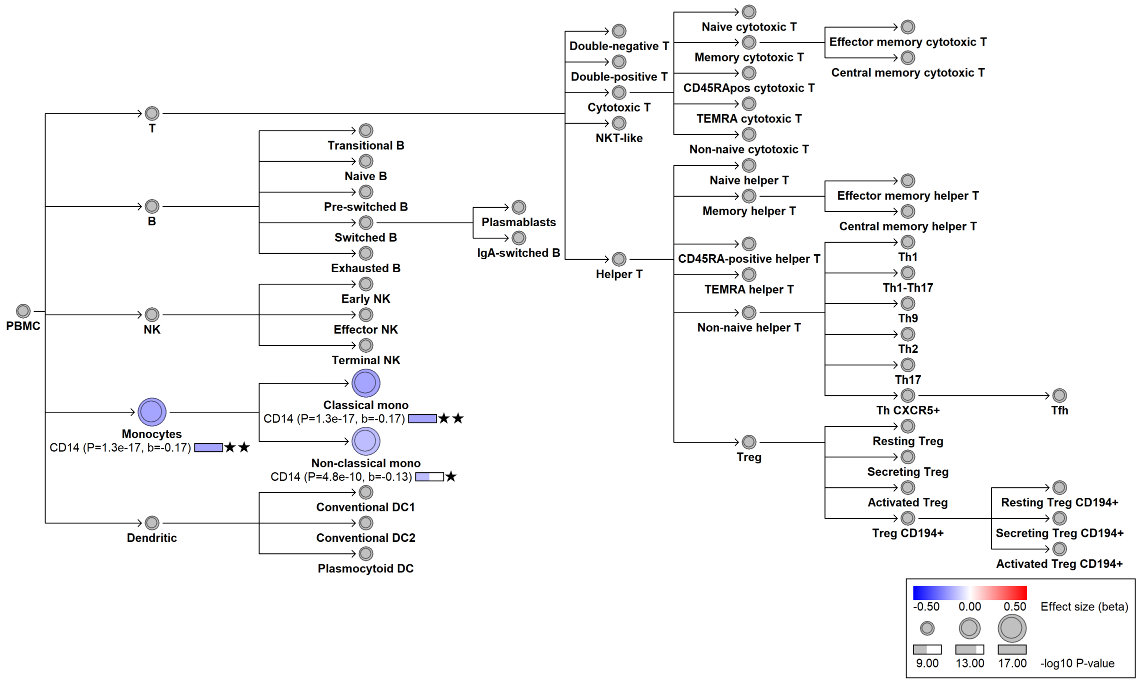

### chr5:158853964[C→A; EAF 7.045%] (rs56364901, cs_52)

Candidate genes: *EBF1*
Diseases associations: NA
Plasma pQTLs: CD22, CD72, CD79B, DRAXIN, FCER2, FCRL1, IGLL1, TCL1A

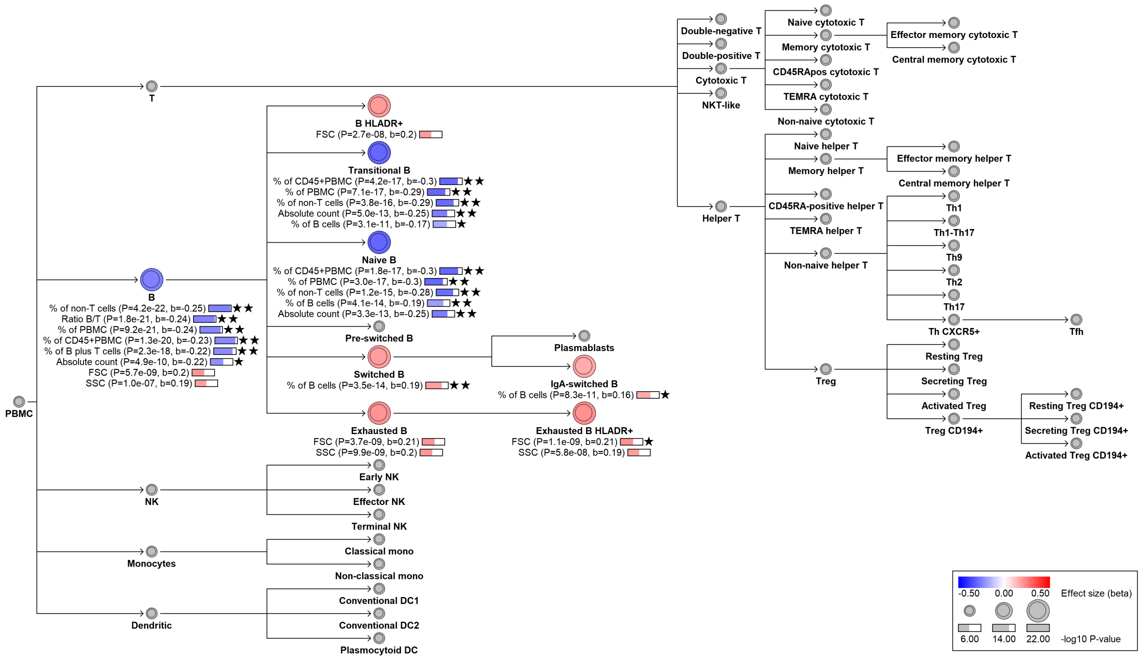

#### chr6:14091451[C→!; EAF 26.833%] (chr6:14091451, cs_88)

Candidate genes: *CD83*
Diseases associations: NA
Plasma pQTLs: NA

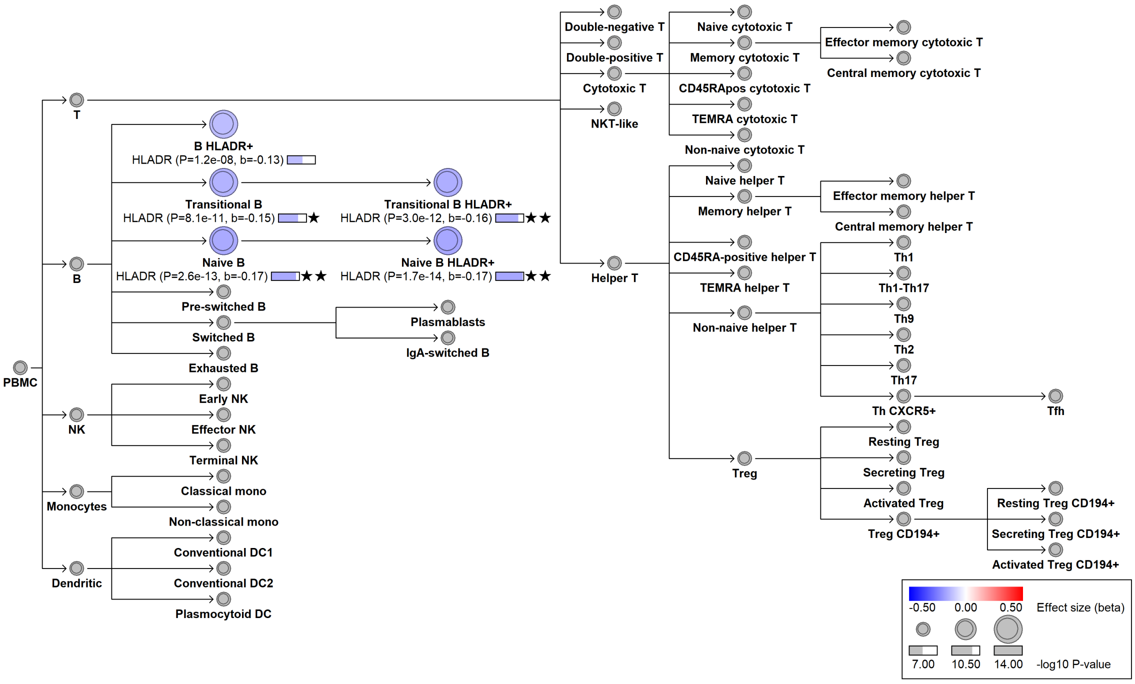

### chr6:14095524[G→A; EAF 5.153%] (rs74405933, cs_209)

Candidate genes: *CD83*
Diseases associations: Rheumatoid arthritis, Systemic lupus erythematosus
Plasma pQTLs: NA

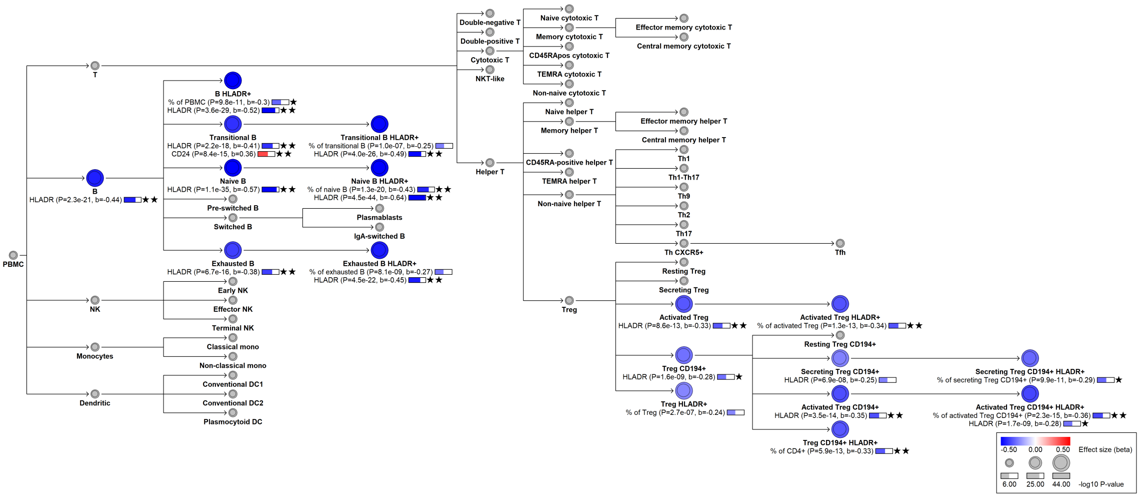

### chr6:15880293[C→G; EAF 8.876%] (rs2066400, cs_203)

Candidate genes: *MYLIP*
Diseases associations: NA
Plasma pQTLs: NA

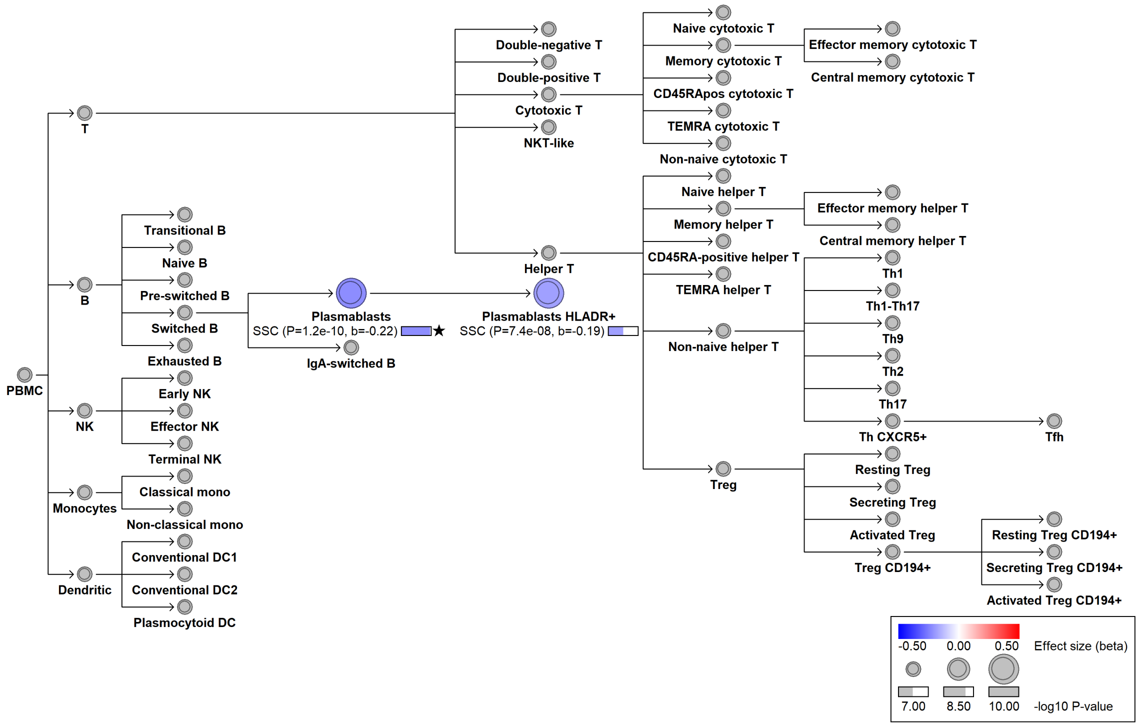

#### chr6:29941733[T→C; EAF 37.008%] (rs9260096, cs_235)

Candidate genes: *HLA region*
Diseases associations: NA
Plasma pQTLs: NA

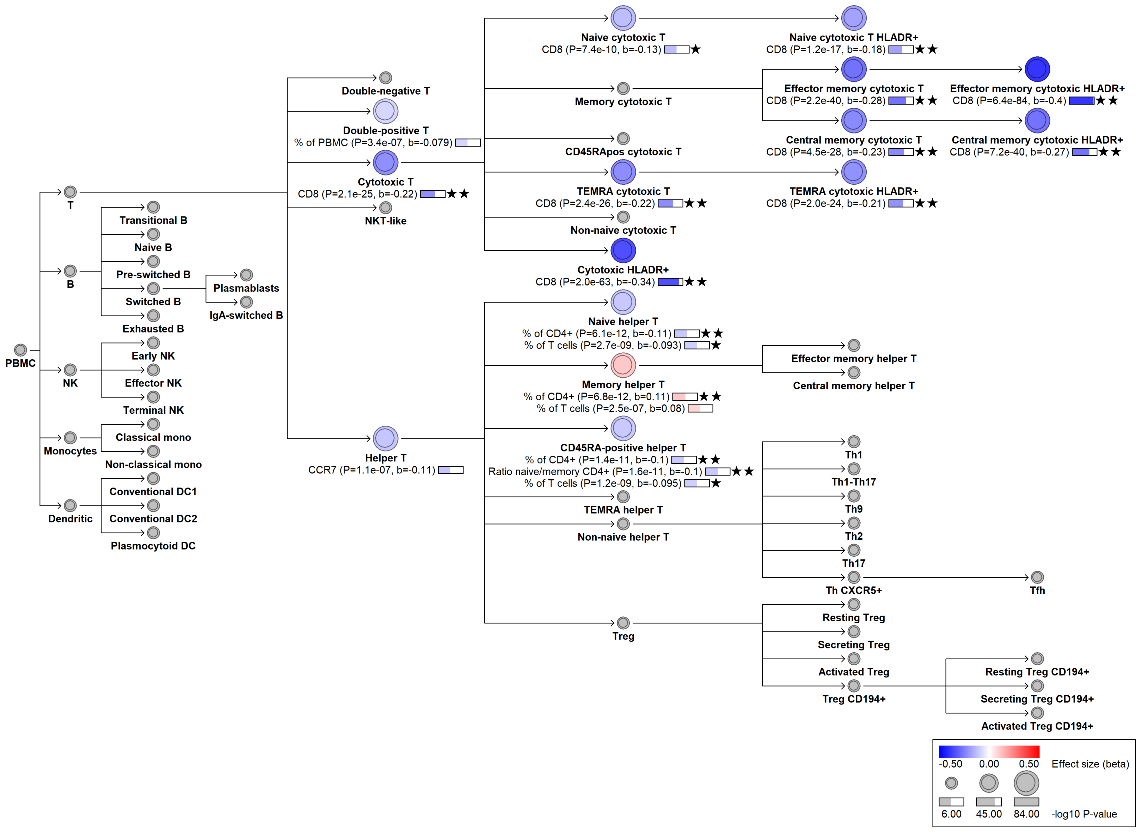

#### chr6:29953145[!→CTAACTAAA; EAF 3.604%] (rs1364270185, cs_236)

Candidate genes: *HLA region*
Diseases associations: NA
Plasma pQTLs: NA

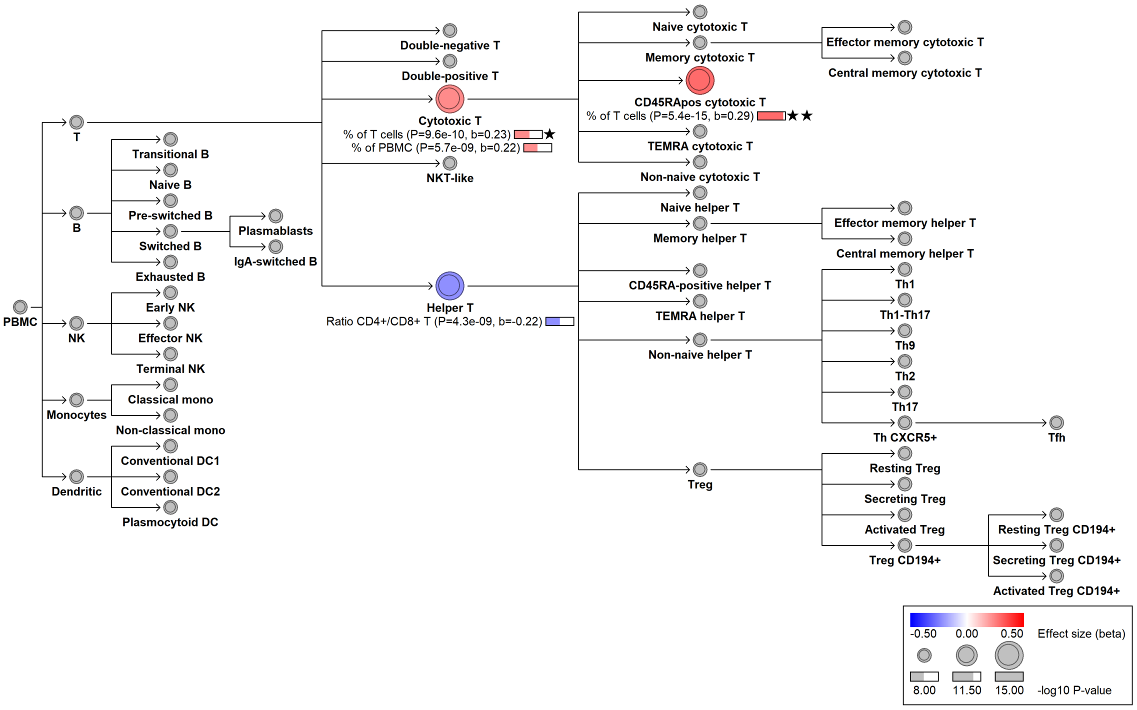

### chr6:31271999[A→C; EAF 28.838%] (rs2074493, cs_237)

Candidate genes: *HLA region*
Diseases associations: NA
Plasma pQTLs: NA

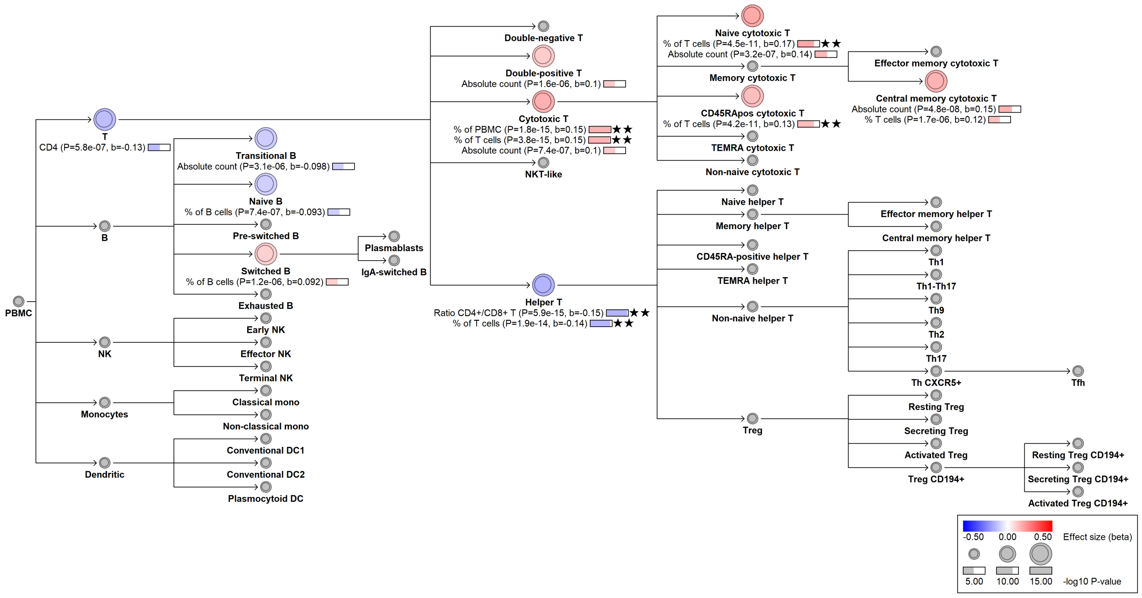

### chr6:31334507[G→C; EAF 20.932%] (rs4947308, cs_238)

Candidate genes: *HLA region*
Diseases associations: NA
Plasma pQTLs: NA

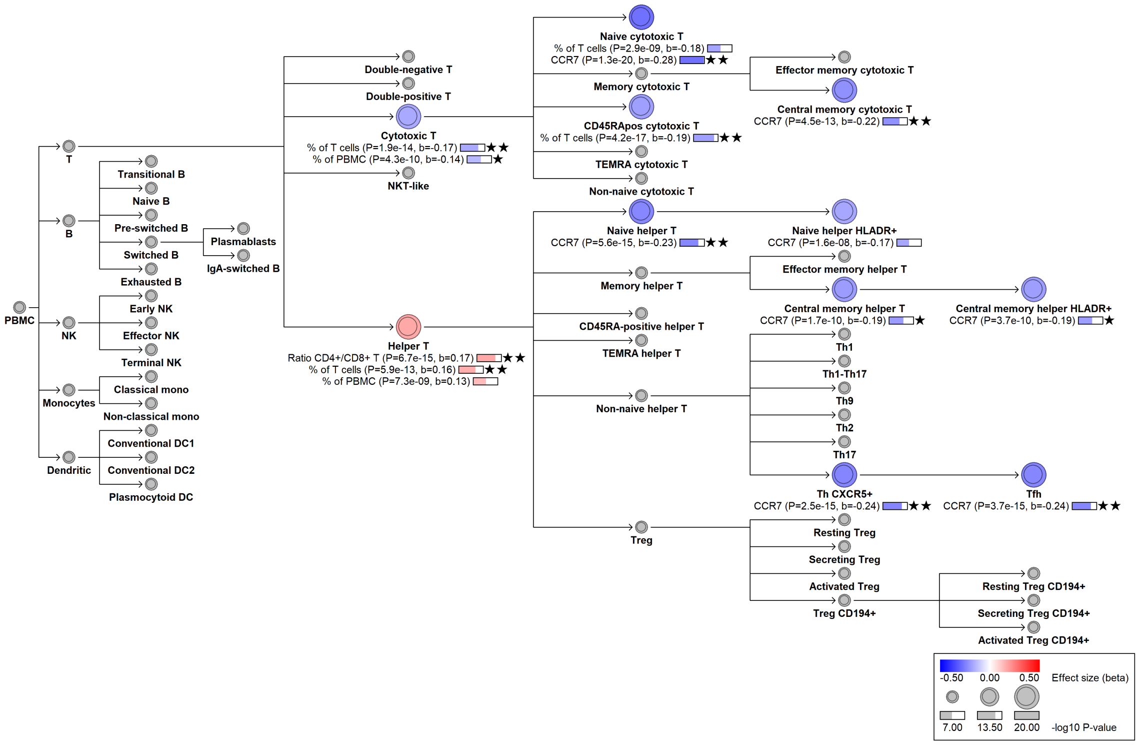

### chr6:31356965[C→T; EAF 15.525%] (rs147324178, cs_239)

Candidate genes: *HLA region*
Diseases associations: NA
Plasma pQTLs: NA

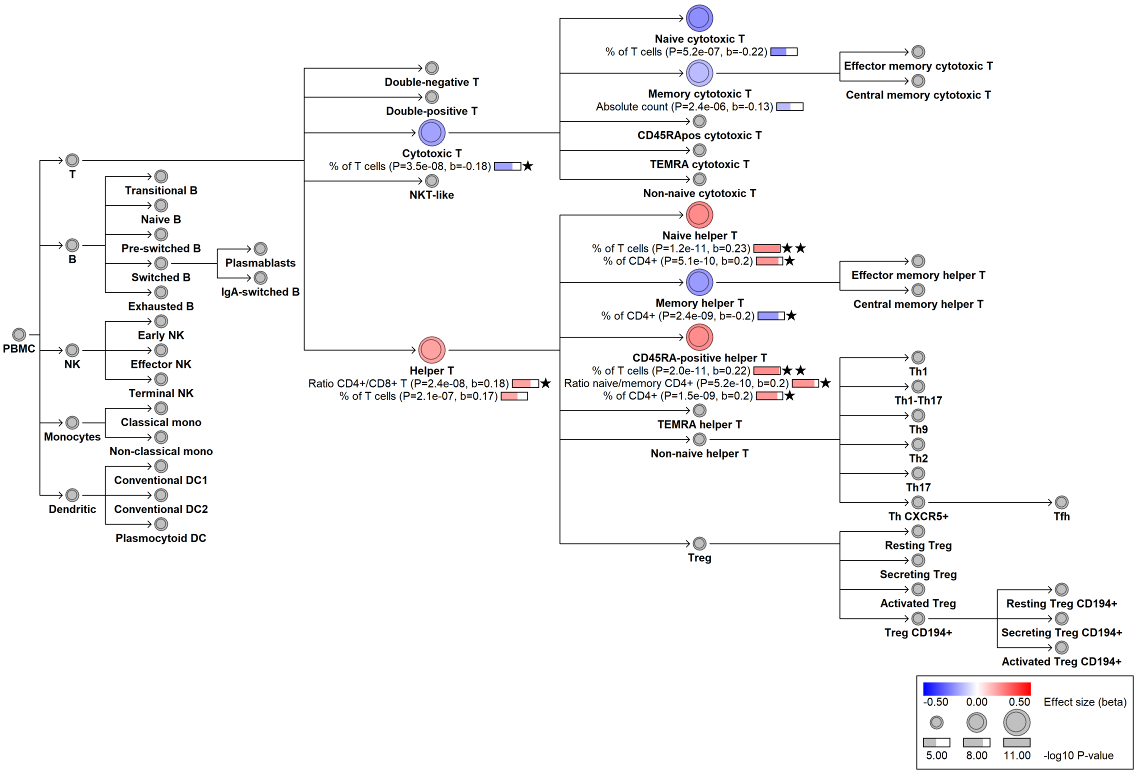

#### chr6:31358074[G→T; EAF 43.376%] (rs28694041, cs_240)

Candidate genes: *HLA region*
Diseases associations: NA
Plasma pQTLs: NA

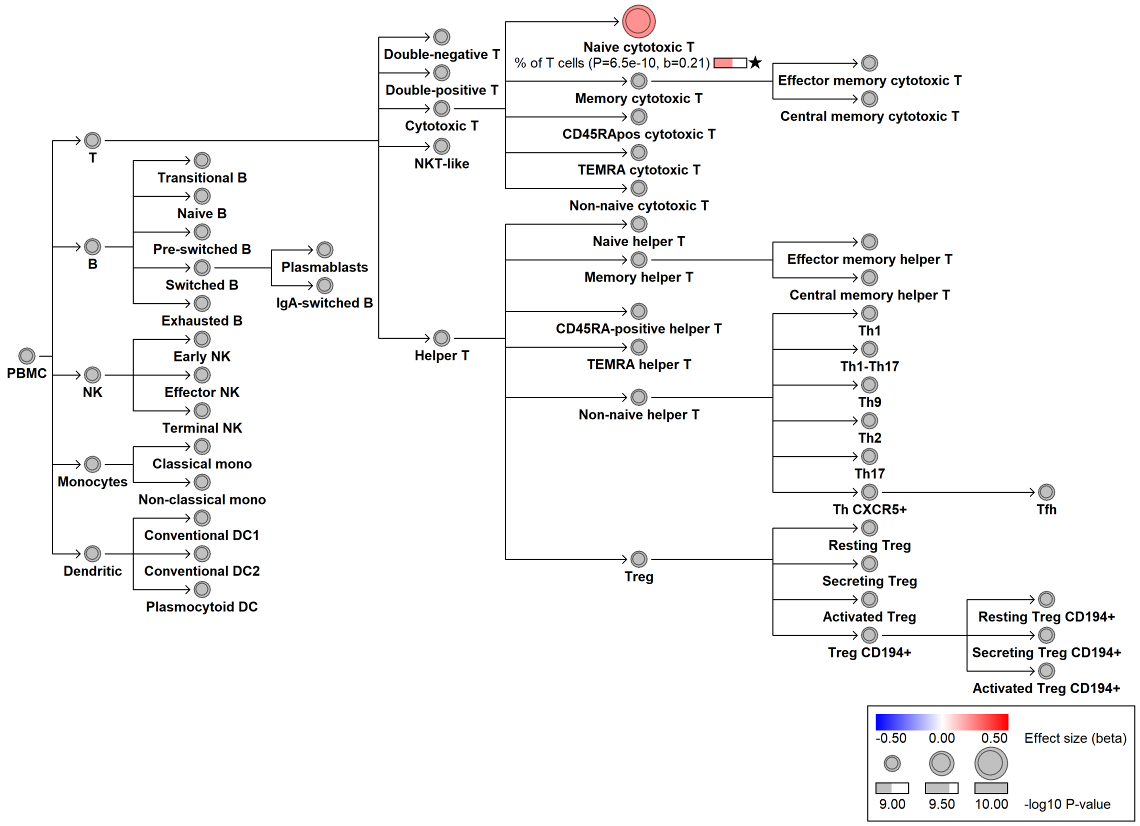

### chr6:31437843[A→G; EAF 24.824%] (rs3128981, cs_241)

Candidate genes: *HLA region*
Diseases associations: NA
Plasma pQTLs: NA

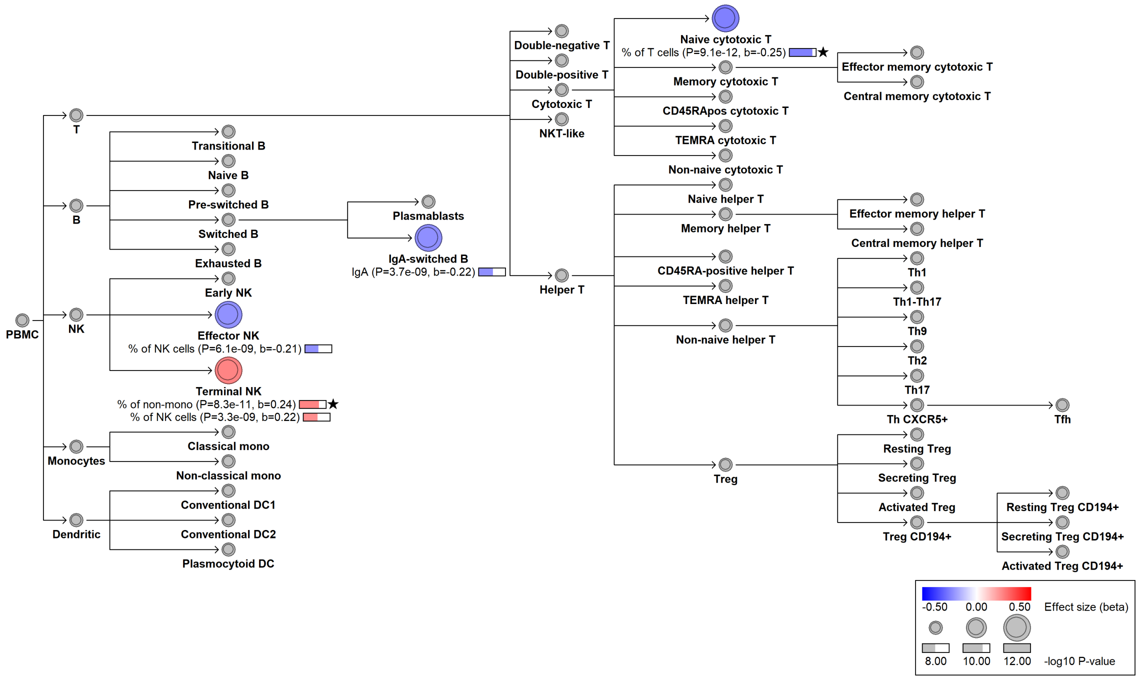

### chr6:32446112[A→C; EAF 44.348%] (rs9268668, cs_242)

Candidate genes: *HLA region*
Diseases associations: NA
Plasma pQTLs: NA

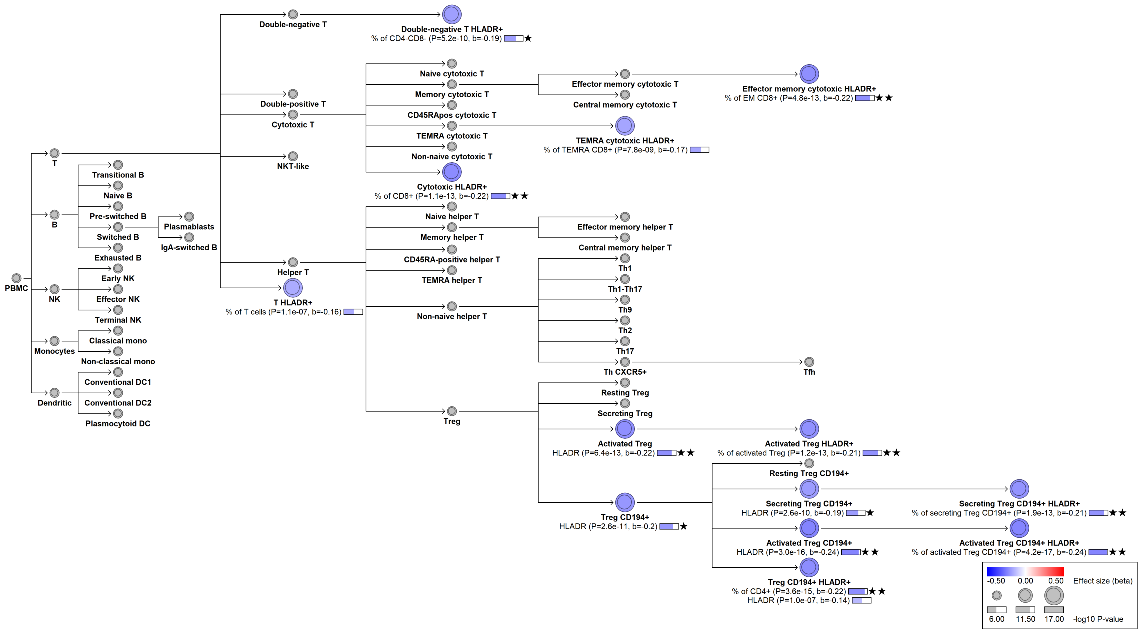

#### chr6:32447452[A→G; EAF 0.524%] (rs115406035, cs_243)

Candidate genes: *HLA region*
Diseases associations: NA
Plasma pQTLs: NA

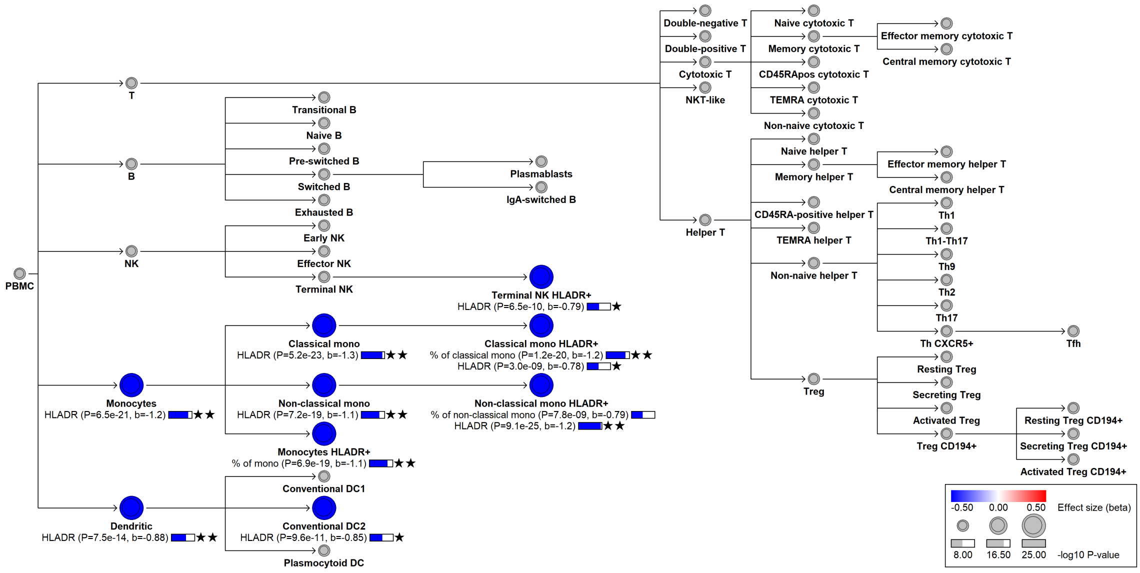

#### chr6:32535144[C→T; EAF 3.819%] (rs116033353, cs_244)

Candidate genes: *HLA region*
Diseases associations: NA
Plasma pQTLs: NA

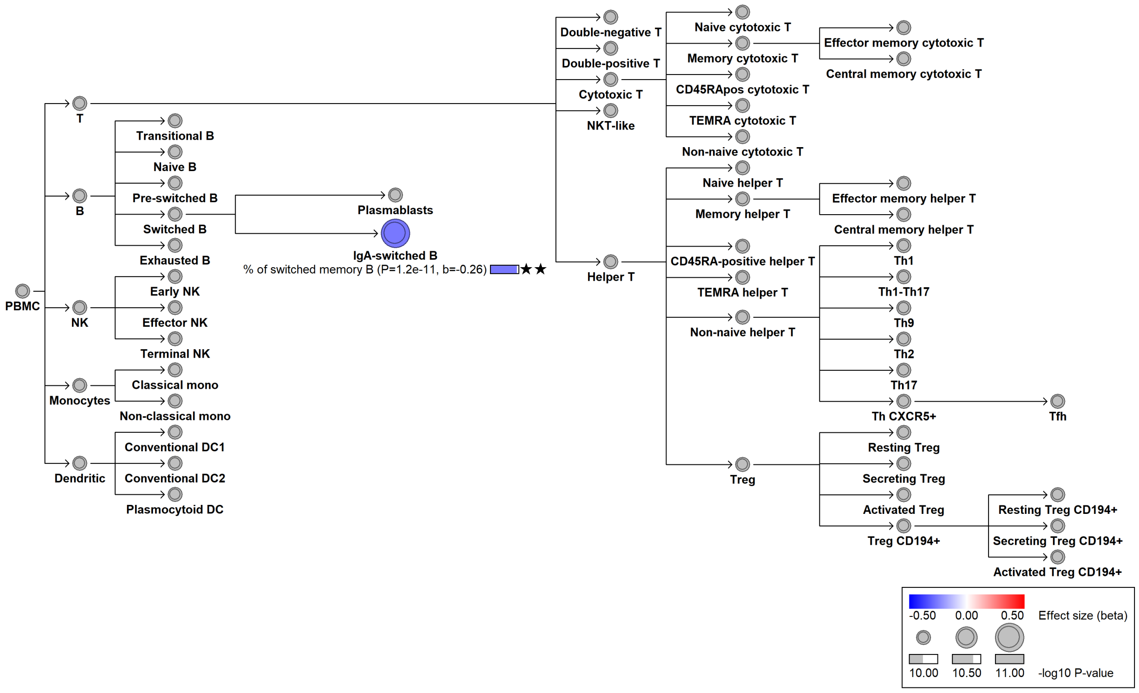

### chr6:32540133[!→AT; EAF 28.724%] (rs778011992, cs_245)

Candidate genes: *HLA region*
Diseases associations: NA
Plasma pQTLs: NA

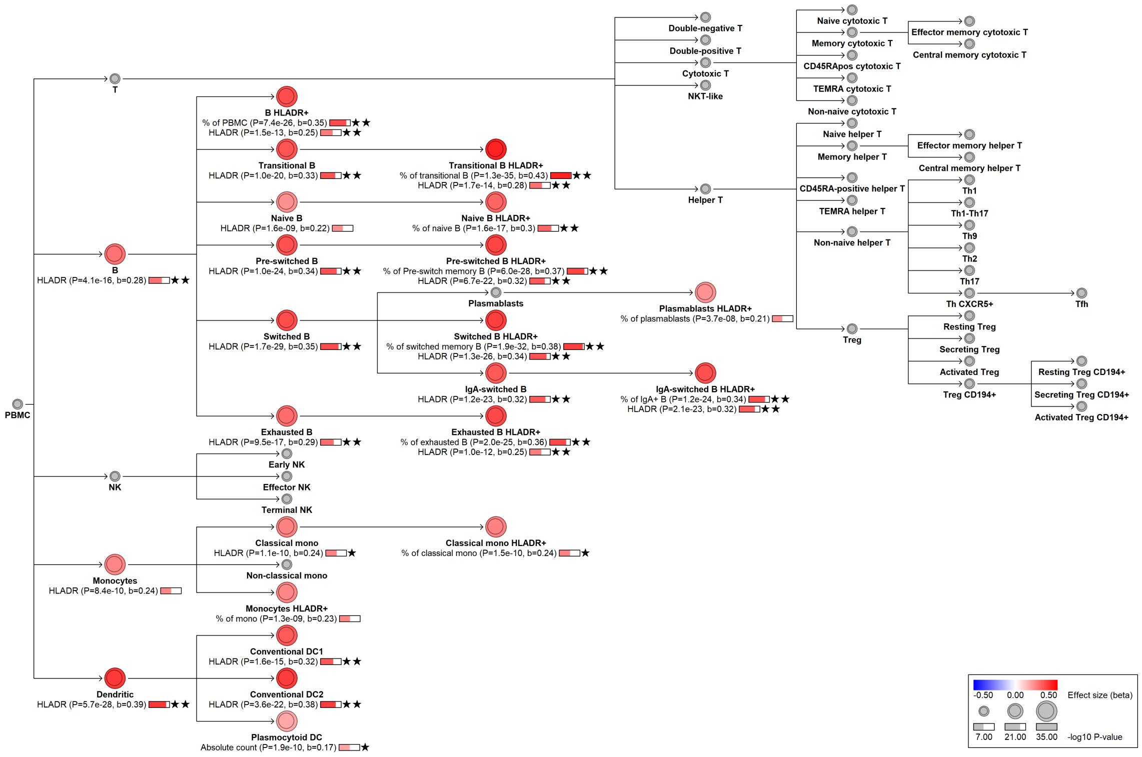

### chr6:32575470[G→T; EAF 1.5%] (rs369001642, cs_246)

Candidate genes: *HLA region*
Diseases associations: NA
Plasma pQTLs: NA

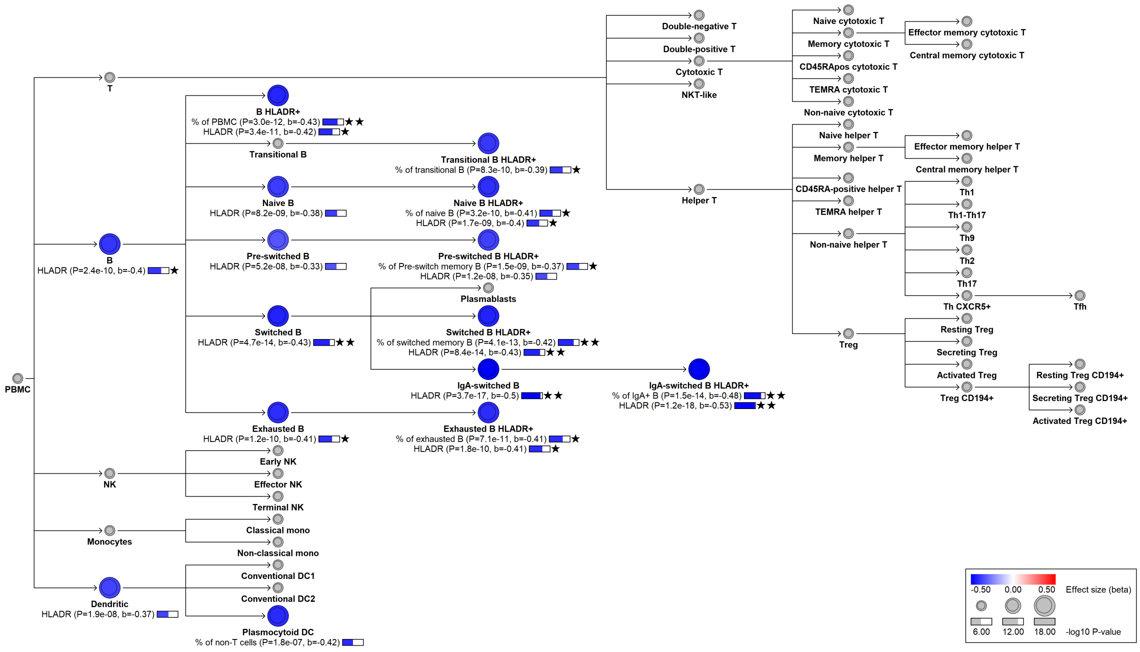

### chr6:32593523[C→T; EAF 49.206%] (rs9270585, cs_247)

Candidate genes: *HLA region*
Diseases associations: NA
Plasma pQTLs: NA

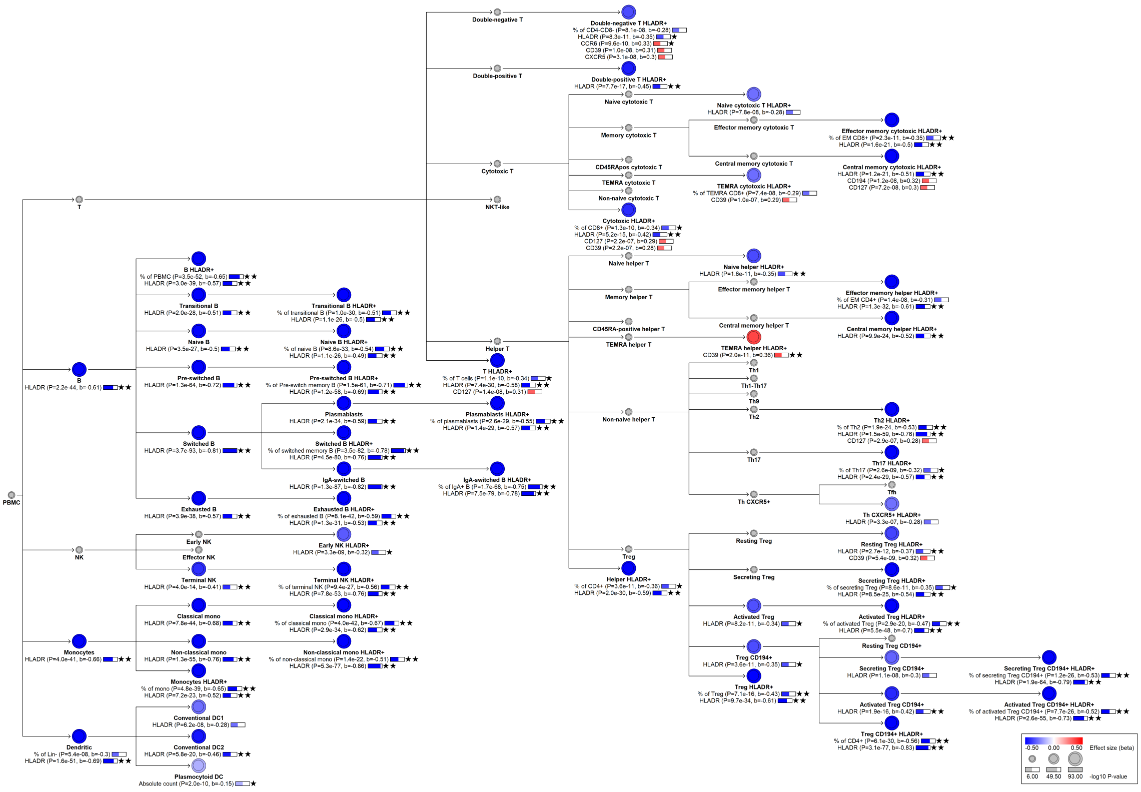

### chr6:32598244[T→G; EAF 33.453%] (rs9270657, cs_248)

Candidate genes: *HLA region*
Diseases associations: NA
Plasma pQTLs: NA

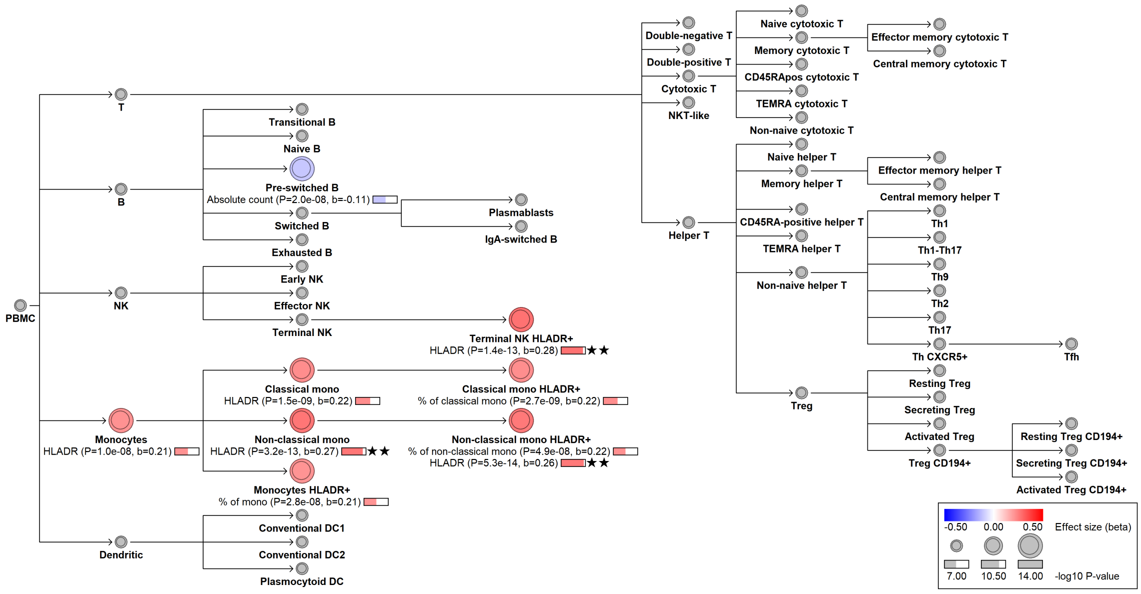

### chr6:32602293[CCAAA→C; EAF 38.098%] (rs113354454, cs_249)

Candidate genes: *HLA region*
Diseases associations: NA
Plasma pQTLs: NA

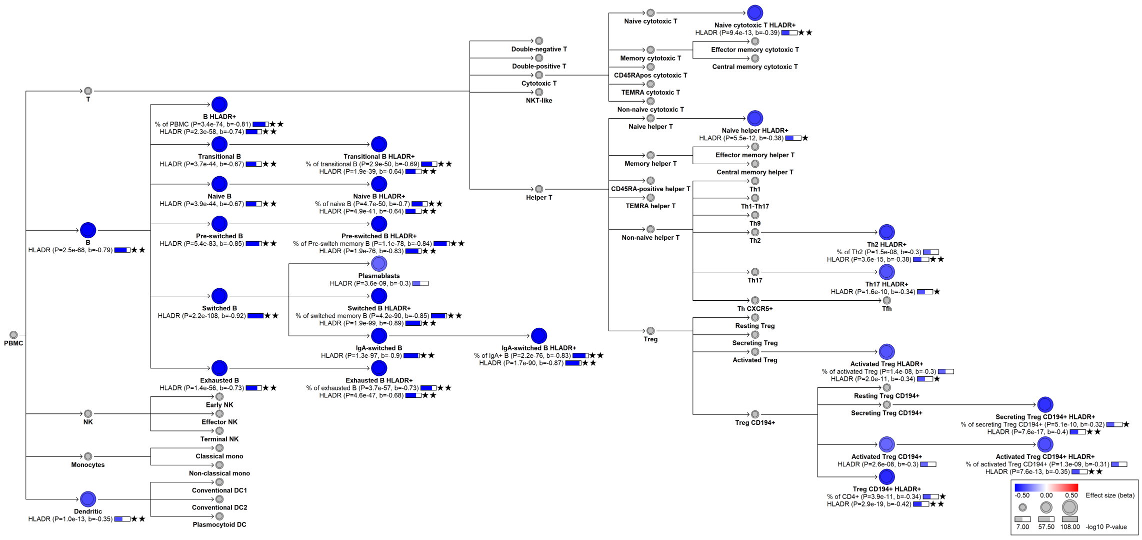

### chr6:32609608[C→T; EAF 18.725%] (rs9271147, cs_250)

Candidate genes: *HLA region*
Diseases associations: NA
Plasma pQTLs: NA

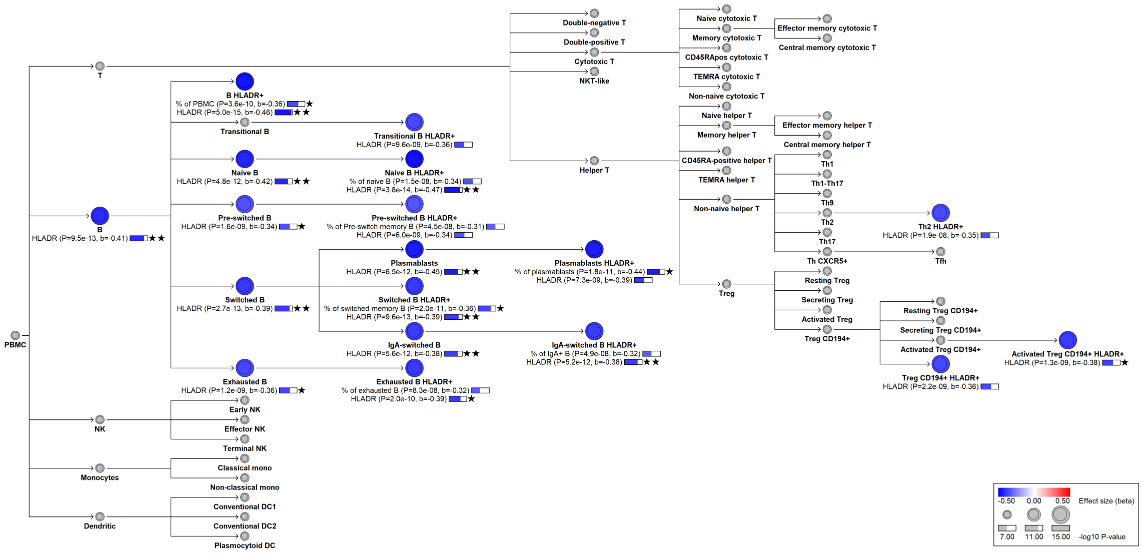

### chr6:32615269[A→G; EAF 47.444%] (rs3129754, cs_251)

Candidate genes: *HLA region*
Diseases associations: NA
Plasma pQTLs: NA

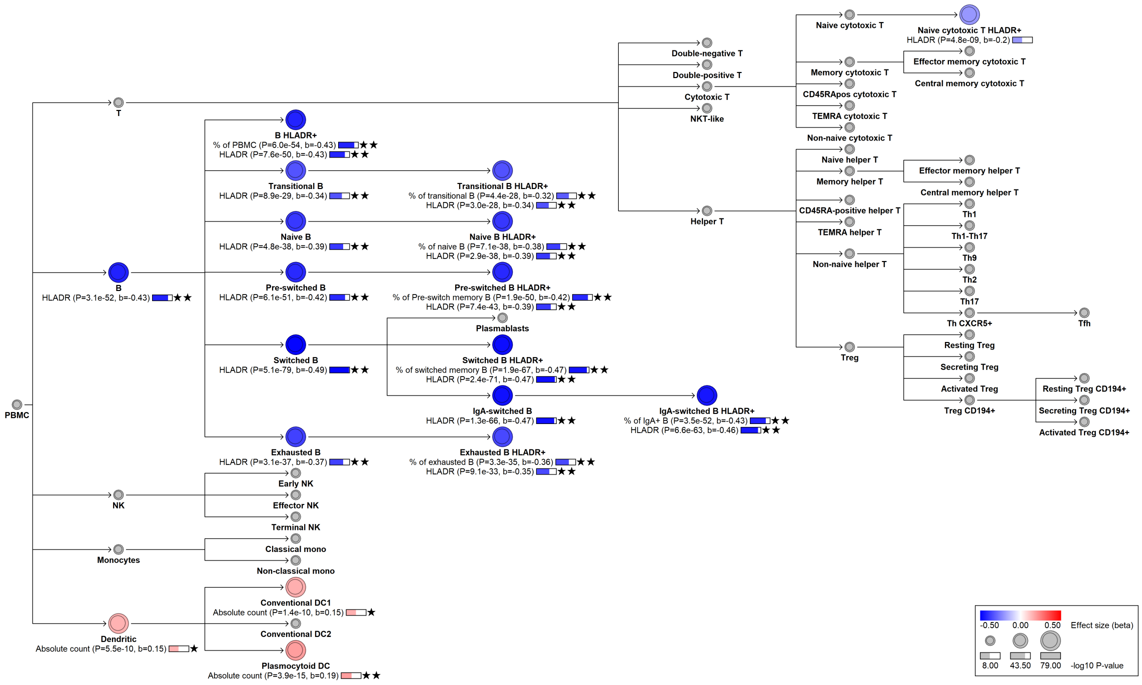

#### chr6:32632698[G→A; EAF 2.695%] (rs114883129, cs_252)

Candidate genes: *HLA region*
Diseases associations: NA
Plasma pQTLs: NA

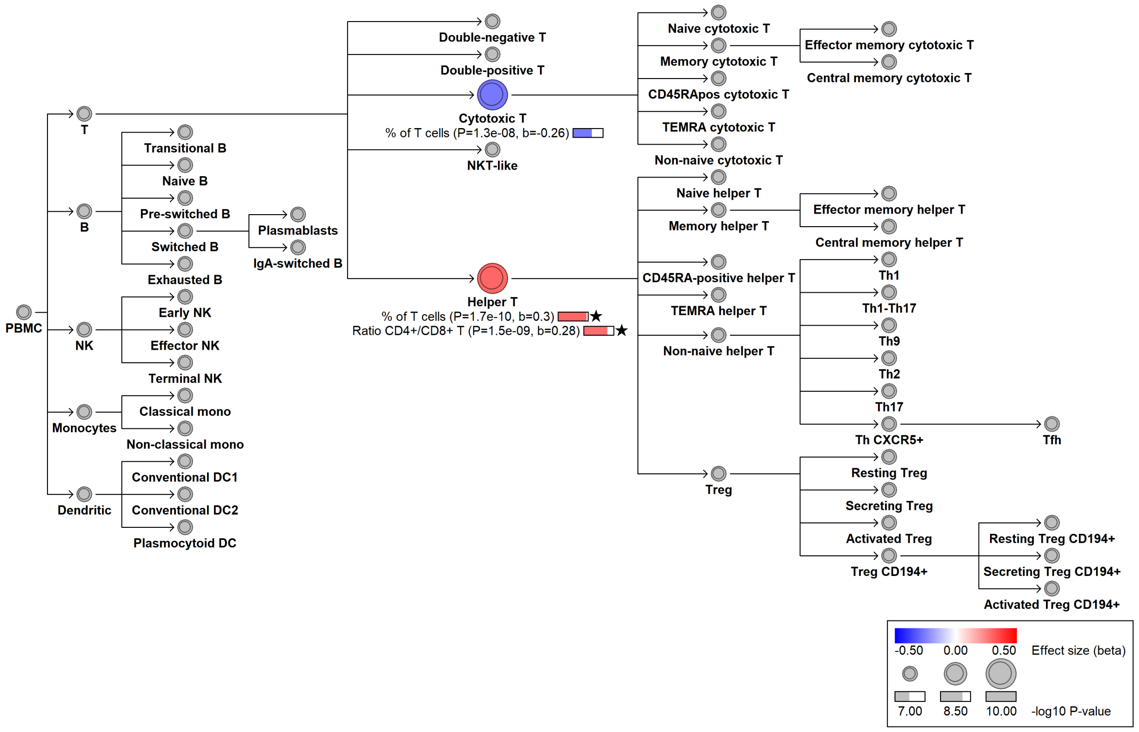

### chr6:32642790[A→T; EAF 7.596%] (rs28400569, cs_253)

Candidate genes: *HLA region*
Diseases associations: NA
Plasma pQTLs: NA

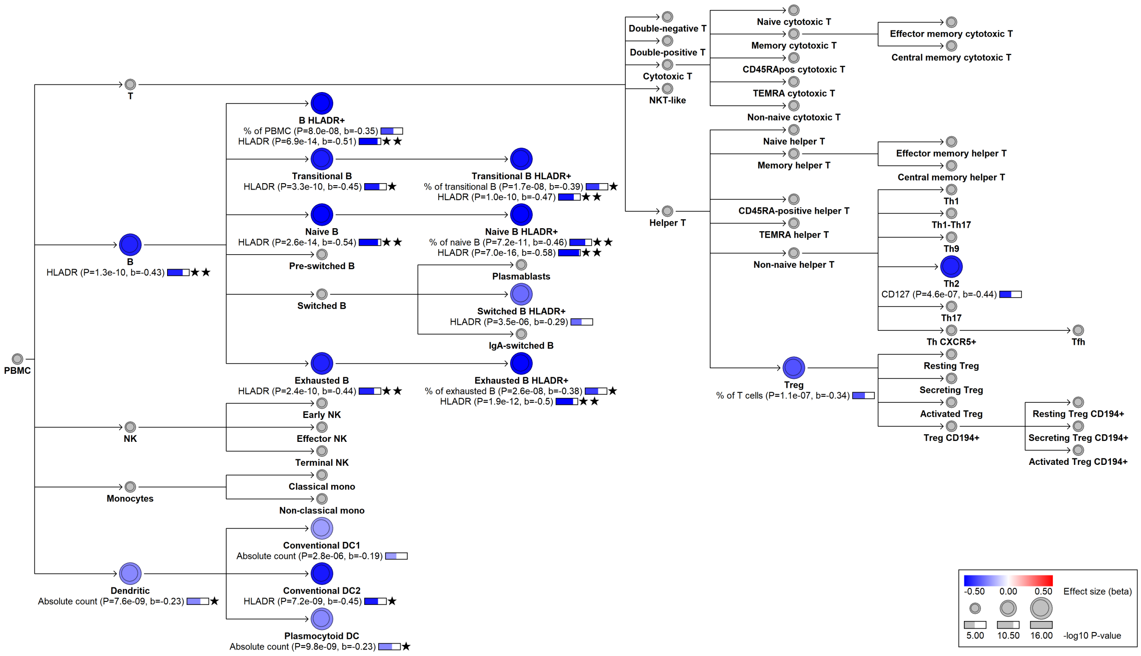

#### chr6:32659207[A→C; EAF 27.771%] (rs4713573, cs_254)

Candidate genes: *HLA region*
Diseases associations: Juvenile rheumatoid arthritis
Plasma pQTLs: NA

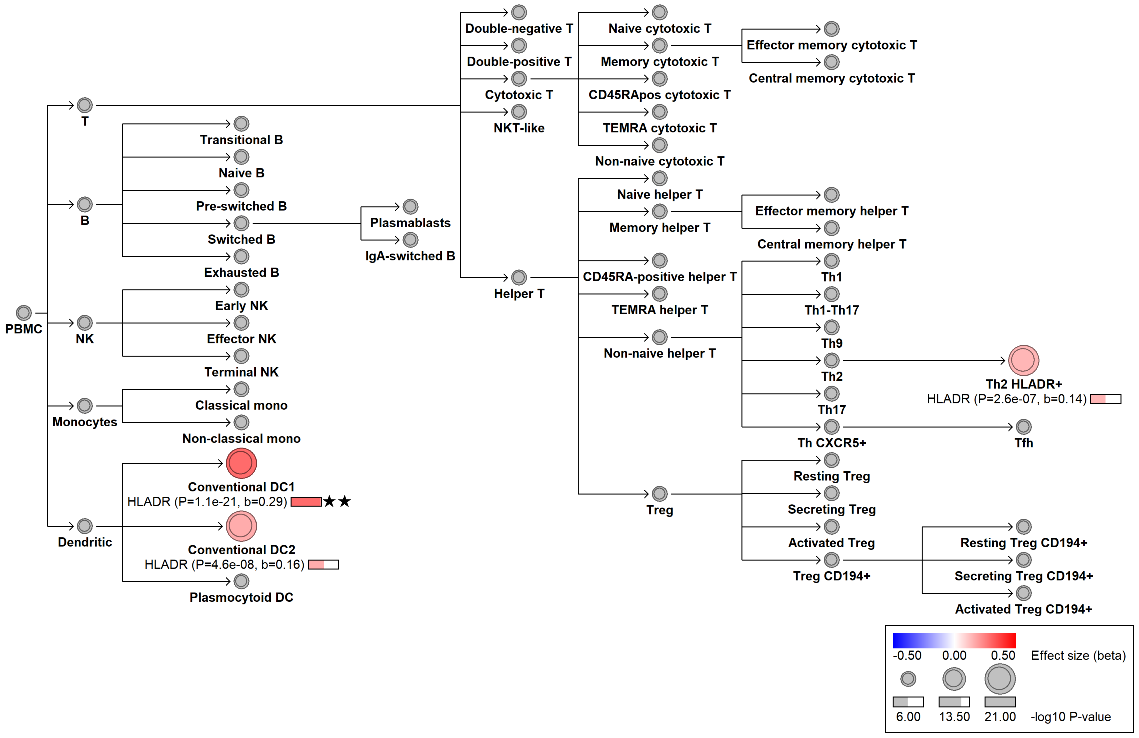

#### chr6:39316203[!→C; EAF 3.492%] (rs75212002, cs_57)

Candidate genes: *KCNK16*
Diseases associations: NA
Plasma pQTLs: NA

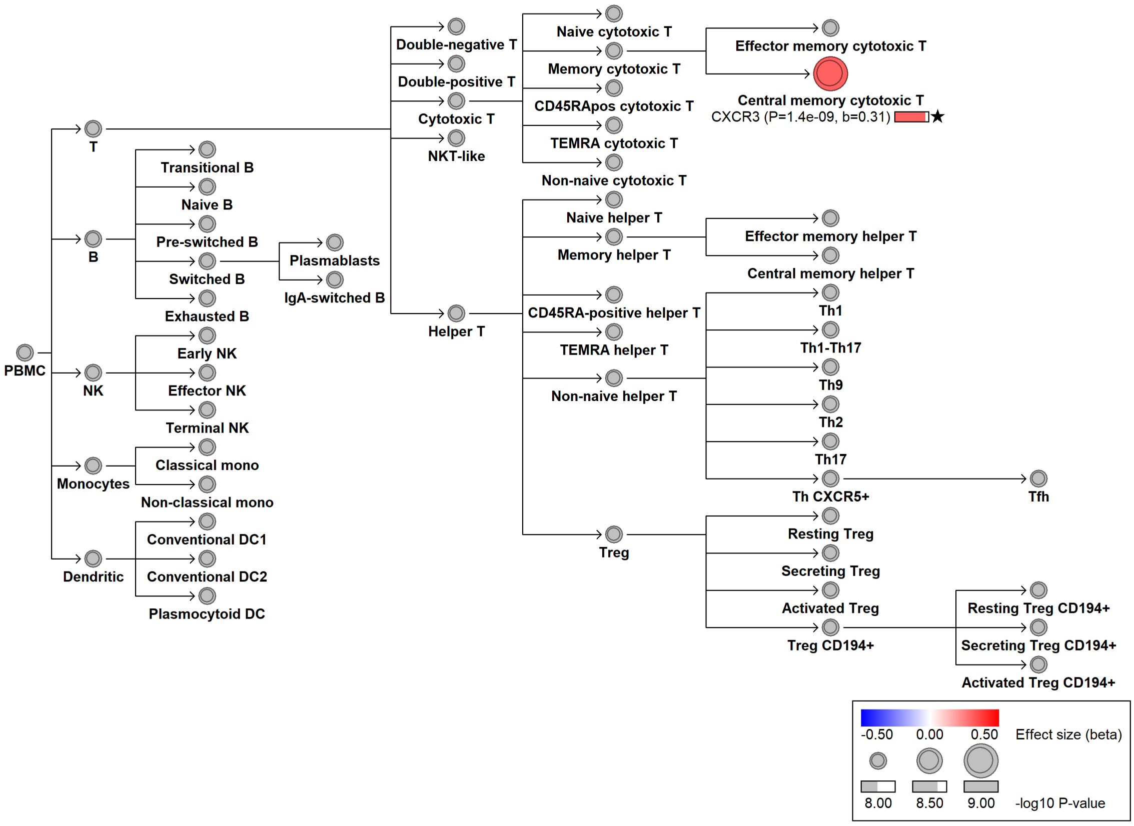

#### chr6:45422749[!→AGGCGGC; EAF 6.163%] (rs771859643, cs_10)

Candidate genes: *RUNX2*
Diseases associations: NA
Plasma pQTLs: NA

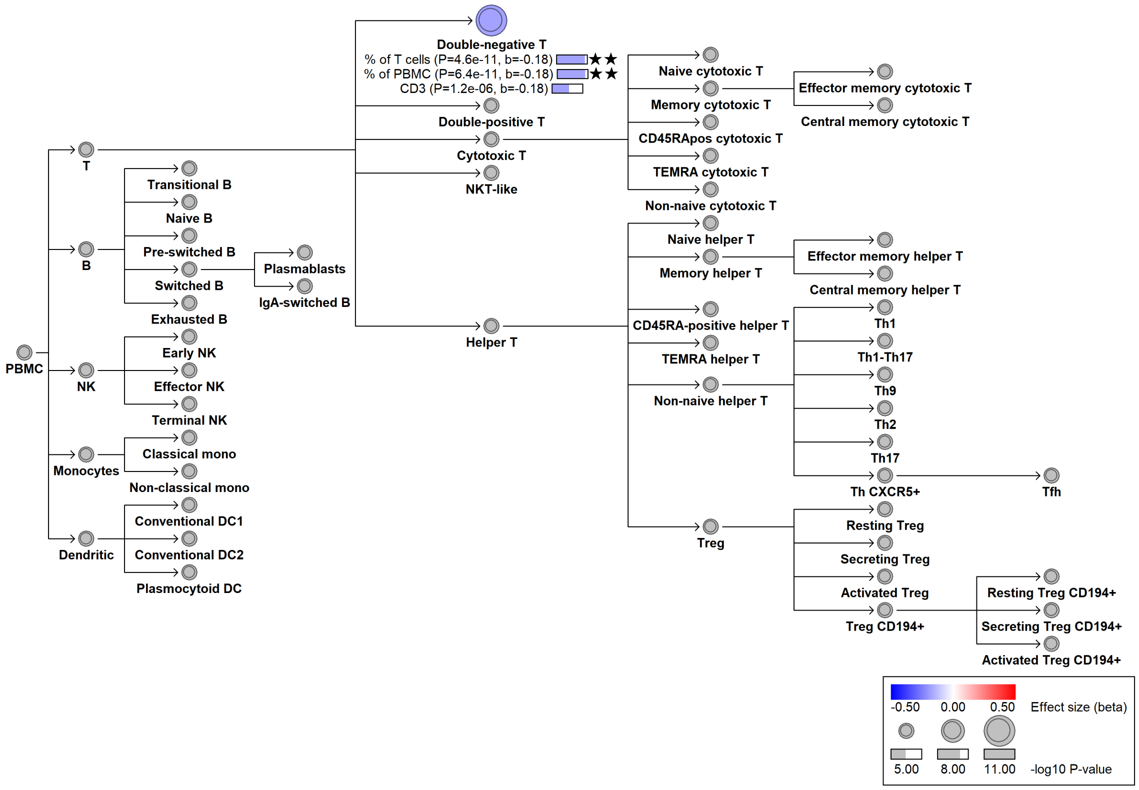

#### chr6:90267049[G→A; EAF 15.578%] (rs72928038, cs_206)

Candidate genes: *BACH2*
Diseases associations: Ankylosing spondylitis, Asthma, Autoimmune thyroiditis, Celiac disease, Crohn's disease, Malignant melanoma, Multiple sclerosis, Non-melanoma skin cancer, Primary sclerosing cholangitis, Psoriasis, Rheumatoid arthritis, Type 1 diabetes, Ulcerative colitis, Vitiligo
Plasma pQTLs: NA

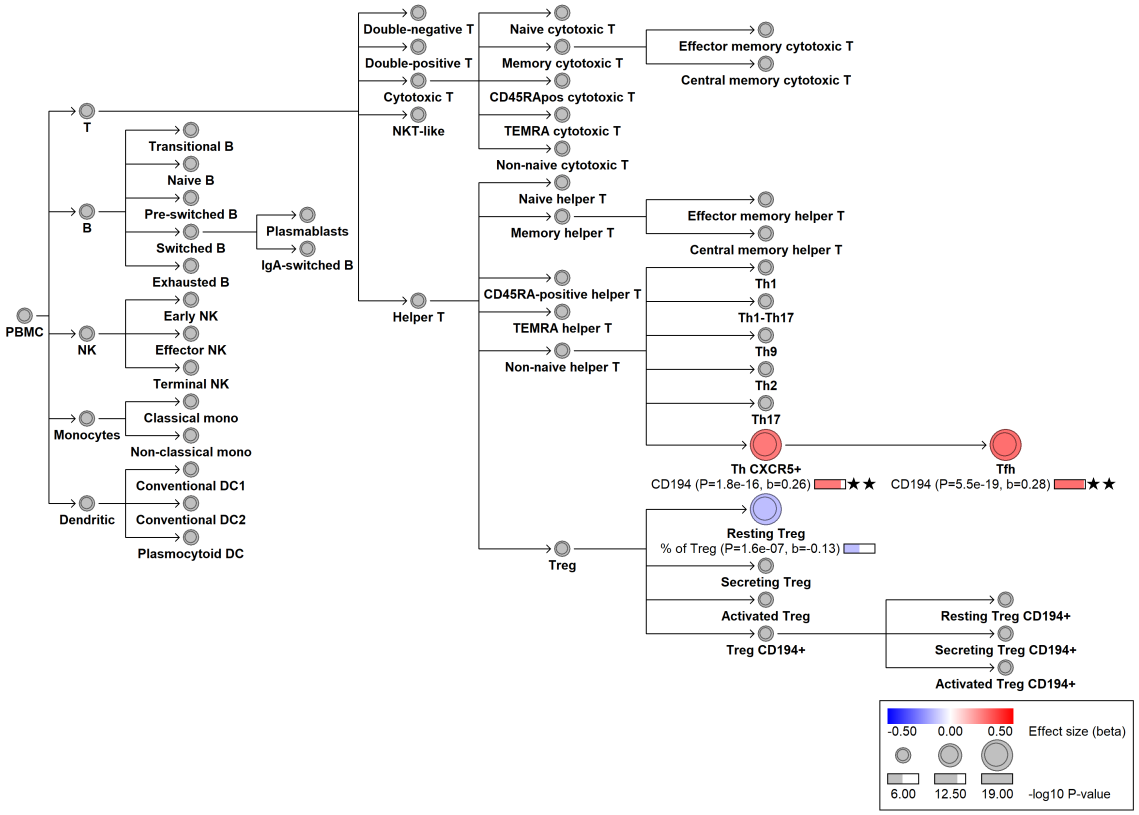

#### chr6:90271938[C→T; EAF 31.239%] (rs45553631, cs_191)

Candidate genes: *BACH2*
Diseases associations: Allergy, Ankylosing spondylitis, Asthma, Chronic obstructive pulmonary disease, Crohn's disease, Eczema, Inflammatory bowel disease, Nasal polyps, Primary sclerosing cholangitis, Psoriasis, Ulcerative colitis
Plasma pQTLs: ANXA1, CA9, CCL17, CCL22, CD207, CRTAP, Composite immunoglobulin trait IgA/IgM, Composite immunoglobulin trait IgM/(IgA*IgG), FCGR1A, FLT3, FLT3LG, GRN, MAPRE3, SPINK2, TGFB2, TMEM70, ZNRD2

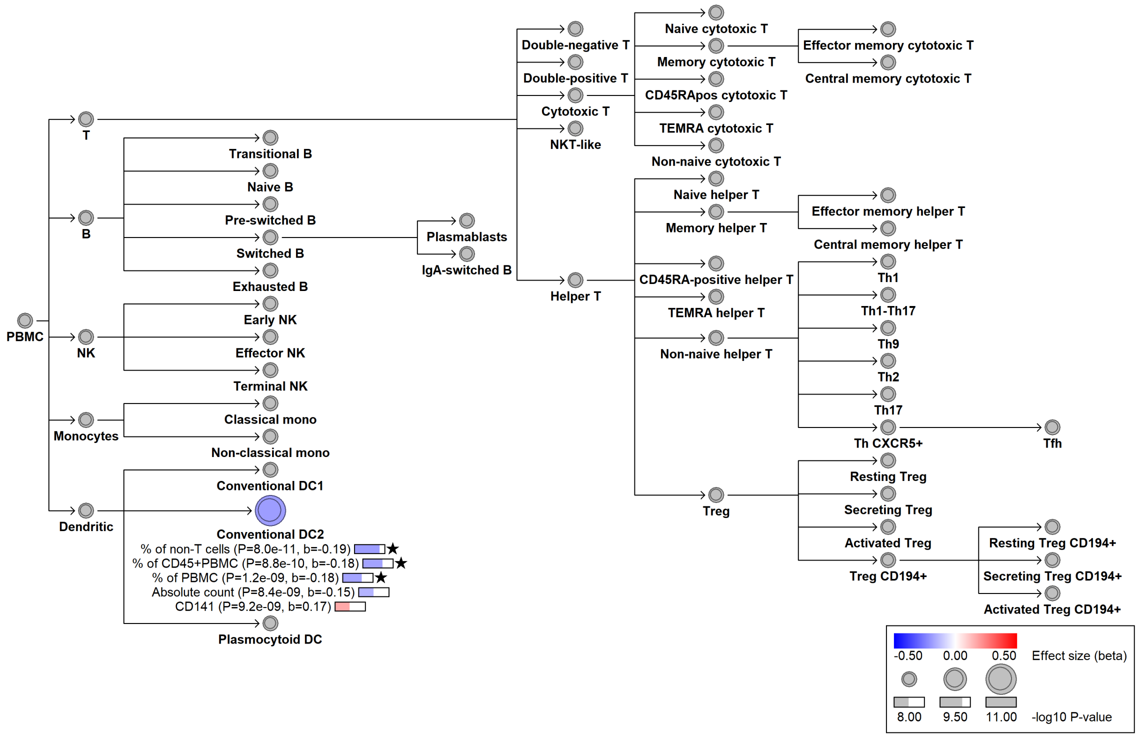

#### chr6:90279522[!→C; EAF 43.087%] (rs10637718, cs_68)

Candidate genes: *BACH2*
Diseases associations: Autoimmune thyroiditis, Systemic lupus erythematosus
Plasma pQTLs: NA

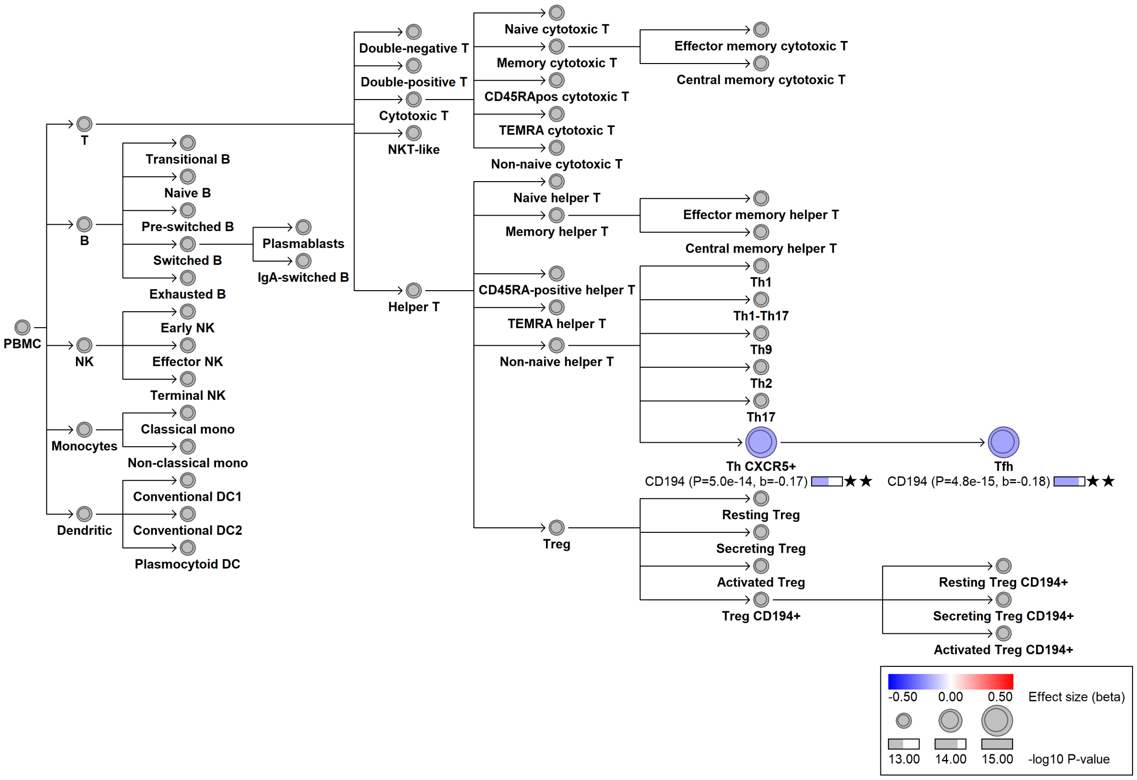

#### chr6:106686130[!→TAC; EAF 6.068%] (rs144171105, cs_58)

Candidate genes: *CD24*
Diseases associations: NA
Plasma pQTLs: NA

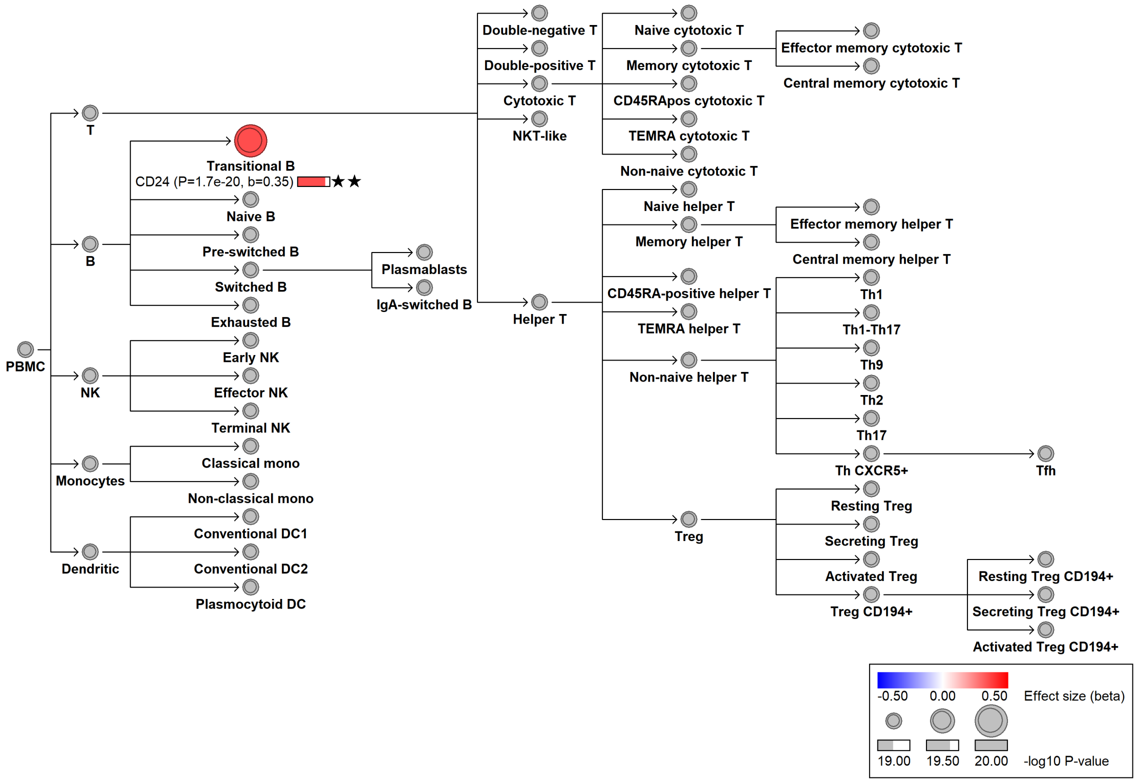

#### chr6:106716201[A→G; EAF 6.373%] (rs9400058, cs_155)

Candidate genes: *CD24*
Diseases associations: NA
Plasma pQTLs: NA

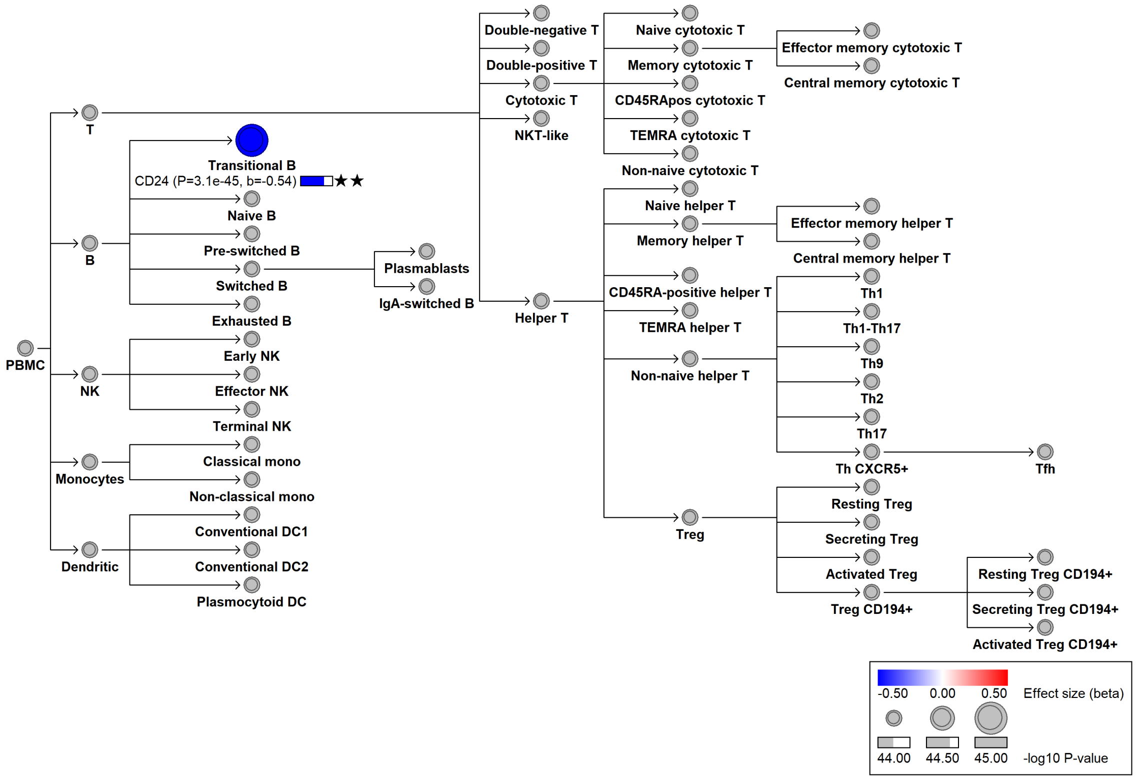

#### chr6:109002316[G→T; EAF 12.144%] (rs2273668, cs_134)

Candidate genes: *SESN1*
Diseases associations: Asthma, Autoimmune thyroiditis, Prostate cancer, Uterine leiomyomata
Plasma pQTLs: NA

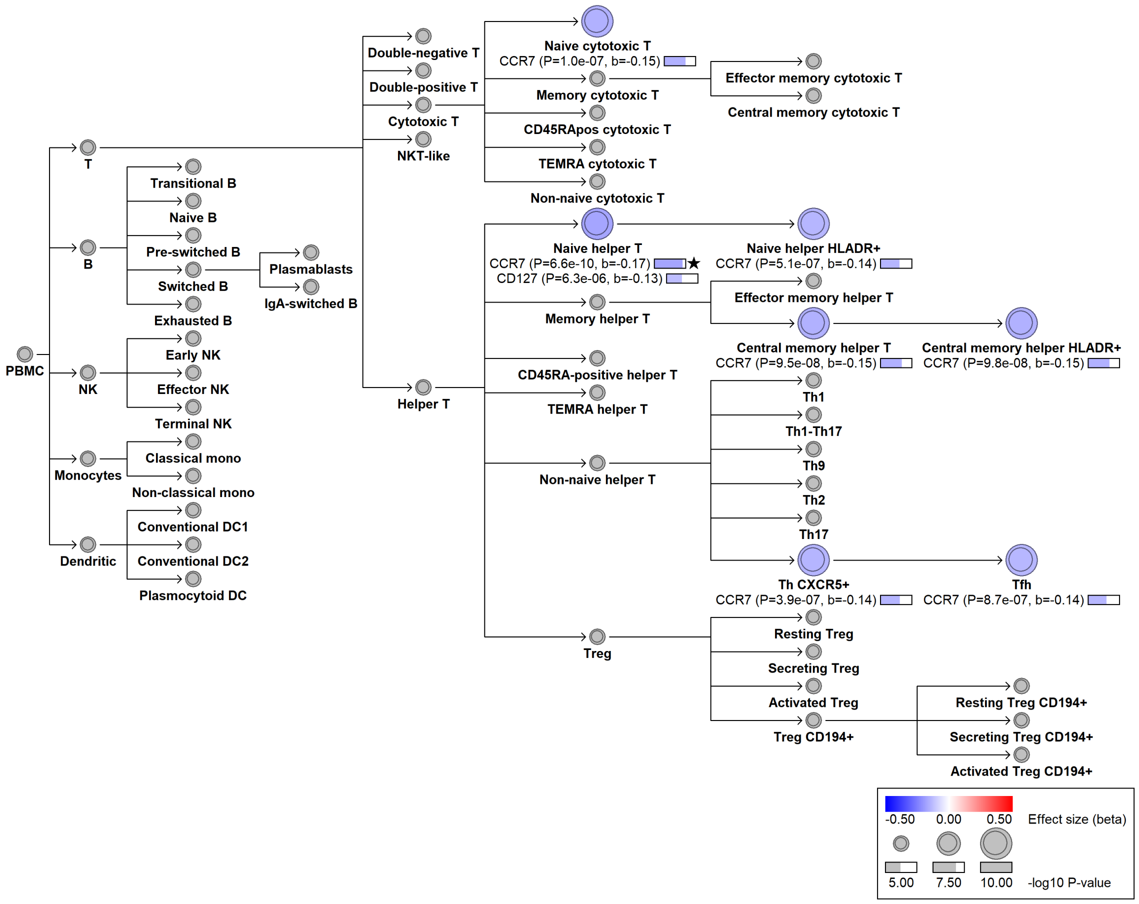

#### chr6:109304676[G→A; EAF 44.521%] (rs1546723, cs_197)

Candidate genes: *CD164*
Diseases associations: NA
Plasma pQTLs: EPO, FLT3LG, IL15, LRRN1, MMP1

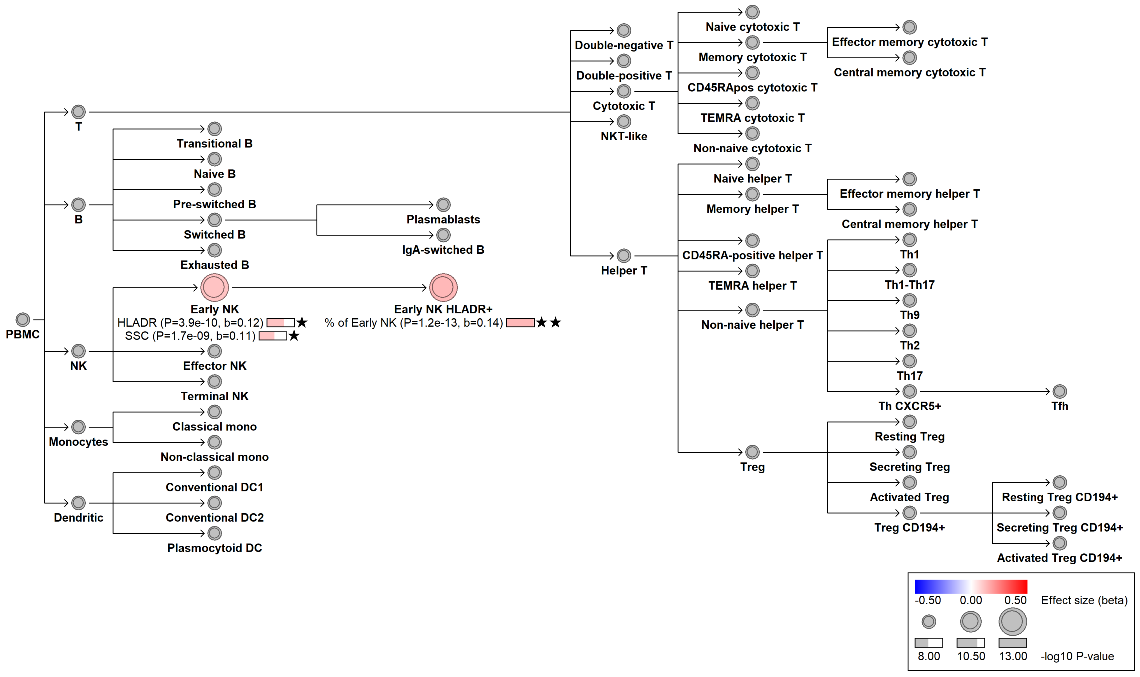

#### chr6:166946901[T→C; EAF 41.357%] (rs2236313, cs_13)

Candidate genes: *CCR6*
Diseases associations: Vitiligo
Plasma pQTLs: NA

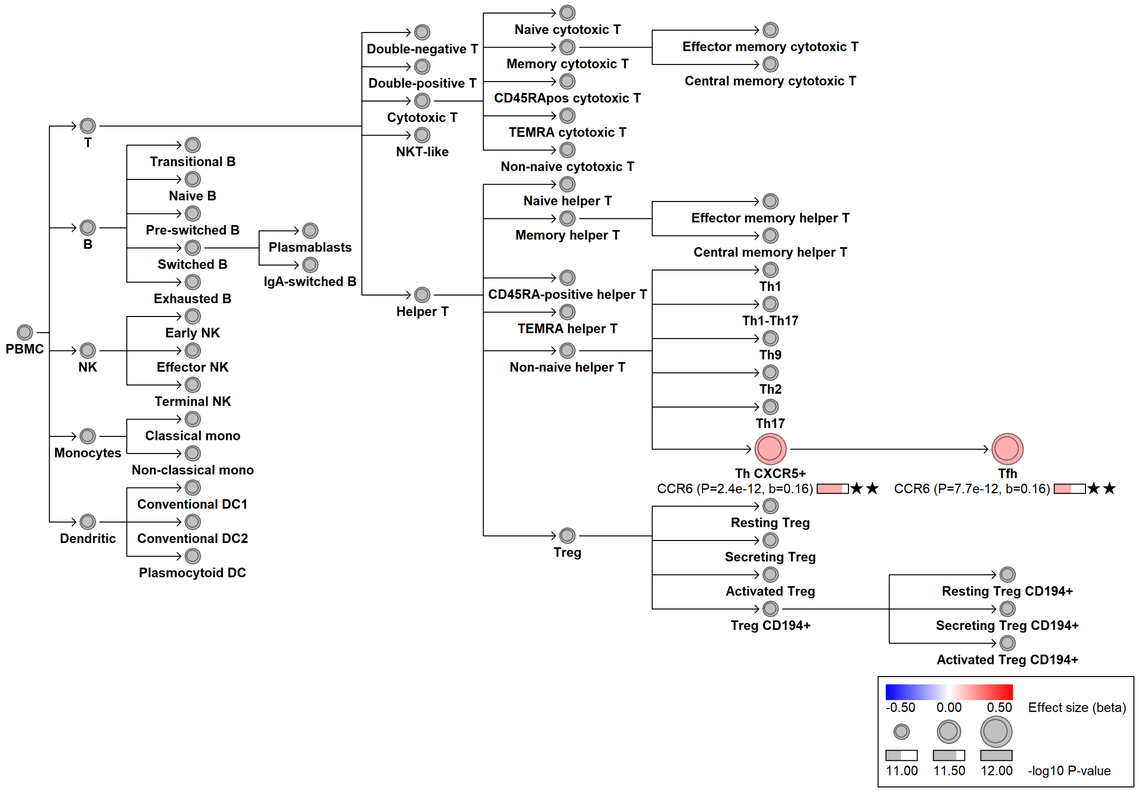

#### chr6:167041651[A→G; EAF 28.435%] (rs162297, cs_204)

Candidate genes: *CCR6*
Diseases associations: NA
Plasma pQTLs: NA

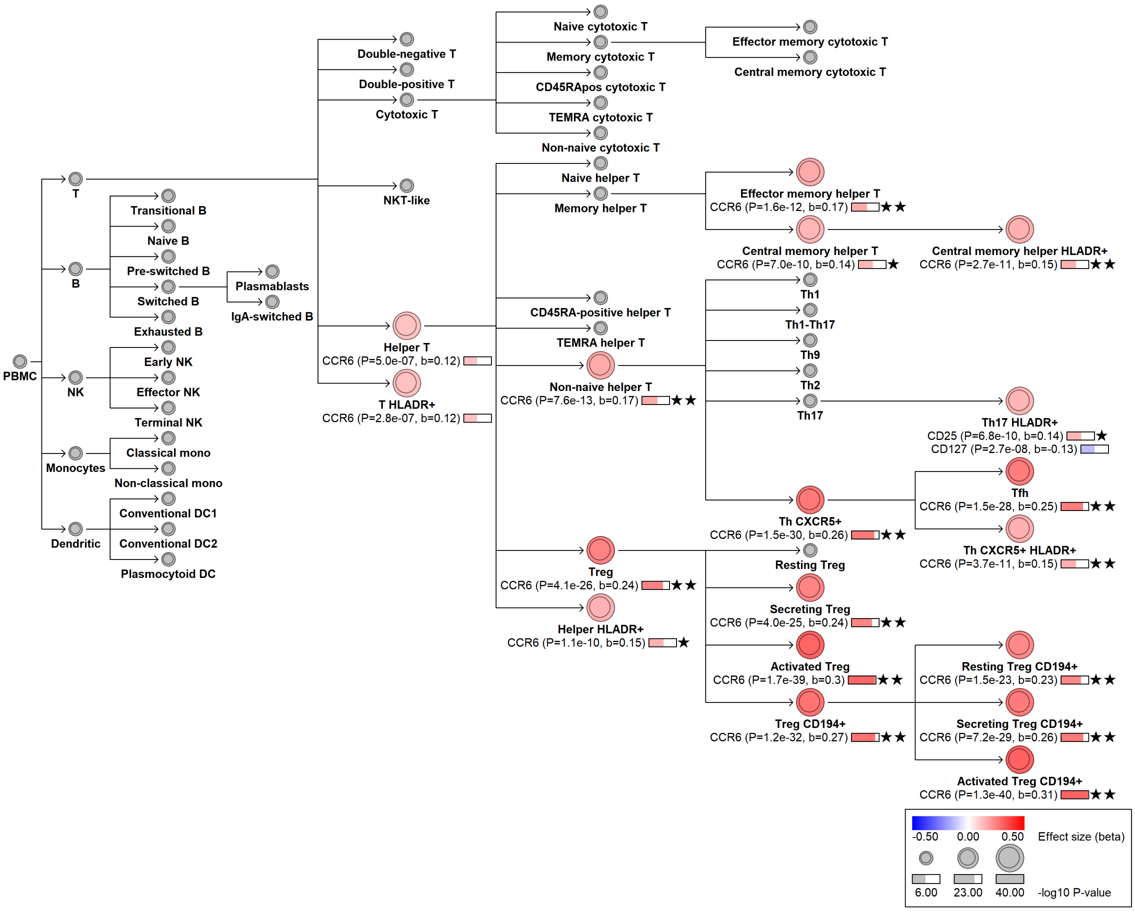

### chr6:167119243[G→A; EAF 44.868%] (rs3093025, cs_8)

Candidate genes: *CCR6*
Diseases associations: Rheumatoid arthritis
Plasma pQTLs: NA

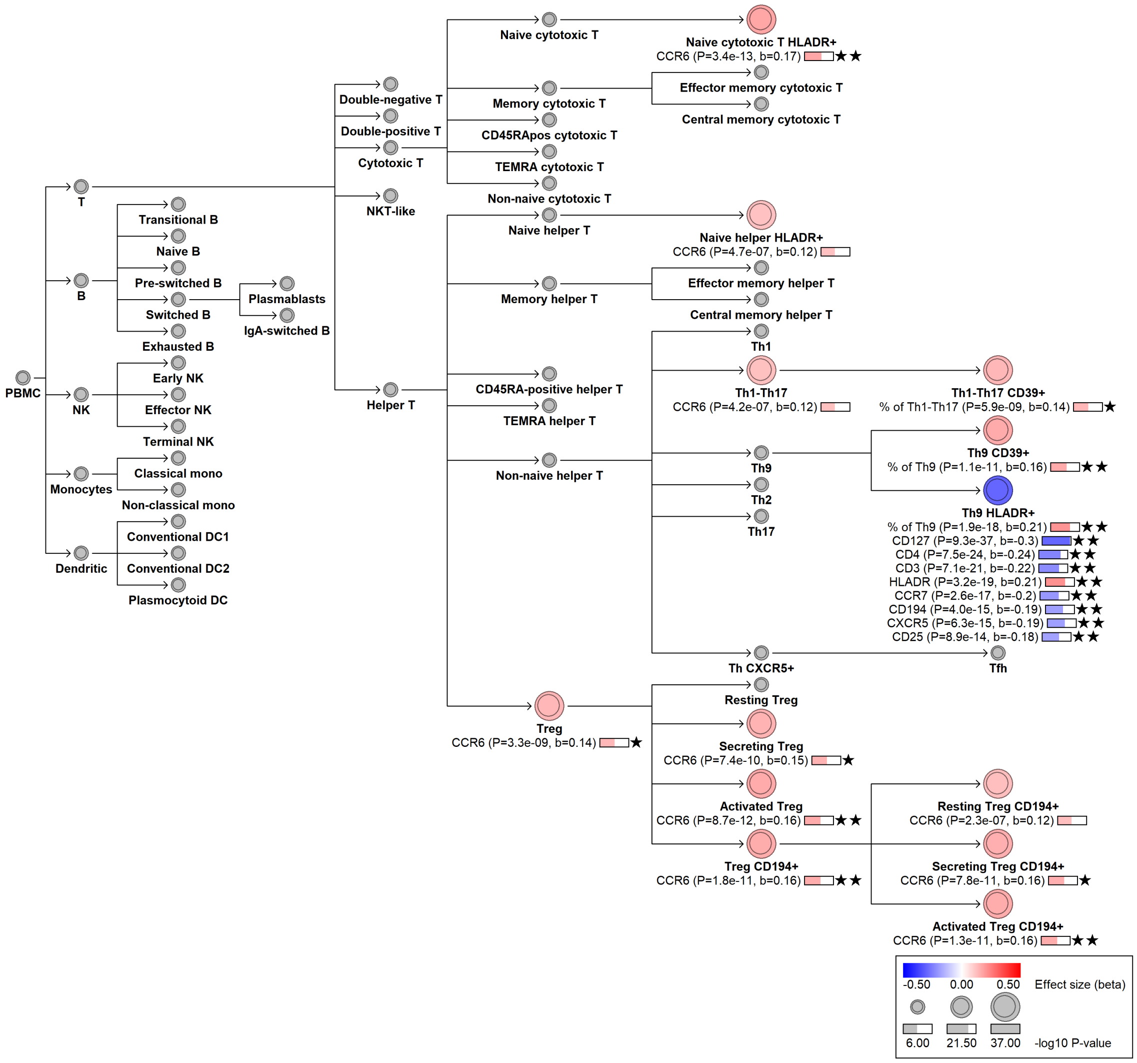

#### chr7:2256917[A→G; EAF 0.517%] (rs144787122, cs_62)

Candidate genes: *SNX8*
Diseases associations: NA
Plasma pQTLs: CD248, FAP, Immunoglobulin G, NOTCH1, OMD

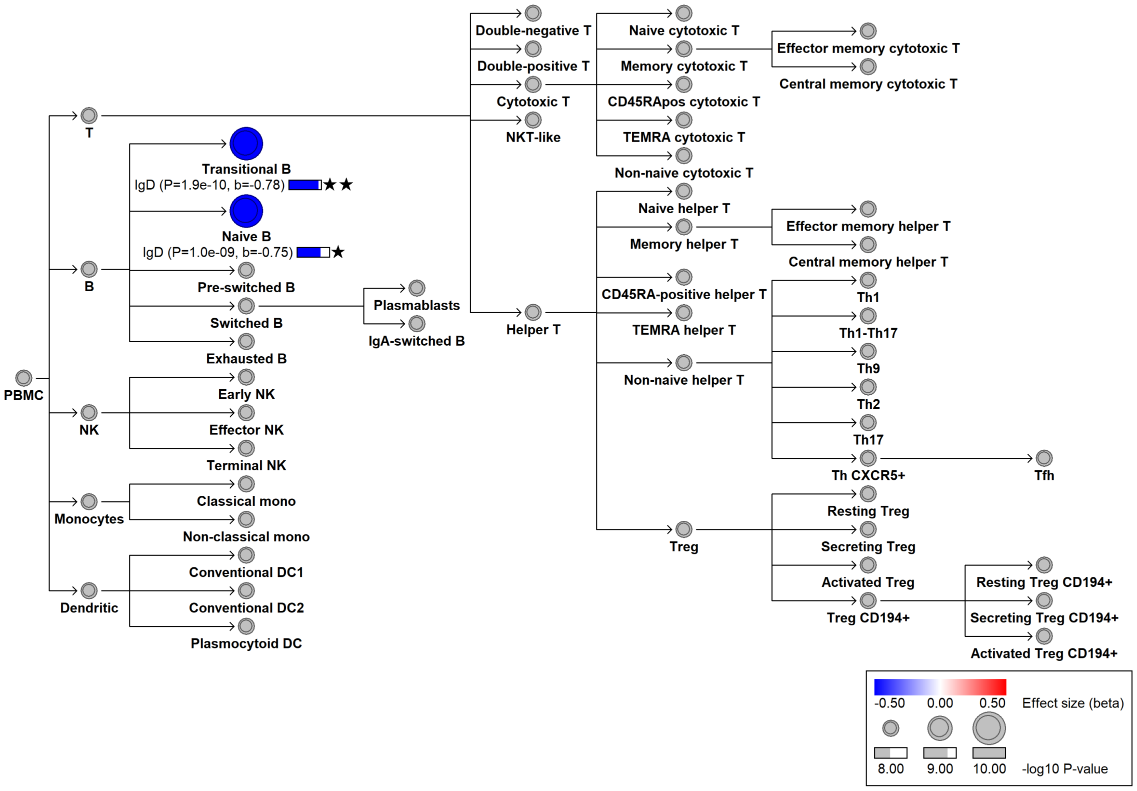

### chr7:2316403[A→C; EAF 26.526%] (rs35642219, cs_34)

Candidate genes: *SNX8*
Diseases associations: NA
Plasma pQTLs: NA

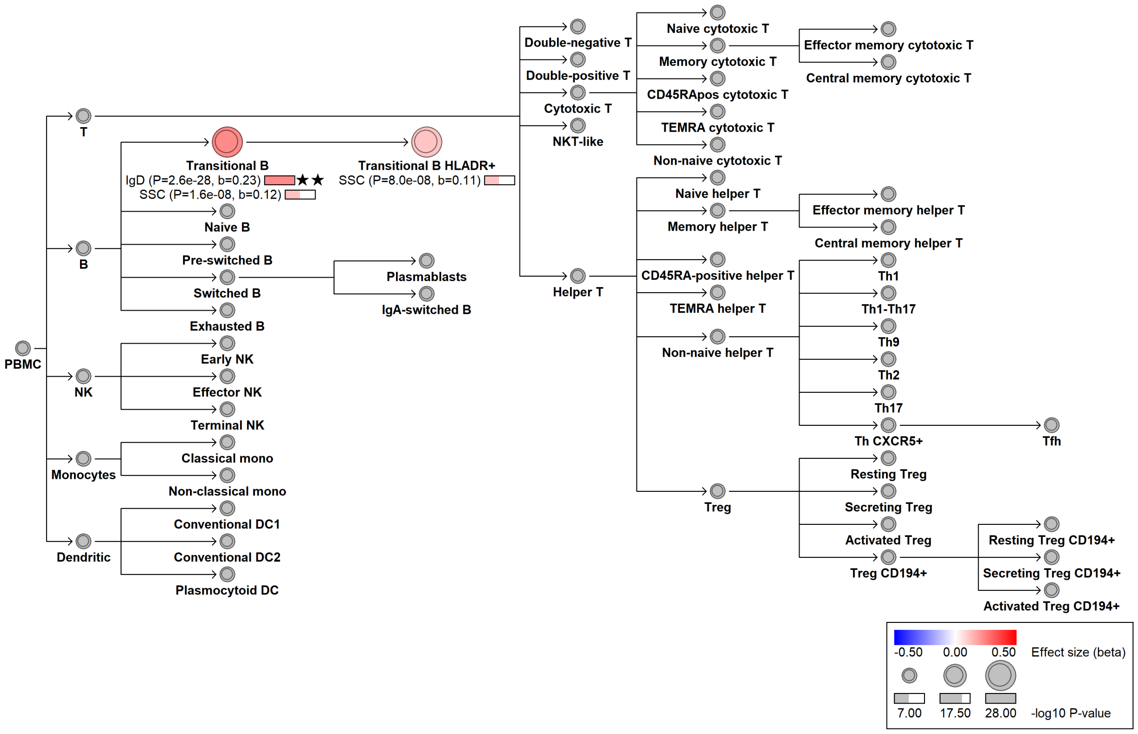

#### chr7:23583181[TTATA→!; EAF 25.149%] (chr7:23583181, cs_14)

Candidate genes: *CCDC126*
Diseases associations: NA
Plasma pQTLs: CCDC126, CR2

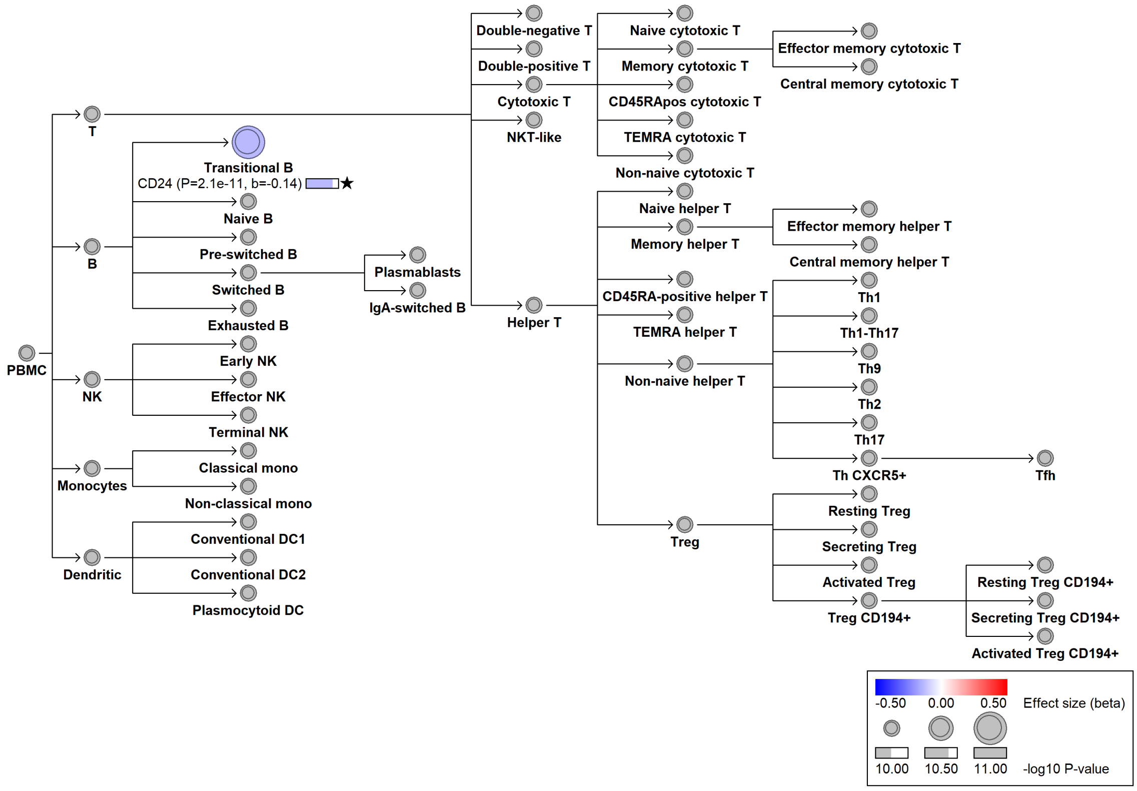

#### chr7:28137682[C→T; EAF 22.259%] (rs11495981, cs_181)

Candidate genes: *JAZF1*
Diseases associations: Allergy, Alzheimer's disease, Asthma, Crohn's disease, Eczema, Multiple sclerosis, Rheumatoid arthritis
Plasma pQTLs: CPM, CTSF

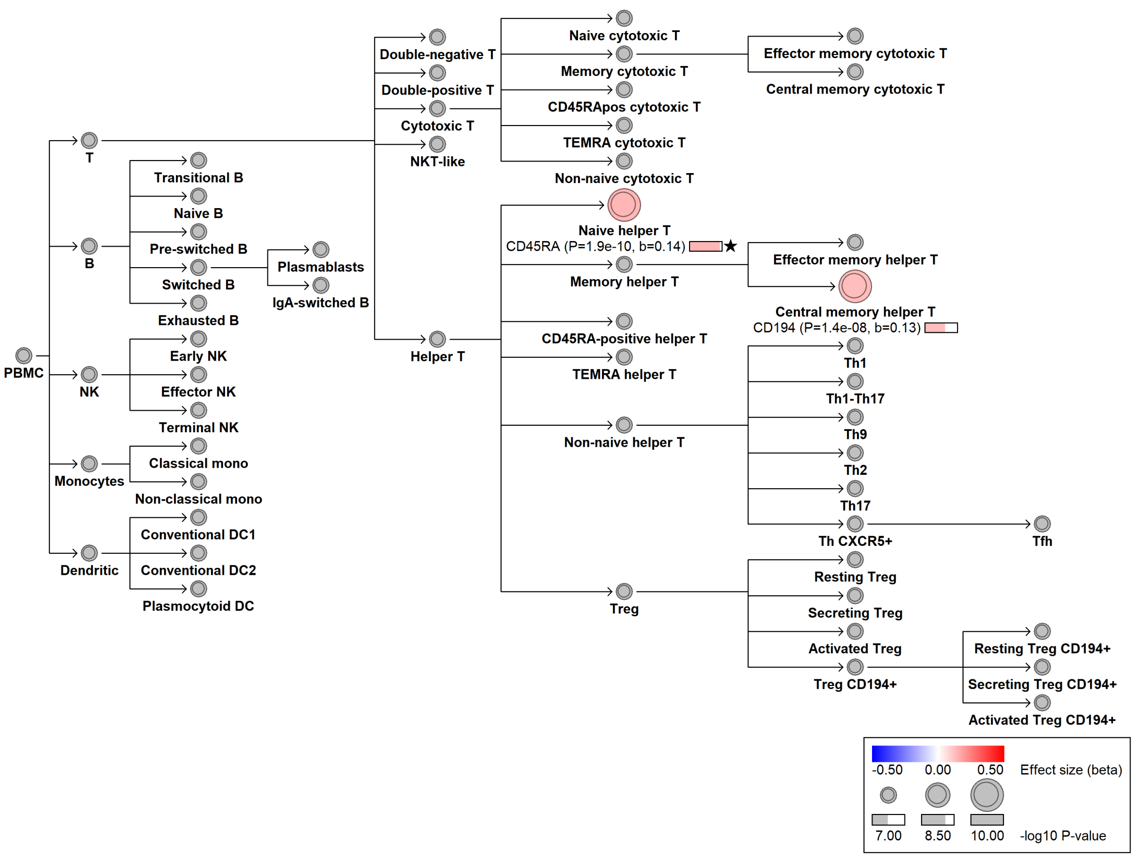

#### chr7:38233206[!→CTCCACCACCACCACTG; EAF 12.361%] (rs200632242, cs_135)

Candidate genes: *TRG region*
Diseases associations: NA
Plasma pQTLs: NA

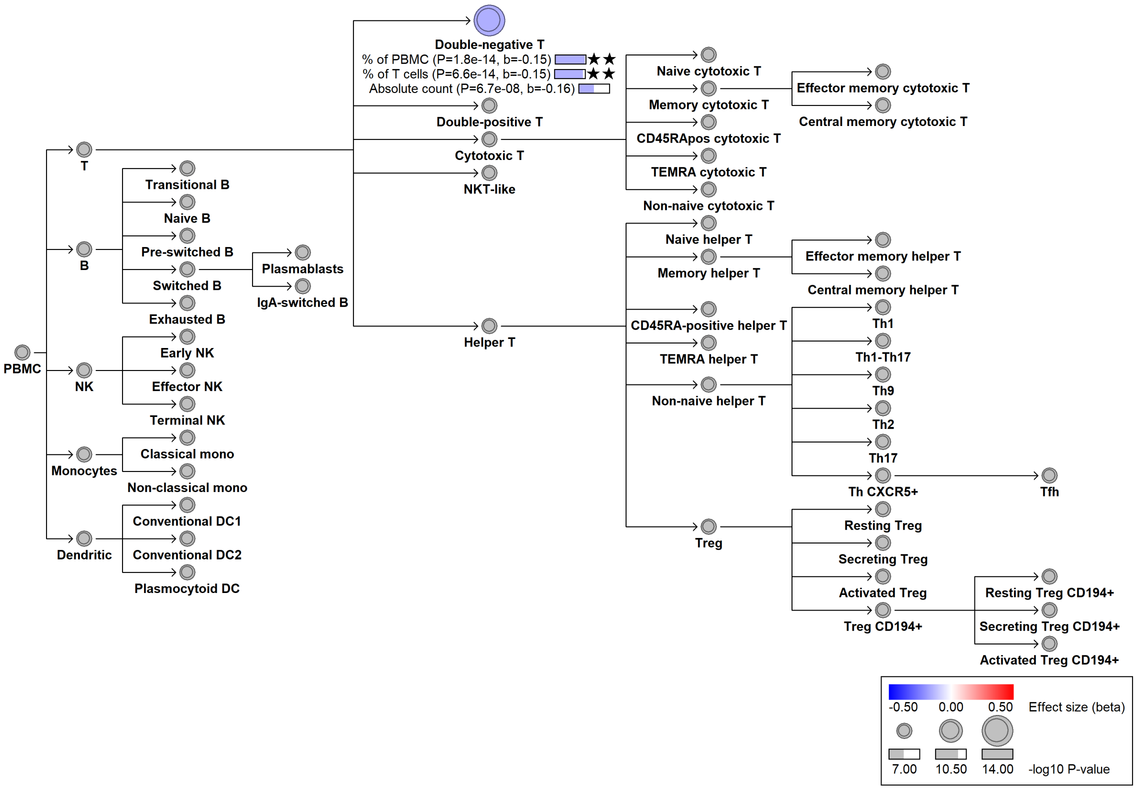

#### chr7:39979216[C→!; EAF 35.851%] (chr7:39979216, cs_231)

Candidate genes: *CDK13*
Diseases associations: NA
Plasma pQTLs: NA

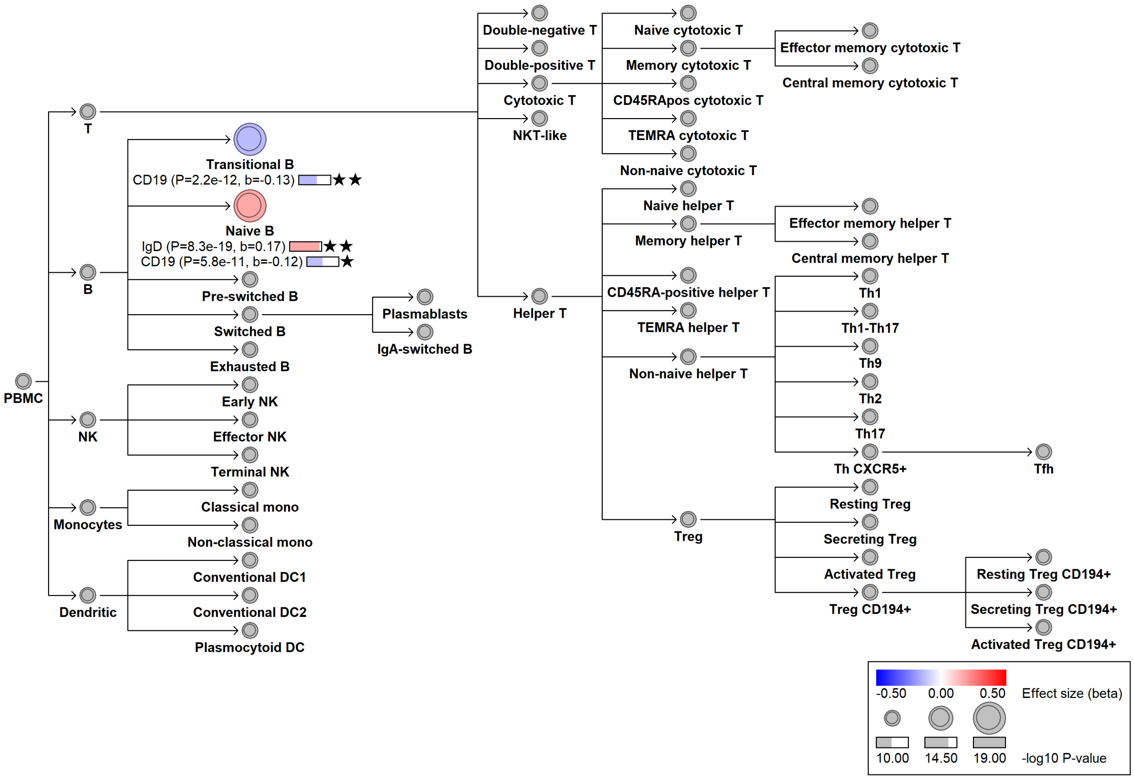

### chr7:49962440[C→T; EAF 2.939%] (rs139839446, cs_145)

Candidate genes: *IKZF1, ZPBP*
Diseases associations: NA
Plasma pQTLs: NA

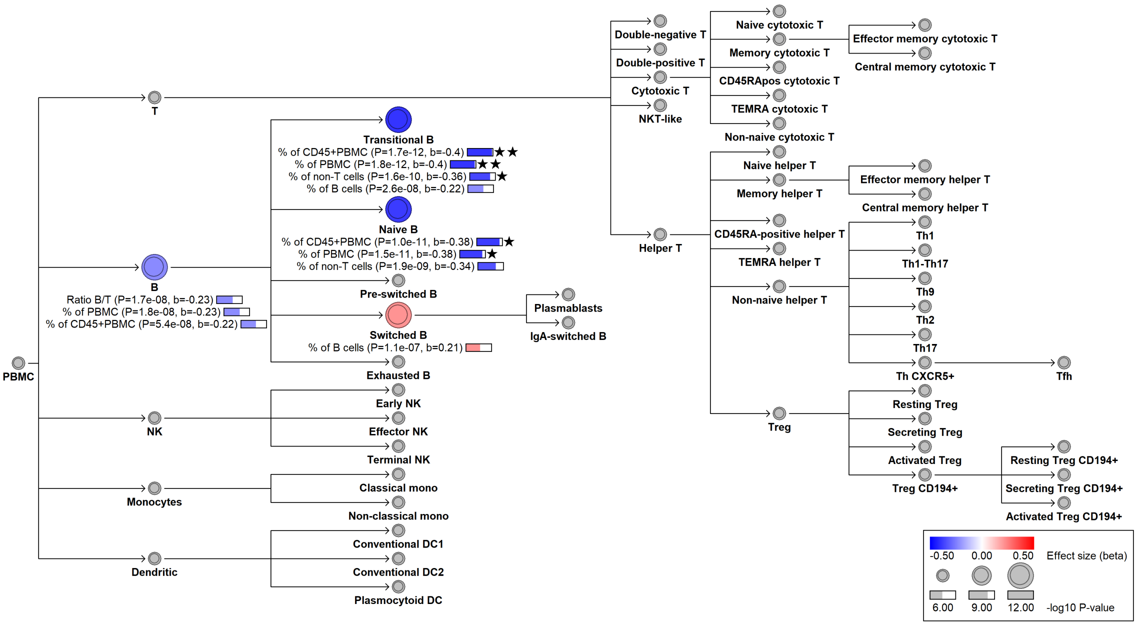

#### chr7:50268931[C→T; EAF 29.735%] (rs876038, cs_210)

Candidate genes: *IKZF1*
Diseases associations: Chronic diseases of tonsils
Plasma pQTLs: ADA2, BST2, CCDC50, CLEC4C, Composite immunoglobulin trait (IgA*IgG*IgM), Composite immunoglobulin trait IgA*IgG, Composite immunoglobulin trait IgA/IgM, Composite immunoglobulin trait IgM/(IgA*IgG), IL3RA, ISG15, Immunoglobulin A, Immunoglobulin G, LAG3, LGALS9, LILRA4, LTF, LY9, MARCO, MZB1, OLFM4, RETN, SIGLEC1, SIGLEC6, SLAMF7

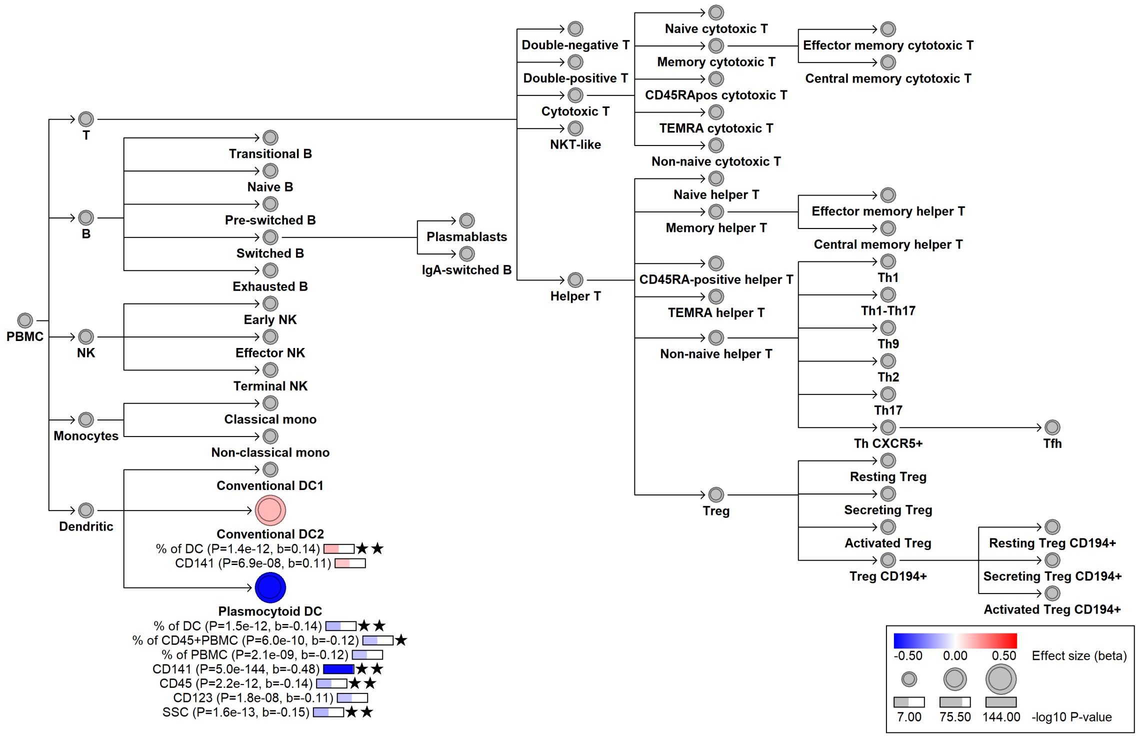

#### chr7:57916522[T→C; EAF 0.6%] (rs200698176, cs_59)

Candidate genes: *ZNF716*
Diseases associations: NA
Plasma pQTLs: NA

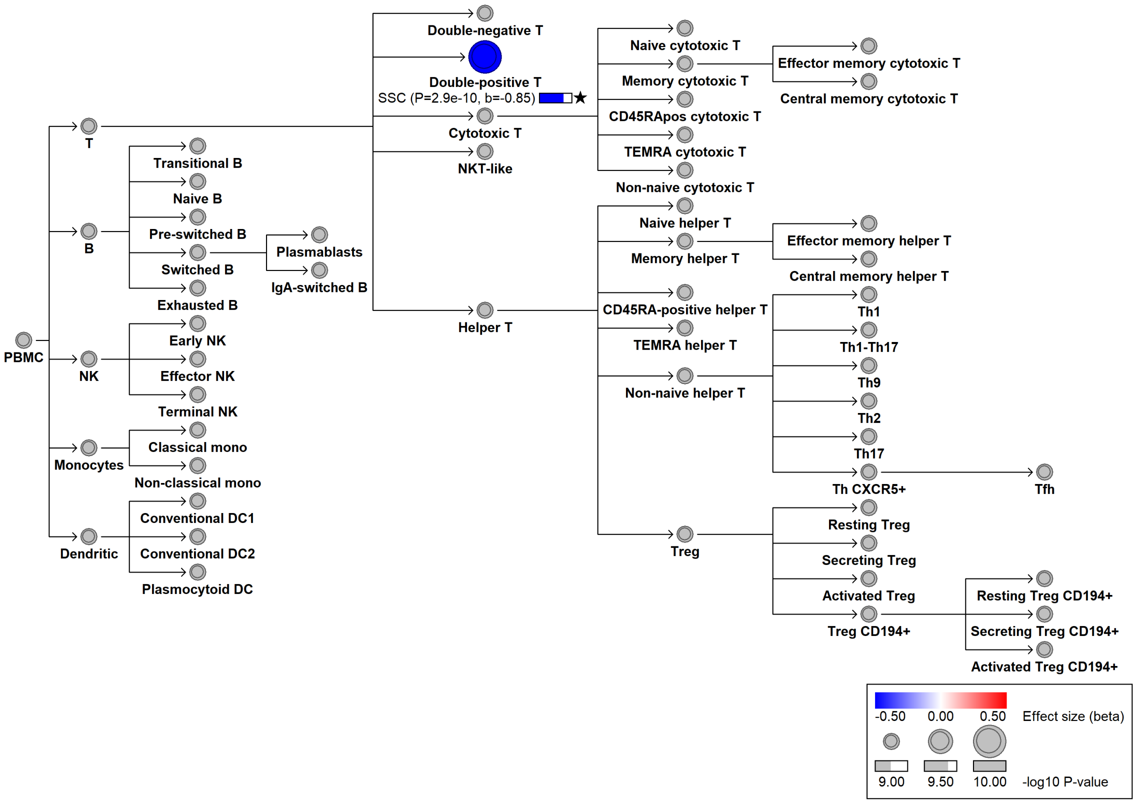

#### chr7:63857581[A→AT; EAF 0.616%] (rs529932388, cs_105)

Candidate genes: *ZNF722*
Diseases associations: NA
Plasma pQTLs: NA

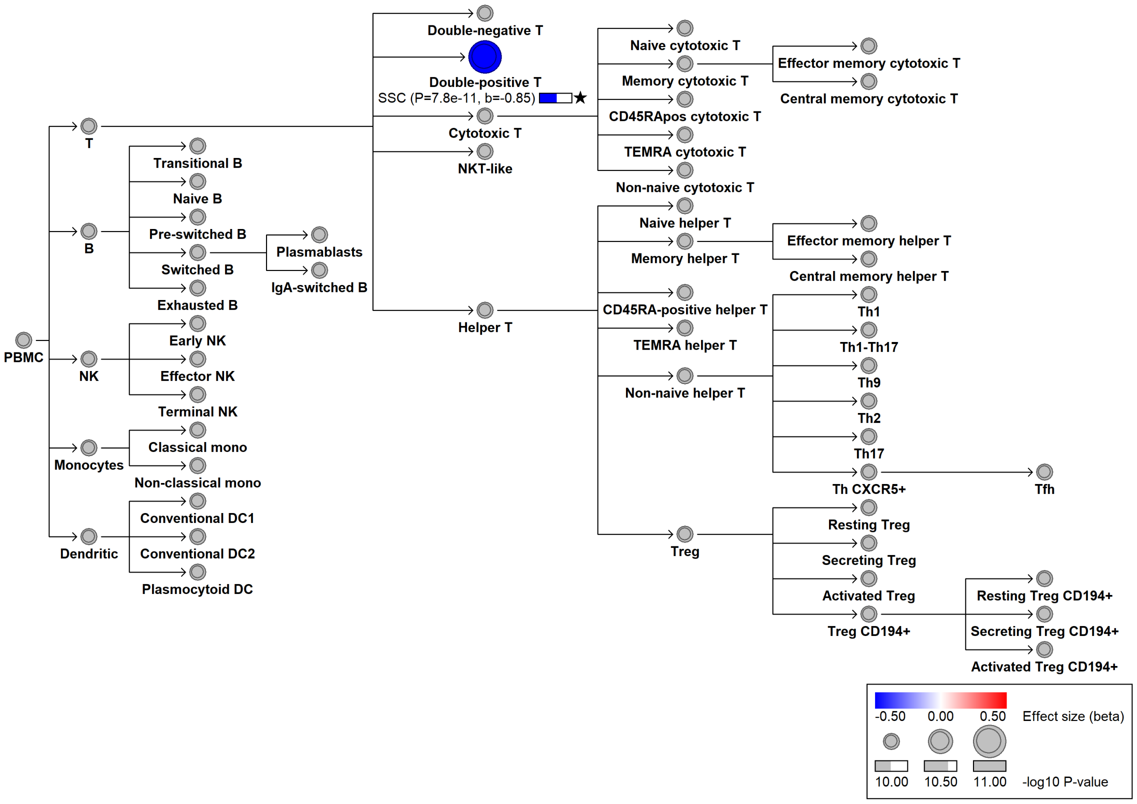

##### chr7:80597569[C→T; EAF 47.757%] (rs7807607, cs_43)

Candidate genes: *CD36*
Diseases associations: NA
Plasma pQTLs: ANGPT1, APP, ARSB, BCL2L11, BDNF, BGN, CALCOCO2, CCL13, CCL14, CCL28, CCL5, CCN2, CD226, CD36, CD40, CD40LG, CD69, CLEC1B, COL2A1, CPXM1, CXCL6, DKK1, EGF, EGFL7, ESAM, F11R, FSTL1, FUT8, GP1BA, HBEGF, HPSE, ITPR1, LILRB4, MDK, MMP1, MPIG6B, NID1, NID2, PDGFA, PDGFB, PF4, PPBP, PSAP, SELP, SERPINE1, SERPINE2, SORT1, SPARC, SPINT2, SUSD1, THPO, TIMP1, TNFSF14, TPP1, TREML1, VEGFA, VEGFC, VSIR

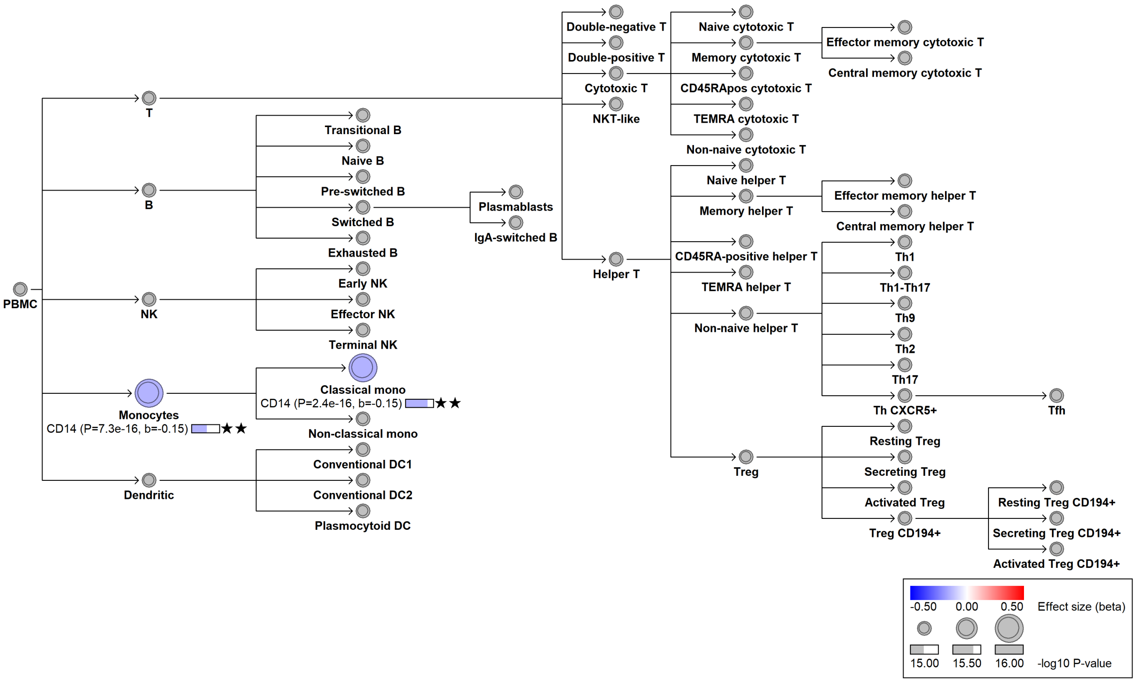

#### chr7:92607515[A→G; EAF 20.938%] (rs4272, cs_208)

Candidate genes: *CDK6*
Diseases associations: Rheumatoid arthritis
Plasma pQTLs: NA

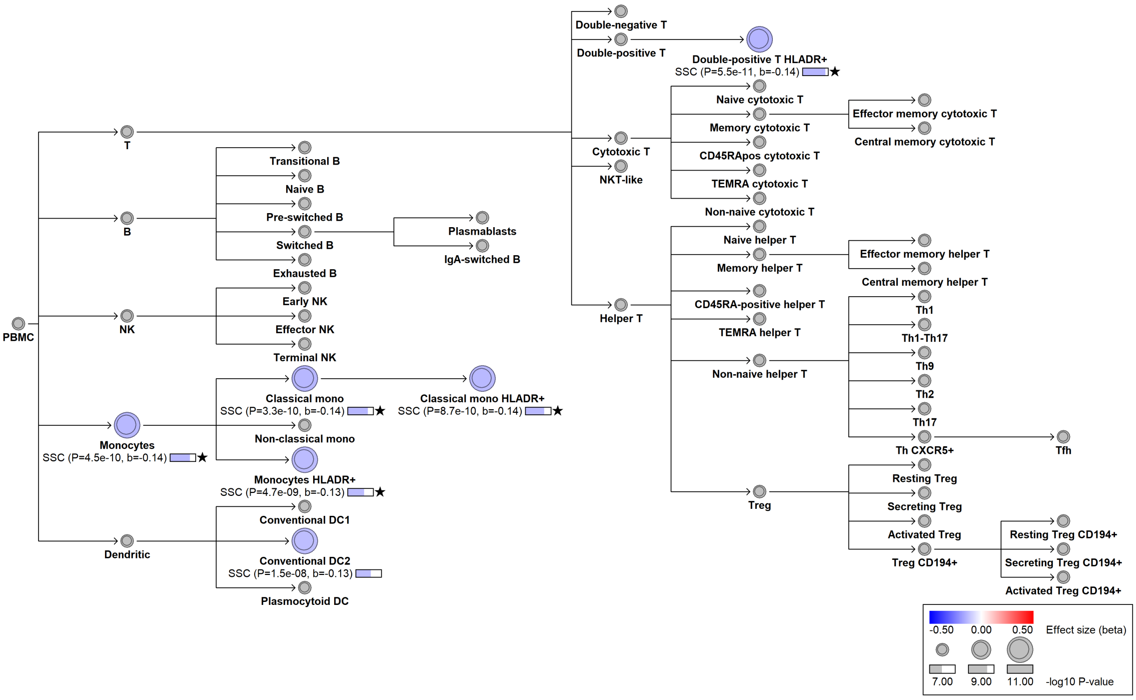

#### chr7:92779056[C→T; EAF 10.001%] (rs445, cs_42)

Candidate genes: *CDK6*
Diseases associations: NA
Plasma pQTLs: CLEC4D, CLEC6A, CRISP3, CSF3, CSF3R, RETN, SELL, TNFRSF10C

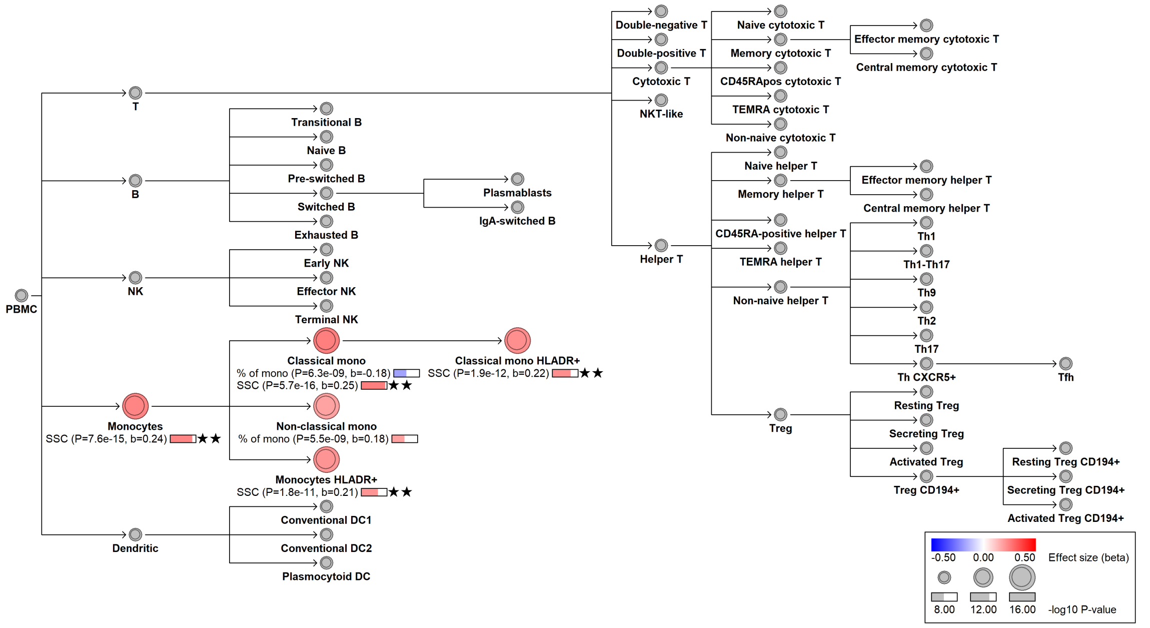

#### chr8:9527294[G→A; EAF 0.14%] (rs529968372, cs_24)

Candidate genes: *NA*
Diseases associations: Osteoarthritis, Systemic lupus erythematosus, Type 2 diabetes
Plasma pQTLs: NA

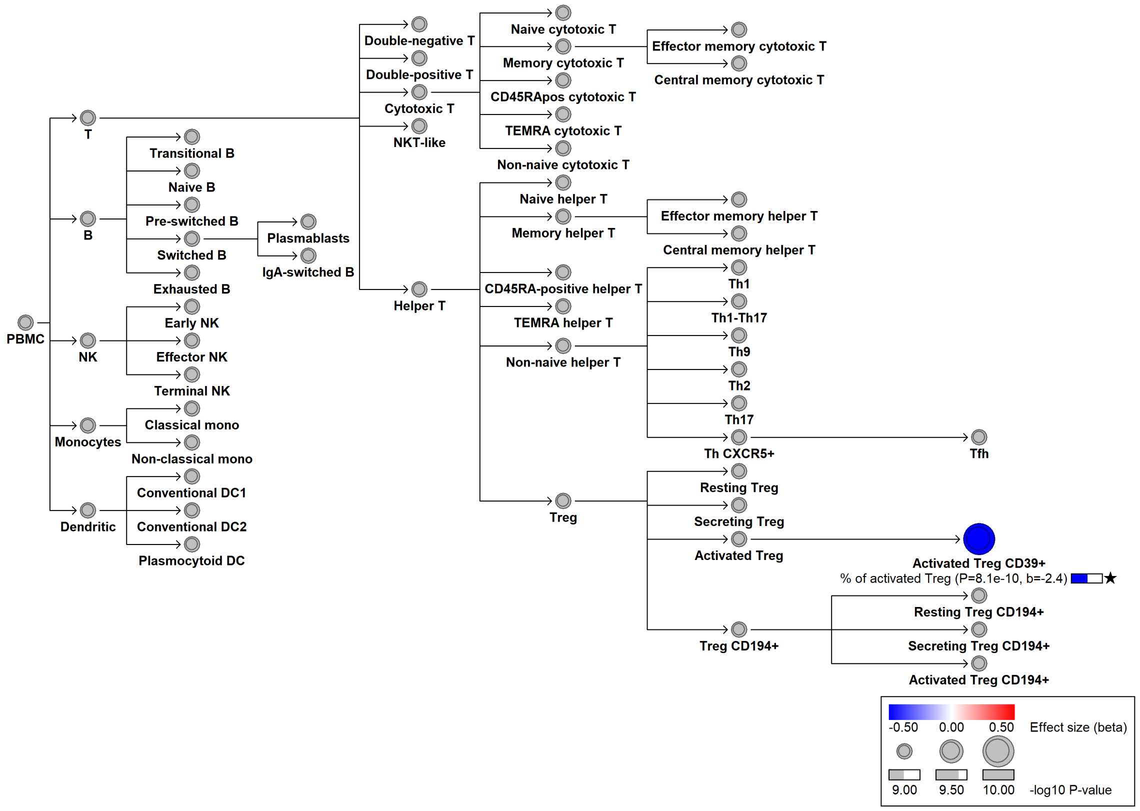

#### chr8:51899783[G→!; EAF 9.989%] (rs3763547, cs_28)

Candidate genes: *PCMTD1*
Diseases associations: NA
Plasma pQTLs: NA

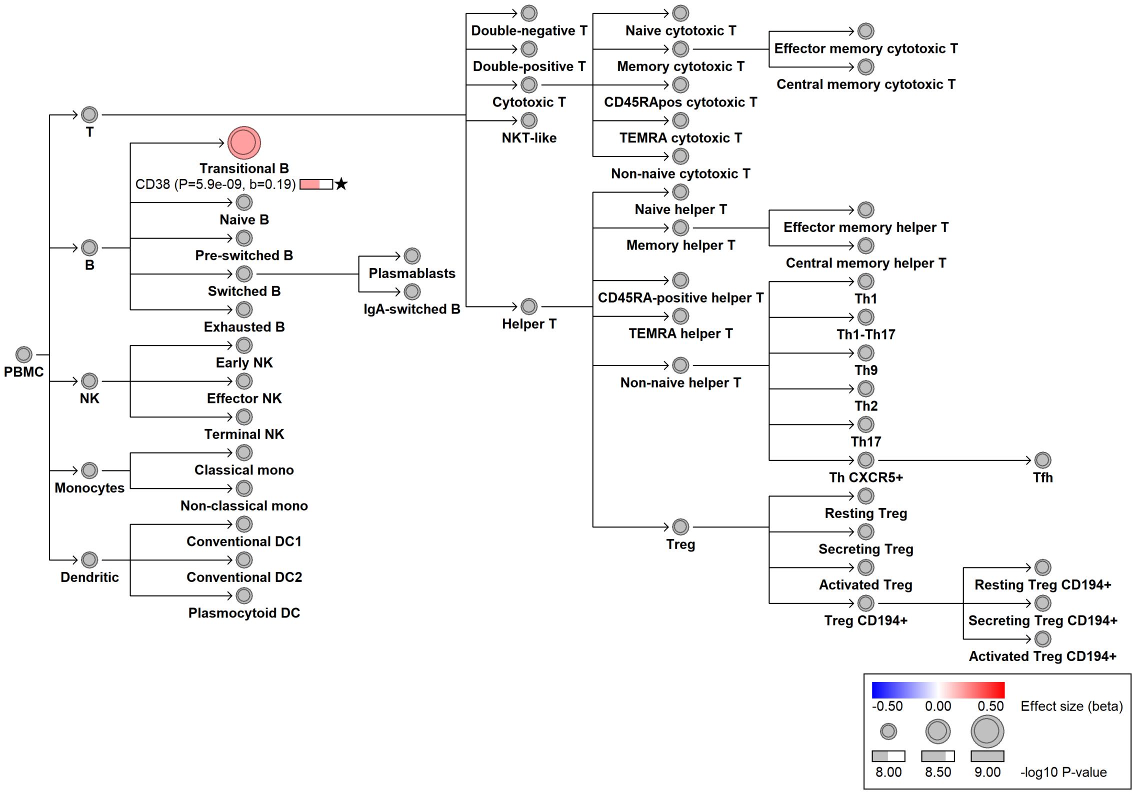

#### chr8:79772646[A→G; EAF 25.249%] (rs2956252, cs_167)

Candidate genes: *HEY1*
Diseases associations: NA
Plasma pQTLs: NA

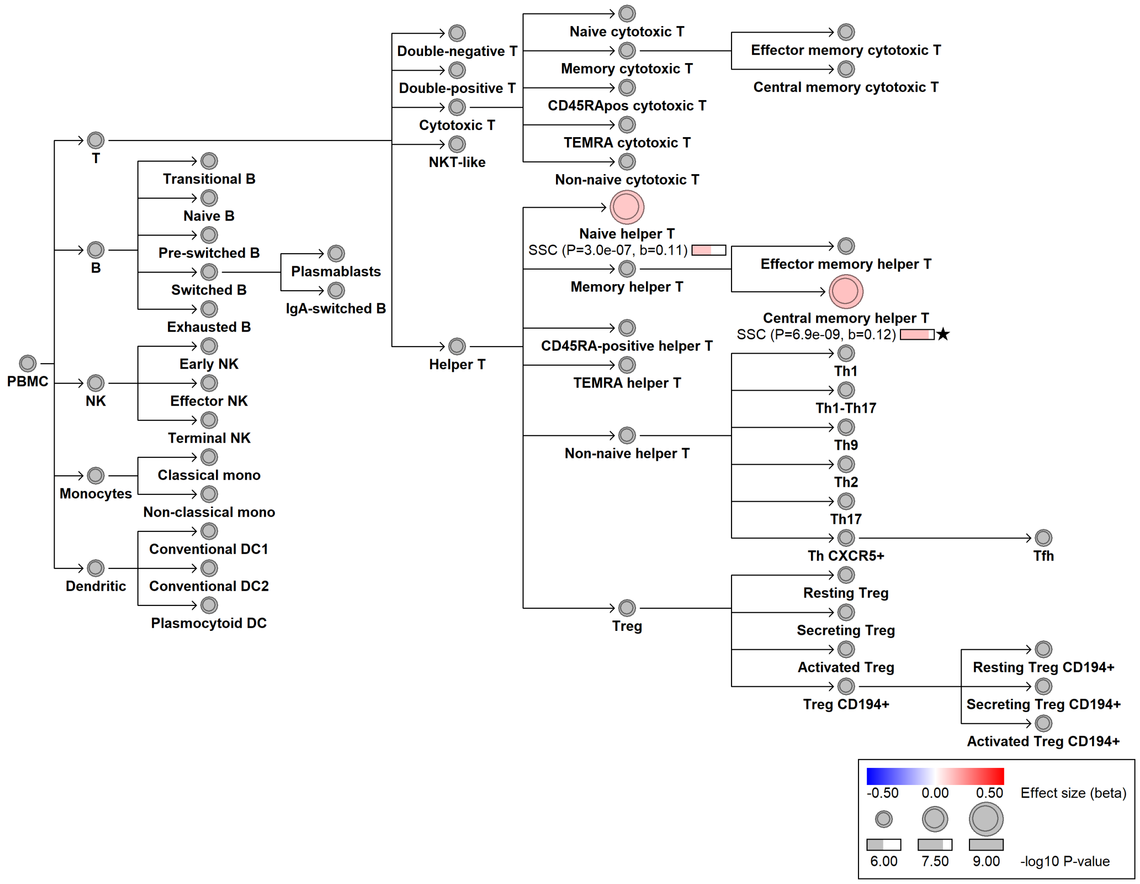

#### chr8:128267743[G→A; EAF 0.906%] (rs146034887, cs_33)

Candidate genes: *MYC*
Diseases associations: NA
Plasma pQTLs: NA

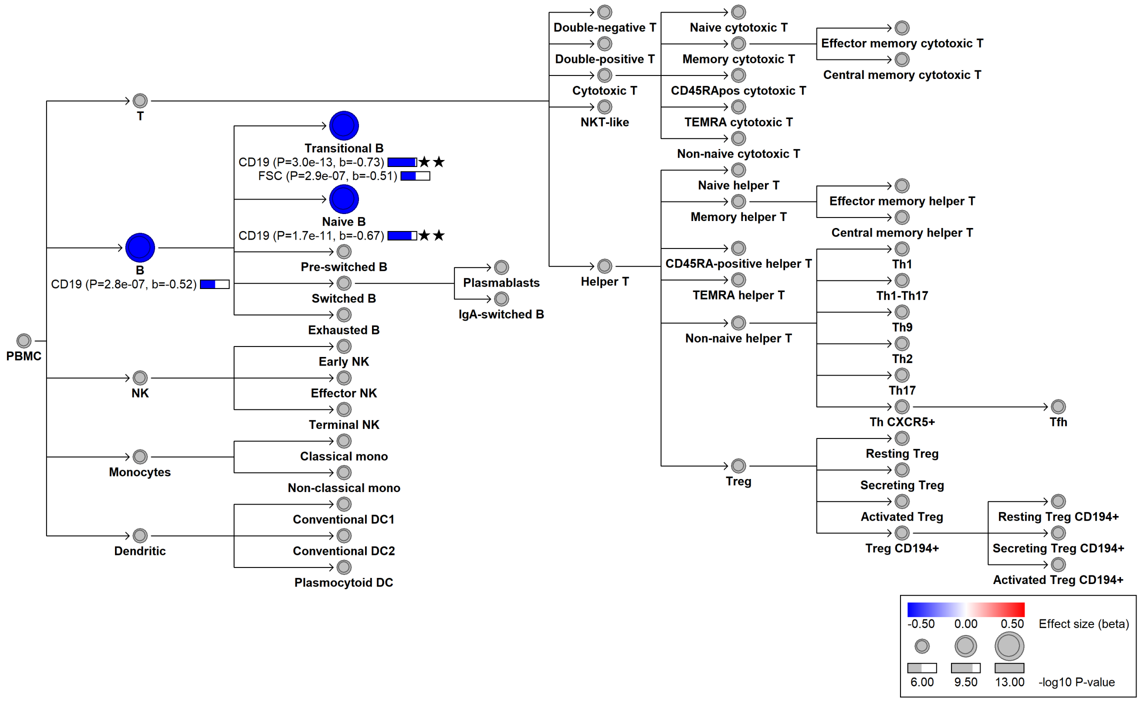

### chr8:129593685[C→T; EAF 45.571%] (rs12544501, cs_211)

Candidate genes: *MYC*
Diseases associations: NA
Plasma pQTLs: NA

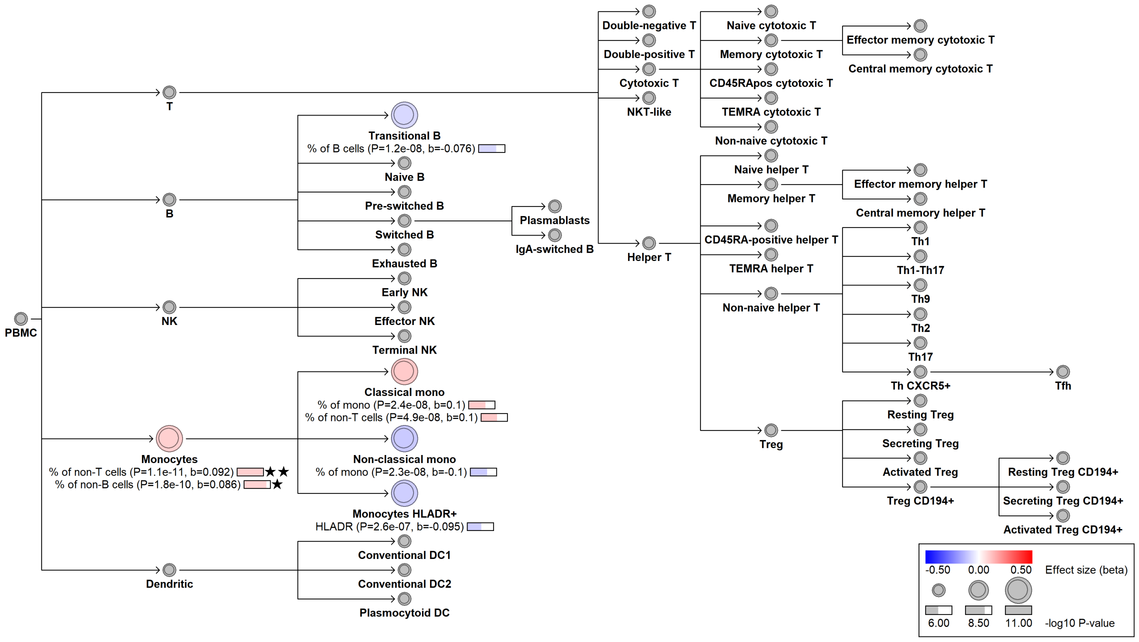

#### chr8:134613158[G→A; EAF 40.915%] (rs4909532, cs_221)

Candidate genes: *ZFAT*
Diseases associations: NA
Plasma pQTLs: NA

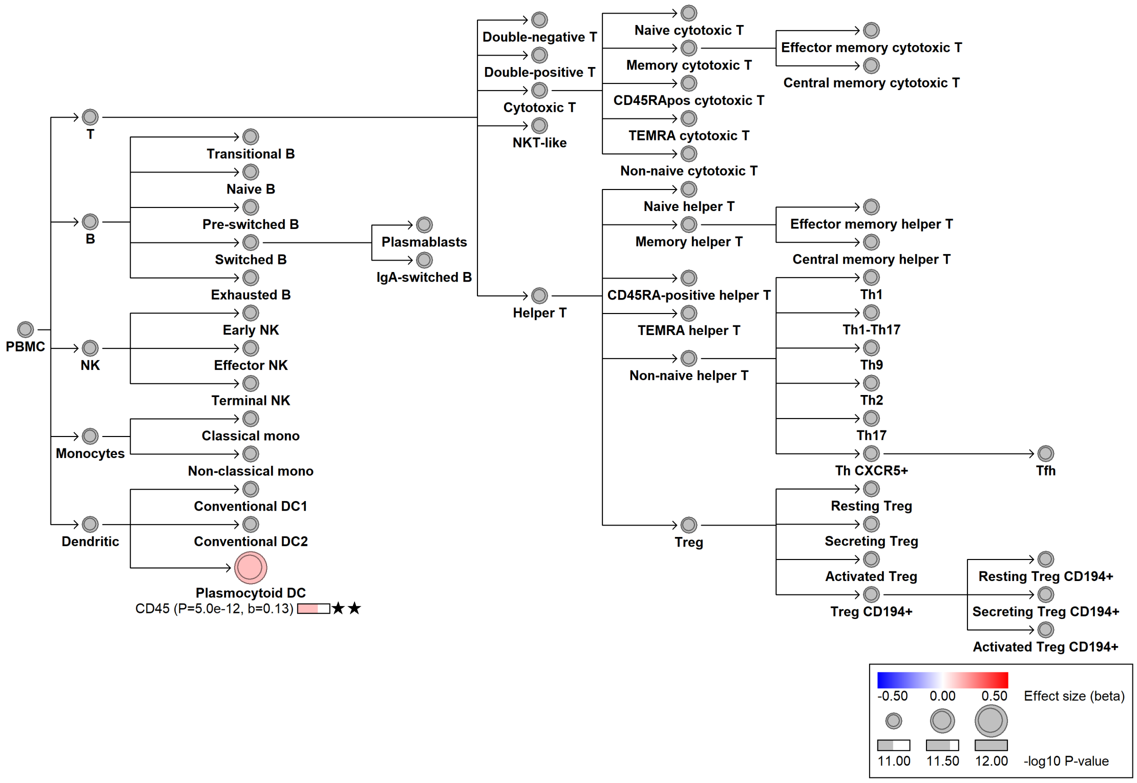

### chr8:143804639[C→T; EAF 5.54%] (rs11542374, cs_256)

Candidate genes: *SCRIB*
Diseases associations: NA
Plasma pQTLs: NA

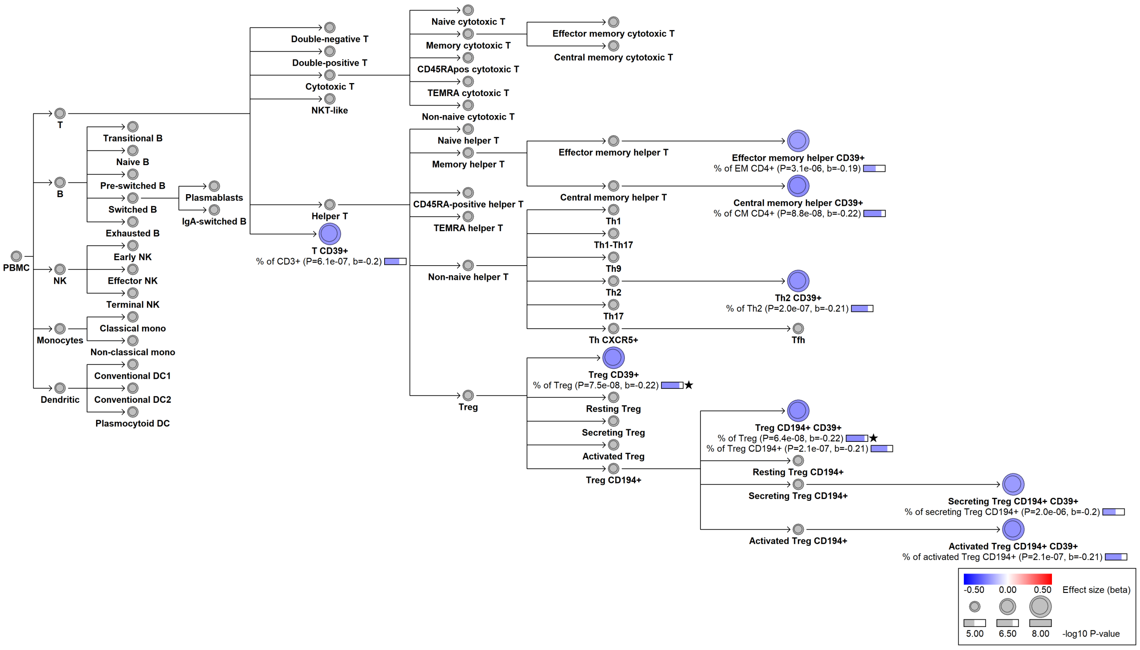

#### chr9:76455225[T→C; EAF 13.632%] (rs57326695, cs_196)

Candidate genes: *GCNT1*
Diseases associations: NA
Plasma pQTLs: NA

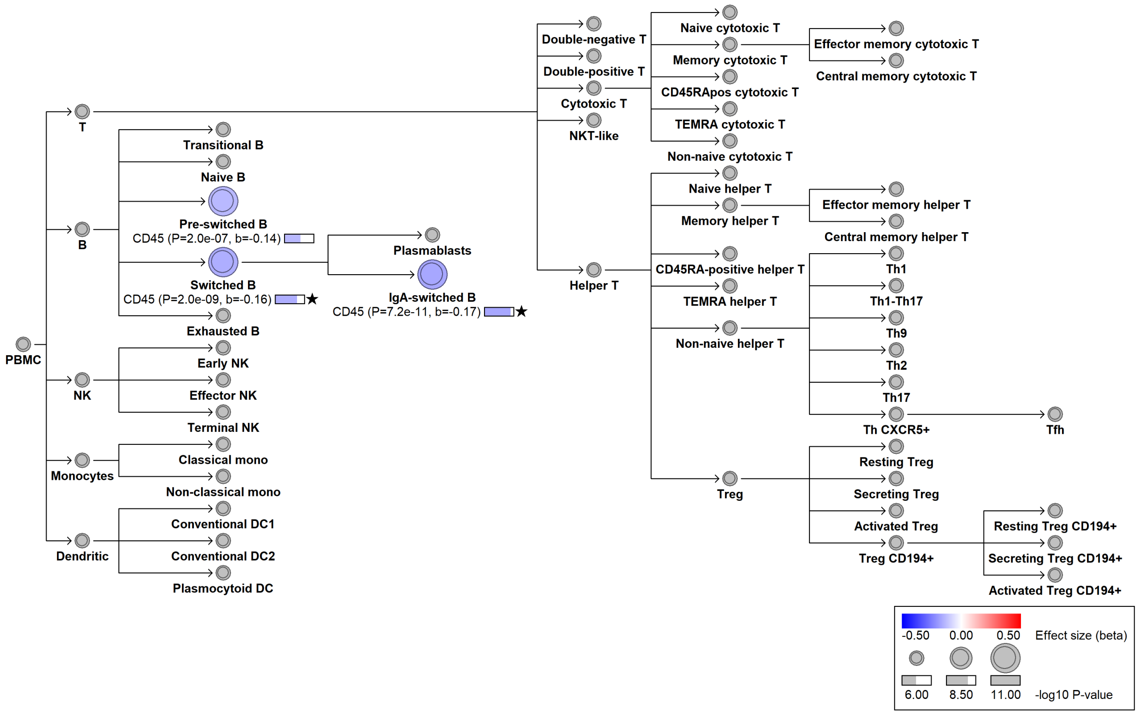

### chr9:87712180[G→A; EAF 48.387%] (rs3128499, cs_32)

Candidate genes: *CTSL*
Diseases associations: NA
Plasma pQTLs: NA

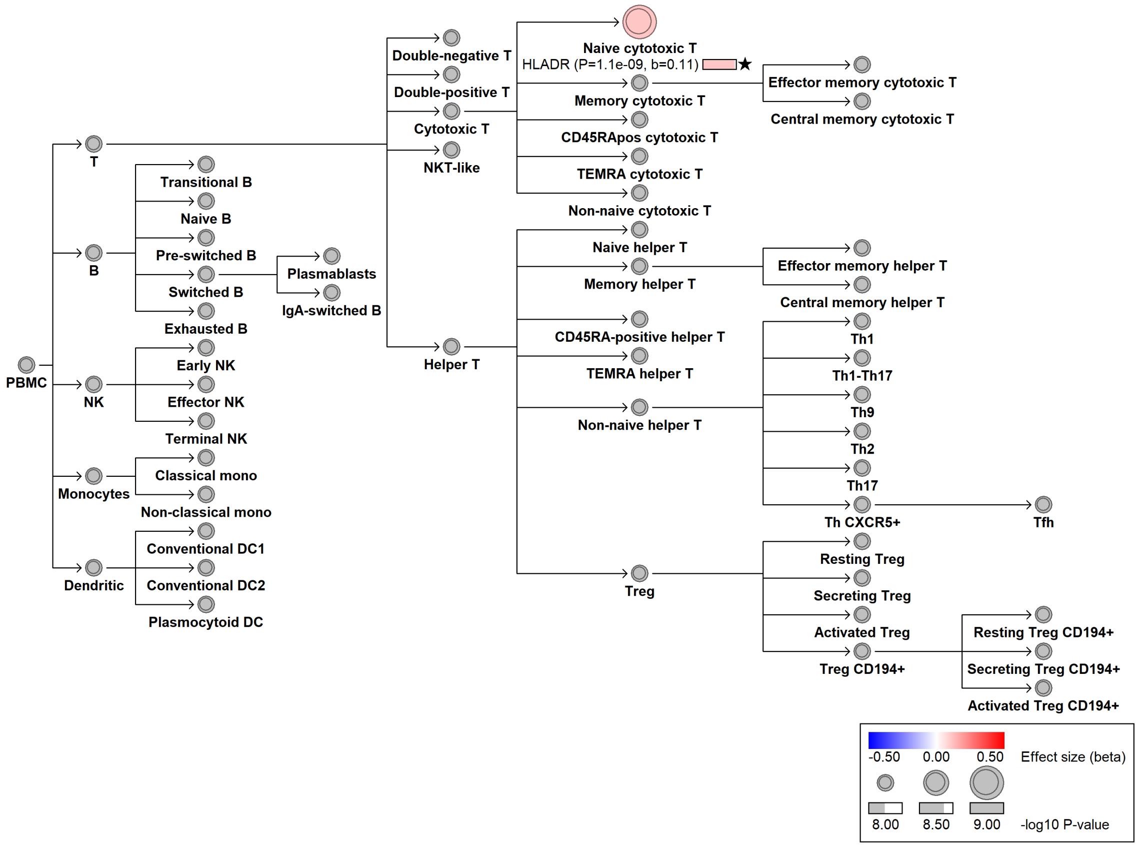

#### chr9:98020373[!→CA; EAF 19.382%] (rs201706603, cs_15)

Candidate genes: *ANP32B*
Diseases associations: Primary biliary cholangitis
Plasma pQTLs: NA

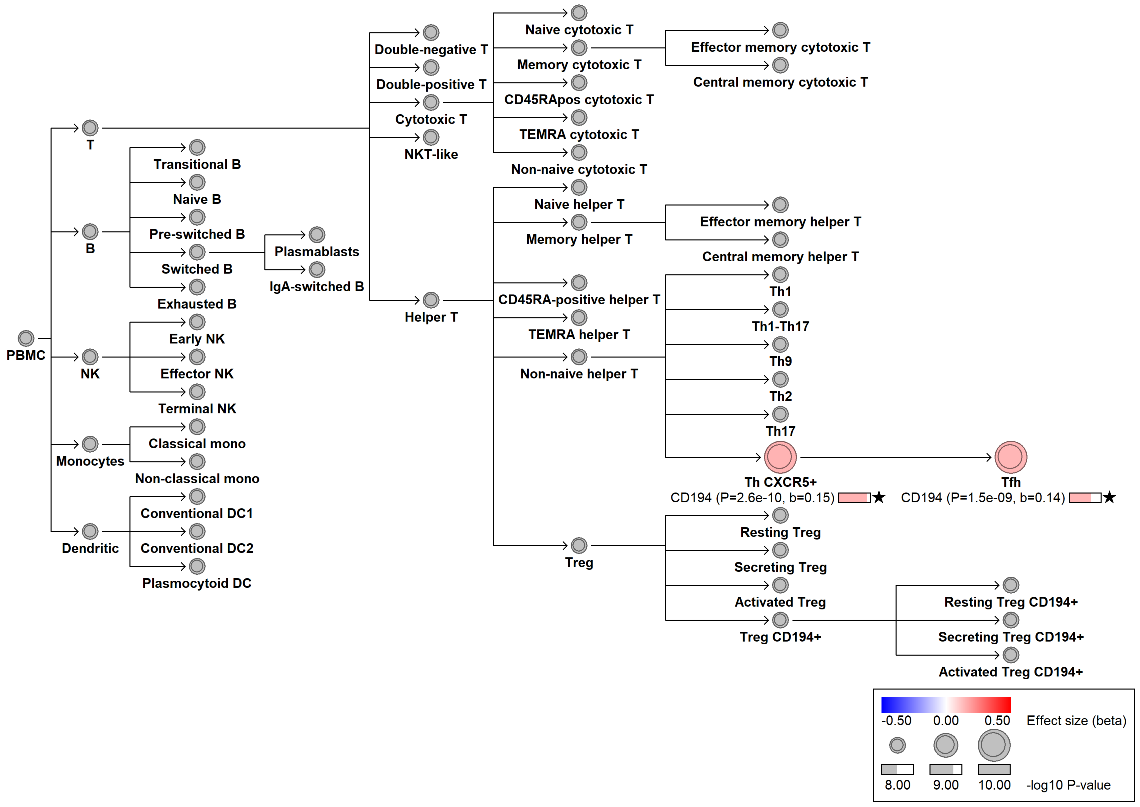

#### chr9:111150273[A→G; EAF 48.675%] (rs10980797, cs_227)

Candidate genes: *LPAR1*
Diseases associations: NA
Plasma pQTLs: LILRA5

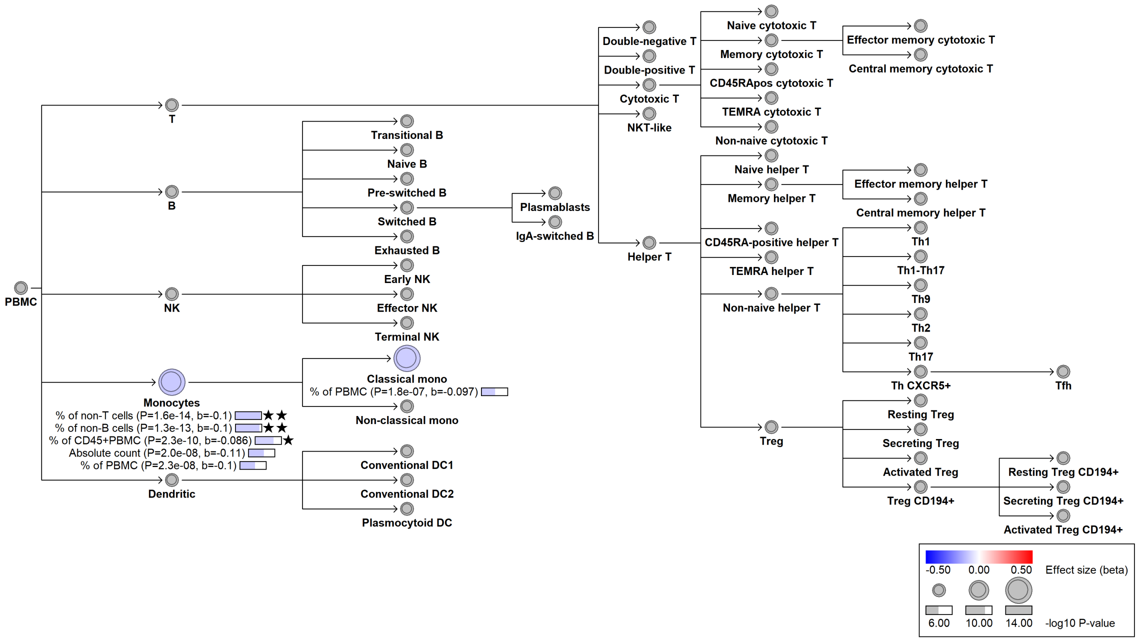

#### chr9:114885236[TGG→!; EAF 46.736%] (rs56325512, cs_262)

Candidate genes: *TNFSF8*
Diseases associations: Ankylosing spondylitis, Crohn's disease, Primary sclerosing cholangitis, Psoriasis, Ulcerative colitis
Plasma pQTLs: NA

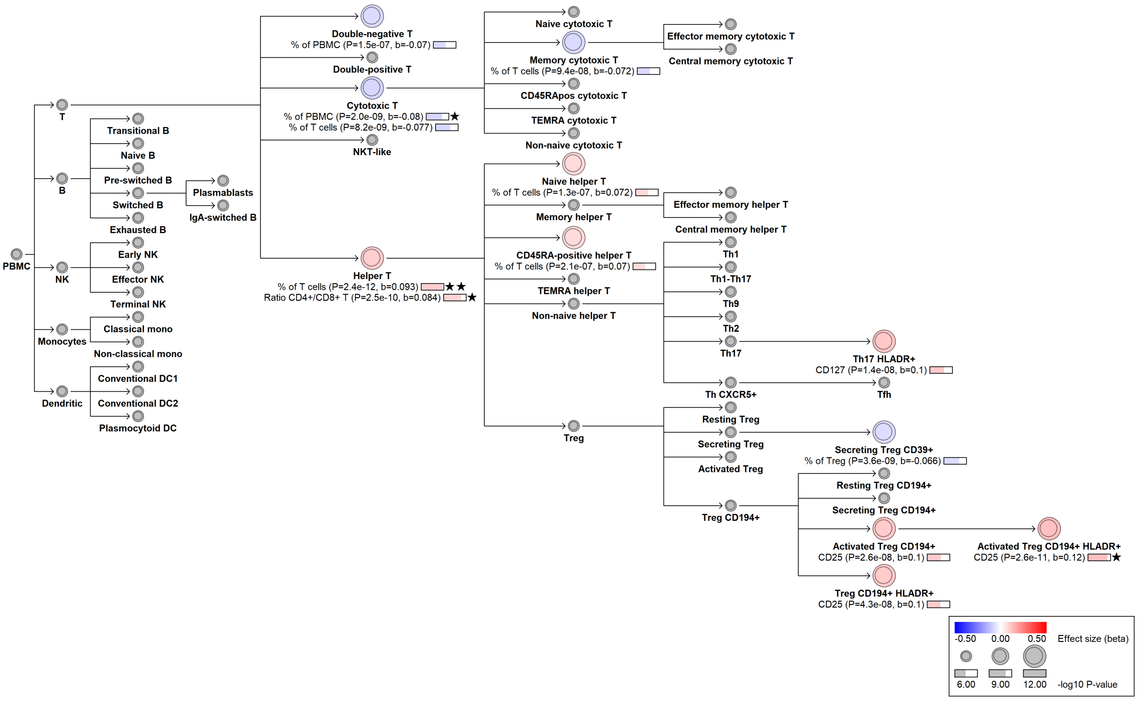

#### chr9:114930602[C→T; EAF 21.292%] (rs3181356, cs_218)

Candidate genes: *TNFSF8*
Diseases associations: NA
Plasma pQTLs: CRTAM, Composite immunoglobulin trait (IgA*IgG*IgM), Composite immunoglobulin trait IgA*IgG, Composite immunoglobulin trait IgA/IgG, Composite immunoglobulin trait IgA/IgM, Immunoglobulin A, LY9, LY96, SLAMF7, TP53BP1

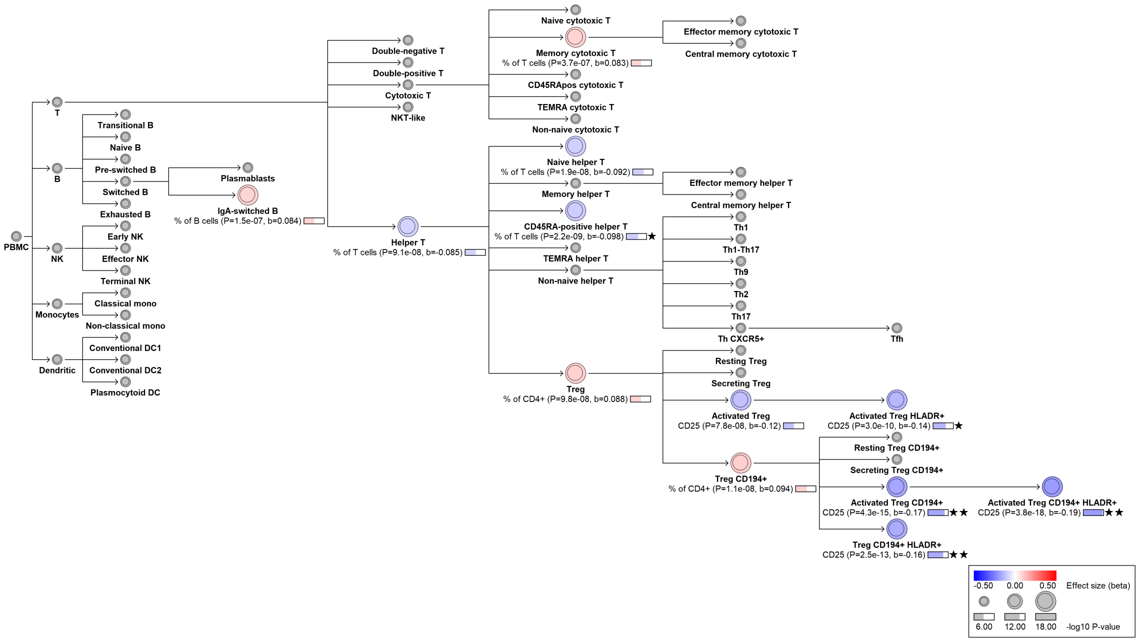

#### chr10:6033868[G→!; EAF 39.861%] (chr10:6033868, cs_140)

Candidate genes: *IL2RA*
Diseases associations: NA
Plasma pQTLs: NA

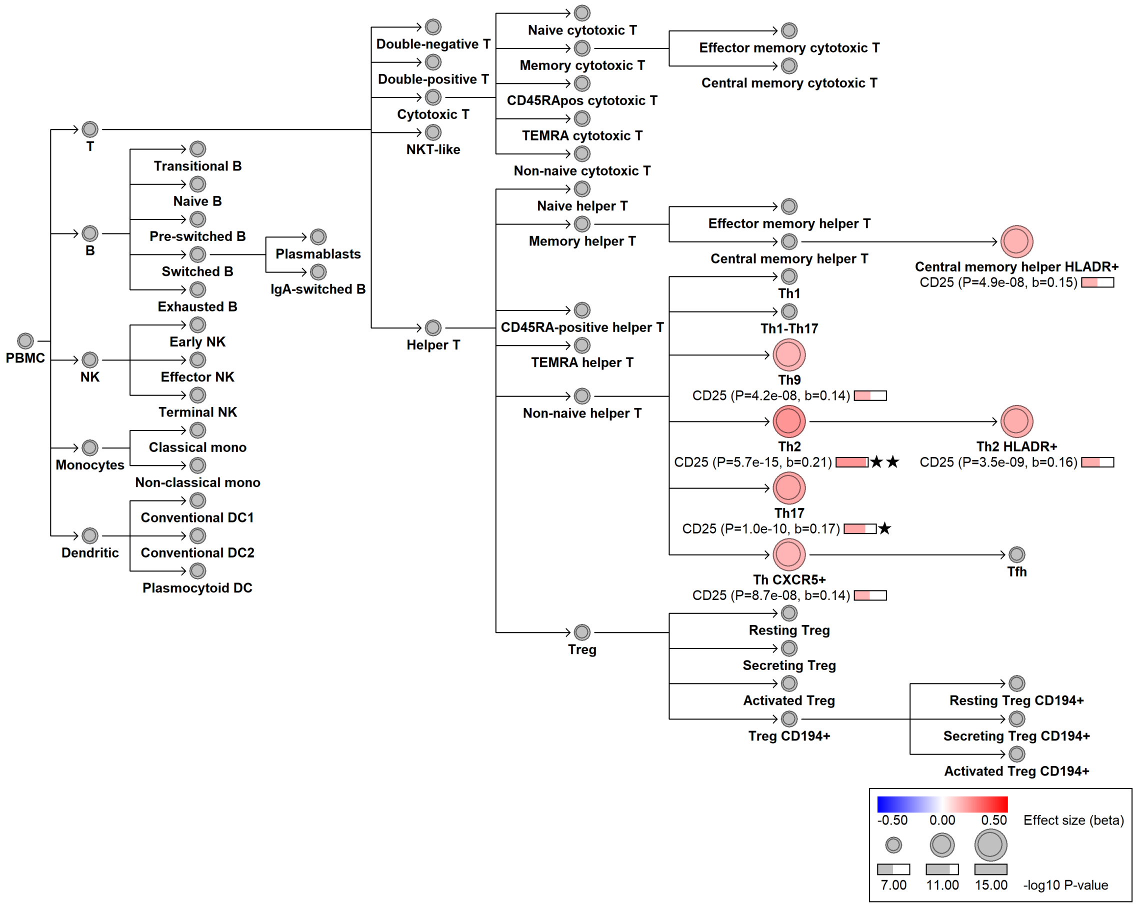

#### chr10:6051176[C→T; EAF 1.287%] (rs12722502, cs_4)

Candidate genes: *IL2RA*
Diseases associations: Allergy, Asthma, Eczema, Nonatopic asthma
Plasma pQTLs: NA

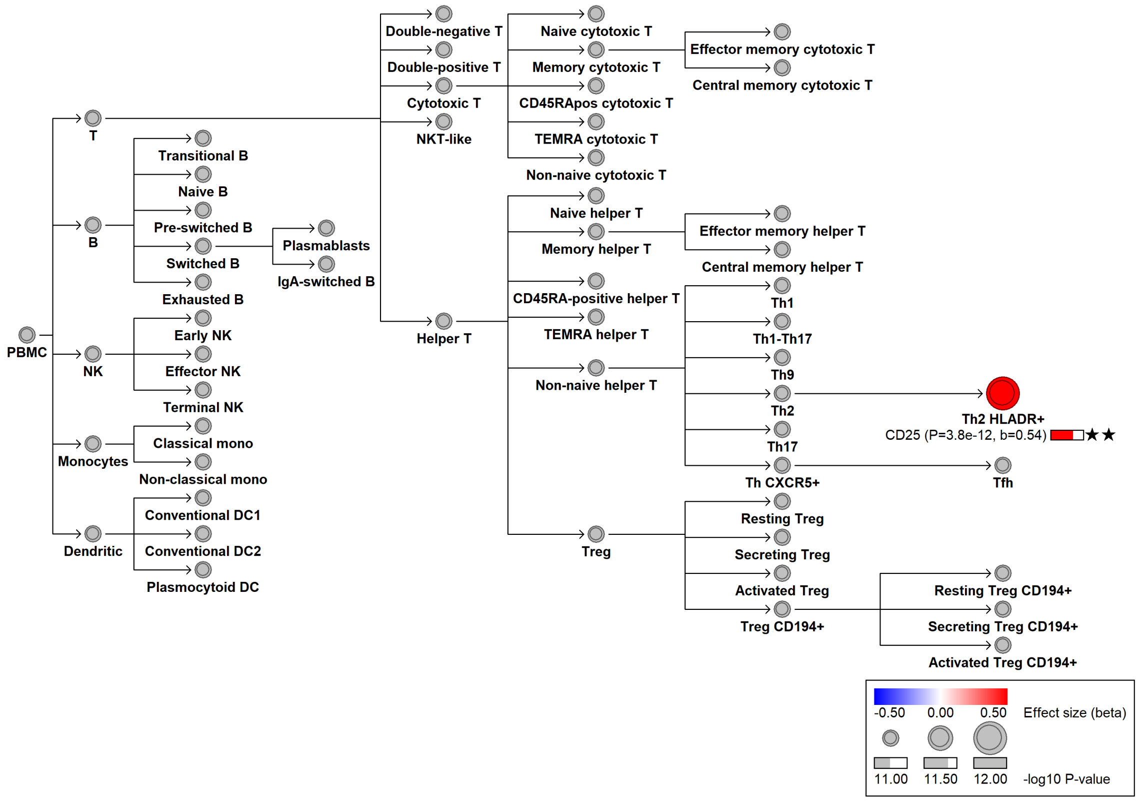

#### chr10:6052734[C→T; EAF 7.707%] (rs61839660, cs_101)

Candidate genes: *IL2RA*
Diseases associations: Allergy, Ankylosing spondylitis, Asthma, Crohn's disease, Eczema, Inflammatory bowel disease, Primary sclerosing cholangitis, Psoriasis, Type 1 diabetes, Ulcerative colitis
Plasma pQTLs: Composite immunoglobulin trait IgA*IgG, Composite immunoglobulin trait IgA/IgG, Composite immunoglobulin trait IgA/IgM, Composite immunoglobulin trait IgM/(IgA*IgG), Immunoglobulin A, LY96, MZB1, SLAMF7, TNFRSF17

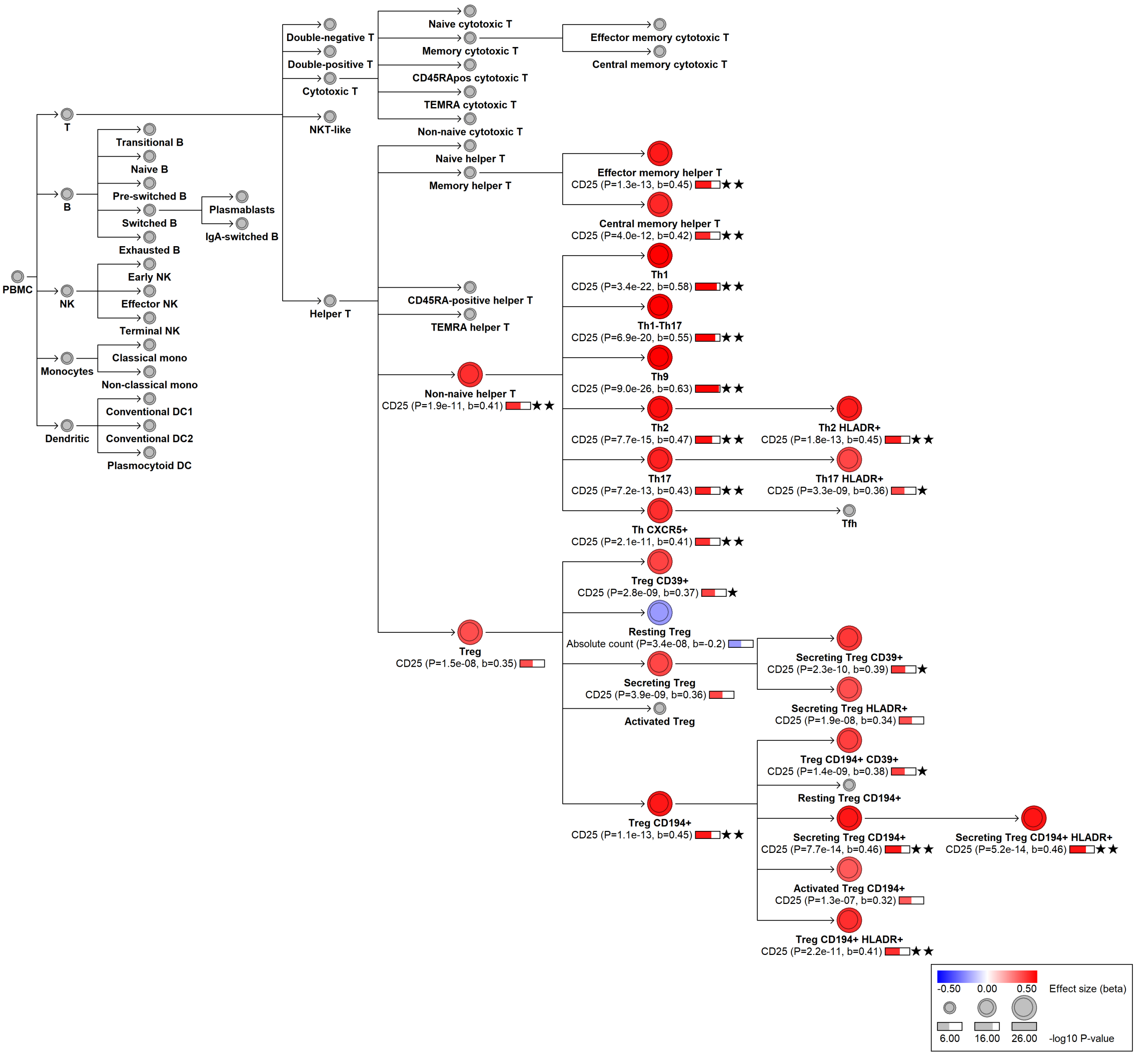

#### chr10:6060794[A→G; EAF 34.075%] (rs3118471, cs_207)

Candidate genes: *IL2RA*
Diseases associations: Ankylosing spondylitis, Autoimmune thyroiditis, Crohn's disease, Primary sclerosing cholangitis, Psoriasis, Rheumatoid arthritis, Systemic lupus erythematosus, Ulcerative colitis
Plasma pQTLs: NA

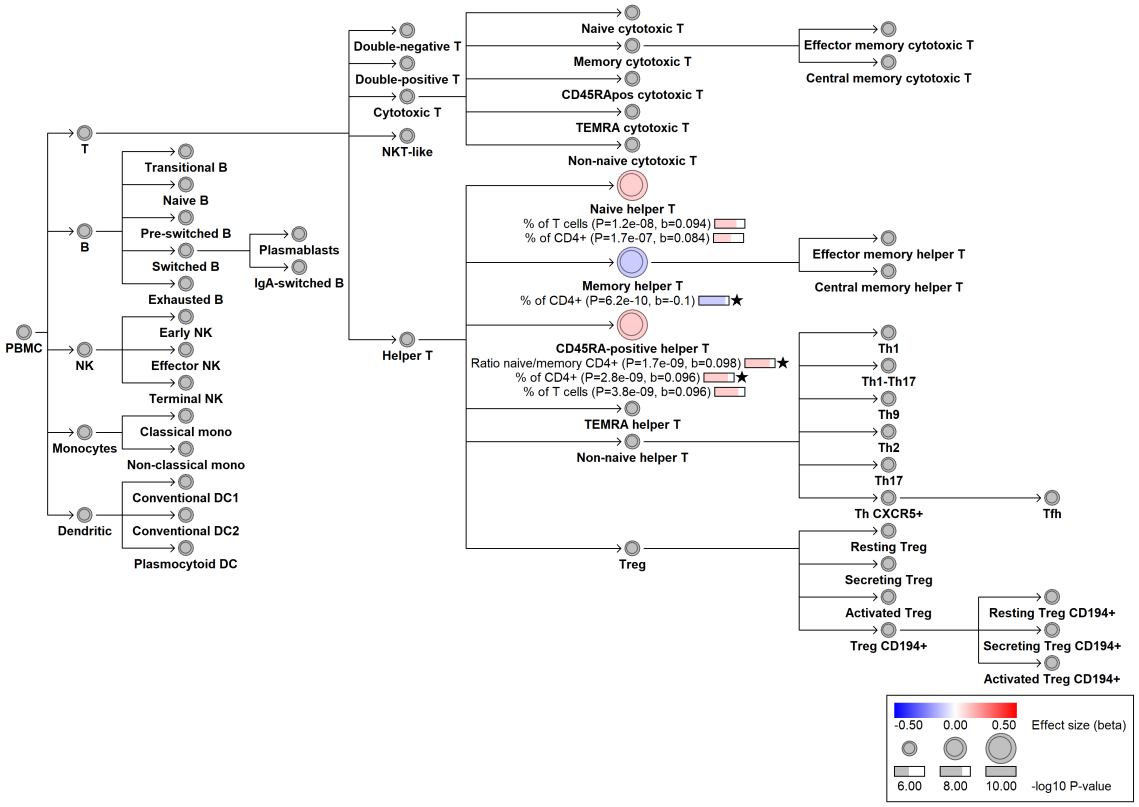

### chr10:6065302[T→C; EAF 19.19%] (rs3118475, cs_35)

Candidate genes: *IL2RA*
Diseases associations: NA
Plasma pQTLs: NA

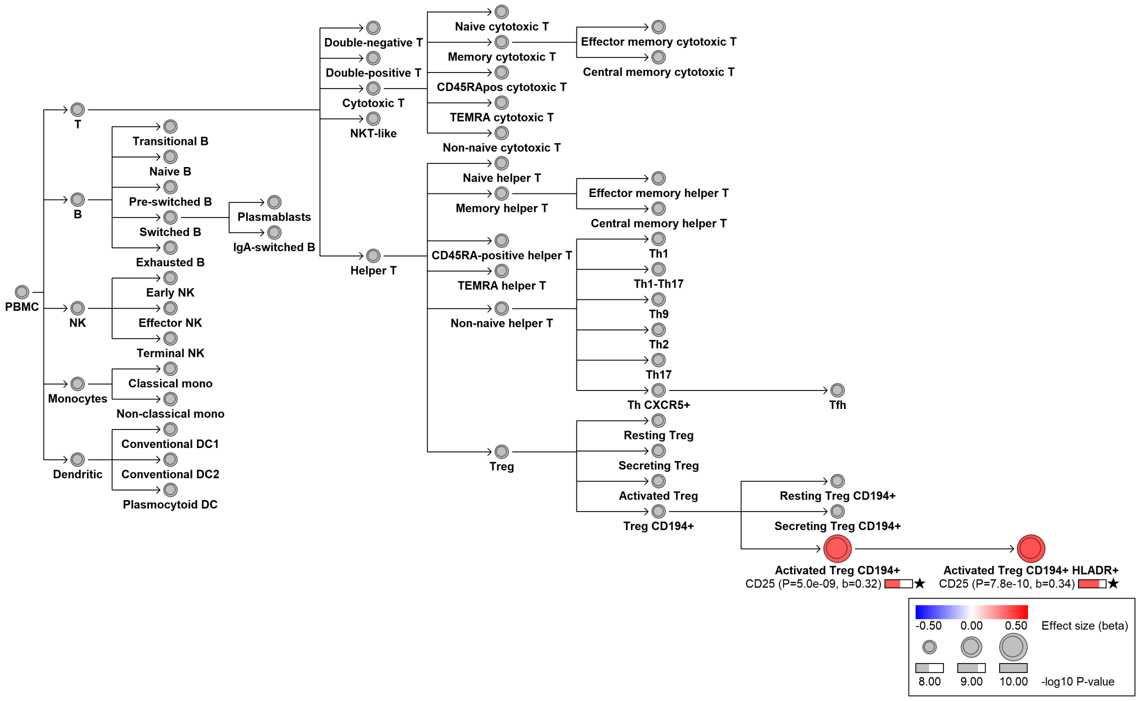

#### chr10:61993723[A→G; EAF 31.514%] (rs9415635, cs_77)

Candidate genes: *ARID5B*
Diseases associations: Childhood ALL
Plasma pQTLs: BLNK, CD22, CD72, CD79B, DRAXIN, FCER2, FCRL1, IGLL1, PCDH9, TCL1A, TNFRSF13C

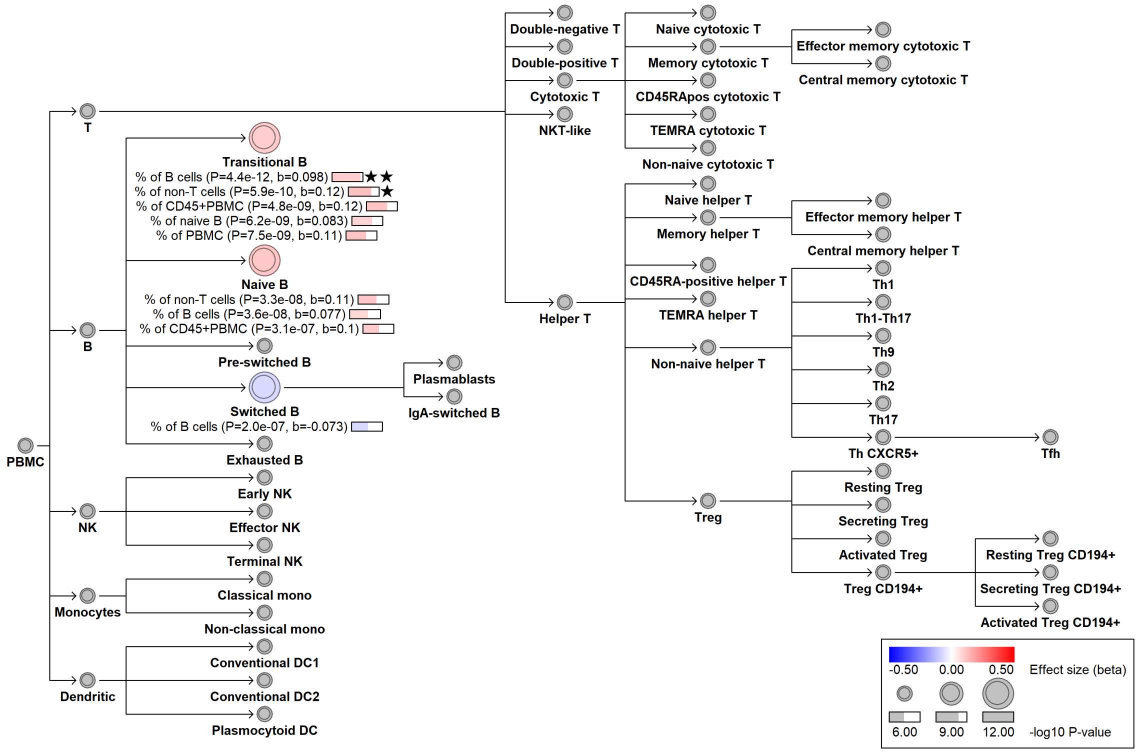

#### chr10:95633718[C→!; EAF 25.035%] (chr10:95633718, cs_17)

Candidate genes: *ENTPD1*
Diseases associations: NA
Plasma pQTLs: NA

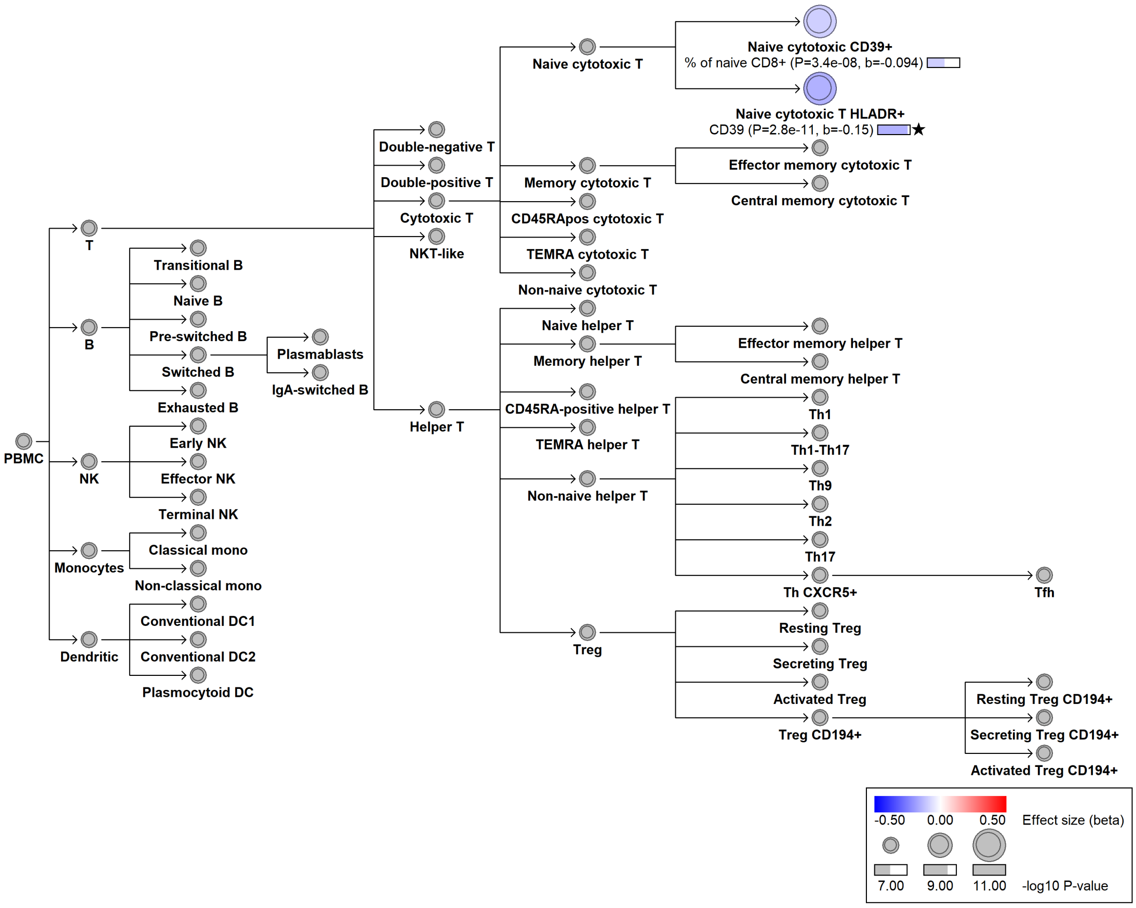

### chr10:95661517[!→A; EAF 34.157%] (rs4918955, cs_217)

Candidate genes: *ENTPD1*
Diseases associations: NA
Plasma pQTLs: NA

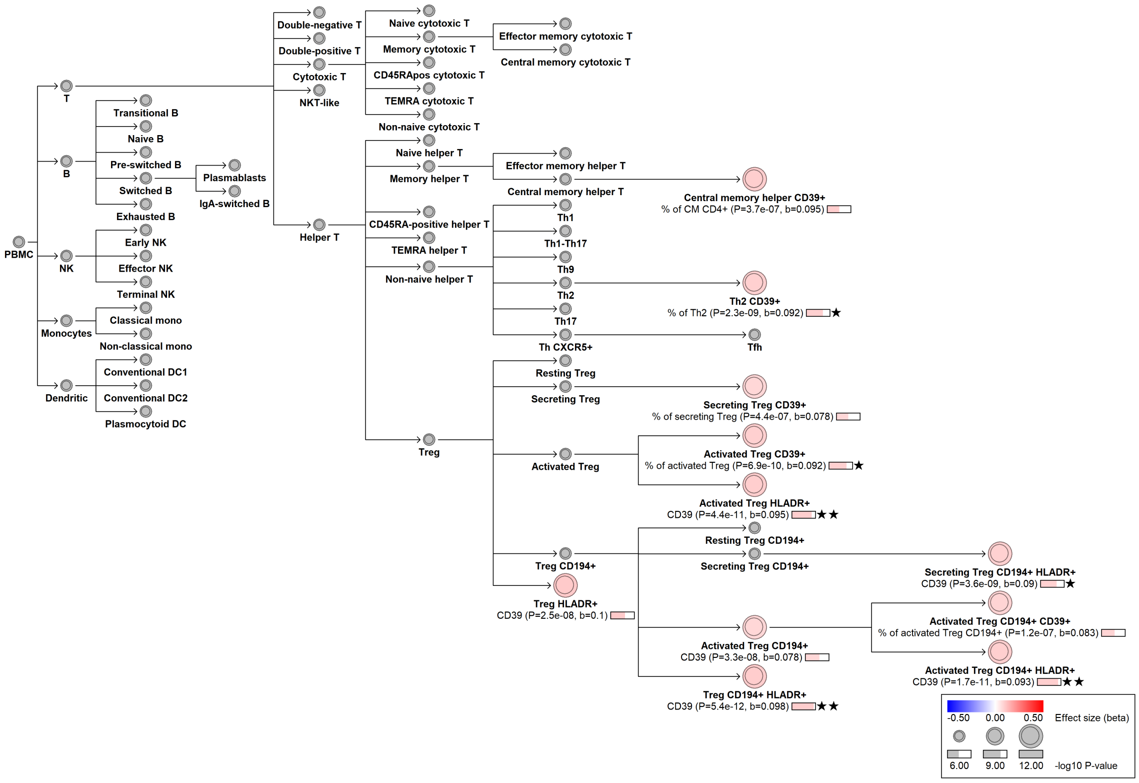

#### chr10:95751710[G→A; EAF 1.853%] (rs7084741, cs_85)

Candidate genes: *ENTPD1*
Diseases associations: NA
Plasma pQTLs: NA

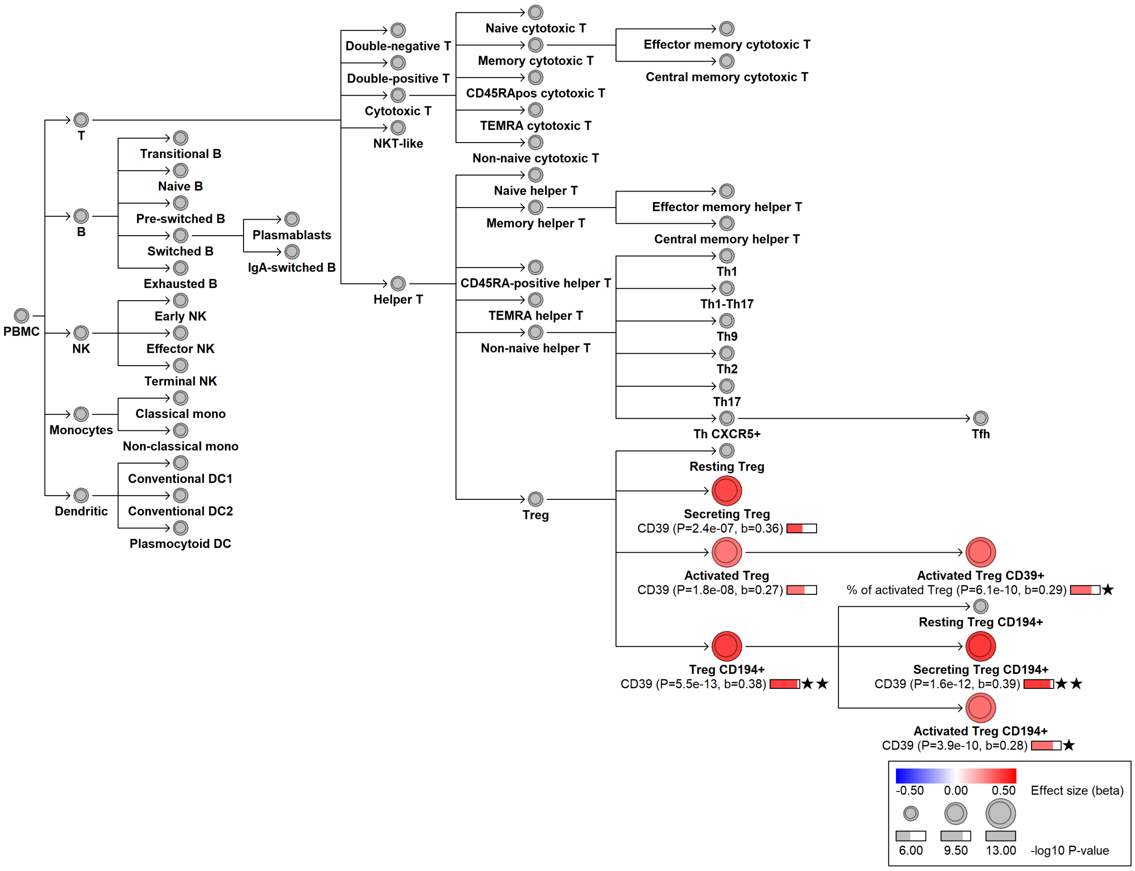

### chr10:95755650[C→G; EAF 1.555%] (rs138691825, cs_79)

Candidate genes: *ENTPD1*
Diseases associations: NA
Plasma pQTLs: NA

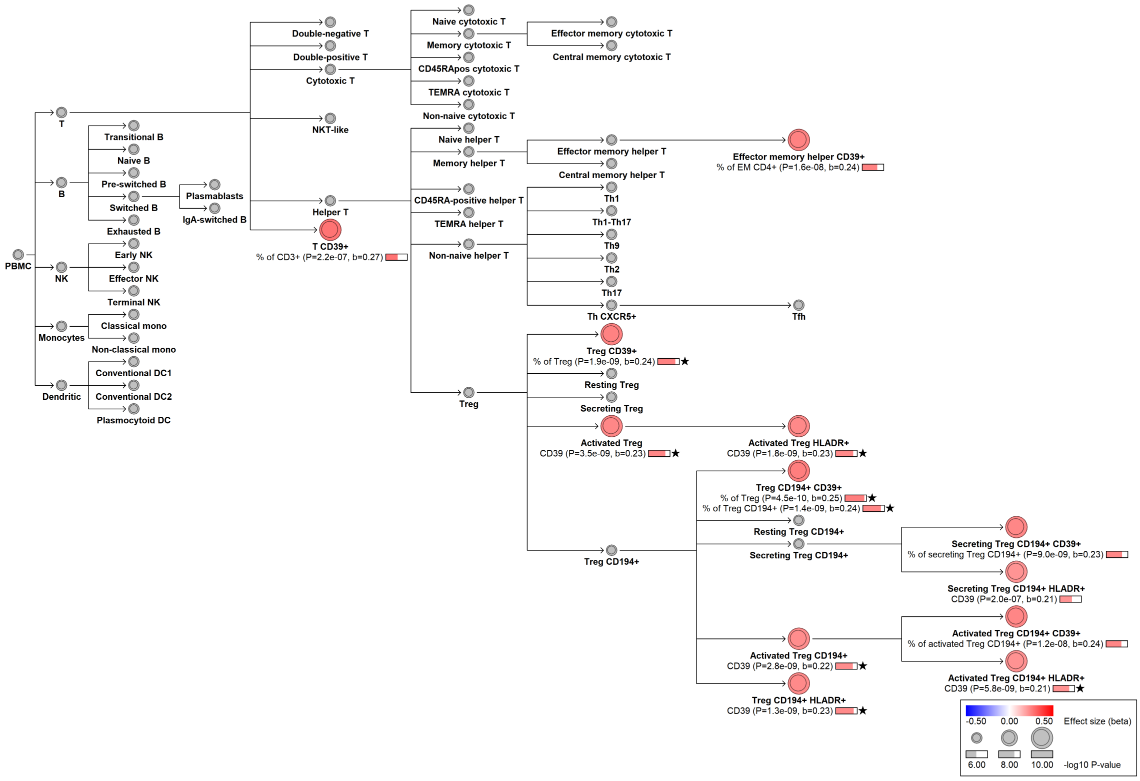

### chr10:95756191[!→TG; EAF 4.07%] (rs149930011, cs_130)

Candidate genes: *ENTPD1*
Diseases associations: NA
Plasma pQTLs: NA

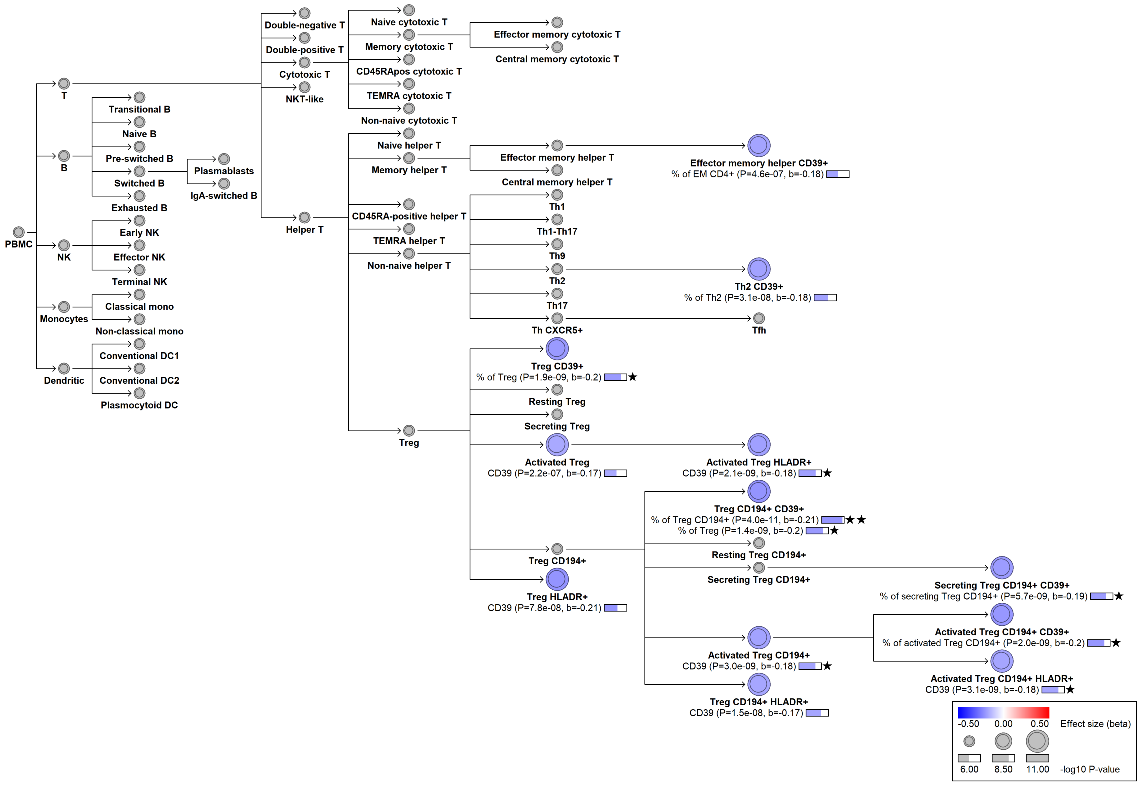

**chr10:95789057[G→A; EAF 45.378%] (rs4918968, cs_186)**

Candidate genes: *ENTPD1*
Diseases associations: NA
Plasma pQTLs: CFP

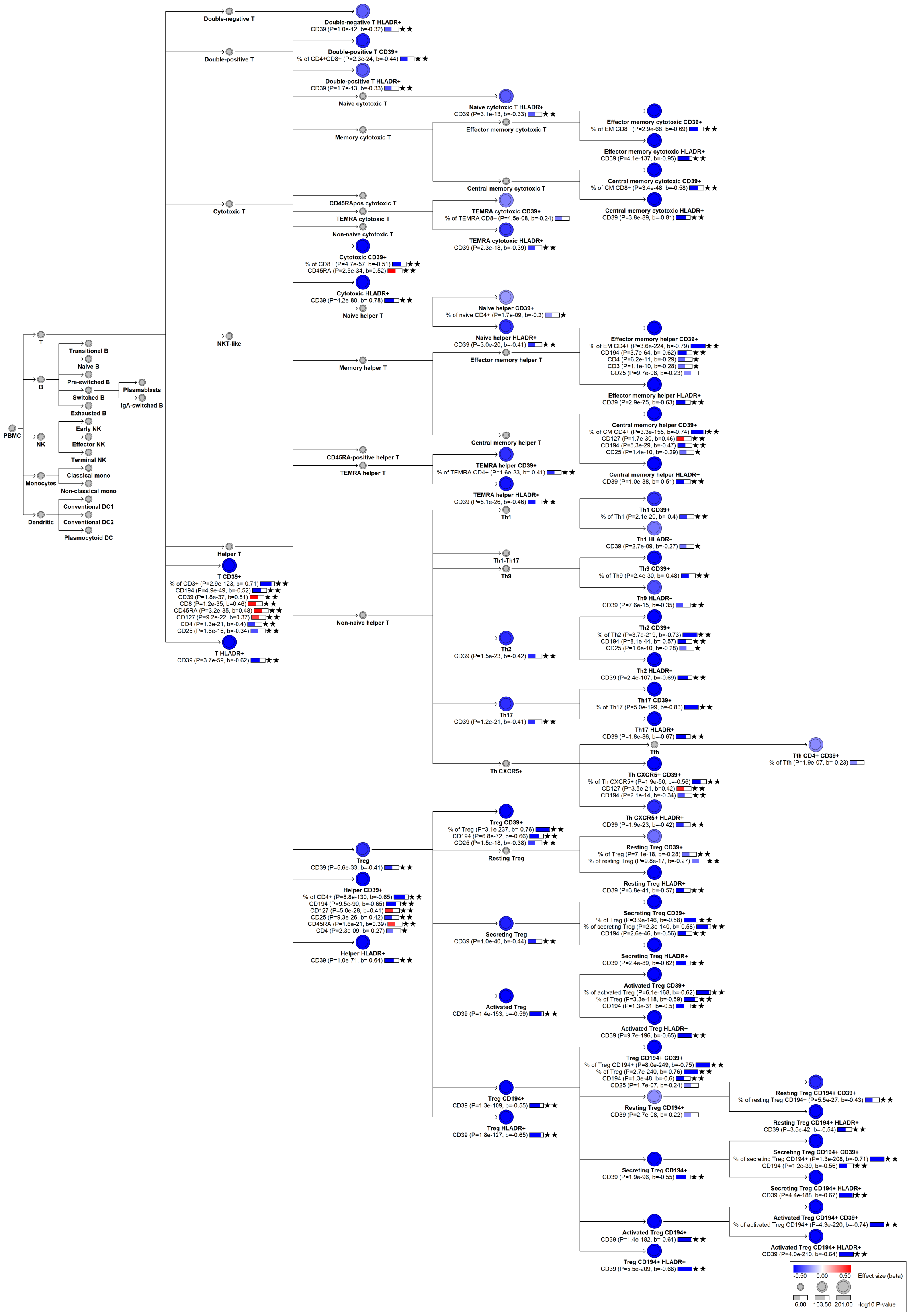

PBMC
T
B
NK
Monocytes
Dendritic
Transitional B
Naive B
Pre-switched B
Switched B
Exhausted B
Plasmablasts
IgA-switched B
Early NK
Effector NK
Terminal NK
Classical mono
Non-classical mono
Conventional DC1
Conventional DC2
Plasmocytoid DC
Double-negative T
Double-positive T
Cytotoxic T
NKT-like
Helper T
T CD39+
% of CD3+ (P=2.9e-123, b=-0.71) ★★
CD194 (P=4.9e-49, b=-0.52) ★★
CD39 (P=1.8e-37, b=0.51) ★★
CD8 (P=1.2e-35, b=0.46) ★★
CD45RA (P=3.2e-35, b=0.48) ★★
CD127 (P=9.2e-22, b=0.37) ★★
CD4 (P=1.3e-21, b=-0.4) ★★
CD25 (P=1.6e-16, b=-0.34) ★★
T HLADR+
CD39 (P=3.7e-59, b=-0.62) ★★
Double-negative T HLADR+
CD39 (P=1.0e-12, b=-0.32) ★★
Double-positive T CD39+
% of CD4+CD8+ (P=2.3e-24, b=-0.44) ★★
Double-positive T HLADR+
CD39 (P=1.7e-13, b=-0.33) ★★
Naive cytotoxic T
Memory cytotoxic T
CD45RApos cytotoxic T
TEMRA cytotoxic T
Non-naive cytotoxic T
Cytotoxic CD39+
% of CD8+ (P=4.7e-57, b=-0.51) ★★
CD45RA (P=2.5e-34, b=0.52) ★★
Cytotoxic HLADR+
CD39 (P=4.2e-80, b=-0.78) ★★
Naive helper T
Memory helper T
CD45RA-positive helper T
TEMRA helper T
Non-naive helper T
Treg
CD39 (P=5.6e-33, b=-0.41) ★★
Helper CD39+
% of CD4+ (P=8.8e-130, b=-0.65) ★★
CD194 (P=9.5e-90, b=-0.65) ★★
CD127 (P=5.0e-28, b=0.41) ★★
CD25 (P=9.3e-26, b=-0.42) ★★
CD45RA (P=1.6e-21, b=0.39) ★★
CD4 (P=2.3e-09, b=-0.27) ★
Helper HLADR+
CD39 (P=1.0e-71, b=-0.64) ★★
Naive cytotoxic T HLADR+
CD39 (P=3.1e-13, b=-0.33) ★★
Effector memory cytotoxic T
Central memory cytotoxic T
TEMRA cytotoxic CD39+
% of TEMRA CD8+ (P=4.5e-08, b=-0.24)
TEMRA cytotoxic HLADR+
CD39 (P=2.3e-18, b=-0.39) ★★
Naive helper CD39+
% of naive CD4+ (P=1.7e-09, b=-0.2) ★
Naive helper HLADR+
CD39 (P=3.0e-20, b=-0.41) ★★
Effector memory helper T
Central memory helper T
TEMRA helper CD39+
% of TEMRA CD4+ (P=1.6e-23, b=-0.41) ★★
TEMRA helper HLADR+
CD39 (P=5.1e-26, b=-0.46) ★★
Th1
Th1-Th17
Th9
Th2
CD39 (P=1.5e-23, b=-0.42) ★★
Th17
CD39 (P=1.2e-21, b=-0.41) ★★
Th CXCR5+
Treg CD39+
% of Treg (P=3.1e-237, b=-0.76) ★★
CD194 (P=6.8e-72, b=-0.66) ★★
CD25 (P=1.5e-18, b=-0.38) ★★
Resting Treg
Secreting Treg
CD39 (P=1.0e-40, b=-0.44) ★★
Activated Treg
CD39 (P=1.4e-153, b=-0.59) ★★
Treg CD194+
CD39 (P=1.3e-109, b=-0.55) ★★
Treg HLADR+
CD39 (P=1.8e-127, b=-0.65) ★★
Effector memory cytotoxic CD39+
% of EM CD8+ (P=2.9e-68, b=-0.69) ★★
Effector memory cytotoxic HLADR+
CD39 (P=4.1e-137, b=-0.95) ★★
Central memory cytotoxic CD39+
% of CM CD8+ (P=3.4e-48, b=-0.58) ★★
Central memory cytotoxic HLADR+
CD39 (P=3.8e-89, b=-0.81) ★★
Effector memory helper CD39+
% of EM CD4+ (P=3.6e-224, b=-0.79) ★★
CD194 (P=3.7e-64, b=-0.62) ★★
CD4 (P=6.2e-11, b=-0.29) ★
CD3 (P=1.1e-10, b=-0.28) ★
CD25 (P=9.7e-08, b=-0.23)
Effector memory helper HLADR+
CD39 (P=2.9e-75, b=-0.63) ★★
Central memory helper CD39+
% of CM CD4+ (P=3.3e-155, b=-0.74) ★★
CD127 (P=1.7e-30, b=0.46) ★★
CD194 (P=5.3e-29, b=-0.47) ★★
CD25 (P=1.4e-10, b=-0.29) ★
Central memory helper HLADR+
CD39 (P=1.0e-38, b=-0.51) ★★
Th1 CD39+
% of Th1 (P=2.1e-20, b=-0.4) ★★
Th1 HLADR+
CD39 (P=2.7e-09, b=-0.27) ★
Th9 CD39+
% of Th9 (P=2.4e-30, b=-0.48) ★★
Th9 HLADR+
CD39 (P=7.6e-15, b=-0.35) ★★
Th2 CD39+
% of Th2 (P=3.7e-219, b=-0.73) ★★
CD194 (P=8.1e-44, b=-0.57) ★★
CD25 (P=1.6e-10, b=-0.28) ★
Th2 HLADR+
CD39 (P=2.4e-107, b=-0.69) ★★
Th17 CD39+
% of Th17 (P=5.0e-199, b=-0.83) ★★
Th17 HLADR+
CD39 (P=1.8e-86, b=-0.67) ★★
Tfh
Th CXCR5+ CD39+
% of Th CXCR5+ (P=1.9e-50, b=-0.56) ★★
CD127 (P=3.5e-21, b=0.42) ★★
CD194 (P=2.1e-14, b=-0.34) ★★
Th CXCR5+ HLADR+
CD39 (P=1.9e-23, b=-0.42) ★★
Tfh CD4+ CD39+
% of Tfh (P=1.9e-07, b=-0.23)
Resting Treg CD39+
% of Treg (P=7.1e-18, b=-0.28) ★★
% of resting Treg (P=9.8e-17, b=-0.27) ★★
Resting Treg HLADR+
CD39 (P=3.8e-41, b=-0.57) ★★
Secreting Treg CD39+
% of Treg (P=3.9e-146, b=-0.58) ★★
% of secreting Treg (P=2.3e-140, b=-0.58) ★★
CD194 (P=2.6e-46, b=-0.56) ★★
Secreting Treg HLADR+
CD39 (P=2.4e-89, b=-0.62) ★★
Activated Treg CD39+
% of activated Treg (P=6.1e-168, b=-0.62) ★★
% of Treg (P=3.3e-118, b=-0.59) ★★
CD194 (P=1.3e-31, b=-0.5) ★★
Activated Treg HLADR+
CD39 (P=9.7e-196, b=-0.65) ★★
Treg CD194+ CD39+
% of Treg CD194+ (P=8.0e-249, b=-0.75) ★★
% of Treg (P=2.7e-240, b=-0.76) ★★
CD194 (P=1.3e-48, b=-0.6) ★★
CD25 (P=1.7e-07, b=-0.24)
Resting Treg CD194+
CD39 (P=2.7e-08, b=-0.22)
Secreting Treg CD194+
CD39 (P=1.9e-96, b=-0.55) ★★
Activated Treg CD194+
CD39 (P=1.4e-182, b=-0.61) ★★
Treg CD194+ HLADR+
CD39 (P=5.5e-209, b=-0.66) ★★
Resting Treg CD194+ CD39+
% of resting Treg CD194+ (P=5.5e-27, b=-0.43) ★★
Resting Treg CD194+ HLADR+
CD39 (P=3.5e-42, b=-0.54) ★★
Secreting Treg CD194+ CD39+
% of secreting Treg CD194+ (P=1.3e-208, b=-0.71) ★★
CD194 (P=1.2e-39, b=-0.56) ★★
Secreting Treg CD194+ HLADR+
CD39 (P=4.4e-188, b=-0.67) ★★
Activated Treg CD194+ CD39+
% of activated Treg CD194+ (P=4.3e-220, b=-0.74) ★★
Activated Treg CD194+ HLADR+
CD39 (P=4.0e-210, b=-0.64) ★★
-0.50 0.00 0.50 Effect size (beta)
6.00 103.50 201.00 -log10 P-value

### chr10:95792577[T→A; EAF 49.648%] (rs10786236, cs_219)

Candidate genes: *ENTPD1*
Diseases associations: NA
Plasma pQTLs: CFP

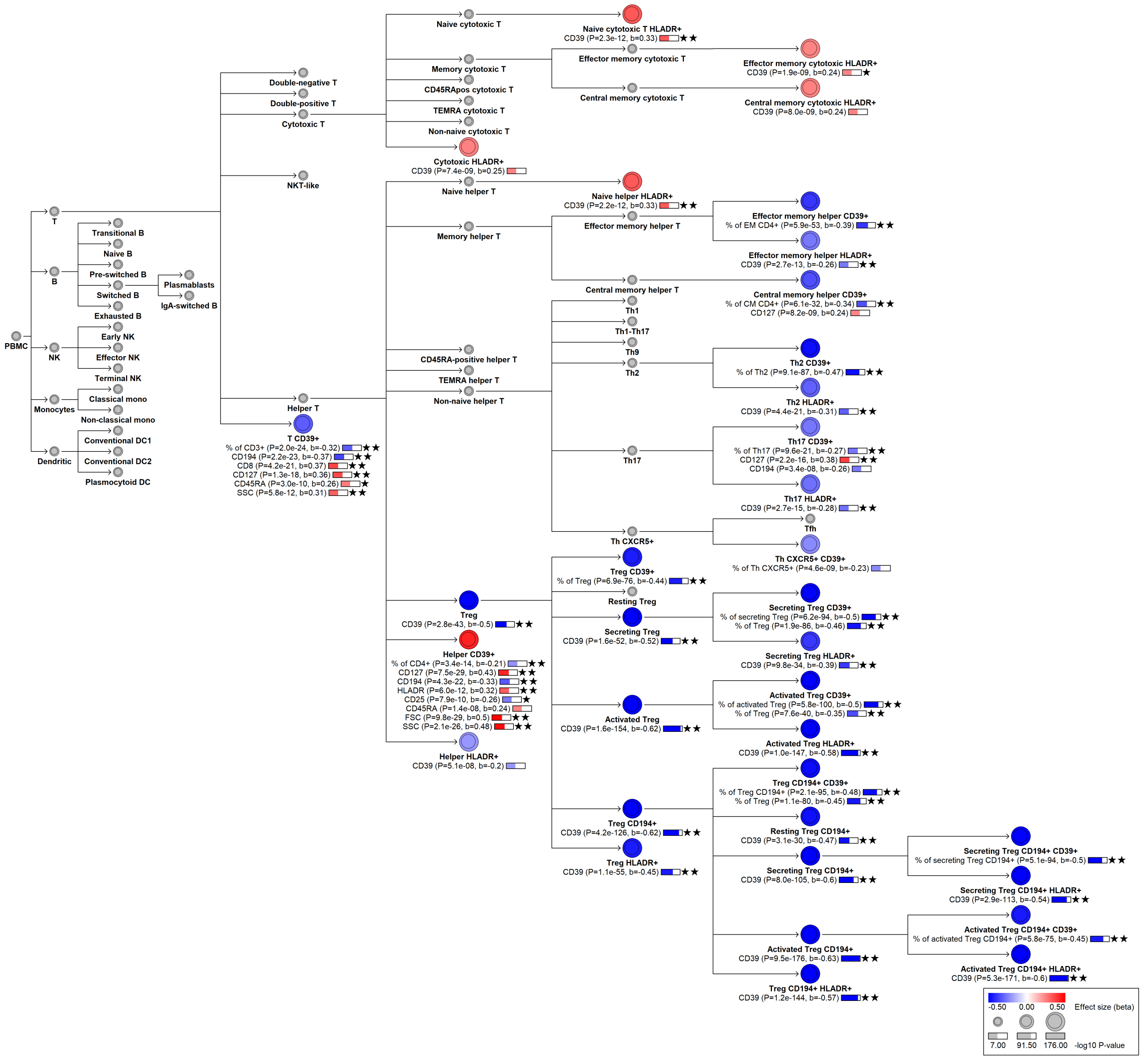

#### chr10:108166738[TTAA→T; EAF 4.374%] (rs144953816, cs_123)

Candidate genes: *NA*
Diseases associations: NA
Plasma pQTLs: NA

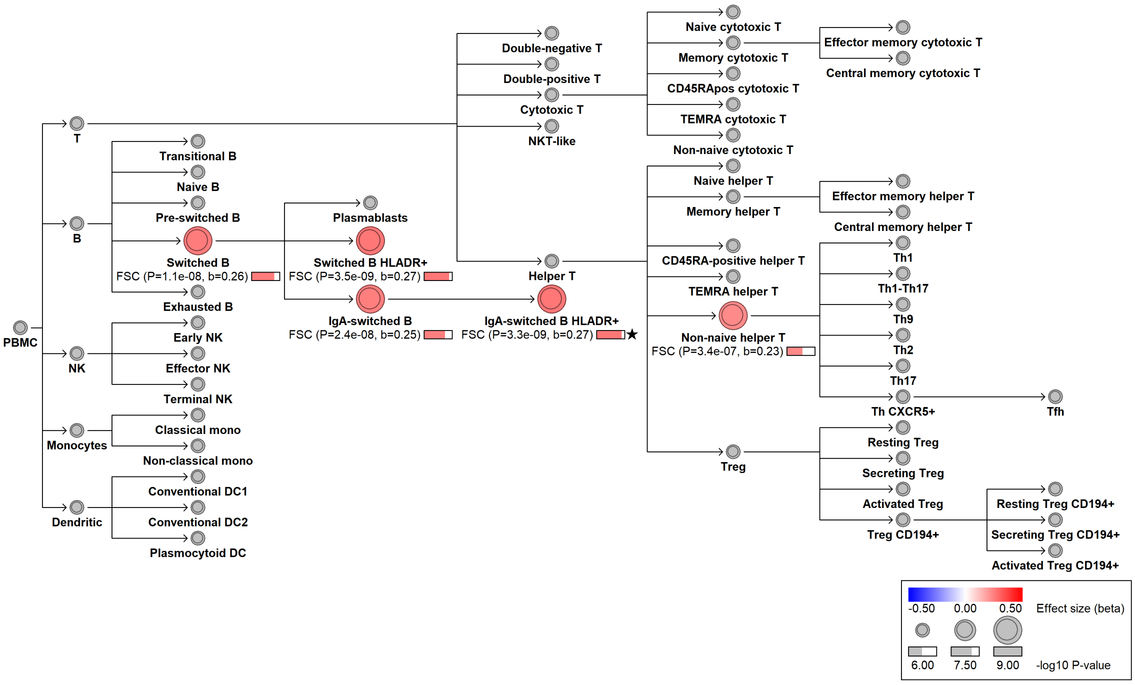

#### chr10:124677669[T→C; EAF 11.537%] (rs10901804, cs_5)

Candidate genes: *FAM53B*
Diseases associations: NA
Plasma pQTLs: NA

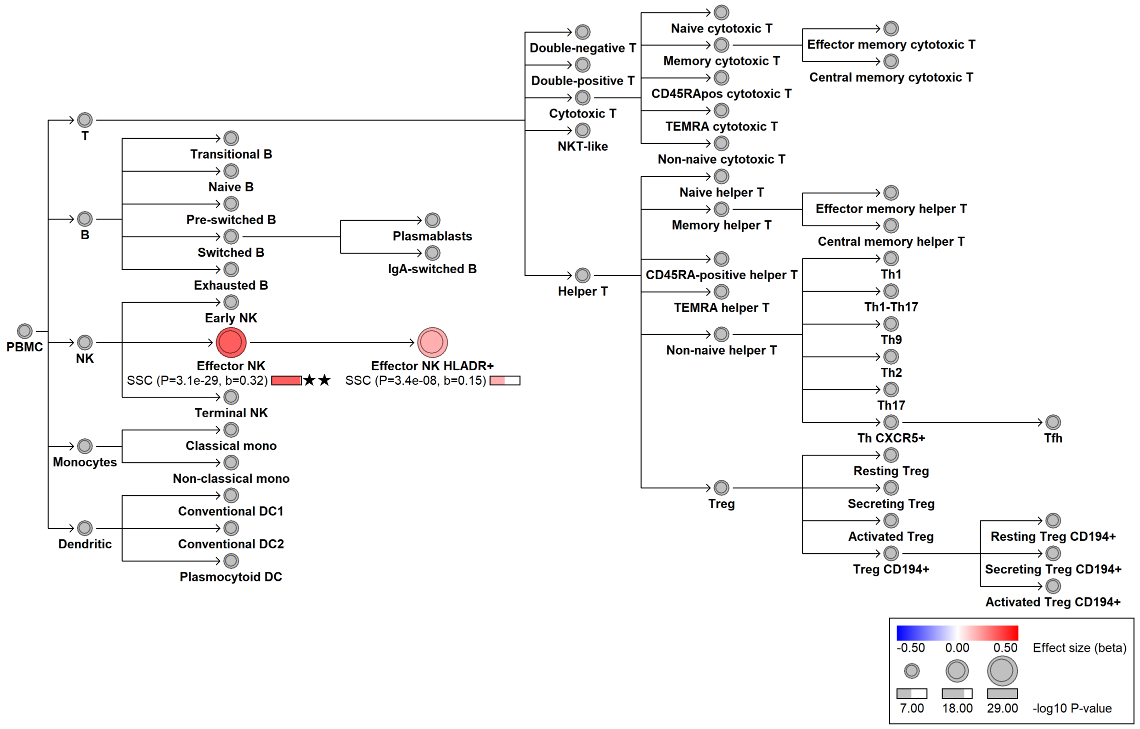

### chr11:2260549[G→A; EAF 12.661%] (rs7929293, cs_163)

Candidate genes: *ASCL2*
Diseases associations: NA
Plasma pQTLs: NA

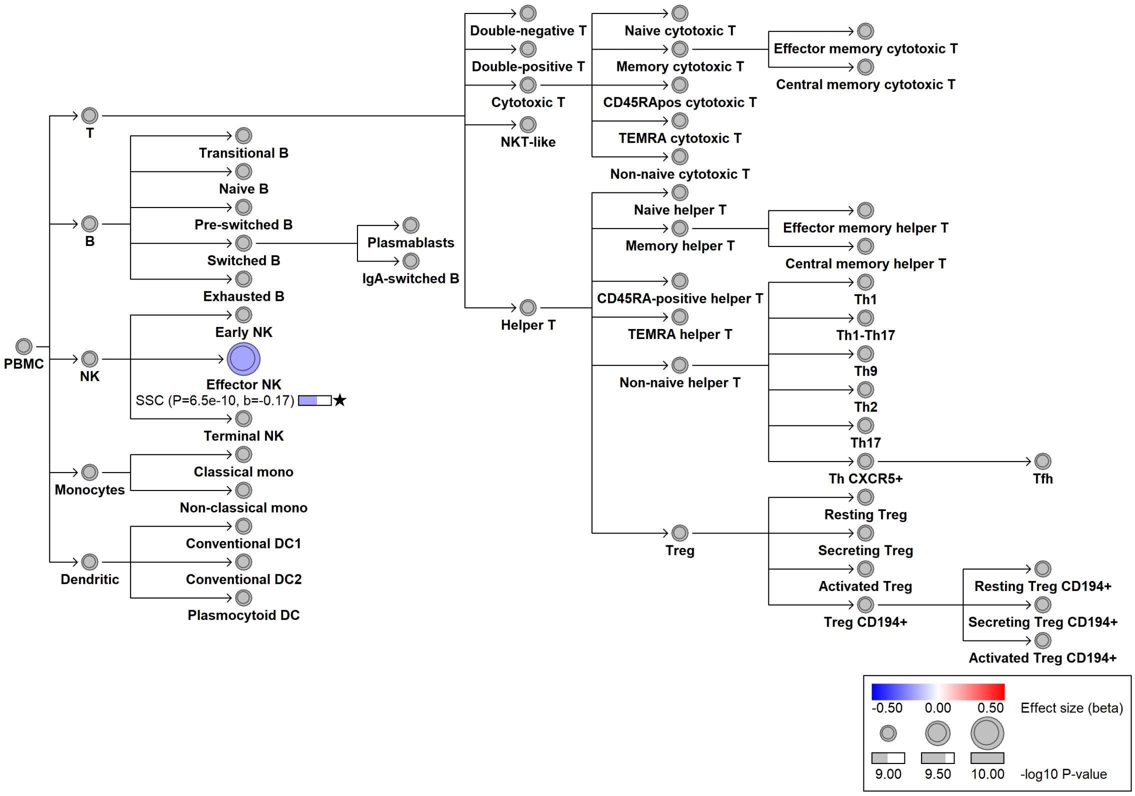

#### chr11:14456981[C→T; EAF 26.924%] (rs2597205, cs_89)

Candidate genes: *GAS5*
Diseases associations: NA
Plasma pQTLs: NA

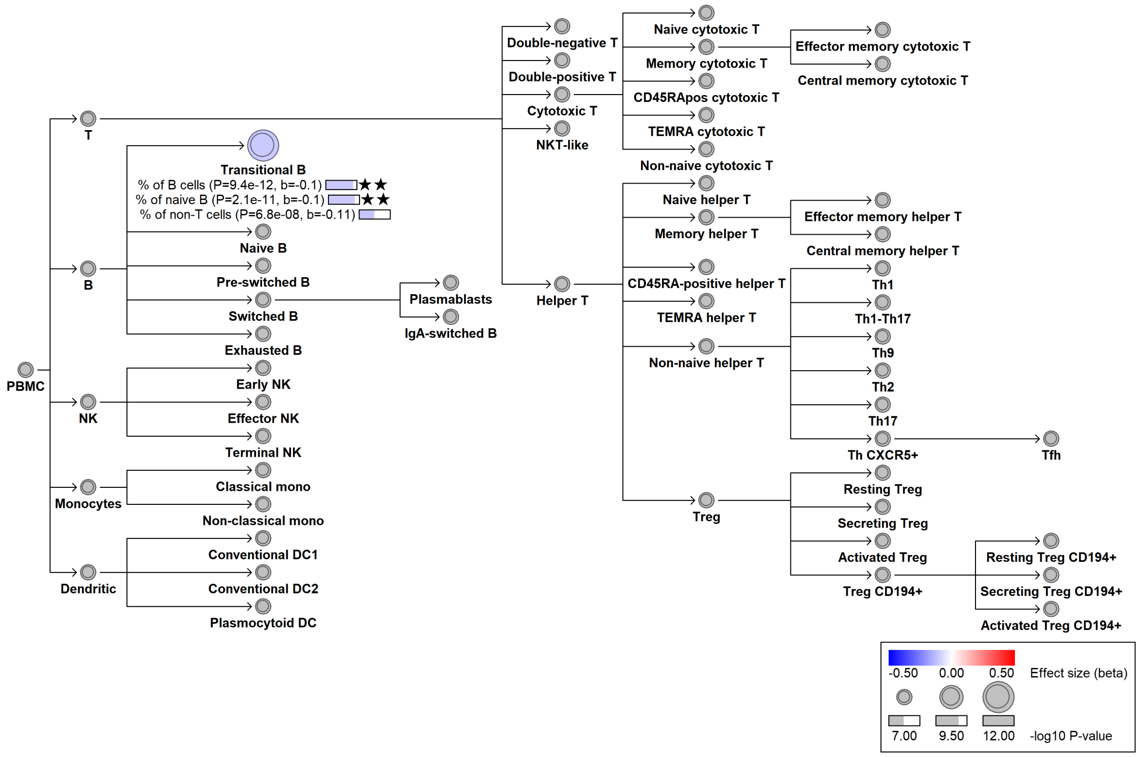

#### chr11:46875895[G→A; EAF 13.885%] (rs2306033, cs_137)

Candidate genes: *NA*
Diseases associations: Anti-NMDA receptor encephalitis, Depression, Glaucoma
Plasma pQTLs: NA

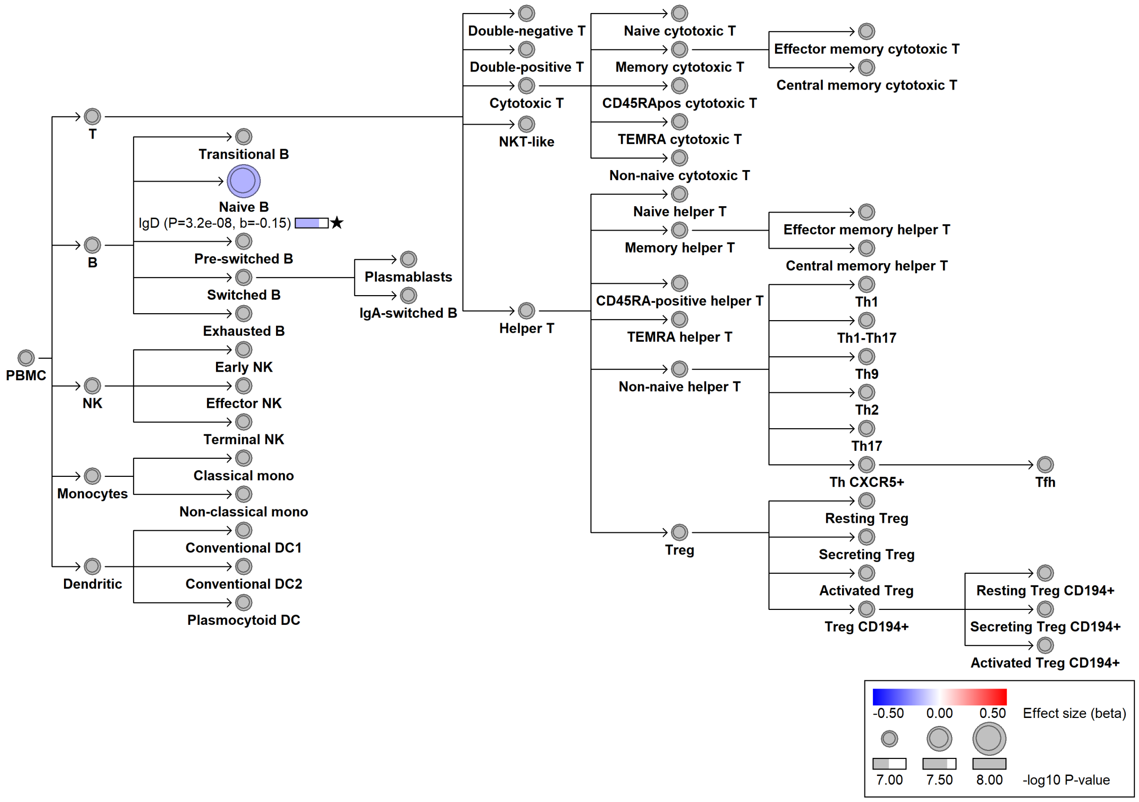

#### chr11:61008737[C→T; EAF 32.537%] (rs11230563, cs_75)

Candidate genes: *CD6*
Diseases associations: Ankylosing spondylitis, Crohn's disease, Inflammatory bowel disease, Primary sclerosing cholangitis, Psoriasis, Ulcerative colitis
Plasma pQTLs: CD6

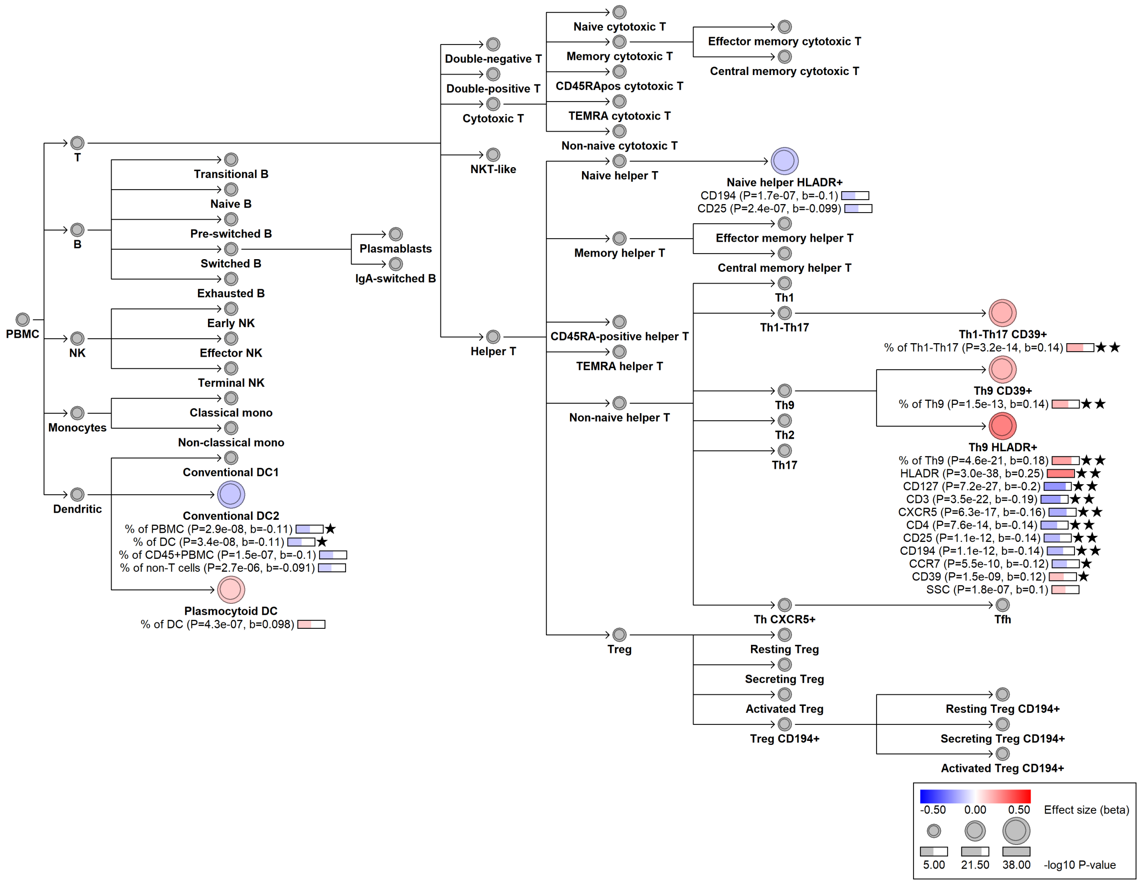

### chr11:112880684[G→A; EAF 3.194%] (rs77738700, cs_27)

Candidate genes: *NCAM1*
Diseases associations: NA
Plasma pQTLs: NA

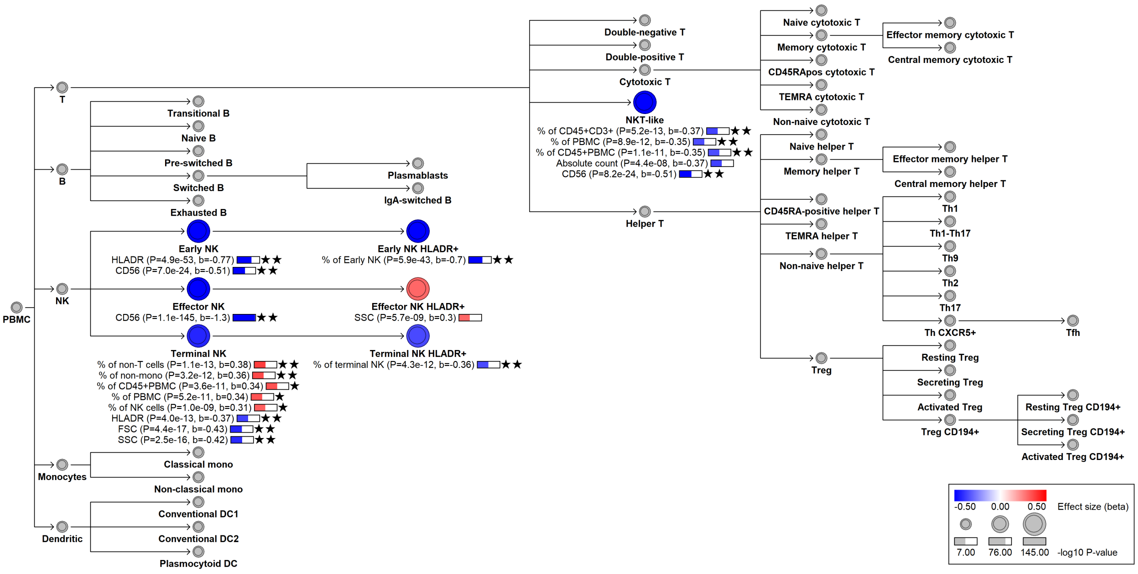

#### chr11:112956992[A→G; EAF 49.046%] (rs4937872, cs_182)

Candidate genes: *NCAM1*
Diseases associations: Depression
Plasma pQTLs: NA

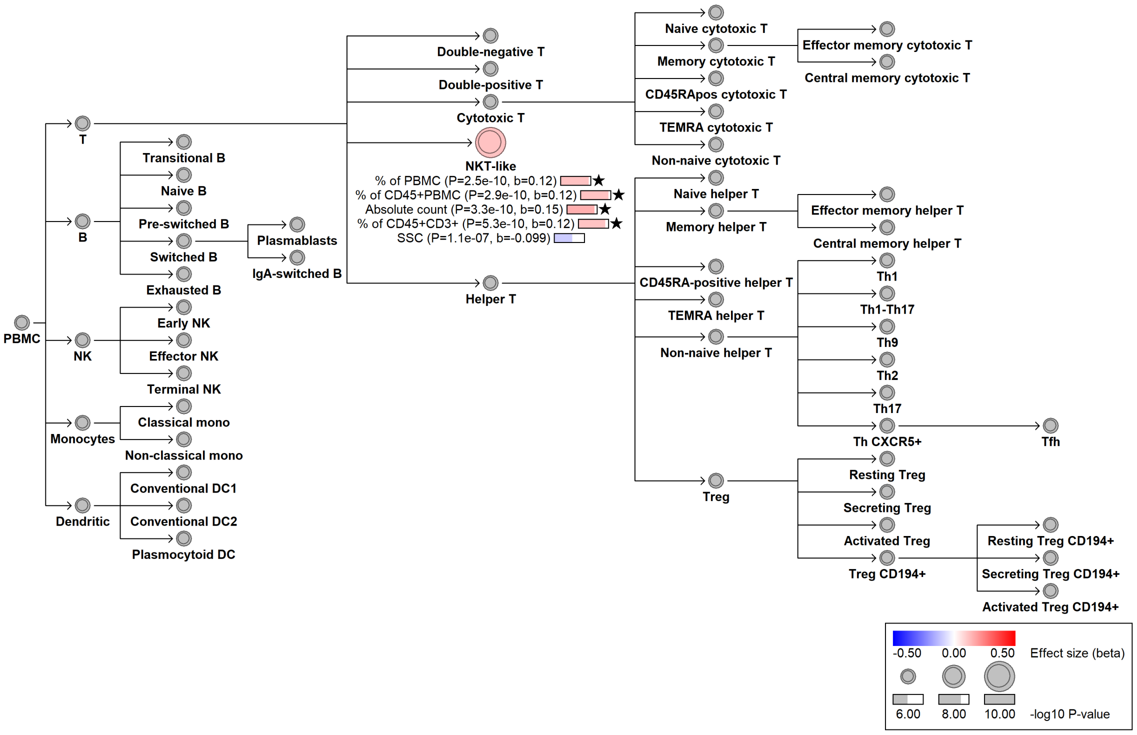

### chr12:6443095[!→GA; EAF 31.772%] (rs34805566, cs_113)

Candidate genes: *CD27*
Diseases associations: NA
Plasma pQTLs: NA

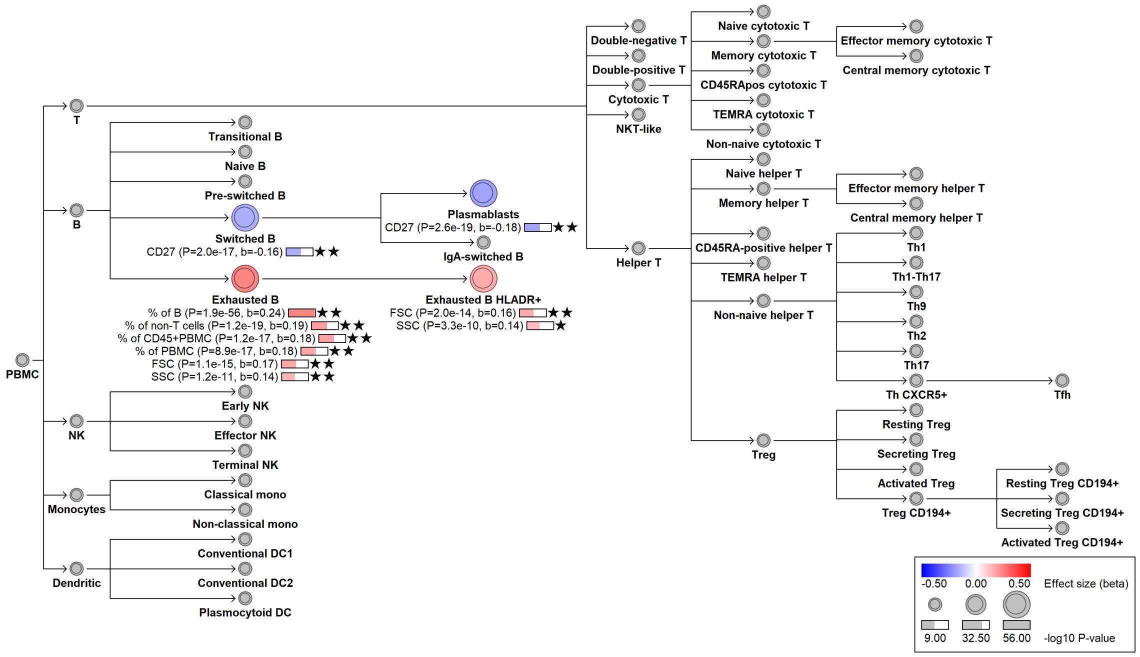

#### chr12:6445462[G→A; EAF 21.595%] (rs25680, cs_106)

Candidate genes: *CD27*
Diseases associations: NA
Plasma pQTLs: CD27

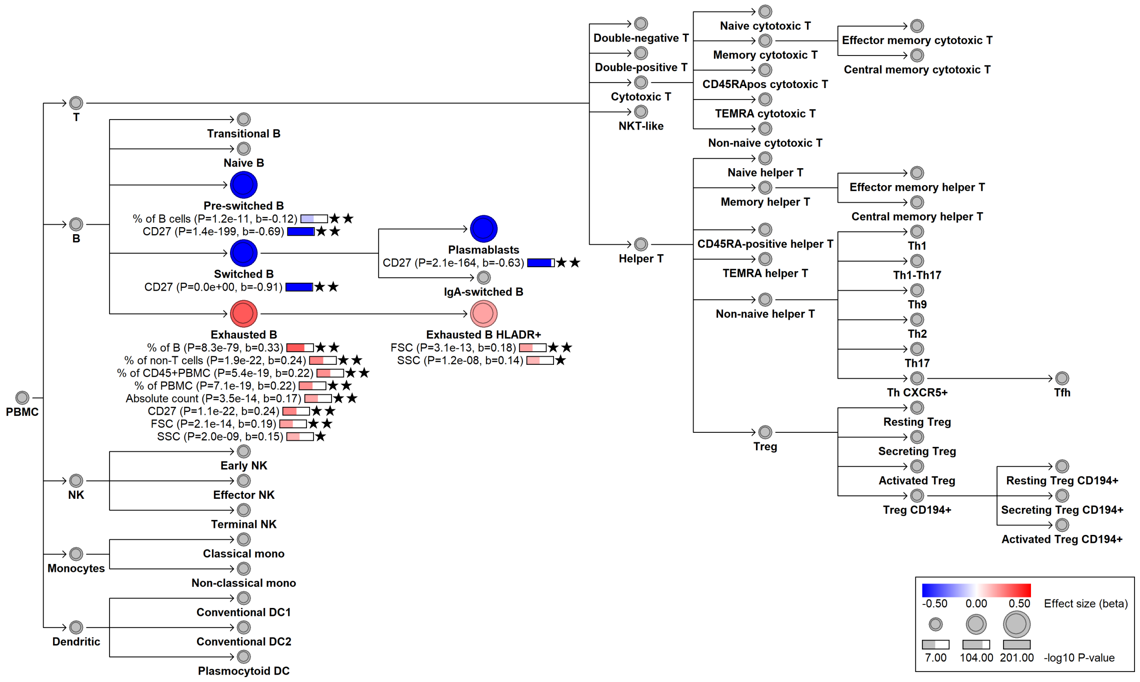

#### chr12:6471148[A→G; EAF 21.605%] (rs12426815, cs_213)

Candidate genes: *CD27*
Diseases associations: NA
Plasma pQTLs: NA

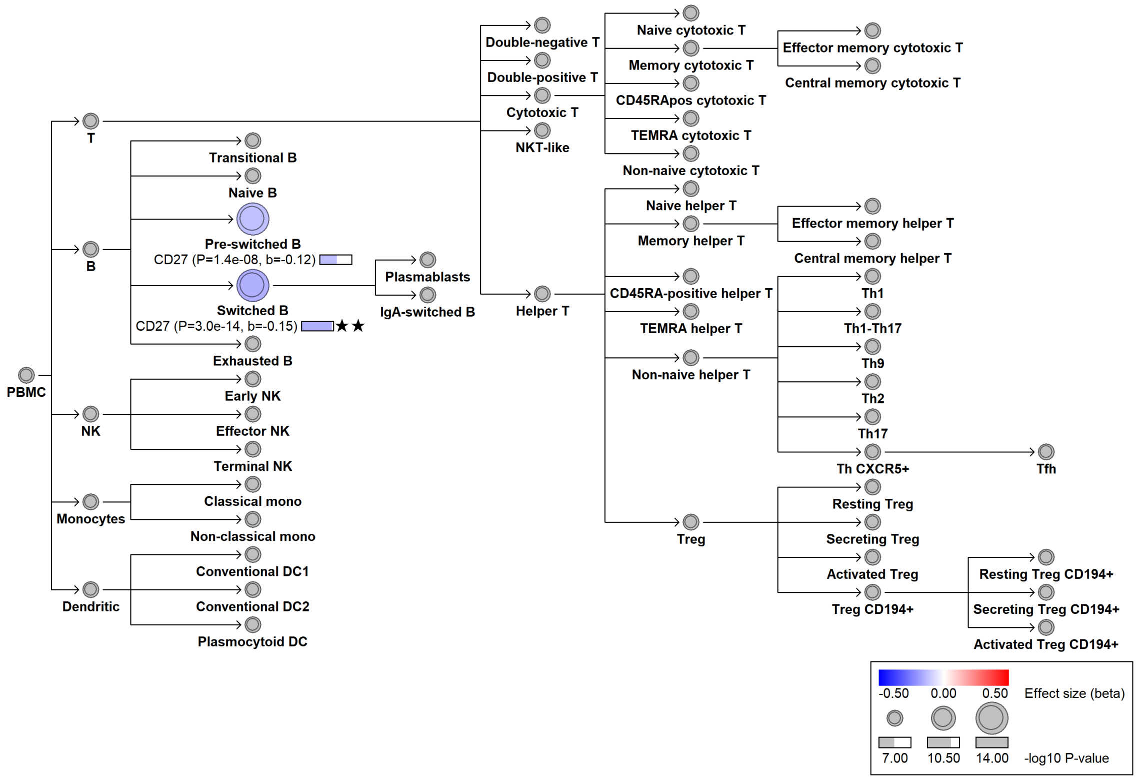

#### chr12:6787028[G→A; EAF 38.239%] (rs1922452, cs_142)

Candidate genes: *CD4*
Diseases associations: NA
Plasma pQTLs: NA

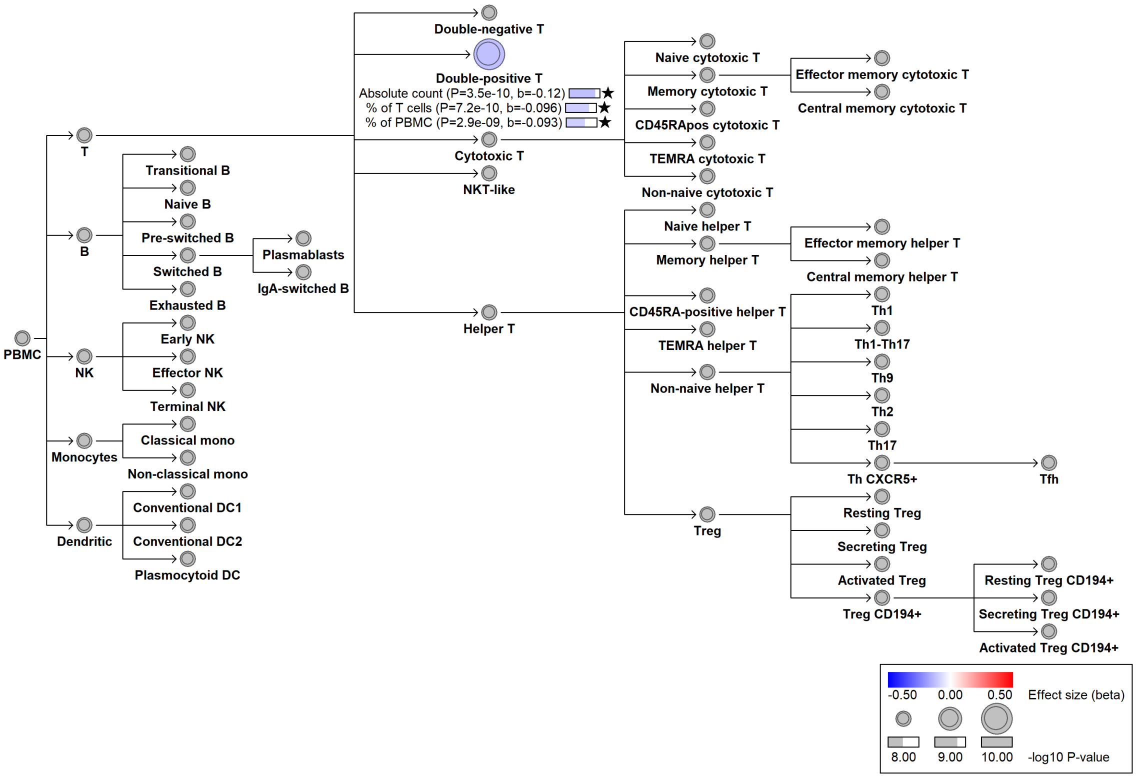

### chr12:6791968[C→T; EAF 32.02%] (rs2707212, cs_22)

Candidate genes: *CD4*
Diseases associations: NA
Plasma pQTLs: NA

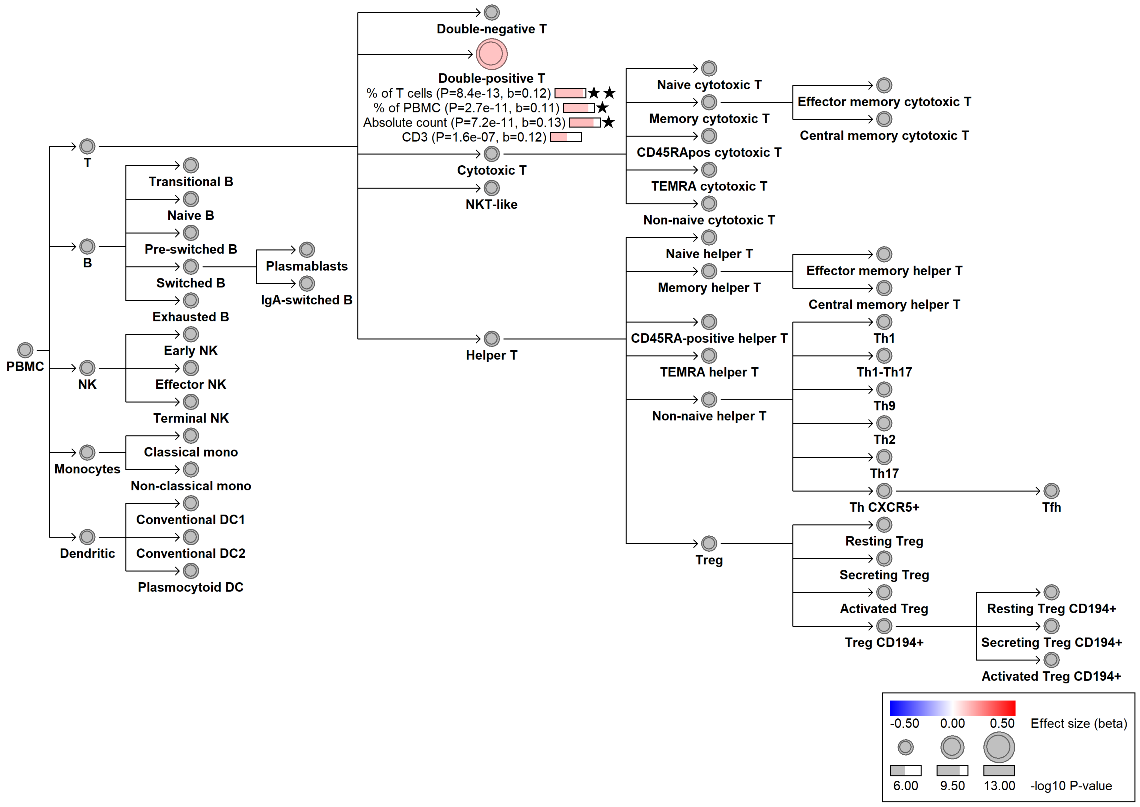

#### chr12:6792800[G→A; EAF 0.506%] (rs117035666, cs_50)

Candidate genes: *CD4*
Diseases associations: NA
Plasma pQTLs: NA

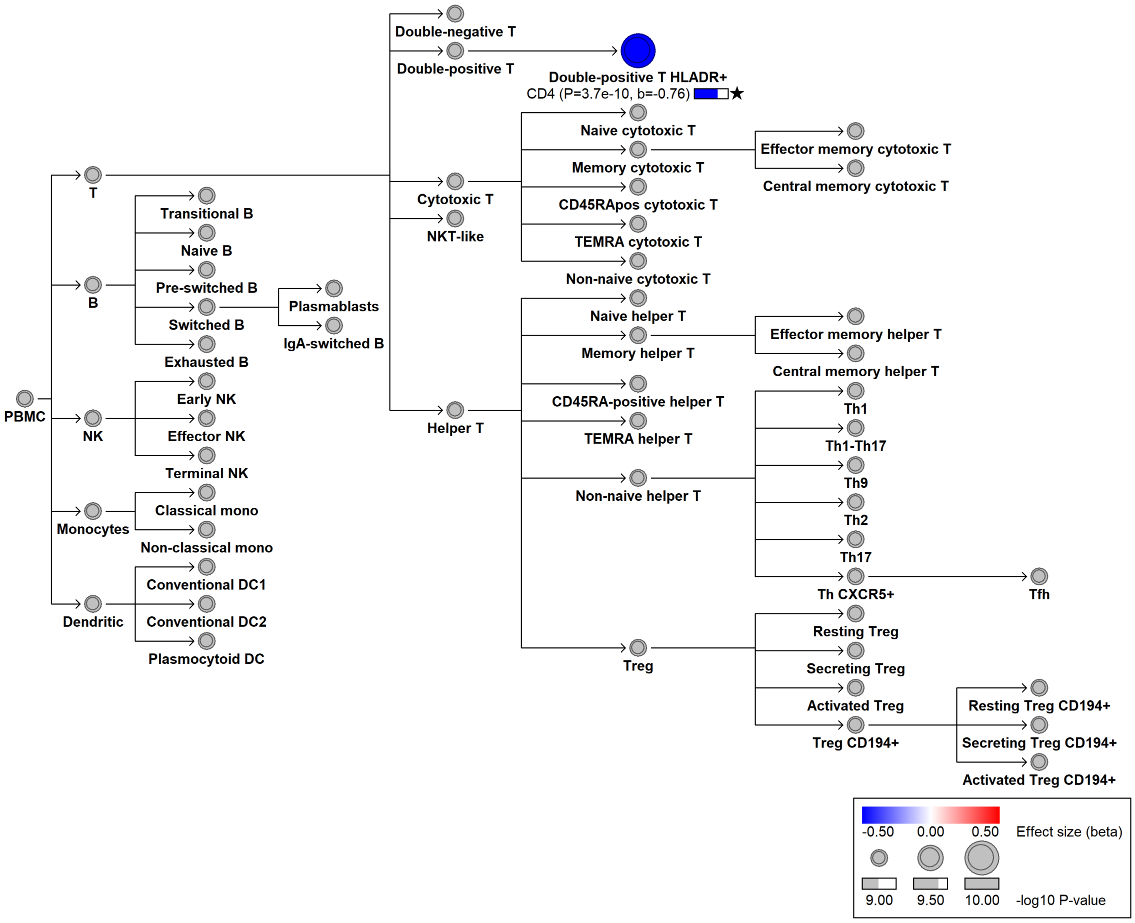

#### chr12:9753255[A→C; EAF 38.369%] (rs917911, cs_84)

Candidate genes: *CD69*
Diseases associations: Type 1 diabetes
Plasma pQTLs: NA

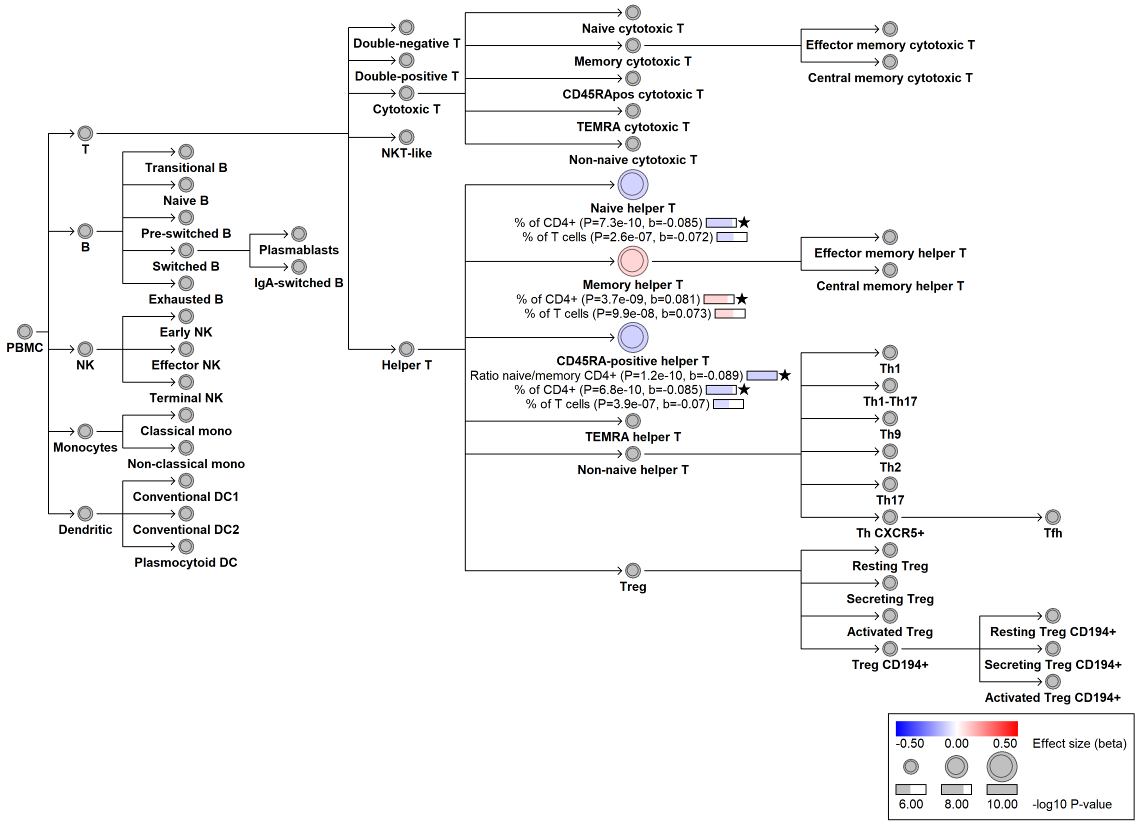

#### chr12:10408358[C→T; EAF 31.645%] (rs2617170, cs_103)

Candidate genes: *KLRC1, KLRC3, KLRC4, KLRK1, YBX3*
Diseases associations: Behcet's disease, Psoriasis
Plasma pQTLs: GZMB, KLRK1

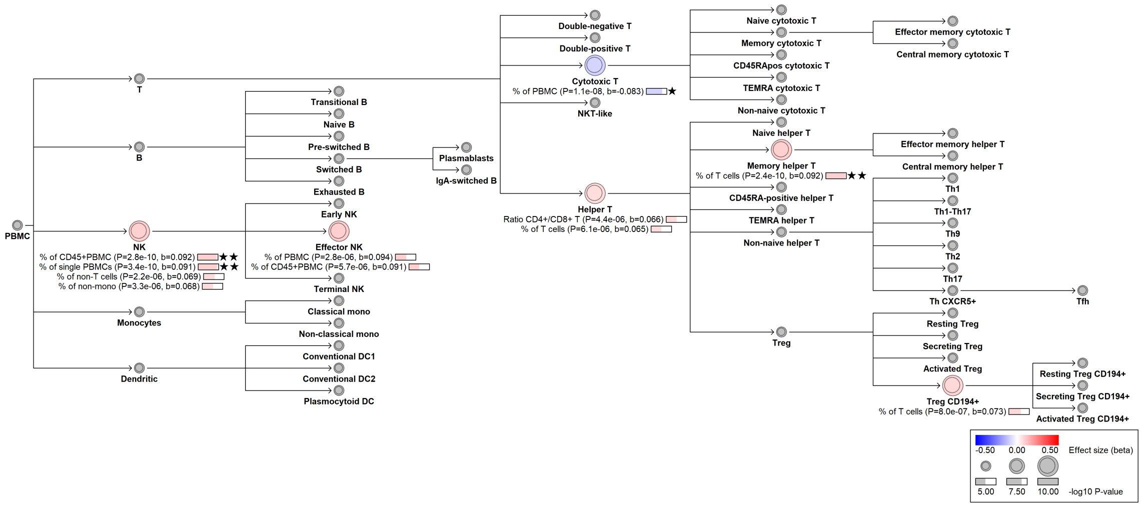

#### chr12:69298386[C→G; EAF 46.718%] (rs317678, cs_184)

Candidate genes: *LYZ*
Diseases associations: NA
Plasma pQTLs: NA

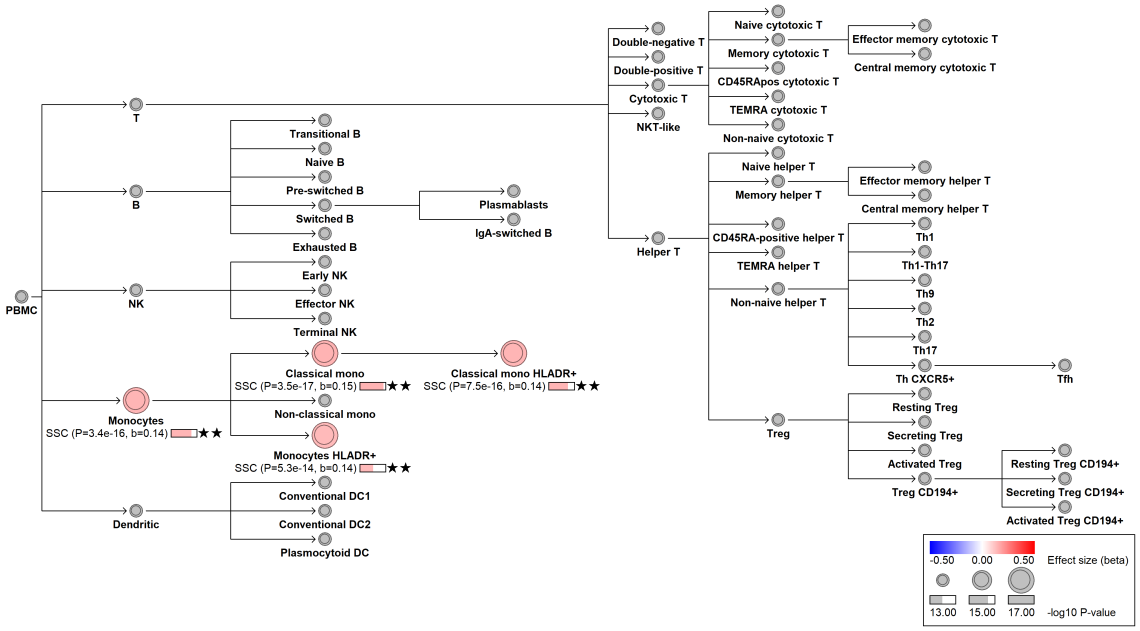

### chr12:69350234[C→A; EAF 6.654%] (rs1800973, cs_187)

Candidate genes: *LYZ*
Diseases associations: NA
Plasma pQTLs: NA

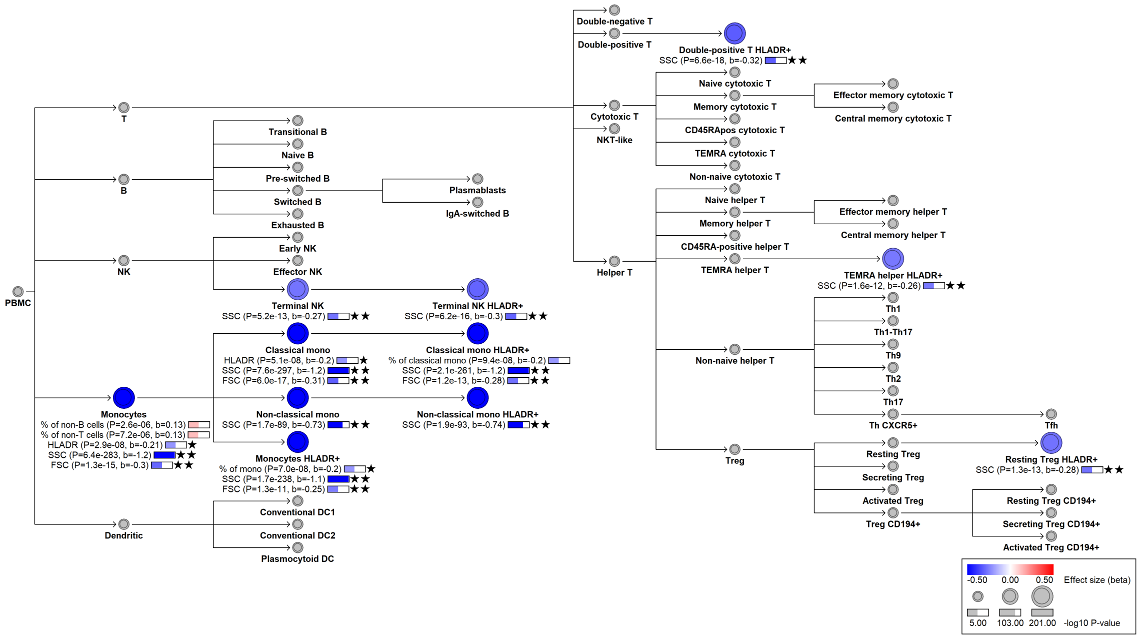

### chr12:69362018[!→TGTTTTTTTTTG; EAF 2.834%] (rs35684976, cs_185)

Candidate genes: *LYZ*
Diseases associations: NA
Plasma pQTLs: NA

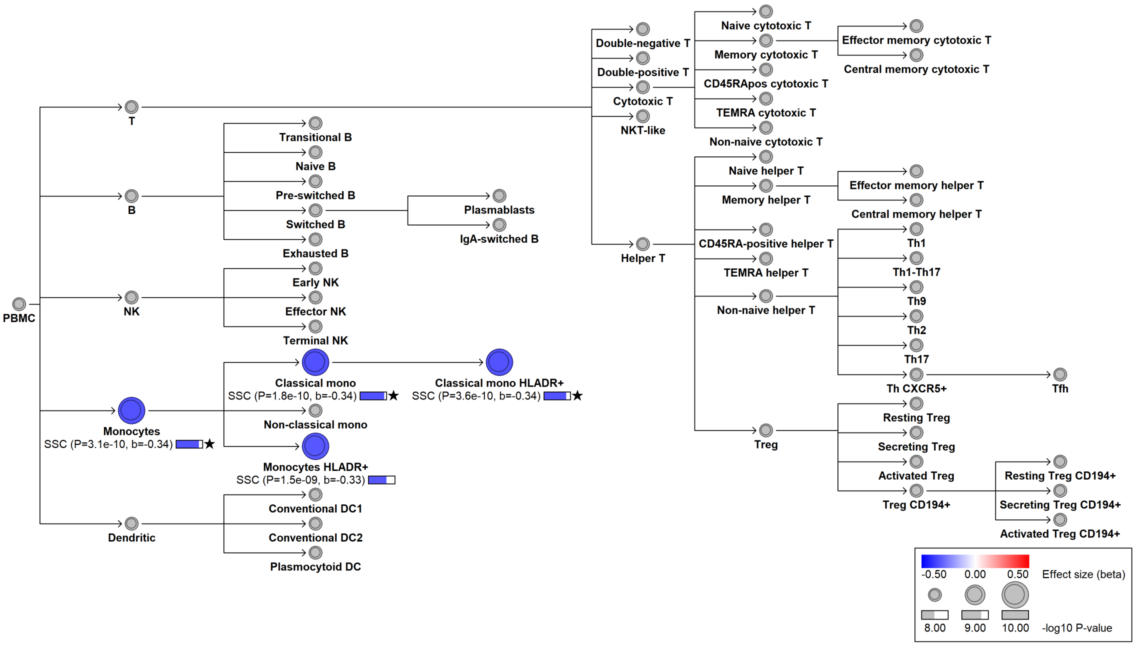

#### chr12:94832806[G→A; EAF 2.936%] (rs118101646, cs_224)

Candidate genes: *NA*
Diseases associations: NA
Plasma pQTLs: NA

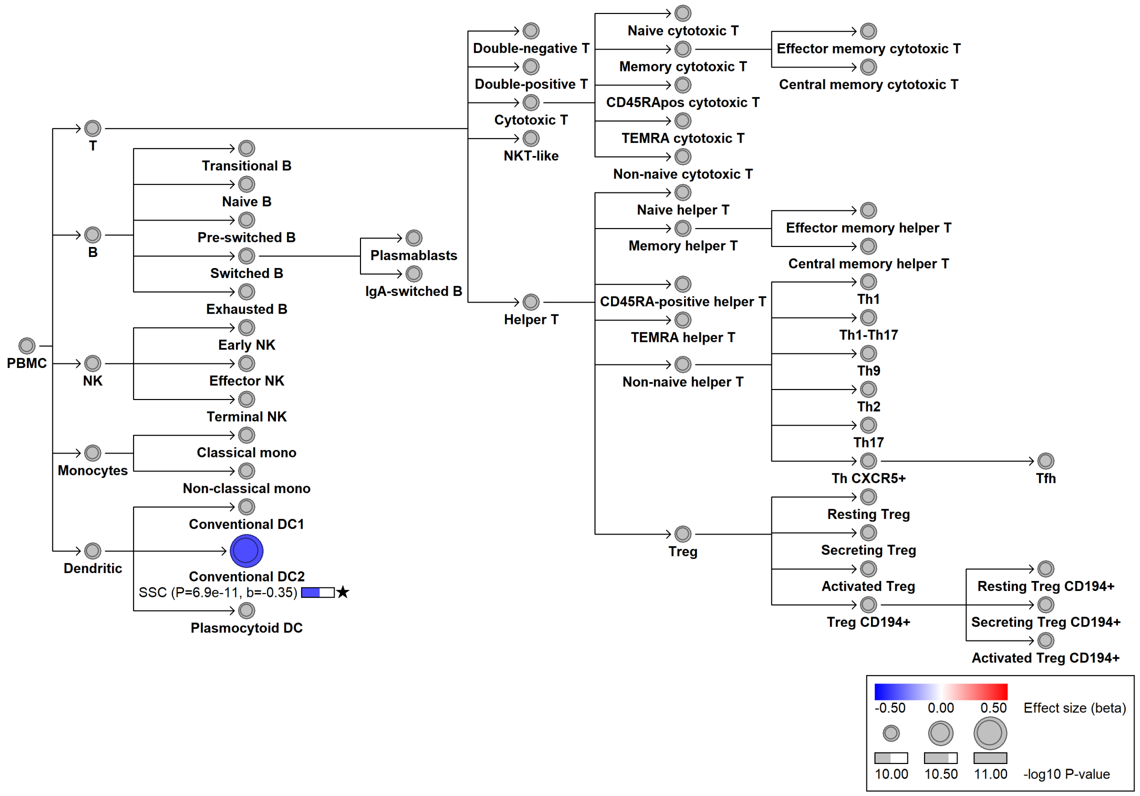

##### chr12:111446804[C→T; EAF 46.218%] (rs3184504, cs_194)

Candidate genes: *SH2B3*
Diseases associations: Allergy, Ankylosing spondylitis, Asthma, Autoimmune thyroiditis, Cardiovascular disease, Celiac disease, Chronic diseases of tonsils, Colorectal cancer, Crohn's disease, Eczema, Endometrial cancer, Inflammatory bowel disease, Multiple sclerosis, Myeloproliferative neoplasms, Preeclampsia, Primary sclerosing cholangitis, Psoriasis, Rheumatoid arthritis, Sarcoidosis, Systemic lupus erythematosus, Type 1 diabetes, Ulcerative colitis, Vitiligo
Plasma pQTLs: AAMDC, AARSD1, ABRAXAS2, ACOT13, ACYP1, ADA, ADAM15, ADAM22, ADGRE1, ADGRE2, ADGRE5, ADGRF5, ADPGK, AGRP, AKT1S1, AKT2, ALDH5A1, AMIGO2, ANXA4, AP1G2, AP2B1, AP3B1, APEX1, APOD, APOL3, ARF6, ARHGAP45, ARHGEF1, ARL2BP, ASAH1, ASPSCR1, ASRGL1, ATG16L1, ATG4A, ATOX1, ATP6V1G1, ATXN2L, ATXN3, B2M, BACH1, BAG4, BAP18, BAX, BCL2, BCL2L1, BCR, BECN1, BLOC1S3, BMP10, BOLA1, BOLA2, BRAP, BTN3A2, BTN3A3, C11orf87, C1QA, C1QTNF3, C5, C7orf50, CACYBP, CALCOCO2, CASC3, CASP3, CASP7, CASP8, CBLN1, CC2D1A, CCDC134, CCDC80, CCL17, CCL18, CCL19, CCL21, CCL22, CCL3, CCL4, CD101, CD160, CD163, CD1C, CD244, CD28, CD2AP, CD300A, CD300C, CD300E, CD3E, CD4, CD40, CD40LG, CD48, CD5, CD5L, CD6, CD69, CD7, CD72, CD74, CD79B, CD84, CD86, CD8A, CD97, CDC26, CDC37, CDC42BPB, CDCP1, CDH17, CDH3, CDH5, CDKN2D, CEACAM8, CECR1, CEP170, CEP20, CETN3, CFP, CHAC2, CHM, CHMP6, CHST11, CHST12, CIAPIN1, CIT, CLC, CLEC1B, CLEC4A, CLEC4D, CLEC6A, CLSTN3, CLU, CMC1, CMIP, CNP, CNPY4, COL1A1, COL28A1, COL3A1, COL5A1, COLEC12, COMMD1, COMP, COMT, COX6C, CPA4, CPE, CPPED1, CPVL, CPXM1, CR1, CR2, CRADD, CREG1, CRELD2, CRH, CRKL, CRTAM, CRYBB1, CRYZL1, CSF1R, CSF2, CSF2RA, CSF3R, CSNK2A1, CST3, CST7, CSTB, CTSD, CTSS, CWC15, CXCL1, CXCL10, CXCL11, CXCL12, CXCL13, CXCL16, CXCL6, CXCL9, CYB5R2, CYR61, Composite immunoglobulin trait (IgA*IgG*IgM), Composite immunoglobulin trait IgA*IgG, Composite immunoglobulin trait IgA/IgG, Composite immunoglobulin trait IgA/IgM, Composite immunoglobulin trait IgM/(IgA*IgG), DAAM1, DBI, DBNL, DCP1A, DCTD, DDHD2, DDX58, DEFA1, DFFA, DGKA, DLL1, DMP1, DNAJB1, DNAJB14, DNAJB2, DNAJC21, DNAJC6, DNM1, DNPH1, DOK1, DPEP2, DPY30, DRG2, DSC2, DTYMK, DXO, DYNLT1, EBI3, ECHS1, EDA, EDIL3, EEF1D, EFCAB2, EFNA4, EGLN1, EHD3, EIF2AK2, EIF4B, EIF4E, EIF4EBP1, EIF4G1, ELAC1, ELOA, ELOB, ENO2, EPO, EREG, ERMAP, ERP29, ESM1, ESYT2, F2R, FABP5, FADD, FAIM3, FAM13A, FARSA, FASLG, FCAMR, FCGR2A, FCGR3A, FCGR3B, FCRL2, FCRL3, FCRL5, FCRL6, FGD3, FGFBP1, FJX1, FKBP14, FKBP4, FKBP5, FLRT2, FLT3, FLT3LG, FMOD, FNTA, FOXO1, FRZB, FUS, FUT8, FXN, FXYD5, GADD45GIP1, GALNT10, GALNT16, GAS7, GBP1, GBP2, GCC1, GCLM, GCNT1, GDF2, GGACT, GGCT, GIPC3, GLB1, GLO1, GLOD4, GMFG, GMPR, GMPR2, GNE, GOLGA3, GOLM2, GORASP2, GP1BA, GP1BB, GP5, GP6, GPKOW, GRAP2, GRHPR, GRPEL1, GRSF1, GSN, GTPBP2, GZMA, GZMB, GZMM, HARS1, HCLS1, HDGF, HDGFL2, HGF, HHEX, HHIP, HLA, HMCN2, HMGCL, HNRNPUL1, HPCAL1, HPSE, HS1BP3, HS6ST1, HSBP1, HSPG2, ICAM1, ICAM2, ICAM4, IDO1, IDUA, IFNL1, IGBP1, IGF1R, IGFLR1, IL12A, IL12B, IL12RB1, IL15, IL15RA, IL16, IL18, IL18BP, IL1RN, IL22RA2, IL27RA, IL2RA, IL2RB, IL2RG, IL32, IL6R, ILKAP, IMMT, IMPACT, ING1, INPP5D, INSR, IPCEF1, IRAK4, IST1, ITGA6, ITGAL, ITGAM, ITGB1BP2, ITGB6, ITGB7, ITPA, Immunoglobulin A, Immunoglobulin G, JAM3, JPT2, KAZN, KIR2DL3, KIR2DL4, KIR2DS2, KITLG, KLRB1, KLRD1, KLRK1, L1CAM, LACTB2, LAG3, LAIR1, LAIR2, LAT, LATS1, LBR, LDLRAP1, LETM1, LGALS3BP,

LGALS8, LHPP, LILRA2, LILRA4, LILRA5, LILRB4, LMNB2, LONP1, LRCH4, LRP11, LRRC59, LRRFIP1, LTA, LTB, LY75, LY9, LYZ, LZTFL1, MAD1L1, MANF, MAP4K5, MAPKAPK2, MARS1, MATN2, MAX, MCAM, MCFD2, MDH1, MDK, MECR, MESD, METAP2, MICB, MIF, MINK1, MMP1, MMP12, MOCS2, MPI, MPIG6B, MRC1, MSRA, MSTN, MTDH, MTHFSD, MTSS1, MTSS2, MYDGF, MYH9, MYOC, NAA80, NAGK, NAP1L4, NARS1, NCK2, NCMAP, NCR1, NCR3, NEDD4L, NEGR1, NELL2, NFE2, NFKB1, NFU1, NHLRC3, NID1, NIT1, NMT1, NOS3, NOTCH1, NPC2, NPL, NPM1, NRGN, NSFL1C, NT5C, NT5E, NTRK2, NTRK3, NUB1, NUCB2, NUDC, NUDT16, NUDT5, NUMB, OGA, OGFR, OIT3, OPLAH, OSM, OTUD7B, OXCT1, PAGR1, PAPPA, PARK7, PARP1, PAXX, PCBP2, PCDH17, PCDH9, PCDHB15, PCOLCE2, PDAP1, PDCD5, PDE5A, PDIA4, PEAR1, PEBP1, PECAM1, PER3, PF4, PGD, PGF, PHACTR2, PHYKPL, PILRA, PILRB, PKD1, PLA2G4A, PLAU, PLAUR, PLCB2, PLEKHO1, PLPBP, PLXNA4, PLXNB3, PLXNC1, PNMA1, POLR2F, POMGNT2, POR, PPCDC, PPIB, PPM1F, PPME1, PPP1CC, PPP1R14A, PPP1R2, PPP2R5A, PRCP, PRDX3, PRG2, PRG3, PRKAR1A, PRKAR2A, PRKRA, PRSS57, PSAP, PSIP1, PSME1, PSME2, PSMG3, PTGES2, PTPRC, PTPRH, PTPRK, PTRHD1, QSOX2, RAB10, RABEP1, RABGAP1L, RANBP1, RBBP4, RBP5, RBPMS2, REEP4, RETN, RGS10, RILP, RILPL2, RNASE3, RNASE6, RNASET2, RNF149, RPE, RRM2, RSPO1, RTN4IP1, RWDD1, S100A10, S100A11, S100A4, SAMD9L, SARG, SCAMP3, SCARA5, SCARF1, SCLY, SCPEP1, SEC31A, SEL1L, SELL, SELP, SELPLG, SEMA3C, SEMA3E, SEMA3F, SEMA4A, SEMA4D, SEMA6B, SEMA7A, SERPINB1, SERPING1, SFRP4, SH2D1A, SH3BP1, SH3GLB2, SHISA5, SIGLEC1, SIGLEC12, SIGLEC14, SIGLEC5, SIGLEC6, SIGLEC7, SIGLEC8, SIRT2, SIRT3, SIT1, SKAP1, SLA2, SLAMF1, SLAMF6, SLAMF8, SLC9A3R1, SLITRK2, SMAD2, SMNDC1, SMS, SNAP29, SNCA, SNX15, SNX5, SPARCL1, SPART, SPINT2, SPRY2, SRP14, SRSF6, ST13, ST3GAL2, STAMBP, STAT1, STAT2, STAT5B, STC2, STIP1, STMN4, STX4, STX7, STX8, SUGT1, SUMF2, SUSD1, SYAP1, TAC1, TACC3, TADA3, TALDO1, TAPBP, TARBP2, TBCA, TBCB, TBCC, TCOF1, TDGF1, TDP1, TGFB1, THBS3, THBS4, THOP1, THTPA, TIA1, TIGIT, TIMM10, TIMM8A, TIMP3, TINAGL1, TLL1, TLR1, TMED8, TMPO, TMSB10, TNF, TNFAIP2, TNFAIP8L2, TNFRSF10B, TNFRSF11B, TNFRSF13B, TNFRSF14, TNFRSF1B, TNFRSF4, TNFRSF6B, TNFRSF8, TNFSF11, TNFSF12, TNFSF13, TNFSF13B, TNFSF14, TNFSF8, TNIP1, TOP2B, TOR1AIP1, TP53I3, TPD52L2, TPK1, TPR, TRA2B, TREML2, TRIAP1, TRIM21, TRIM24, TRIM25, TRIM58, TSC22D1, TSPYL1, TWF2, TXLNA, TXN, TXNDC5, TXNDC9, TYMP, UBAC1, UBE2L6, UBXN1, UFD1, UROD, USP25, USP8, UST, VAMP8, VAV3, VBP1, VCAM1, VNN2, VPS28, VPS4B, VSIR, VTI1A, WARS, WASHC3, WFDC8, WFIKKN1, XCL1, XIAP, YARS1, YJU2, YTHDF3, YWHAQ, ZBTB17, ZFYVE19, ZHX2

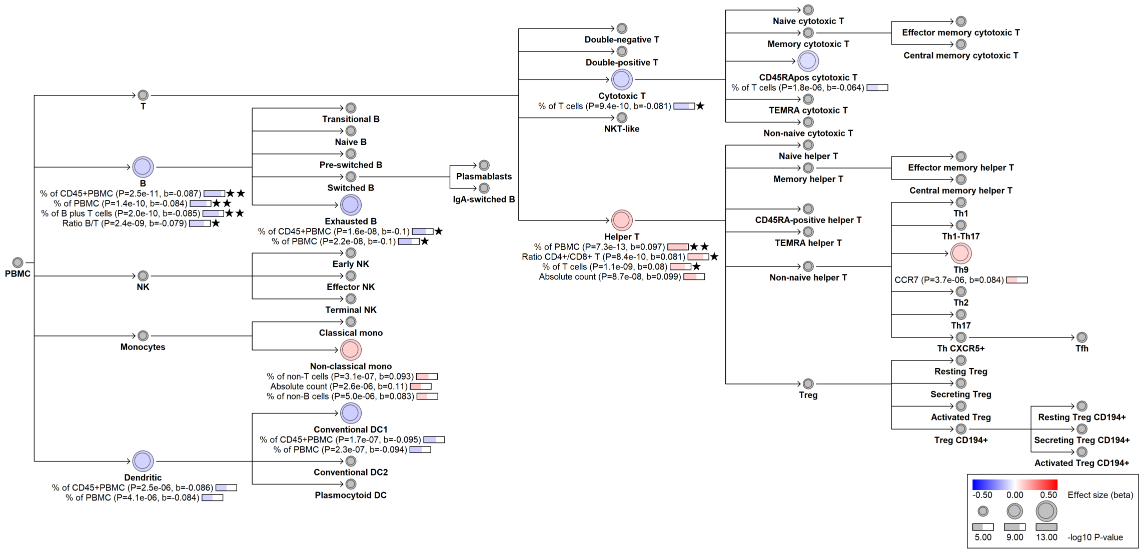
PBMC
T
B
% of CD45+PBMC (P=2.5e-11, b=-0.087) ★★
% of PBMC (P=1.4e-10, b=-0.084) ★★
% of B plus T cells (P=2.0e-10, b=-0.085) ★★
Ratio B/T (P=2.4e-09, b=-0.079) ★
NK
Monocytes
Dendritic
% of CD45+PBMC (P=2.5e-06, b=-0.086)
% of PBMC (P=4.1e-06, b=-0.084)
Transitional B
Naive B
Pre-switched B
Switched B
Exhausted B
% of CD45+PBMC (P=1.6e-08, b=-0.1) ★
% of PBMC (P=2.2e-08, b=-0.1) ★
Plasmablasts
IgA-switched B
Early NK
Effector NK
Terminal NK
Classical mono
Non-classical mono
% of non-T cells (P=3.1e-07, b=0.093)
Absolute count (P=2.6e-06, b=0.11)
% of non-B cells (P=5.0e-06, b=0.083)
Conventional DC1
% of CD45+PBMC (P=1.7e-07, b=-0.095)
% of PBMC (P=2.3e-07, b=-0.094)
Conventional DC2
Plasmocytoid DC
Double-negative T
Double-positive T
Cytotoxic T
% of T cells (P=9.4e-10, b=-0.081) ★
NKT-like
Helper T
% of PBMC (P=7.3e-13, b=0.097) ★★
Ratio CD4+/CD8+ T (P=8.4e-10, b=0.081) ★
% of T cells (P=1.1e-09, b=0.08) ★
Absolute count (P=8.7e-08, b=0.099)
Naive cytotoxic T
Memory cytotoxic T
CD45RApos cytotoxic T
% of T cells (P=1.8e-06, b=-0.064)
TEMRA cytotoxic T
Non-naive cytotoxic T
Effector memory cytotoxic T
Central memory cytotoxic T
Naive helper T
Memory helper T
CD45RA-positive helper T
TEMRA helper T
Non-naive helper T
Treg
Effector memory helper T
Central memory helper T
Th1
Th1-Th17
Th9
CCR7 (P=3.7e-06, b=0.084)
Th2
Th17
Th CXCR5+
Tfh
Resting Treg
Secreting Treg
Activated Treg
Treg CD194+
Resting Treg CD194+
Secreting Treg CD194+
Activated Treg CD194+
-0.50 0.00 0.50 Effect size (beta)
5.00 9.00 13.00 -log10 P-value

#### chr12:120906687[G→T; EAF 43.073%] (rs2701177, cs_205)

Candidate genes: *SPPL3*
Diseases associations: Allergy, Asthma, Depression, Eczema, Systemic lupus erythematosus
Plasma pQTLs: CD22, CSF1R, IL12A, LYVE1, SEMA7A

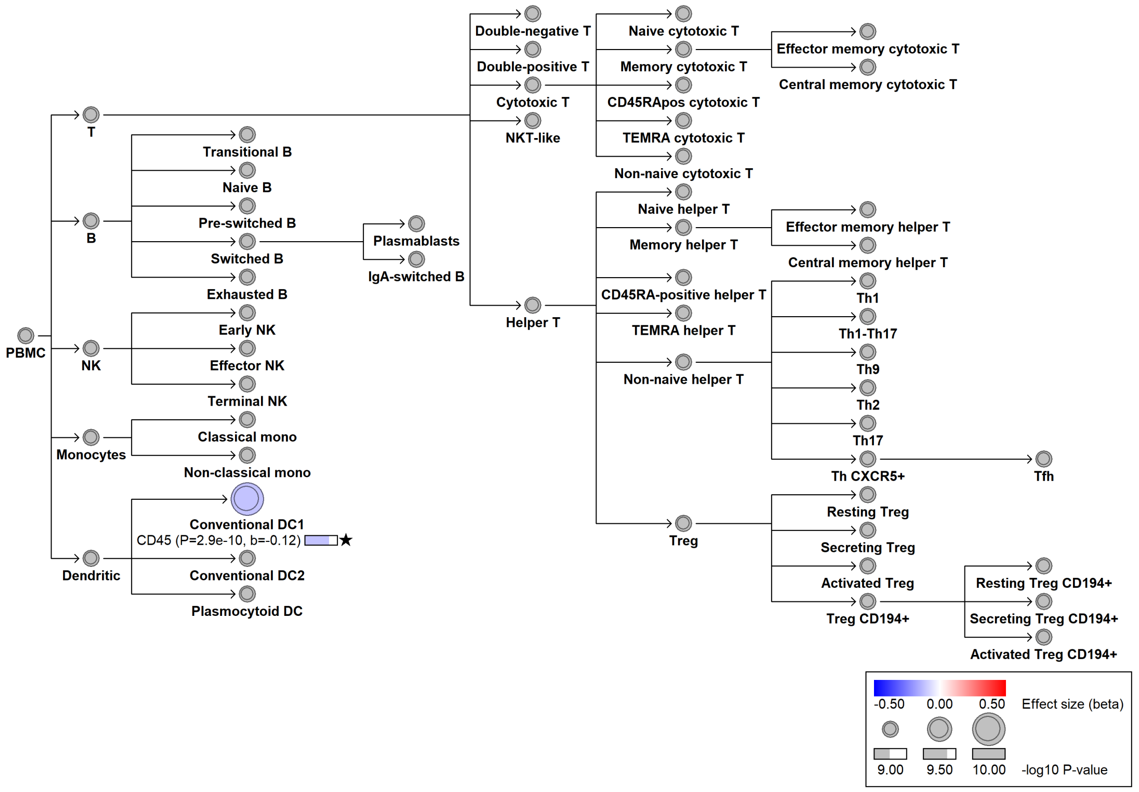

#### chr13:28029870[T→C; EAF 1.592%] (rs76428106, cs_3)

Candidate genes: *FLT3*
Diseases associations: Autoimmune thyroiditis, Rheumatoid arthritis
Plasma pQTLs: ADA2, ADAM8, BLNK, C1QTNF1, CCDC50, CCL2, CCL21, CD160, CD1C, CD207, CD22, CD244, CD27, CD300C, CD300E, CD48, CD72, CD79B, CD80, CD83, CD86, CLEC11A, CLEC4C, CLEC4D, CLEC5A, CLEC6A, CLEC7A, CR2, CRISP3, CSF1R, CSF2RA, Composite immunoglobulin trait (IgA*IgG*IgM), Composite immunoglobulin trait IgA*IgG, Composite immunoglobulin trait IgM/(IgA*IgG), FAIM3, FCAR, FCER2, FCRL1, FCRL2, FCRL5, FLT3LG, GRN, ICAM3, IFNLR1, IGLL1, IL12A, IL12B, IL15, IL18BP, IL6R, ITGA5, Immunoglobulin A, Immunoglobulin G, KPNA2, LAG3, LCN2, LILRA2, LILRA4, LILRA5, LY9, LYZ, MAT2B, MPO, MZB1, NCR1, OSCAR, PDCD1, PGLYRP1, PRTN3, PTPRC, QPCT, RAB26, RETN, RNASE2, RNASE3, RNASE6, SELL, SELPLG, SEMA4A, SIGLEC10, SIGLEC6, SIRPB1, SLAMF7, SPINK2, TCL1A, TNFRSF13C, TNFRSF8, VCAM1

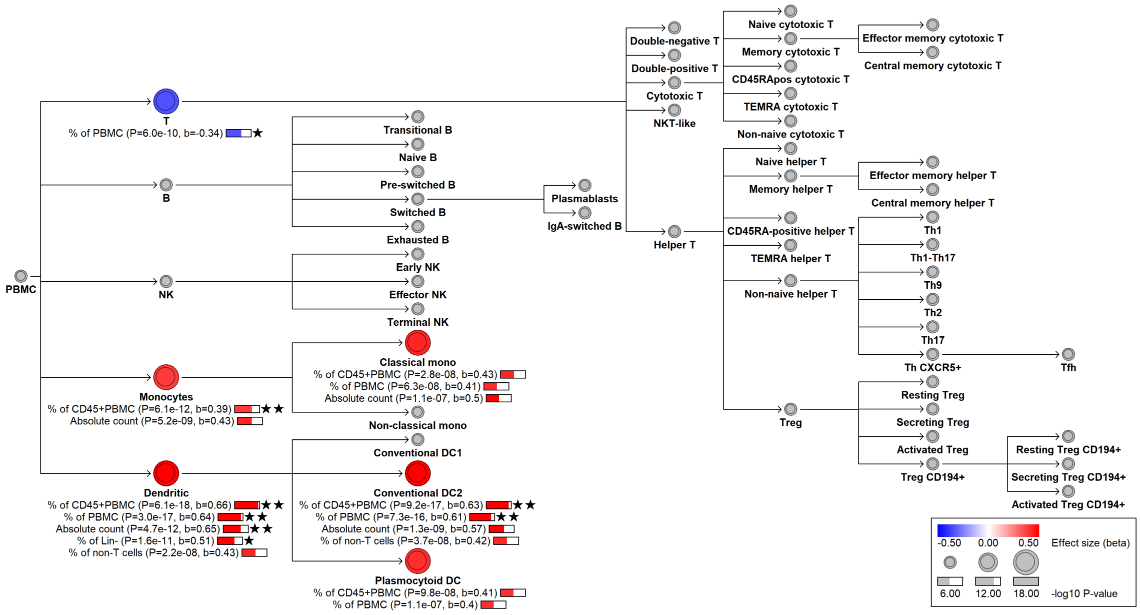

### chr13:32286287[T→C; EAF 1.169%] (rs75684916, cs_82)

Candidate genes: *FRY*
Diseases associations: NA
Plasma pQTLs: NA

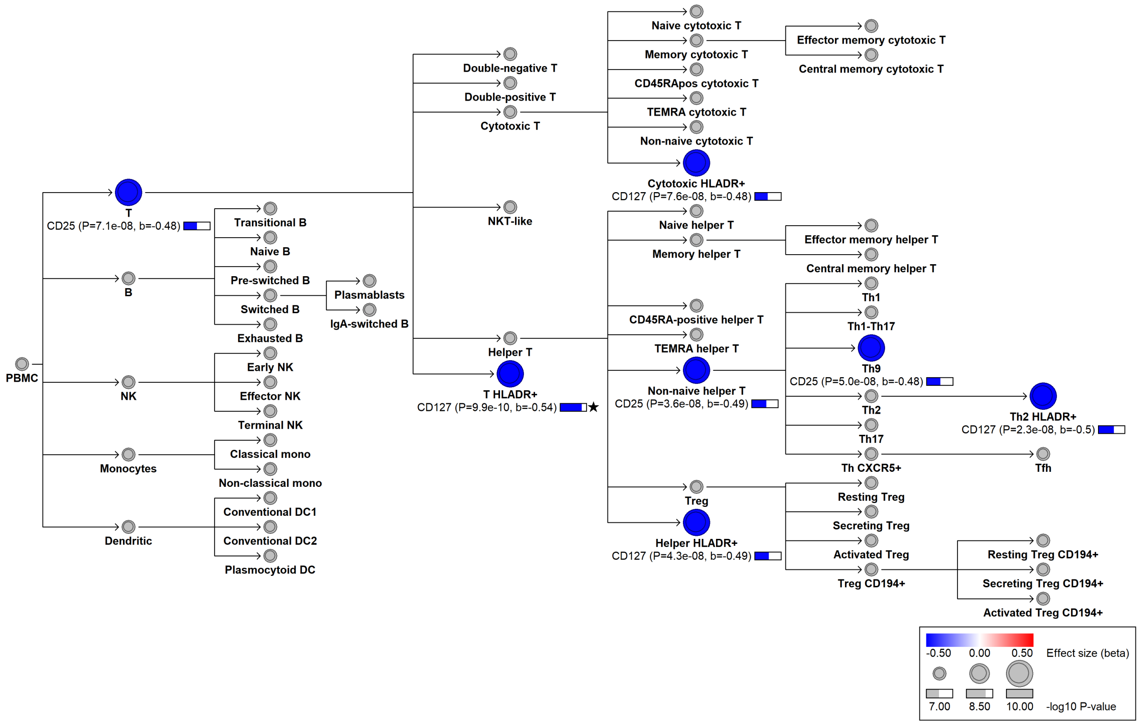

#### chr13:50418947[!→C; EAF 3.887%] (rs143280072, cs_152)

Candidate genes: *DLEU1*
Diseases associations: NA
Plasma pQTLs: NA

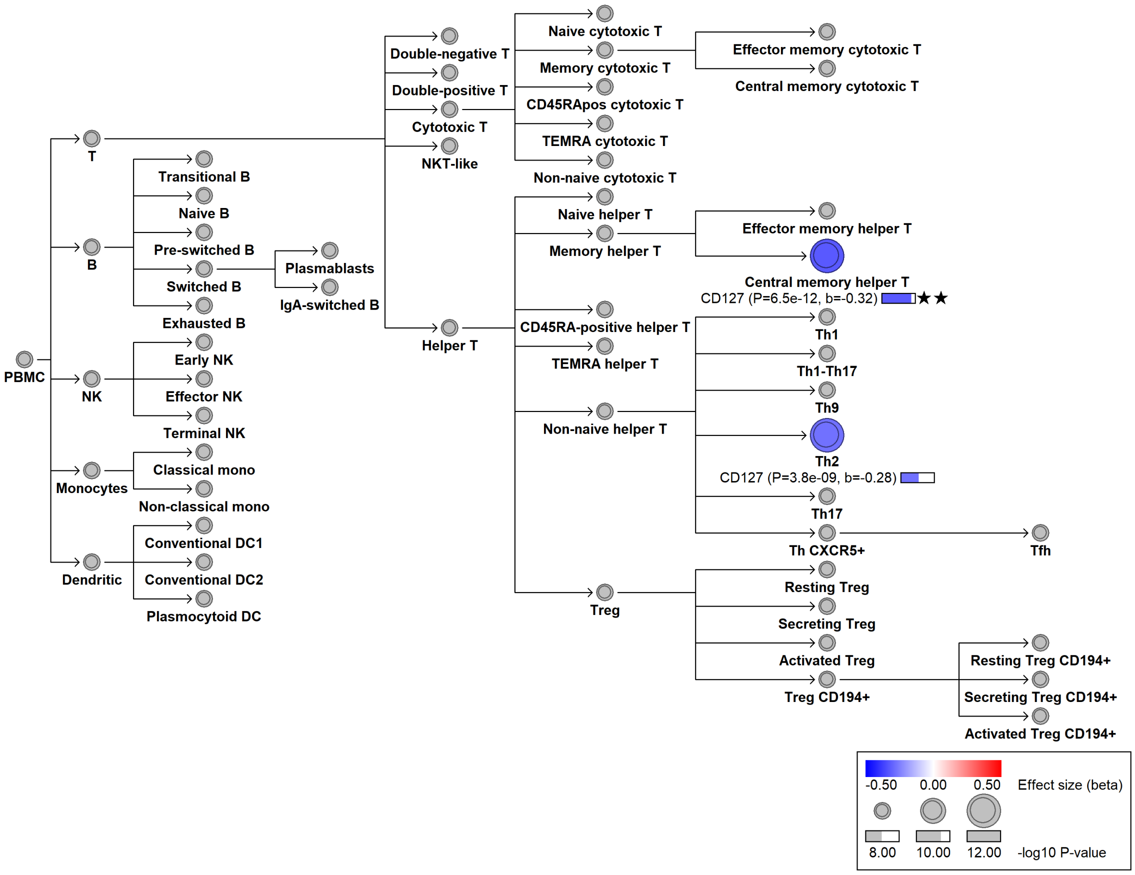

#### chr13:99432709[A→T; EAF 25.73%] (rs9517724, cs_83)

Candidate genes: *LNCARGI*
Diseases associations: NA
Plasma pQTLs: NA

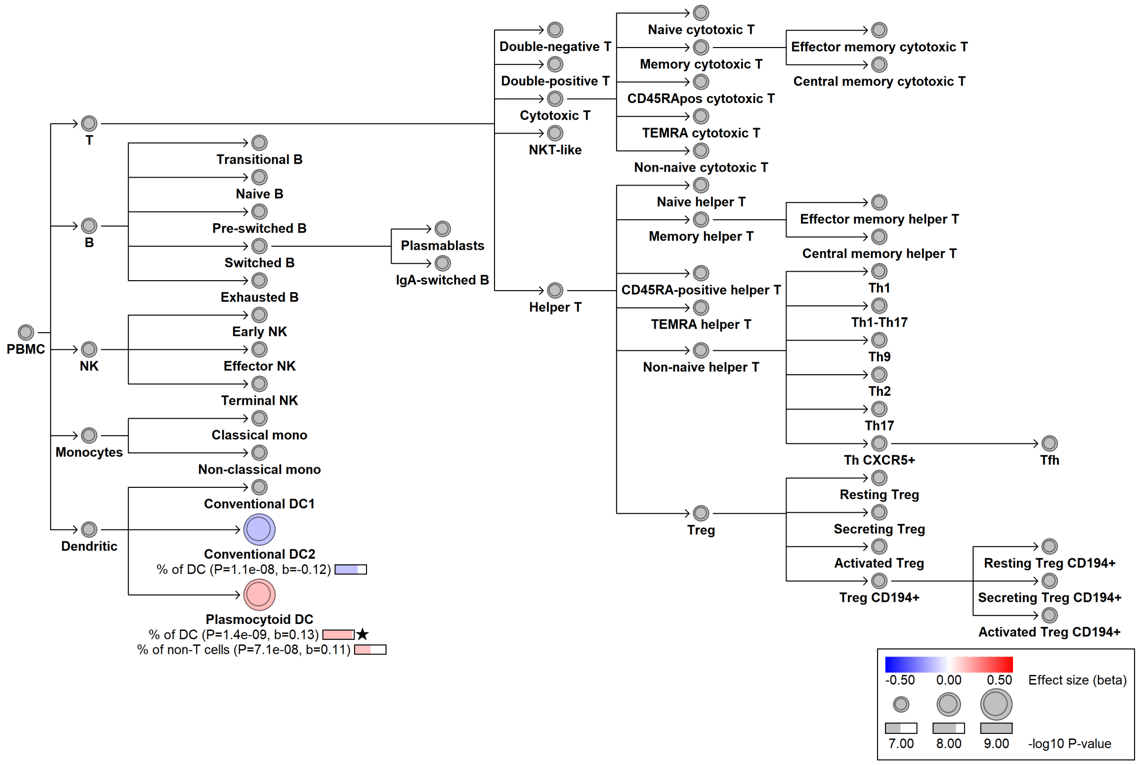

#### chr13:108308032[TGCTG→T; EAF 2.099%] (rs200748895, cs_53)

Candidate genes: *TNFSF13B*
Diseases associations: Chronic diseases of tonsils
Plasma pQTLs: C1QL2, CD27, CD5L, CD72, CXCL13, Composite immunoglobulin trait (IgA*IgG*IgM), Composite immunoglobulin trait IgA*IgG, FAIM3, FCER2, FCRL1, FCRL2, FCRL3, FCRL5, Immunoglobulin G, Immunoglobulin M, LTA, LTB, MZB1, TNFRSF13B, TNFRSF13C, TNFRSF17, TNFSF13B

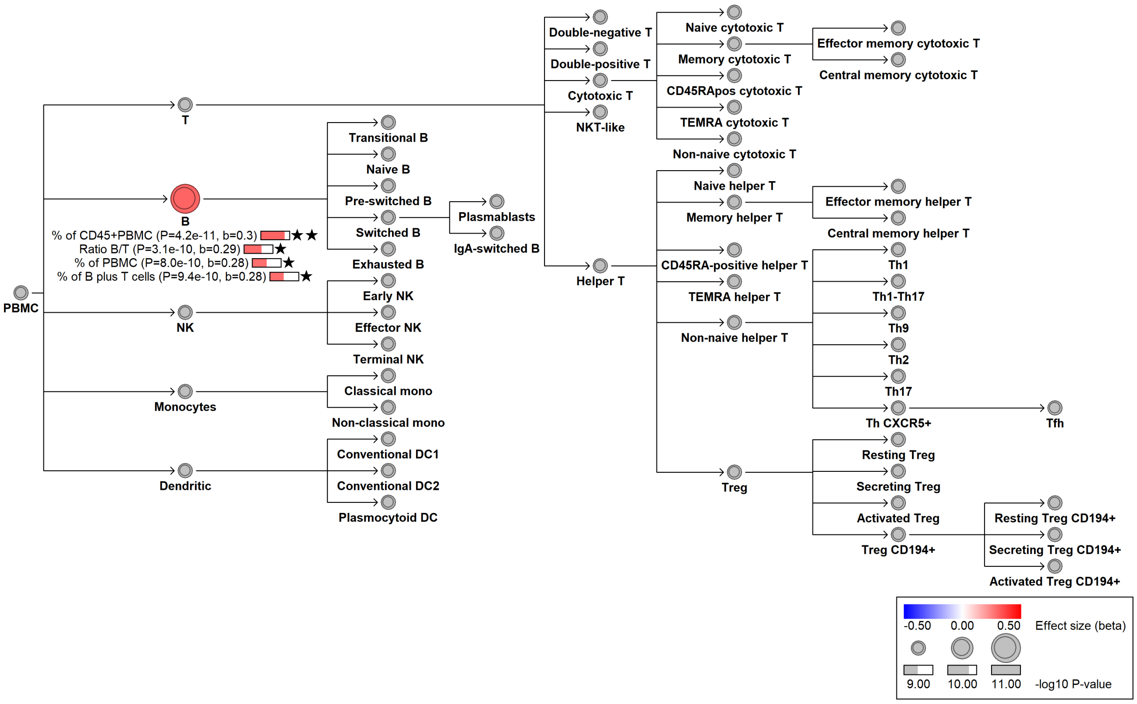

#### chr14:35340149[!→C; EAF 18.242%] (rs12436188, cs_30)

Candidate genes: *NFKBIA*
Diseases associations: NA
Plasma pQTLs: NA

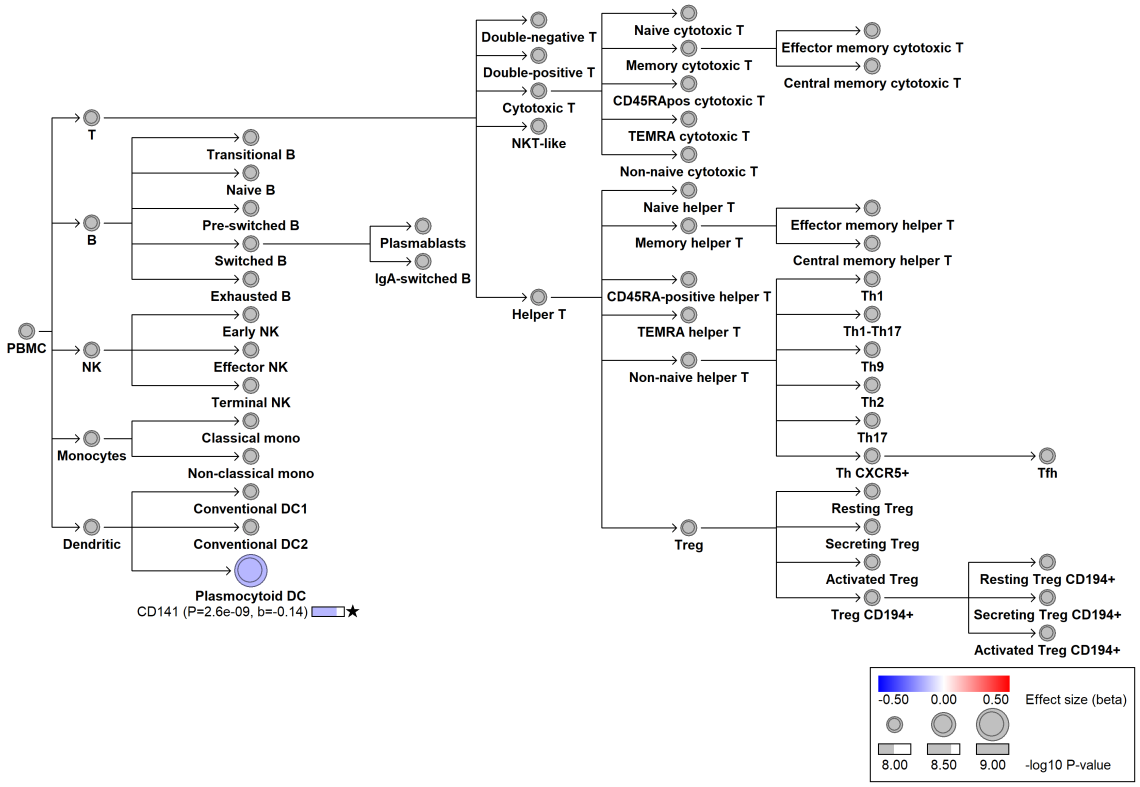

### chr14:105588119[G→A; EAF 3.831%] (rs61984162, cs_133)

Candidate genes: *IGHA2*
Diseases associations: NA
Plasma pQTLs: NA

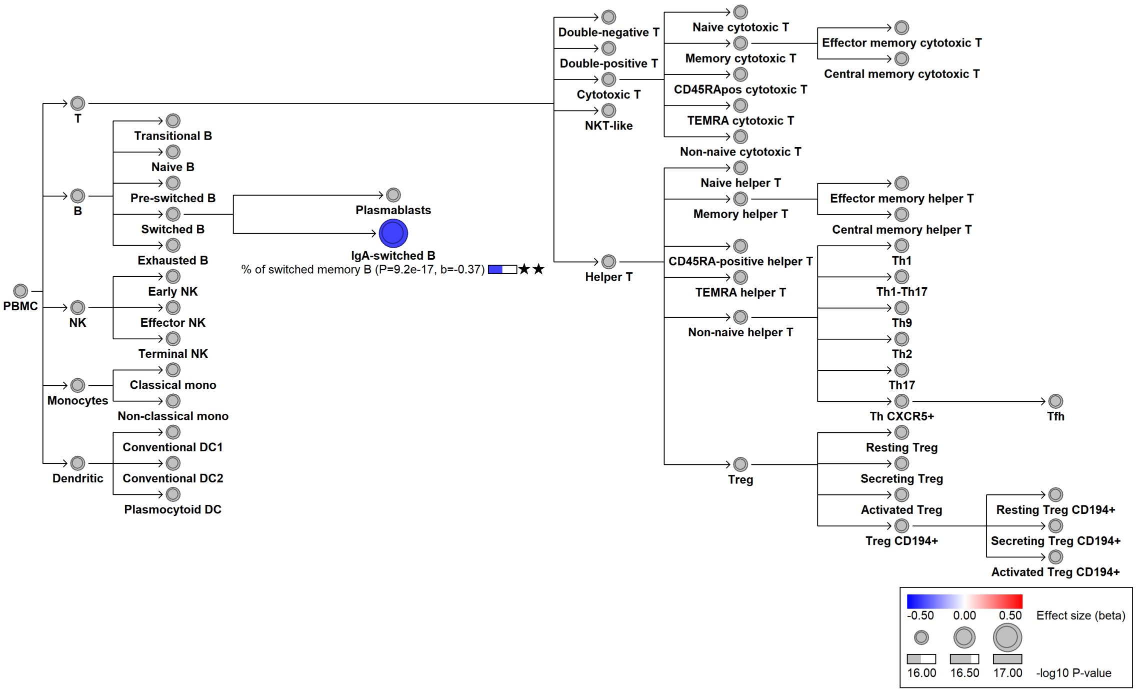

#### chr15:40110818[G→GC; EAF 46.946%] (rs36081508, cs_69)

Candidate genes: *BMF*
Diseases associations: Chronic lymphocytic leukemia
Plasma pQTLs: Composite immunoglobulin trait IgA/IgG, Composite immunoglobulin trait IgA/IgM, Composite immunoglobulin trait IgM/(IgA*IgG), Immunoglobulin A

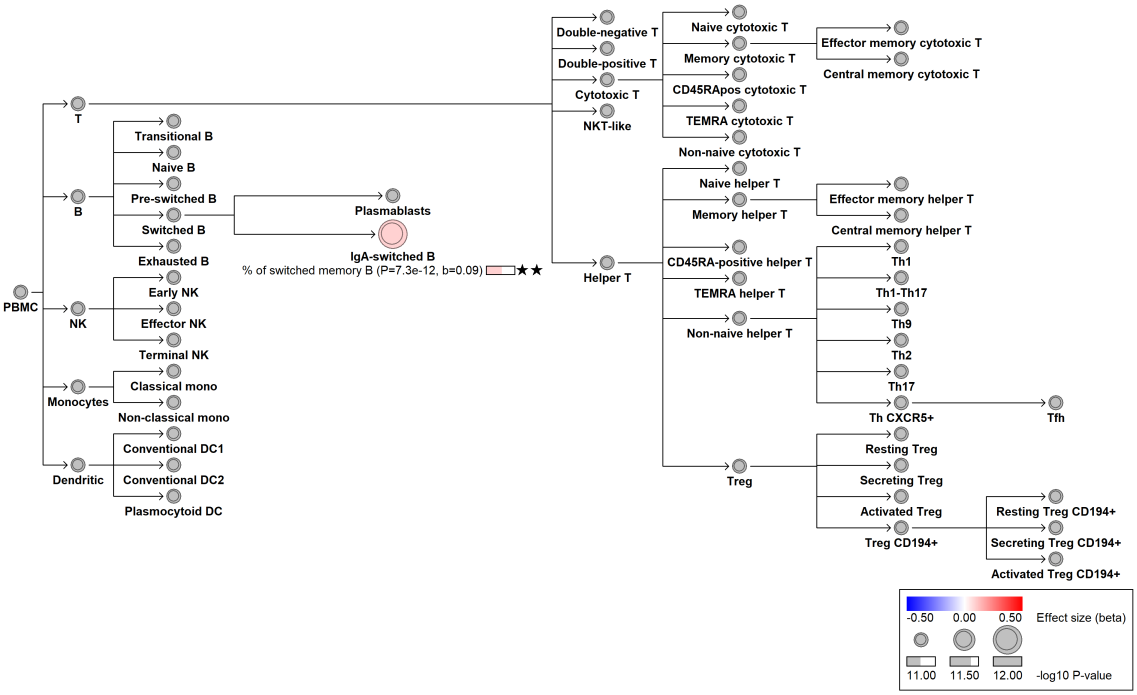

#### chr15:40624039[A→G; EAF 40.19%] (rs17747633, cs_115)

Candidate genes: *CCDC32*
Diseases associations: NA
Plasma pQTLs: Composite immunoglobulin trait IgG/IgM, Immunoglobulin M

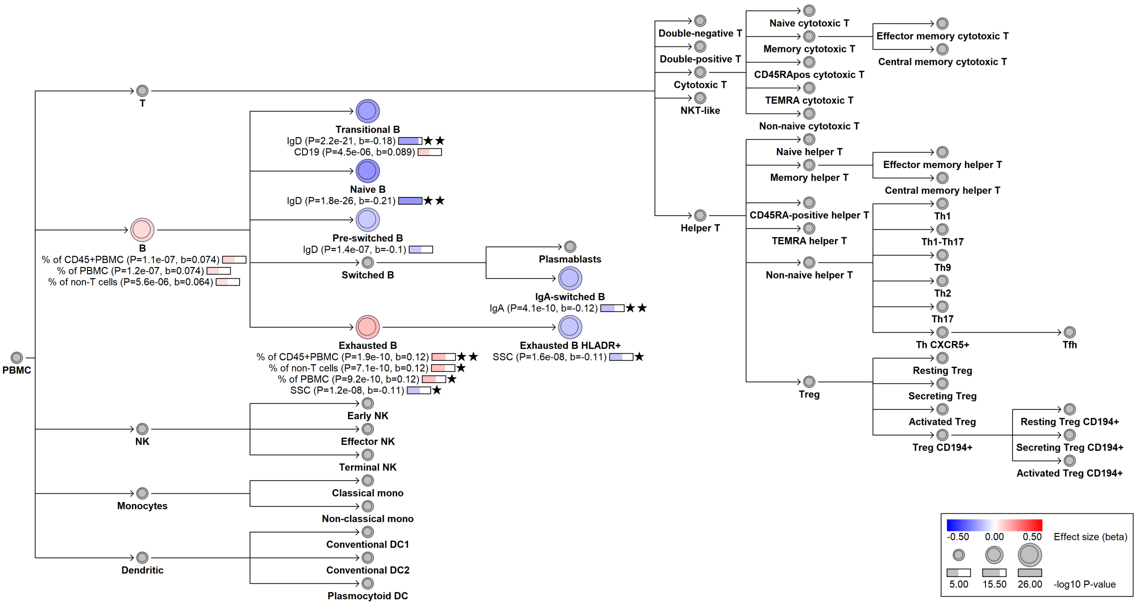

#### chr15:60765034[G→!; EAF 14.288%] (chr15:60765034, cs_147)

Candidate genes: *RORA*
Diseases associations: Allergy, Asthma, Eczema, Eosinophilic esophagitis, Inflammatory bowel disease
Plasma pQTLs: NA

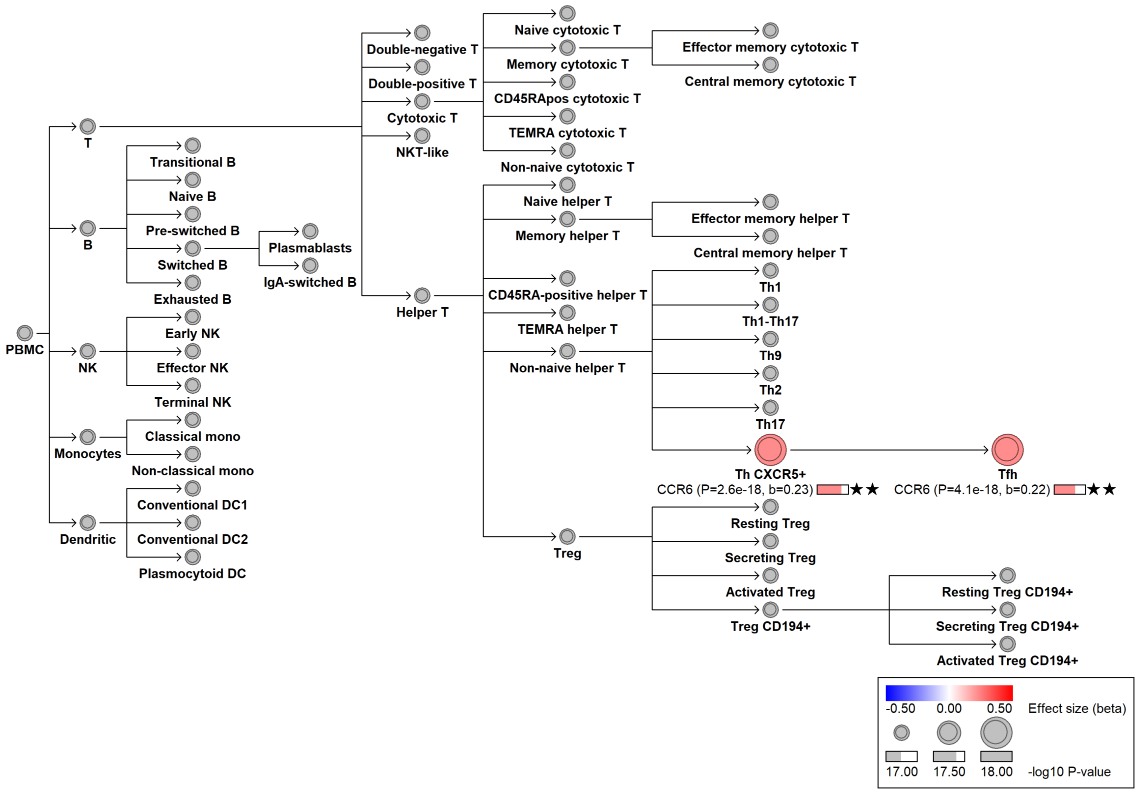

#### chr15:70439674[A→G; EAF 40.247%] (rs35026629, cs_111)

Candidate genes: *NA*
Diseases associations: NA
Plasma pQTLs: NA

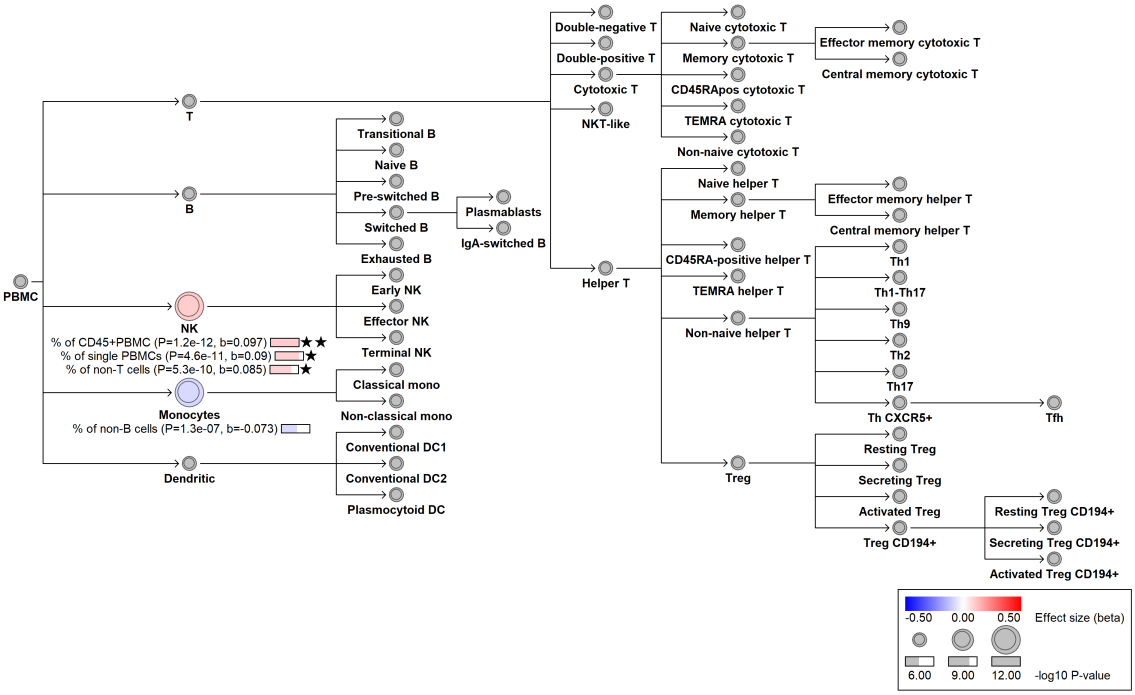

### chr15:79971003[A→C; EAF 25.822%] (rs1138358, cs_132)

Candidate genes: *BCL2A1*

Diseases associations: NA

Plasma pQTLs: CD83, IDO1, IL12A, IL12B, LILRA5, WARS

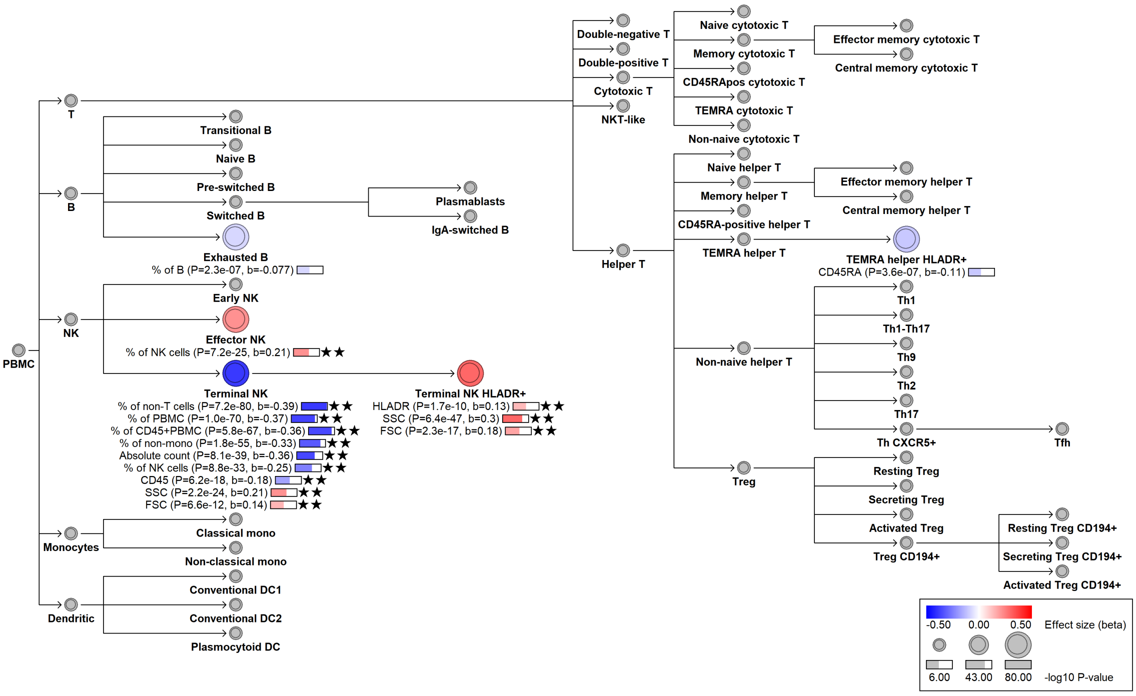

#### chr16:377479[C→T; EAF 45.003%] (rs11248931, cs_258)

Candidate genes: *PGAP6*

Diseases associations: NA

Plasma pQTLs: ART3, CFC1, CNTN1, CNTN3, EFNA4, EFNA5, LYPD3, NCAM2, NTM, TCTN3, UMOD

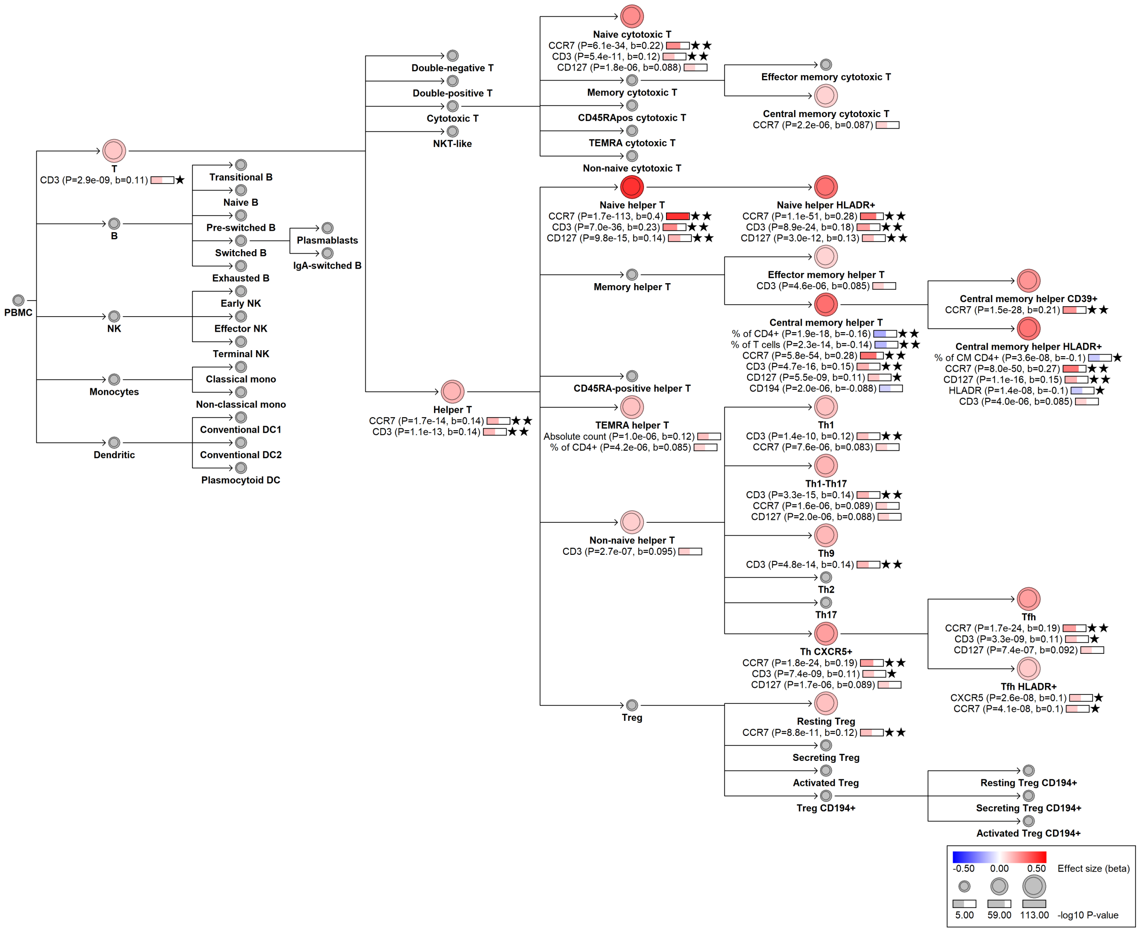

#### chr16:380304[T→C; EAF 42.284%] (rs12921174, cs_229)

Candidate genes: *PGAP6*
Diseases associations: NA
Plasma pQTLs: EFNA4

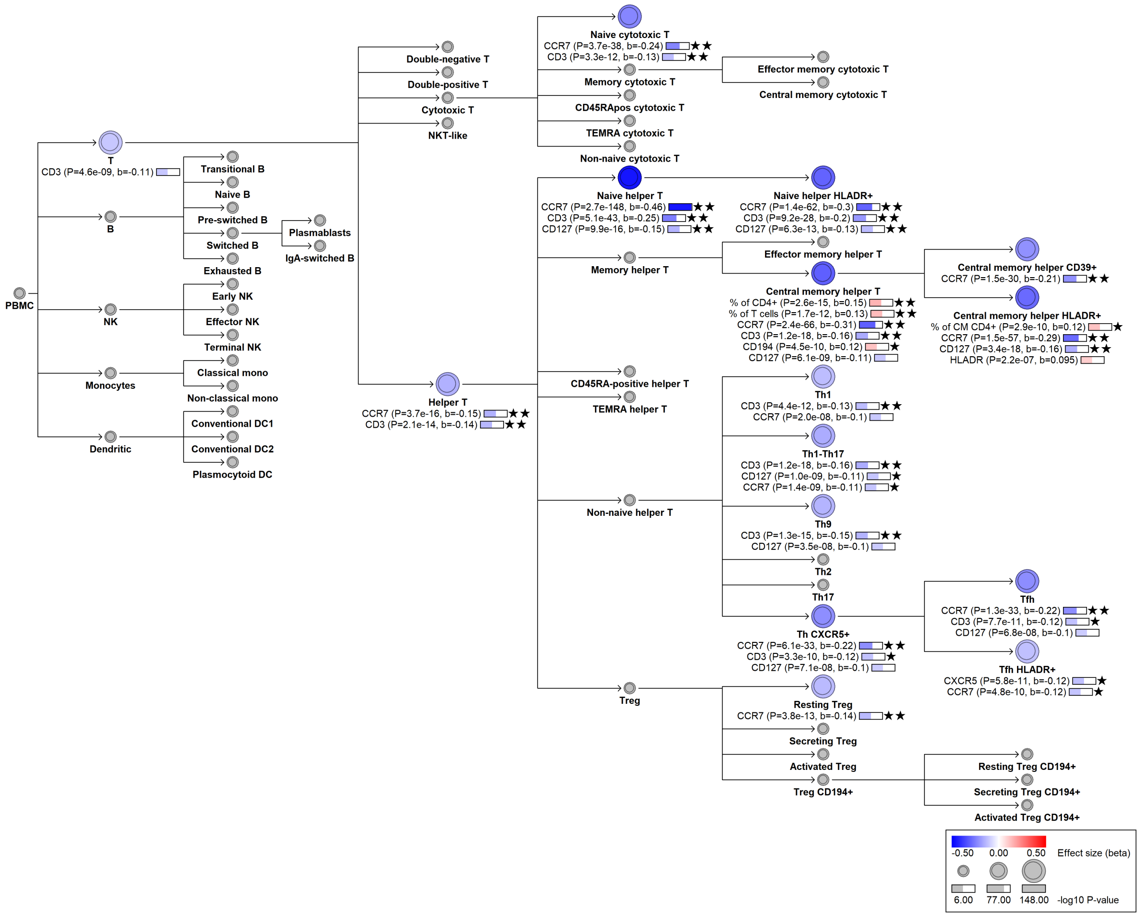

#### chr16:10877045[A→!; EAF 24.905%] (rs3087456, cs_108)

Candidate genes: *CIITA*
Diseases associations: NA
Plasma pQTLs: CD5L, CD74, IL18RAP

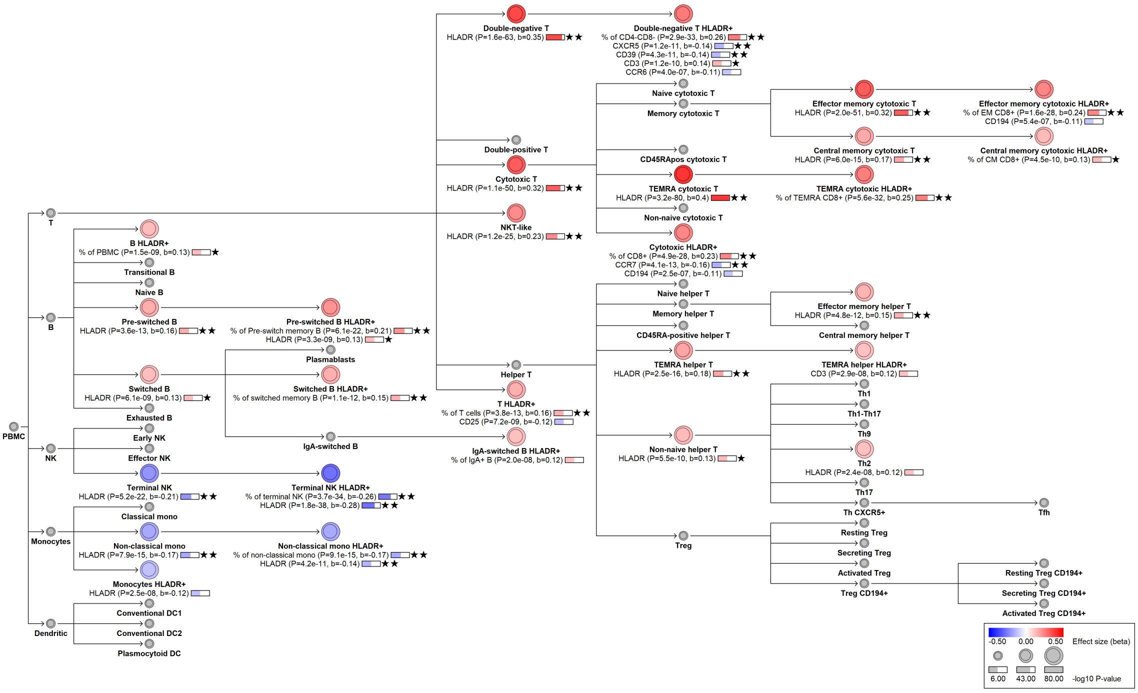

### chr16:10895362[C→G; EAF 1.856%] (rs2229317, cs_76)

Candidate genes: *CIITA*
Diseases associations: NA
Plasma pQTLs: NA

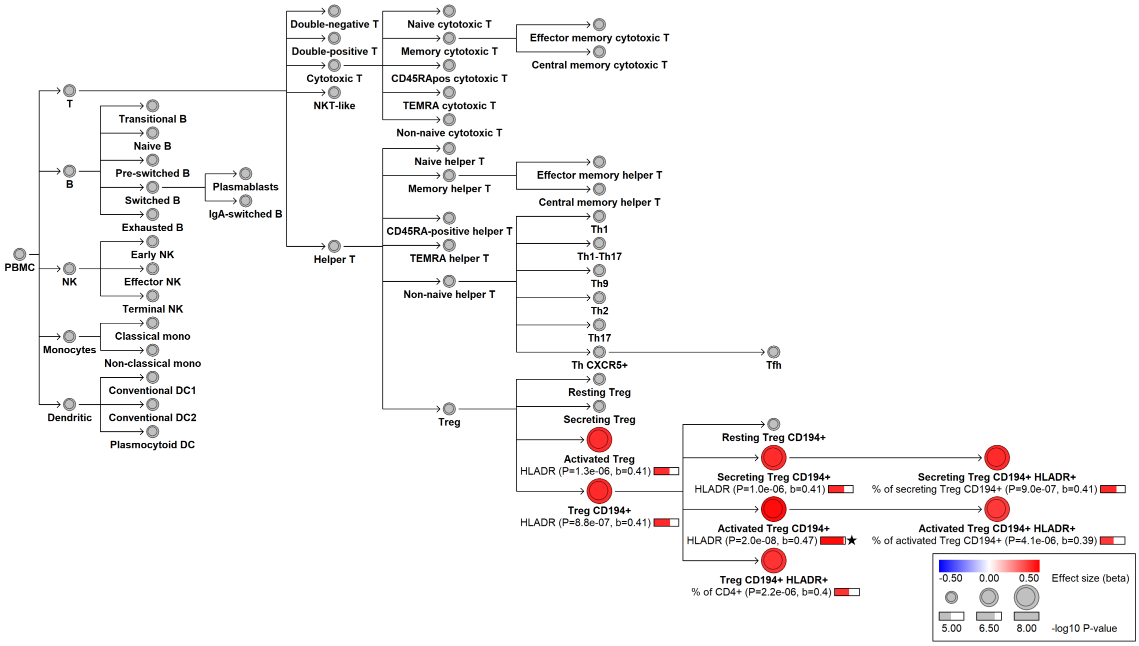

### chr16:10910506[C→!; EAF 16.213%] (rs6498126, cs_97)

Candidate genes: *CIITA*
Diseases associations: NA
Plasma pQTLs: NA

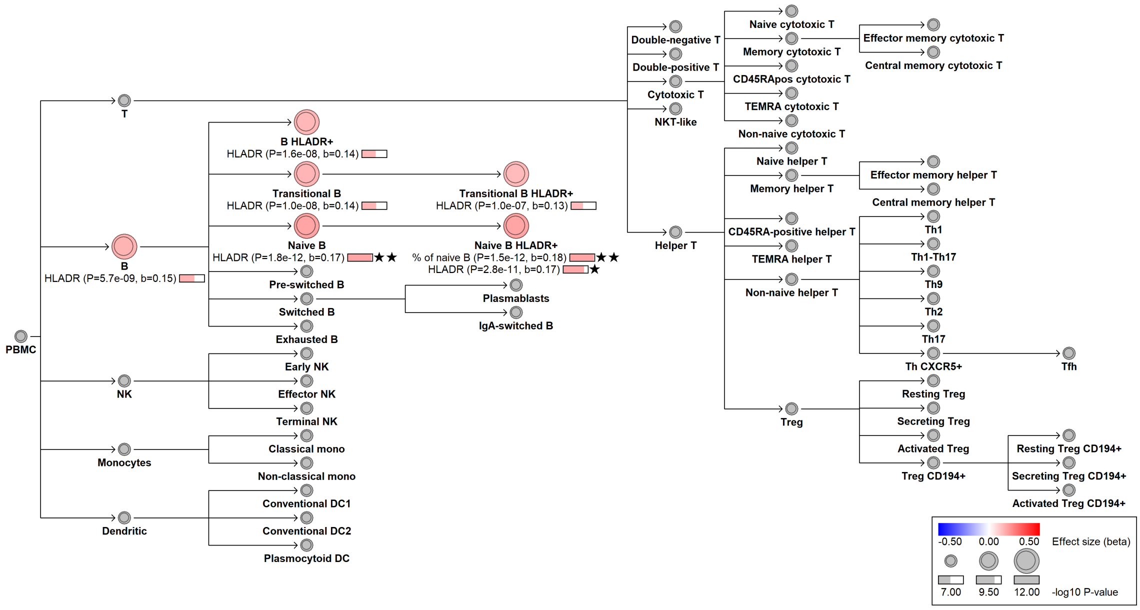

### chr16:10911032[CA→!; EAF 25.043%] (rs397801599, cs_189)

Candidate genes: *CIITA*
Diseases associations: NA
Plasma pQTLs: NA

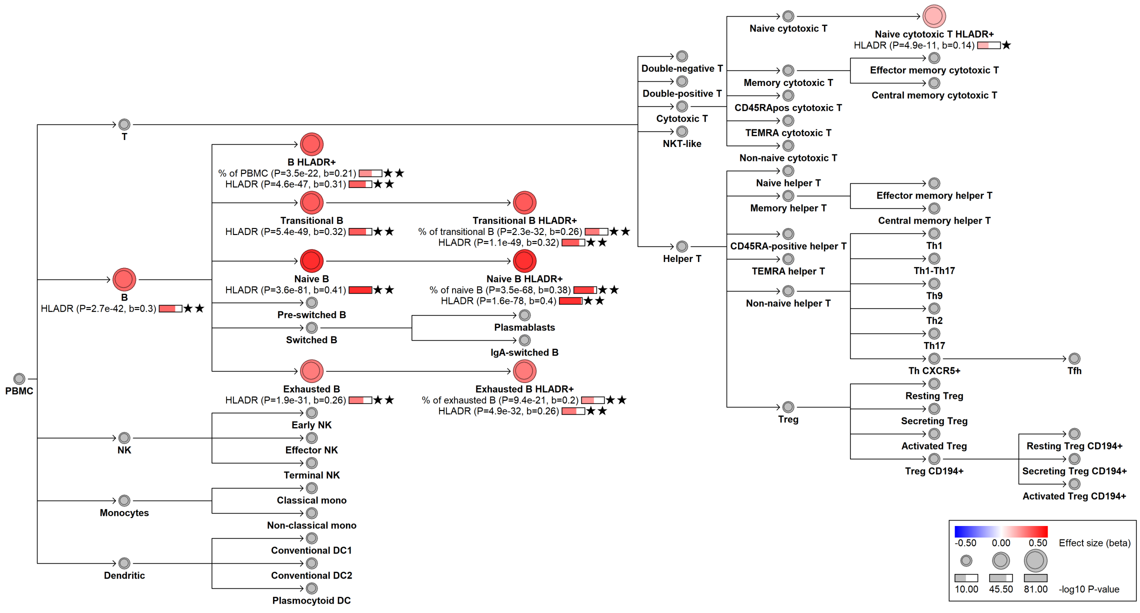

#### chr16:19699568[C→T; EAF 2.065%] (rs150300279, cs_216)

Candidate genes: *VPS35L*
Diseases associations: NA
Plasma pQTLs: NA

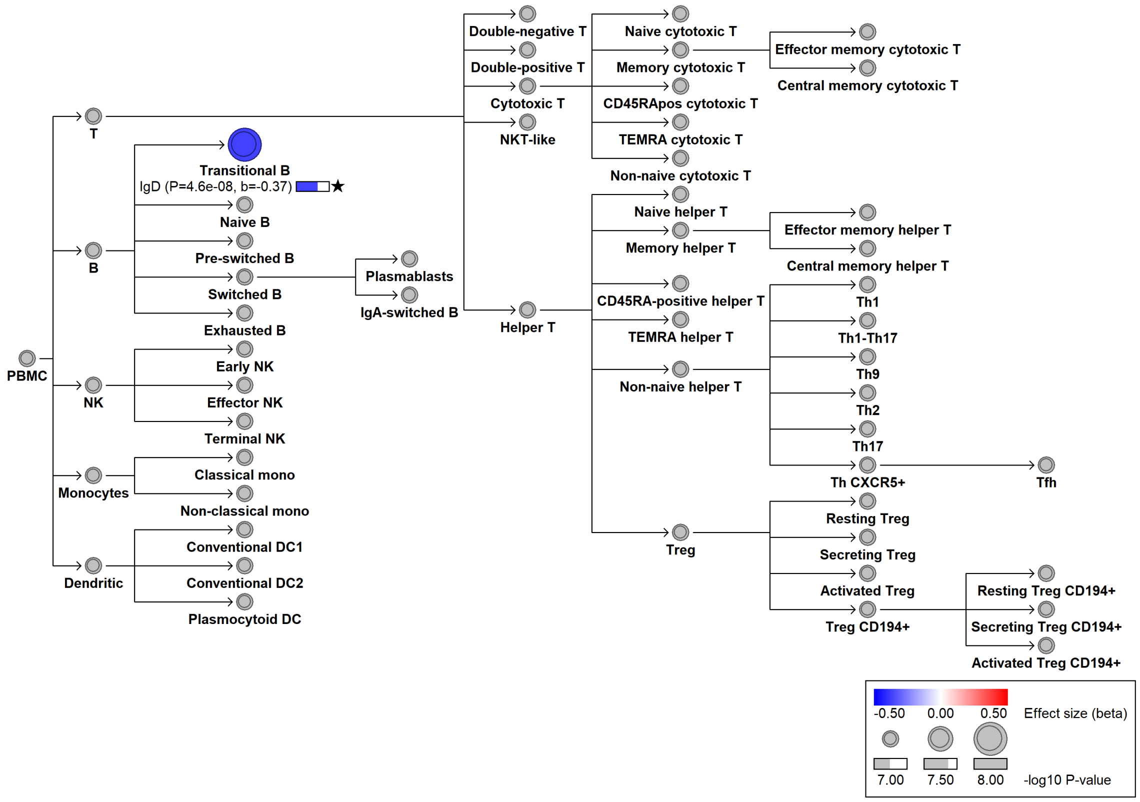

#### chr16:28932936[G→A; EAF 1.05%] (rs142342927, cs_166)

Candidate genes: *CD19*
Diseases associations: NA
Plasma pQTLs: NA

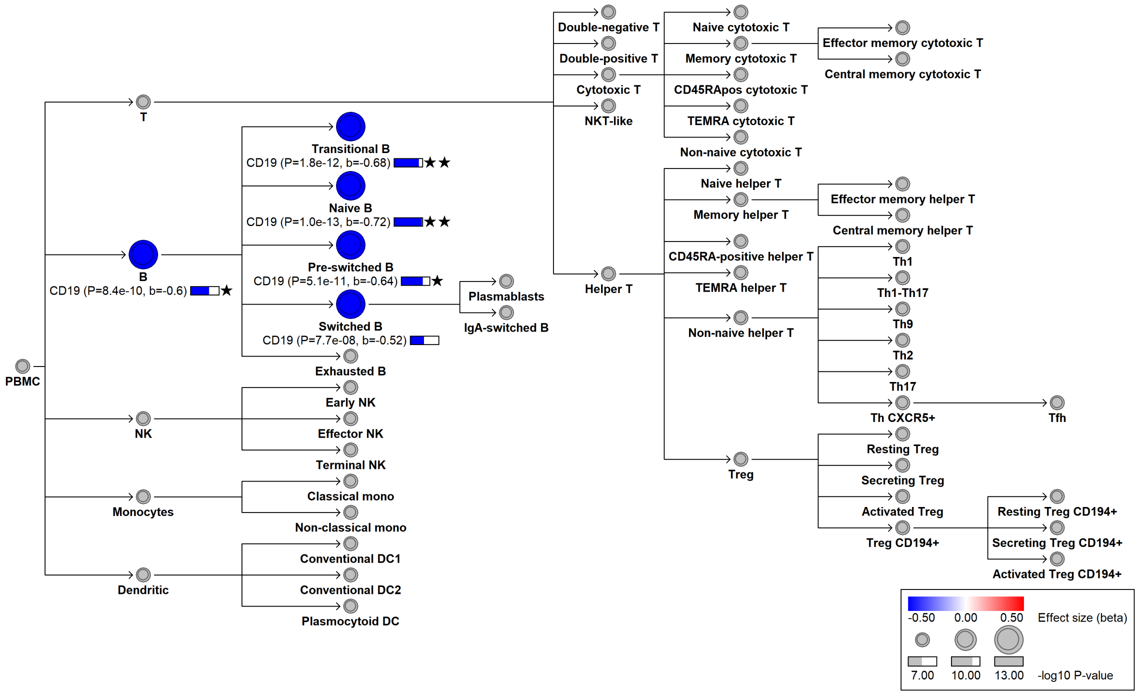

#### chr16:28977312[TAAAAAAAAAA→!; EAF 3.72E+01%] (rs11297402, cs_56)

Candidate genes: *CD19*
Diseases associations: Chronic diseases of tonsils, Parkinson’s disease, Type 2 diabetes
Plasma pQTLs: NA

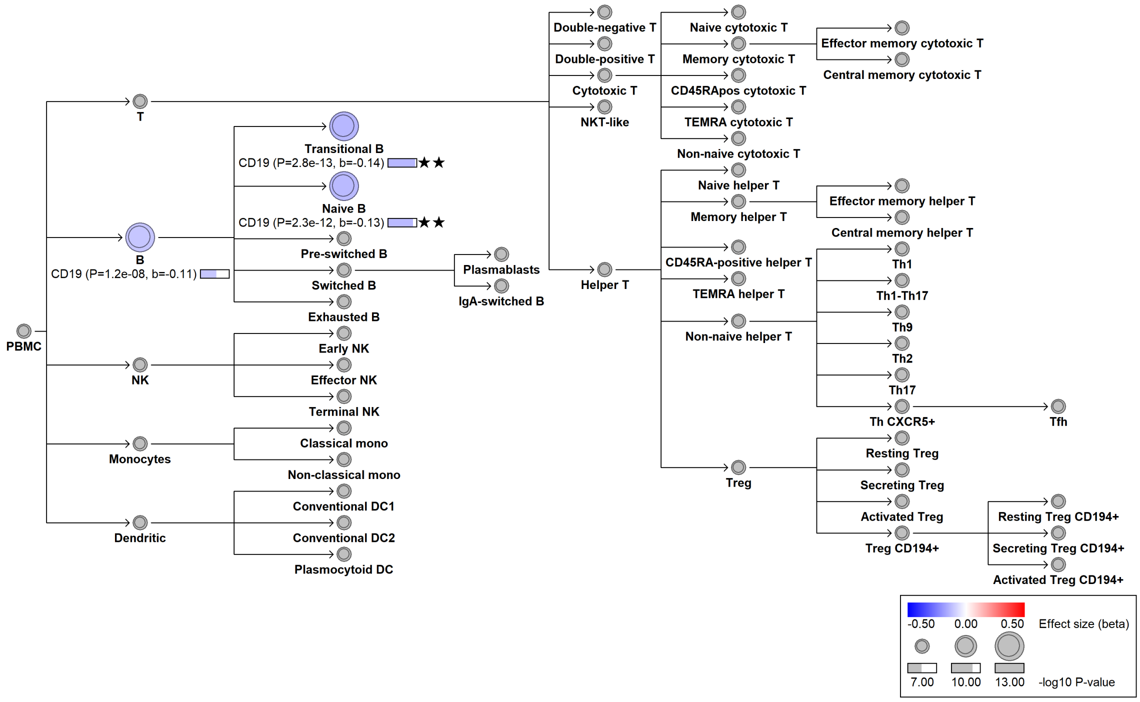

#### chr16:85961830[T→C; EAF 9.954%] (rs113646461, cs_78)

Candidate genes: *IRF8*
Diseases associations: Crohn's disease, Inflammatory bowel disease
Plasma pQTLs: NCR1

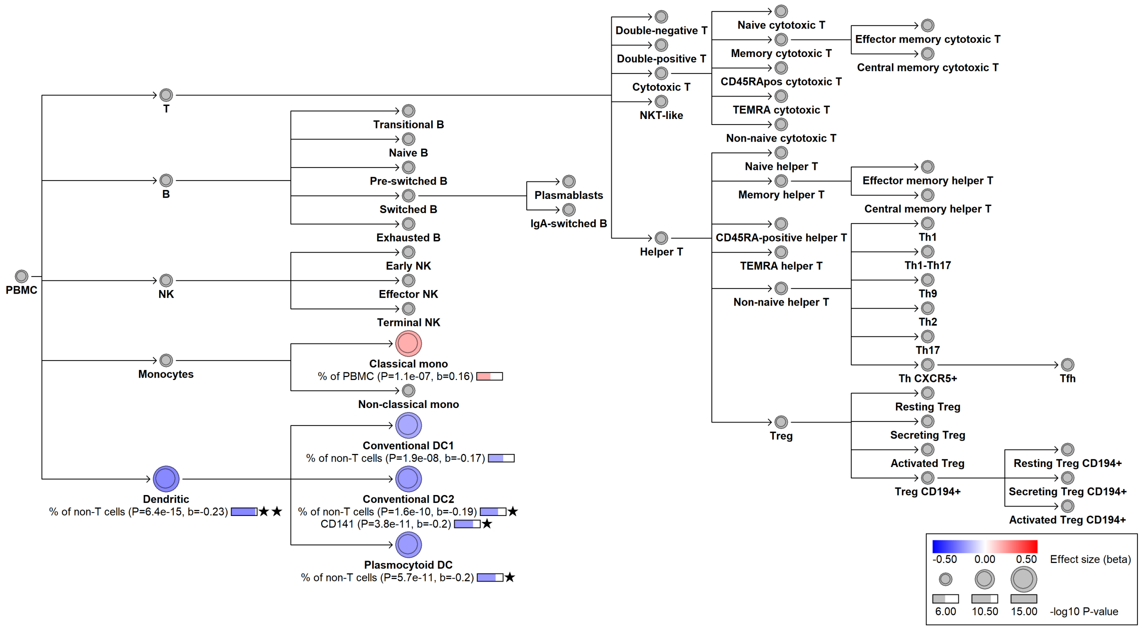

#### chr16:85982795[C→G; EAF 23.174%] (rs9927316, cs_136)

Candidate genes: *IRF8*
Diseases associations: Rheumatoid arthritis
Plasma pQTLs: FCER2, IGLL1, IL22RA2

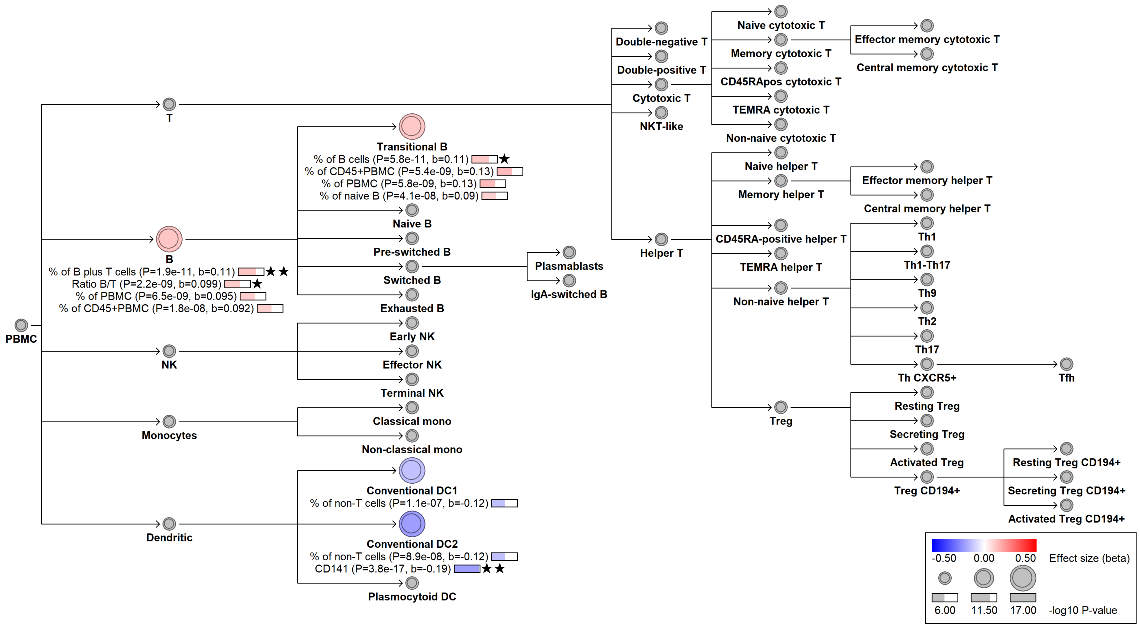

#### chr16:88978825[C→T; EAF 15.329%] (rs62045817, cs_144)

Candidate genes: *CBFA2T3*
Diseases associations: NA
Plasma pQTLs: CLEC4C, CRTAP, Composite immunoglobulin trait (IgA*IgG*IgM), Immunoglobulin M, KIF22, LILRA4, PIKFYVE, SIGLEC6

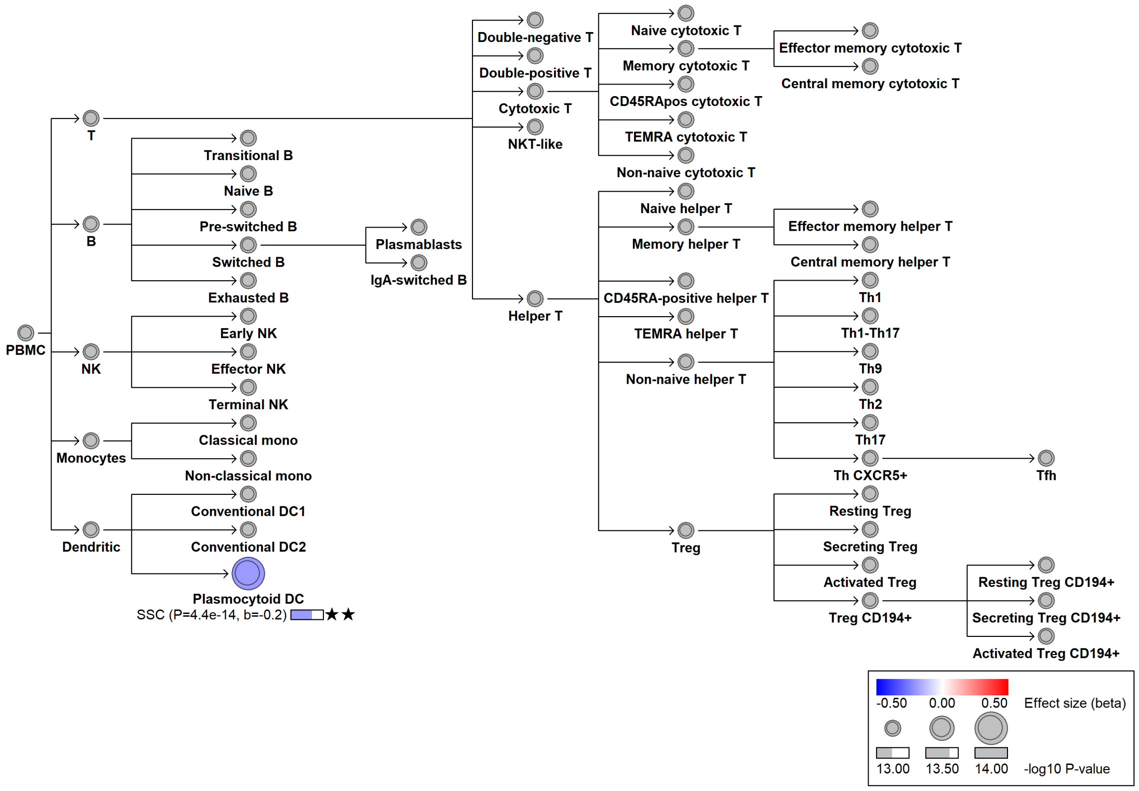

### chr17:2812079[C→T; EAF 48.483%] (rs8073448, cs_80)

Candidate genes: *RAP1GAP2*
Diseases associations: NA
Plasma pQTLs: TCL1A

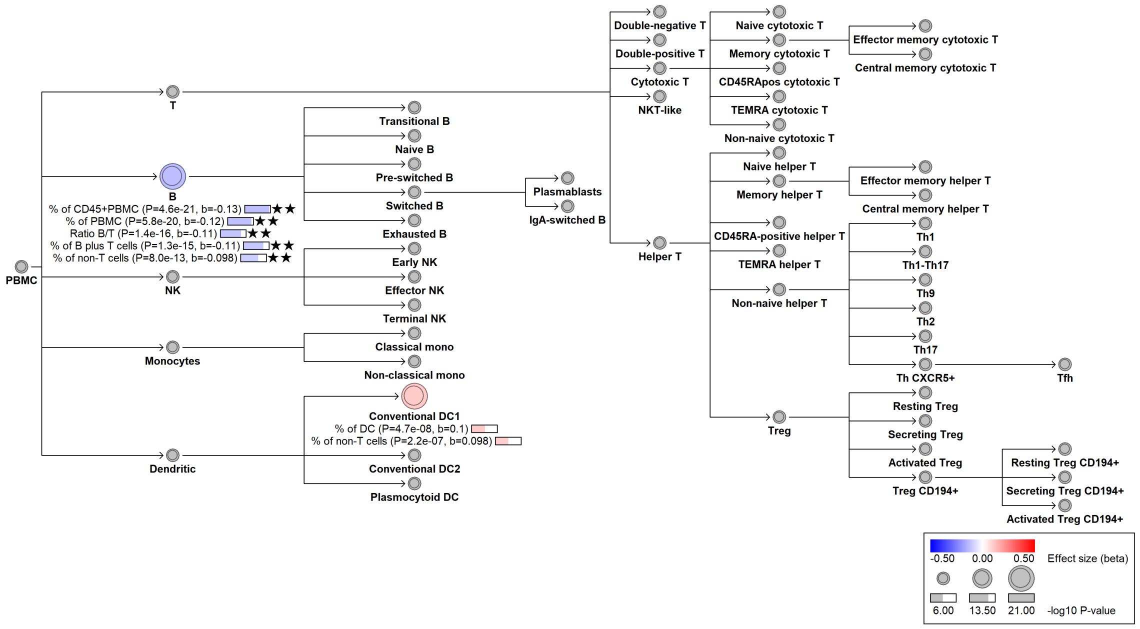

#### chr17:16940415[G→T; EAF 0.876%] (rs72553883, cs_214)

Candidate genes: *TNFRSF13B*
Diseases associations: Chronic diseases of tonsils, Common variable immunodeficiency
Plasma pQTLs: CR2, Composite immunoglobulin trait (IgA*IgG*IgM), Composite immunoglobulin trait IgA*IgG, FCRL1, FCRL4, TNFSF13B

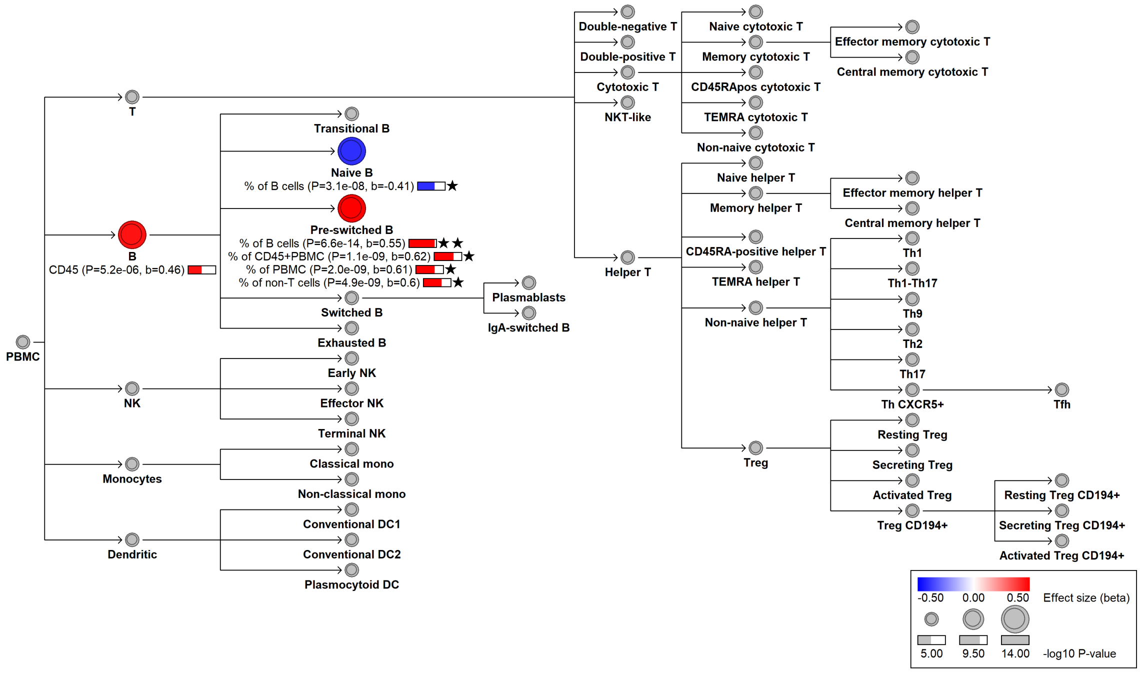

#### chr17:16948873[A→G; EAF 0.342%] (rs34557412, cs_117)

Candidate genes: *TNFRSF13B*
Diseases associations: Chronic diseases of tonsils, Common variable immunodeficiency
Plasma pQTLs: Composite immunoglobulin trait (IgA*IgG*IgM), Composite immunoglobulin trait IgA*IgG, Composite immunoglobulin trait IgA/IgG, Composite immunoglobulin trait IgM/(IgA*IgG), Immunoglobulin A, Immunoglobulin G

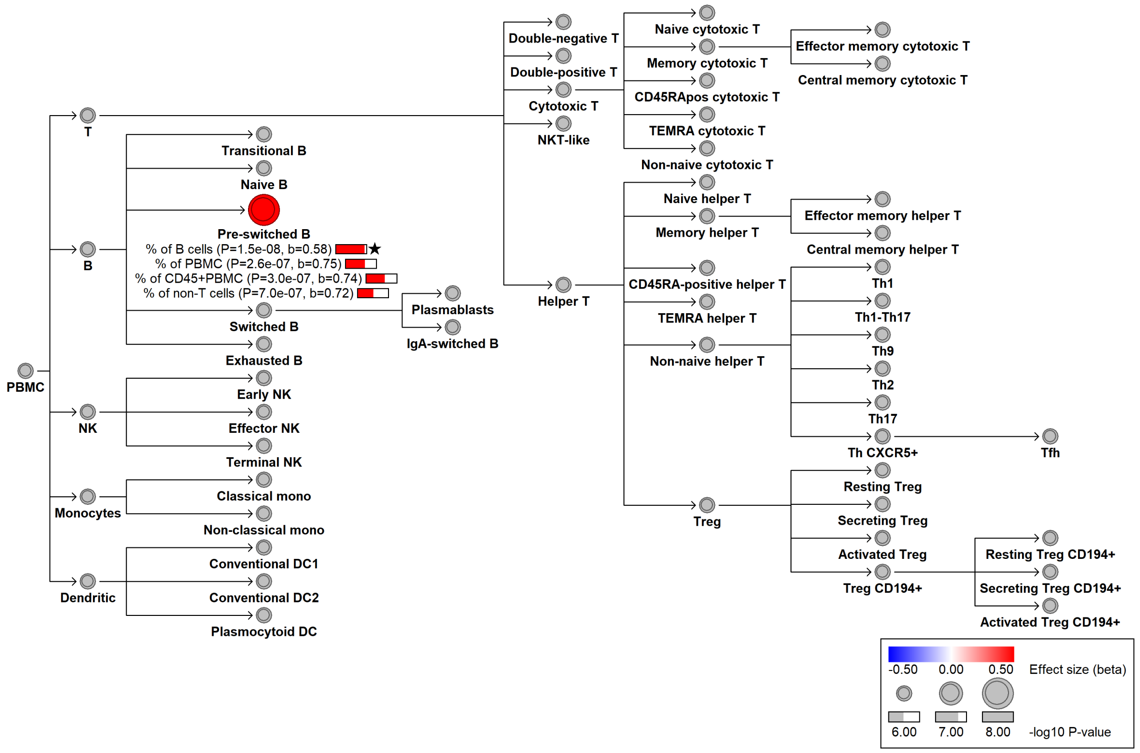

#### chr17:17669723[T→C; EAF 47.655%] (rs2350632, cs_161)

Candidate genes: *RAI1*
Diseases associations: NA
Plasma pQTLs: CBLIF

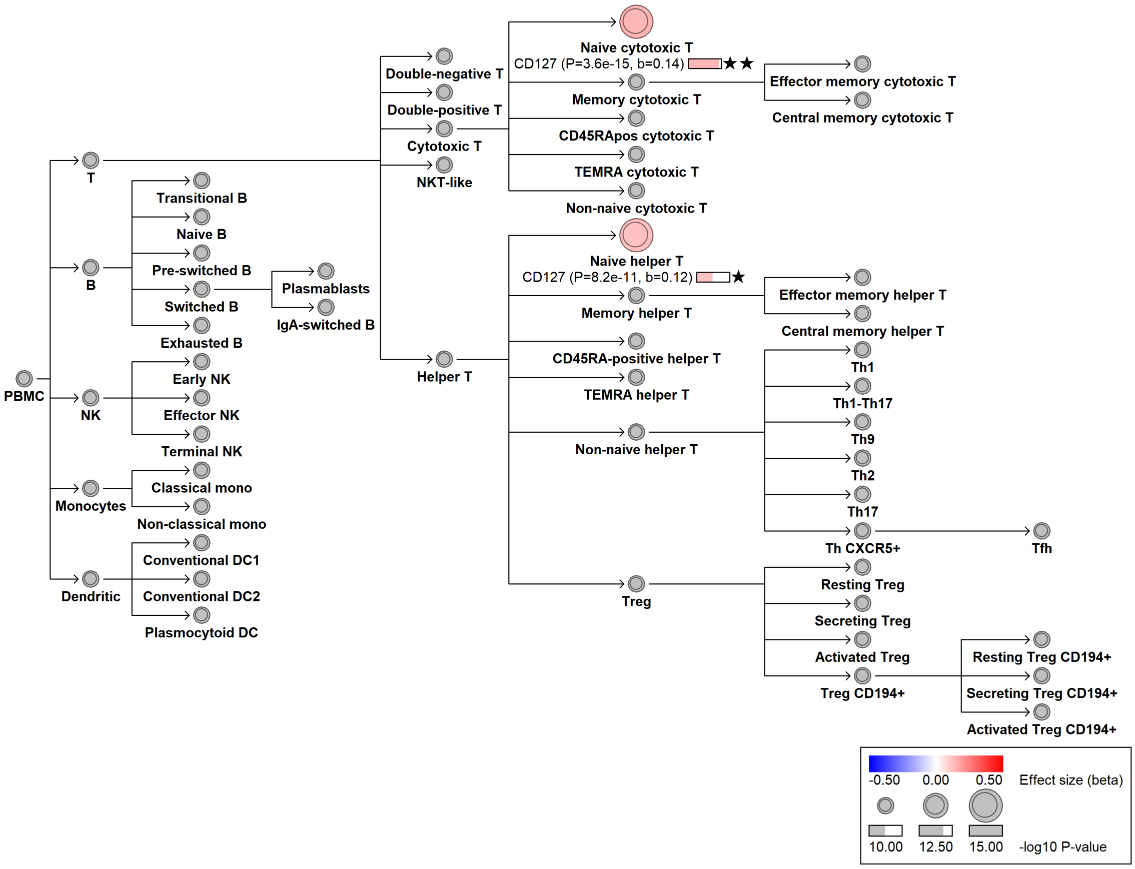

#### chr17:35476573[G→A; EAF 36.056%] (rs58745116, cs_143)

Candidate genes: *SLFN12L*
Diseases associations: NA
Plasma pQTLs: CD160, FASLG, FCRL6, FGFBP2, GZMB, IDO1, IGF1R, IL2RB, INSR, ITGB2, KLRD1, NCR1, SIGLEC7

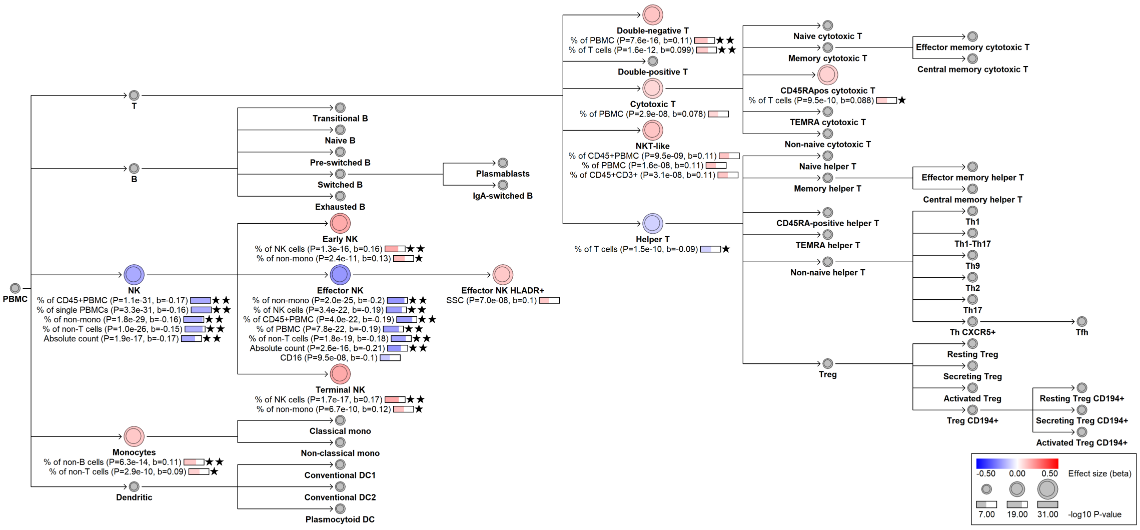

### chr17:35487739[T→C; EAF 7.975%] (rs4796089, cs_176)

Candidate genes: *SLFN12L*
Diseases associations: NA
Plasma pQTLs: NA

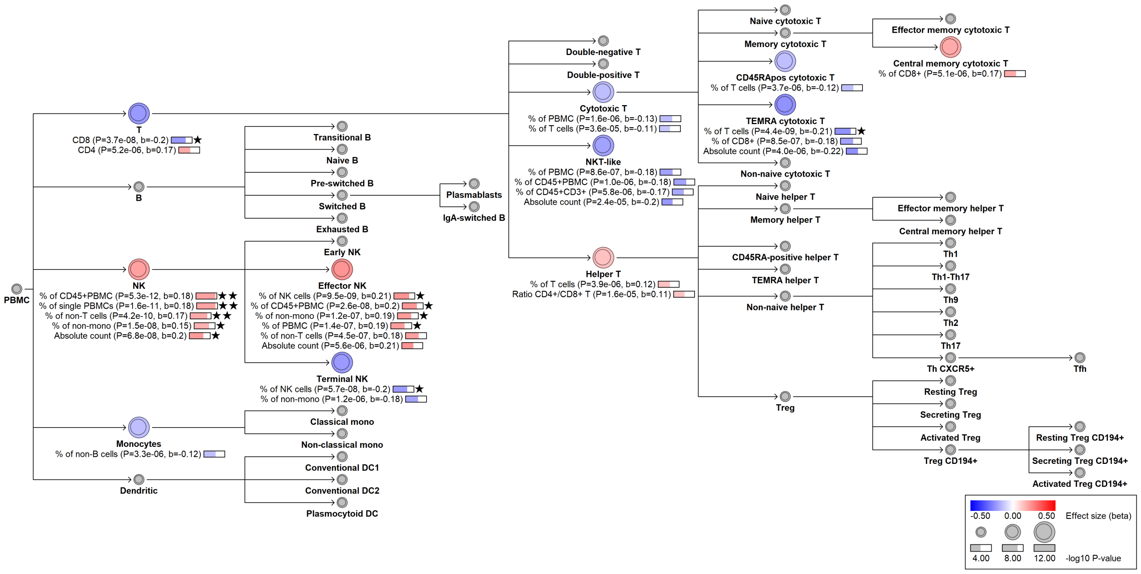

#### chr17:39863888[C→T; EAF 19.839%] (rs9635726, cs_255)

Candidate genes: *IKZF3, GSDMB, ORMDL3*
Diseases associations: Asthma, Cardiovascular disease, Primary biliary cholangitis
Plasma pQTLs: NA

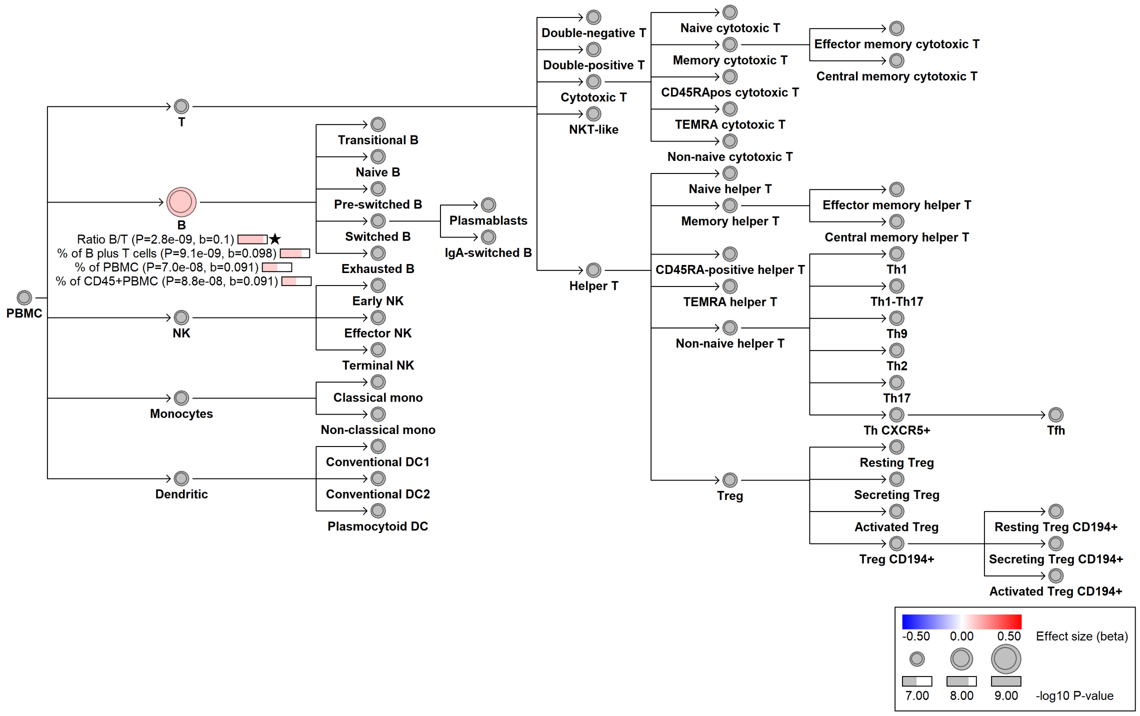

#### chr17:39965740[G→A; EAF 44.987%] (rs3894194, cs_168)

Candidate genes: *GSDMA*
Diseases associations: Allergy, Asthma, Cardiovascular disease, Systemic sclerosis
Plasma pQTLs: FABP9

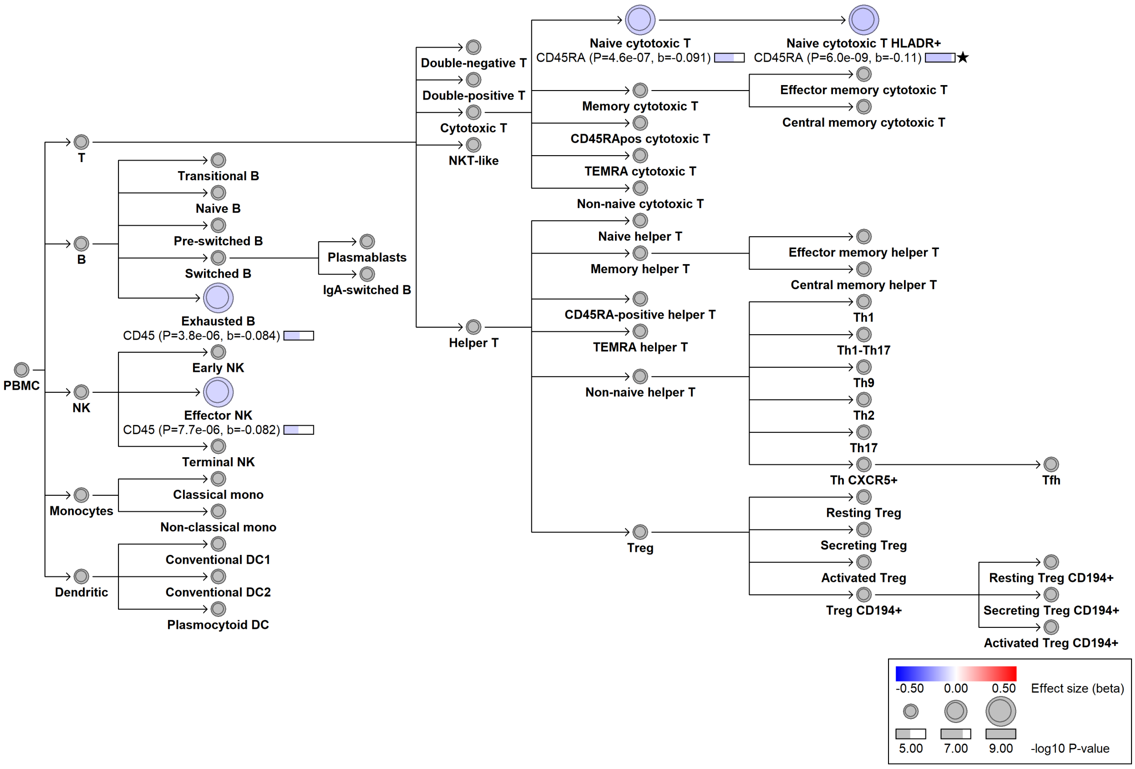

#### chr17:40608272[T→A; EAF 1.152%] (rs112401631, cs_201)

Candidate genes: *CCR7*
Diseases associations: Allergy, Asthma, Eczema
Plasma pQTLs: GZMK

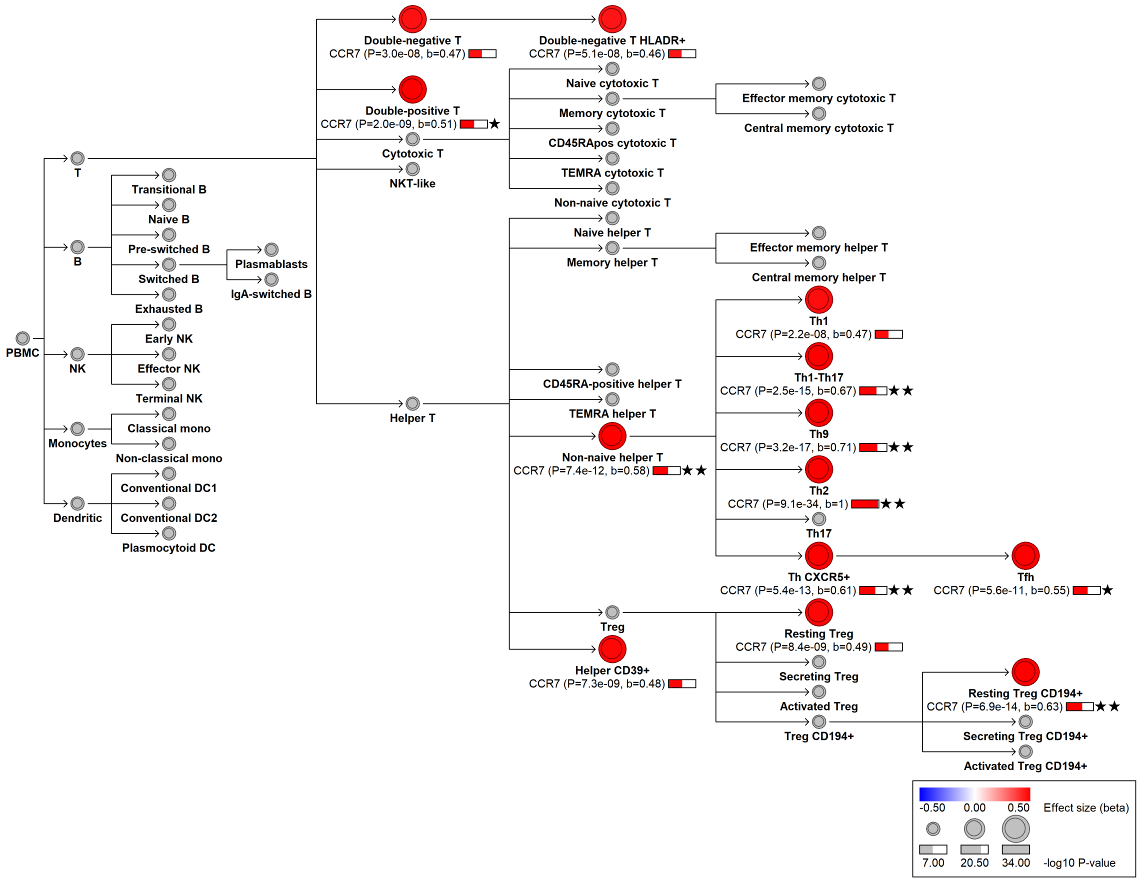

#### chr17:40624926[C→A; EAF 36.339%] (rs1013971, cs_228)

Candidate genes: *CCR7*
Diseases associations: Allergy, Asthma, Eczema, Inflammatory bowel disease, Type 1 diabetes, Ulcerative colitis
Plasma pQTLs: NA

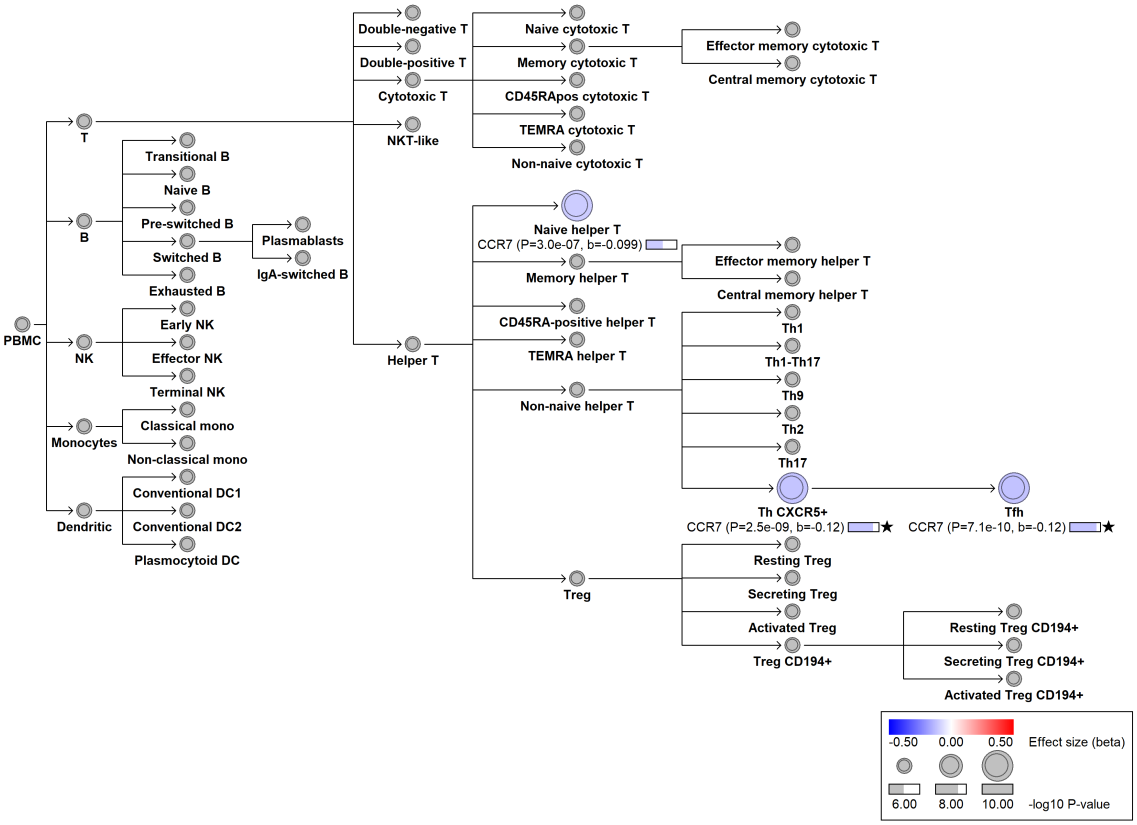

#### chr17:45854491[GAGAGAC→!; EAF 28.931%] (chr17:45854491, cs_37)

Candidate genes: *NA*
Diseases associations: Breast cancer, Chronic obstructive pulmonary disease, Depression, Glaucoma, Osteoarthritis, Ovarian cancer, Parkinson's disease, Primary biliary cholangitis, Progressive supranuclear palsy, Pulmonary fibrosis, Systemic lupus erythematosus, Type 1 diabetes
Plasma pQTLs: NA

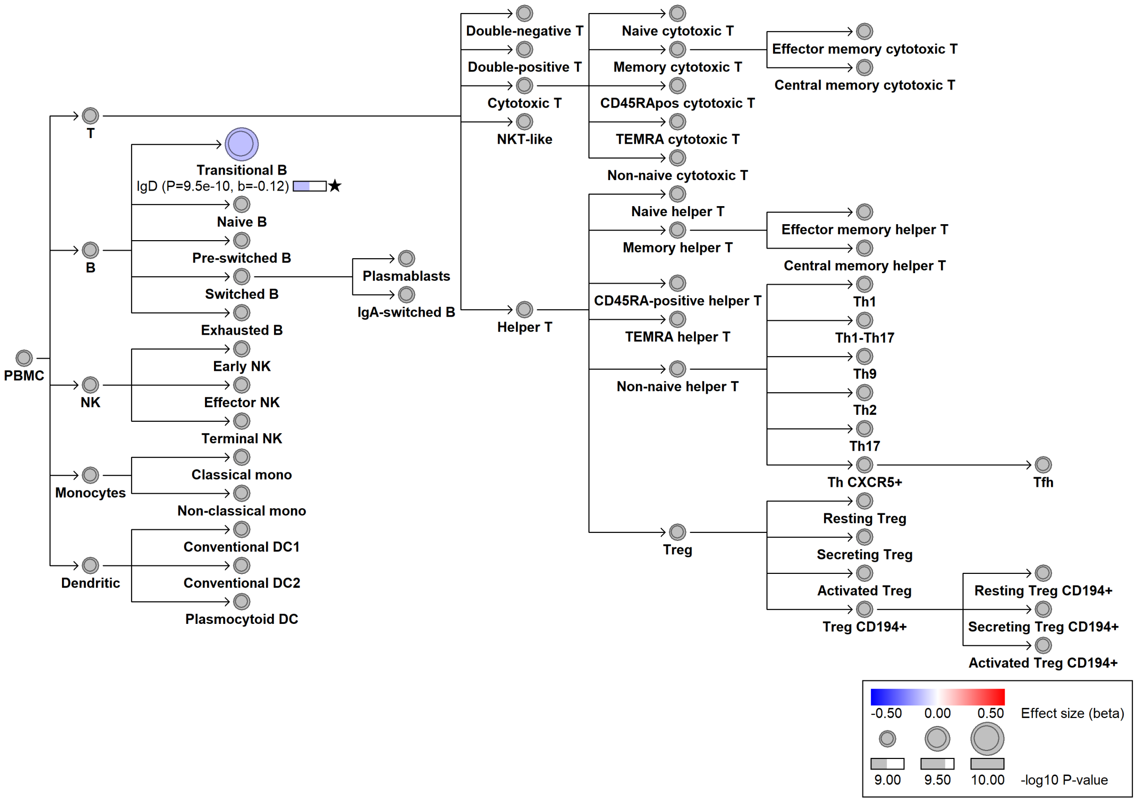

### chr17:58278036[G→A; EAF 2.946%] (rs28730837, cs_212)

Candidate genes: *MPO*
Diseases associations: Myeloperoxidase deficiency
Plasma pQTLs: NA

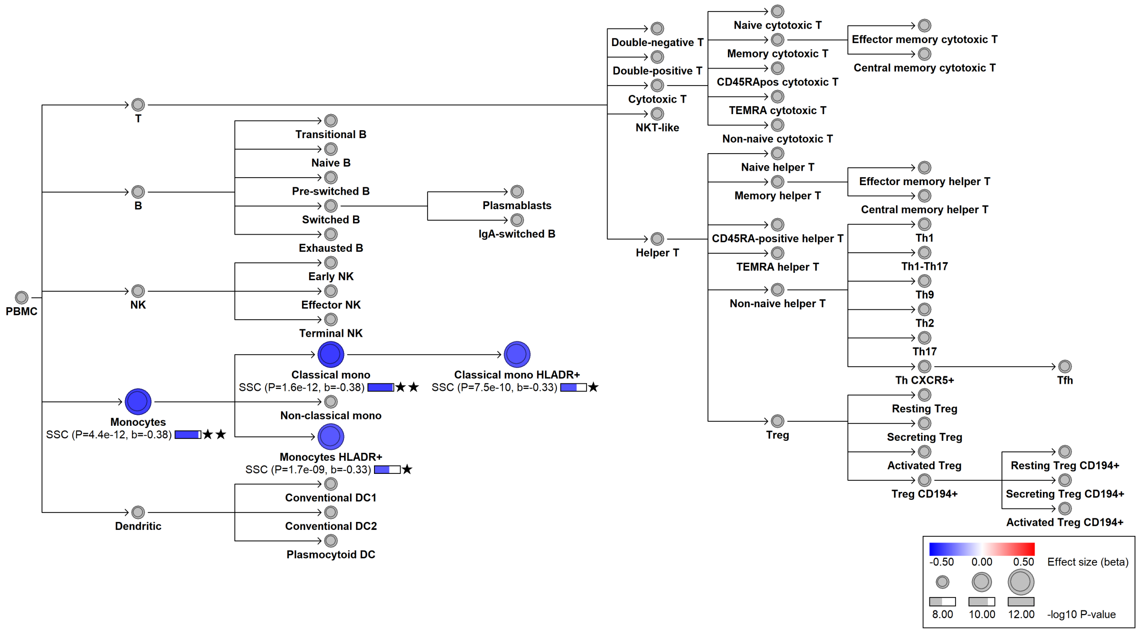

#### chr17:58279141[A→G; EAF 1.465%] (rs56378716, cs_20)

Candidate genes: *MPO*
Diseases associations: Myeloperoxidase deficiency
Plasma pQTLs: CEACAM6, CEACAM8, ELANE, ICAM3, MANSC1, OLFM4, PRSS57, PRTN3

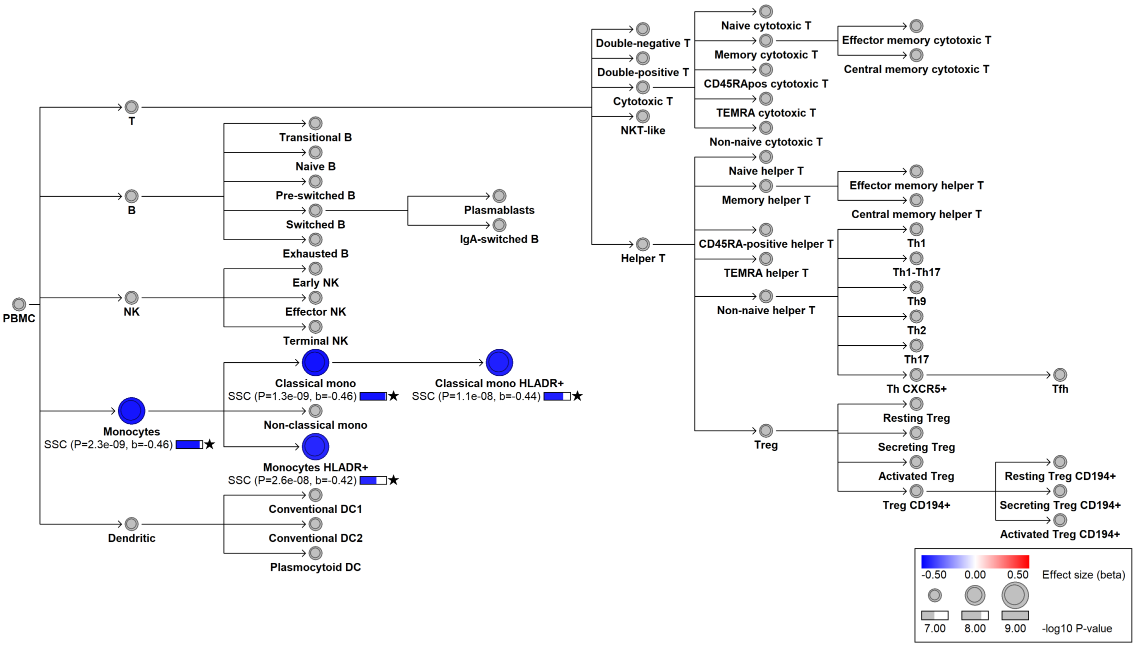

#### chr17:63937688[G→T; EAF 36.758%] (rs4968674, cs_25)

Candidate genes: *CD79B*
Diseases associations: NA
Plasma pQTLs: CD79B, Composite immunoglobulin trait IgA/IgG, Immunoglobulin A, Immunoglobulin G

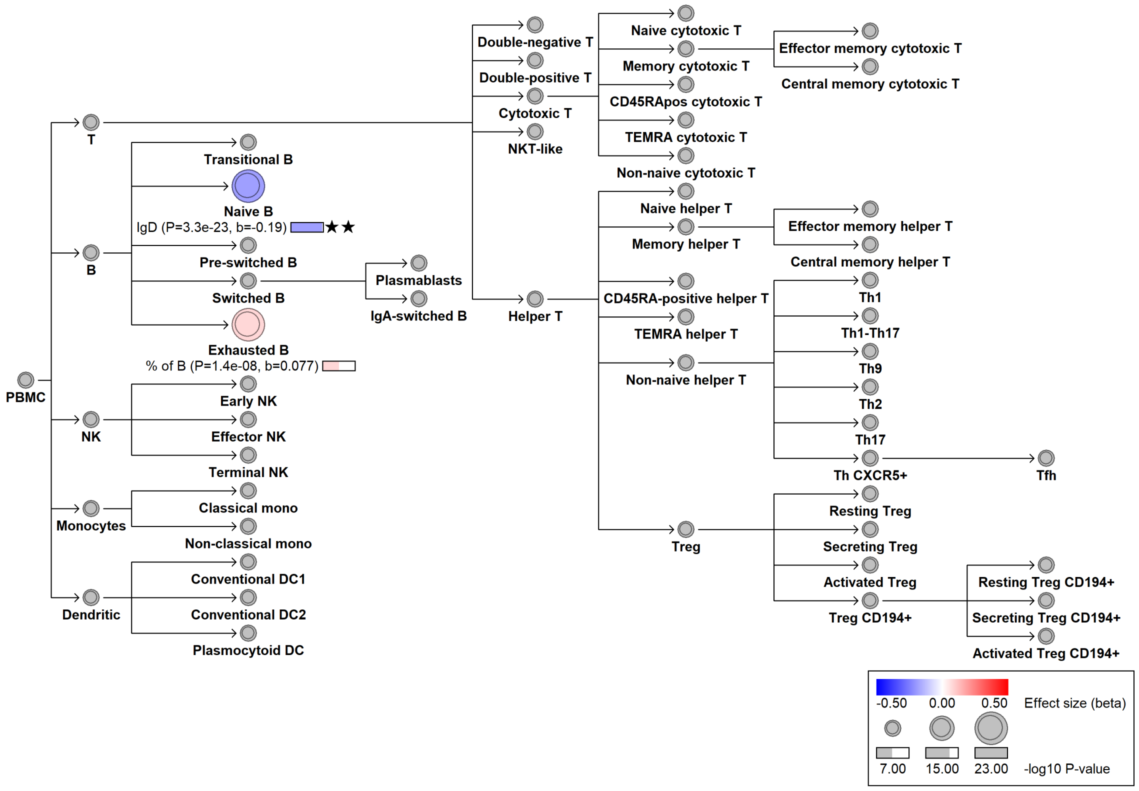

#### chr17:78125237[G→A; EAF 8.885%] (rs12449858, cs_222)

Candidate genes: *TMC6*
Diseases associations: NA
Plasma pQTLs: TCL1A

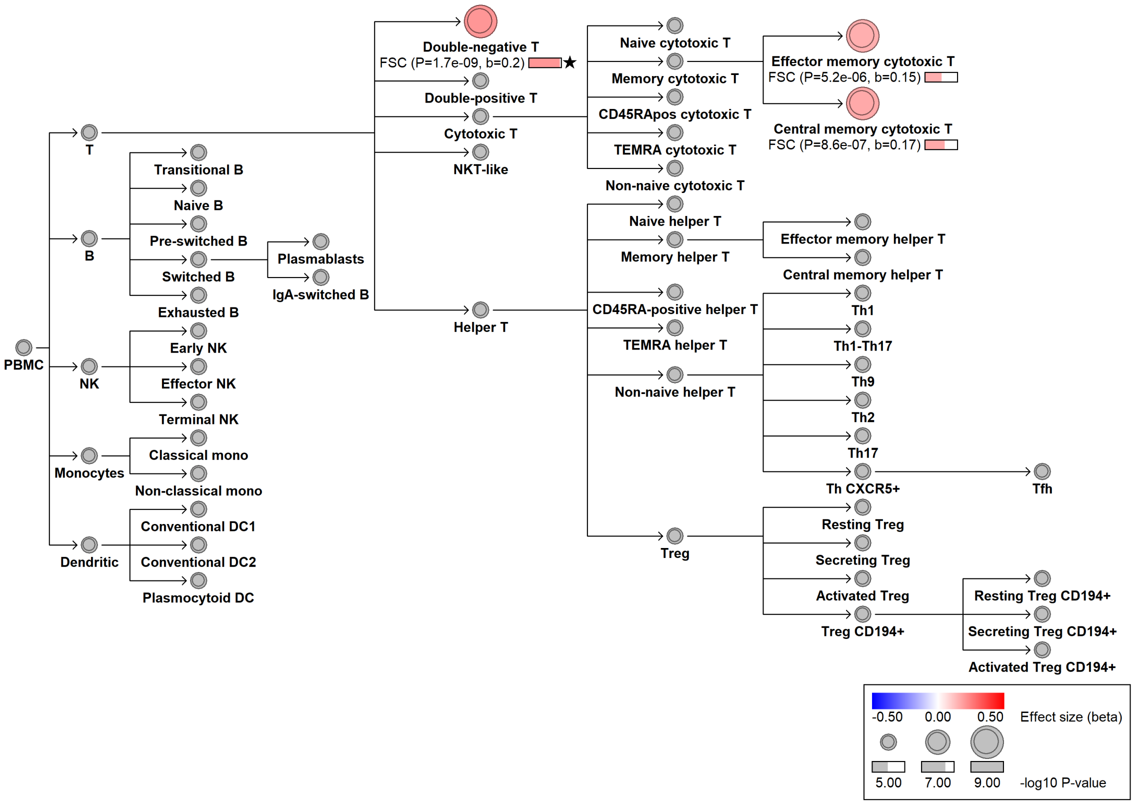

#### chr17:78134494[A→T; EAF 48.465%] (rs7208422, cs_16)

Candidate genes: *TMC8*
Diseases associations: NA
Plasma pQTLs: CD6, CD7, EZR, PAG1, SH2D1A, SIT1, SKAP1

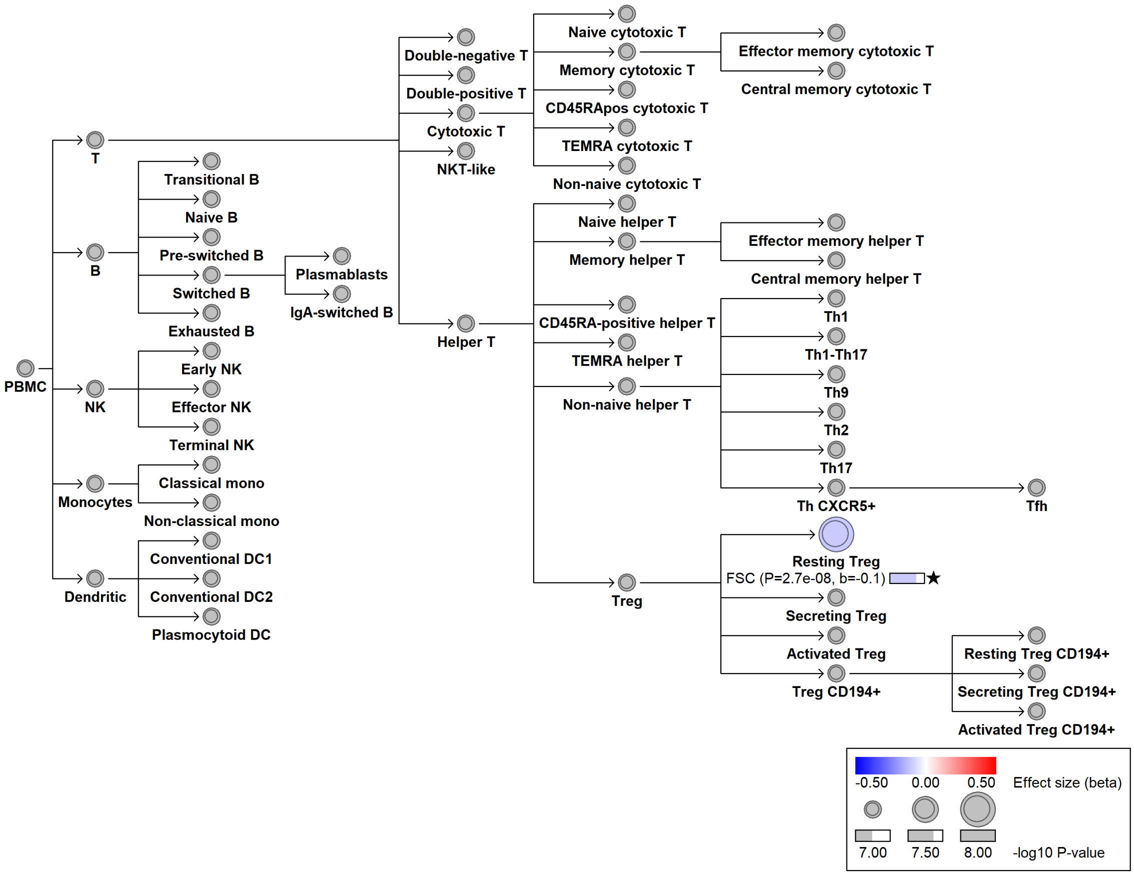

#### chr18:2002725[T→C; EAF 47.47%] (rs1940647, cs_202)

Candidate genes: *NA*
Diseases associations: NA
Plasma pQTLs: NA

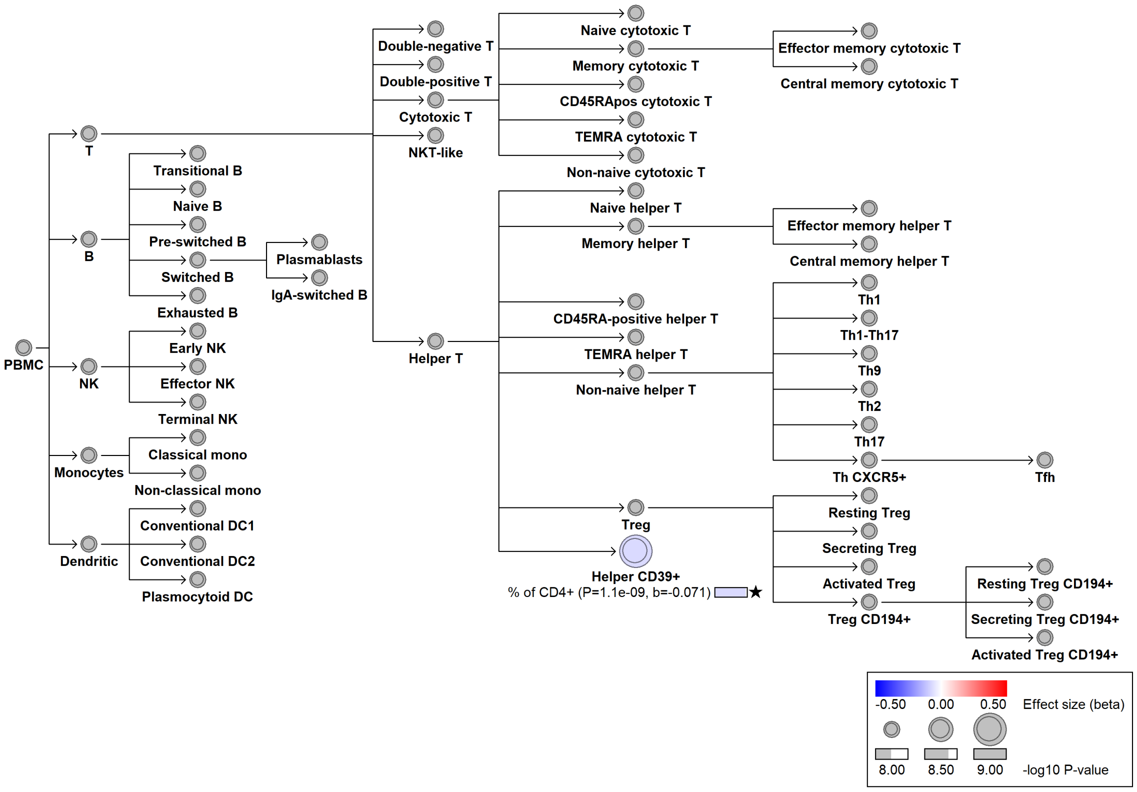

#### chr18:63193013[C→T; EAF 28.637%] (rs12457700, cs_65)

Candidate genes: *BCL2*
Diseases associations: Adolescent idiopathic scoliosis
Plasma pQTLs: NA

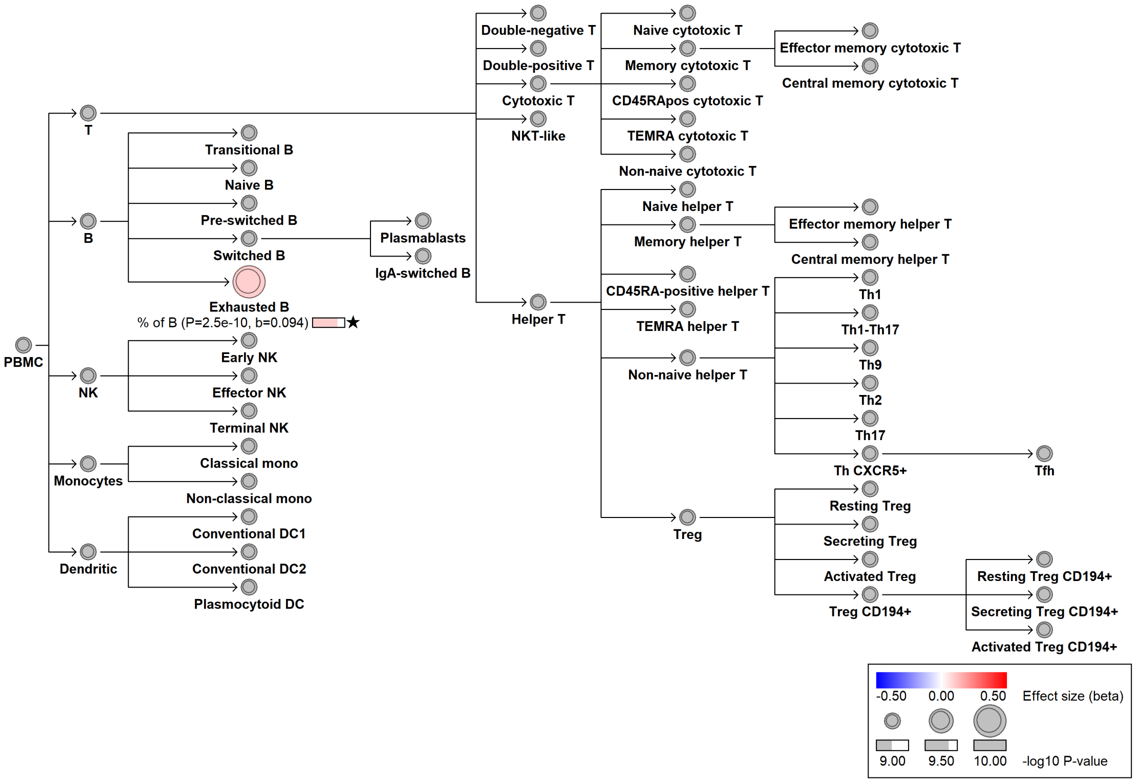

### chr19:851976[G→A; EAF 23.411%] (rs2007647, cs_180)

Candidate genes: *ELANE*
Diseases associations: NA
Plasma pQTLs: NA

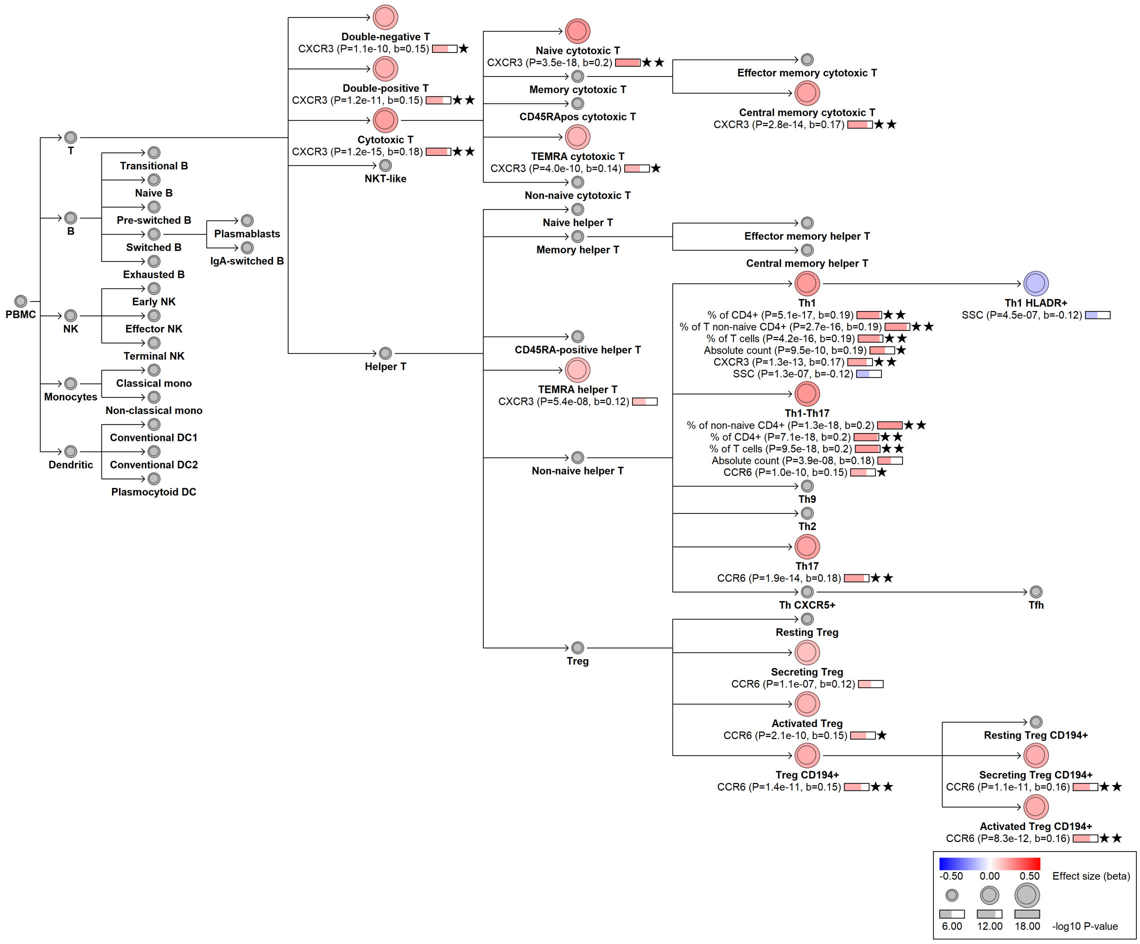

#### chr19:7350159[G→A; EAF 3.471%] (rs62111672, cs_192)

Candidate genes: *ARHGEF18*
Diseases associations: NA
Plasma pQTLs: NA

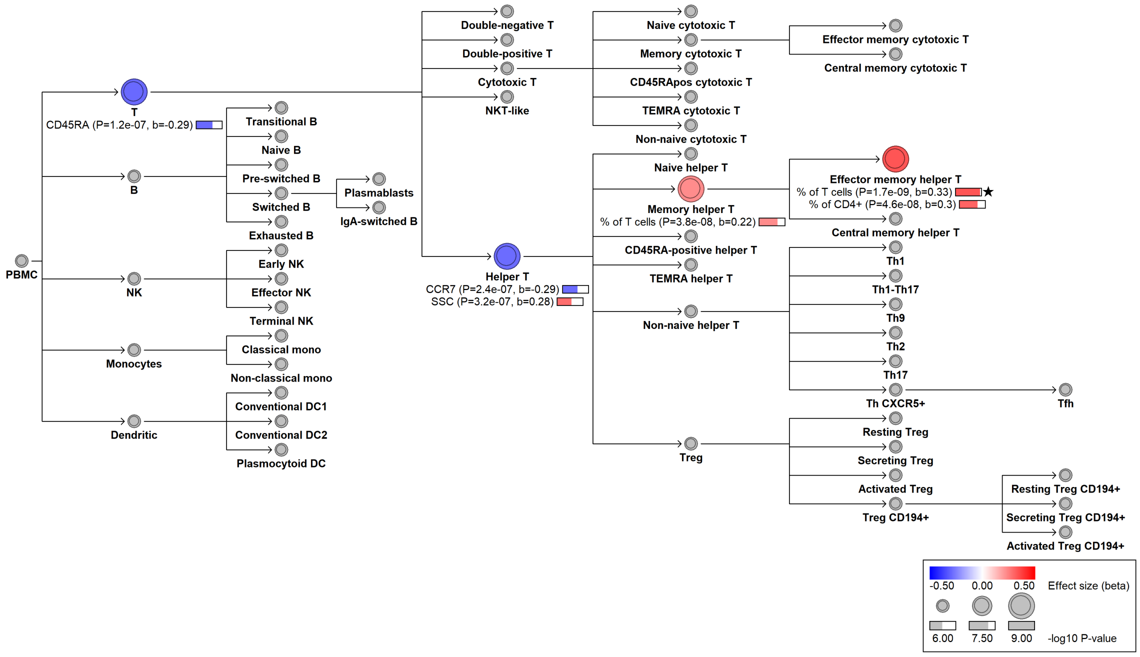

#### chr19:8677438[C→T; EAF 14.469%] (rs2918299, cs_120)

Candidate genes: *NFILZ*
Diseases associations: Allergy, Asthma, Chronic diseases of tonsils, Eczema, Endometriosis
Plasma pQTLs: NA

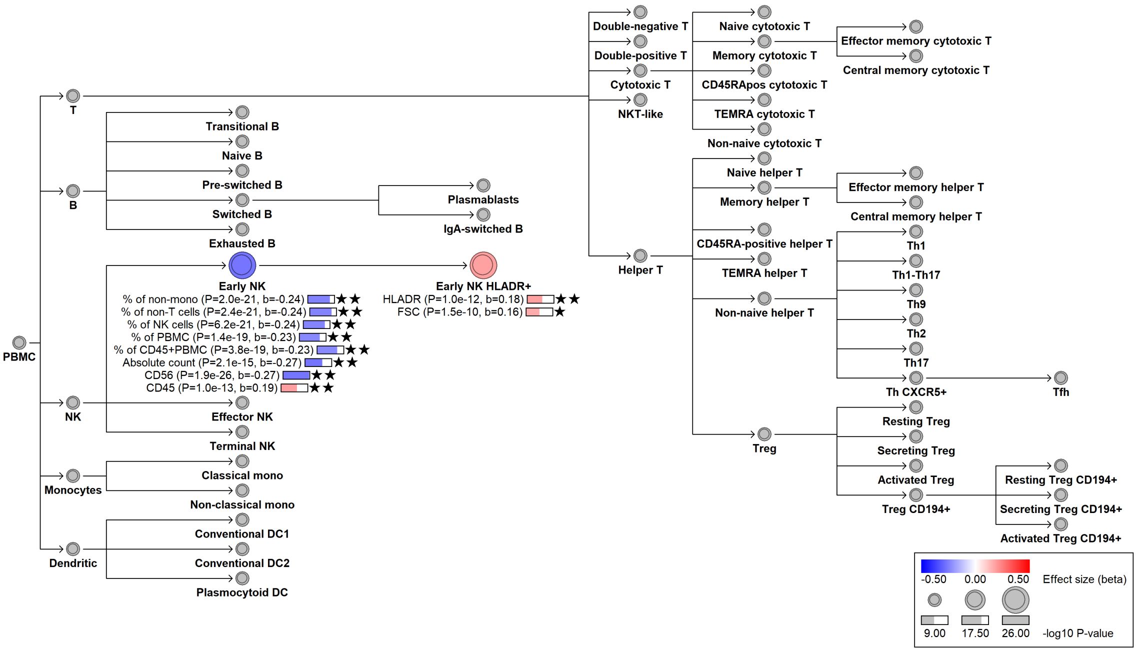

#### chr19:16327617[GC→G; EAF 39.147%] (rs35663614, cs_200)

Candidate genes: *KLF2*
Diseases associations: NA
Plasma pQTLs: INSR

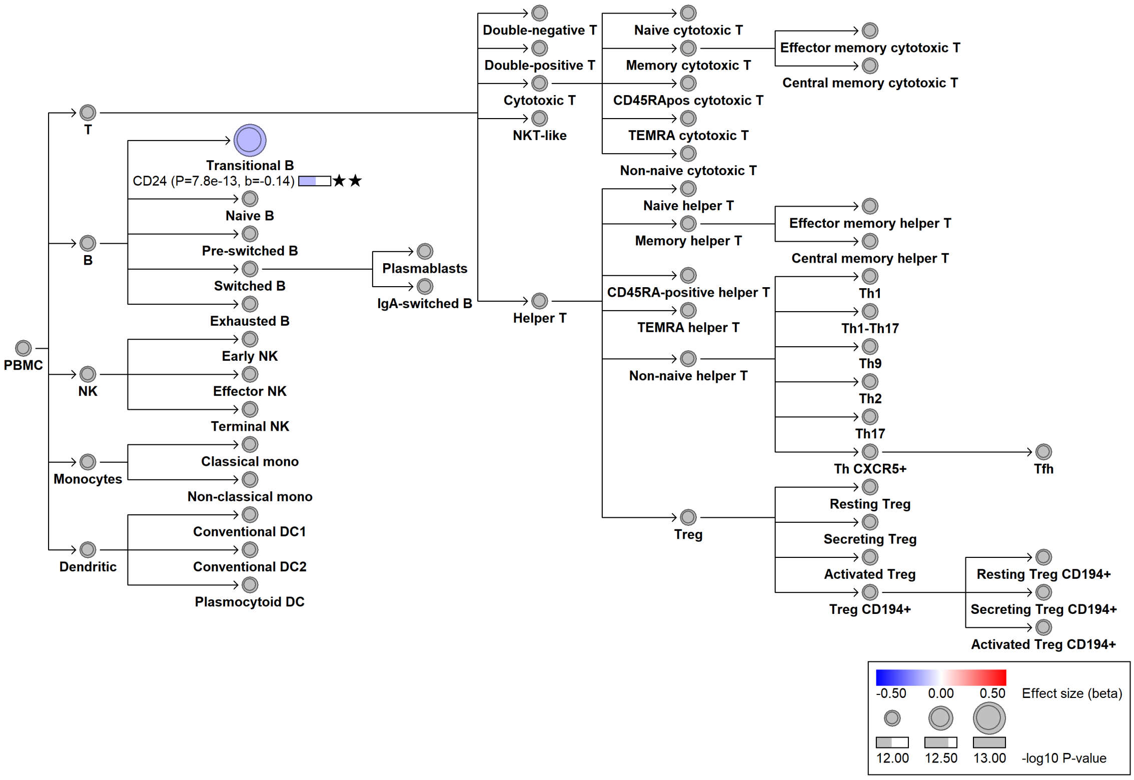

### chr19:17549491[A→G; EAF 46.672%] (rs11666267, cs_129)

Candidate genes: *NIBAN3*
Diseases associations: NA
Plasma pQTLs: CD5L, FCER2, Immunoglobulin M, TCL1A, TNFRSF13C

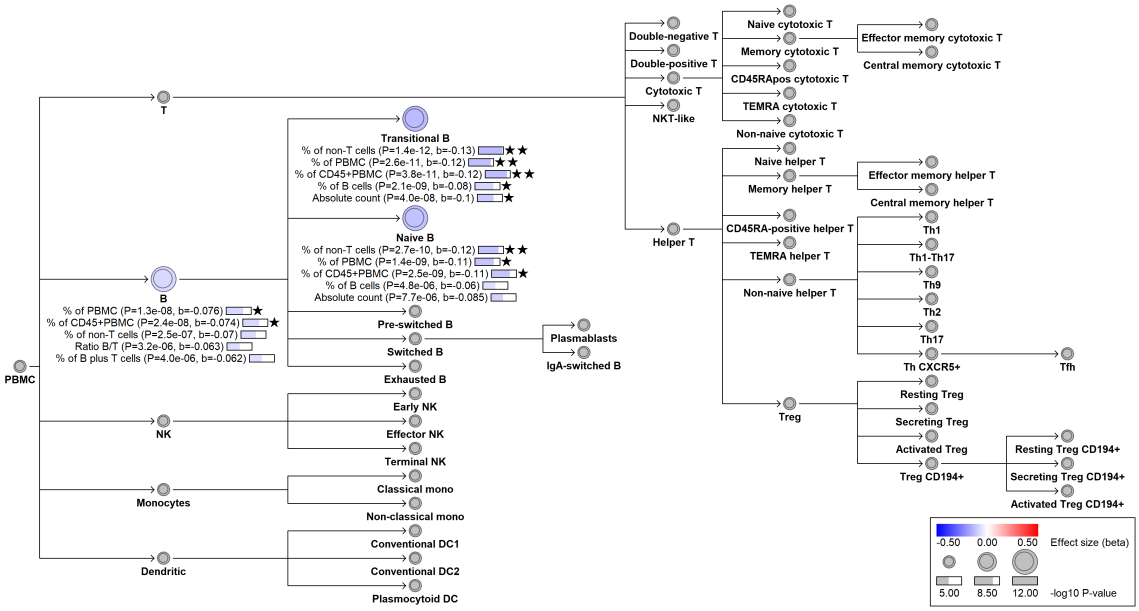

### chr19:18175134[!→A; EAF 22.327%] (rs11554159, cs_226)

Candidate genes: *IFI30*
Diseases associations: Multiple sclerosis
Plasma pQTLs: AFM, CTSH, SIGLEC1

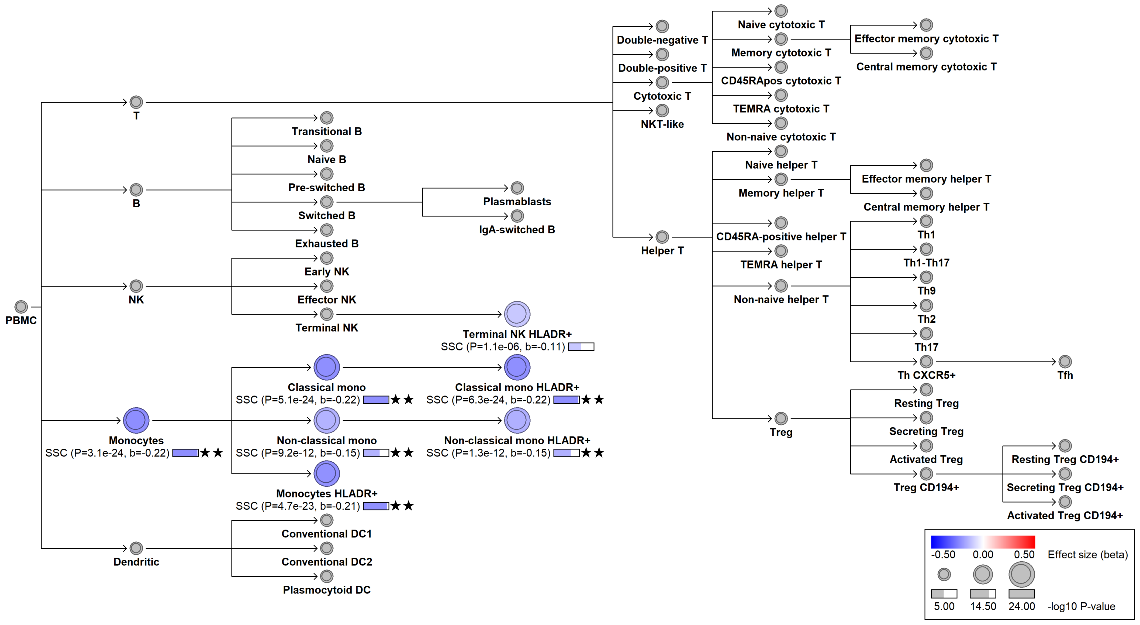

#### chr19:33263642[C→T; EAF 9.946%] (rs78744187, cs_150)

Candidate genes: *CEBPA*
Diseases associations: NA
Plasma pQTLs: EPO, KIT, SIGLEC6, TPSAB1, TPSB2, TPSD1

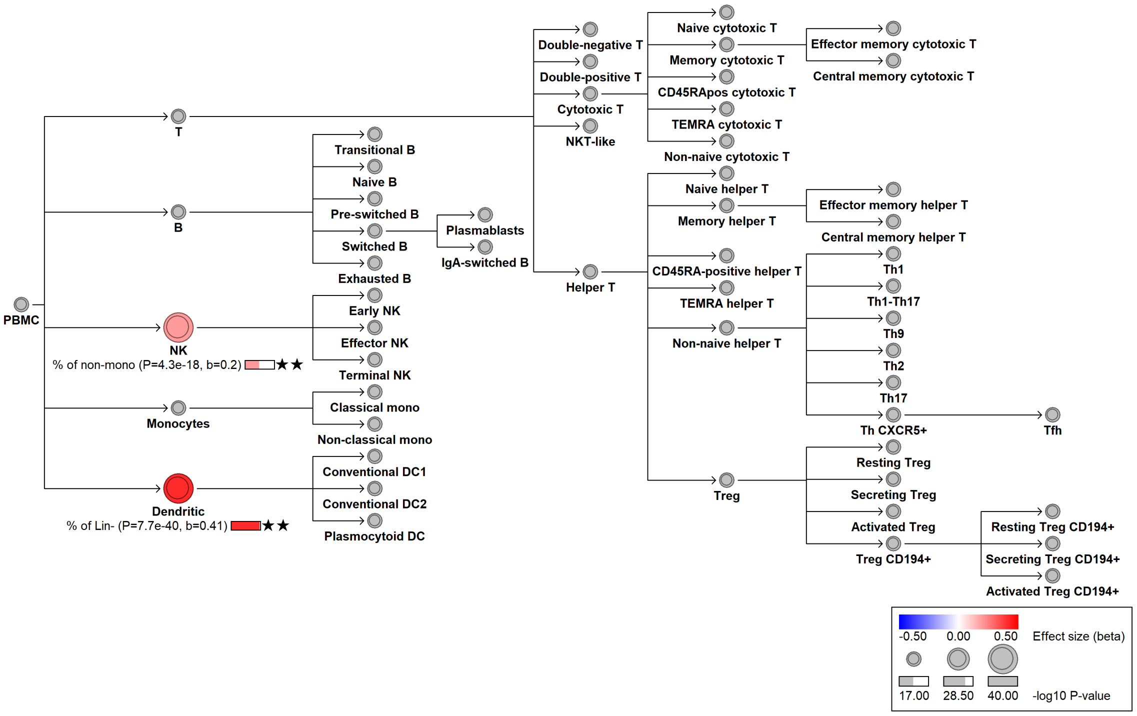

#### chr19:35169605[A→!; EAF 20.89%] (rs12110, cs_72)

Candidate genes: *FXYD1, FXYD5, FXYD7*
Diseases associations: NA
Plasma pQTLs: CEACAM6, CEL, CNN1, FXYD5, ITM2A, SELPLG

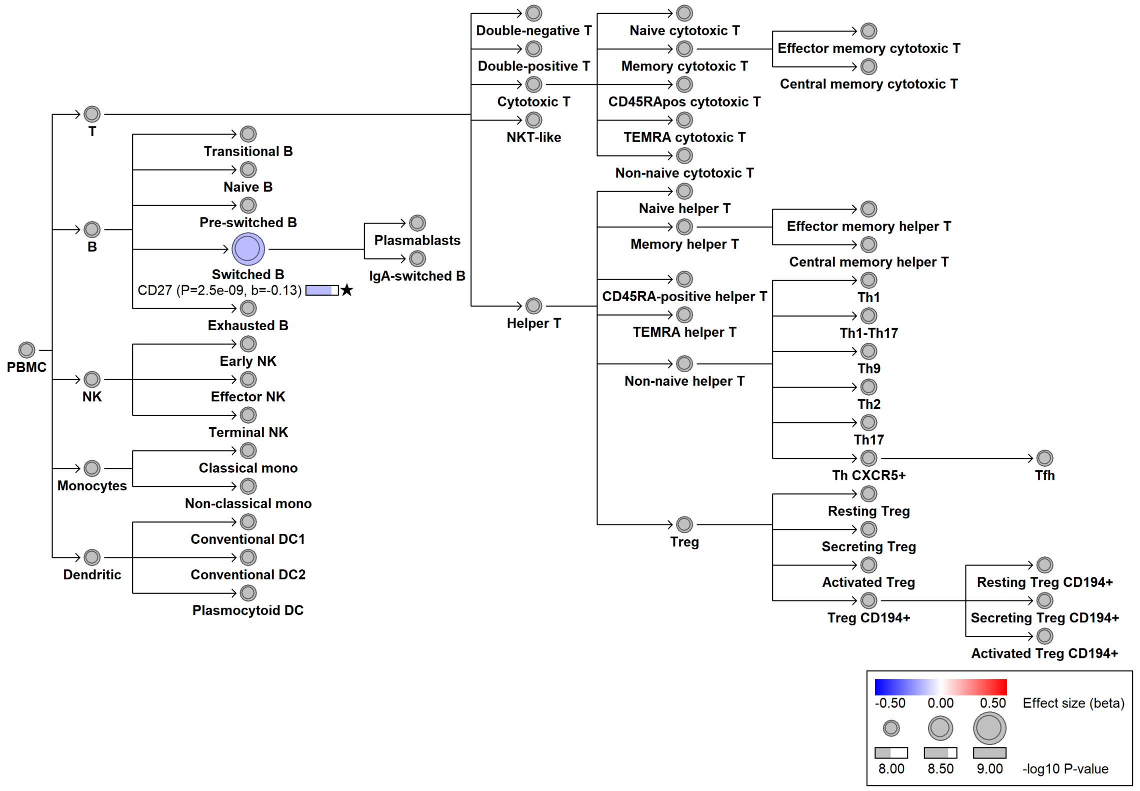

#### chr19:51577724[C→T; EAF 1.053%] (rs558374434, cs_179)

Candidate genes: *NA*
Diseases associations: NA
Plasma pQTLs: NA

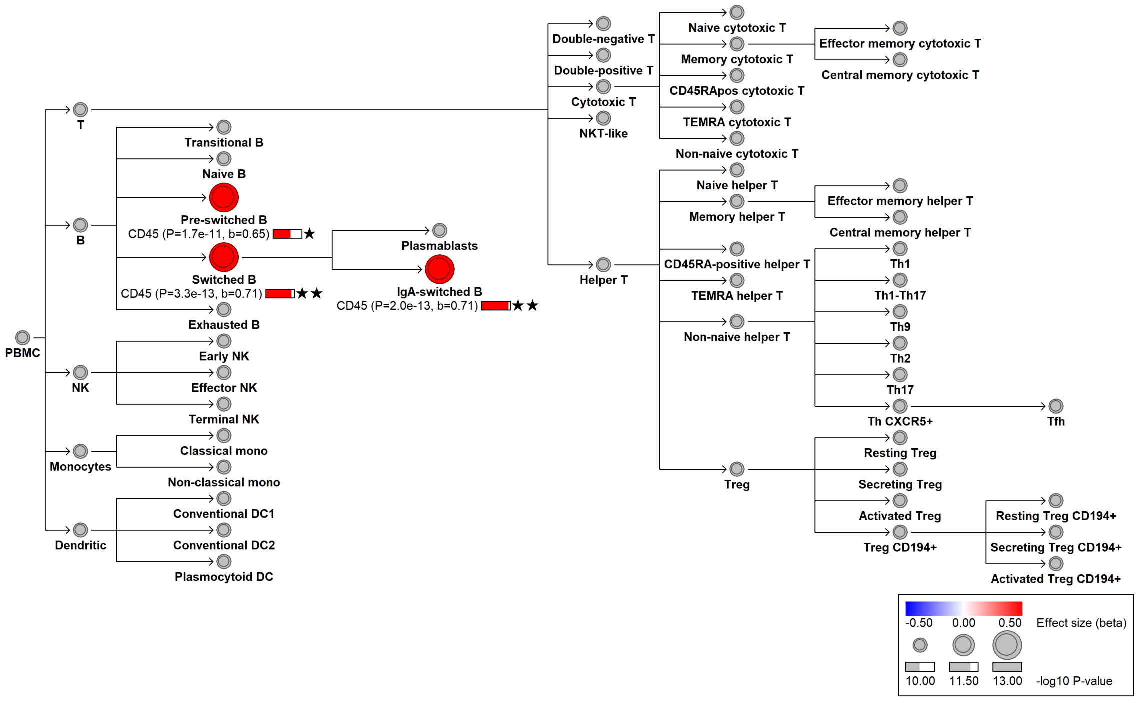

#### chr19:52840865[!→G; EAF 44.83%] (rs10419826, cs_158)

Candidate genes: *ZNF468*
Diseases associations: NA
Plasma pQTLs: NA

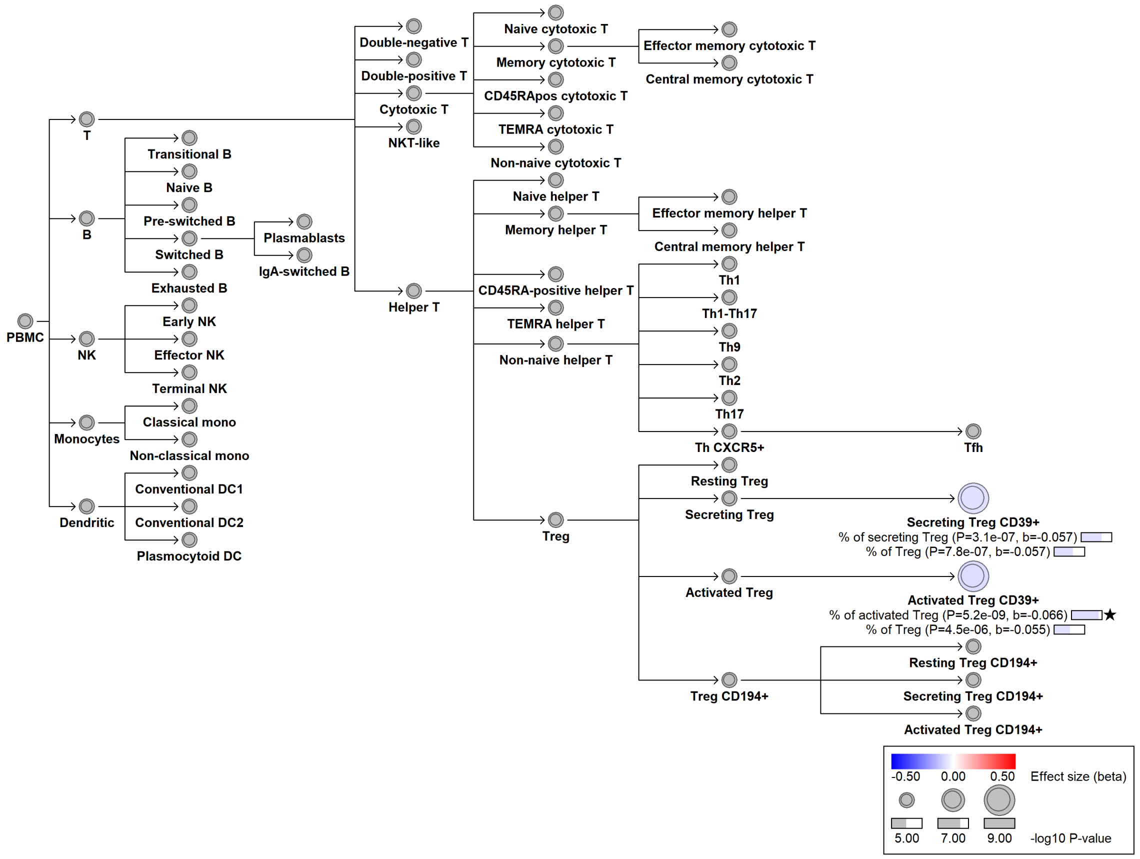

#### chr19:53810453[G→C; EAF 7.711%] (rs34971363, cs_70)

Candidate genes: *NLRP12*
Diseases associations: NA
Plasma pQTLs: NA

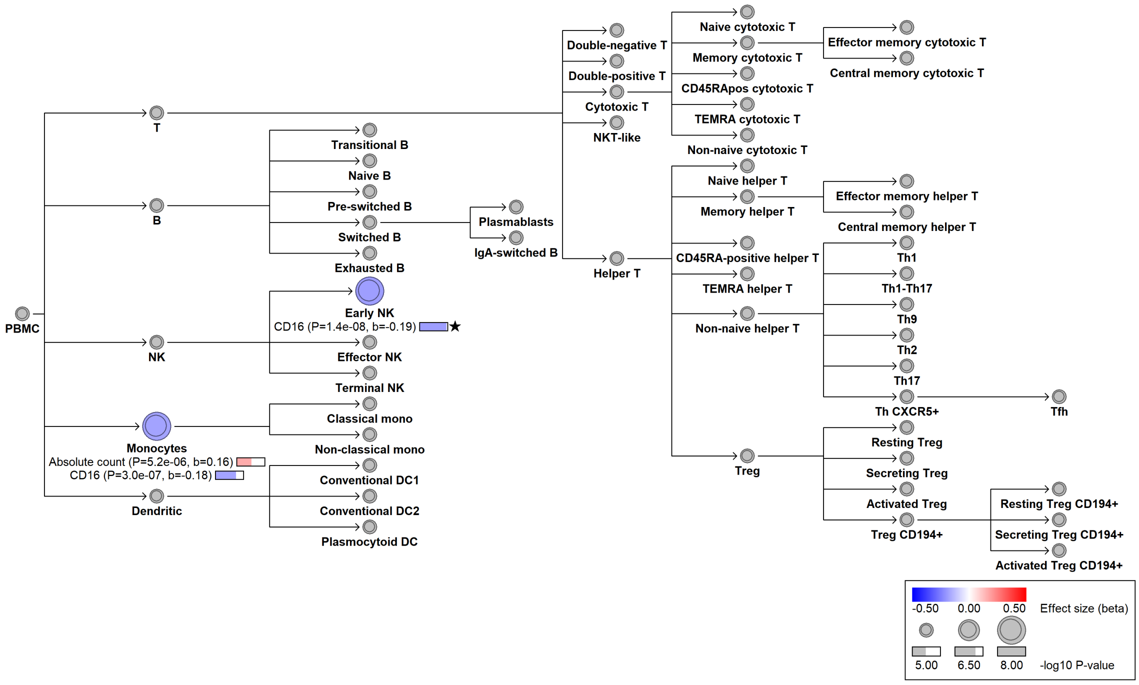

##### chr19:53824059[C→A; EAF 22.657%] (rs34436714, cs_9)

Candidate genes: *NLRP12*
Diseases associations: NA
Plasma pQTLs: AIF1, ANXA3, APBB1IP, APEX1, BAG4, BCL2L15, BID, CAPG, CASP1, CHCHD10, CLC, DGCR6, EGLN1, ELOA, FEN1, FGR, GIMAP7, GIMAP8, HCLS1, HNMT, IDO1, IL1B, IL1RN, KLF4, LGALS1, LRCH4, NCF2, NEDD9, PARP1, PIKFYVE, PXN, RBP7, S100A11, S100A12, S100P, SMNDC1, TIGAR, TMSB10, TRIAP1, ZBP1

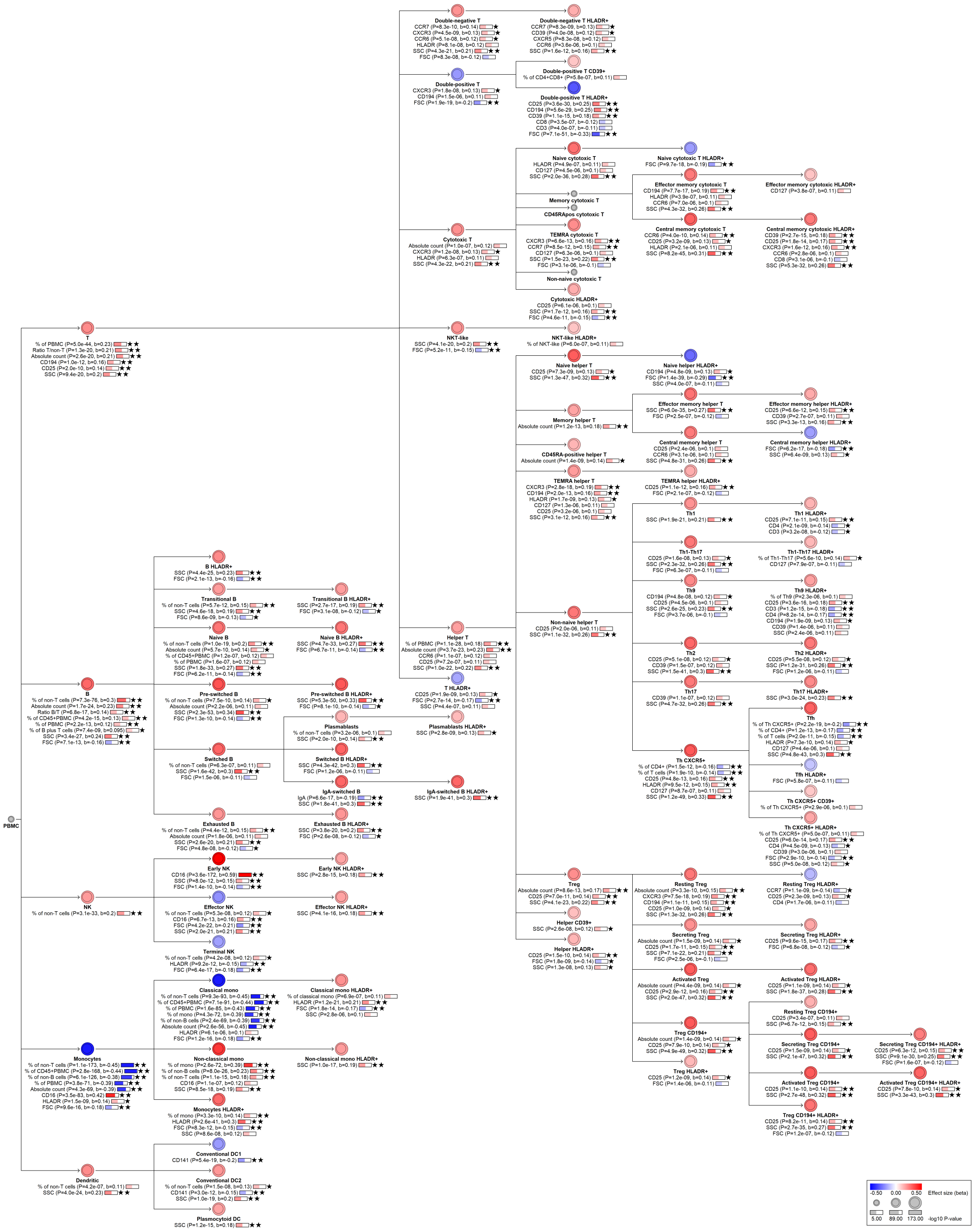
-0.50 0.00 0.50 Effect size (beta)
5.00 89.00 173.00 -log10 P-value

#### chr19:54241789[C→T; EAF 13.565%] (rs1052966, cs_154)

Candidate genes: *LILRA4, LILRA6, LILRB3, LILRB5*
Diseases associations: NA
Plasma pQTLs: NA

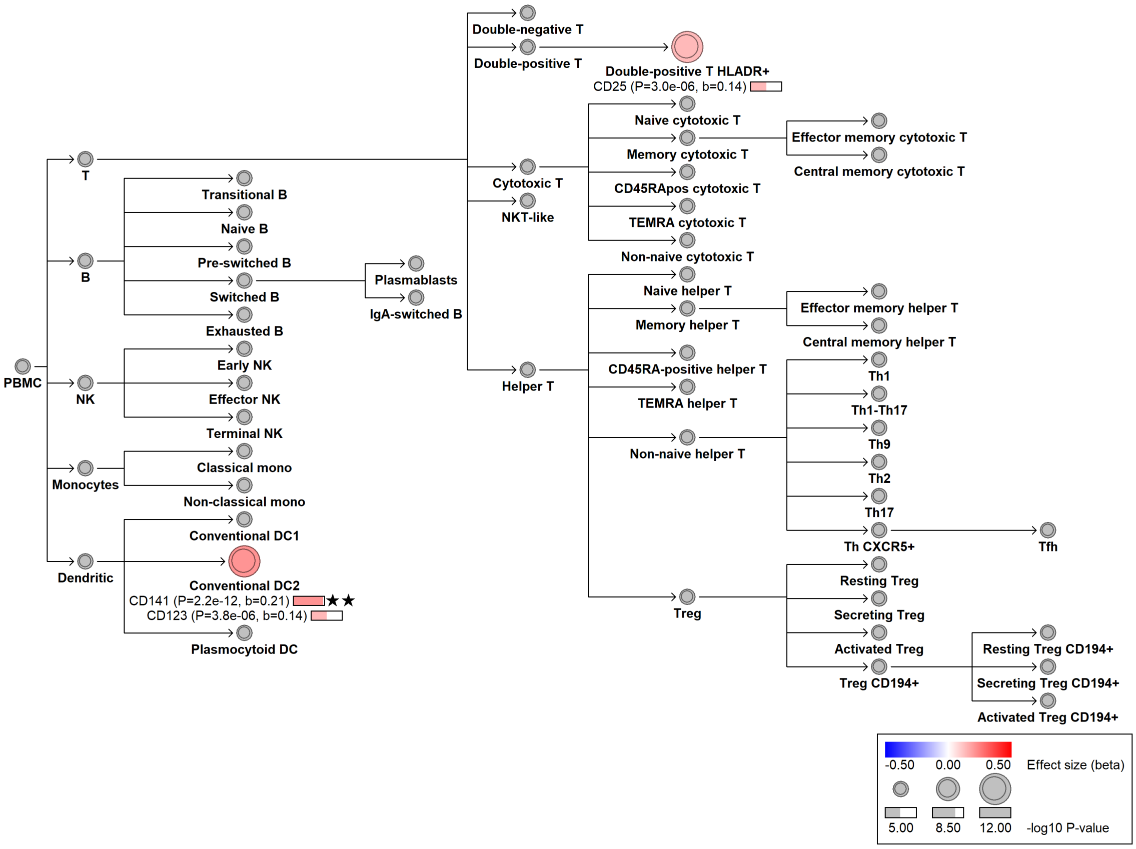

#### chr20:17848636[G→A; EAF 36.181%] (rs2745851, cs_54)

Candidate genes: *RRBP1, SNX5*
Diseases associations: NA
Plasma pQTLs: NA

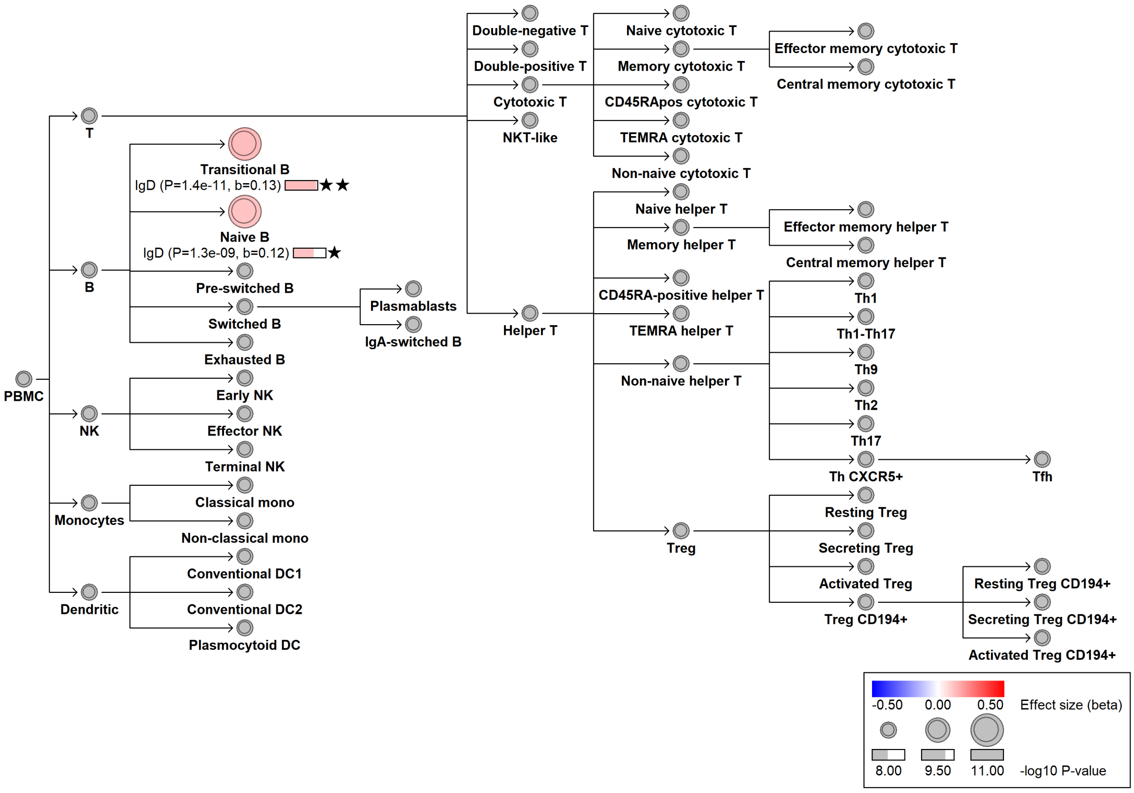

#### chr20:22712498[G→T; EAF 27.752%] (rs2404489, cs_73)

Candidate genes: *THBD*
Diseases associations: NA
Plasma pQTLs: NA

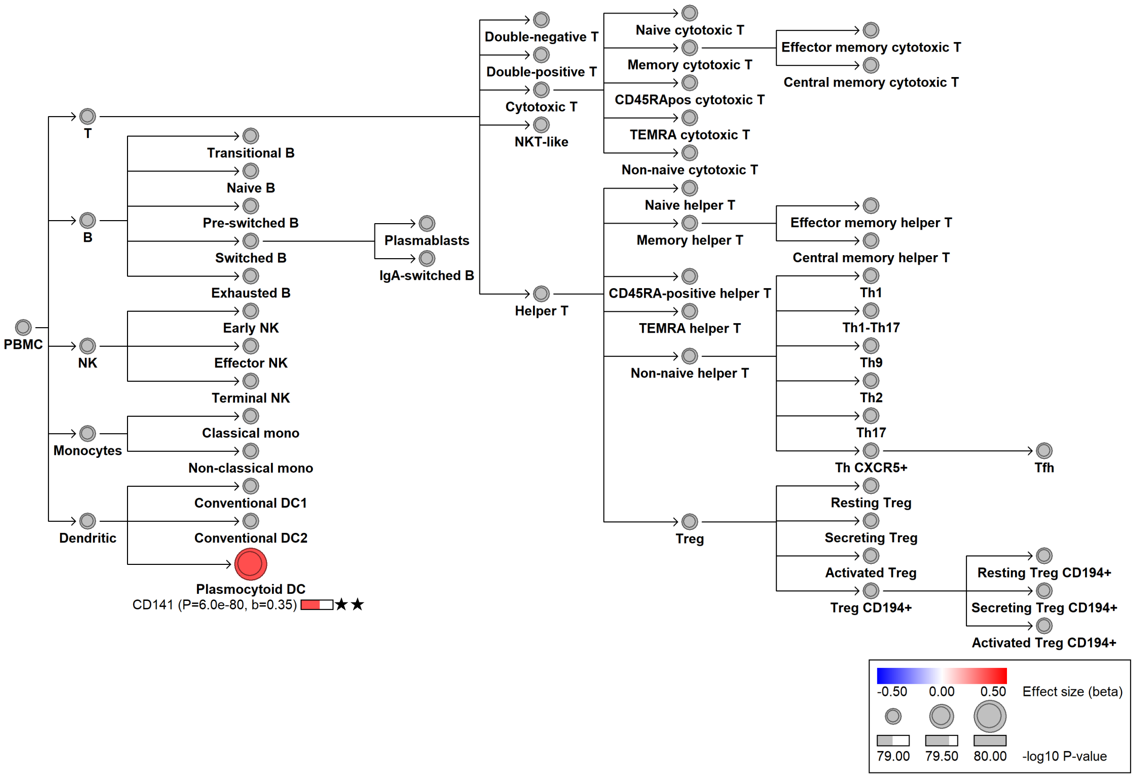

#### chr20:23033049[T→C; EAF 0.324%] (rs561123609, cs_91)

Candidate genes: *THBD*
Diseases associations: NA
Plasma pQTLs: NA

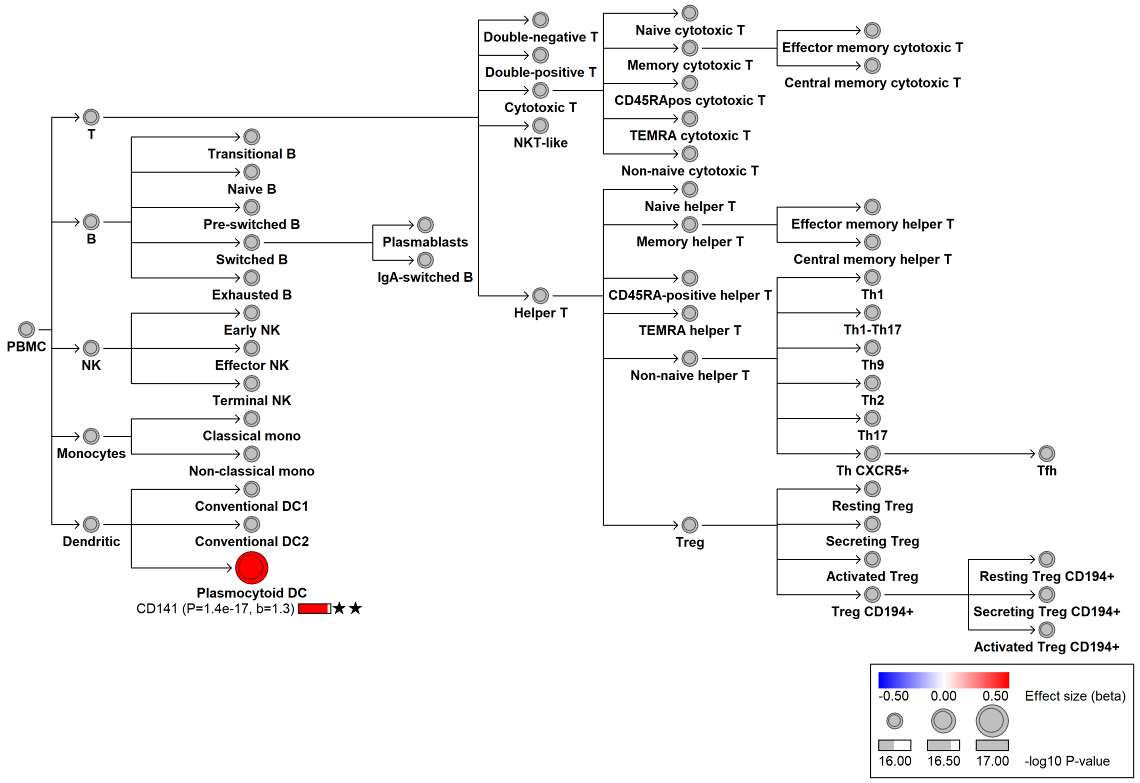

#### chr20:23084705[G→A; EAF 25.596%] (rs3746732, cs_183)

Candidate genes: *THBD, CD93*
Diseases associations: NA
Plasma pQTLs: NA

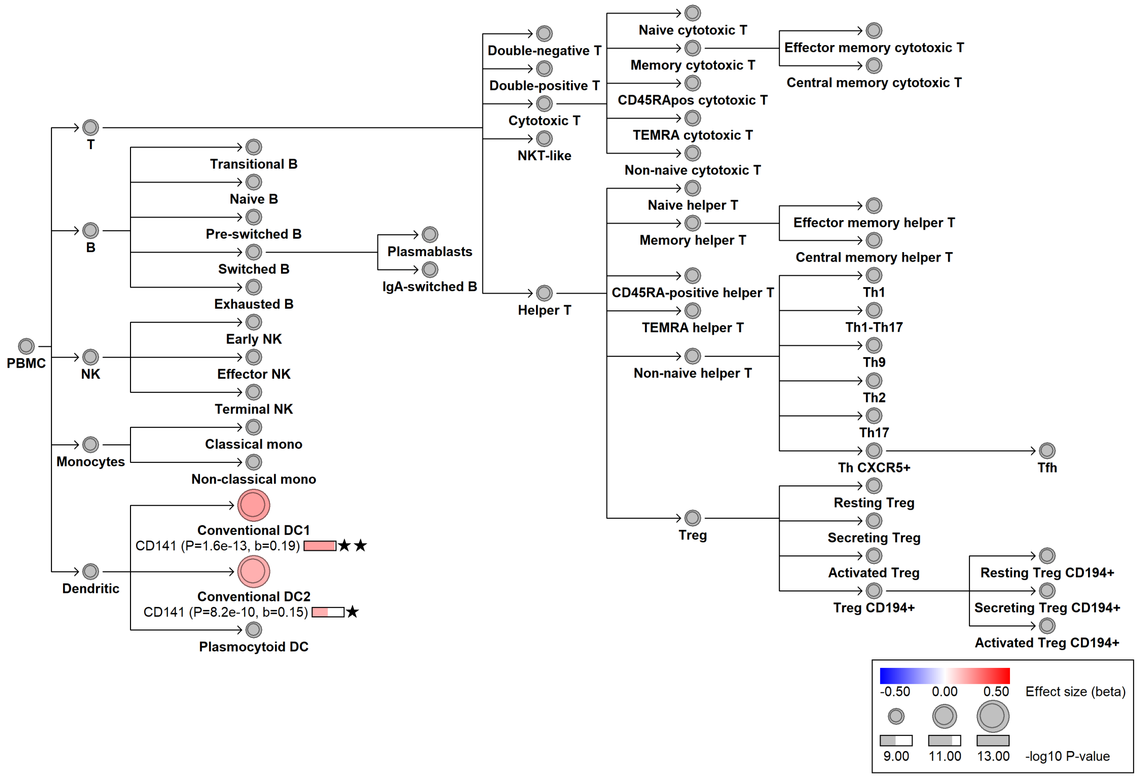

#### chr20:23112819[G→A; EAF 9.982%] (rs844849, cs_18)

Candidate genes: *THBD, CD93*
Diseases associations: NA
Plasma pQTLs: NA

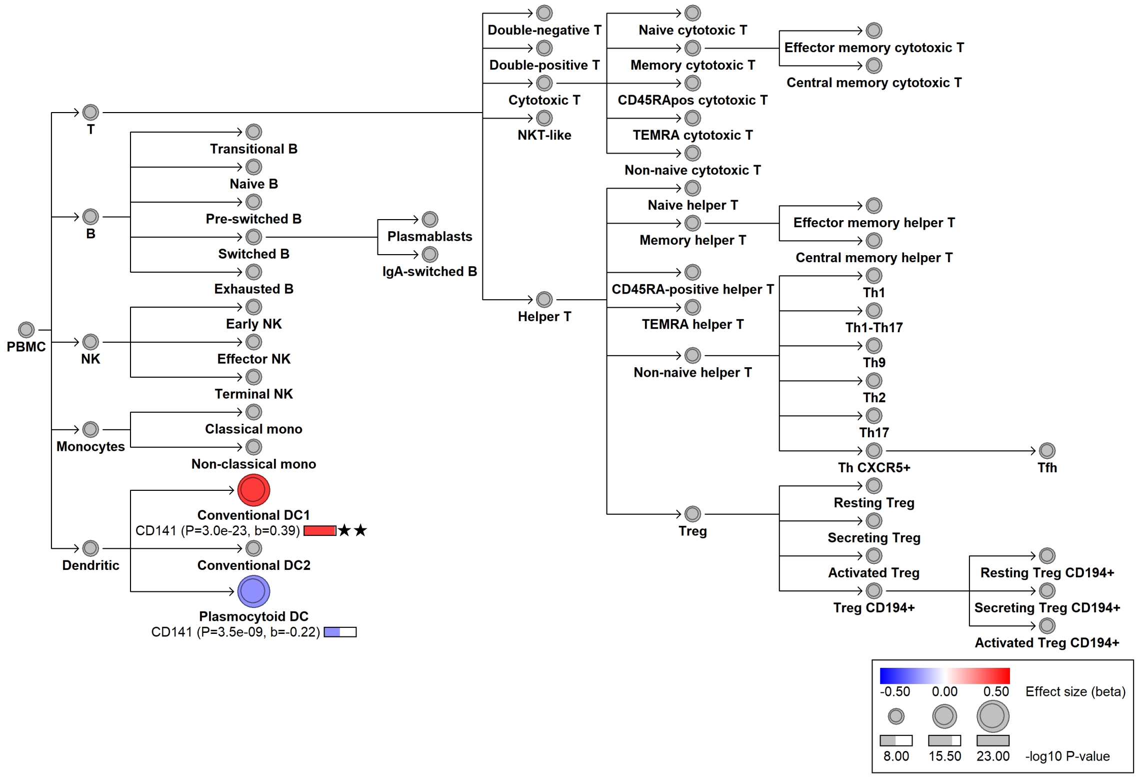

#### chr20:23119854[T→!; EAF 27.743%] (chr20:23119854, cs_31)

Candidate genes: *THBD*
Diseases associations: NA
Plasma pQTLs: NA

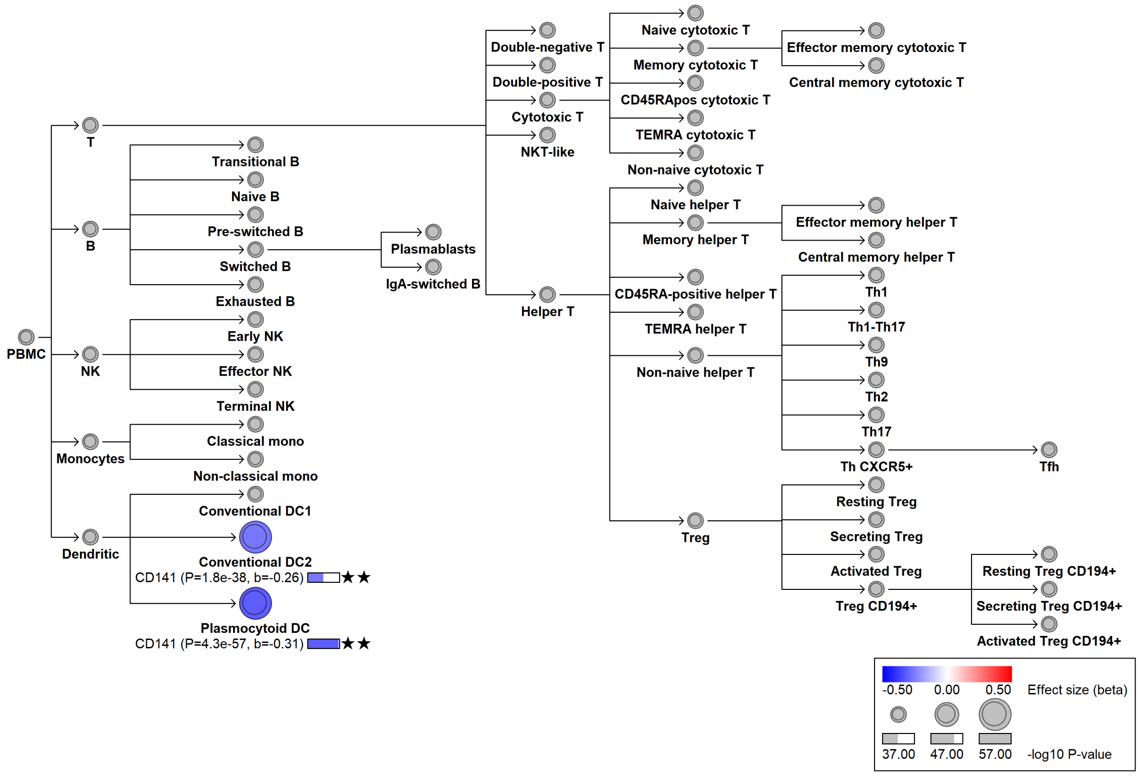

#### chr20:23123022[G→A; EAF 2.615%] (rs80182235, cs_61)

Candidate genes: *THBD*
Diseases associations: NA
Plasma pQTLs: NA

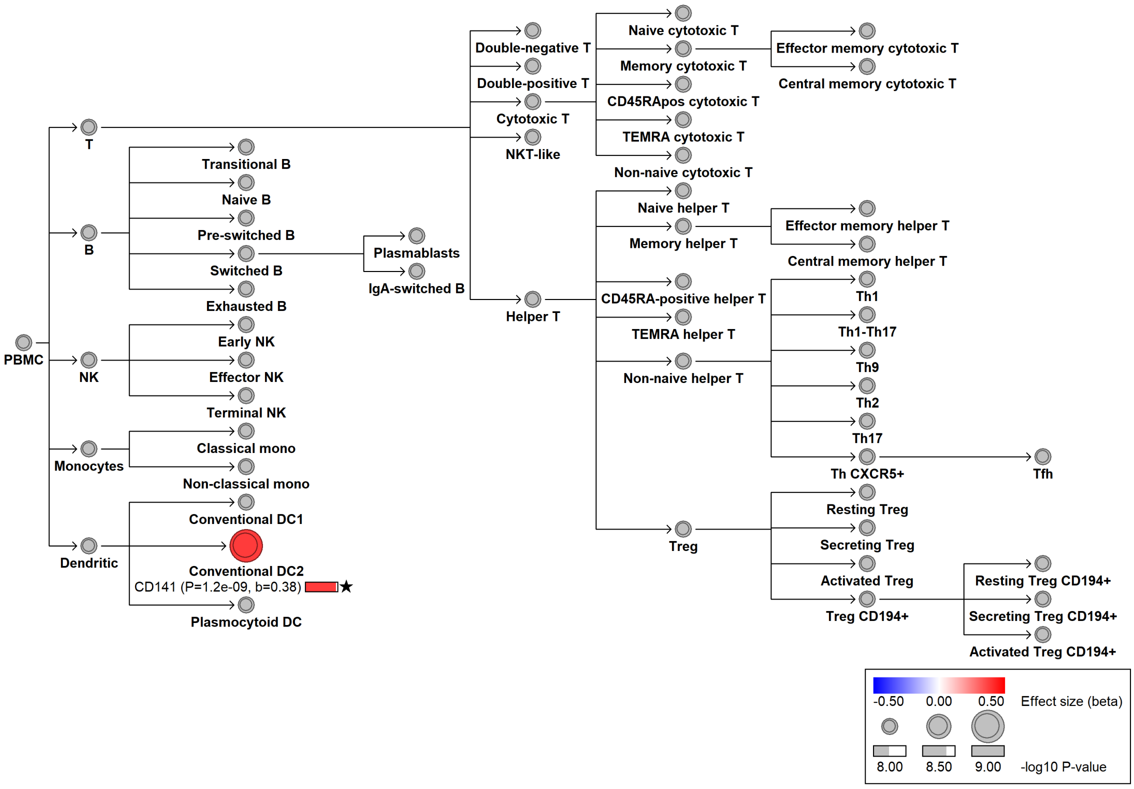

#### chr20:23159856[G→C; EAF 48.744%] (rs999307, cs_63)

Candidate genes: *THBD*
Diseases associations: NA
Plasma pQTLs: NA

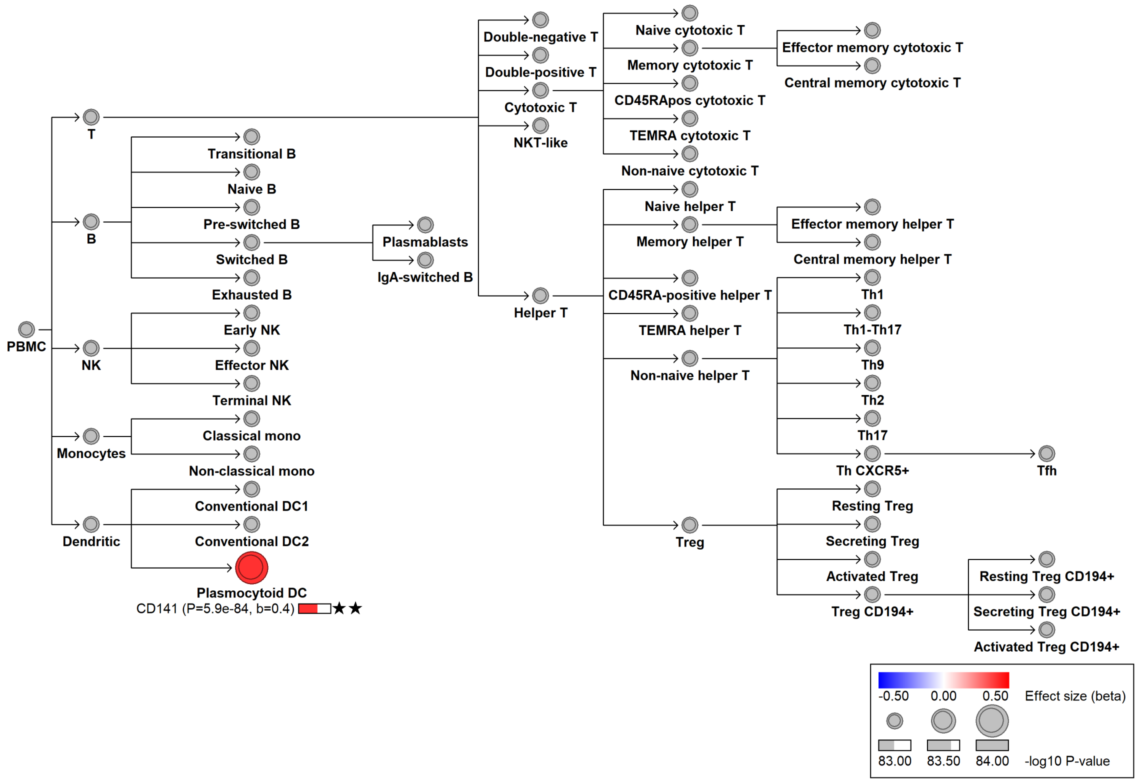

#### chr20:23226600[T→!; EAF 7.332%] (chr20:23226600, cs_2)

Candidate genes: *THBD*
Diseases associations: NA
Plasma pQTLs: NA

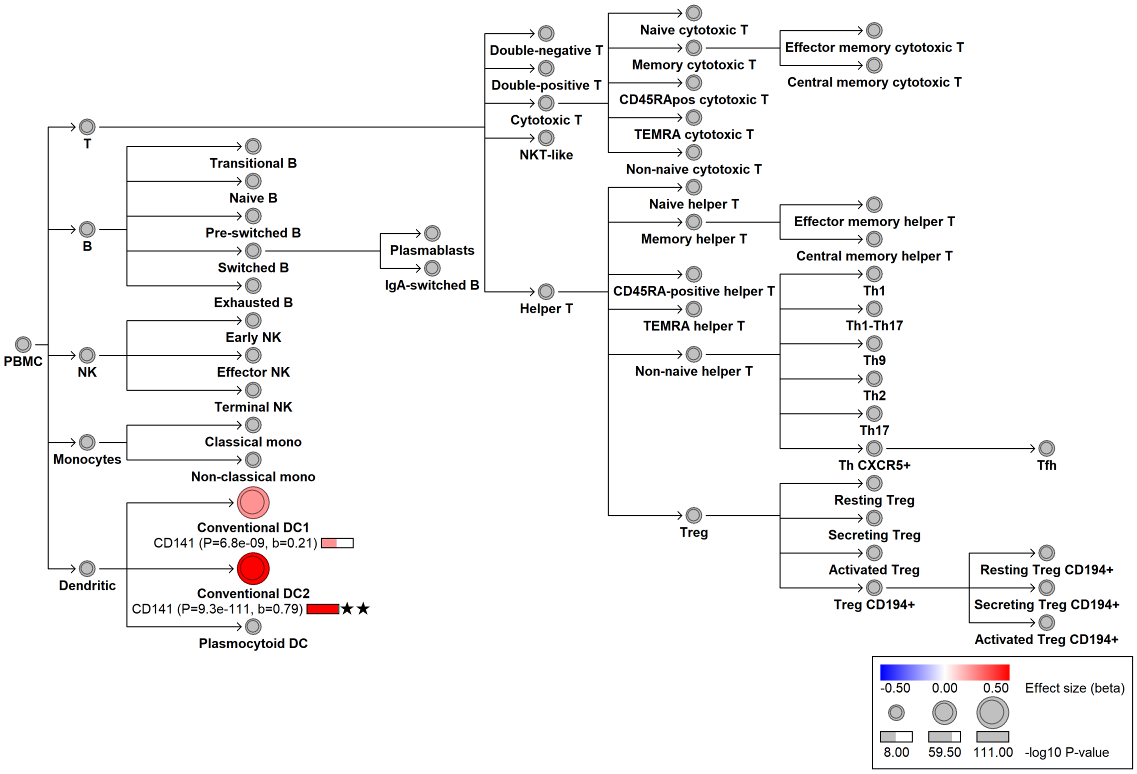

### chr20:23231235[G→!; EAF 14.42%] (chr20:23231235, cs_153)

Candidate genes: *THBD*
Diseases associations: NA
Plasma pQTLs: NA

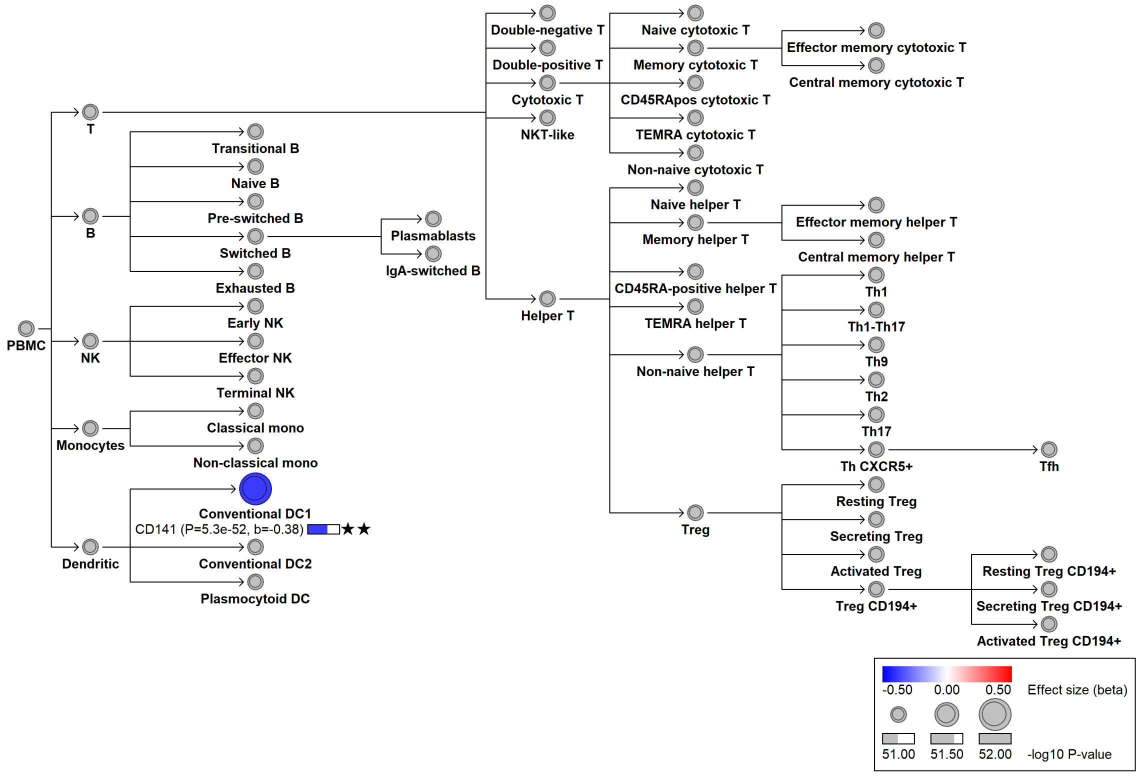

#### chr20:46119308[G→T; EAF 24.479%] (rs4810485, cs_122)

Candidate genes: *CD40*
Diseases associations: Ankylosing spondylitis, Chronic hepatitis B infection, Crohn's disease, Inflammatory bowel disease, Kawasaki disease, Multiple sclerosis, Primary sclerosing cholangitis, Psoriasis, Rheumatoid arthritis, Systemic lupus erythematosus, Ulcerative colitis
Plasma pQTLs: CD22, CD40LG, FCER2, FCRL1, TNFSF13B

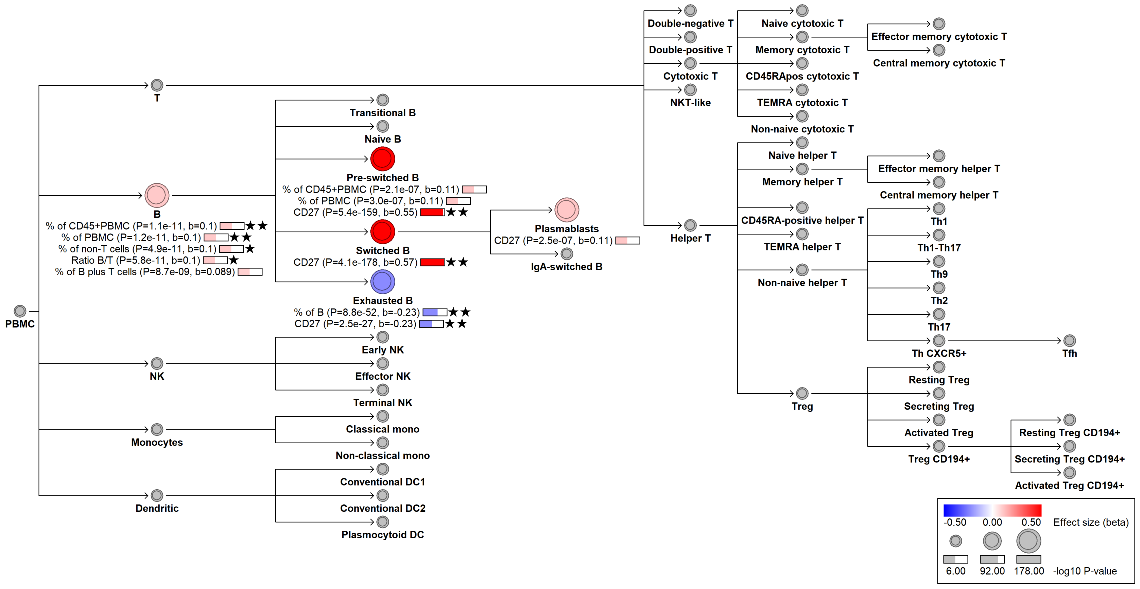

#### chr20:46126737[G→C; EAF 1.658%] (rs41282788, cs_90)

Candidate genes: *CD40*
Diseases associations: NA
Plasma pQTLs: NA

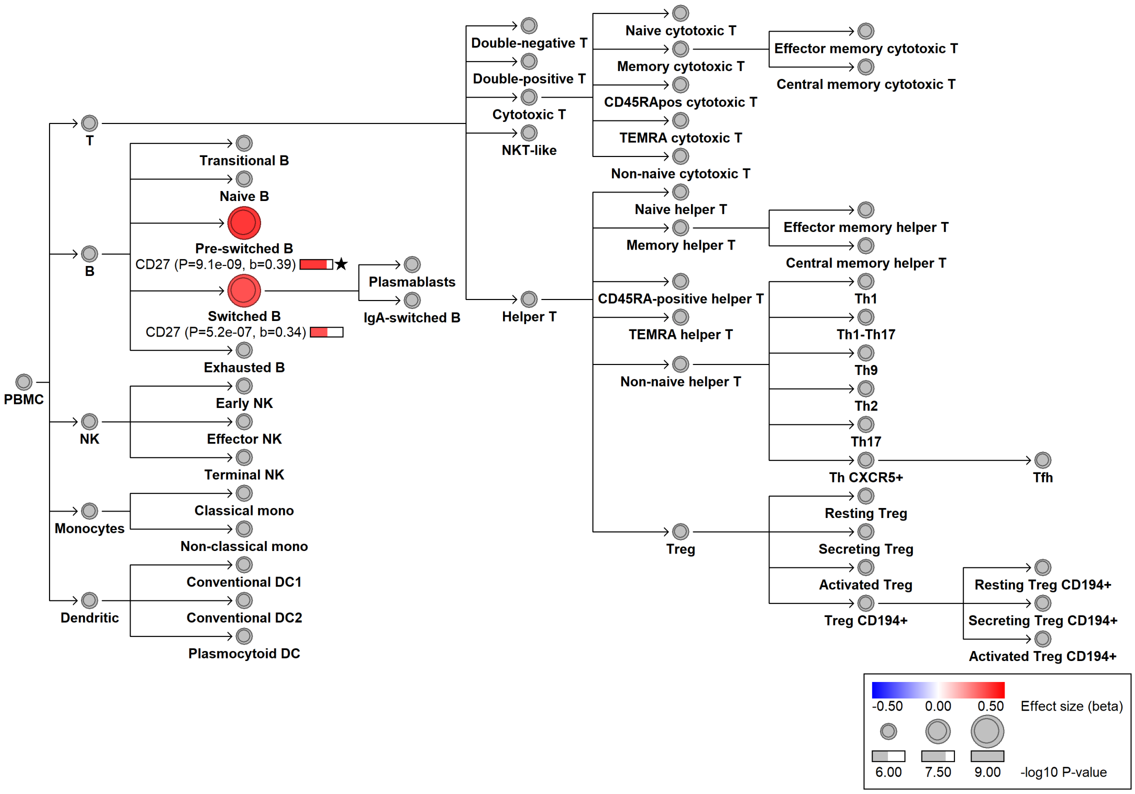

#### chr22:23580365[G→A; EAF 8.614%] (rs9624216, cs_124)

Candidate genes: *IGLL1*
Diseases associations: NA
Plasma pQTLs: IGLL1

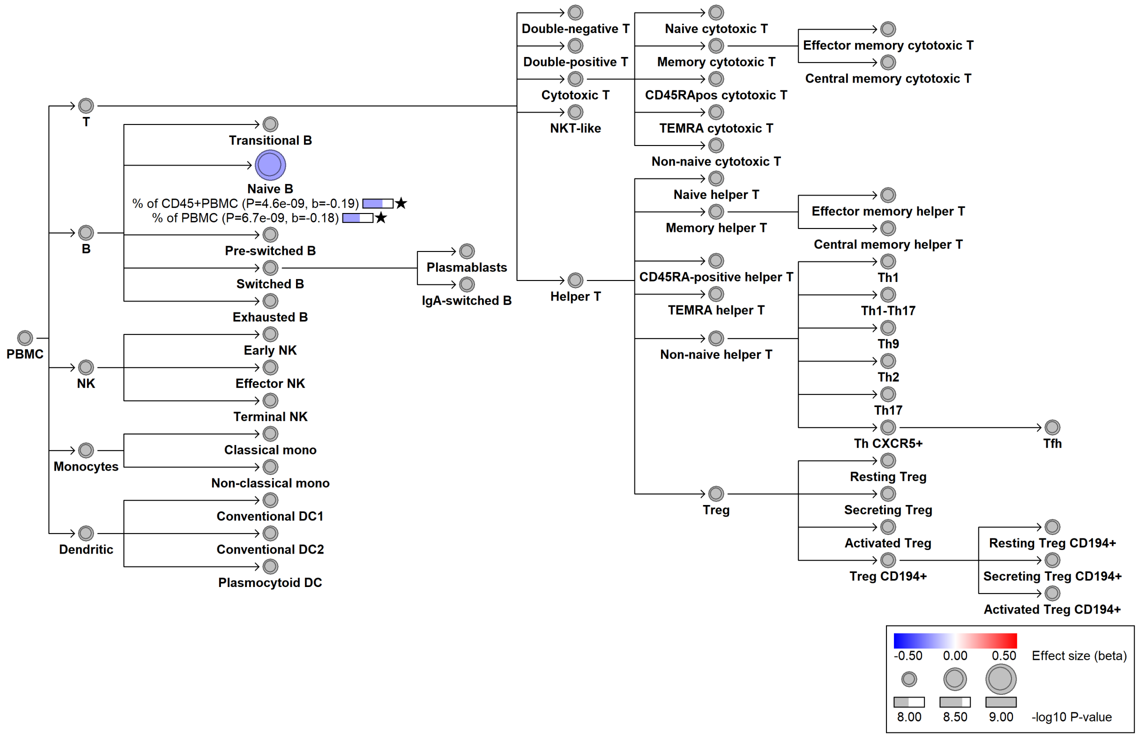

#### chr22:30196498[!→C; EAF 46.49%] (chr22:30196498, cs_170)

Candidate genes: *LIF*
Diseases associations: Autoimmune thyroiditis, Chronic diseases of tonsils, Crohn's disease, IgA nephropathy, Inflammatory bowel disease, Ulcerative colitis
Plasma pQTLs: CCL19, CCL21, CXCL13, Composite immunoglobulin trait (IgA*IgG*IgM), Composite immunoglobulin trait IgA*IgG, Composite immunoglobulin trait IgA/IgG, Composite immunoglobulin trait IgA/IgM, Composite immunoglobulin trait IgM/(IgA*IgG), FAIM3, FCRL5, Immunoglobulin A, TMPRSS11D, TNFRSF13B, TNFRSF17

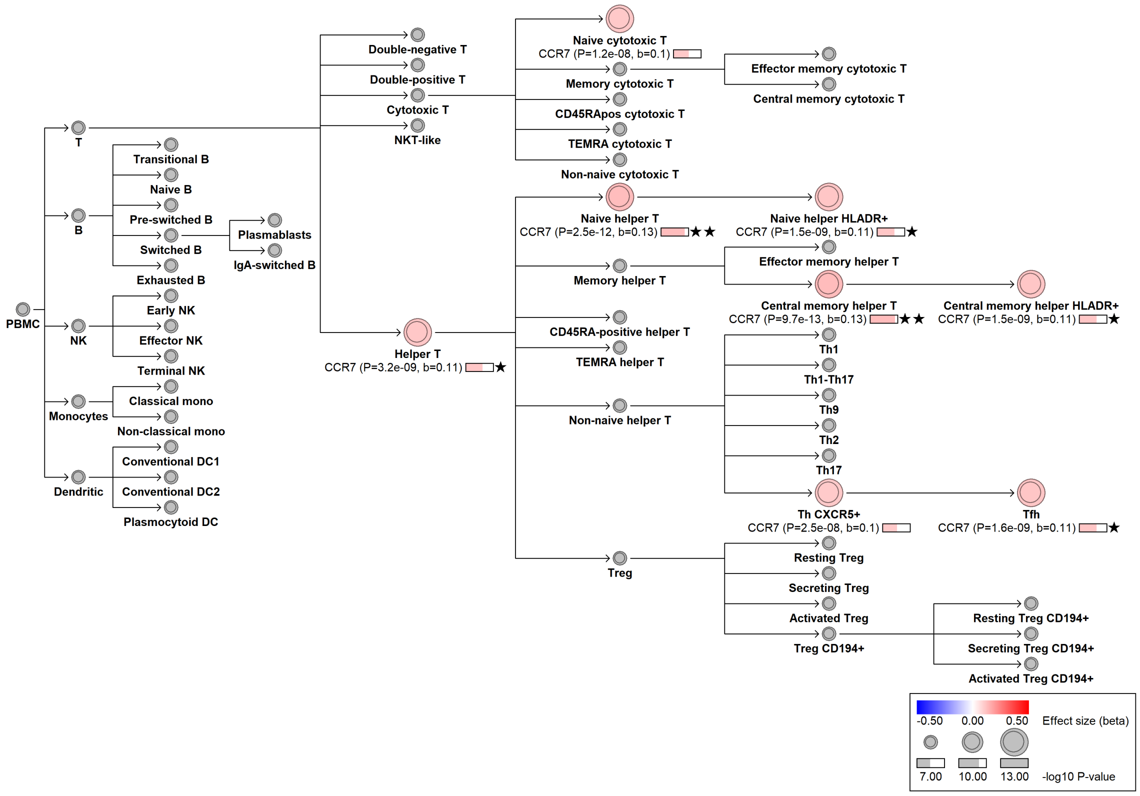

#### chr22:36160775[T→G; EAF 16.861%] (rs132653, cs_178)

Candidate genes: *APOL3*
Diseases associations: NA
Plasma pQTLs: APOL3, DCXR, FAM213A, KRT18, POMGNT2, SCLY

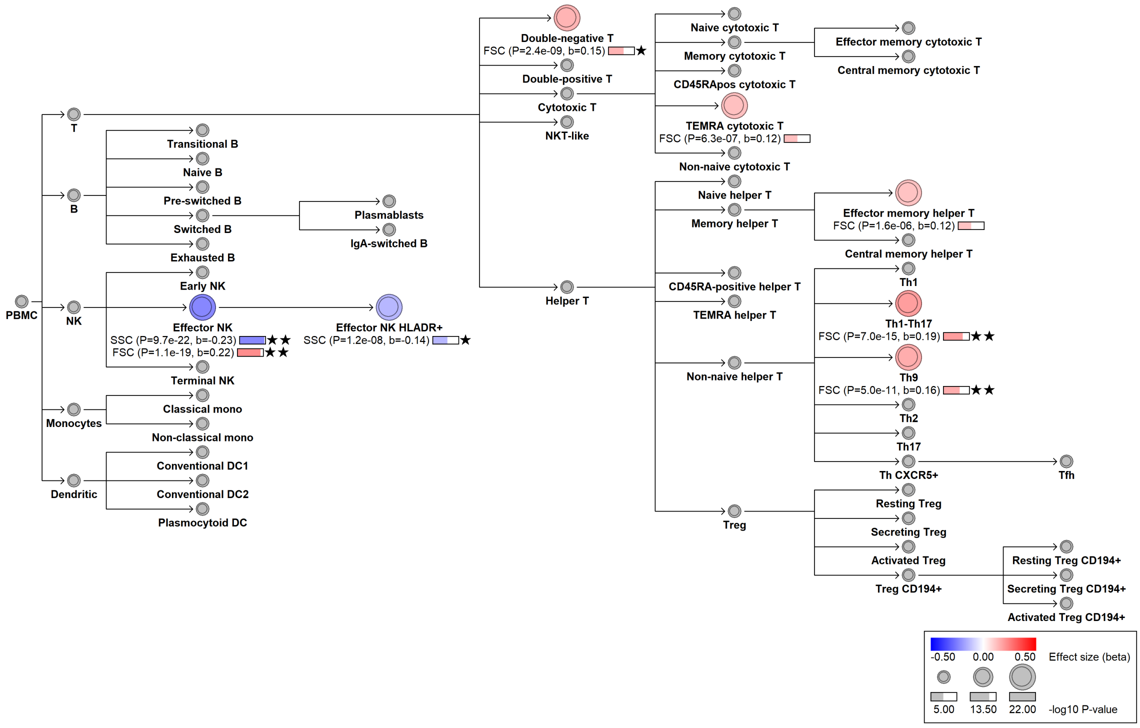

#### chr22:39449893[A→C; EAF 24.017%] (rs1005522, cs_121)

Candidate genes: *MGAT3*
Diseases associations: NA
Plasma pQTLs: CD22, L1CAM, PCDH17, PRTG

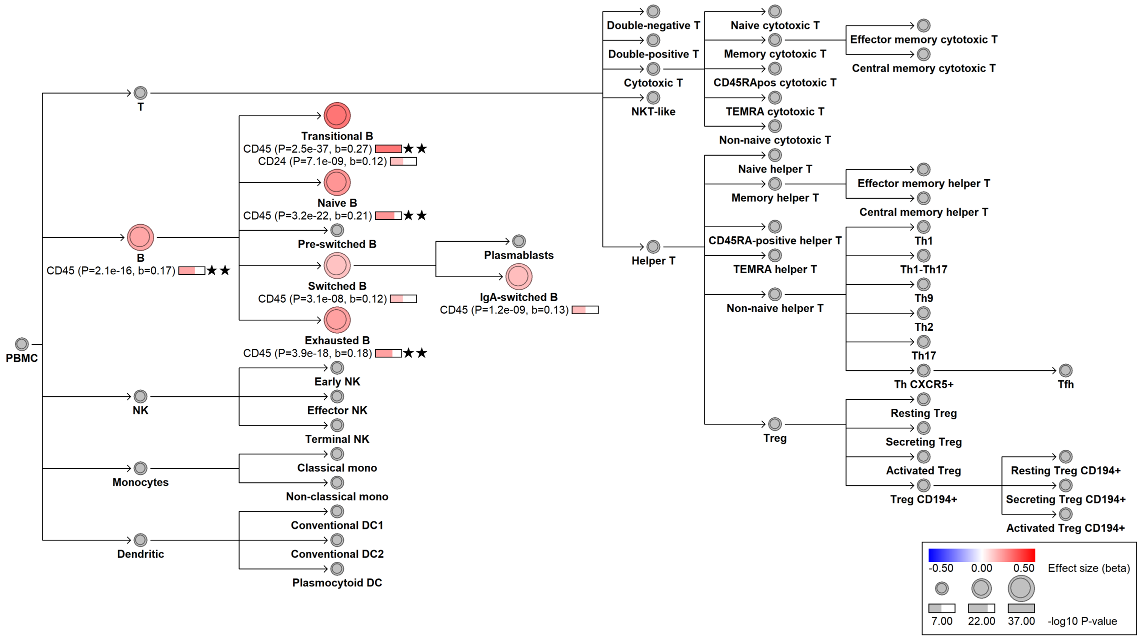

### chr22:41926712[G→C; EAF 9.075%] (rs77874543, cs_81)

Candidate genes: *TNFRSF13C*
Diseases associations: Common variable immunodeficiency
Plasma pQTLs: NA

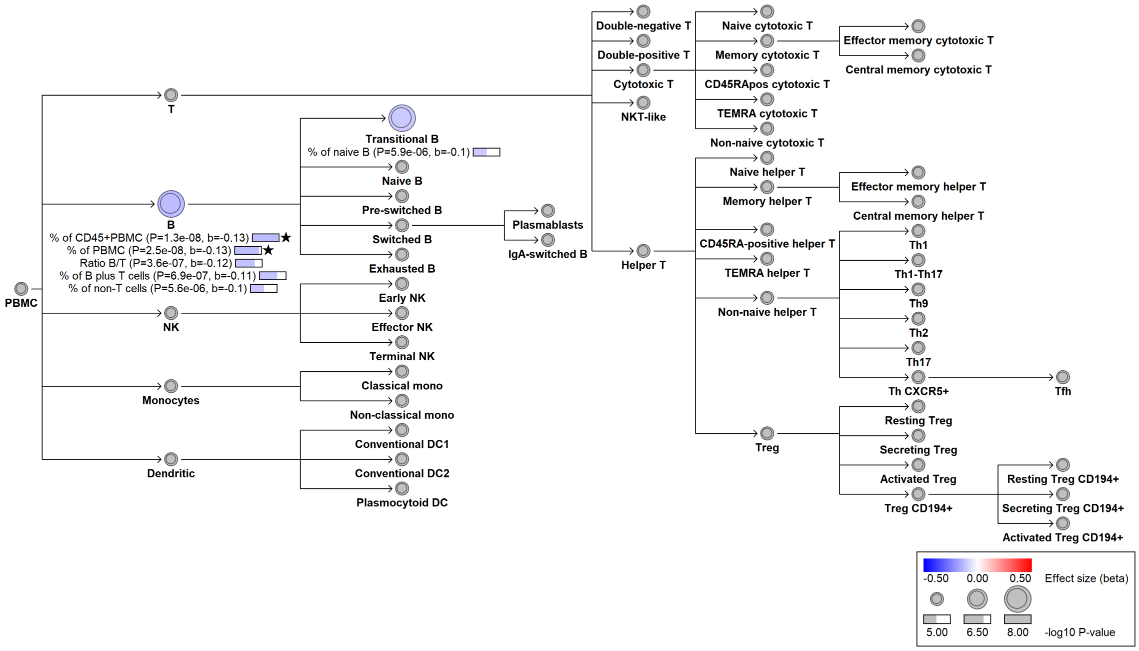

#### chr22:41929013[G→A; EAF 1.816%] (rs75743846, cs_164)

Candidate genes: *TNFRSF13C*
Diseases associations: NA
Plasma pQTLs: FCER2, TCL1A

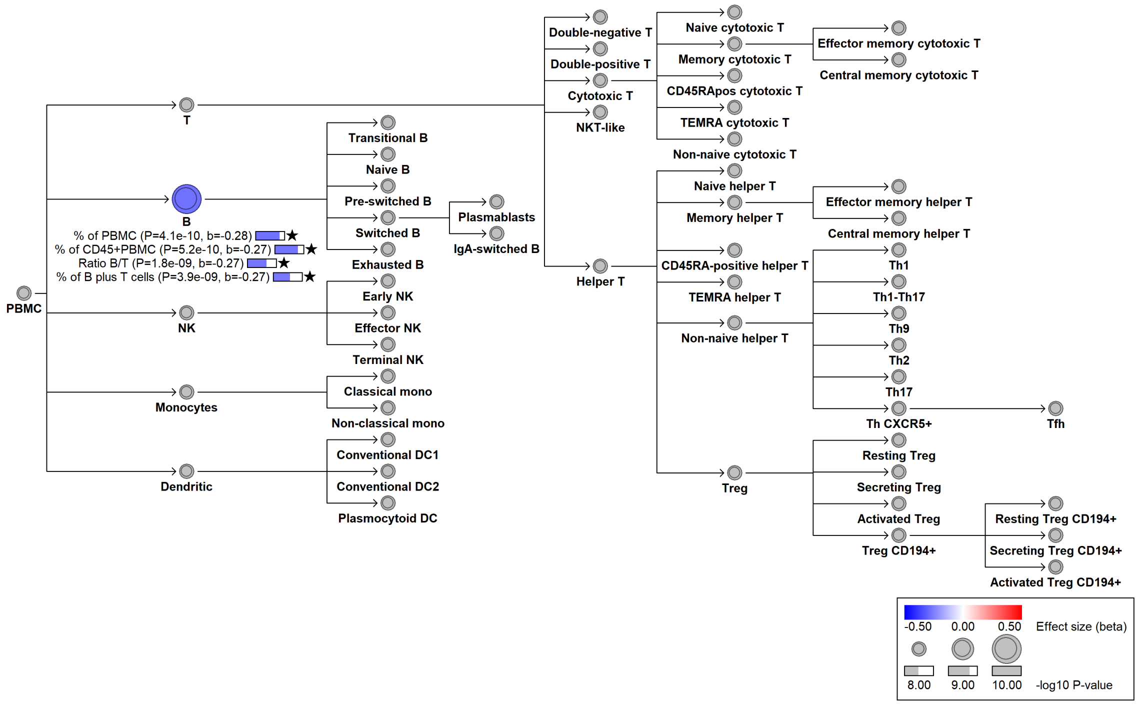

#### chr22:44027084[G→T; EAF 15.686%] (rs13057455, cs_36)

Candidate genes: *PARVB*
Diseases associations: NA
Plasma pQTLs: NA

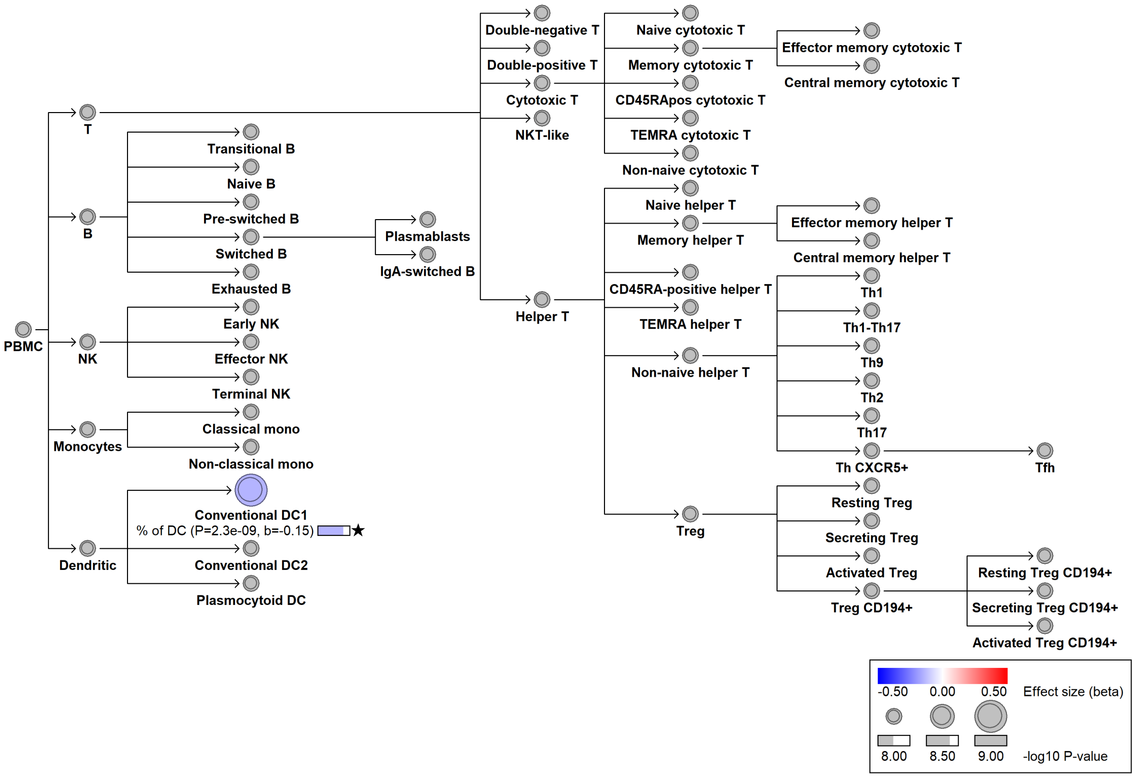

#### chrX:104121909[!→T; EAF 9.686%] (chrX:104121909, cs_234)

Candidate genes: *SLC25A53*
Diseases associations: NA
Plasma pQTLs: NA

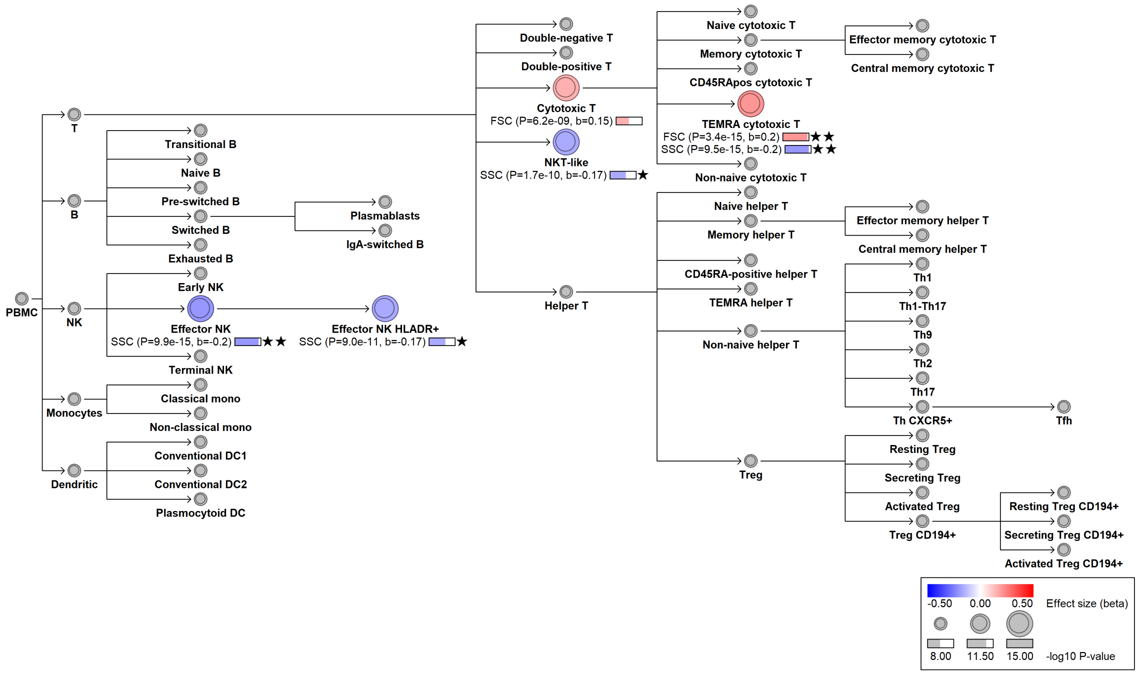
